## Supplemental File 4 for "Integrated view and comparative analysis of baseline protein expression in mouse and rat tissues"

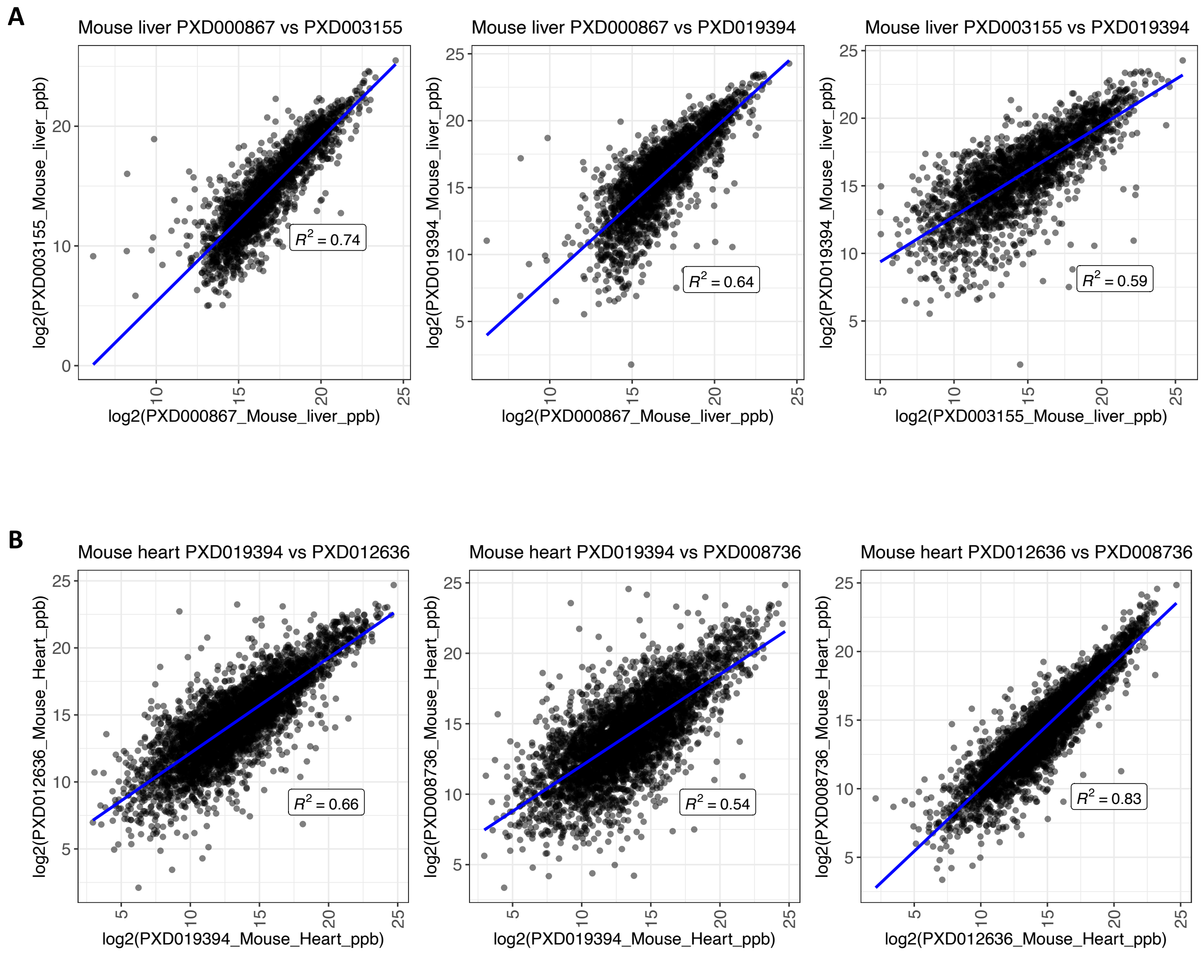
A
Mouse liver PXD000867 vs PXD003155
log2(PXD003155_Mouse_liver_ppb)
log2(PXD000867_Mouse_liver_ppb)
$R^2 = 0.74$
Mouse liver PXD000867 vs PXD019394
log2(PXD019394_Mouse_liver_ppb)
log2(PXD000867_Mouse_liver_ppb)
$R^2 = 0.64$
Mouse liver PXD003155 vs PXD019394
log2(PXD019394_Mouse_liver_ppb)
log2(PXD003155_Mouse_liver_ppb)
$R^2 = 0.59$
B
Mouse heart PXD019394 vs PXD012636
log2(PXD012636_Mouse_Heart_ppb)
log2(PXD019394_Mouse_Heart_ppb)
$R^2 = 0.66$
Mouse heart PXD019394 vs PXD008736
log2(PXD008736_Mouse_Heart_ppb)
log2(PXD019394_Mouse_Heart_ppb)
$R^2 = 0.54$
Mouse heart PXD012636 vs PXD008736
log2(PXD008736_Mouse_Heart_ppb)
log2(PXD012636_Mouse_Heart_ppb)
$R^2 = 0.83$

**C**

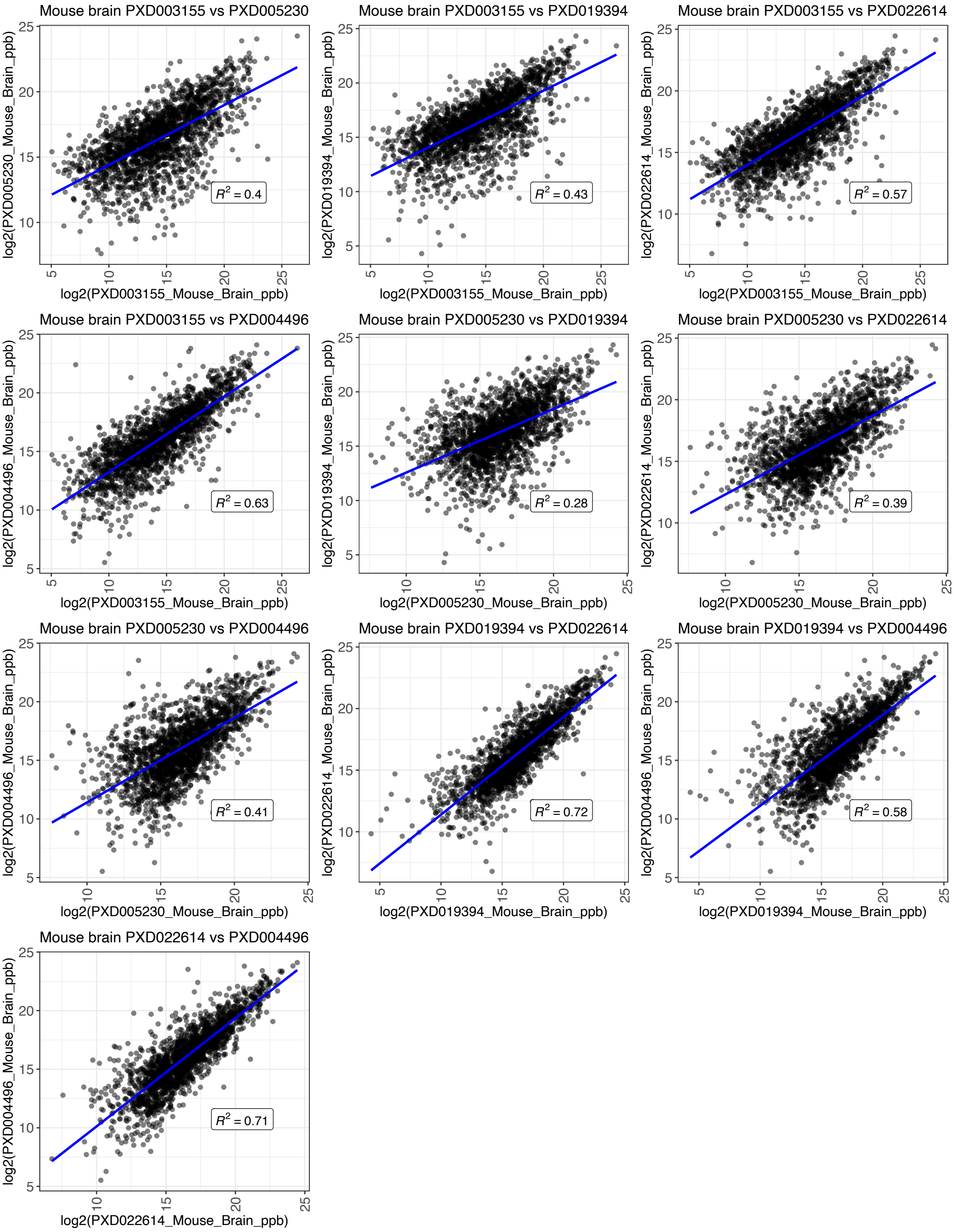
Mouse brain PXD003155 vs PXD005230
log2(PXD005230_Mouse_Brain_ppb)
log2(PXD003155_Mouse_Brain_ppb)
$R^2 = 0.4$
Mouse brain PXD003155 vs PXD019394
log2(PXD019394_Mouse_Brain_ppb)
log2(PXD003155_Mouse_Brain_ppb)
$R^2 = 0.43$
Mouse brain PXD003155 vs PXD022614
log2(PXD022614_Mouse_Brain_ppb)
log2(PXD003155_Mouse_Brain_ppb)
$R^2 = 0.57$
Mouse brain PXD003155 vs PXD004496
log2(PXD004496_Mouse_Brain_ppb)
log2(PXD003155_Mouse_Brain_ppb)
$R^2 = 0.63$
Mouse brain PXD005230 vs PXD019394
log2(PXD019394_Mouse_Brain_ppb)
log2(PXD005230_Mouse_Brain_ppb)
$R^2 = 0.28$
Mouse brain PXD005230 vs PXD022614
log2(PXD022614_Mouse_Brain_ppb)
log2(PXD005230_Mouse_Brain_ppb)
$R^2 = 0.39$
Mouse brain PXD005230 vs PXD004496
log2(PXD004496_Mouse_Brain_ppb)
log2(PXD005230_Mouse_Brain_ppb)
$R^2 = 0.41$
Mouse brain PXD019394 vs PXD022614
log2(PXD022614_Mouse_Brain_ppb)
log2(PXD019394_Mouse_Brain_ppb)
$R^2 = 0.72$
Mouse brain PXD019394 vs PXD004496
log2(PXD004496_Mouse_Brain_ppb)
log2(PXD019394_Mouse_Brain_ppb)
$R^2 = 0.58$
Mouse brain PXD022614 vs PXD004496
log2(PXD004496_Mouse_Brain_ppb)
log2(PXD022614_Mouse_Brain_ppb)
$R^2 = 0.71$

**D**

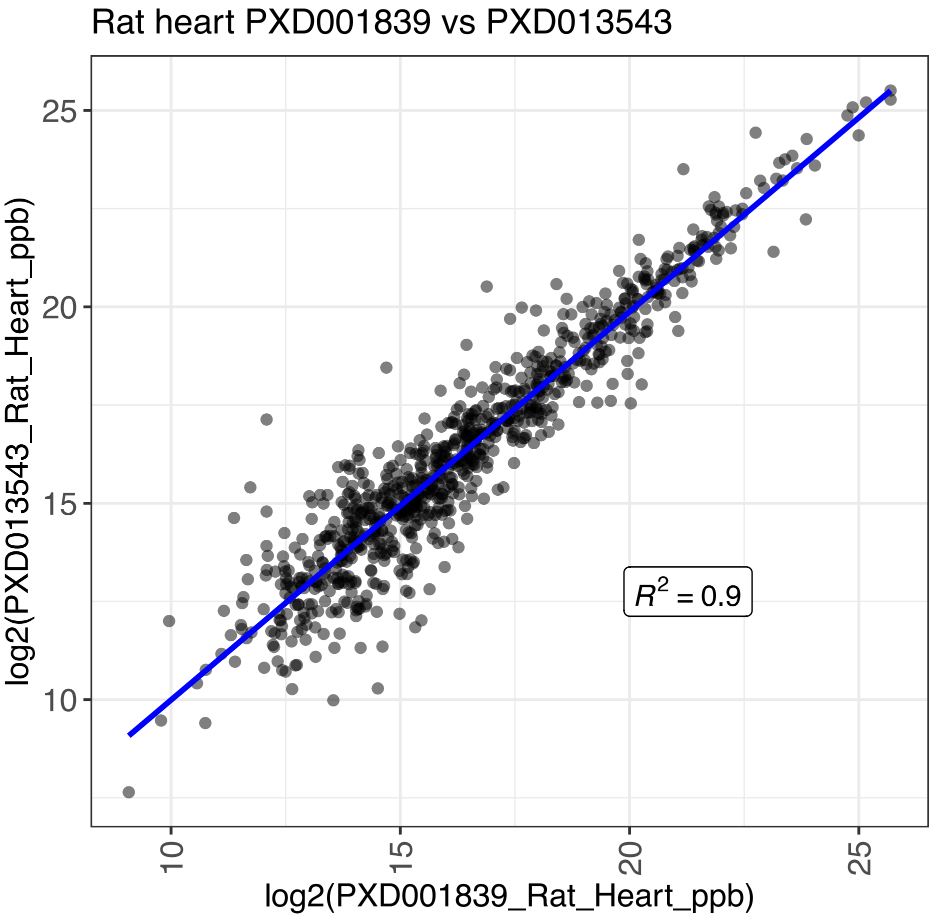


**Supplementary Figure S1**. Correlation of protein expression between datasets for (**A**) mouse liver, (**B**) mouse heart, (**C**) mouse brain and (**D**) rat heart.

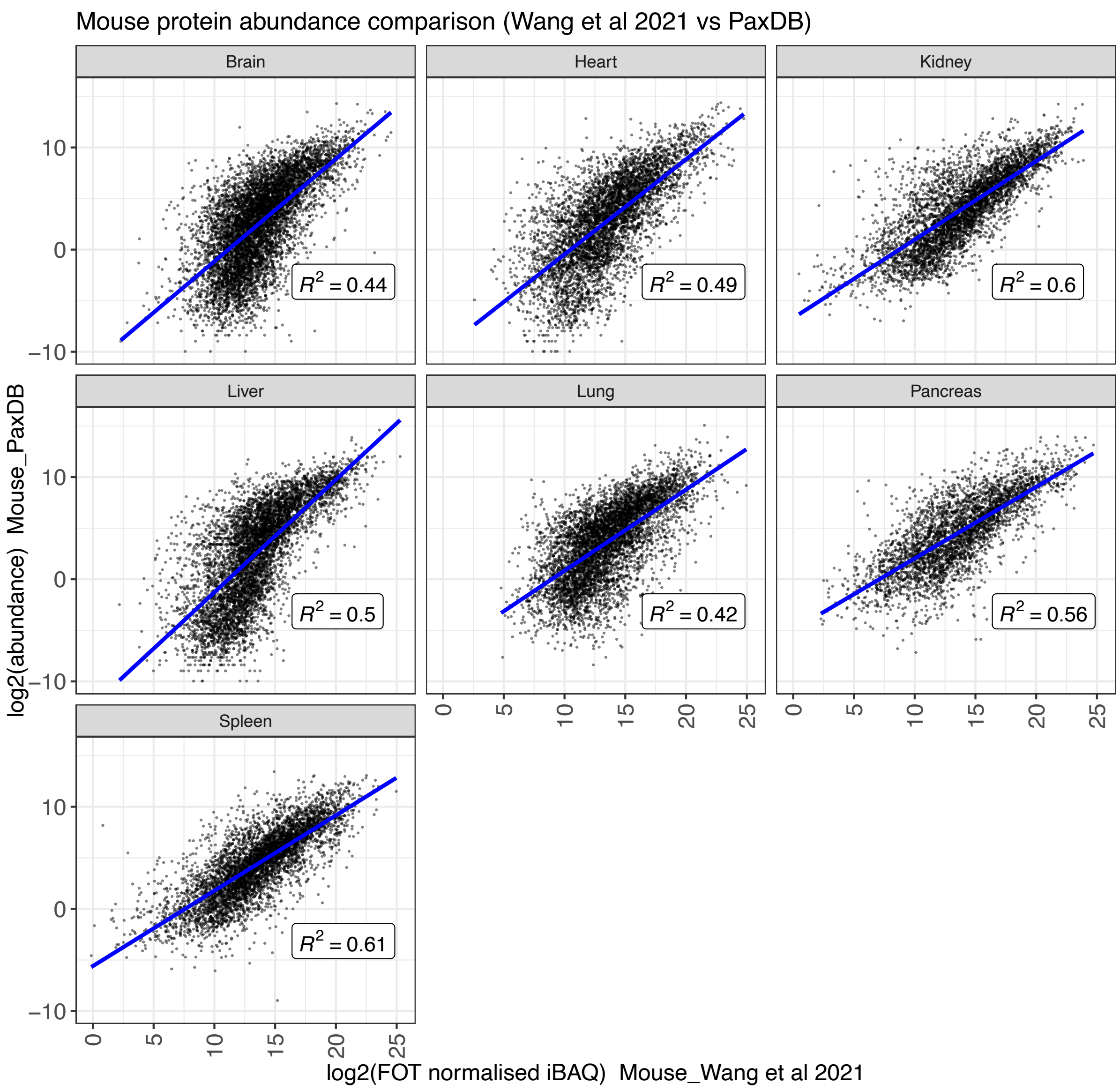


**Supplementary Figure S2**. Comparison of mouse protein abundances in various organs between this study (iBAQ) and PaxDB (spectral counting).

A

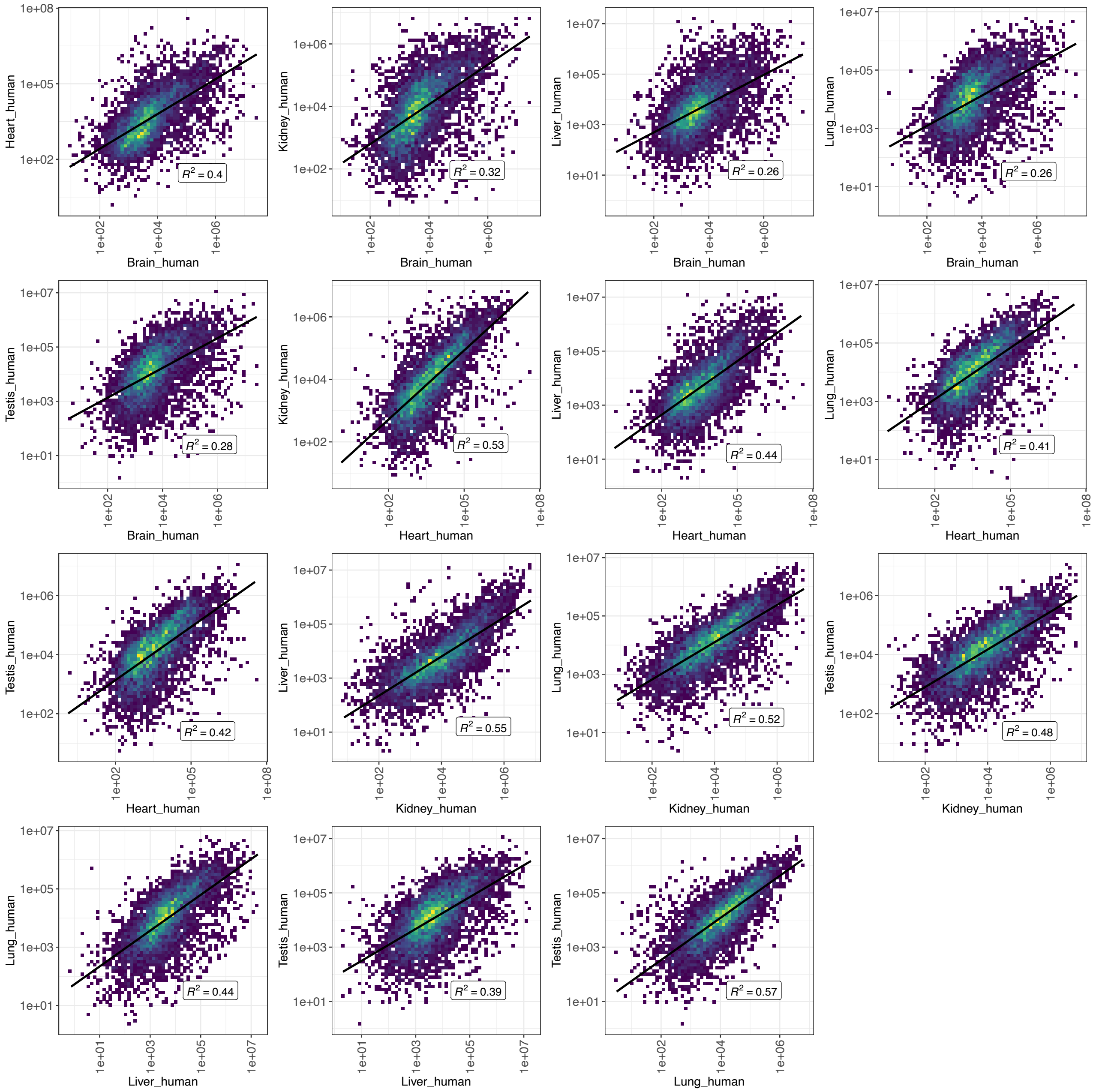

Heart_human
Brain_human
$R^2 = 0.4$
Kidney_human
Brain_human
$R^2 = 0.32$
Liver_human
Brain_human
$R^2 = 0.26$
Lung_human
Brain_human
$R^2 = 0.26$
Testis_human
Brain_human
$R^2 = 0.28$
Kidney_human
Heart_human
$R^2 = 0.53$
Liver_human
Heart_human
$R^2 = 0.44$
Lung_human
Heart_human
$R^2 = 0.41$
Testis_human
Heart_human
$R^2 = 0.42$
Liver_human
Kidney_human
$R^2 = 0.55$
Lung_human
Kidney_human
$R^2 = 0.52$
Testis_human
Kidney_human
$R^2 = 0.48$
Lung_human
Liver_human
$R^2 = 0.44$
Testis_human
Liver_human
$R^2 = 0.39$
Testis_human
Lung_human
$R^2 = 0.57$

B

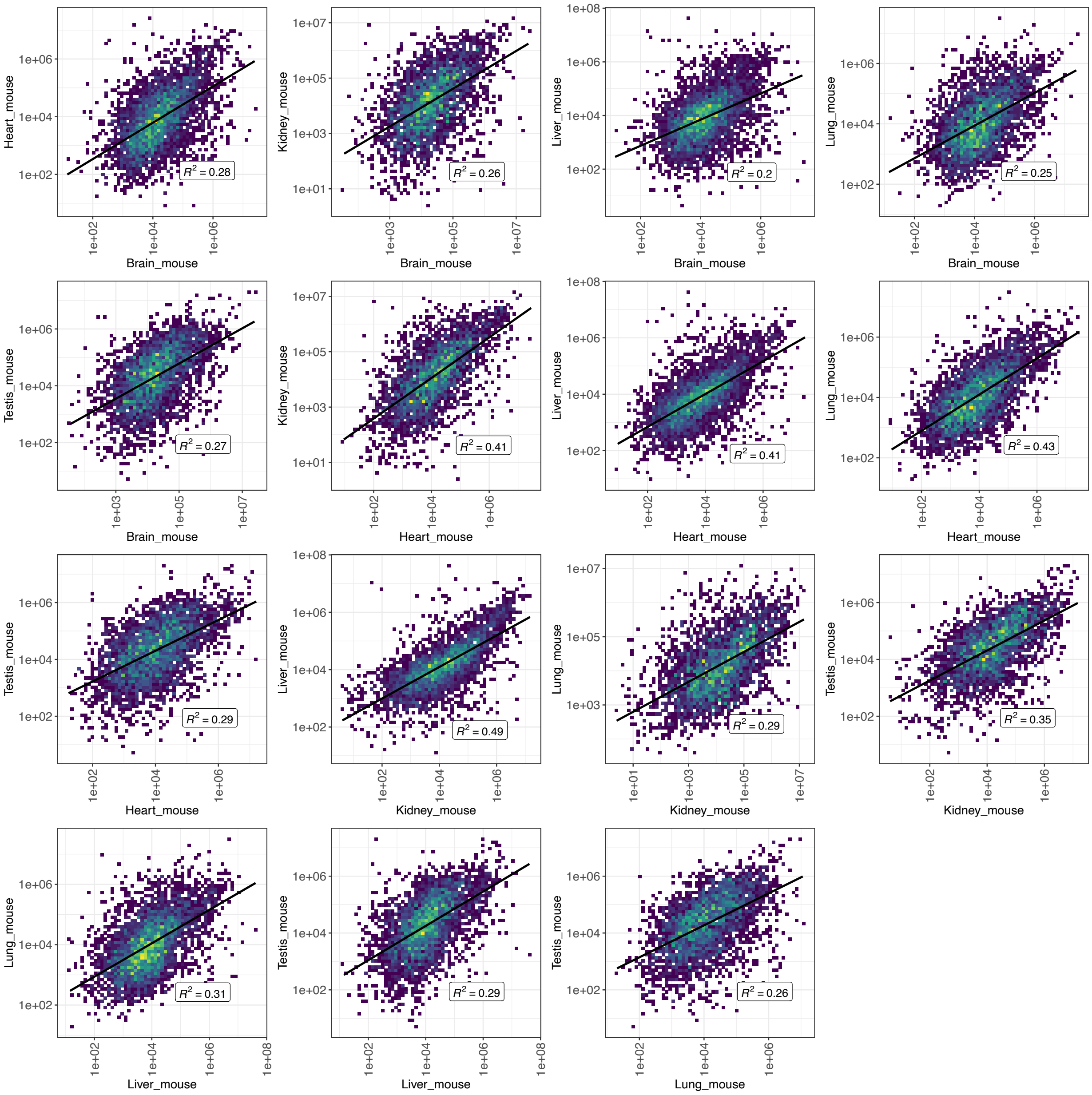
Heart_mouse
Brain_mouse
$R^2 = 0.28$
Kidney_mouse
Brain_mouse
$R^2 = 0.26$
Liver_mouse
Brain_mouse
$R^2 = 0.2$
Lung_mouse
Brain_mouse
$R^2 = 0.25$
Testis_mouse
Brain_mouse
$R^2 = 0.27$
Kidney_mouse
Heart_mouse
$R^2 = 0.41$
Liver_mouse
Heart_mouse
$R^2 = 0.41$
Lung_mouse
Heart_mouse
$R^2 = 0.43$
Testis_mouse
Heart_mouse
$R^2 = 0.29$
Liver_mouse
Kidney_mouse
$R^2 = 0.49$
Lung_mouse
Kidney_mouse
$R^2 = 0.29$
Testis_mouse
Kidney_mouse
$R^2 = 0.35$
Lung_mouse
Liver_mouse
$R^2 = 0.31$
Testis_mouse
Liver_mouse
$R^2 = 0.29$
Testis_mouse
Lung_mouse
$R^2 = 0.26$

**C**

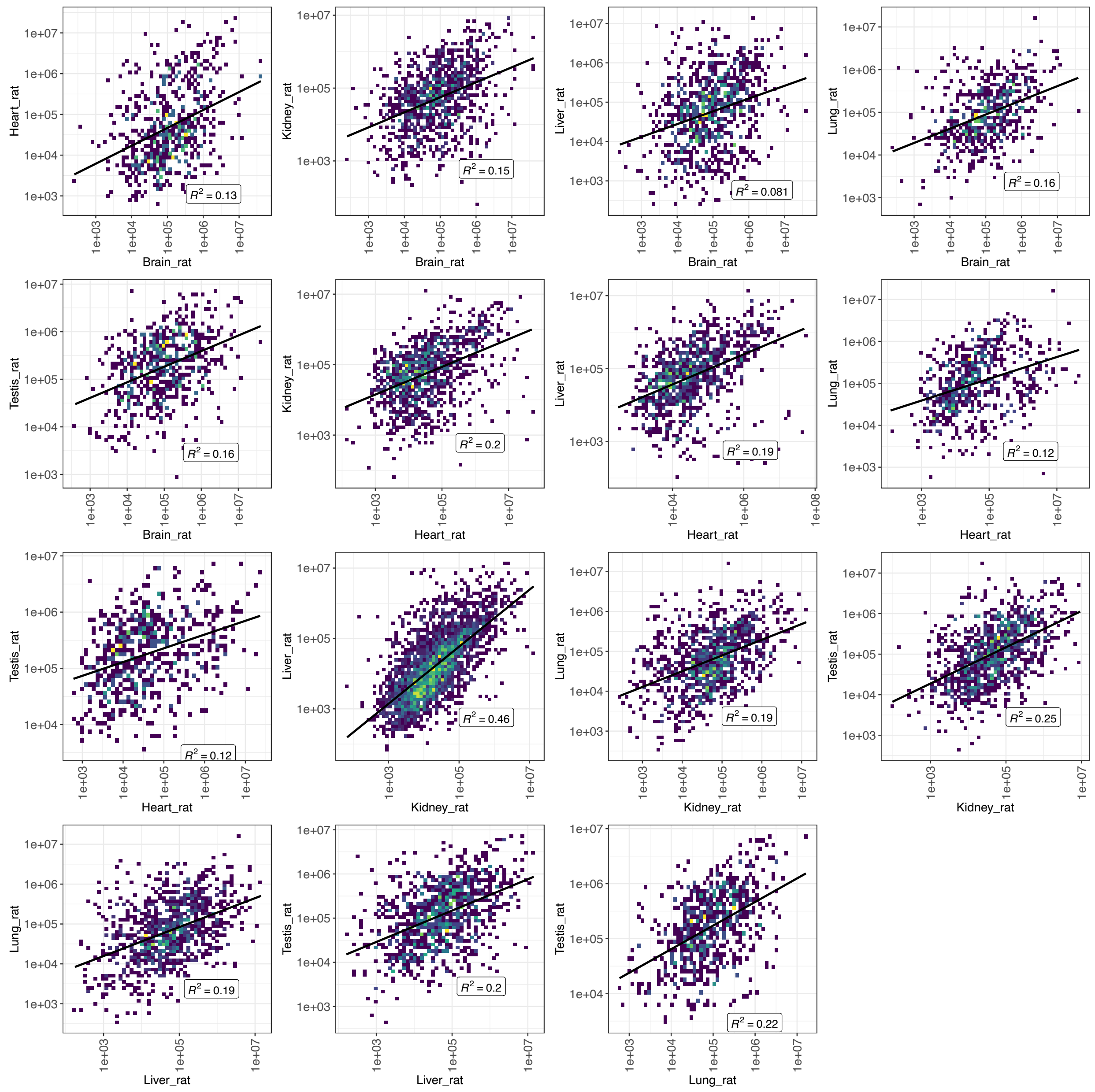


**Supplementary Figure S3**. Correlation of protein expression between organs within (A) human, (B) mouse and (C) rat.
