## Supplemental File 8 for "Integrated view and comparative analysis of baseline protein expression in mouse and rat tissues"

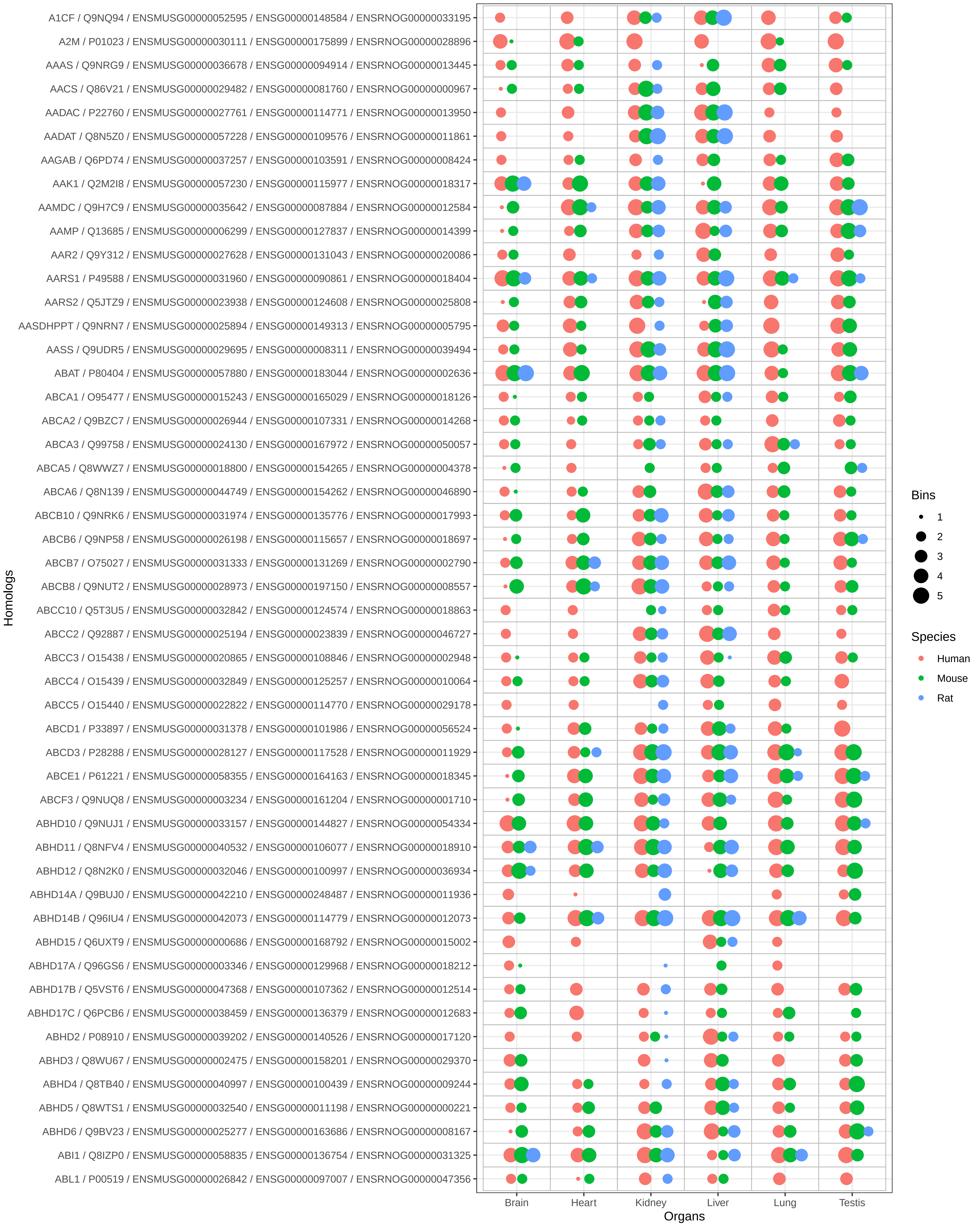
A1CF / Q9NQ94 / ENSMUSG00000052595 / ENSG00000148584 / ENSRNOG00000033195
A2M / P01023 / ENSMUSG00000030111 / ENSG00000175899 / ENSRNOG00000028896
AAAS / Q9NRG9 / ENSMUSG00000036678 / ENSG00000094914 / ENSRNOG00000013445
AACS / Q86V21 / ENSMUSG00000029482 / ENSG00000081760 / ENSRNOG00000000967
AADAC / P22760 / ENSMUSG00000027761 / ENSG00000114771 / ENSRNOG00000013950
AADAT / Q8N5Z0 / ENSMUSG00000057228 / ENSG00000109576 / ENSRNOG00000011861
AAGAB / Q6PD74 / ENSMUSG00000037257 / ENSG00000103591 / ENSRNOG00000008424
AAK1 / Q2M2I8 / ENSMUSG00000057230 / ENSG00000115977 / ENSRNOG00000018317
AAMDC / Q9H7C9 / ENSMUSG00000035642 / ENSG00000087884 / ENSRNOG00000012584
AAMP / Q13685 / ENSMUSG00000006299 / ENSG00000127837 / ENSRNOG00000014399
AAR2 / Q9Y312 / ENSMUSG00000027628 / ENSG00000131043 / ENSRNOG00000020086
AARS1 / P49588 / ENSMUSG00000031960 / ENSG00000090861 / ENSRNOG00000018404
AARS2 / Q5JTZ9 / ENSMUSG00000023938 / ENSG00000124608 / ENSRNOG00000025808
AASDHPPT / Q9NRN7 / ENSMUSG00000025894 / ENSG00000149313 / ENSRNOG00000005795
AASS / Q9UDR5 / ENSMUSG00000029695 / ENSG00000008311 / ENSRNOG00000039494
ABAT / P80404 / ENSMUSG00000057880 / ENSG00000183044 / ENSRNOG00000002636
ABCA1 / O95477 / ENSMUSG00000015243 / ENSG00000165029 / ENSRNOG00000018126
ABCA2 / Q9BZC7 / ENSMUSG00000026944 / ENSG00000107331 / ENSRNOG00000014268
ABCA3 / Q99758 / ENSMUSG00000024130 / ENSG00000167972 / ENSRNOG00000050057
ABCA5 / Q8WWZ7 / ENSMUSG00000018800 / ENSG00000154265 / ENSRNOG00000004378
ABCA6 / Q8N139 / ENSMUSG00000044749 / ENSG00000154262 / ENSRNOG00000046890
ABCB10 / Q9NRK6 / ENSMUSG00000031974 / ENSG00000135776 / ENSRNOG00000017993
ABCB6 / Q9NP58 / ENSMUSG00000026198 / ENSG00000115657 / ENSRNOG00000018697
ABCB7 / O75027 / ENSMUSG00000031333 / ENSG00000131269 / ENSRNOG00000002790
ABCB8 / Q9NUT2 / ENSMUSG00000028973 / ENSG00000197150 / ENSRNOG00000008557
ABCC10 / Q5T3U5 / ENSMUSG00000032842 / ENSG00000124574 / ENSRNOG00000018863
ABCC2 / Q92887 / ENSMUSG00000025194 / ENSG00000023839 / ENSRNOG00000046727
ABCC3 / O15438 / ENSMUSG00000020865 / ENSG00000108846 / ENSRNOG00000002948
ABCC4 / O15439 / ENSMUSG00000032849 / ENSG00000125257 / ENSRNOG00000010064
ABCC5 / O15440 / ENSMUSG00000022822 / ENSG00000114770 / ENSRNOG00000029178
ABCD1 / P33897 / ENSMUSG00000031378 / ENSG00000101986 / ENSRNOG00000056524
ABCD3 / P28288 / ENSMUSG00000028127 / ENSG00000117528 / ENSRNOG00000011929
ABCE1 / P61221 / ENSMUSG00000058355 / ENSG00000164163 / ENSRNOG00000018345
ABCF3 / Q9NUQ8 / ENSMUSG00000003234 / ENSG00000161204 / ENSRNOG00000001710
ABHD10 / Q9NUJ1 / ENSMUSG00000033157 / ENSG00000144827 / ENSRNOG00000054334
ABHD11 / Q8NFV4 / ENSMUSG00000040532 / ENSG00000106077 / ENSRNOG00000018910
ABHD12 / Q8N2K0 / ENSMUSG00000032046 / ENSG00000100997 / ENSRNOG00000036934
ABHD14A / Q9BUJ0 / ENSMUSG00000042210 / ENSG00000248487 / ENSRNOG00000011936
ABHD14B / Q96IU4 / ENSMUSG00000042073 / ENSG00000114779 / ENSRNOG00000012073
ABHD15 / Q6UXT9 / ENSMUSG00000000686 / ENSG00000168792 / ENSRNOG00000015002
ABHD17A / Q96GS6 / ENSMUSG00000003346 / ENSG00000129968 / ENSRNOG00000018212
ABHD17B / Q5VST6 / ENSMUSG00000047368 / ENSG00000107362 / ENSRNOG00000012514
ABHD17C / Q6PCB6 / ENSMUSG00000038459 / ENSG00000136379 / ENSRNOG00000012683
ABHD2 / P08910 / ENSMUSG00000039202 / ENSG00000140526 / ENSRNOG00000017120
ABHD3 / Q8WU67 / ENSMUSG00000002475 / ENSG00000158201 / ENSRNOG00000029370
ABHD4 / Q8TB40 / ENSMUSG00000040997 / ENSG00000100439 / ENSRNOG00000009244
ABHD5 / Q8WTS1 / ENSMUSG00000032540 / ENSG00000011198 / ENSRNOG00000000221
ABHD6 / Q9BV23 / ENSMUSG00000025277 / ENSG00000163686 / ENSRNOG00000008167
ABI1 / Q8IZP0 / ENSMUSG00000058835 / ENSG00000136754 / ENSRNOG00000031325
ABL1 / P00519 / ENSMUSG00000026842 / ENSG00000097007 / ENSRNOG00000047356
Homologs
Brain
Heart
Kidney
Liver
Lung
Testis
Organs
Bins
1
2
3
4
5
Species
Human
Mouse
Rat

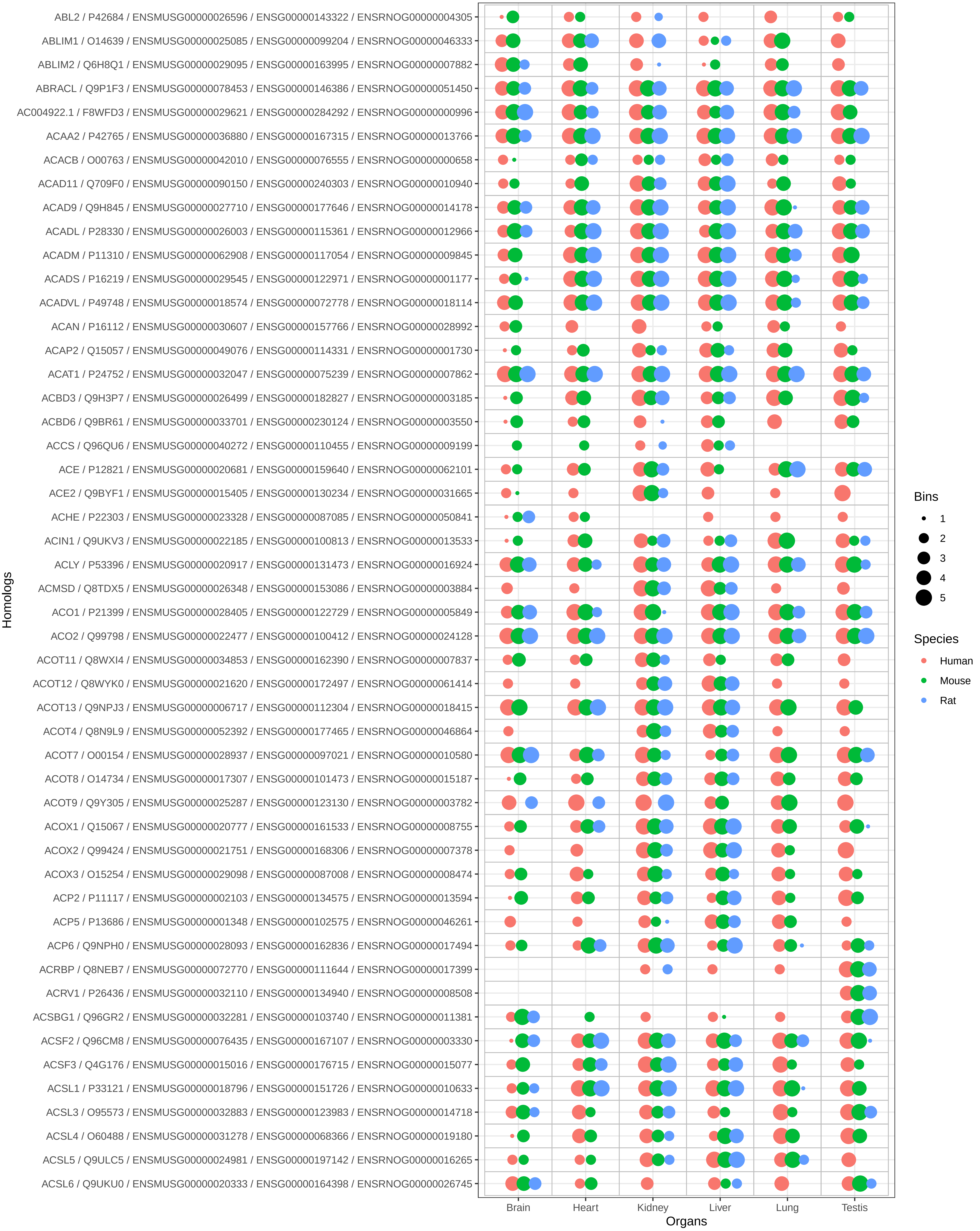
ABL2 / P42684 / ENSMUSG00000026596 / ENSG00000143322 / ENSRNOG00000004305
ABLIM1 / O14639 / ENSMUSG00000025085 / ENSG00000099204 / ENSRNOG00000046333
ABLIM2 / Q6H8Q1 / ENSMUSG00000029095 / ENSG00000163995 / ENSRNOG00000007882
ABRACL / Q9P1F3 / ENSMUSG00000078453 / ENSG00000146386 / ENSRNOG00000051450
AC004922.1 / F8WFD3 / ENSMUSG00000029621 / ENSG00000284292 / ENSRNOG00000000996
ACAA2 / P42765 / ENSMUSG00000036880 / ENSG00000167315 / ENSRNOG00000013766
ACACB / O00763 / ENSMUSG00000042010 / ENSG00000076555 / ENSRNOG00000000658
ACAD11 / Q709F0 / ENSMUSG00000090150 / ENSG00000240303 / ENSRNOG00000010940
ACAD9 / Q9H845 / ENSMUSG00000027710 / ENSG00000177646 / ENSRNOG00000014178
ACADL / P28330 / ENSMUSG00000026003 / ENSG00000115361 / ENSRNOG00000012966
ACADM / P11310 / ENSMUSG00000062908 / ENSG00000117054 / ENSRNOG00000009845
ACADS / P16219 / ENSMUSG00000029545 / ENSG00000122971 / ENSRNOG00000001177
ACADVL / P49748 / ENSMUSG00000018574 / ENSG00000072778 / ENSRNOG00000018114
ACAN / P16112 / ENSMUSG00000030607 / ENSG00000157766 / ENSRNOG00000028992
ACAP2 / Q15057 / ENSMUSG00000049076 / ENSG00000114331 / ENSRNOG00000001730
ACAT1 / P24752 / ENSMUSG00000032047 / ENSG00000075239 / ENSRNOG00000007862
ACBD3 / Q9H3P7 / ENSMUSG00000026499 / ENSG00000182827 / ENSRNOG00000003185
ACBD6 / Q9BR61 / ENSMUSG00000033701 / ENSG00000230124 / ENSRNOG00000003550
ACCS / Q96QU6 / ENSMUSG00000040272 / ENSG00000110455 / ENSRNOG00000009199
ACE / P12821 / ENSMUSG00000020681 / ENSG00000159640 / ENSRNOG00000062101
ACE2 / Q9BYF1 / ENSMUSG00000015405 / ENSG00000130234 / ENSRNOG00000031665
ACHE / P22303 / ENSMUSG00000023328 / ENSG00000087085 / ENSRNOG00000050841
ACIN1 / Q9UKV3 / ENSMUSG00000022185 / ENSG00000100813 / ENSRNOG00000013533
ACLY / P53396 / ENSMUSG00000020917 / ENSG00000131473 / ENSRNOG00000016924
ACMSD / Q8TDX5 / ENSMUSG00000026348 / ENSG00000153086 / ENSRNOG00000003884
ACO1 / P21399 / ENSMUSG00000028405 / ENSG00000122729 / ENSRNOG00000005849
ACO2 / Q99798 / ENSMUSG00000022477 / ENSG00000100412 / ENSRNOG00000024128
ACOT11 / Q8WXI4 / ENSMUSG00000034853 / ENSG00000162390 / ENSRNOG00000007837
ACOT12 / Q8WYK0 / ENSMUSG00000021620 / ENSG00000172497 / ENSRNOG00000061414
ACOT13 / Q9NPJ3 / ENSMUSG00000006717 / ENSG00000112304 / ENSRNOG00000018415
ACOT4 / Q8N9L9 / ENSMUSG00000052392 / ENSG00000177465 / ENSRNOG00000046864
ACOT7 / O00154 / ENSMUSG00000028937 / ENSG00000097021 / ENSRNOG00000010580
ACOT8 / O14734 / ENSMUSG00000017307 / ENSG00000101473 / ENSRNOG00000015187
ACOT9 / Q9Y305 / ENSMUSG00000025287 / ENSG00000123130 / ENSRNOG00000003782
ACOX1 / Q15067 / ENSMUSG00000020777 / ENSG00000161533 / ENSRNOG00000008755
ACOX2 / Q99424 / ENSMUSG00000021751 / ENSG00000168306 / ENSRNOG00000007378
ACOX3 / O15254 / ENSMUSG00000029098 / ENSG00000087008 / ENSRNOG00000008474
ACP2 / P11117 / ENSMUSG00000002103 / ENSG00000134575 / ENSRNOG00000013594
ACP5 / P13686 / ENSMUSG00000001348 / ENSG00000102575 / ENSRNOG00000046261
ACP6 / Q9NPH0 / ENSMUSG00000028093 / ENSG00000162836 / ENSRNOG00000017494
ACRBP / Q8NEB7 / ENSMUSG00000072770 / ENSG00000111644 / ENSRNOG00000017399
ACRV1 / P26436 / ENSMUSG00000032110 / ENSG00000134940 / ENSRNOG00000008508
ACSBG1 / Q96GR2 / ENSMUSG00000032281 / ENSG00000103740 / ENSRNOG00000011381
ACSF2 / Q96CM8 / ENSMUSG00000076435 / ENSG00000167107 / ENSRNOG00000003330
ACSF3 / Q4G176 / ENSMUSG00000015016 / ENSG00000176715 / ENSRNOG00000015077
ACSL1 / P33121 / ENSMUSG00000018796 / ENSG00000151726 / ENSRNOG00000010633
ACSL3 / O95573 / ENSMUSG00000032883 / ENSG00000123983 / ENSRNOG00000014718
ACSL4 / O60488 / ENSMUSG00000031278 / ENSG00000068366 / ENSRNOG00000019180
ACSL5 / Q9ULC5 / ENSMUSG00000024981 / ENSG00000197142 / ENSRNOG00000016265
ACSL6 / Q9UKU0 / ENSMUSG00000020333 / ENSG00000164398 / ENSRNOG00000026745
Homologs
Brain
Heart
Kidney
Liver
Lung
Testis
Organs
Bins
1
2
3
4
5
Species
Human
Mouse
Rat

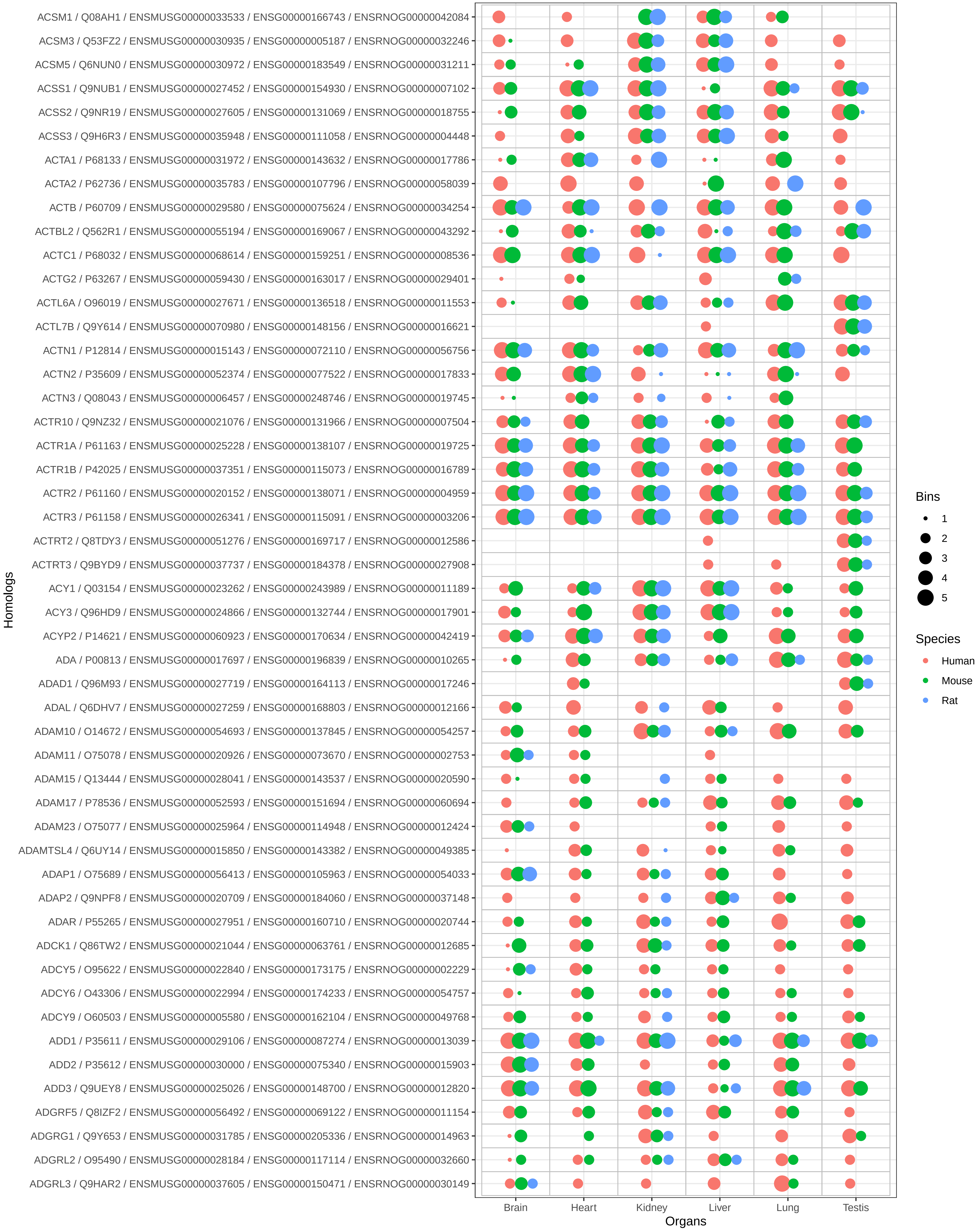
ACSM1 / Q08AH1 / ENSMUSG00000033533 / ENSG00000166743 / ENSRNOG00000042084
ACSM3 / Q53FZ2 / ENSMUSG00000030935 / ENSG00000005187 / ENSRNOG00000032246
ACSM5 / Q6NUN0 / ENSMUSG00000030972 / ENSG00000183549 / ENSRNOG00000031211
ACSS1 / Q9NUB1 / ENSMUSG00000027452 / ENSG00000154930 / ENSRNOG00000007102
ACSS2 / Q9NR19 / ENSMUSG00000027605 / ENSG00000131069 / ENSRNOG00000018755
ACSS3 / Q9H6R3 / ENSMUSG00000035948 / ENSG00000111058 / ENSRNOG00000004448
ACTA1 / P68133 / ENSMUSG00000031972 / ENSG00000143632 / ENSRNOG00000017786
ACTA2 / P62736 / ENSMUSG00000035783 / ENSG00000107796 / ENSRNOG00000058039
ACTB / P60709 / ENSMUSG00000029580 / ENSG00000075624 / ENSRNOG00000034254
ACTBL2 / Q562R1 / ENSMUSG00000055194 / ENSG00000169067 / ENSRNOG00000043292
ACTC1 / P68032 / ENSMUSG00000068614 / ENSG00000159251 / ENSRNOG00000008536
ACTG2 / P63267 / ENSMUSG00000059430 / ENSG00000163017 / ENSRNOG00000029401
ACTL6A / O96019 / ENSMUSG00000027671 / ENSG00000136518 / ENSRNOG00000011553
ACTL7B / Q9Y614 / ENSMUSG00000070980 / ENSG00000148156 / ENSRNOG00000016621
ACTN1 / P12814 / ENSMUSG00000015143 / ENSG00000072110 / ENSRNOG00000056756
ACTN2 / P35609 / ENSMUSG00000052374 / ENSG00000077522 / ENSRNOG00000017833
ACTN3 / Q08043 / ENSMUSG00000006457 / ENSG00000248746 / ENSRNOG00000019745
ACTR10 / Q9NZ32 / ENSMUSG00000021076 / ENSG00000131966 / ENSRNOG00000007504
ACTR1A / P61163 / ENSMUSG00000025228 / ENSG00000138107 / ENSRNOG00000019725
ACTR1B / P42025 / ENSMUSG00000037351 / ENSG00000115073 / ENSRNOG00000016789
ACTR2 / P61160 / ENSMUSG00000020152 / ENSG00000138071 / ENSRNOG00000004959
ACTR3 / P61158 / ENSMUSG00000026341 / ENSG00000115091 / ENSRNOG00000003206
ACTRT2 / Q8TDY3 / ENSMUSG00000051276 / ENSG00000169717 / ENSRNOG00000012586
ACTRT3 / Q9BYD9 / ENSMUSG00000037737 / ENSG00000184378 / ENSRNOG00000027908
ACY1 / Q03154 / ENSMUSG00000023262 / ENSG00000243989 / ENSRNOG00000011189
ACY3 / Q96HD9 / ENSMUSG00000024866 / ENSG00000132744 / ENSRNOG00000017901
ACYP2 / P14621 / ENSMUSG00000060923 / ENSG00000170634 / ENSRNOG00000042419
ADA / P00813 / ENSMUSG00000017697 / ENSG00000196839 / ENSRNOG00000010265
ADAD1 / Q96M93 / ENSMUSG00000027719 / ENSG00000164113 / ENSRNOG00000017246
ADAL / Q6DHV7 / ENSMUSG00000027259 / ENSG00000168803 / ENSRNOG00000012166
ADAM10 / O14672 / ENSMUSG00000054693 / ENSG00000137845 / ENSRNOG00000054257
ADAM11 / O75078 / ENSMUSG00000020926 / ENSG00000073670 / ENSRNOG00000002753
ADAM15 / Q13444 / ENSMUSG00000028041 / ENSG00000143537 / ENSRNOG00000020590
ADAM17 / P78536 / ENSMUSG00000052593 / ENSG00000151694 / ENSRNOG00000060694
ADAM23 / O75077 / ENSMUSG00000025964 / ENSG00000114948 / ENSRNOG00000012424
ADAMTSL4 / Q6UY14 / ENSMUSG00000015850 / ENSG00000143382 / ENSRNOG00000049385
ADAP1 / O75689 / ENSMUSG00000056413 / ENSG00000105963 / ENSRNOG00000054033
ADAP2 / Q9NPF8 / ENSMUSG00000020709 / ENSG00000184060 / ENSRNOG00000037148
ADAR / P55265 / ENSMUSG00000027951 / ENSG00000160710 / ENSRNOG00000020744
ADCK1 / Q86TW2 / ENSMUSG00000021044 / ENSG00000063761 / ENSRNOG00000012685
ADCY5 / O95622 / ENSMUSG00000022840 / ENSG00000173175 / ENSRNOG00000002229
ADCY6 / O43306 / ENSMUSG00000022994 / ENSG00000174233 / ENSRNOG00000054757
ADCY9 / O60503 / ENSMUSG00000005580 / ENSG00000162104 / ENSRNOG00000049768
ADD1 / P35611 / ENSMUSG00000029106 / ENSG00000087274 / ENSRNOG00000013039
ADD2 / P35612 / ENSMUSG00000030000 / ENSG00000075340 / ENSRNOG00000015903
ADD3 / Q9UEY8 / ENSMUSG00000025026 / ENSG00000148700 / ENSRNOG00000012820
ADGRF5 / Q8IZF2 / ENSMUSG00000056492 / ENSG00000069122 / ENSRNOG00000011154
ADGRG1 / Q9Y653 / ENSMUSG00000031785 / ENSG00000205336 / ENSRNOG00000014963
ADGRL2 / O95490 / ENSMUSG00000028184 / ENSG00000117114 / ENSRNOG00000032660
ADGRL3 / Q9HAR2 / ENSMUSG00000037605 / ENSG00000150471 / ENSRNOG00000030149
Homologs
Brain
Heart
Kidney
Liver
Lung
Testis
Organs
Bins
1
2
3
4
5
Species
Human
Mouse
Rat

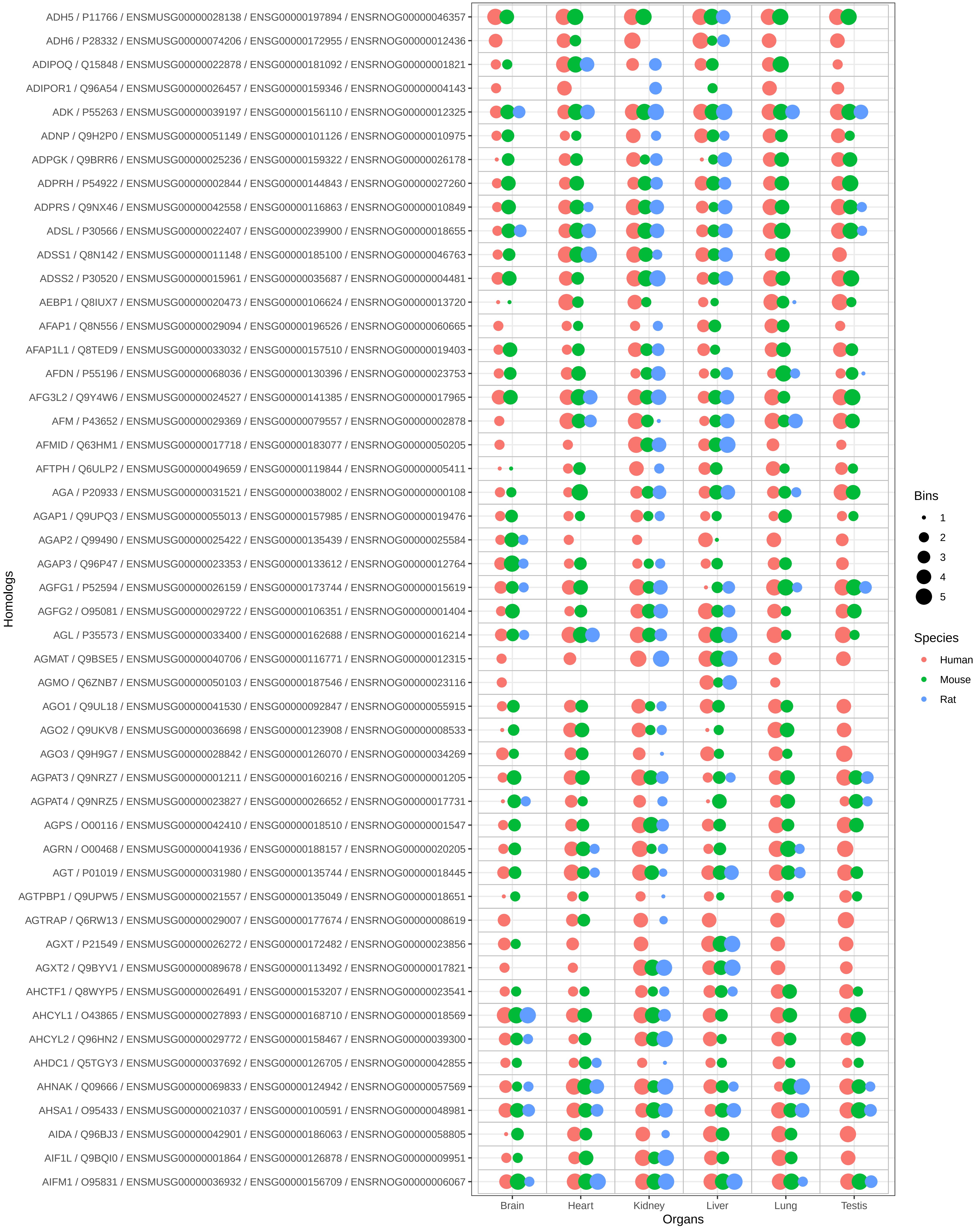
ADH5 / P11766 / ENSMUSG00000028138 / ENSG00000197894 / ENSRNOG00000046357
ADH6 / P28332 / ENSMUSG00000074206 / ENSG00000172955 / ENSRNOG00000012436
ADIPOQ / Q15848 / ENSMUSG00000022878 / ENSG00000181092 / ENSRNOG00000001821
ADIPOR1 / Q96A54 / ENSMUSG00000026457 / ENSG00000159346 / ENSRNOG00000004143
ADK / P55263 / ENSMUSG00000039197 / ENSG00000156110 / ENSRNOG00000012325
ADNP / Q9H2P0 / ENSMUSG00000051149 / ENSG00000101126 / ENSRNOG00000010975
ADPGK / Q9BRR6 / ENSMUSG00000025236 / ENSG00000159322 / ENSRNOG00000026178
ADPRH / P54922 / ENSMUSG00000002844 / ENSG00000144843 / ENSRNOG00000027260
ADPRS / Q9NX46 / ENSMUSG00000042558 / ENSG00000116863 / ENSRNOG00000010849
ADSL / P30566 / ENSMUSG00000022407 / ENSG00000239900 / ENSRNOG00000018655
ADSS1 / Q8N142 / ENSMUSG00000011148 / ENSG00000185100 / ENSRNOG00000046763
ADSS2 / P30520 / ENSMUSG00000015961 / ENSG00000035687 / ENSRNOG00000004481
AEBP1 / Q8IUX7 / ENSMUSG00000020473 / ENSG00000106624 / ENSRNOG00000013720
AFAP1 / Q8N556 / ENSMUSG00000029094 / ENSG00000196526 / ENSRNOG00000060665
AFAP1L1 / Q8TED9 / ENSMUSG00000033032 / ENSG00000157510 / ENSRNOG00000019403
AFDN / P55196 / ENSMUSG00000068036 / ENSG00000130396 / ENSRNOG00000023753
AFG3L2 / Q9Y4W6 / ENSMUSG00000024527 / ENSG00000141385 / ENSRNOG00000017965
AFM / P43652 / ENSMUSG00000029369 / ENSG00000079557 / ENSRNOG00000002878
AFMID / Q63HM1 / ENSMUSG00000017718 / ENSG00000183077 / ENSRNOG00000050205
AFTPH / Q6ULP2 / ENSMUSG00000049659 / ENSG00000119844 / ENSRNOG00000005411
AGA / P20933 / ENSMUSG00000031521 / ENSG00000038002 / ENSRNOG00000000108
AGAP1 / Q9UPQ3 / ENSMUSG00000055013 / ENSG00000157985 / ENSRNOG00000019476
AGAP2 / Q99490 / ENSMUSG00000025422 / ENSG00000135439 / ENSRNOG00000025584
AGAP3 / Q96P47 / ENSMUSG00000023353 / ENSG00000133612 / ENSRNOG00000012764
AGFG1 / P52594 / ENSMUSG00000026159 / ENSG00000173744 / ENSRNOG00000015619
AGFG2 / O95081 / ENSMUSG00000029722 / ENSG00000106351 / ENSRNOG00000001404
AGL / P35573 / ENSMUSG00000033400 / ENSG00000162688 / ENSRNOG00000016214
AGMAT / Q9BSE5 / ENSMUSG00000040706 / ENSG00000116771 / ENSRNOG00000012315
AGMO / Q6ZNB7 / ENSMUSG00000050103 / ENSG00000187546 / ENSRNOG00000023116
AGO1 / Q9UL18 / ENSMUSG00000041530 / ENSG00000092847 / ENSRNOG00000055915
AGO2 / Q9UKV8 / ENSMUSG00000036698 / ENSG00000123908 / ENSRNOG00000008533
AGO3 / Q9H9G7 / ENSMUSG00000028842 / ENSG00000126070 / ENSRNOG00000034269
AGPAT3 / Q9NRZ7 / ENSMUSG00000001211 / ENSG00000160216 / ENSRNOG00000001205
AGPAT4 / Q9NRZ5 / ENSMUSG00000023827 / ENSG00000026652 / ENSRNOG00000017731
AGPS / O00116 / ENSMUSG00000042410 / ENSG00000018510 / ENSRNOG00000001547
AGRN / O00468 / ENSMUSG00000041936 / ENSG00000188157 / ENSRNOG00000020205
AGT / P01019 / ENSMUSG00000031980 / ENSG00000135744 / ENSRNOG00000018445
AGTPBP1 / Q9UPW5 / ENSMUSG00000021557 / ENSG00000135049 / ENSRNOG00000018651
AGTRAP / Q6RW13 / ENSMUSG00000029007 / ENSG00000177674 / ENSRNOG00000008619
AGXT / P21549 / ENSMUSG00000026272 / ENSG00000172482 / ENSRNOG00000023856
AGXT2 / Q9BYV1 / ENSMUSG00000089678 / ENSG00000113492 / ENSRNOG00000017821
AHCTF1 / Q8WYP5 / ENSMUSG00000026491 / ENSG00000153207 / ENSRNOG00000023541
AHCYL1 / O43865 / ENSMUSG00000027893 / ENSG00000168710 / ENSRNOG00000018569
AHCYL2 / Q96HN2 / ENSMUSG00000029772 / ENSG00000158467 / ENSRNOG00000039300
AHDC1 / Q5TGY3 / ENSMUSG00000037692 / ENSG00000126705 / ENSRNOG00000042855
AHNAK / Q09666 / ENSMUSG00000069833 / ENSG00000124942 / ENSRNOG00000057569
AHSA1 / O95433 / ENSMUSG00000021037 / ENSG00000100591 / ENSRNOG00000048981
AIDA / Q96BJ3 / ENSMUSG00000042901 / ENSG00000186063 / ENSRNOG00000058805
AIF1L / Q9BQI0 / ENSMUSG00000001864 / ENSG00000126878 / ENSRNOG00000009951
AIFM1 / O95831 / ENSMUSG00000036932 / ENSG00000156709 / ENSRNOG00000006067
Homologs
Brain
Heart
Kidney
Liver
Lung
Testis
Organs
Bins
1
2
3
4
5
Species
Human
Mouse
Rat

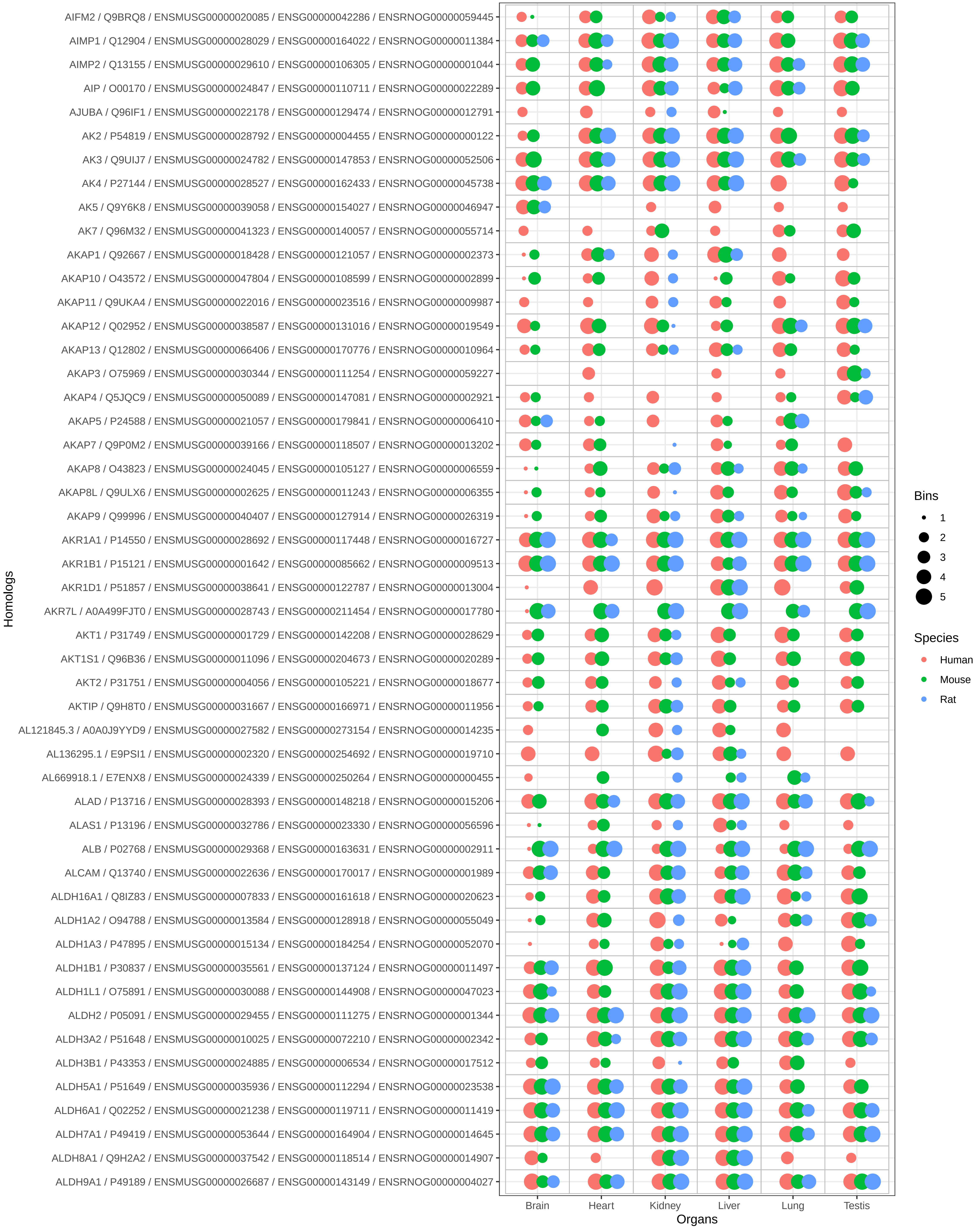
AIFM2 / Q9BRQ8 / ENSMUSG00000020085 / ENSG00000042286 / ENSRNOG00000059445
AIMP1 / Q12904 / ENSMUSG00000028029 / ENSG00000164022 / ENSRNOG00000011384
AIMP2 / Q13155 / ENSMUSG00000029610 / ENSG00000106305 / ENSRNOG00000001044
AIP / O00170 / ENSMUSG00000024847 / ENSG00000110711 / ENSRNOG00000022289
AJUBA / Q96IF1 / ENSMUSG00000022178 / ENSG00000129474 / ENSRNOG00000012791
AK2 / P54819 / ENSMUSG00000028792 / ENSG00000004455 / ENSRNOG00000000122
AK3 / Q9UIJ7 / ENSMUSG00000024782 / ENSG00000147853 / ENSRNOG00000052506
AK4 / P27144 / ENSMUSG00000028527 / ENSG00000162433 / ENSRNOG00000045738
AK5 / Q9Y6K8 / ENSMUSG00000039058 / ENSG00000154027 / ENSRNOG00000046947
AK7 / Q96M32 / ENSMUSG00000041323 / ENSG00000140057 / ENSRNOG00000055714
AKAP1 / Q92667 / ENSMUSG00000018428 / ENSG00000121057 / ENSRNOG00000002373
AKAP10 / O43572 / ENSMUSG00000047804 / ENSG00000108599 / ENSRNOG00000002899
AKAP11 / Q9UKA4 / ENSMUSG00000022016 / ENSG00000023516 / ENSRNOG00000009987
AKAP12 / Q02952 / ENSMUSG00000038587 / ENSG00000131016 / ENSRNOG00000019549
AKAP13 / Q12802 / ENSMUSG00000066406 / ENSG00000170776 / ENSRNOG00000010964
AKAP3 / O75969 / ENSMUSG00000030344 / ENSG00000111254 / ENSRNOG00000059227
AKAP4 / Q5JQC9 / ENSMUSG00000050089 / ENSG00000147081 / ENSRNOG00000002921
AKAP5 / P24588 / ENSMUSG00000021057 / ENSG00000179841 / ENSRNOG00000006410
AKAP7 / Q9P0M2 / ENSMUSG00000039166 / ENSG00000118507 / ENSRNOG00000013202
AKAP8 / O43823 / ENSMUSG00000024045 / ENSG00000105127 / ENSRNOG00000006559
AKAP8L / Q9ULX6 / ENSMUSG00000002625 / ENSG00000011243 / ENSRNOG00000006355
AKAP9 / Q99996 / ENSMUSG00000040407 / ENSG00000127914 / ENSRNOG00000026319
AKR1A1 / P14550 / ENSMUSG00000028692 / ENSG00000117448 / ENSRNOG00000016727
AKR1B1 / P15121 / ENSMUSG00000001642 / ENSG00000085662 / ENSRNOG00000009513
AKR1D1 / P51857 / ENSMUSG00000038641 / ENSG00000122787 / ENSRNOG00000013004
AKR7L / A0A499FJT0 / ENSMUSG00000028743 / ENSG00000211454 / ENSRNOG00000017780
AKT1 / P31749 / ENSMUSG00000001729 / ENSG00000142208 / ENSRNOG00000028629
AKT1S1 / Q96B36 / ENSMUSG00000011096 / ENSG00000204673 / ENSRNOG00000020289
AKT2 / P31751 / ENSMUSG00000004056 / ENSG00000105221 / ENSRNOG00000018677
AKTIP / Q9H8T0 / ENSMUSG00000031667 / ENSG00000166971 / ENSRNOG00000011956
AL121845.3 / A0A0J9YYD9 / ENSMUSG00000027582 / ENSG00000273154 / ENSRNOG00000014235
AL136295.1 / E9PSI1 / ENSMUSG00000002320 / ENSG00000254692 / ENSRNOG00000019710
AL669918.1 / E7ENX8 / ENSMUSG00000024339 / ENSG00000250264 / ENSRNOG00000000455
ALAD / P13716 / ENSMUSG00000028393 / ENSG00000148218 / ENSRNOG00000015206
ALAS1 / P13196 / ENSMUSG00000032786 / ENSG00000023330 / ENSRNOG00000056596
ALB / P02768 / ENSMUSG00000029368 / ENSG00000163631 / ENSRNOG00000002911
ALCAM / Q13740 / ENSMUSG00000022636 / ENSG00000170017 / ENSRNOG00000001989
ALDH16A1 / Q8IZ83 / ENSMUSG00000007833 / ENSG00000161618 / ENSRNOG00000020623
ALDH1A2 / O94788 / ENSMUSG00000013584 / ENSG00000128918 / ENSRNOG00000055049
ALDH1A3 / P47895 / ENSMUSG00000015134 / ENSG00000184254 / ENSRNOG00000052070
ALDH1B1 / P30837 / ENSMUSG00000035561 / ENSG00000137124 / ENSRNOG00000011497
ALDH1L1 / O75891 / ENSMUSG00000030088 / ENSG00000144908 / ENSRNOG00000047023
ALDH2 / P05091 / ENSMUSG00000029455 / ENSG00000111275 / ENSRNOG00000001344
ALDH3A2 / P51648 / ENSMUSG00000010025 / ENSG00000072210 / ENSRNOG00000002342
ALDH3B1 / P43353 / ENSMUSG00000024885 / ENSG00000006534 / ENSRNOG00000017512
ALDH5A1 / P51649 / ENSMUSG00000035936 / ENSG00000112294 / ENSRNOG00000023538
ALDH6A1 / Q02252 / ENSMUSG00000021238 / ENSG00000119711 / ENSRNOG00000011419
ALDH7A1 / P49419 / ENSMUSG00000053644 / ENSG00000164904 / ENSRNOG00000014645
ALDH8A1 / Q9H2A2 / ENSMUSG00000037542 / ENSG00000118514 / ENSRNOG00000014907
ALDH9A1 / P49189 / ENSMUSG00000026687 / ENSG00000143149 / ENSRNOG00000004027
Homologs
Brain
Heart
Kidney
Liver
Lung
Testis
Organs
Bins
1
2
3
4
5
Species
Human
Mouse
Rat

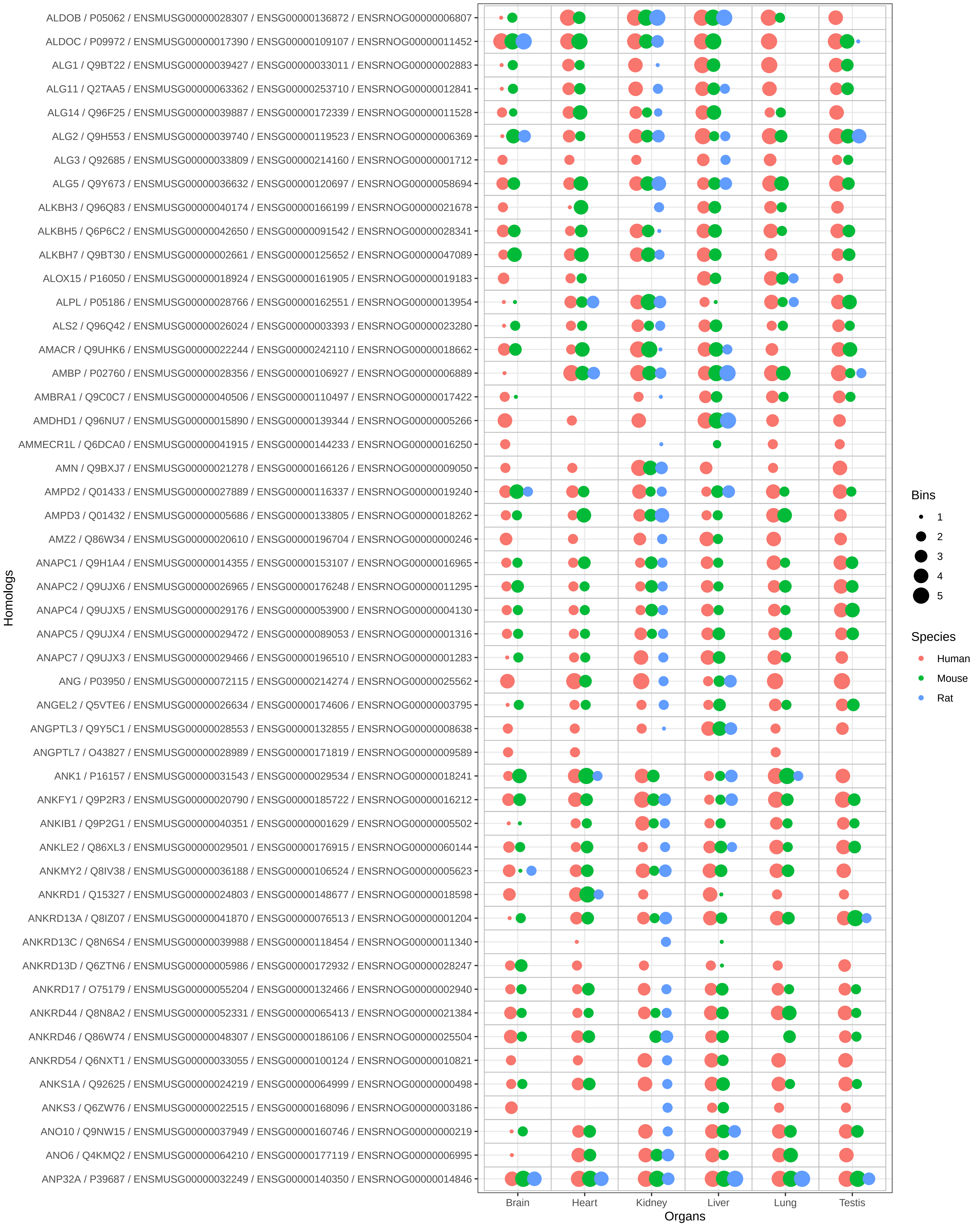
ALDOB / P05062 / ENSMUSG00000028307 / ENSG00000136872 / ENSRNOG00000006807
ALDOC / P09972 / ENSMUSG00000017390 / ENSG00000109107 / ENSRNOG00000011452
ALG1 / Q9BT22 / ENSMUSG00000039427 / ENSG00000033011 / ENSRNOG00000002883
ALG11 / Q2TAA5 / ENSMUSG00000063362 / ENSG00000253710 / ENSRNOG00000012841
ALG14 / Q96F25 / ENSMUSG00000039887 / ENSG00000172339 / ENSRNOG00000011528
ALG2 / Q9H553 / ENSMUSG00000039740 / ENSG00000119523 / ENSRNOG00000006369
ALG3 / Q92685 / ENSMUSG00000033809 / ENSG00000214160 / ENSRNOG00000001712
ALG5 / Q9Y673 / ENSMUSG00000036632 / ENSG00000120697 / ENSRNOG00000058694
ALKBH3 / Q96Q83 / ENSMUSG00000040174 / ENSG00000166199 / ENSRNOG00000021678
ALKBH5 / Q6P6C2 / ENSMUSG00000042650 / ENSG00000091542 / ENSRNOG00000028341
ALKBH7 / Q9BT30 / ENSMUSG00000002661 / ENSG00000125652 / ENSRNOG00000047089
ALOX15 / P16050 / ENSMUSG00000018924 / ENSG00000161905 / ENSRNOG00000019183
ALPL / P05186 / ENSMUSG00000028766 / ENSG00000162551 / ENSRNOG00000013954
ALS2 / Q96Q42 / ENSMUSG00000026024 / ENSG00000003393 / ENSRNOG00000023280
AMACR / Q9UHK6 / ENSMUSG00000022244 / ENSG00000242110 / ENSRNOG00000018662
AMBP / P02760 / ENSMUSG00000028356 / ENSG00000106927 / ENSRNOG00000006889
AMBRA1 / Q9C0C7 / ENSMUSG00000040506 / ENSG00000110497 / ENSRNOG00000017422
AMDHD1 / Q96NU7 / ENSMUSG00000015890 / ENSG00000139344 / ENSRNOG00000005266
AMMECR1L / Q6DCA0 / ENSMUSG00000041915 / ENSG00000144233 / ENSRNOG00000016250
AMN / Q9BXJ7 / ENSMUSG00000021278 / ENSG00000166126 / ENSRNOG00000009050
AMPD2 / Q01433 / ENSMUSG00000027889 / ENSG00000116337 / ENSRNOG00000019240
AMPD3 / Q01432 / ENSMUSG00000005686 / ENSG00000133805 / ENSRNOG00000018262
AMZ2 / Q86W34 / ENSMUSG00000020610 / ENSG00000196704 / ENSRNOG00000000246
ANAPC1 / Q9H1A4 / ENSMUSG00000014355 / ENSG00000153107 / ENSRNOG00000016965
ANAPC2 / Q9UJX6 / ENSMUSG00000026965 / ENSG00000176248 / ENSRNOG00000011295
ANAPC4 / Q9UJX5 / ENSMUSG00000029176 / ENSG00000053900 / ENSRNOG00000004130
ANAPC5 / Q9UJX4 / ENSMUSG00000029472 / ENSG00000089053 / ENSRNOG00000001316
ANAPC7 / Q9UJX3 / ENSMUSG00000029466 / ENSG00000196510 / ENSRNOG00000001283
ANG / P03950 / ENSMUSG00000072115 / ENSG00000214274 / ENSRNOG00000025562
ANGEL2 / Q5VTE6 / ENSMUSG00000026634 / ENSG00000174606 / ENSRNOG00000003795
ANGPTL3 / Q9Y5C1 / ENSMUSG00000028553 / ENSG00000132855 / ENSRNOG00000008638
ANGPTL7 / O43827 / ENSMUSG00000028989 / ENSG00000171819 / ENSRNOG00000009589
ANK1 / P16157 / ENSMUSG00000031543 / ENSG00000029534 / ENSRNOG00000018241
ANKFY1 / Q9P2R3 / ENSMUSG00000020790 / ENSG00000185722 / ENSRNOG00000016212
ANKIB1 / Q9P2G1 / ENSMUSG00000040351 / ENSG00000001629 / ENSRNOG00000005502
ANKLE2 / Q86XL3 / ENSMUSG00000029501 / ENSG00000176915 / ENSRNOG00000060144
ANKMY2 / Q8IV38 / ENSMUSG00000036188 / ENSG00000106524 / ENSRNOG00000005623
ANKRD1 / Q15327 / ENSMUSG00000024803 / ENSG00000148677 / ENSRNOG00000018598
ANKRD13A / Q8IZ07 / ENSMUSG00000041870 / ENSG00000076513 / ENSRNOG00000001204
ANKRD13C / Q8N6S4 / ENSMUSG00000039988 / ENSG00000118454 / ENSRNOG00000011340
ANKRD13D / Q6ZTN6 / ENSMUSG00000005986 / ENSG00000172932 / ENSRNOG00000028247
ANKRD17 / O75179 / ENSMUSG00000055204 / ENSG00000132466 / ENSRNOG00000002940
ANKRD44 / Q8N8A2 / ENSMUSG00000052331 / ENSG00000065413 / ENSRNOG00000021384
ANKRD46 / Q86W74 / ENSMUSG00000048307 / ENSG00000186106 / ENSRNOG00000025504
ANKRD54 / Q6NXT1 / ENSMUSG00000033055 / ENSG00000100124 / ENSRNOG00000010821
ANKS1A / Q92625 / ENSMUSG00000024219 / ENSG00000064999 / ENSRNOG00000000498
ANKS3 / Q6ZW76 / ENSMUSG00000022515 / ENSG00000168096 / ENSRNOG00000003186
ANO10 / Q9NW15 / ENSMUSG00000037949 / ENSG00000160746 / ENSRNOG00000000219
ANO6 / Q4KMQ2 / ENSMUSG00000064210 / ENSG00000177119 / ENSRNOG00000006995
ANP32A / P39687 / ENSMUSG00000032249 / ENSG00000140350 / ENSRNOG00000014846
Homologs
Brain
Heart
Kidney
Liver
Lung
Testis
Organs
Bins
1
2
3
4
5
Species
Human
Mouse
Rat

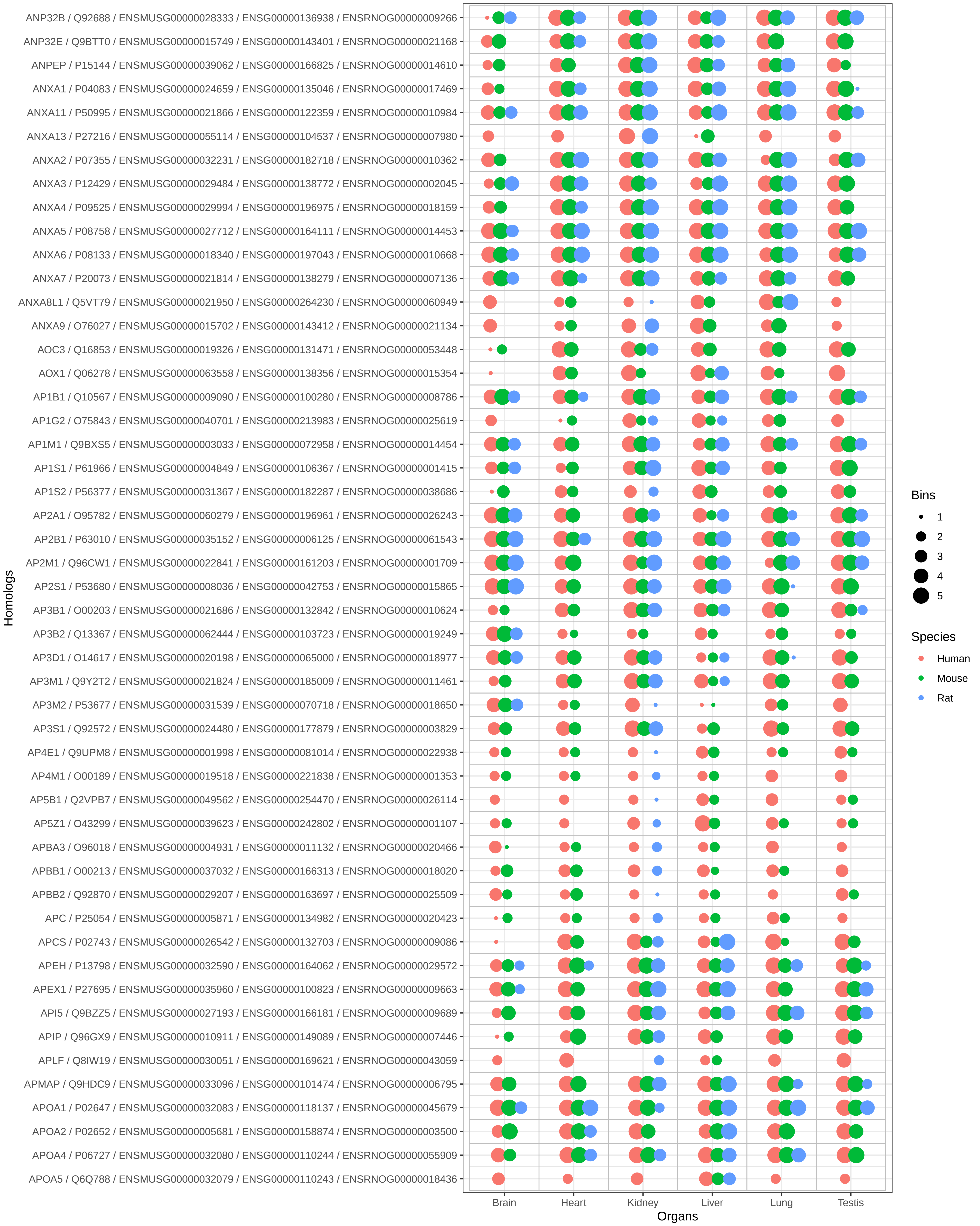

ANP32B / Q92688 / ENSMUSG00000028333 / ENSG00000136938 / ENSRNOG00000009266
ANP32E / Q9BTT0 / ENSMUSG00000015749 / ENSG00000143401 / ENSRNOG00000021168
ANPEP / P15144 / ENSMUSG00000039062 / ENSG00000166825 / ENSRNOG00000014610
ANXA1 / P04083 / ENSMUSG00000024659 / ENSG00000135046 / ENSRNOG00000017469
ANXA11 / P50995 / ENSMUSG00000021866 / ENSG00000122359 / ENSRNOG00000010984
ANXA13 / P27216 / ENSMUSG00000055114 / ENSG00000104537 / ENSRNOG00000007980
ANXA2 / P07355 / ENSMUSG00000032231 / ENSG00000182718 / ENSRNOG00000010362
ANXA3 / P12429 / ENSMUSG00000029484 / ENSG00000138772 / ENSRNOG00000002045
ANXA4 / P09525 / ENSMUSG00000029994 / ENSG00000196975 / ENSRNOG00000018159
ANXA5 / P08758 / ENSMUSG00000027712 / ENSG00000164111 / ENSRNOG00000014453
ANXA6 / P08133 / ENSMUSG00000018340 / ENSG00000197043 / ENSRNOG00000010668
ANXA7 / P20073 / ENSMUSG00000021814 / ENSG00000138279 / ENSRNOG00000007136
ANXA8L1 / Q5VT79 / ENSMUSG00000021950 / ENSG00000264230 / ENSRNOG00000060949
ANXA9 / O76027 / ENSMUSG00000015702 / ENSG00000143412 / ENSRNOG00000021134
AOC3 / Q16853 / ENSMUSG00000019326 / ENSG00000131471 / ENSRNOG00000053448
AOX1 / Q06278 / ENSMUSG00000063558 / ENSG00000138356 / ENSRNOG00000015354
AP1B1 / Q10567 / ENSMUSG00000009090 / ENSG00000100280 / ENSRNOG00000008786
AP1G2 / O75843 / ENSMUSG00000040701 / ENSG00000213983 / ENSRNOG00000025619
AP1M1 / Q9BXS5 / ENSMUSG00000003033 / ENSG00000072958 / ENSRNOG00000014454
AP1S1 / P61966 / ENSMUSG00000004849 / ENSG00000106367 / ENSRNOG00000001415
AP1S2 / P56377 / ENSMUSG00000031367 / ENSG00000182287 / ENSRNOG00000038686
AP2A1 / O95782 / ENSMUSG00000060279 / ENSG00000196961 / ENSRNOG00000026243
AP2B1 / P63010 / ENSMUSG00000035152 / ENSG00000006125 / ENSRNOG00000061543
AP2M1 / Q96CW1 / ENSMUSG00000022841 / ENSG00000161203 / ENSRNOG00000001709
AP2S1 / P53680 / ENSMUSG00000008036 / ENSG00000042753 / ENSRNOG00000015865
AP3B1 / O00203 / ENSMUSG00000021686 / ENSG00000132842 / ENSRNOG00000010624
AP3B2 / Q13367 / ENSMUSG00000062444 / ENSG00000103723 / ENSRNOG00000019249
AP3D1 / O14617 / ENSMUSG00000020198 / ENSG00000065000 / ENSRNOG00000018977
AP3M1 / Q9Y2T2 / ENSMUSG00000021824 / ENSG00000185009 / ENSRNOG00000011461
AP3M2 / P53677 / ENSMUSG00000031539 / ENSG00000070718 / ENSRNOG00000018650
AP3S1 / Q92572 / ENSMUSG00000024480 / ENSG00000177879 / ENSRNOG00000003829
AP4E1 / Q9UPM8 / ENSMUSG00000001998 / ENSG00000081014 / ENSRNOG00000022938
AP4M1 / O00189 / ENSMUSG00000019518 / ENSG00000221838 / ENSRNOG00000001353
AP5B1 / Q2VPB7 / ENSMUSG00000049562 / ENSG00000254470 / ENSRNOG00000026114
AP5Z1 / O43299 / ENSMUSG00000039623 / ENSG00000242802 / ENSRNOG00000001107
APBA3 / O96018 / ENSMUSG00000004931 / ENSG00000011132 / ENSRNOG00000020466
APBB1 / O00213 / ENSMUSG00000037032 / ENSG00000166313 / ENSRNOG00000018020
APBB2 / Q92870 / ENSMUSG00000029207 / ENSG00000163697 / ENSRNOG00000025509
APC / P25054 / ENSMUSG00000005871 / ENSG00000134982 / ENSRNOG00000020423
APCS / P02743 / ENSMUSG00000026542 / ENSG00000132703 / ENSRNOG00000009086
APEH / P13798 / ENSMUSG00000032590 / ENSG00000164062 / ENSRNOG00000029572
APEX1 / P27695 / ENSMUSG00000035960 / ENSG00000100823 / ENSRNOG00000009663
API5 / Q9BZZ5 / ENSMUSG00000027193 / ENSG00000166181 / ENSRNOG00000009689
APIP / Q96GX9 / ENSMUSG00000010911 / ENSG00000149089 / ENSRNOG00000007446
APLF / Q8IW19 / ENSMUSG00000030051 / ENSG00000169621 / ENSRNOG00000043059
APMAP / Q9HDC9 / ENSMUSG00000033096 / ENSG00000101474 / ENSRNOG00000006795
APOA1 / P02647 / ENSMUSG00000032083 / ENSG00000118137 / ENSRNOG00000045679
APOA2 / P02652 / ENSMUSG00000005681 / ENSG00000158874 / ENSRNOG00000003500
APOA4 / P06727 / ENSMUSG00000032080 / ENSG00000110244 / ENSRNOG00000055909
APOA5 / Q6Q788 / ENSMUSG00000032079 / ENSG00000110243 / ENSRNOG00000018436
Homologs
Brain
Heart
Kidney
Liver
Lung
Testis
Organs
Bins
1
2
3
4
5
Species
Human
Mouse
Rat

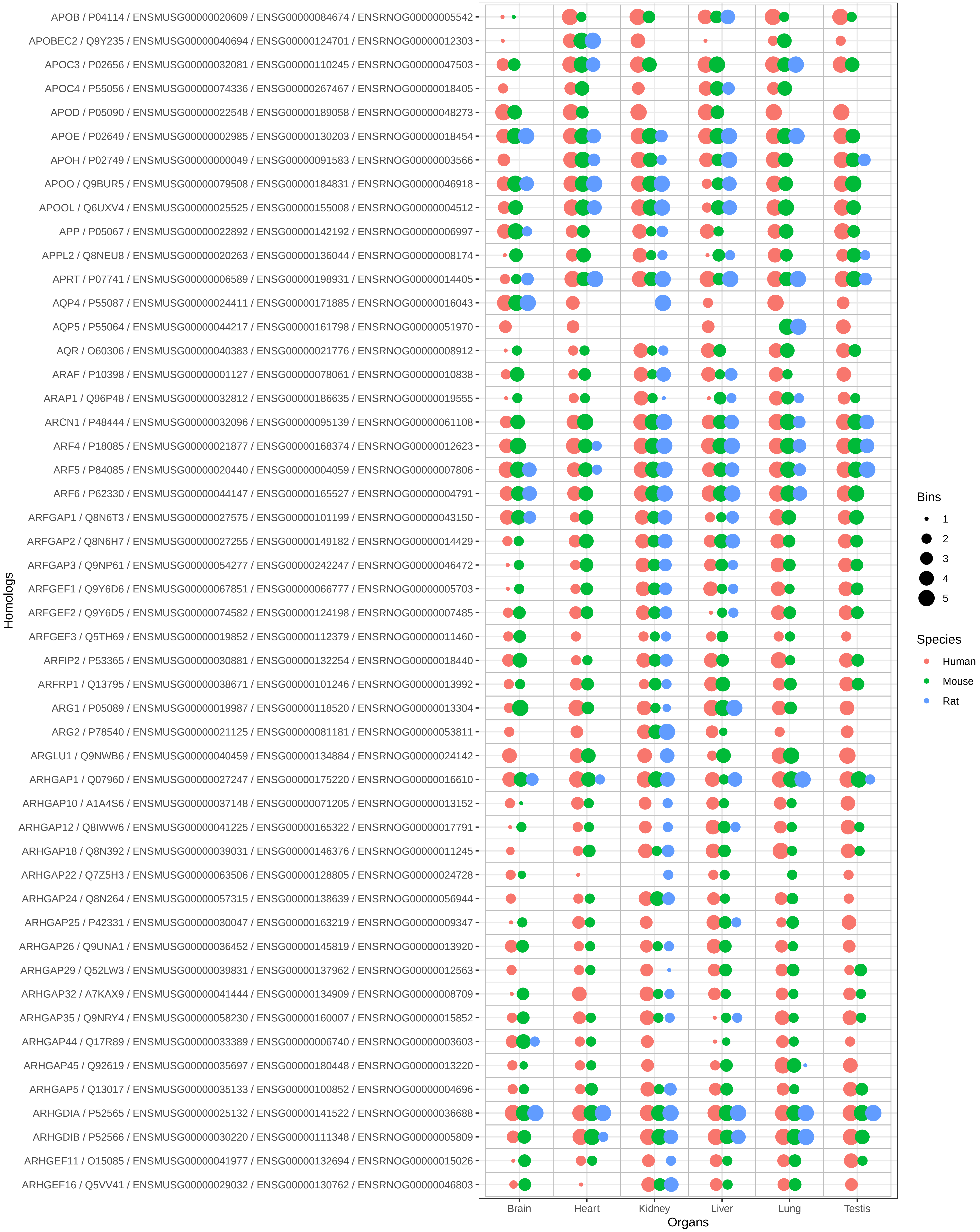

APOB / P04114 / ENSMUSG00000020609 / ENSG00000084674 / ENSRNOG00000005542
APOBEC2 / Q9Y235 / ENSMUSG00000040694 / ENSG00000124701 / ENSRNOG00000012303
APOC3 / P02656 / ENSMUSG00000032081 / ENSG00000110245 / ENSRNOG00000047503
APOC4 / P55056 / ENSMUSG00000074336 / ENSG00000267467 / ENSRNOG00000018405
APOD / P05090 / ENSMUSG00000022548 / ENSG00000189058 / ENSRNOG00000048273
APOE / P02649 / ENSMUSG00000002985 / ENSG00000130203 / ENSRNOG00000018454
APOH / P02749 / ENSMUSG00000000049 / ENSG00000091583 / ENSRNOG00000003566
APOO / Q9BUR5 / ENSMUSG00000079508 / ENSG00000184831 / ENSRNOG00000046918
APOOL / Q6UXV4 / ENSMUSG00000025525 / ENSG00000155008 / ENSRNOG00000004512
APP / P05067 / ENSMUSG00000022892 / ENSG00000142192 / ENSRNOG00000006997
APPL2 / Q8NEU8 / ENSMUSG00000020263 / ENSG00000136044 / ENSRNOG00000008174
APRT / P07741 / ENSMUSG00000006589 / ENSG00000198931 / ENSRNOG00000014405
AQP4 / P55087 / ENSMUSG00000024411 / ENSG00000171885 / ENSRNOG00000016043
AQP5 / P55064 / ENSMUSG00000044217 / ENSG00000161798 / ENSRNOG00000051970
AQR / O60306 / ENSMUSG00000040383 / ENSG00000021776 / ENSRNOG00000008912
ARAF / P10398 / ENSMUSG00000001127 / ENSG00000078061 / ENSRNOG00000010838
ARAP1 / Q96P48 / ENSMUSG00000032812 / ENSG00000186635 / ENSRNOG00000019555
ARCN1 / P48444 / ENSMUSG00000032096 / ENSG00000095139 / ENSRNOG00000061108
ARF4 / P18085 / ENSMUSG00000021877 / ENSG00000168374 / ENSRNOG00000012623
ARF5 / P84085 / ENSMUSG00000020440 / ENSG00000004059 / ENSRNOG00000007806
ARF6 / P62330 / ENSMUSG00000044147 / ENSG00000165527 / ENSRNOG00000004791
ARFGAP1 / Q8N6T3 / ENSMUSG00000027575 / ENSG00000101199 / ENSRNOG00000043150
ARFGAP2 / Q8N6H7 / ENSMUSG00000027255 / ENSG00000149182 / ENSRNOG00000014429
ARFGAP3 / Q9NP61 / ENSMUSG00000054277 / ENSG00000242247 / ENSRNOG00000046472
ARFGEF1 / Q9Y6D6 / ENSMUSG00000067851 / ENSG00000066777 / ENSRNOG00000005703
ARFGEF2 / Q9Y6D5 / ENSMUSG00000074582 / ENSG00000124198 / ENSRNOG00000007485
ARFGEF3 / Q5TH69 / ENSMUSG00000019852 / ENSG00000112379 / ENSRNOG00000011460
ARFIP2 / P53365 / ENSMUSG00000030881 / ENSG00000132254 / ENSRNOG00000018440
ARFRP1 / Q13795 / ENSMUSG00000038671 / ENSG00000101246 / ENSRNOG00000013992
ARG1 / P05089 / ENSMUSG00000019987 / ENSG00000118520 / ENSRNOG00000013304
ARG2 / P78540 / ENSMUSG00000021125 / ENSG00000081181 / ENSRNOG00000053811
ARGLU1 / Q9NWB6 / ENSMUSG00000040459 / ENSG00000134884 / ENSRNOG00000024142
ARHGAP1 / Q07960 / ENSMUSG00000027247 / ENSG00000175220 / ENSRNOG00000016610
ARHGAP10 / A1A4S6 / ENSMUSG00000037148 / ENSG00000071205 / ENSRNOG00000013152
ARHGAP12 / Q8IWW6 / ENSMUSG00000041225 / ENSG00000165322 / ENSRNOG00000017791
ARHGAP18 / Q8N392 / ENSMUSG00000039031 / ENSG00000146376 / ENSRNOG00000011245
ARHGAP22 / Q7Z5H3 / ENSMUSG00000063506 / ENSG00000128805 / ENSRNOG00000024728
ARHGAP24 / Q8N264 / ENSMUSG00000057315 / ENSG00000138639 / ENSRNOG00000056944
ARHGAP25 / P42331 / ENSMUSG00000030047 / ENSG00000163219 / ENSRNOG00000009347
ARHGAP26 / Q9UNA1 / ENSMUSG00000036452 / ENSG00000145819 / ENSRNOG00000013920
ARHGAP29 / Q52LW3 / ENSMUSG00000039831 / ENSG00000137962 / ENSRNOG00000012563
ARHGAP32 / A7KAX9 / ENSMUSG00000041444 / ENSG00000134909 / ENSRNOG00000008709
ARHGAP35 / Q9NRY4 / ENSMUSG00000058230 / ENSG00000160007 / ENSRNOG00000015852
ARHGAP44 / Q17R89 / ENSMUSG00000033389 / ENSG00000006740 / ENSRNOG00000003603
ARHGAP45 / Q92619 / ENSMUSG00000035697 / ENSG00000180448 / ENSRNOG00000013220
ARHGAP5 / Q13017 / ENSMUSG00000035133 / ENSG00000100852 / ENSRNOG00000004696
ARHGDIA / P52565 / ENSMUSG00000025132 / ENSG00000141522 / ENSRNOG00000036688
ARHGDIB / P52566 / ENSMUSG00000030220 / ENSG00000111348 / ENSRNOG00000005809
ARHGEF11 / O15085 / ENSMUSG00000041977 / ENSG00000132694 / ENSRNOG00000015026
ARHGEF16 / Q5VV41 / ENSMUSG00000029032 / ENSG00000130762 / ENSRNOG00000046803
Homologs
Brain
Heart
Kidney
Liver
Lung
Testis
Organs
Bins
1
2
3
4
5
Species
Human
Mouse
Rat

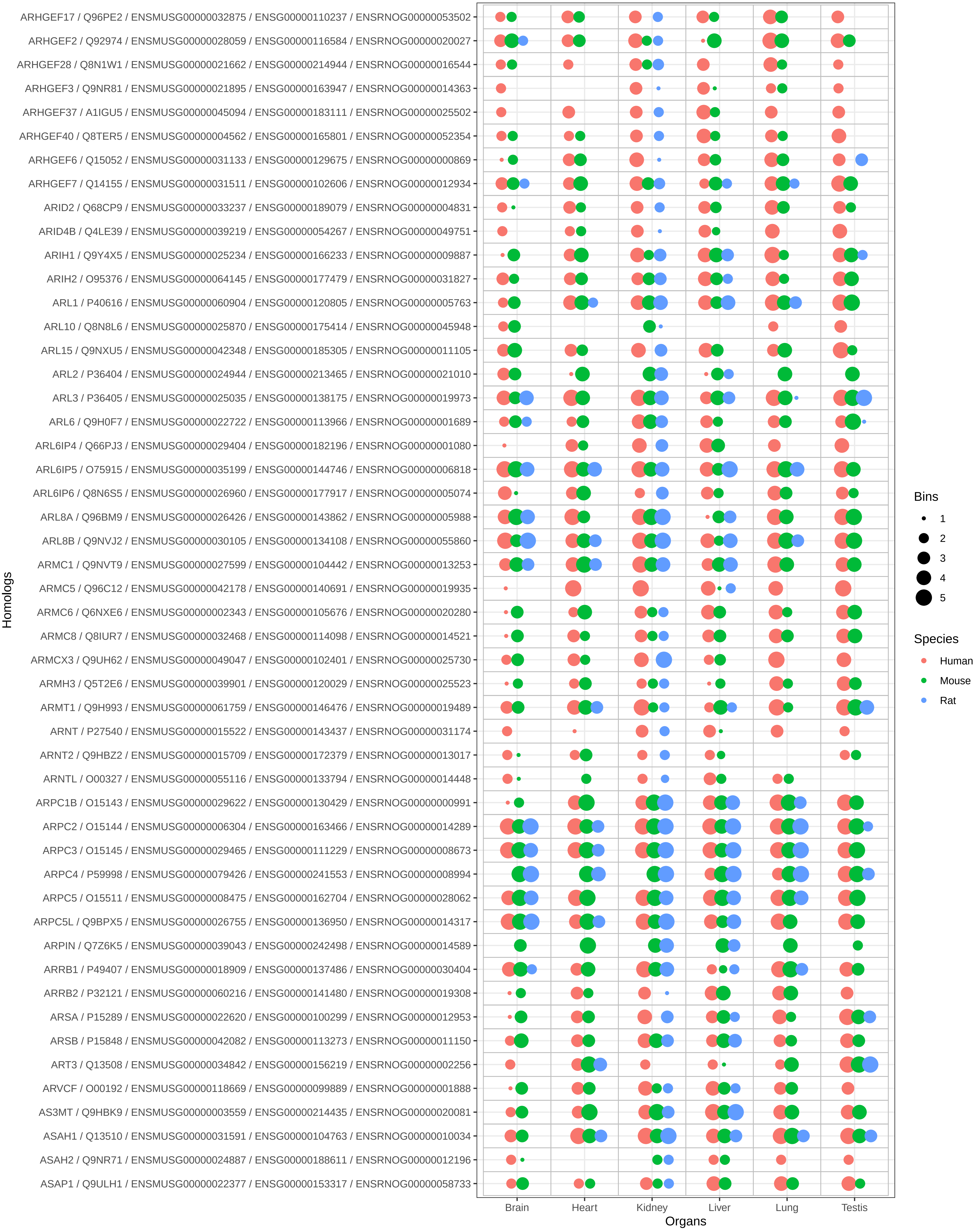
ARHGEF17 / Q96PE2 / ENSMUSG00000032875 / ENSG00000110237 / ENSRNOG00000053502
ARHGEF2 / Q92974 / ENSMUSG00000028059 / ENSG00000116584 / ENSRNOG00000020027
ARHGEF28 / Q8N1W1 / ENSMUSG00000021662 / ENSG00000214944 / ENSRNOG00000016544
ARHGEF3 / Q9NR81 / ENSMUSG00000021895 / ENSG00000163947 / ENSRNOG00000014363
ARHGEF37 / A1IGU5 / ENSMUSG00000045094 / ENSG00000183111 / ENSRNOG00000025502
ARHGEF40 / Q8TER5 / ENSMUSG00000004562 / ENSG00000165801 / ENSRNOG00000052354
ARHGEF6 / Q15052 / ENSMUSG00000031133 / ENSG00000129675 / ENSRNOG00000000869
ARHGEF7 / Q14155 / ENSMUSG00000031511 / ENSG00000102606 / ENSRNOG00000012934
ARID2 / Q68CP9 / ENSMUSG00000033237 / ENSG00000189079 / ENSRNOG00000004831
ARID4B / Q4LE39 / ENSMUSG00000039219 / ENSG00000054267 / ENSRNOG00000049751
ARIH1 / Q9Y4X5 / ENSMUSG00000025234 / ENSG00000166233 / ENSRNOG00000009887
ARIH2 / O95376 / ENSMUSG00000064145 / ENSG00000177479 / ENSRNOG00000031827
ARL1 / P40616 / ENSMUSG00000060904 / ENSG00000120805 / ENSRNOG00000005763
ARL10 / Q8N8L6 / ENSMUSG00000025870 / ENSG00000175414 / ENSRNOG00000045948
ARL15 / Q9NXU5 / ENSMUSG00000042348 / ENSG00000185305 / ENSRNOG00000011105
ARL2 / P36404 / ENSMUSG00000024944 / ENSG00000213465 / ENSRNOG00000021010
ARL3 / P36405 / ENSMUSG00000025035 / ENSG00000138175 / ENSRNOG00000019973
ARL6 / Q9H0F7 / ENSMUSG00000022722 / ENSG00000113966 / ENSRNOG00000001689
ARL6IP4 / Q66PJ3 / ENSMUSG00000029404 / ENSG00000182196 / ENSRNOG00000001080
ARL6IP5 / O75915 / ENSMUSG00000035199 / ENSG00000144746 / ENSRNOG00000006818
ARL6IP6 / Q8N6S5 / ENSMUSG00000026960 / ENSG00000177917 / ENSRNOG00000005074
ARL8A / Q96BM9 / ENSMUSG00000026426 / ENSG00000143862 / ENSRNOG00000005988
ARL8B / Q9NVJ2 / ENSMUSG00000030105 / ENSG00000134108 / ENSRNOG00000055860
ARMC1 / Q9NVT9 / ENSMUSG00000027599 / ENSG00000104442 / ENSRNOG00000013253
ARMC5 / Q96C12 / ENSMUSG00000042178 / ENSG00000140691 / ENSRNOG00000019935
ARMC6 / Q6NXE6 / ENSMUSG00000002343 / ENSG00000105676 / ENSRNOG00000020280
ARMC8 / Q8IUR7 / ENSMUSG00000032468 / ENSG00000114098 / ENSRNOG00000014521
ARMCX3 / Q9UH62 / ENSMUSG00000049047 / ENSG00000102401 / ENSRNOG00000025730
ARMH3 / Q5T2E6 / ENSMUSG00000039901 / ENSG00000120029 / ENSRNOG00000025523
ARMT1 / Q9H993 / ENSMUSG00000061759 / ENSG00000146476 / ENSRNOG00000019489
ARNT / P27540 / ENSMUSG00000015522 / ENSG00000143437 / ENSRNOG00000031174
ARNT2 / Q9HBZ2 / ENSMUSG00000015709 / ENSG00000172379 / ENSRNOG00000013017
ARNTL / O00327 / ENSMUSG00000055116 / ENSG00000133794 / ENSRNOG00000014448
ARPC1B / O15143 / ENSMUSG00000029622 / ENSG00000130429 / ENSRNOG00000000991
ARPC2 / O15144 / ENSMUSG00000006304 / ENSG00000163466 / ENSRNOG00000014289
ARPC3 / O15145 / ENSMUSG00000029465 / ENSG00000111229 / ENSRNOG00000008673
ARPC4 / P59998 / ENSMUSG00000079426 / ENSG00000241553 / ENSRNOG00000008994
ARPC5 / O15511 / ENSMUSG00000008475 / ENSG00000162704 / ENSRNOG00000028062
ARPC5L / Q9BPX5 / ENSMUSG00000026755 / ENSG00000136950 / ENSRNOG00000014317
ARPIN / Q7Z6K5 / ENSMUSG00000039043 / ENSG00000242498 / ENSRNOG00000014589
ARRB1 / P49407 / ENSMUSG00000018909 / ENSG00000137486 / ENSRNOG00000030404
ARRB2 / P32121 / ENSMUSG00000060216 / ENSG00000141480 / ENSRNOG00000019308
ARSA / P15289 / ENSMUSG00000022620 / ENSG00000100299 / ENSRNOG00000012953
ARSB / P15848 / ENSMUSG00000042082 / ENSG00000113273 / ENSRNOG00000011150
ART3 / Q13508 / ENSMUSG00000034842 / ENSG00000156219 / ENSRNOG00000002256
ARVCF / O00192 / ENSMUSG00000118669 / ENSG00000099889 / ENSRNOG00000001888
AS3MT / Q9HBK9 / ENSMUSG00000003559 / ENSG00000214435 / ENSRNOG00000020081
ASAH1 / Q13510 / ENSMUSG00000031591 / ENSG00000104763 / ENSRNOG00000010034
ASAH2 / Q9NR71 / ENSMUSG00000024887 / ENSG00000188611 / ENSRNOG00000012196
ASAP1 / Q9ULH1 / ENSMUSG00000022377 / ENSG00000153317 / ENSRNOG00000058733
Homologs
Brain
Heart
Kidney
Liver
Lung
Testis
Organs
Bins
1
2
3
4
5
Species
Human
Mouse
Rat

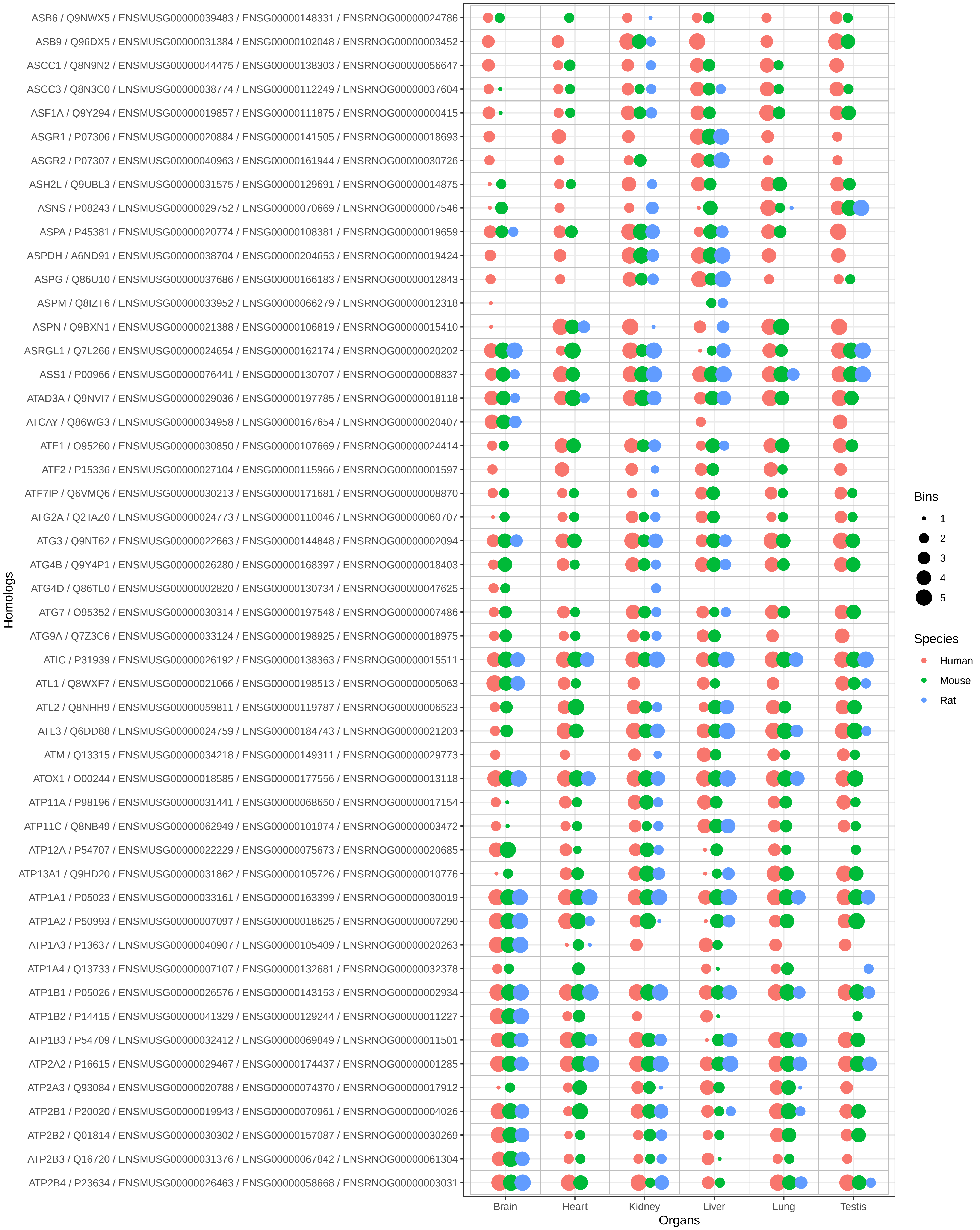
ASB6 / Q9NWX5 / ENSMUSG00000039483 / ENSG00000148331 / ENSRNOG00000024786
ASB9 / Q96DX5 / ENSMUSG00000031384 / ENSG00000102048 / ENSRNOG00000003452
ASCC1 / Q8N9N2 / ENSMUSG00000044475 / ENSG00000138303 / ENSRNOG00000056647
ASCC3 / Q8N3C0 / ENSMUSG00000038774 / ENSG00000112249 / ENSRNOG00000037604
ASF1A / Q9Y294 / ENSMUSG00000019857 / ENSG00000111875 / ENSRNOG00000000415
ASGR1 / P07306 / ENSMUSG00000020884 / ENSG00000141505 / ENSRNOG00000018693
ASGR2 / P07307 / ENSMUSG00000040963 / ENSG00000161944 / ENSRNOG00000030726
ASH2L / Q9UBL3 / ENSMUSG00000031575 / ENSG00000129691 / ENSRNOG00000014875
ASNS / P08243 / ENSMUSG00000029752 / ENSG00000070669 / ENSRNOG00000007546
ASPA / P45381 / ENSMUSG00000020774 / ENSG00000108381 / ENSRNOG00000019659
ASPDH / A6ND91 / ENSMUSG00000038704 / ENSG00000204653 / ENSRNOG00000019424
ASPG / Q86U10 / ENSMUSG00000037686 / ENSG00000166183 / ENSRNOG00000012843
ASPM / Q8IZT6 / ENSMUSG00000033952 / ENSG00000066279 / ENSRNOG00000012318
ASPN / Q9BXN1 / ENSMUSG00000021388 / ENSG00000106819 / ENSRNOG00000015410
ASRGL1 / Q7L266 / ENSMUSG00000024654 / ENSG00000162174 / ENSRNOG00000020202
ASS1 / P00966 / ENSMUSG00000076441 / ENSG00000130707 / ENSRNOG00000008837
ATAD3A / Q9NVI7 / ENSMUSG00000029036 / ENSG00000197785 / ENSRNOG00000018118
ATCAY / Q86WG3 / ENSMUSG00000034958 / ENSG00000167654 / ENSRNOG00000020407
ATE1 / O95260 / ENSMUSG00000030850 / ENSG00000107669 / ENSRNOG00000024414
ATF2 / P15336 / ENSMUSG00000027104 / ENSG00000115966 / ENSRNOG00000001597
ATF7IP / Q6VMQ6 / ENSMUSG00000030213 / ENSG00000171681 / ENSRNOG00000008870
ATG2A / Q2TAZ0 / ENSMUSG00000024773 / ENSG00000110046 / ENSRNOG00000060707
ATG3 / Q9NT62 / ENSMUSG00000022663 / ENSG00000144848 / ENSRNOG00000002094
ATG4B / Q9Y4P1 / ENSMUSG00000026280 / ENSG00000168397 / ENSRNOG00000018403
ATG4D / Q86TL0 / ENSMUSG00000002820 / ENSG00000130734 / ENSRNOG00000047625
ATG7 / O95352 / ENSMUSG00000030314 / ENSG00000197548 / ENSRNOG00000007486
ATG9A / Q7Z3C6 / ENSMUSG00000033124 / ENSG00000198925 / ENSRNOG00000018975
ATIC / P31939 / ENSMUSG00000026192 / ENSG00000138363 / ENSRNOG00000015511
ATL1 / Q8WXF7 / ENSMUSG00000021066 / ENSG00000198513 / ENSRNOG00000005063
ATL2 / Q8NHH9 / ENSMUSG00000059811 / ENSG00000119787 / ENSRNOG00000006523
ATL3 / Q6DD88 / ENSMUSG00000024759 / ENSG00000184743 / ENSRNOG00000021203
ATM / Q13315 / ENSMUSG00000034218 / ENSG00000149311 / ENSRNOG00000029773
ATOX1 / O00244 / ENSMUSG00000018585 / ENSG00000177556 / ENSRNOG00000013118
ATP11A / P98196 / ENSMUSG00000031441 / ENSG00000068650 / ENSRNOG00000017154
ATP11C / Q8NB49 / ENSMUSG00000062949 / ENSG00000101974 / ENSRNOG00000003472
ATP12A / P54707 / ENSMUSG00000022229 / ENSG00000075673 / ENSRNOG00000020685
ATP13A1 / Q9HD20 / ENSMUSG00000031862 / ENSG00000105726 / ENSRNOG00000010776
ATP1A1 / P05023 / ENSMUSG00000033161 / ENSG00000163399 / ENSRNOG00000030019
ATP1A2 / P50993 / ENSMUSG00000007097 / ENSG00000018625 / ENSRNOG00000007290
ATP1A3 / P13637 / ENSMUSG00000040907 / ENSG00000105409 / ENSRNOG00000020263
ATP1A4 / Q13733 / ENSMUSG00000007107 / ENSG00000132681 / ENSRNOG00000032378
ATP1B1 / P05026 / ENSMUSG00000026576 / ENSG00000143153 / ENSRNOG00000002934
ATP1B2 / P14415 / ENSMUSG00000041329 / ENSG00000129244 / ENSRNOG00000011227
ATP1B3 / P54709 / ENSMUSG00000032412 / ENSG00000069849 / ENSRNOG00000011501
ATP2A2 / P16615 / ENSMUSG00000029467 / ENSG00000174437 / ENSRNOG00000001285
ATP2A3 / Q93084 / ENSMUSG00000020788 / ENSG00000074370 / ENSRNOG00000017912
ATP2B1 / P20020 / ENSMUSG00000019943 / ENSG00000070961 / ENSRNOG00000004026
ATP2B2 / Q01814 / ENSMUSG00000030302 / ENSG00000157087 / ENSRNOG00000030269
ATP2B3 / Q16720 / ENSMUSG00000031376 / ENSG00000067842 / ENSRNOG00000061304
ATP2B4 / P23634 / ENSMUSG00000026463 / ENSG00000058668 / ENSRNOG00000003031
Homologs
Brain
Heart
Kidney
Liver
Lung
Testis
Organs
Bins
1
2
3
4
5
Species
Human
Mouse
Rat

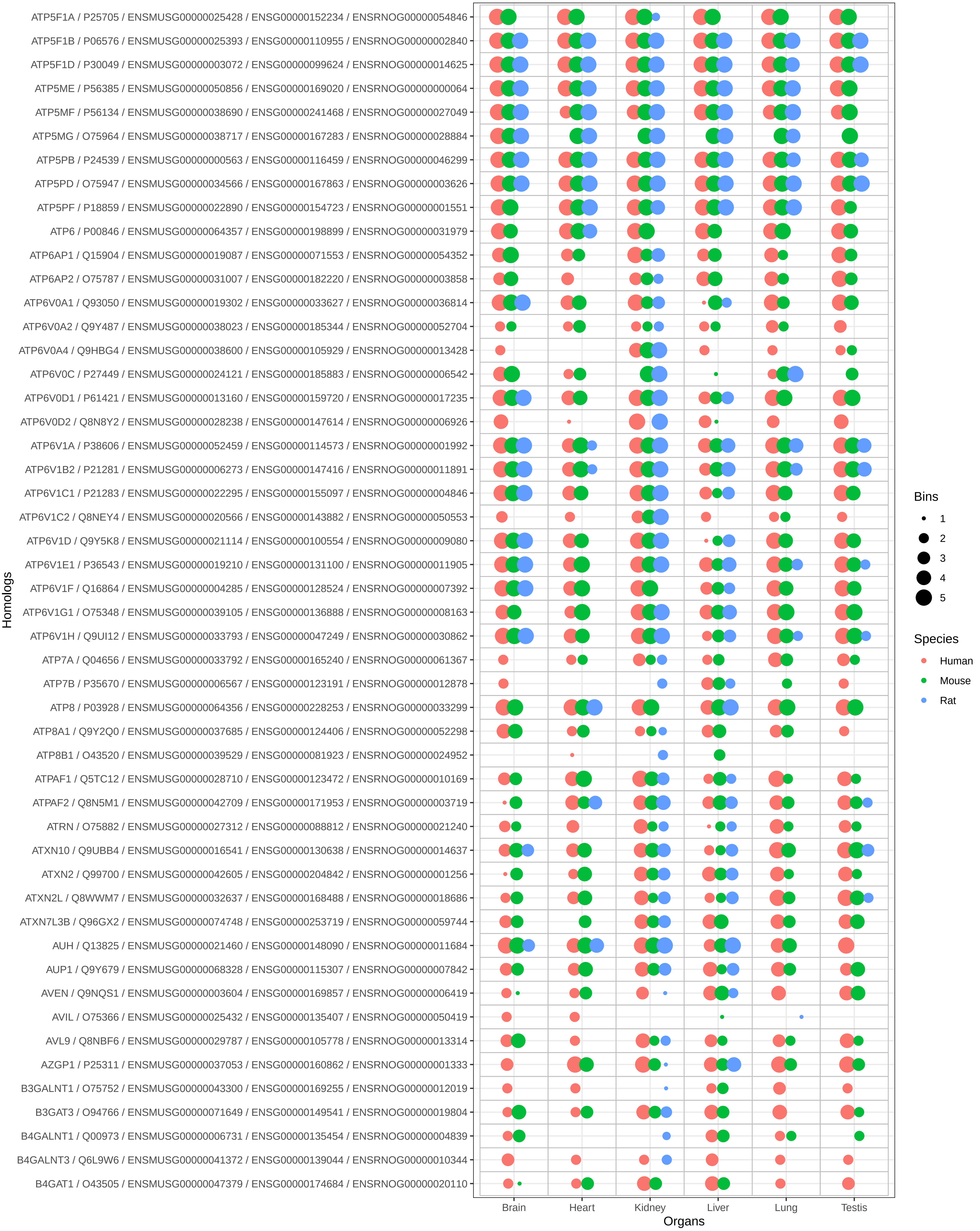
ATP5F1A / P25705 / ENSMUSG00000025428 / ENSG00000152234 / ENSRNOG00000054846
ATP5F1B / P06576 / ENSMUSG00000025393 / ENSG00000110955 / ENSRNOG00000002840
ATP5F1D / P30049 / ENSMUSG00000003072 / ENSG00000099624 / ENSRNOG00000014625
ATP5ME / P56385 / ENSMUSG00000050856 / ENSG00000169020 / ENSRNOG00000000064
ATP5MF / P56134 / ENSMUSG00000038690 / ENSG00000241468 / ENSRNOG00000027049
ATP5MG / O75964 / ENSMUSG00000038717 / ENSG00000167283 / ENSRNOG00000028884
ATP5PB / P24539 / ENSMUSG00000000563 / ENSG00000116459 / ENSRNOG00000046299
ATP5PD / O75947 / ENSMUSG00000034566 / ENSG00000167863 / ENSRNOG00000003626
ATP5PF / P18859 / ENSMUSG00000022890 / ENSG00000154723 / ENSRNOG00000001551
ATP6 / P00846 / ENSMUSG00000064357 / ENSG00000198899 / ENSRNOG00000031979
ATP6AP1 / Q15904 / ENSMUSG00000019087 / ENSG00000071553 / ENSRNOG00000054352
ATP6AP2 / O75787 / ENSMUSG00000031007 / ENSG00000182220 / ENSRNOG00000003858
ATP6V0A1 / Q93050 / ENSMUSG00000019302 / ENSG00000033627 / ENSRNOG00000036814
ATP6V0A2 / Q9Y487 / ENSMUSG00000038023 / ENSG00000185344 / ENSRNOG00000052704
ATP6V0A4 / Q9HBG4 / ENSMUSG00000038600 / ENSG00000105929 / ENSRNOG00000013428
ATP6V0C / P27449 / ENSMUSG00000024121 / ENSG00000185883 / ENSRNOG00000006542
ATP6V0D1 / P61421 / ENSMUSG00000013160 / ENSG00000159720 / ENSRNOG00000017235
ATP6V0D2 / Q8N8Y2 / ENSMUSG00000028238 / ENSG00000147614 / ENSRNOG00000006926
ATP6V1A / P38606 / ENSMUSG00000052459 / ENSG00000114573 / ENSRNOG00000001992
ATP6V1B2 / P21281 / ENSMUSG00000006273 / ENSG00000147416 / ENSRNOG00000011891
ATP6V1C1 / P21283 / ENSMUSG00000022295 / ENSG00000155097 / ENSRNOG00000004846
ATP6V1C2 / Q8NEY4 / ENSMUSG00000020566 / ENSG00000143882 / ENSRNOG00000050553
ATP6V1D / Q9Y5K8 / ENSMUSG00000021114 / ENSG00000100554 / ENSRNOG00000009080
ATP6V1E1 / P36543 / ENSMUSG00000019210 / ENSG00000131100 / ENSRNOG00000011905
ATP6V1F / Q16864 / ENSMUSG00000004285 / ENSG00000128524 / ENSRNOG00000007392
ATP6V1G1 / O75348 / ENSMUSG00000039105 / ENSG00000136888 / ENSRNOG00000008163
ATP6V1H / Q9UI12 / ENSMUSG00000033793 / ENSG00000047249 / ENSRNOG00000030862
ATP7A / Q04656 / ENSMUSG00000033792 / ENSG00000165240 / ENSRNOG00000061367
ATP7B / P35670 / ENSMUSG00000006567 / ENSG00000123191 / ENSRNOG00000012878
ATP8 / P03928 / ENSMUSG00000064356 / ENSG00000228253 / ENSRNOG00000033299
ATP8A1 / Q9Y2Q0 / ENSMUSG00000037685 / ENSG00000124406 / ENSRNOG00000052298
ATP8B1 / O43520 / ENSMUSG00000039529 / ENSG00000081923 / ENSRNOG00000024952
ATPAF1 / Q5TC12 / ENSMUSG00000028710 / ENSG00000123472 / ENSRNOG00000010169
ATPAF2 / Q8N5M1 / ENSMUSG00000042709 / ENSG00000171953 / ENSRNOG00000003719
ATRN / O75882 / ENSMUSG00000027312 / ENSG00000088812 / ENSRNOG00000021240
ATXN10 / Q9UBB4 / ENSMUSG00000016541 / ENSG00000130638 / ENSRNOG00000014637
ATXN2 / Q99700 / ENSMUSG00000042605 / ENSG00000204842 / ENSRNOG00000001256
ATXN2L / Q8WWM7 / ENSMUSG00000032637 / ENSG00000168488 / ENSRNOG00000018686
ATXN7L3B / Q96GX2 / ENSMUSG00000074748 / ENSG00000253719 / ENSRNOG00000059744
AUH / Q13825 / ENSMUSG00000021460 / ENSG00000148090 / ENSRNOG00000011684
AUP1 / Q9Y679 / ENSMUSG00000068328 / ENSG00000115307 / ENSRNOG00000007842
AVEN / Q9NQS1 / ENSMUSG00000003604 / ENSG00000169857 / ENSRNOG00000006419
AVIL / O75366 / ENSMUSG00000025432 / ENSG00000135407 / ENSRNOG00000050419
AVL9 / Q8NBF6 / ENSMUSG00000029787 / ENSG00000105778 / ENSRNOG00000013314
AZGP1 / P25311 / ENSMUSG00000037053 / ENSG00000160862 / ENSRNOG00000001333
B3GALNT1 / O75752 / ENSMUSG00000043300 / ENSG00000169255 / ENSRNOG00000012019
B3GAT3 / O94766 / ENSMUSG00000071649 / ENSG00000149541 / ENSRNOG00000019804
B4GALNT1 / Q00973 / ENSMUSG00000006731 / ENSG00000135454 / ENSRNOG00000004839
B4GALNT3 / Q6L9W6 / ENSMUSG00000041372 / ENSG00000139044 / ENSRNOG00000010344
B4GAT1 / O43505 / ENSMUSG00000047379 / ENSG00000174684 / ENSRNOG00000020110
Homologs
Brain
Heart
Kidney
Liver
Lung
Testis
Organs
Bins
1
2
3
4
5
Species
Human
Mouse
Rat

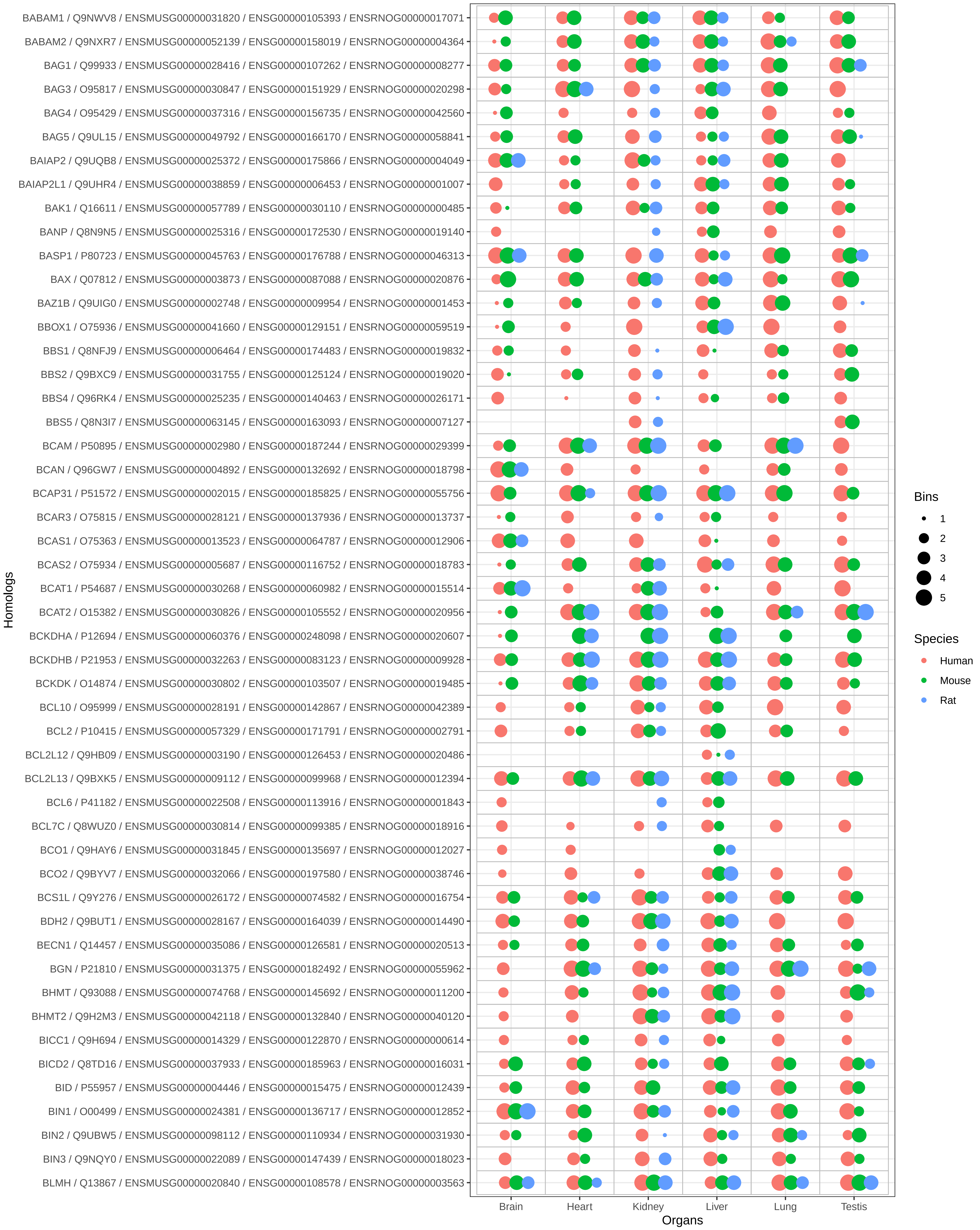
BABAM1 / Q9NWV8 / ENSMUSG00000031820 / ENSG00000105393 / ENSRNOG00000017071
BABAM2 / Q9NXR7 / ENSMUSG00000052139 / ENSG00000158019 / ENSRNOG00000004364
BAG1 / Q99933 / ENSMUSG00000028416 / ENSG00000107262 / ENSRNOG00000008277
BAG3 / O95817 / ENSMUSG00000030847 / ENSG00000151929 / ENSRNOG00000020298
BAG4 / O95429 / ENSMUSG00000037316 / ENSG00000156735 / ENSRNOG00000042560
BAG5 / Q9UL15 / ENSMUSG00000049792 / ENSG00000166170 / ENSRNOG00000058841
BAIAP2 / Q9UQB8 / ENSMUSG00000025372 / ENSG00000175866 / ENSRNOG00000004049
BAIAP2L1 / Q9UHR4 / ENSMUSG00000038859 / ENSG00000006453 / ENSRNOG00000001007
BAK1 / Q16611 / ENSMUSG00000057789 / ENSG00000030110 / ENSRNOG00000000485
BANP / Q8N9N5 / ENSMUSG00000025316 / ENSG00000172530 / ENSRNOG00000019140
BASP1 / P80723 / ENSMUSG00000045763 / ENSG00000176788 / ENSRNOG00000046313
BAX / Q07812 / ENSMUSG00000003873 / ENSG00000087088 / ENSRNOG00000020876
BAZ1B / Q9UIG0 / ENSMUSG00000002748 / ENSG00000009954 / ENSRNOG00000001453
BBOX1 / O75936 / ENSMUSG00000041660 / ENSG00000129151 / ENSRNOG00000059519
BBS1 / Q8NFJ9 / ENSMUSG00000006464 / ENSG00000174483 / ENSRNOG00000019832
BBS2 / Q9BXC9 / ENSMUSG00000031755 / ENSG00000125124 / ENSRNOG00000019020
BBS4 / Q96RK4 / ENSMUSG00000025235 / ENSG00000140463 / ENSRNOG00000026171
BBS5 / Q8N3I7 / ENSMUSG00000063145 / ENSG00000163093 / ENSRNOG00000007127
BCAM / P50895 / ENSMUSG00000002980 / ENSG00000187244 / ENSRNOG00000029399
BCAN / Q96GW7 / ENSMUSG00000004892 / ENSG00000132692 / ENSRNOG00000018798
BCAP31 / P51572 / ENSMUSG00000002015 / ENSG00000185825 / ENSRNOG00000055756
BCAR3 / O75815 / ENSMUSG00000028121 / ENSG00000137936 / ENSRNOG00000013737
BCAS1 / O75363 / ENSMUSG00000013523 / ENSG00000064787 / ENSRNOG00000012906
BCAS2 / O75934 / ENSMUSG00000005687 / ENSG00000116752 / ENSRNOG00000018783
BCAT1 / P54687 / ENSMUSG00000030268 / ENSG00000060982 / ENSRNOG00000015514
BCAT2 / O15382 / ENSMUSG00000030826 / ENSG00000105552 / ENSRNOG00000020956
BCKDHA / P12694 / ENSMUSG00000060376 / ENSG00000248098 / ENSRNOG00000020607
BCKDHB / P21953 / ENSMUSG00000032263 / ENSG00000083123 / ENSRNOG00000009928
BCKDK / O14874 / ENSMUSG00000030802 / ENSG00000103507 / ENSRNOG00000019485
BCL10 / O95999 / ENSMUSG00000028191 / ENSG00000142867 / ENSRNOG00000042389
BCL2 / P10415 / ENSMUSG00000057329 / ENSG00000171791 / ENSRNOG00000002791
BCL2L12 / Q9HB09 / ENSMUSG00000003190 / ENSG00000126453 / ENSRNOG00000020486
BCL2L13 / Q9BXK5 / ENSMUSG00000009112 / ENSG00000099968 / ENSRNOG00000012394
BCL6 / P41182 / ENSMUSG00000022508 / ENSG00000113916 / ENSRNOG00000001843
BCL7C / Q8WUZ0 / ENSMUSG00000030814 / ENSG00000099385 / ENSRNOG00000018916
BCO1 / Q9HAY6 / ENSMUSG00000031845 / ENSG00000135697 / ENSRNOG00000012027
BCO2 / Q9BYV7 / ENSMUSG00000032066 / ENSG00000197580 / ENSRNOG00000038746
BCS1L / Q9Y276 / ENSMUSG00000026172 / ENSG00000074582 / ENSRNOG00000016754
BDH2 / Q9BUT1 / ENSMUSG00000028167 / ENSG00000164039 / ENSRNOG00000014490
BECN1 / Q14457 / ENSMUSG00000035086 / ENSG00000126581 / ENSRNOG00000020513
BGN / P21810 / ENSMUSG00000031375 / ENSG00000182492 / ENSRNOG00000055962
BHMT / Q93088 / ENSMUSG00000074768 / ENSG00000145692 / ENSRNOG00000011200
BHMT2 / Q9H2M3 / ENSMUSG00000042118 / ENSG00000132840 / ENSRNOG00000040120
BICC1 / Q9H694 / ENSMUSG00000014329 / ENSG00000122870 / ENSRNOG00000000614
BICD2 / Q8TD16 / ENSMUSG00000037933 / ENSG00000185963 / ENSRNOG00000016031
BID / P55957 / ENSMUSG00000004446 / ENSG00000015475 / ENSRNOG00000012439
BIN1 / O00499 / ENSMUSG00000024381 / ENSG00000136717 / ENSRNOG00000012852
BIN2 / Q9UBW5 / ENSMUSG00000098112 / ENSG00000110934 / ENSRNOG00000031930
BIN3 / Q9NQY0 / ENSMUSG00000022089 / ENSG00000147439 / ENSRNOG00000018023
BLMH / Q13867 / ENSMUSG00000020840 / ENSG00000108578 / ENSRNOG00000003563
Homologs
Brain
Heart
Kidney
Liver
Lung
Testis
Organs
Bins
1
2
3
4
5
Species
Human
Mouse
Rat

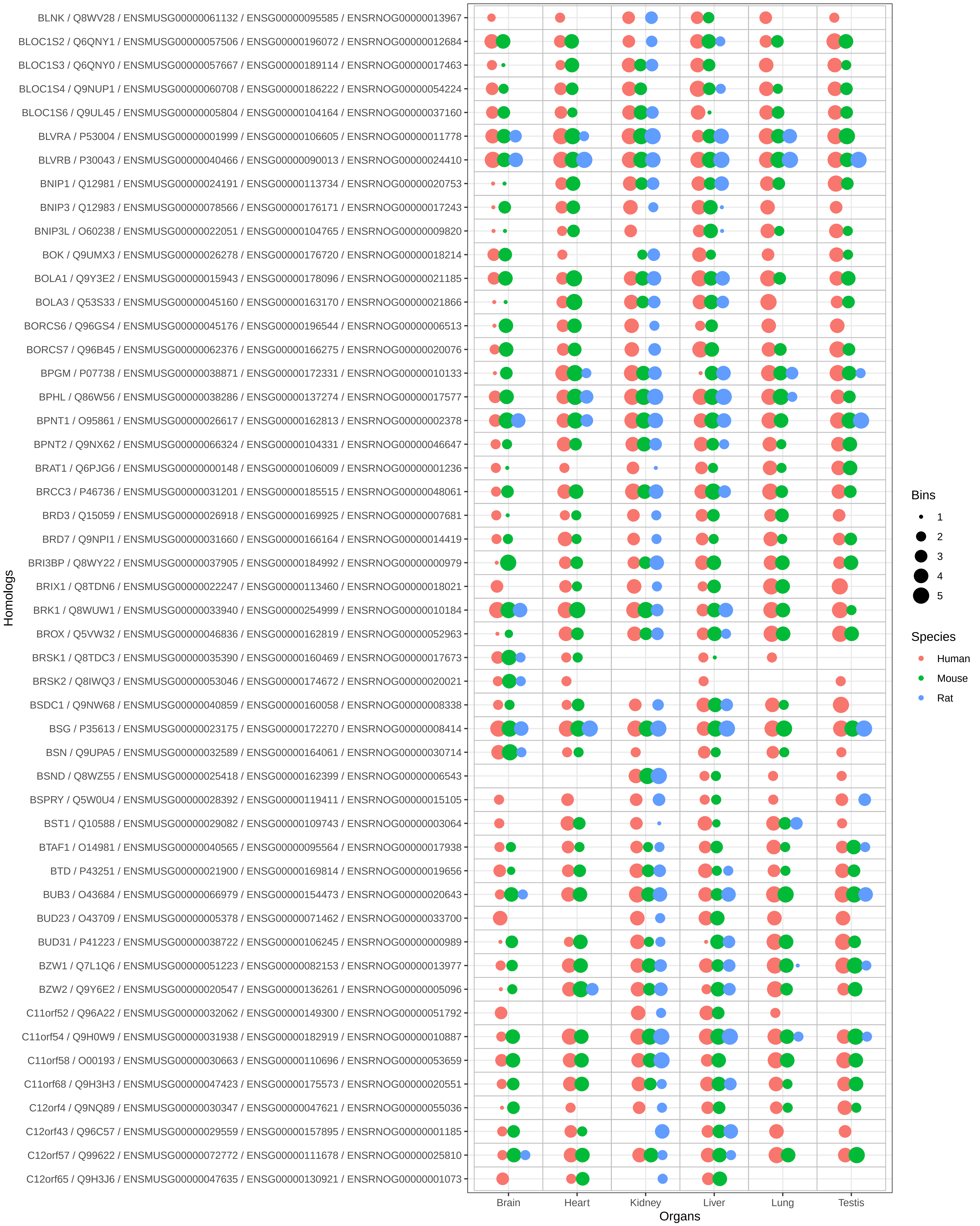
BLNK / Q8WV28 / ENSMUSG00000061132 / ENSG00000095585 / ENSRNOG00000013967
BLOC1S2 / Q6QNY1 / ENSMUSG00000057506 / ENSG00000196072 / ENSRNOG00000012684
BLOC1S3 / Q6QNY0 / ENSMUSG00000057667 / ENSG00000189114 / ENSRNOG00000017463
BLOC1S4 / Q9NUP1 / ENSMUSG00000060708 / ENSG00000186222 / ENSRNOG00000054224
BLOC1S6 / Q9UL45 / ENSMUSG00000005804 / ENSG00000104164 / ENSRNOG00000037160
BLVRA / P53004 / ENSMUSG00000001999 / ENSG00000106605 / ENSRNOG00000011778
BLVRB / P30043 / ENSMUSG00000040466 / ENSG00000090013 / ENSRNOG00000024410
BNIP1 / Q12981 / ENSMUSG00000024191 / ENSG00000113734 / ENSRNOG00000020753
BNIP3 / Q12983 / ENSMUSG00000078566 / ENSG00000176171 / ENSRNOG00000017243
BNIP3L / O60238 / ENSMUSG00000022051 / ENSG00000104765 / ENSRNOG00000009820
BOK / Q9UMX3 / ENSMUSG00000026278 / ENSG00000176720 / ENSRNOG00000018214
BOLA1 / Q9Y3E2 / ENSMUSG00000015943 / ENSG00000178096 / ENSRNOG00000021185
BOLA3 / Q53S33 / ENSMUSG00000045160 / ENSG00000163170 / ENSRNOG00000021866
BORCS6 / Q96GS4 / ENSMUSG00000045176 / ENSG00000196544 / ENSRNOG00000006513
BORCS7 / Q96B45 / ENSMUSG00000062376 / ENSG00000166275 / ENSRNOG00000020076
BPGM / P07738 / ENSMUSG00000038871 / ENSG00000172331 / ENSRNOG00000010133
BPHL / Q86W56 / ENSMUSG00000038286 / ENSG00000137274 / ENSRNOG00000017577
BPNT1 / O95861 / ENSMUSG00000026617 / ENSG00000162813 / ENSRNOG00000002378
BPNT2 / Q9NX62 / ENSMUSG00000066324 / ENSG00000104331 / ENSRNOG00000046647
BRAT1 / Q6PJG6 / ENSMUSG00000000148 / ENSG00000106009 / ENSRNOG00000001236
BRCC3 / P46736 / ENSMUSG00000031201 / ENSG00000185515 / ENSRNOG00000048061
BRD3 / Q15059 / ENSMUSG00000026918 / ENSG00000169925 / ENSRNOG00000007681
BRD7 / Q9NPI1 / ENSMUSG00000031660 / ENSG00000166164 / ENSRNOG00000014419
BRI3BP / Q8WY22 / ENSMUSG00000037905 / ENSG00000184992 / ENSRNOG00000000979
BRIX1 / Q8TDN6 / ENSMUSG00000022247 / ENSG00000113460 / ENSRNOG00000018021
BRK1 / Q8WUW1 / ENSMUSG00000033940 / ENSG00000254999 / ENSRNOG00000010184
BROX / Q5VW32 / ENSMUSG00000046836 / ENSG00000162819 / ENSRNOG00000052963
BRSK1 / Q8TDC3 / ENSMUSG00000035390 / ENSG00000160469 / ENSRNOG00000017673
BRSK2 / Q8IWQ3 / ENSMUSG00000053046 / ENSG00000174672 / ENSRNOG00000020021
BSDC1 / Q9NW68 / ENSMUSG00000040859 / ENSG00000160058 / ENSRNOG00000008338
BSG / P35613 / ENSMUSG00000023175 / ENSG00000172270 / ENSRNOG00000008414
BSN / Q9UPA5 / ENSMUSG00000032589 / ENSG00000164061 / ENSRNOG00000030714
BSND / Q8WZ55 / ENSMUSG00000025418 / ENSG00000162399 / ENSRNOG00000006543
BSPRY / Q5W0U4 / ENSMUSG00000028392 / ENSG00000119411 / ENSRNOG00000015105
BST1 / Q10588 / ENSMUSG00000029082 / ENSG00000109743 / ENSRNOG00000003064
BTAF1 / O14981 / ENSMUSG00000040565 / ENSG00000095564 / ENSRNOG00000017938
BTD / P43251 / ENSMUSG00000021900 / ENSG00000169814 / ENSRNOG00000019656
BUB3 / O43684 / ENSMUSG00000066979 / ENSG00000154473 / ENSRNOG00000020643
BUD23 / O43709 / ENSMUSG00000005378 / ENSG00000071462 / ENSRNOG00000033700
BUD31 / P41223 / ENSMUSG00000038722 / ENSG00000106245 / ENSRNOG00000000989
BZW1 / Q7L1Q6 / ENSMUSG00000051223 / ENSG00000082153 / ENSRNOG00000013977
BZW2 / Q9Y6E2 / ENSMUSG00000020547 / ENSG00000136261 / ENSRNOG00000005096
C11orf52 / Q96A22 / ENSMUSG00000032062 / ENSG00000149300 / ENSRNOG00000051792
C11orf54 / Q9H0W9 / ENSMUSG00000031938 / ENSG00000182919 / ENSRNOG00000010887
C11orf58 / O00193 / ENSMUSG00000030663 / ENSG00000110696 / ENSRNOG00000053659
C11orf68 / Q9H3H3 / ENSMUSG00000047423 / ENSG00000175573 / ENSRNOG00000020551
C12orf4 / Q9NQ89 / ENSMUSG00000030347 / ENSG00000047621 / ENSRNOG00000055036
C12orf43 / Q96C57 / ENSMUSG00000029559 / ENSG00000157895 / ENSRNOG00000001185
C12orf57 / Q99622 / ENSMUSG00000072772 / ENSG00000111678 / ENSRNOG00000025810
C12orf65 / Q9H3J6 / ENSMUSG00000047635 / ENSG00000130921 / ENSRNOG00000001073
Homologs
Brain
Heart
Kidney
Liver
Lung
Testis
Organs
Bins
1
2
3
4
5
Species
Human
Mouse
Rat

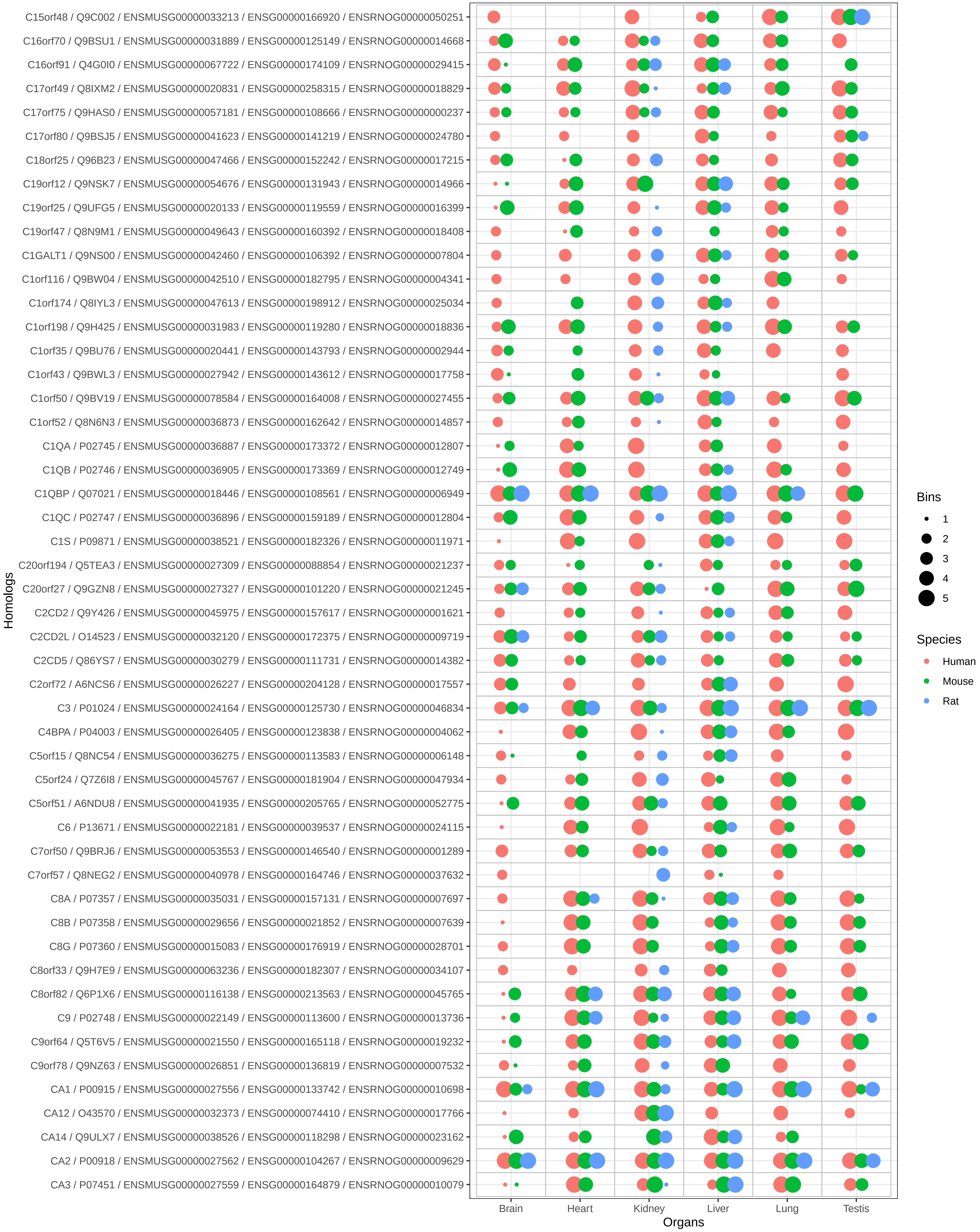
C15orf48 / Q9C002 / ENSMUSG00000033213 / ENSG00000166920 / ENSRNOG00000050251
C16orf70 / Q9BSU1 / ENSMUSG00000031889 / ENSG00000125149 / ENSRNOG00000014668
C16orf91 / Q4G0I0 / ENSMUSG00000067722 / ENSG00000174109 / ENSRNOG00000029415
C17orf49 / Q8IXM2 / ENSMUSG00000020831 / ENSG00000258315 / ENSRNOG00000018829
C17orf75 / Q9HAS0 / ENSMUSG00000057181 / ENSG00000108666 / ENSRNOG00000000237
C17orf80 / Q9BSJ5 / ENSMUSG00000041623 / ENSG00000141219 / ENSRNOG00000024780
C18orf25 / Q96B23 / ENSMUSG00000047466 / ENSG00000152242 / ENSRNOG00000017215
C19orf12 / Q9NSK7 / ENSMUSG00000054676 / ENSG00000131943 / ENSRNOG00000014966
C19orf25 / Q9UFG5 / ENSMUSG00000020133 / ENSG00000119559 / ENSRNOG00000016399
C19orf47 / Q8N9M1 / ENSMUSG00000049643 / ENSG00000160392 / ENSRNOG00000018408
C1GALT1 / Q9NS00 / ENSMUSG00000042460 / ENSG00000106392 / ENSRNOG00000007804
C1orf116 / Q9BW04 / ENSMUSG00000042510 / ENSG00000182795 / ENSRNOG00000004341
C1orf174 / Q8IYL3 / ENSMUSG00000047613 / ENSG00000198912 / ENSRNOG00000025034
C1orf198 / Q9H425 / ENSMUSG00000031983 / ENSG00000119280 / ENSRNOG00000018836
C1orf35 / Q9BU76 / ENSMUSG00000020441 / ENSG00000143793 / ENSRNOG00000002944
C1orf43 / Q9BWL3 / ENSMUSG00000027942 / ENSG00000143612 / ENSRNOG00000017758
C1orf50 / Q9BV19 / ENSMUSG00000078584 / ENSG00000164008 / ENSRNOG00000027455
C1orf52 / Q8N6N3 / ENSMUSG00000036873 / ENSG00000162642 / ENSRNOG00000014857
C1QA / P02745 / ENSMUSG00000036887 / ENSG00000173372 / ENSRNOG00000012807
C1QB / P02746 / ENSMUSG00000036905 / ENSG00000173369 / ENSRNOG00000012749
C1QBP / Q07021 / ENSMUSG00000018446 / ENSG00000108561 / ENSRNOG00000006949
C1QC / P02747 / ENSMUSG00000036896 / ENSG00000159189 / ENSRNOG00000012804
C1S / P09871 / ENSMUSG00000038521 / ENSG00000182326 / ENSRNOG00000011971
C20orf194 / Q5TEA3 / ENSMUSG00000027309 / ENSG00000088854 / ENSRNOG00000021237
C20orf27 / Q9GZN8 / ENSMUSG00000027327 / ENSG00000101220 / ENSRNOG00000021245
C2CD2 / Q9Y426 / ENSMUSG00000045975 / ENSG00000157617 / ENSRNOG00000001621
C2CD2L / O14523 / ENSMUSG00000032120 / ENSG00000172375 / ENSRNOG00000009719
C2CD5 / Q86YS7 / ENSMUSG00000030279 / ENSG00000111731 / ENSRNOG00000014382
C2orf72 / A6NCS6 / ENSMUSG00000026227 / ENSG00000204128 / ENSRNOG00000017557
C3 / P01024 / ENSMUSG00000024164 / ENSG00000125730 / ENSRNOG00000046834
C4BPA / P04003 / ENSMUSG00000026405 / ENSG00000123838 / ENSRNOG00000004062
C5orf15 / Q8NC54 / ENSMUSG00000036275 / ENSG00000113583 / ENSRNOG00000006148
C5orf24 / Q7Z6I8 / ENSMUSG00000045767 / ENSG00000181904 / ENSRNOG00000047934
C5orf51 / A6NDU8 / ENSMUSG00000041935 / ENSG00000205765 / ENSRNOG00000052775
C6 / P13671 / ENSMUSG00000022181 / ENSG00000039537 / ENSRNOG00000024115
C7orf50 / Q9BRJ6 / ENSMUSG00000053553 / ENSG00000146540 / ENSRNOG00000001289
C7orf57 / Q8NEG2 / ENSMUSG00000040978 / ENSG00000164746 / ENSRNOG00000037632
C8A / P07357 / ENSMUSG00000035031 / ENSG00000157131 / ENSRNOG00000007697
C8B / P07358 / ENSMUSG00000029656 / ENSG00000021852 / ENSRNOG00000007639
C8G / P07360 / ENSMUSG00000015083 / ENSG00000176919 / ENSRNOG00000028701
C8orf33 / Q9H7E9 / ENSMUSG00000063236 / ENSG00000182307 / ENSRNOG00000034107
C8orf82 / Q6P1X6 / ENSMUSG00000116138 / ENSG00000213563 / ENSRNOG00000045765
C9 / P02748 / ENSMUSG00000022149 / ENSG00000113600 / ENSRNOG00000013736
C9orf64 / Q5T6V5 / ENSMUSG00000021550 / ENSG00000165118 / ENSRNOG00000019232
C9orf78 / Q9NZ63 / ENSMUSG00000026851 / ENSG00000136819 / ENSRNOG00000007532
CA1 / P00915 / ENSMUSG00000027556 / ENSG00000133742 / ENSRNOG00000010698
CA12 / O43570 / ENSMUSG00000032373 / ENSG00000074410 / ENSRNOG00000017766
CA14 / Q9ULX7 / ENSMUSG00000038526 / ENSG00000118298 / ENSRNOG00000023162
CA2 / P00918 / ENSMUSG00000027562 / ENSG00000104267 / ENSRNOG00000009629
CA3 / P07451 / ENSMUSG00000027559 / ENSG00000164879 / ENSRNOG00000010079
Homologs
Brain
Heart
Kidney
Liver
Lung
Testis
Organs
Bins
1
2
3
4
5
Species
Human
Mouse
Rat

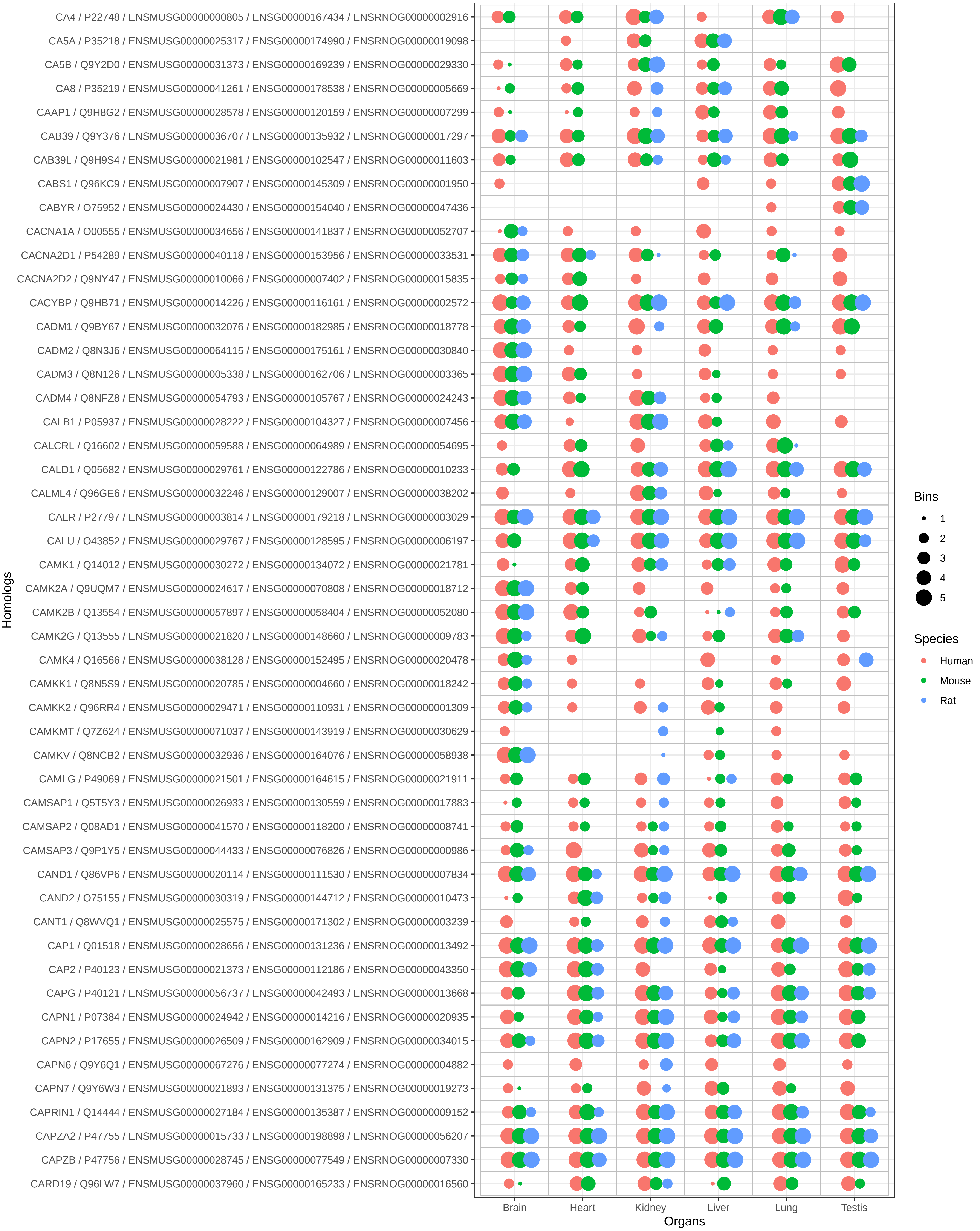
CA4 / P22748 / ENSMUSG00000000805 / ENSG00000167434 / ENSRNOG00000002916
CA5A / P35218 / ENSMUSG00000025317 / ENSG00000174990 / ENSRNOG00000019098
CA5B / Q9Y2D0 / ENSMUSG00000031373 / ENSG00000169239 / ENSRNOG00000029330
CA8 / P35219 / ENSMUSG00000041261 / ENSG00000178538 / ENSRNOG00000005669
CAAP1 / Q9H8G2 / ENSMUSG00000028578 / ENSG00000120159 / ENSRNOG00000007299
CAB39 / Q9Y376 / ENSMUSG00000036707 / ENSG00000135932 / ENSRNOG00000017297
CAB39L / Q9H9S4 / ENSMUSG00000021981 / ENSG00000102547 / ENSRNOG00000011603
CABS1 / Q96KC9 / ENSMUSG00000007907 / ENSG00000145309 / ENSRNOG00000001950
CABYR / O75952 / ENSMUSG00000024430 / ENSG00000154040 / ENSRNOG00000047436
CACNA1A / O00555 / ENSMUSG00000034656 / ENSG00000141837 / ENSRNOG00000052707
CACNA2D1 / P54289 / ENSMUSG00000040118 / ENSG00000153956 / ENSRNOG00000033531
CACNA2D2 / Q9NY47 / ENSMUSG00000010066 / ENSG00000007402 / ENSRNOG00000015835
CACYBP / Q9HB71 / ENSMUSG00000014226 / ENSG00000116161 / ENSRNOG00000002572
CADM1 / Q9BY67 / ENSMUSG00000032076 / ENSG00000182985 / ENSRNOG00000018778
CADM2 / Q8N3J6 / ENSMUSG00000064115 / ENSG00000175161 / ENSRNOG00000030840
CADM3 / Q8N126 / ENSMUSG00000005338 / ENSG00000162706 / ENSRNOG00000003365
CADM4 / Q8NFZ8 / ENSMUSG00000054793 / ENSG00000105767 / ENSRNOG00000024243
CALB1 / P05937 / ENSMUSG00000028222 / ENSG00000104327 / ENSRNOG00000007456
CALCRL / Q16602 / ENSMUSG00000059588 / ENSG00000064989 / ENSRNOG00000054695
CALD1 / Q05682 / ENSMUSG00000029761 / ENSG00000122786 / ENSRNOG00000010233
CALML4 / Q96GE6 / ENSMUSG00000032246 / ENSG00000129007 / ENSRNOG00000038202
CALR / P27797 / ENSMUSG00000003814 / ENSG00000179218 / ENSRNOG00000003029
CALU / O43852 / ENSMUSG00000029767 / ENSG00000128595 / ENSRNOG00000006197
CAMK1 / Q14012 / ENSMUSG00000030272 / ENSG00000134072 / ENSRNOG00000021781
CAMK2A / Q9UQM7 / ENSMUSG00000024617 / ENSG00000070808 / ENSRNOG00000018712
CAMK2B / Q13554 / ENSMUSG00000057897 / ENSG00000058404 / ENSRNOG00000052080
CAMK2G / Q13555 / ENSMUSG00000021820 / ENSG00000148660 / ENSRNOG00000009783
CAMK4 / Q16566 / ENSMUSG00000038128 / ENSG00000152495 / ENSRNOG00000020478
CAMKK1 / Q8N5S9 / ENSMUSG00000020785 / ENSG00000004660 / ENSRNOG00000018242
CAMKK2 / Q96RR4 / ENSMUSG00000029471 / ENSG00000110931 / ENSRNOG00000001309
CAMKMT / Q7Z624 / ENSMUSG00000071037 / ENSG00000143919 / ENSRNOG00000030629
CAMKV / Q8NCB2 / ENSMUSG00000032936 / ENSG00000164076 / ENSRNOG00000058938
CAMLG / P49069 / ENSMUSG00000021501 / ENSG00000164615 / ENSRNOG00000021911
CAMSAP1 / Q5T5Y3 / ENSMUSG00000026933 / ENSG00000130559 / ENSRNOG00000017883
CAMSAP2 / Q08AD1 / ENSMUSG00000041570 / ENSG00000118200 / ENSRNOG00000008741
CAMSAP3 / Q9P1Y5 / ENSMUSG00000044433 / ENSG00000076826 / ENSRNOG00000000986
CAND1 / Q86VP6 / ENSMUSG00000020114 / ENSG00000111530 / ENSRNOG00000007834
CAND2 / O75155 / ENSMUSG00000030319 / ENSG00000144712 / ENSRNOG00000010473
CANT1 / Q8WVQ1 / ENSMUSG00000025575 / ENSG00000171302 / ENSRNOG00000003239
CAP1 / Q01518 / ENSMUSG00000028656 / ENSG00000131236 / ENSRNOG00000013492
CAP2 / P40123 / ENSMUSG00000021373 / ENSG00000112186 / ENSRNOG00000043350
CAPG / P40121 / ENSMUSG00000056737 / ENSG00000042493 / ENSRNOG00000013668
CAPN1 / P07384 / ENSMUSG00000024942 / ENSG00000014216 / ENSRNOG00000020935
CAPN2 / P17655 / ENSMUSG00000026509 / ENSG00000162909 / ENSRNOG00000034015
CAPN6 / Q9Y6Q1 / ENSMUSG00000067276 / ENSG00000077274 / ENSRNOG00000004882
CAPN7 / Q9Y6W3 / ENSMUSG00000021893 / ENSG00000131375 / ENSRNOG00000019273
CAPRIN1 / Q14444 / ENSMUSG00000027184 / ENSG00000135387 / ENSRNOG00000009152
CAPZA2 / P47755 / ENSMUSG00000015733 / ENSG00000198898 / ENSRNOG00000056207
CAPZB / P47756 / ENSMUSG00000028745 / ENSG00000077549 / ENSRNOG00000007330
CARD19 / Q96LW7 / ENSMUSG00000037960 / ENSG00000165233 / ENSRNOG00000016560
Homologs
Brain
Heart
Kidney
Liver
Lung
Testis
Organs
Bins
1
2
3
4
5
Species
Human
Mouse
Rat

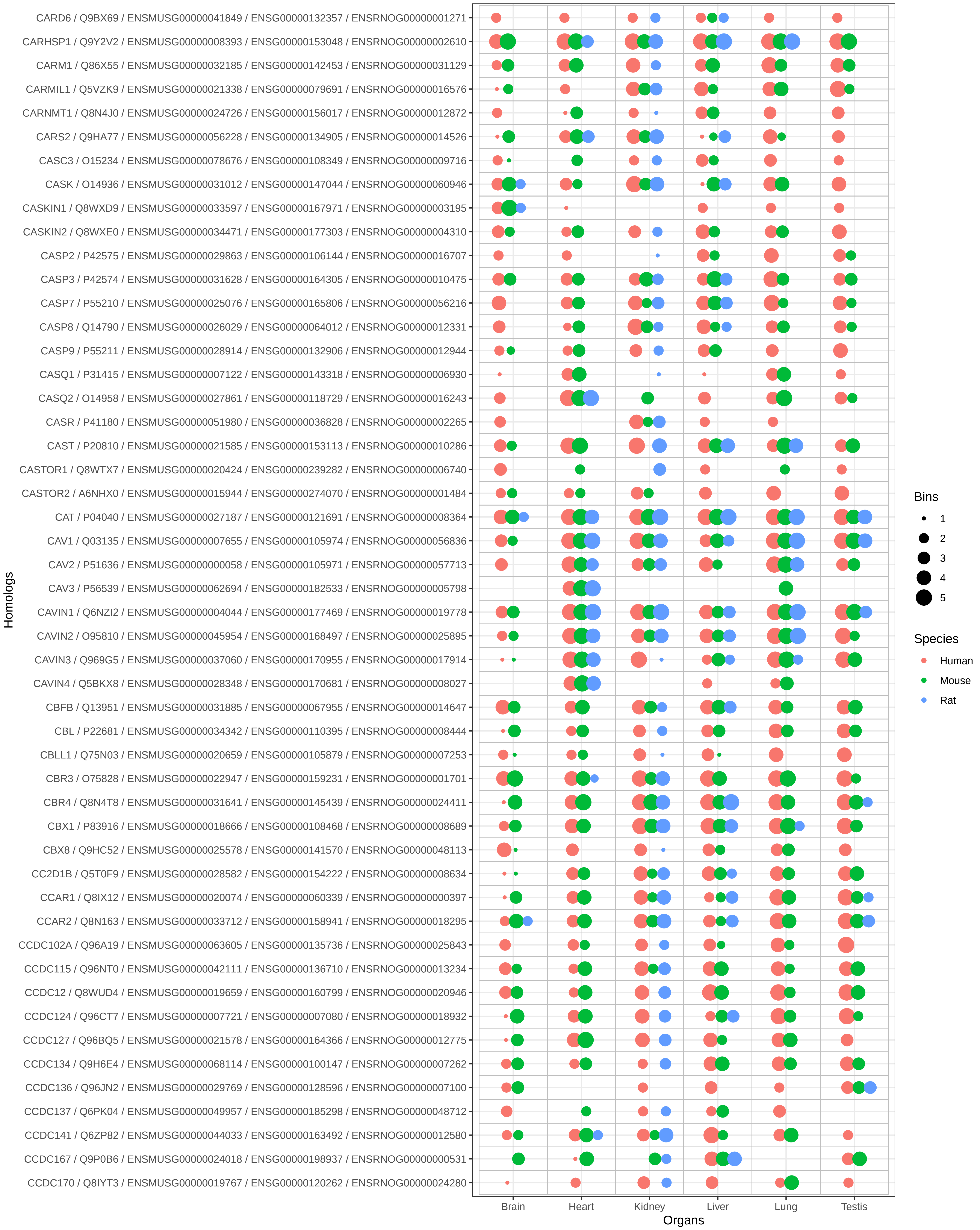
CARD6 / Q9BX69 / ENSMUSG00000041849 / ENSG00000132357 / ENSRNOG00000001271
CARHSP1 / Q9Y2V2 / ENSMUSG00000008393 / ENSG00000153048 / ENSRNOG00000002610
CARM1 / Q86X55 / ENSMUSG00000032185 / ENSG00000142453 / ENSRNOG00000031129
CARMIL1 / Q5VZK9 / ENSMUSG00000021338 / ENSG00000079691 / ENSRNOG00000016576
CARNMT1 / Q8N4J0 / ENSMUSG00000024726 / ENSG00000156017 / ENSRNOG00000012872
CARS2 / Q9HA77 / ENSMUSG00000056228 / ENSG00000134905 / ENSRNOG00000014526
CASC3 / O15234 / ENSMUSG00000078676 / ENSG00000108349 / ENSRNOG00000009716
CASK / O14936 / ENSMUSG00000031012 / ENSG00000147044 / ENSRNOG00000060946
CASKIN1 / Q8WXD9 / ENSMUSG00000033597 / ENSG00000167971 / ENSRNOG00000003195
CASKIN2 / Q8WXE0 / ENSMUSG00000034471 / ENSG00000177303 / ENSRNOG00000004310
CASP2 / P42575 / ENSMUSG00000029863 / ENSG00000106144 / ENSRNOG00000016707
CASP3 / P42574 / ENSMUSG00000031628 / ENSG00000164305 / ENSRNOG00000010475
CASP7 / P55210 / ENSMUSG00000025076 / ENSG00000165806 / ENSRNOG00000056216
CASP8 / Q14790 / ENSMUSG00000026029 / ENSG00000064012 / ENSRNOG00000012331
CASP9 / P55211 / ENSMUSG00000028914 / ENSG00000132906 / ENSRNOG00000012944
CASQ1 / P31415 / ENSMUSG00000007122 / ENSG00000143318 / ENSRNOG00000006930
CASQ2 / O14958 / ENSMUSG00000027861 / ENSG00000118729 / ENSRNOG00000016243
CASR / P41180 / ENSMUSG00000051980 / ENSG00000036828 / ENSRNOG00000002265
CAST / P20810 / ENSMUSG00000021585 / ENSG00000153113 / ENSRNOG00000010286
CASTOR1 / Q8WTX7 / ENSMUSG00000020424 / ENSG00000239282 / ENSRNOG00000006740
CASTOR2 / A6NHX0 / ENSMUSG00000015944 / ENSG00000274070 / ENSRNOG00000001484
CAT / P04040 / ENSMUSG00000027187 / ENSG00000121691 / ENSRNOG00000008364
CAV1 / Q03135 / ENSMUSG00000007655 / ENSG00000105974 / ENSRNOG00000056836
CAV2 / P51636 / ENSMUSG00000000058 / ENSG00000105971 / ENSRNOG00000057713
CAV3 / P56539 / ENSMUSG00000062694 / ENSG00000182533 / ENSRNOG00000005798
CAVIN1 / Q6NZI2 / ENSMUSG00000004044 / ENSG00000177469 / ENSRNOG00000019778
CAVIN2 / O95810 / ENSMUSG00000045954 / ENSG00000168497 / ENSRNOG00000025895
CAVIN3 / Q969G5 / ENSMUSG00000037060 / ENSG00000170955 / ENSRNOG00000017914
CAVIN4 / Q5BKX8 / ENSMUSG00000028348 / ENSG00000170681 / ENSRNOG00000008027
CBFB / Q13951 / ENSMUSG00000031885 / ENSG00000067955 / ENSRNOG00000014647
CBL / P22681 / ENSMUSG00000034342 / ENSG00000110395 / ENSRNOG00000008444
CBLL1 / Q75N03 / ENSMUSG00000020659 / ENSG00000105879 / ENSRNOG00000007253
CBR3 / O75828 / ENSMUSG00000022947 / ENSG00000159231 / ENSRNOG00000001701
CBR4 / Q8N4T8 / ENSMUSG00000031641 / ENSG00000145439 / ENSRNOG00000024411
CBX1 / P83916 / ENSMUSG00000018666 / ENSG00000108468 / ENSRNOG00000008689
CBX8 / Q9HC52 / ENSMUSG00000025578 / ENSG00000141570 / ENSRNOG00000048113
CC2D1B / Q5T0F9 / ENSMUSG00000028582 / ENSG00000154222 / ENSRNOG00000008634
CCAR1 / Q8IX12 / ENSMUSG00000020074 / ENSG00000060339 / ENSRNOG00000000397
CCAR2 / Q8N163 / ENSMUSG00000033712 / ENSG00000158941 / ENSRNOG00000018295
CCDC102A / Q96A19 / ENSMUSG00000063605 / ENSG00000135736 / ENSRNOG00000025843
CCDC115 / Q96NT0 / ENSMUSG00000042111 / ENSG00000136710 / ENSRNOG00000013234
CCDC12 / Q8WUD4 / ENSMUSG00000019659 / ENSG00000160799 / ENSRNOG00000020946
CCDC124 / Q96CT7 / ENSMUSG00000007721 / ENSG00000007080 / ENSRNOG00000018932
CCDC127 / Q96BQ5 / ENSMUSG00000021578 / ENSG00000164366 / ENSRNOG00000012775
CCDC134 / Q9H6E4 / ENSMUSG00000068114 / ENSG00000100147 / ENSRNOG00000007262
CCDC136 / Q96JN2 / ENSMUSG00000029769 / ENSG00000128596 / ENSRNOG00000007100
CCDC137 / Q6PK04 / ENSMUSG00000049957 / ENSG00000185298 / ENSRNOG00000048712
CCDC141 / Q6ZP82 / ENSMUSG00000044033 / ENSG00000163492 / ENSRNOG00000012580
CCDC167 / Q9P0B6 / ENSMUSG00000024018 / ENSG00000198937 / ENSRNOG00000000531
CCDC170 / Q8IYT3 / ENSMUSG00000019767 / ENSG00000120262 / ENSRNOG00000024280
Homologs
Brain
Heart
Kidney
Liver
Lung
Testis
Organs
Bins
1
2
3
4
5
Species
Human
Mouse
Rat

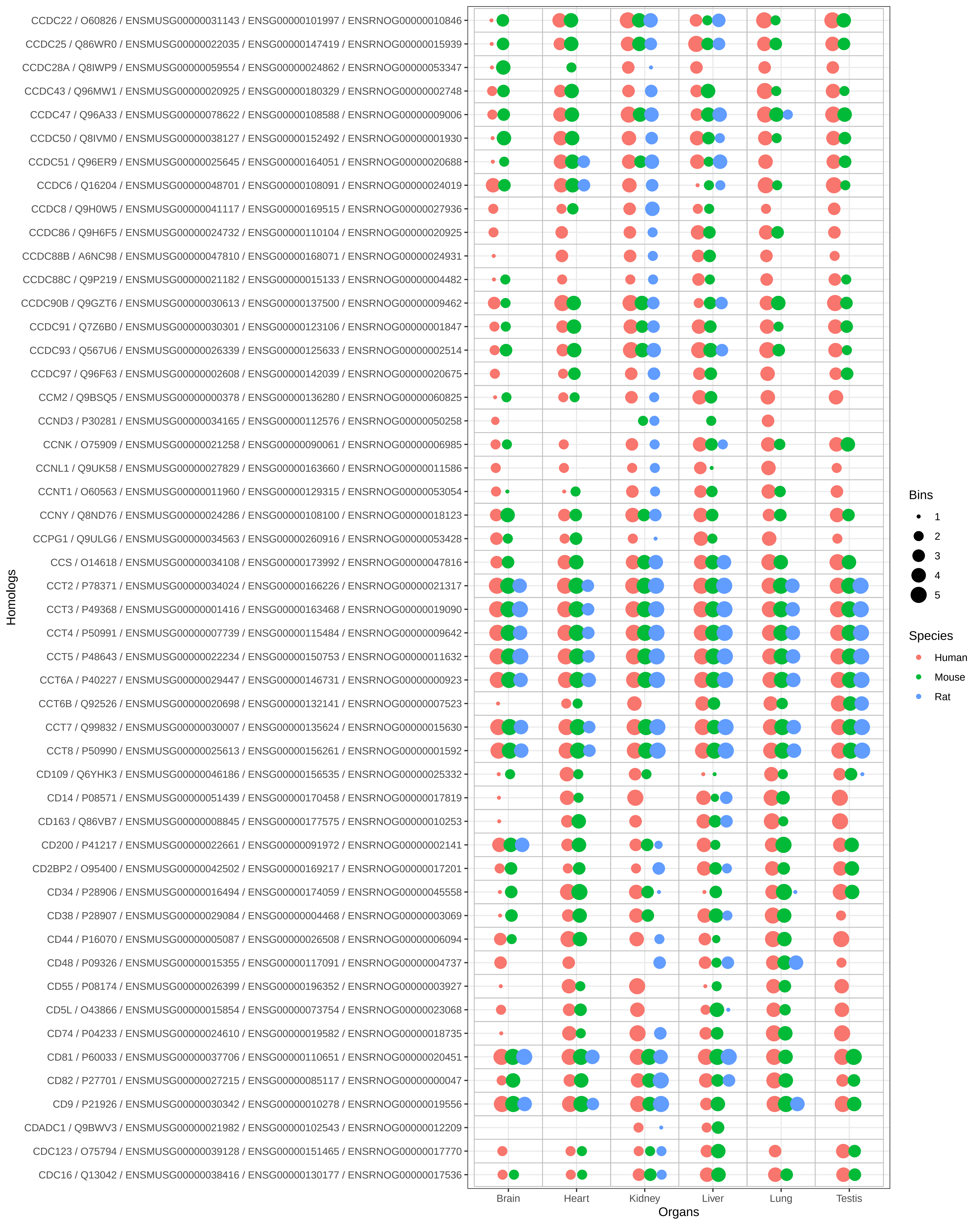
CCDC22 / O60826 / ENSMUSG00000031143 / ENSG00000101997 / ENSRNOG00000010846
CCDC25 / Q86WR0 / ENSMUSG00000022035 / ENSG00000147419 / ENSRNOG00000015939
CCDC28A / Q8IWP9 / ENSMUSG00000059554 / ENSG00000024862 / ENSRNOG00000053347
CCDC43 / Q96MW1 / ENSMUSG00000020925 / ENSG00000180329 / ENSRNOG00000002748
CCDC47 / Q96A33 / ENSMUSG00000078622 / ENSG00000108588 / ENSRNOG00000009006
CCDC50 / Q8IVM0 / ENSMUSG00000038127 / ENSG00000152492 / ENSRNOG00000001930
CCDC51 / Q96ER9 / ENSMUSG00000025645 / ENSG00000164051 / ENSRNOG00000020688
CCDC6 / Q16204 / ENSMUSG00000048701 / ENSG00000108091 / ENSRNOG00000024019
CCDC8 / Q9H0W5 / ENSMUSG00000041117 / ENSG00000169515 / ENSRNOG00000027936
CCDC86 / Q9H6F5 / ENSMUSG00000024732 / ENSG00000110104 / ENSRNOG00000020925
CCDC88B / A6NC98 / ENSMUSG00000047810 / ENSG00000168071 / ENSRNOG00000024931
CCDC88C / Q9P219 / ENSMUSG00000021182 / ENSG00000015133 / ENSRNOG00000004482
CCDC90B / Q9GZT6 / ENSMUSG00000030613 / ENSG00000137500 / ENSRNOG00000009462
CCDC91 / Q7Z6B0 / ENSMUSG00000030301 / ENSG00000123106 / ENSRNOG00000001847
CCDC93 / Q567U6 / ENSMUSG00000026339 / ENSG00000125633 / ENSRNOG00000002514
CCDC97 / Q96F63 / ENSMUSG00000002608 / ENSG00000142039 / ENSRNOG00000020675
CCM2 / Q9BSQ5 / ENSMUSG00000000378 / ENSG00000136280 / ENSRNOG00000060825
CCND3 / P30281 / ENSMUSG00000034165 / ENSG00000112576 / ENSRNOG00000050258
CCNK / O75909 / ENSMUSG00000021258 / ENSG00000090061 / ENSRNOG00000006985
CCNL1 / Q9UK58 / ENSMUSG00000027829 / ENSG00000163660 / ENSRNOG00000011586
CCNT1 / O60563 / ENSMUSG00000011960 / ENSG00000129315 / ENSRNOG00000053054
CCNY / Q8ND76 / ENSMUSG00000024286 / ENSG00000108100 / ENSRNOG00000018123
CCPG1 / Q9ULG6 / ENSMUSG00000034563 / ENSG00000260916 / ENSRNOG00000053428
CCS / O14618 / ENSMUSG00000034108 / ENSG00000173992 / ENSRNOG00000047816
CCT2 / P78371 / ENSMUSG00000034024 / ENSG00000166226 / ENSRNOG00000021317
CCT3 / P49368 / ENSMUSG00000001416 / ENSG00000163468 / ENSRNOG00000019090
CCT4 / P50991 / ENSMUSG00000007739 / ENSG00000115484 / ENSRNOG00000009642
CCT5 / P48643 / ENSMUSG00000022234 / ENSG00000150753 / ENSRNOG00000011632
CCT6A / P40227 / ENSMUSG00000029447 / ENSG00000146731 / ENSRNOG00000000923
CCT6B / Q92526 / ENSMUSG00000020698 / ENSG00000132141 / ENSRNOG00000007523
CCT7 / Q99832 / ENSMUSG00000030007 / ENSG00000135624 / ENSRNOG00000015630
CCT8 / P50990 / ENSMUSG00000025613 / ENSG00000156261 / ENSRNOG00000001592
CD109 / Q6YHK3 / ENSMUSG00000046186 / ENSG00000156535 / ENSRNOG00000025332
CD14 / P08571 / ENSMUSG00000051439 / ENSG00000170458 / ENSRNOG00000017819
CD163 / Q86VB7 / ENSMUSG00000008845 / ENSG00000177575 / ENSRNOG00000010253
CD200 / P41217 / ENSMUSG00000022661 / ENSG00000091972 / ENSRNOG00000002141
CD2BP2 / O95400 / ENSMUSG00000042502 / ENSG00000169217 / ENSRNOG00000017201
CD34 / P28906 / ENSMUSG00000016494 / ENSG00000174059 / ENSRNOG00000045558
CD38 / P28907 / ENSMUSG00000029084 / ENSG00000004468 / ENSRNOG00000003069
CD44 / P16070 / ENSMUSG00000005087 / ENSG00000026508 / ENSRNOG00000006094
CD48 / P09326 / ENSMUSG00000015355 / ENSG00000117091 / ENSRNOG00000004737
CD55 / P08174 / ENSMUSG00000026399 / ENSG00000196352 / ENSRNOG00000003927
CD5L / O43866 / ENSMUSG00000015854 / ENSG00000073754 / ENSRNOG00000023068
CD74 / P04233 / ENSMUSG00000024610 / ENSG00000019582 / ENSRNOG00000018735
CD81 / P60033 / ENSMUSG00000037706 / ENSG00000110651 / ENSRNOG00000020451
CD82 / P27701 / ENSMUSG00000027215 / ENSG00000085117 / ENSRNOG00000000047
CD9 / P21926 / ENSMUSG00000030342 / ENSG00000010278 / ENSRNOG00000019556
CDADC1 / Q9BWV3 / ENSMUSG00000021982 / ENSG00000102543 / ENSRNOG00000012209
CDC123 / O75794 / ENSMUSG00000039128 / ENSG00000151465 / ENSRNOG00000017770
CDC16 / Q13042 / ENSMUSG00000038416 / ENSG00000130177 / ENSRNOG00000017536
Homologs
Brain
Heart
Kidney
Liver
Lung
Testis
Organs
Bins
1
2
3
4
5
Species
Human
Mouse
Rat

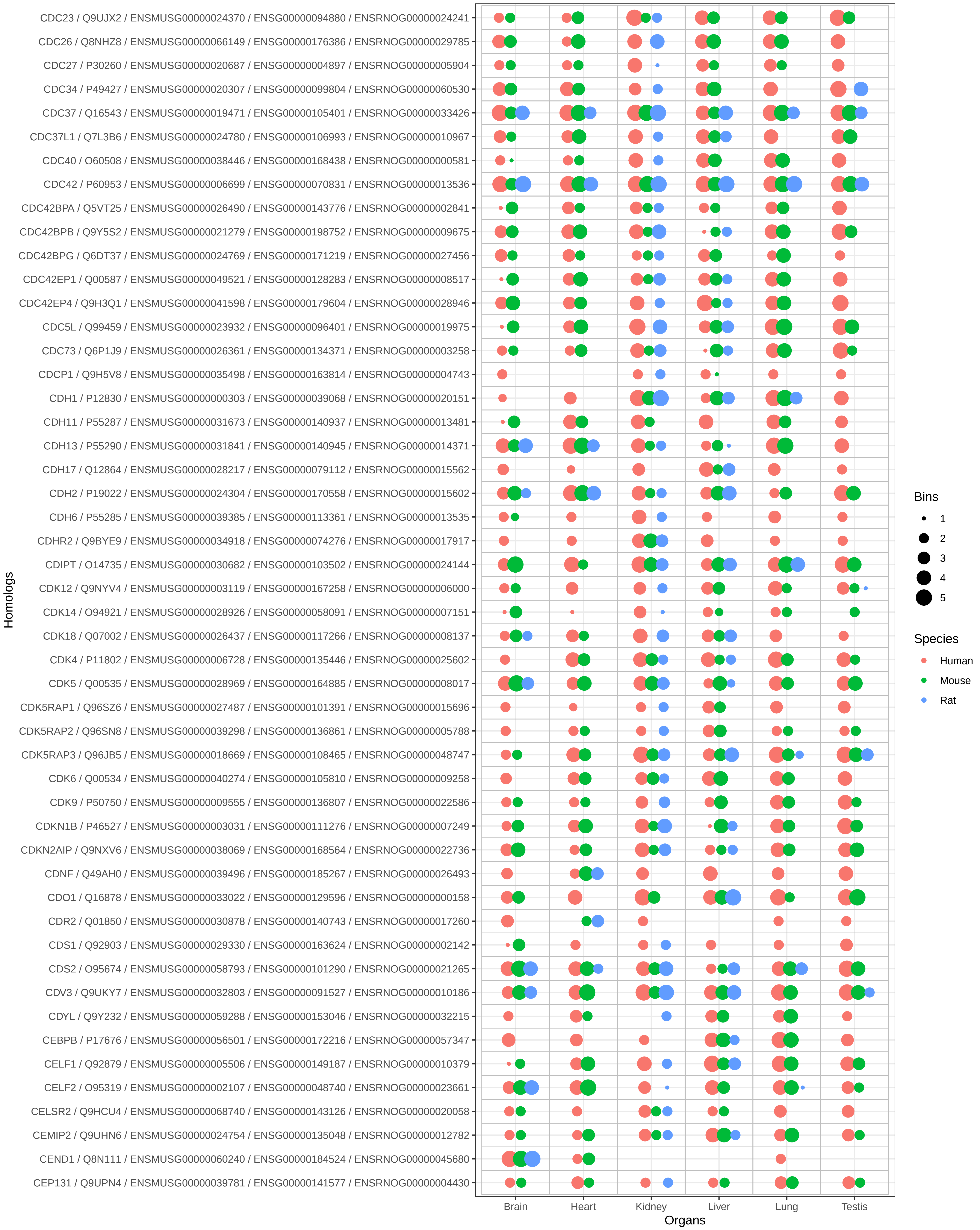

CDC23 / Q9UJX2 / ENSMUSG00000024370 / ENSG00000094880 / ENSRNOG00000024241
CDC26 / Q8NHZ8 / ENSMUSG00000066149 / ENSG00000176386 / ENSRNOG00000029785
CDC27 / P30260 / ENSMUSG00000020687 / ENSG00000004897 / ENSRNOG00000005904
CDC34 / P49427 / ENSMUSG00000020307 / ENSG00000099804 / ENSRNOG00000060530
CDC37 / Q16543 / ENSMUSG00000019471 / ENSG00000105401 / ENSRNOG00000033426
CDC37L1 / Q7L3B6 / ENSMUSG00000024780 / ENSG00000106993 / ENSRNOG00000010967
CDC40 / O60508 / ENSMUSG00000038446 / ENSG00000168438 / ENSRNOG00000000581
CDC42 / P60953 / ENSMUSG00000006699 / ENSG00000070831 / ENSRNOG00000013536
CDC42BPA / Q5VT25 / ENSMUSG00000026490 / ENSG00000143776 / ENSRNOG00000002841
CDC42BPB / Q9Y5S2 / ENSMUSG00000021279 / ENSG00000198752 / ENSRNOG00000009675
CDC42BPG / Q6DT37 / ENSMUSG00000024769 / ENSG00000171219 / ENSRNOG00000027456
CDC42EP1 / Q00587 / ENSMUSG00000049521 / ENSG00000128283 / ENSRNOG00000008517
CDC42EP4 / Q9H3Q1 / ENSMUSG00000041598 / ENSG00000179604 / ENSRNOG00000028946
CDC5L / Q99459 / ENSMUSG00000023932 / ENSG00000096401 / ENSRNOG00000019975
CDC73 / Q6P1J9 / ENSMUSG00000026361 / ENSG00000134371 / ENSRNOG00000003258
CDCP1 / Q9H5V8 / ENSMUSG00000035498 / ENSG00000163814 / ENSRNOG00000004743
CDH1 / P12830 / ENSMUSG00000000303 / ENSG00000039068 / ENSRNOG00000020151
CDH11 / P55287 / ENSMUSG00000031673 / ENSG00000140937 / ENSRNOG00000013481
CDH13 / P55290 / ENSMUSG00000031841 / ENSG00000140945 / ENSRNOG00000014371
CDH17 / Q12864 / ENSMUSG00000028217 / ENSG00000079112 / ENSRNOG00000015562
CDH2 / P19022 / ENSMUSG00000024304 / ENSG00000170558 / ENSRNOG00000015602
CDH6 / P55285 / ENSMUSG00000039385 / ENSG00000113361 / ENSRNOG00000013535
CDHR2 / Q9BYE9 / ENSMUSG00000034918 / ENSG00000074276 / ENSRNOG00000017917
CDIPT / O14735 / ENSMUSG00000030682 / ENSG00000103502 / ENSRNOG00000024144
CDK12 / Q9NYV4 / ENSMUSG00000003119 / ENSG00000167258 / ENSRNOG00000006000
CDK14 / O94921 / ENSMUSG00000028926 / ENSG00000058091 / ENSRNOG00000007151
CDK18 / Q07002 / ENSMUSG00000026437 / ENSG00000117266 / ENSRNOG00000008137
CDK4 / P11802 / ENSMUSG00000006728 / ENSG00000135446 / ENSRNOG00000025602
CDK5 / Q00535 / ENSMUSG00000028969 / ENSG00000164885 / ENSRNOG00000008017
CDK5RAP1 / Q96SZ6 / ENSMUSG00000027487 / ENSG00000101391 / ENSRNOG00000015696
CDK5RAP2 / Q96SN8 / ENSMUSG00000039298 / ENSG00000136861 / ENSRNOG00000005788
CDK5RAP3 / Q96JB5 / ENSMUSG00000018669 / ENSG00000108465 / ENSRNOG00000048747
CDK6 / Q00534 / ENSMUSG00000040274 / ENSG00000105810 / ENSRNOG00000009258
CDK9 / P50750 / ENSMUSG00000009555 / ENSG00000136807 / ENSRNOG00000022586
CDKN1B / P46527 / ENSMUSG00000003031 / ENSG00000111276 / ENSRNOG00000007249
CDKN2AIP / Q9NXV6 / ENSMUSG00000038069 / ENSG00000168564 / ENSRNOG00000022736
CDNF / Q49AH0 / ENSMUSG00000039496 / ENSG00000185267 / ENSRNOG00000026493
CDO1 / Q16878 / ENSMUSG00000033022 / ENSG00000129596 / ENSRNOG00000000158
CDR2 / Q01850 / ENSMUSG00000030878 / ENSG00000140743 / ENSRNOG00000017260
CDS1 / Q92903 / ENSMUSG00000029330 / ENSG00000163624 / ENSRNOG00000002142
CDS2 / O95674 / ENSMUSG00000058793 / ENSG00000101290 / ENSRNOG00000021265
CDV3 / Q9UKY7 / ENSMUSG00000032803 / ENSG00000091527 / ENSRNOG00000010186
CDYL / Q9Y232 / ENSMUSG00000059288 / ENSG00000153046 / ENSRNOG00000032215
CEBPB / P17676 / ENSMUSG00000056501 / ENSG00000172216 / ENSRNOG00000057347
CELF1 / Q92879 / ENSMUSG00000005506 / ENSG00000149187 / ENSRNOG00000010379
CELF2 / O95319 / ENSMUSG00000002107 / ENSG00000048740 / ENSRNOG00000023661
CELSR2 / Q9HCU4 / ENSMUSG00000068740 / ENSG00000143126 / ENSRNOG00000020058
CEMIP2 / Q9UHN6 / ENSMUSG00000024754 / ENSG00000135048 / ENSRNOG00000012782
CEND1 / Q8N111 / ENSMUSG00000060240 / ENSG00000184524 / ENSRNOG00000045680
CEP131 / Q9UPN4 / ENSMUSG00000039781 / ENSG00000141577 / ENSRNOG00000004430
Homologs
Brain
Heart
Kidney
Liver
Lung
Testis
Organs
Bins
1
2
3
4
5
Species
Human
Mouse
Rat

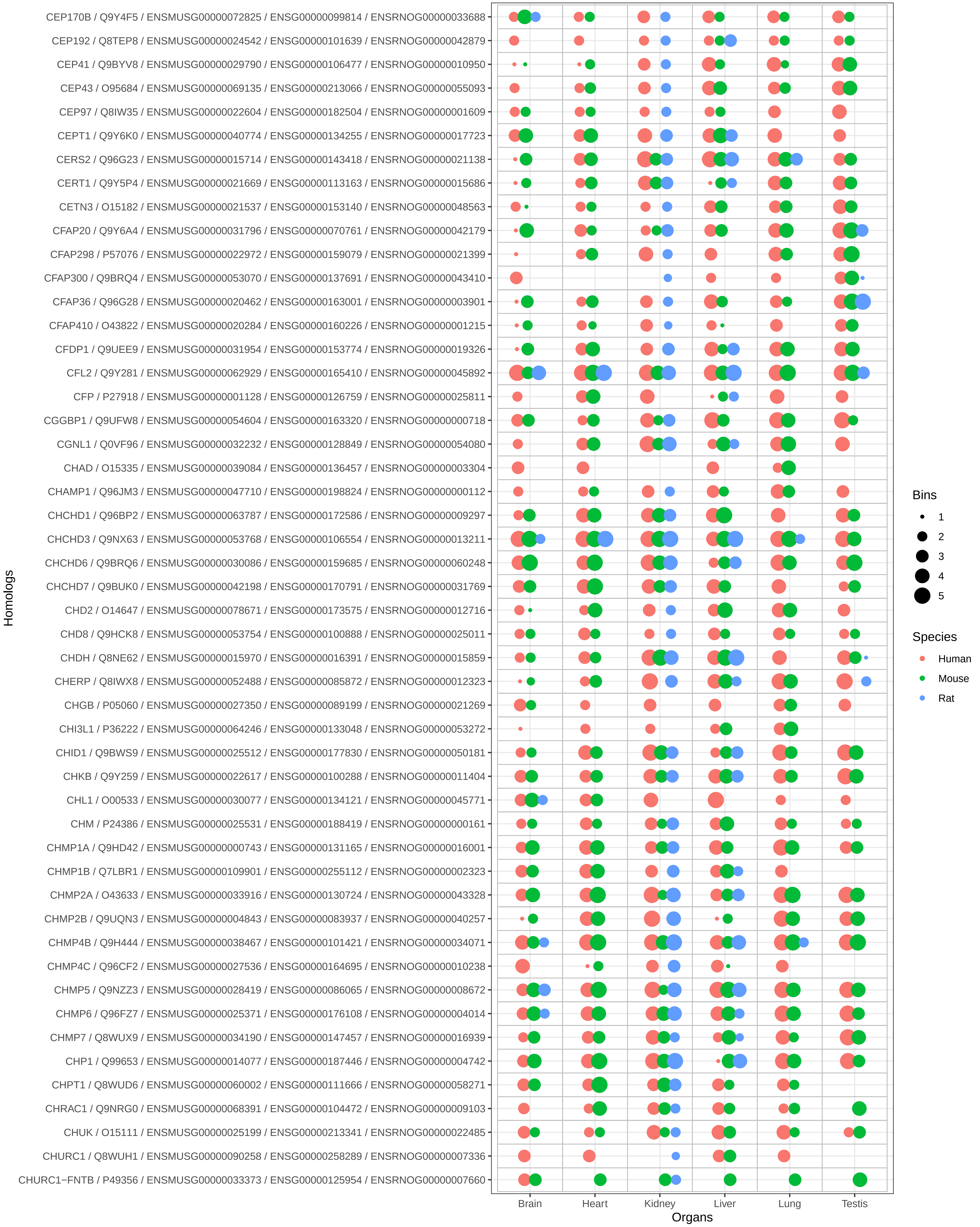
CEP170B / Q9Y4F5 / ENSMUSG00000072825 / ENSG00000099814 / ENSRNOG00000033688
CEP192 / Q8TEP8 / ENSMUSG00000024542 / ENSG00000101639 / ENSRNOG00000042879
CEP41 / Q9BYV8 / ENSMUSG00000029790 / ENSG00000106477 / ENSRNOG00000010950
CEP43 / O95684 / ENSMUSG00000069135 / ENSG00000213066 / ENSRNOG00000055093
CEP97 / Q8IW35 / ENSMUSG00000022604 / ENSG00000182504 / ENSRNOG00000001609
CEPT1 / Q9Y6K0 / ENSMUSG00000040774 / ENSG00000134255 / ENSRNOG00000017723
CERS2 / Q96G23 / ENSMUSG00000015714 / ENSG00000143418 / ENSRNOG00000021138
CERT1 / Q9Y5P4 / ENSMUSG00000021669 / ENSG00000113163 / ENSRNOG00000015686
CETN3 / O15182 / ENSMUSG00000021537 / ENSG00000153140 / ENSRNOG00000048563
CFAP20 / Q9Y6A4 / ENSMUSG00000031796 / ENSG00000070761 / ENSRNOG00000042179
CFAP298 / P57076 / ENSMUSG00000022972 / ENSG00000159079 / ENSRNOG00000021399
CFAP300 / Q9BRQ4 / ENSMUSG00000053070 / ENSG00000137691 / ENSRNOG00000043410
CFAP36 / Q96G28 / ENSMUSG00000020462 / ENSG00000163001 / ENSRNOG00000003901
CFAP410 / O43822 / ENSMUSG00000020284 / ENSG00000160226 / ENSRNOG00000001215
CFDP1 / Q9UEE9 / ENSMUSG00000031954 / ENSG00000153774 / ENSRNOG00000019326
CFL2 / Q9Y281 / ENSMUSG00000062929 / ENSG00000165410 / ENSRNOG00000045892
CFP / P27918 / ENSMUSG00000001128 / ENSG00000126759 / ENSRNOG00000025811
CGGBP1 / Q9UFW8 / ENSMUSG00000054604 / ENSG00000163320 / ENSRNOG00000000718
CGNL1 / Q0VF96 / ENSMUSG00000032232 / ENSG00000128849 / ENSRNOG00000054080
CHAD / O15335 / ENSMUSG00000039084 / ENSG00000136457 / ENSRNOG00000003304
CHAMP1 / Q96JM3 / ENSMUSG00000047710 / ENSG00000198824 / ENSRNOG00000000112
CHCHD1 / Q96BP2 / ENSMUSG00000063787 / ENSG00000172586 / ENSRNOG00000009297
CHCHD3 / Q9NX63 / ENSMUSG00000053768 / ENSG00000106554 / ENSRNOG00000013211
CHCHD6 / Q9BRQ6 / ENSMUSG00000030086 / ENSG00000159685 / ENSRNOG00000060248
CHCHD7 / Q9BUK0 / ENSMUSG00000042198 / ENSG00000170791 / ENSRNOG00000031769
CHD2 / O14647 / ENSMUSG00000078671 / ENSG00000173575 / ENSRNOG00000012716
CHD8 / Q9HCK8 / ENSMUSG00000053754 / ENSG00000100888 / ENSRNOG00000025011
CHDH / Q8NE62 / ENSMUSG00000015970 / ENSG00000016391 / ENSRNOG00000015859
CHERP / Q8IWX8 / ENSMUSG00000052488 / ENSG00000085872 / ENSRNOG00000012323
CHGB / P05060 / ENSMUSG00000027350 / ENSG00000089199 / ENSRNOG00000021269
CHI3L1 / P36222 / ENSMUSG00000064246 / ENSG00000133048 / ENSRNOG00000053272
CHID1 / Q9BWS9 / ENSMUSG00000025512 / ENSG00000177830 / ENSRNOG00000050181
CHKB / Q9Y259 / ENSMUSG00000022617 / ENSG00000100288 / ENSRNOG00000011404
CHL1 / O00533 / ENSMUSG00000030077 / ENSG00000134121 / ENSRNOG00000045771
CHM / P24386 / ENSMUSG00000025531 / ENSG00000188419 / ENSRNOG00000000161
CHMP1A / Q9HD42 / ENSMUSG00000000743 / ENSG00000131165 / ENSRNOG00000016001
CHMP1B / Q7LBR1 / ENSMUSG00000109901 / ENSG00000255112 / ENSRNOG00000002323
CHMP2A / O43633 / ENSMUSG00000033916 / ENSG00000130724 / ENSRNOG00000043328
CHMP2B / Q9UQN3 / ENSMUSG00000004843 / ENSG00000083937 / ENSRNOG00000040257
CHMP4B / Q9H444 / ENSMUSG00000038467 / ENSG00000101421 / ENSRNOG00000034071
CHMP4C / Q96CF2 / ENSMUSG00000027536 / ENSG00000164695 / ENSRNOG00000010238
CHMP5 / Q9NZZ3 / ENSMUSG00000028419 / ENSG00000086065 / ENSRNOG00000008672
CHMP6 / Q96FZ7 / ENSMUSG00000025371 / ENSG00000176108 / ENSRNOG00000004014
CHMP7 / Q8WUX9 / ENSMUSG00000034190 / ENSG00000147457 / ENSRNOG00000016939
CHP1 / Q99653 / ENSMUSG00000014077 / ENSG00000187446 / ENSRNOG00000004742
CHPT1 / Q8WUD6 / ENSMUSG00000060002 / ENSG00000111666 / ENSRNOG00000058271
CHRAC1 / Q9NRG0 / ENSMUSG00000068391 / ENSG00000104472 / ENSRNOG00000009103
CHUK / O15111 / ENSMUSG00000025199 / ENSG00000213341 / ENSRNOG00000022485
CHURC1 / Q8WUH1 / ENSMUSG00000090258 / ENSG00000258289 / ENSRNOG00000007336
CHURC1−FNTB / P49356 / ENSMUSG00000033373 / ENSG00000125954 / ENSRNOG00000007660
Homologs
Brain
Heart
Kidney
Liver
Lung
Testis
Organs
Bins
1
2
3
4
5
Species
Human
Mouse
Rat

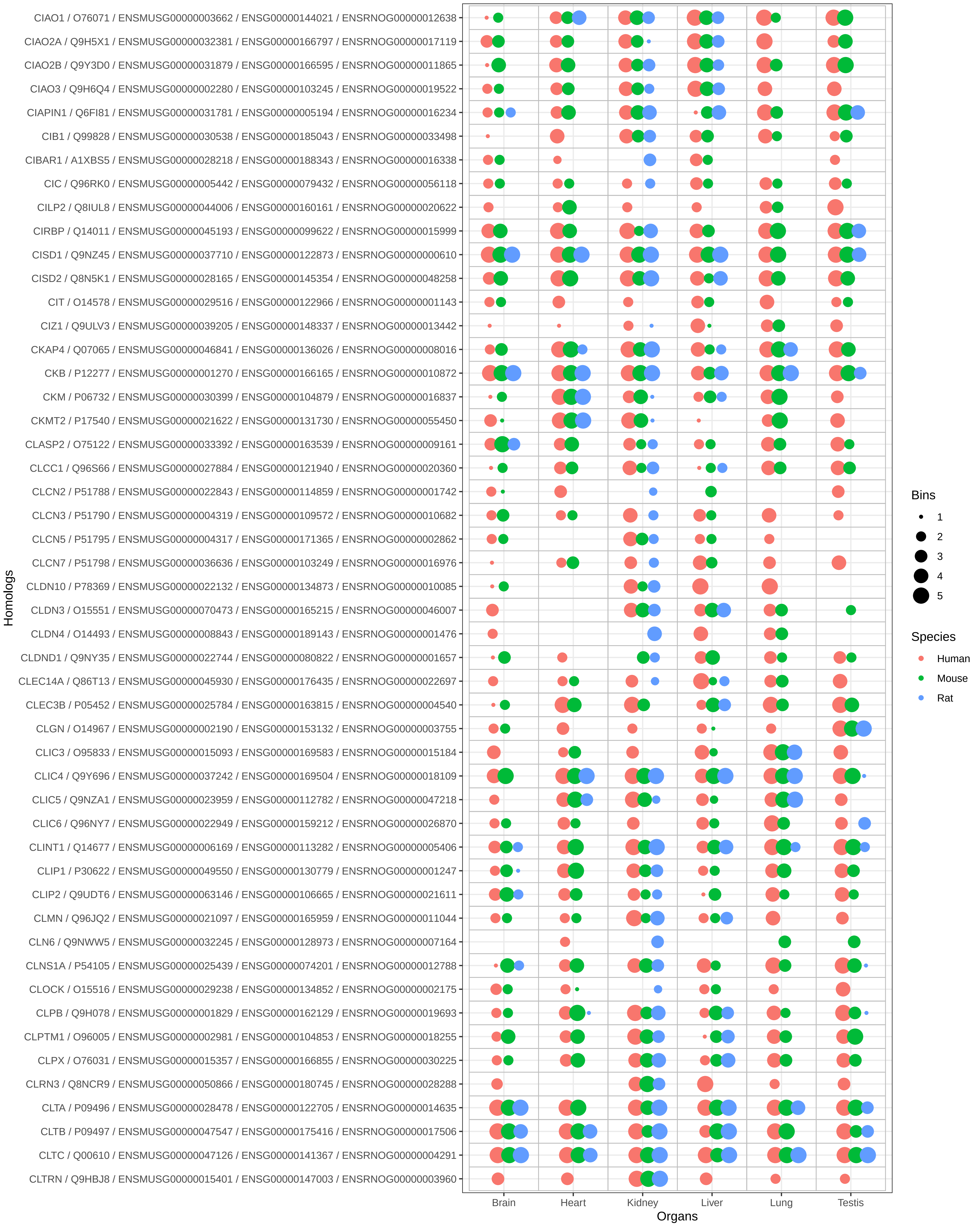
CIAO1 / O76071 / ENSMUSG00000003662 / ENSG00000144021 / ENSRNOG00000012638
CIAO2A / Q9H5X1 / ENSMUSG00000032381 / ENSG00000166797 / ENSRNOG00000017119
CIAO2B / Q9Y3D0 / ENSMUSG00000031879 / ENSG00000166595 / ENSRNOG00000011865
CIAO3 / Q9H6Q4 / ENSMUSG00000002280 / ENSG00000103245 / ENSRNOG00000019522
CIAPIN1 / Q6FI81 / ENSMUSG00000031781 / ENSG00000005194 / ENSRNOG00000016234
CIB1 / Q99828 / ENSMUSG00000030538 / ENSG00000185043 / ENSRNOG00000033498
CIBAR1 / A1XBS5 / ENSMUSG00000028218 / ENSG00000188343 / ENSRNOG00000016338
CIC / Q96RK0 / ENSMUSG00000005442 / ENSG00000079432 / ENSRNOG00000056118
CILP2 / Q8IUL8 / ENSMUSG00000044006 / ENSG00000160161 / ENSRNOG00000020622
CIRBP / Q14011 / ENSMUSG00000045193 / ENSG00000099622 / ENSRNOG00000015999
CISD1 / Q9NZ45 / ENSMUSG00000037710 / ENSG00000122873 / ENSRNOG00000000610
CISD2 / Q8N5K1 / ENSMUSG00000028165 / ENSG00000145354 / ENSRNOG00000048258
CIT / O14578 / ENSMUSG00000029516 / ENSG00000122966 / ENSRNOG00000001143
CIZ1 / Q9ULV3 / ENSMUSG00000039205 / ENSG00000148337 / ENSRNOG00000013442
CKAP4 / Q07065 / ENSMUSG00000046841 / ENSG00000136026 / ENSRNOG00000008016
CKB / P12277 / ENSMUSG00000001270 / ENSG00000166165 / ENSRNOG00000010872
CKM / P06732 / ENSMUSG00000030399 / ENSG00000104879 / ENSRNOG00000016837
CKMT2 / P17540 / ENSMUSG00000021622 / ENSG00000131730 / ENSRNOG00000055450
CLASP2 / O75122 / ENSMUSG00000033392 / ENSG00000163539 / ENSRNOG00000009161
CLCC1 / Q96S66 / ENSMUSG00000027884 / ENSG00000121940 / ENSRNOG00000020360
CLCN2 / P51788 / ENSMUSG00000022843 / ENSG00000114859 / ENSRNOG00000001742
CLCN3 / P51790 / ENSMUSG00000004319 / ENSG00000109572 / ENSRNOG00000010682
CLCN5 / P51795 / ENSMUSG00000004317 / ENSG00000171365 / ENSRNOG00000002862
CLCN7 / P51798 / ENSMUSG00000036636 / ENSG00000103249 / ENSRNOG00000016976
CLDN10 / P78369 / ENSMUSG00000022132 / ENSG00000134873 / ENSRNOG00000010085
CLDN3 / O15551 / ENSMUSG00000070473 / ENSG00000165215 / ENSRNOG00000046007
CLDN4 / O14493 / ENSMUSG00000008843 / ENSG00000189143 / ENSRNOG00000001476
CLDND1 / Q9NY35 / ENSMUSG00000022744 / ENSG00000080822 / ENSRNOG00000001657
CLEC14A / Q86T13 / ENSMUSG00000045930 / ENSG00000176435 / ENSRNOG00000022697
CLEC3B / P05452 / ENSMUSG00000025784 / ENSG00000163815 / ENSRNOG00000004540
CLGN / O14967 / ENSMUSG00000002190 / ENSG00000153132 / ENSRNOG00000003755
CLIC3 / O95833 / ENSMUSG00000015093 / ENSG00000169583 / ENSRNOG00000015184
CLIC4 / Q9Y696 / ENSMUSG00000037242 / ENSG00000169504 / ENSRNOG00000018109
CLIC5 / Q9NZA1 / ENSMUSG00000023959 / ENSG00000112782 / ENSRNOG00000047218
CLIC6 / Q96NY7 / ENSMUSG00000022949 / ENSG00000159212 / ENSRNOG00000026870
CLINT1 / Q14677 / ENSMUSG00000006169 / ENSG00000113282 / ENSRNOG00000005406
CLIP1 / P30622 / ENSMUSG00000049550 / ENSG00000130779 / ENSRNOG00000001247
CLIP2 / Q9UDT6 / ENSMUSG00000063146 / ENSG00000106665 / ENSRNOG00000021611
CLMN / Q96JQ2 / ENSMUSG00000021097 / ENSG00000165959 / ENSRNOG00000011044
CLN6 / Q9NWW5 / ENSMUSG00000032245 / ENSG00000128973 / ENSRNOG00000007164
CLNS1A / P54105 / ENSMUSG00000025439 / ENSG00000074201 / ENSRNOG00000012788
CLOCK / O15516 / ENSMUSG00000029238 / ENSG00000134852 / ENSRNOG00000002175
CLPB / Q9H078 / ENSMUSG00000001829 / ENSG00000162129 / ENSRNOG00000019693
CLPTM1 / O96005 / ENSMUSG00000002981 / ENSG00000104853 / ENSRNOG00000018255
CLPX / O76031 / ENSMUSG00000015357 / ENSG00000166855 / ENSRNOG00000030225
CLRN3 / Q8NCR9 / ENSMUSG00000050866 / ENSG00000180745 / ENSRNOG00000028288
CLTA / P09496 / ENSMUSG00000028478 / ENSG00000122705 / ENSRNOG00000014635
CLTB / P09497 / ENSMUSG00000047547 / ENSG00000175416 / ENSRNOG00000017506
CLTC / Q00610 / ENSMUSG00000047126 / ENSG00000141367 / ENSRNOG00000004291
CLTRN / Q9HBJ8 / ENSMUSG00000015401 / ENSG00000147003 / ENSRNOG00000003960
Homologs
Brain
Heart
Kidney
Liver
Lung
Testis
Organs
Bins
1
2
3
4
5
Species
Human
Mouse
Rat

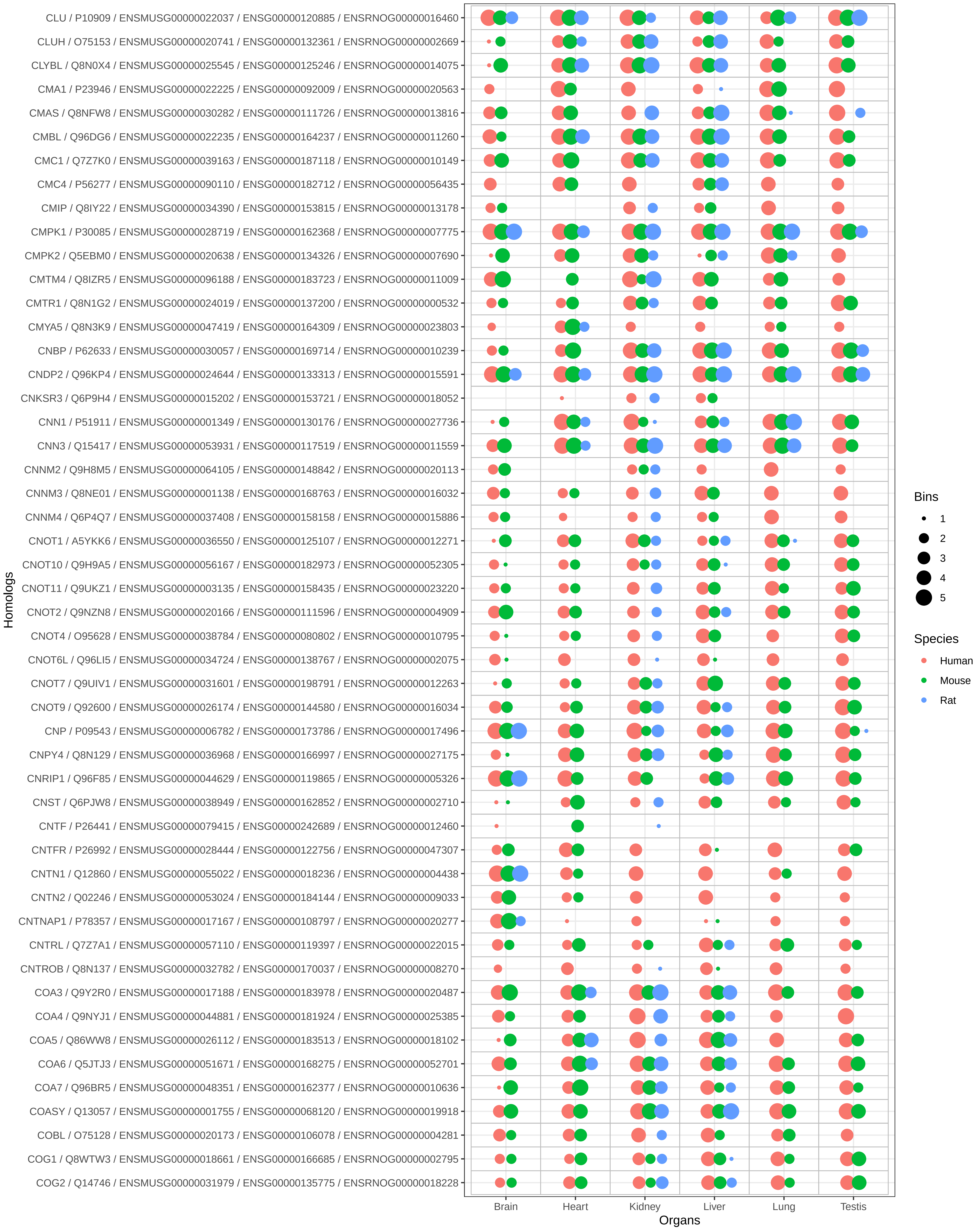
CLU / P10909 / ENSMUSG00000022037 / ENSG00000120885 / ENSRNOG00000016460
CLUH / O75153 / ENSMUSG00000020741 / ENSG00000132361 / ENSRNOG00000002669
CLYBL / Q8N0X4 / ENSMUSG00000025545 / ENSG00000125246 / ENSRNOG00000014075
CMA1 / P23946 / ENSMUSG00000022225 / ENSG00000092009 / ENSRNOG00000020563
CMAS / Q8NFW8 / ENSMUSG00000030282 / ENSG00000111726 / ENSRNOG00000013816
CMBL / Q96DG6 / ENSMUSG00000022235 / ENSG00000164237 / ENSRNOG00000011260
CMC1 / Q7Z7K0 / ENSMUSG00000039163 / ENSG00000187118 / ENSRNOG00000010149
CMC4 / P56277 / ENSMUSG00000090110 / ENSG00000182712 / ENSRNOG00000056435
CMIP / Q8IY22 / ENSMUSG00000034390 / ENSG00000153815 / ENSRNOG00000013178
CMPK1 / P30085 / ENSMUSG00000028719 / ENSG00000162368 / ENSRNOG00000007775
CMPK2 / Q5EBM0 / ENSMUSG00000020638 / ENSG00000134326 / ENSRNOG00000007690
CMTM4 / Q8IZR5 / ENSMUSG00000096188 / ENSG00000183723 / ENSRNOG00000011009
CMTR1 / Q8N1G2 / ENSMUSG00000024019 / ENSG00000137200 / ENSRNOG00000000532
CMYA5 / Q8N3K9 / ENSMUSG00000047419 / ENSG00000164309 / ENSRNOG00000023803
CNBP / P62633 / ENSMUSG00000030057 / ENSG00000169714 / ENSRNOG00000010239
CNDP2 / Q96KP4 / ENSMUSG00000024644 / ENSG00000133313 / ENSRNOG00000015591
CNKSR3 / Q6P9H4 / ENSMUSG00000015202 / ENSG00000153721 / ENSRNOG00000018052
CNN1 / P51911 / ENSMUSG00000001349 / ENSG00000130176 / ENSRNOG00000027736
CNN3 / Q15417 / ENSMUSG00000053931 / ENSG00000117519 / ENSRNOG00000011559
CNNM2 / Q9H8M5 / ENSMUSG00000064105 / ENSG00000148842 / ENSRNOG00000020113
CNNM3 / Q8NE01 / ENSMUSG00000001138 / ENSG00000168763 / ENSRNOG00000016032
CNNM4 / Q6P4Q7 / ENSMUSG00000037408 / ENSG00000158158 / ENSRNOG00000015886
CNOT1 / A5YKK6 / ENSMUSG00000036550 / ENSG00000125107 / ENSRNOG00000012271
CNOT10 / Q9H9A5 / ENSMUSG00000056167 / ENSG00000182973 / ENSRNOG00000052305
CNOT11 / Q9UKZ1 / ENSMUSG00000003135 / ENSG00000158435 / ENSRNOG00000023220
CNOT2 / Q9NZN8 / ENSMUSG00000020166 / ENSG00000111596 / ENSRNOG00000004909
CNOT4 / O95628 / ENSMUSG00000038784 / ENSG00000080802 / ENSRNOG00000010795
CNOT6L / Q96LI5 / ENSMUSG00000034724 / ENSG00000138767 / ENSRNOG00000002075
CNOT7 / Q9UIV1 / ENSMUSG00000031601 / ENSG00000198791 / ENSRNOG00000012263
CNOT9 / Q92600 / ENSMUSG00000026174 / ENSG00000144580 / ENSRNOG00000016034
CNP / P09543 / ENSMUSG00000006782 / ENSG00000173786 / ENSRNOG00000017496
CNPY4 / Q8N129 / ENSMUSG00000036968 / ENSG00000166997 / ENSRNOG00000027175
CNRIP1 / Q96F85 / ENSMUSG00000044629 / ENSG00000119865 / ENSRNOG00000005326
CNST / Q6PJW8 / ENSMUSG00000038949 / ENSG00000162852 / ENSRNOG00000002710
CNTF / P26441 / ENSMUSG00000079415 / ENSG00000242689 / ENSRNOG00000012460
CNTFR / P26992 / ENSMUSG00000028444 / ENSG00000122756 / ENSRNOG00000047307
CNTN1 / Q12860 / ENSMUSG00000055022 / ENSG00000018236 / ENSRNOG00000004438
CNTN2 / Q02246 / ENSMUSG00000053024 / ENSG00000184144 / ENSRNOG00000009033
CNTNAP1 / P78357 / ENSMUSG00000017167 / ENSG00000108797 / ENSRNOG00000020277
CNTRL / Q7Z7A1 / ENSMUSG00000057110 / ENSG00000119397 / ENSRNOG00000022015
CNTROB / Q8N137 / ENSMUSG00000032782 / ENSG00000170037 / ENSRNOG00000008270
COA3 / Q9Y2R0 / ENSMUSG00000017188 / ENSG00000183978 / ENSRNOG00000020487
COA4 / Q9NYJ1 / ENSMUSG00000044881 / ENSG00000181924 / ENSRNOG00000025385
COA5 / Q86WW8 / ENSMUSG00000026112 / ENSG00000183513 / ENSRNOG00000018102
COA6 / Q5JTJ3 / ENSMUSG00000051671 / ENSG00000168275 / ENSRNOG00000052701
COA7 / Q96BR5 / ENSMUSG00000048351 / ENSG00000162377 / ENSRNOG00000010636
COASY / Q13057 / ENSMUSG00000001755 / ENSG00000068120 / ENSRNOG00000019918
COBL / O75128 / ENSMUSG00000020173 / ENSG00000106078 / ENSRNOG00000004281
COG1 / Q8WTW3 / ENSMUSG00000018661 / ENSG00000166685 / ENSRNOG00000002795
COG2 / Q14746 / ENSMUSG00000031979 / ENSG00000135775 / ENSRNOG00000018228
Homologs
Brain
Heart
Kidney
Liver
Lung
Testis
Organs
Bins
1
2
3
4
5
Species
Human
Mouse
Rat

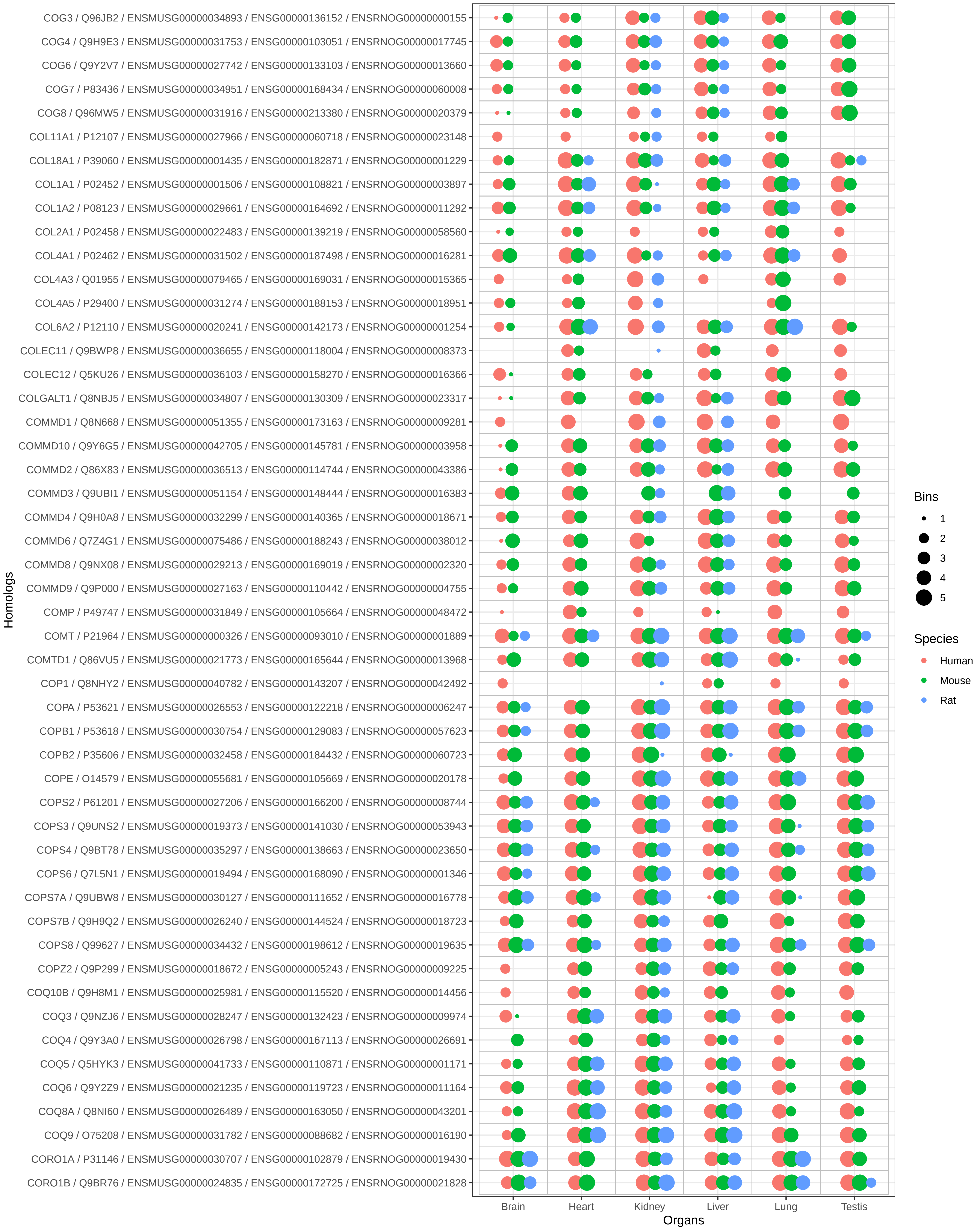

COG3 / Q96JB2 / ENSMUSG00000034893 / ENSG00000136152 / ENSRNOG00000000155
COG4 / Q9H9E3 / ENSMUSG00000031753 / ENSG00000103051 / ENSRNOG00000017745
COG6 / Q9Y2V7 / ENSMUSG00000027742 / ENSG00000133103 / ENSRNOG00000013660
COG7 / P83436 / ENSMUSG00000034951 / ENSG00000168434 / ENSRNOG00000060008
COG8 / Q96MW5 / ENSMUSG00000031916 / ENSG00000213380 / ENSRNOG00000020379
COL11A1 / P12107 / ENSMUSG00000027966 / ENSG00000060718 / ENSRNOG00000023148
COL18A1 / P39060 / ENSMUSG00000001435 / ENSG00000182871 / ENSRNOG00000001229
COL1A1 / P02452 / ENSMUSG00000001506 / ENSG00000108821 / ENSRNOG00000003897
COL1A2 / P08123 / ENSMUSG00000029661 / ENSG00000164692 / ENSRNOG00000011292
COL2A1 / P02458 / ENSMUSG00000022483 / ENSG00000139219 / ENSRNOG00000058560
COL4A1 / P02462 / ENSMUSG00000031502 / ENSG00000187498 / ENSRNOG00000016281
COL4A3 / Q01955 / ENSMUSG00000079465 / ENSG00000169031 / ENSRNOG00000015365
COL4A5 / P29400 / ENSMUSG00000031274 / ENSG00000188153 / ENSRNOG00000018951
COL6A2 / P12110 / ENSMUSG00000020241 / ENSG00000142173 / ENSRNOG00000001254
COLEC11 / Q9BWP8 / ENSMUSG00000036655 / ENSG00000118004 / ENSRNOG00000008373
COLEC12 / Q5KU26 / ENSMUSG00000036103 / ENSG00000158270 / ENSRNOG00000016366
COLGALT1 / Q8NBJ5 / ENSMUSG00000034807 / ENSG00000130309 / ENSRNOG00000023317
COMMD1 / Q8N668 / ENSMUSG00000051355 / ENSG00000173163 / ENSRNOG00000009281
COMMD10 / Q9Y6G5 / ENSMUSG00000042705 / ENSG00000145781 / ENSRNOG00000003958
COMMD2 / Q86X83 / ENSMUSG00000036513 / ENSG00000114744 / ENSRNOG00000043386
COMMD3 / Q9UBI1 / ENSMUSG00000051154 / ENSG00000148444 / ENSRNOG00000016383
COMMD4 / Q9H0A8 / ENSMUSG00000032299 / ENSG00000140365 / ENSRNOG00000018671
COMMD6 / Q7Z4G1 / ENSMUSG00000075486 / ENSG00000188243 / ENSRNOG00000038012
COMMD8 / Q9NX08 / ENSMUSG00000029213 / ENSG00000169019 / ENSRNOG00000002320
COMMD9 / Q9P000 / ENSMUSG00000027163 / ENSG00000110442 / ENSRNOG00000004755
COMP / P49747 / ENSMUSG00000031849 / ENSG00000105664 / ENSRNOG00000048472
COMT / P21964 / ENSMUSG00000000326 / ENSG00000093010 / ENSRNOG00000001889
COMTD1 / Q86VU5 / ENSMUSG00000021773 / ENSG00000165644 / ENSRNOG00000013968
COP1 / Q8NHY2 / ENSMUSG00000040782 / ENSG00000143207 / ENSRNOG00000042492
COPA / P53621 / ENSMUSG00000026553 / ENSG00000122218 / ENSRNOG00000006247
COPB1 / P53618 / ENSMUSG00000030754 / ENSG00000129083 / ENSRNOG00000057623
COPB2 / P35606 / ENSMUSG00000032458 / ENSG00000184432 / ENSRNOG00000060723
COPE / O14579 / ENSMUSG00000055681 / ENSG00000105669 / ENSRNOG00000020178
COPS2 / P61201 / ENSMUSG00000027206 / ENSG00000166200 / ENSRNOG00000008744
COPS3 / Q9UNS2 / ENSMUSG00000019373 / ENSG00000141030 / ENSRNOG00000053943
COPS4 / Q9BT78 / ENSMUSG00000035297 / ENSG00000138663 / ENSRNOG00000023650
COPS6 / Q7L5N1 / ENSMUSG00000019494 / ENSG00000168090 / ENSRNOG00000001346
COPS7A / Q9UBW8 / ENSMUSG00000030127 / ENSG00000111652 / ENSRNOG00000016778
COPS7B / Q9H9Q2 / ENSMUSG00000026240 / ENSG00000144524 / ENSRNOG00000018723
COPS8 / Q99627 / ENSMUSG00000034432 / ENSG00000198612 / ENSRNOG00000019635
COPZ2 / Q9P299 / ENSMUSG00000018672 / ENSG00000005243 / ENSRNOG00000009225
COQ10B / Q9H8M1 / ENSMUSG00000025981 / ENSG00000115520 / ENSRNOG00000014456
COQ3 / Q9NZJ6 / ENSMUSG00000028247 / ENSG00000132423 / ENSRNOG00000009974
COQ4 / Q9Y3A0 / ENSMUSG00000026798 / ENSG00000167113 / ENSRNOG00000026691
COQ5 / Q5HYK3 / ENSMUSG00000041733 / ENSG00000110871 / ENSRNOG00000001171
COQ6 / Q9Y2Z9 / ENSMUSG00000021235 / ENSG00000119723 / ENSRNOG00000011164
COQ8A / Q8NI60 / ENSMUSG00000026489 / ENSG00000163050 / ENSRNOG00000043201
COQ9 / O75208 / ENSMUSG00000031782 / ENSG00000088682 / ENSRNOG00000016190
CORO1A / P31146 / ENSMUSG00000030707 / ENSG00000102879 / ENSRNOG00000019430
CORO1B / Q9BR76 / ENSMUSG00000024835 / ENSG00000172725 / ENSRNOG00000021828
Homologs
Brain
Heart
Kidney
Liver
Lung
Testis
Organs
Bins
1
2
3
4
5
Species
Human
Mouse
Rat

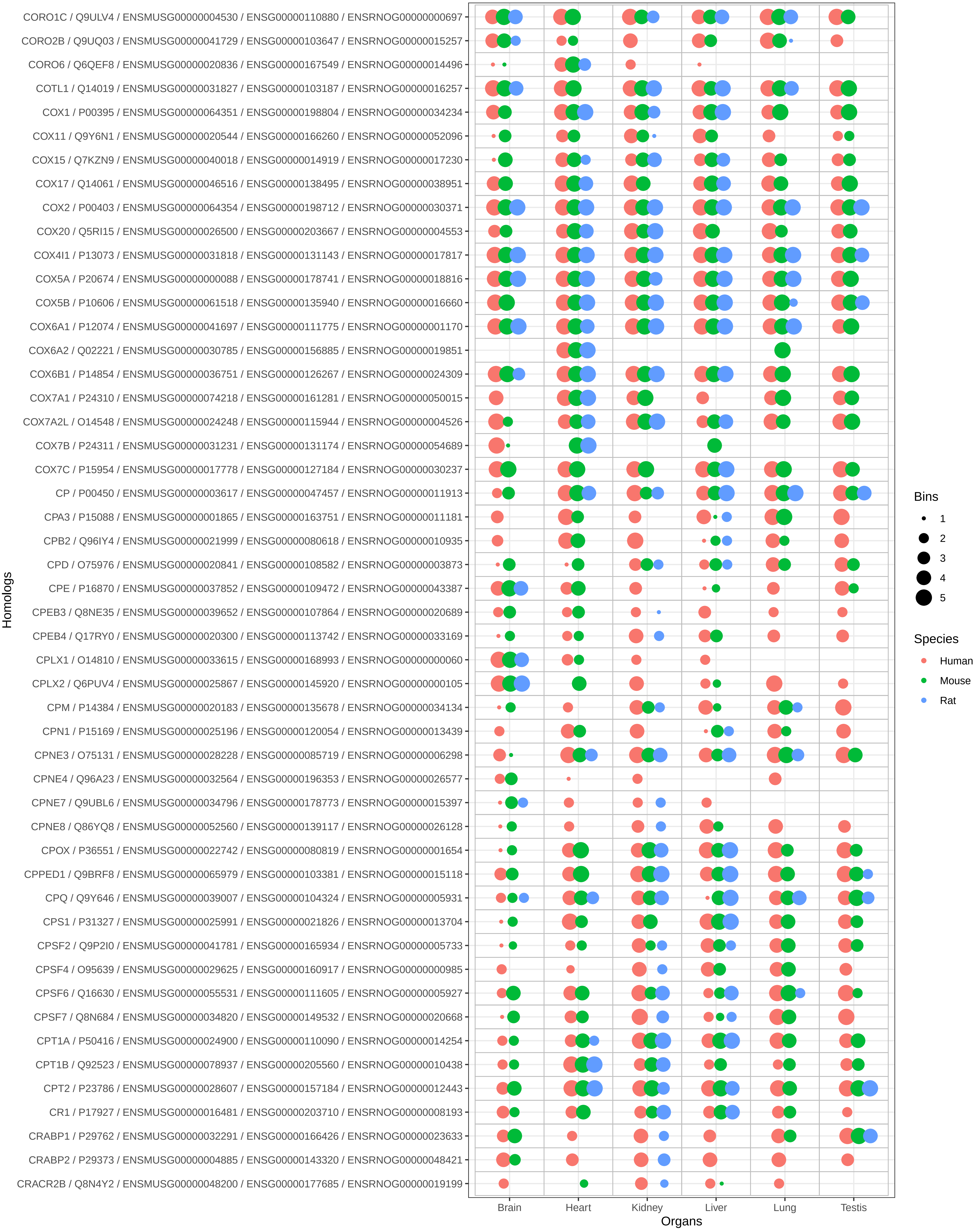

CORO1C / Q9ULV4 / ENSMUSG00000004530 / ENSG00000110880 / ENSRNOG00000000697
CORO2B / Q9UQ03 / ENSMUSG00000041729 / ENSG00000103647 / ENSRNOG00000015257
CORO6 / Q6QEF8 / ENSMUSG00000020836 / ENSG00000167549 / ENSRNOG00000014496
COTL1 / Q14019 / ENSMUSG00000031827 / ENSG00000103187 / ENSRNOG00000016257
COX1 / P00395 / ENSMUSG00000064351 / ENSG00000198804 / ENSRNOG00000034234
COX11 / Q9Y6N1 / ENSMUSG00000020544 / ENSG00000166260 / ENSRNOG00000052096
COX15 / Q7KZN9 / ENSMUSG00000040018 / ENSG00000014919 / ENSRNOG00000017230
COX17 / Q14061 / ENSMUSG00000046516 / ENSG00000138495 / ENSRNOG00000038951
COX2 / P00403 / ENSMUSG00000064354 / ENSG00000198712 / ENSRNOG00000030371
COX20 / Q5RI15 / ENSMUSG00000026500 / ENSG00000203667 / ENSRNOG00000004553
COX4I1 / P13073 / ENSMUSG00000031818 / ENSG00000131143 / ENSRNOG00000017817
COX5A / P20674 / ENSMUSG00000000088 / ENSG00000178741 / ENSRNOG00000018816
COX5B / P10606 / ENSMUSG00000061518 / ENSG00000135940 / ENSRNOG00000016660
COX6A1 / P12074 / ENSMUSG00000041697 / ENSG00000111775 / ENSRNOG00000001170
COX6A2 / Q02221 / ENSMUSG00000030785 / ENSG00000156885 / ENSRNOG00000019851
COX6B1 / P14854 / ENSMUSG00000036751 / ENSG00000126267 / ENSRNOG00000024309
COX7A1 / P24310 / ENSMUSG00000074218 / ENSG00000161281 / ENSRNOG00000050015
COX7A2L / O14548 / ENSMUSG00000024248 / ENSG00000115944 / ENSRNOG00000004526
COX7B / P24311 / ENSMUSG00000031231 / ENSG00000131174 / ENSRNOG00000054689
COX7C / P15954 / ENSMUSG00000017778 / ENSG00000127184 / ENSRNOG00000030237
CP / P00450 / ENSMUSG00000003617 / ENSG00000047457 / ENSRNOG00000011913
CPA3 / P15088 / ENSMUSG00000001865 / ENSG00000163751 / ENSRNOG00000011181
CPB2 / Q96IY4 / ENSMUSG00000021999 / ENSG00000080618 / ENSRNOG00000010935
CPD / O75976 / ENSMUSG00000020841 / ENSG00000108582 / ENSRNOG00000003873
CPE / P16870 / ENSMUSG00000037852 / ENSG00000109472 / ENSRNOG00000043387
CPEB3 / Q8NE35 / ENSMUSG00000039652 / ENSG00000107864 / ENSRNOG00000020689
CPEB4 / Q17RY0 / ENSMUSG00000020300 / ENSG00000113742 / ENSRNOG00000033169
CPLX1 / O14810 / ENSMUSG00000033615 / ENSG00000168993 / ENSRNOG00000000060
CPLX2 / Q6PUV4 / ENSMUSG00000025867 / ENSG00000145920 / ENSRNOG00000000105
CPM / P14384 / ENSMUSG00000020183 / ENSG00000135678 / ENSRNOG00000034134
CPN1 / P15169 / ENSMUSG00000025196 / ENSG00000120054 / ENSRNOG00000013439
CPNE3 / O75131 / ENSMUSG00000028228 / ENSG00000085719 / ENSRNOG00000006298
CPNE4 / Q96A23 / ENSMUSG00000032564 / ENSG00000196353 / ENSRNOG00000026577
CPNE7 / Q9UBL6 / ENSMUSG00000034796 / ENSG00000178773 / ENSRNOG00000015397
CPNE8 / Q86YQ8 / ENSMUSG00000052560 / ENSG00000139117 / ENSRNOG00000026128
CPOX / P36551 / ENSMUSG00000022742 / ENSG00000080819 / ENSRNOG00000001654
CPPED1 / Q9BRF8 / ENSMUSG00000065979 / ENSG00000103381 / ENSRNOG00000015118
CPQ / Q9Y646 / ENSMUSG00000039007 / ENSG00000104324 / ENSRNOG00000005931
CPS1 / P31327 / ENSMUSG00000025991 / ENSG00000021826 / ENSRNOG00000013704
CPSF2 / Q9P2I0 / ENSMUSG00000041781 / ENSG00000165934 / ENSRNOG00000005733
CPSF4 / O95639 / ENSMUSG00000029625 / ENSG00000160917 / ENSRNOG00000000985
CPSF6 / Q16630 / ENSMUSG00000055531 / ENSG00000111605 / ENSRNOG00000005927
CPSF7 / Q8N684 / ENSMUSG00000034820 / ENSG00000149532 / ENSRNOG00000020668
CPT1A / P50416 / ENSMUSG00000024900 / ENSG00000110090 / ENSRNOG00000014254
CPT1B / Q92523 / ENSMUSG00000078937 / ENSG00000205560 / ENSRNOG00000010438
CPT2 / P23786 / ENSMUSG00000028607 / ENSG00000157184 / ENSRNOG00000012443
CR1 / P17927 / ENSMUSG00000016481 / ENSG00000203710 / ENSRNOG00000008193
CRABP1 / P29762 / ENSMUSG00000032291 / ENSG00000166426 / ENSRNOG00000023633
CRABP2 / P29373 / ENSMUSG00000004885 / ENSG00000143320 / ENSRNOG00000048421
CRACR2B / Q8N4Y2 / ENSMUSG00000048200 / ENSG00000177685 / ENSRNOG00000019199
Homologs
Brain
Heart
Kidney
Liver
Lung
Testis
Organs
Bins
1
2
3
4
5
Species
Human
Mouse
Rat

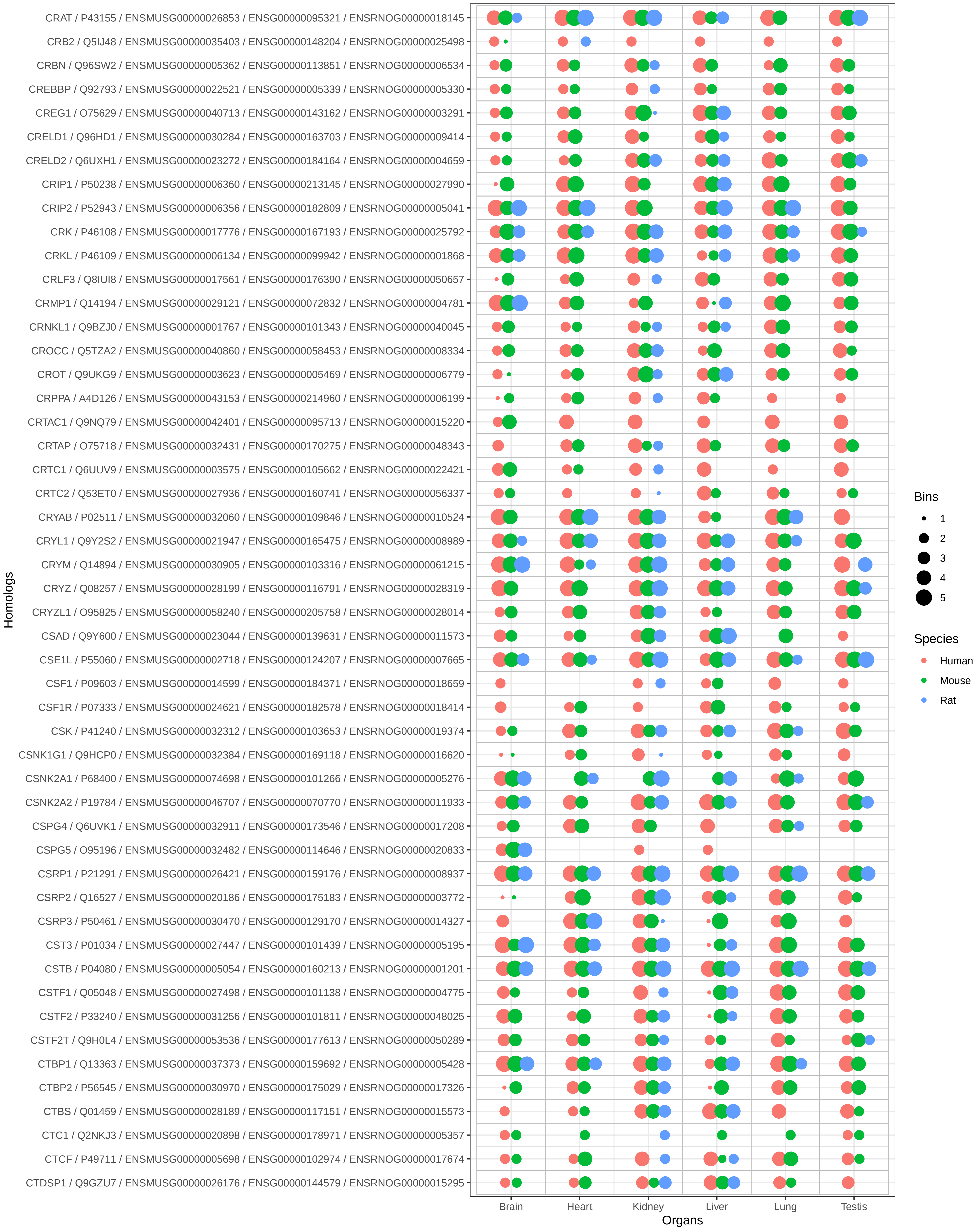
CRAT / P43155 / ENSMUSG00000026853 / ENSG00000095321 / ENSRNOG00000018145
CRB2 / Q5IJ48 / ENSMUSG00000035403 / ENSG00000148204 / ENSRNOG00000025498
CRBN / Q96SW2 / ENSMUSG00000005362 / ENSG00000113851 / ENSRNOG00000006534
CREBBP / Q92793 / ENSMUSG00000022521 / ENSG00000005339 / ENSRNOG00000005330
CREG1 / O75629 / ENSMUSG00000040713 / ENSG00000143162 / ENSRNOG00000003291
CRELD1 / Q96HD1 / ENSMUSG00000030284 / ENSG00000163703 / ENSRNOG00000009414
CRELD2 / Q6UXH1 / ENSMUSG00000023272 / ENSG00000184164 / ENSRNOG00000004659
CRIP1 / P50238 / ENSMUSG00000006360 / ENSG00000213145 / ENSRNOG00000027990
CRIP2 / P52943 / ENSMUSG00000006356 / ENSG00000182809 / ENSRNOG00000005041
CRK / P46108 / ENSMUSG00000017776 / ENSG00000167193 / ENSRNOG00000025792
CRKL / P46109 / ENSMUSG00000006134 / ENSG00000099942 / ENSRNOG00000001868
CRLF3 / Q8IUI8 / ENSMUSG00000017561 / ENSG00000176390 / ENSRNOG00000050657
CRMP1 / Q14194 / ENSMUSG00000029121 / ENSG00000072832 / ENSRNOG00000004781
CRNKL1 / Q9BZJ0 / ENSMUSG00000001767 / ENSG00000101343 / ENSRNOG00000040045
CROCC / Q5TZA2 / ENSMUSG00000040860 / ENSG00000058453 / ENSRNOG00000008334
CROT / Q9UKG9 / ENSMUSG00000003623 / ENSG00000005469 / ENSRNOG00000006779
CRPPA / A4D126 / ENSMUSG00000043153 / ENSG00000214960 / ENSRNOG00000006199
CRTAC1 / Q9NQ79 / ENSMUSG00000042401 / ENSG00000095713 / ENSRNOG00000015220
CRTAP / O75718 / ENSMUSG00000032431 / ENSG00000170275 / ENSRNOG00000048343
CRTC1 / Q6UUV9 / ENSMUSG00000003575 / ENSG00000105662 / ENSRNOG00000022421
CRTC2 / Q53ET0 / ENSMUSG00000027936 / ENSG00000160741 / ENSRNOG00000056337
CRYAB / P02511 / ENSMUSG00000032060 / ENSG00000109846 / ENSRNOG00000010524
CRYL1 / Q9Y2S2 / ENSMUSG00000021947 / ENSG00000165475 / ENSRNOG00000008989
CRYM / Q14894 / ENSMUSG00000030905 / ENSG00000103316 / ENSRNOG00000061215
CRYZ / Q08257 / ENSMUSG00000028199 / ENSG00000116791 / ENSRNOG00000028319
CRYZL1 / O95825 / ENSMUSG00000058240 / ENSG00000205758 / ENSRNOG00000028014
CSAD / Q9Y600 / ENSMUSG00000023044 / ENSG00000139631 / ENSRNOG00000011573
CSE1L / P55060 / ENSMUSG00000002718 / ENSG00000124207 / ENSRNOG00000007665
CSF1 / P09603 / ENSMUSG00000014599 / ENSG00000184371 / ENSRNOG00000018659
CSF1R / P07333 / ENSMUSG00000024621 / ENSG00000182578 / ENSRNOG00000018414
CSK / P41240 / ENSMUSG00000032312 / ENSG00000103653 / ENSRNOG00000019374
CSNK1G1 / Q9HCP0 / ENSMUSG00000032384 / ENSG00000169118 / ENSRNOG00000016620
CSNK2A1 / P68400 / ENSMUSG00000074698 / ENSG00000101266 / ENSRNOG00000005276
CSNK2A2 / P19784 / ENSMUSG00000046707 / ENSG00000070770 / ENSRNOG00000011933
CSPG4 / Q6UVK1 / ENSMUSG00000032911 / ENSG00000173546 / ENSRNOG00000017208
CSPG5 / O95196 / ENSMUSG00000032482 / ENSG00000114646 / ENSRNOG00000020833
CSRP1 / P21291 / ENSMUSG00000026421 / ENSG00000159176 / ENSRNOG00000008937
CSRP2 / Q16527 / ENSMUSG00000020186 / ENSG00000175183 / ENSRNOG00000003772
CSRP3 / P50461 / ENSMUSG00000030470 / ENSG00000129170 / ENSRNOG00000014327
CST3 / P01034 / ENSMUSG00000027447 / ENSG00000101439 / ENSRNOG00000005195
CSTB / P04080 / ENSMUSG00000005054 / ENSG00000160213 / ENSRNOG00000001201
CSTF1 / Q05048 / ENSMUSG00000027498 / ENSG00000101138 / ENSRNOG00000004775
CSTF2 / P33240 / ENSMUSG00000031256 / ENSG00000101811 / ENSRNOG00000048025
CSTF2T / Q9H0L4 / ENSMUSG00000053536 / ENSG00000177613 / ENSRNOG00000050289
CTBP1 / Q13363 / ENSMUSG00000037373 / ENSG00000159692 / ENSRNOG00000005428
CTBP2 / P56545 / ENSMUSG00000030970 / ENSG00000175029 / ENSRNOG00000017326
CTBS / Q01459 / ENSMUSG00000028189 / ENSG00000117151 / ENSRNOG00000015573
CTC1 / Q2NKJ3 / ENSMUSG00000020898 / ENSG00000178971 / ENSRNOG00000005357
CTCF / P49711 / ENSMUSG00000005698 / ENSG00000102974 / ENSRNOG00000017674
CTDSP1 / Q9GZU7 / ENSMUSG00000026176 / ENSG00000144579 / ENSRNOG00000015295
Homologs
Brain
Heart
Kidney
Liver
Lung
Testis
Organs
Bins
1
2
3
4
5
Species
Human
Mouse
Rat

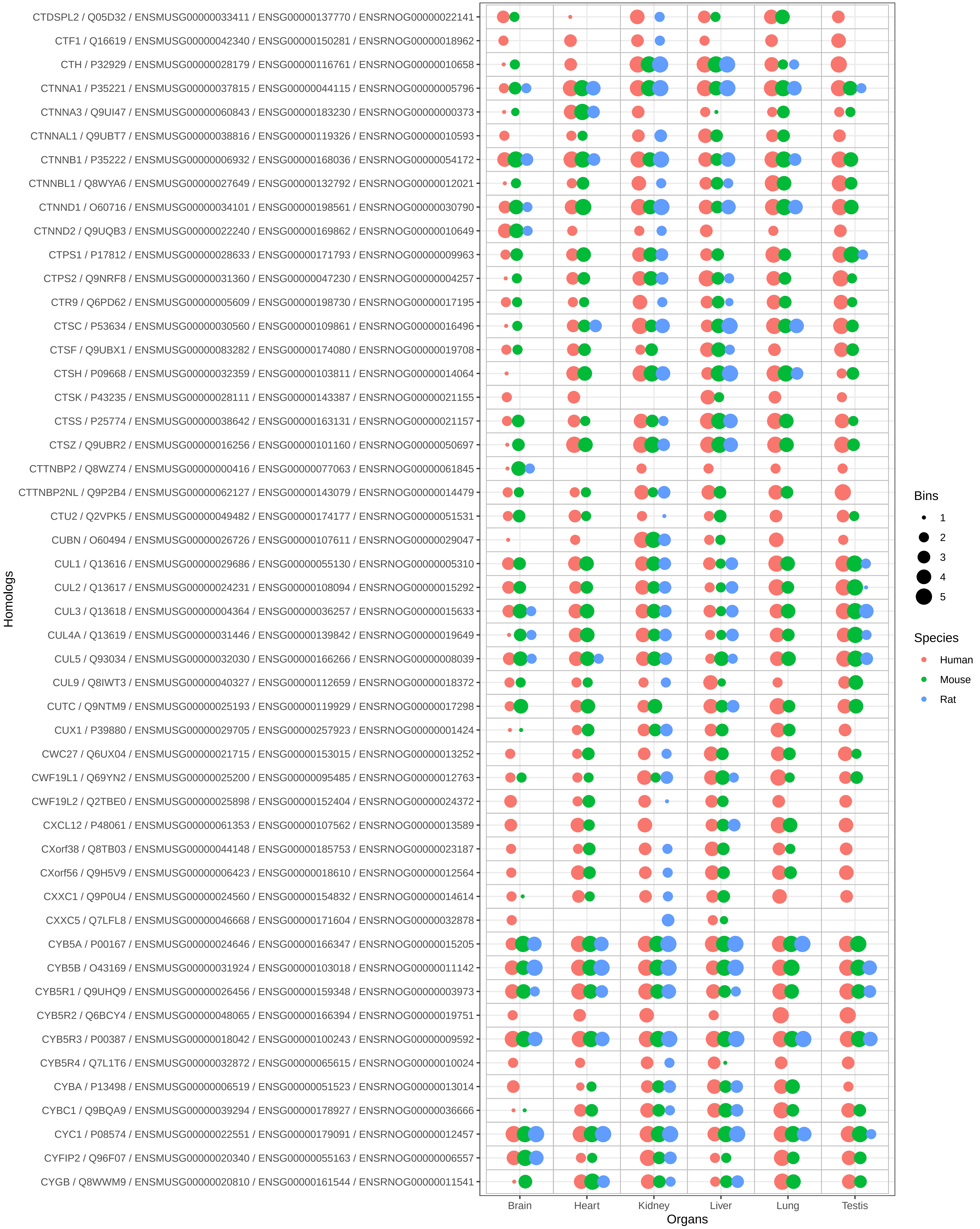
CTDSPL2 / Q05D32 / ENSMUSG00000033411 / ENSG00000137770 / ENSRNOG00000022141
CTF1 / Q16619 / ENSMUSG00000042340 / ENSG00000150281 / ENSRNOG00000018962
CTH / P32929 / ENSMUSG00000028179 / ENSG00000116761 / ENSRNOG00000010658
CTNNA1 / P35221 / ENSMUSG00000037815 / ENSG00000044115 / ENSRNOG00000005796
CTNNA3 / Q9UI47 / ENSMUSG00000060843 / ENSG00000183230 / ENSRNOG00000000373
CTNNAL1 / Q9UBT7 / ENSMUSG00000038816 / ENSG00000119326 / ENSRNOG00000010593
CTNNB1 / P35222 / ENSMUSG00000006932 / ENSG00000168036 / ENSRNOG00000054172
CTNNBL1 / Q8WYA6 / ENSMUSG00000027649 / ENSG00000132792 / ENSRNOG00000012021
CTNND1 / O60716 / ENSMUSG00000034101 / ENSG00000198561 / ENSRNOG00000030790
CTNND2 / Q9UQB3 / ENSMUSG00000022240 / ENSG00000169862 / ENSRNOG00000010649
CTPS1 / P17812 / ENSMUSG00000028633 / ENSG00000171793 / ENSRNOG00000009963
CTPS2 / Q9NRF8 / ENSMUSG00000031360 / ENSG00000047230 / ENSRNOG00000004257
CTR9 / Q6PD62 / ENSMUSG00000005609 / ENSG00000198730 / ENSRNOG00000017195
CTSC / P53634 / ENSMUSG00000030560 / ENSG00000109861 / ENSRNOG00000016496
CTSF / Q9UBX1 / ENSMUSG00000083282 / ENSG00000174080 / ENSRNOG00000019708
CTSH / P09668 / ENSMUSG00000032359 / ENSG00000103811 / ENSRNOG00000014064
CTSK / P43235 / ENSMUSG00000028111 / ENSG00000143387 / ENSRNOG00000021155
CTSS / P25774 / ENSMUSG00000038642 / ENSG00000163131 / ENSRNOG00000021157
CTSZ / Q9UBR2 / ENSMUSG00000016256 / ENSG00000101160 / ENSRNOG00000050697
CTTNBP2 / Q8WZ74 / ENSMUSG00000000416 / ENSG00000077063 / ENSRNOG00000061845
CTTNBP2NL / Q9P2B4 / ENSMUSG00000062127 / ENSG00000143079 / ENSRNOG00000014479
CTU2 / Q2VPK5 / ENSMUSG00000049482 / ENSG00000174177 / ENSRNOG00000051531
CUBN / O60494 / ENSMUSG00000026726 / ENSG00000107611 / ENSRNOG00000029047
CUL1 / Q13616 / ENSMUSG00000029686 / ENSG00000055130 / ENSRNOG00000005310
CUL2 / Q13617 / ENSMUSG00000024231 / ENSG00000108094 / ENSRNOG00000015292
CUL3 / Q13618 / ENSMUSG00000004364 / ENSG00000036257 / ENSRNOG00000015633
CUL4A / Q13619 / ENSMUSG00000031446 / ENSG00000139842 / ENSRNOG00000019649
CUL5 / Q93034 / ENSMUSG00000032030 / ENSG00000166266 / ENSRNOG00000008039
CUL9 / Q8IWT3 / ENSMUSG00000040327 / ENSG00000112659 / ENSRNOG00000018372
CUTC / Q9NTM9 / ENSMUSG00000025193 / ENSG00000119929 / ENSRNOG00000017298
CUX1 / P39880 / ENSMUSG00000029705 / ENSG00000257923 / ENSRNOG00000001424
CWC27 / Q6UX04 / ENSMUSG00000021715 / ENSG00000153015 / ENSRNOG00000013252
CWF19L1 / Q69YN2 / ENSMUSG00000025200 / ENSG00000095485 / ENSRNOG00000012763
CWF19L2 / Q2TBE0 / ENSMUSG00000025898 / ENSG00000152404 / ENSRNOG00000024372
CXCL12 / P48061 / ENSMUSG00000061353 / ENSG00000107562 / ENSRNOG00000013589
CXorf38 / Q8TB03 / ENSMUSG00000044148 / ENSG00000185753 / ENSRNOG00000023187
CXorf56 / Q9H5V9 / ENSMUSG00000006423 / ENSG00000018610 / ENSRNOG00000012564
CXXC1 / Q9P0U4 / ENSMUSG00000024560 / ENSG00000154832 / ENSRNOG00000014614
CXXC5 / Q7LFL8 / ENSMUSG00000046668 / ENSG00000171604 / ENSRNOG00000032878
CYB5A / P00167 / ENSMUSG00000024646 / ENSG00000166347 / ENSRNOG00000015205
CYB5B / O43169 / ENSMUSG00000031924 / ENSG00000103018 / ENSRNOG00000011142
CYB5R1 / Q9UHQ9 / ENSMUSG00000026456 / ENSG00000159348 / ENSRNOG00000003973
CYB5R2 / Q6BCY4 / ENSMUSG00000048065 / ENSG00000166394 / ENSRNOG00000019751
CYB5R3 / P00387 / ENSMUSG00000018042 / ENSG00000100243 / ENSRNOG00000009592
CYB5R4 / Q7L1T6 / ENSMUSG00000032872 / ENSG00000065615 / ENSRNOG00000010024
CYBA / P13498 / ENSMUSG00000006519 / ENSG00000051523 / ENSRNOG00000013014
CYBC1 / Q9BQA9 / ENSMUSG00000039294 / ENSG00000178927 / ENSRNOG00000036666
CYC1 / P08574 / ENSMUSG00000022551 / ENSG00000179091 / ENSRNOG00000012457
CYFIP2 / Q96F07 / ENSMUSG00000020340 / ENSG00000055163 / ENSRNOG00000006557
CYGB / Q8WWM9 / ENSMUSG00000020810 / ENSG00000161544 / ENSRNOG00000011541
Homologs
Brain
Heart
Kidney
Liver
Lung
Testis
Organs
Bins
1
2
3
4
5
Species
Human
Mouse
Rat

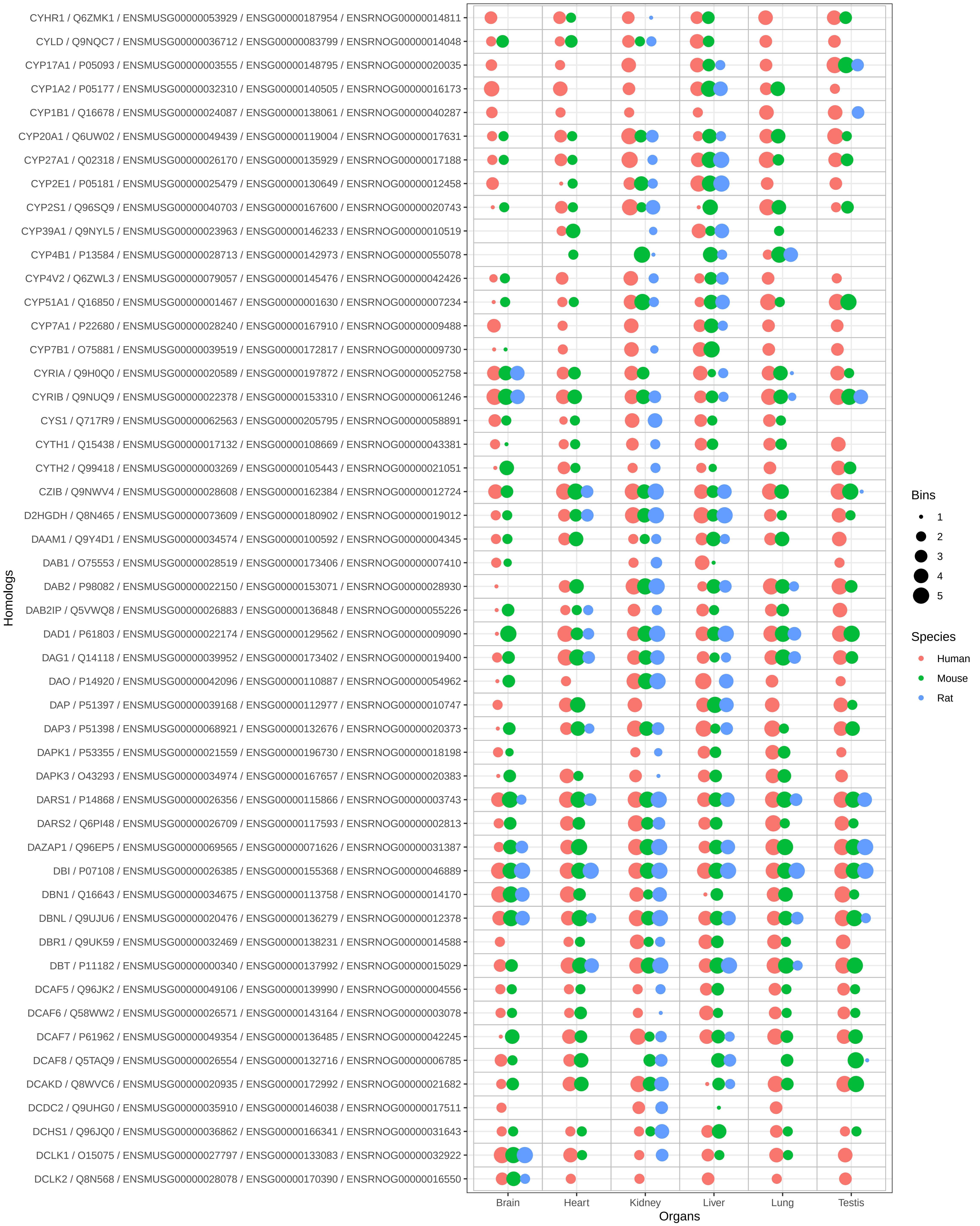

CYHR1 / Q6ZMK1 / ENSMUSG00000053929 / ENSG00000187954 / ENSRNOG00000014811
CYLD / Q9NQC7 / ENSMUSG00000036712 / ENSG00000083799 / ENSRNOG00000014048
CYP17A1 / P05093 / ENSMUSG00000003555 / ENSG00000148795 / ENSRNOG00000020035
CYP1A2 / P05177 / ENSMUSG00000032310 / ENSG00000140505 / ENSRNOG00000016173
CYP1B1 / Q16678 / ENSMUSG00000024087 / ENSG00000138061 / ENSRNOG00000040287
CYP20A1 / Q6UW02 / ENSMUSG00000049439 / ENSG00000119004 / ENSRNOG00000017631
CYP27A1 / Q02318 / ENSMUSG00000026170 / ENSG00000135929 / ENSRNOG00000017188
CYP2E1 / P05181 / ENSMUSG00000025479 / ENSG00000130649 / ENSRNOG00000012458
CYP2S1 / Q96SQ9 / ENSMUSG00000040703 / ENSG00000167600 / ENSRNOG00000020743
CYP39A1 / Q9NYL5 / ENSMUSG00000023963 / ENSG00000146233 / ENSRNOG00000010519
CYP4B1 / P13584 / ENSMUSG00000028713 / ENSG00000142973 / ENSRNOG00000055078
CYP4V2 / Q6ZWL3 / ENSMUSG00000079057 / ENSG00000145476 / ENSRNOG00000042426
CYP51A1 / Q16850 / ENSMUSG00000001467 / ENSG00000001630 / ENSRNOG00000007234
CYP7A1 / P22680 / ENSMUSG00000028240 / ENSG00000167910 / ENSRNOG00000009488
CYP7B1 / O75881 / ENSMUSG00000039519 / ENSG00000172817 / ENSRNOG00000009730
CYRIA / Q9H0Q0 / ENSMUSG00000020589 / ENSG00000197872 / ENSRNOG00000052758
CYRIB / Q9NUQ9 / ENSMUSG00000022378 / ENSG00000153310 / ENSRNOG00000061246
CYS1 / Q717R9 / ENSMUSG00000062563 / ENSG00000205795 / ENSRNOG00000058891
CYTH1 / Q15438 / ENSMUSG00000017132 / ENSG00000108669 / ENSRNOG00000043381
CYTH2 / Q99418 / ENSMUSG00000003269 / ENSG00000105443 / ENSRNOG00000021051
CZIB / Q9NWV4 / ENSMUSG00000028608 / ENSG00000162384 / ENSRNOG00000012724
D2HGDH / Q8N465 / ENSMUSG00000073609 / ENSG00000180902 / ENSRNOG00000019012
DAAM1 / Q9Y4D1 / ENSMUSG00000034574 / ENSG00000100592 / ENSRNOG00000004345
DAB1 / O75553 / ENSMUSG00000028519 / ENSG00000173406 / ENSRNOG00000007410
DAB2 / P98082 / ENSMUSG00000022150 / ENSG00000153071 / ENSRNOG00000028930
DAB2IP / Q5VWQ8 / ENSMUSG00000026883 / ENSG00000136848 / ENSRNOG00000055226
DAD1 / P61803 / ENSMUSG00000022174 / ENSG00000129562 / ENSRNOG00000009090
DAG1 / Q14118 / ENSMUSG00000039952 / ENSG00000173402 / ENSRNOG00000019400
DAO / P14920 / ENSMUSG00000042096 / ENSG00000110887 / ENSRNOG00000054962
DAP / P51397 / ENSMUSG00000039168 / ENSG00000112977 / ENSRNOG00000010747
DAP3 / P51398 / ENSMUSG00000068921 / ENSG00000132676 / ENSRNOG00000020373
DAPK1 / P53355 / ENSMUSG00000021559 / ENSG00000196730 / ENSRNOG00000018198
DAPK3 / O43293 / ENSMUSG00000034974 / ENSG00000167657 / ENSRNOG00000020383
DARS1 / P14868 / ENSMUSG00000026356 / ENSG00000115866 / ENSRNOG00000003743
DARS2 / Q6PI48 / ENSMUSG00000026709 / ENSG00000117593 / ENSRNOG00000002813
DAZAP1 / Q96EP5 / ENSMUSG00000069565 / ENSG00000071626 / ENSRNOG00000031387
DBI / P07108 / ENSMUSG00000026385 / ENSG00000155368 / ENSRNOG00000046889
DBN1 / Q16643 / ENSMUSG00000034675 / ENSG00000113758 / ENSRNOG00000014170
DBNL / Q9UJU6 / ENSMUSG00000020476 / ENSG00000136279 / ENSRNOG00000012378
DBR1 / Q9UK59 / ENSMUSG00000032469 / ENSG00000138231 / ENSRNOG00000014588
DBT / P11182 / ENSMUSG00000000340 / ENSG00000137992 / ENSRNOG00000015029
DCAF5 / Q96JK2 / ENSMUSG00000049106 / ENSG00000139990 / ENSRNOG00000004556
DCAF6 / Q58WW2 / ENSMUSG00000026571 / ENSG00000143164 / ENSRNOG00000003078
DCAF7 / P61962 / ENSMUSG00000049354 / ENSG00000136485 / ENSRNOG00000042245
DCAF8 / Q5TAQ9 / ENSMUSG00000026554 / ENSG00000132716 / ENSRNOG00000006785
DCAKD / Q8WVC6 / ENSMUSG00000020935 / ENSG00000172992 / ENSRNOG00000021682
DCDC2 / Q9UHG0 / ENSMUSG00000035910 / ENSG00000146038 / ENSRNOG00000017511
DCHS1 / Q96JQ0 / ENSMUSG00000036862 / ENSG00000166341 / ENSRNOG00000031643
DCLK1 / O15075 / ENSMUSG00000027797 / ENSG00000133083 / ENSRNOG00000032922
DCLK2 / Q8N568 / ENSMUSG00000028078 / ENSG00000170390 / ENSRNOG00000016550
Homologs
Brain
Heart
Kidney
Liver
Lung
Testis
Organs
Bins
1
2
3
4
5
Species
Human
Mouse
Rat

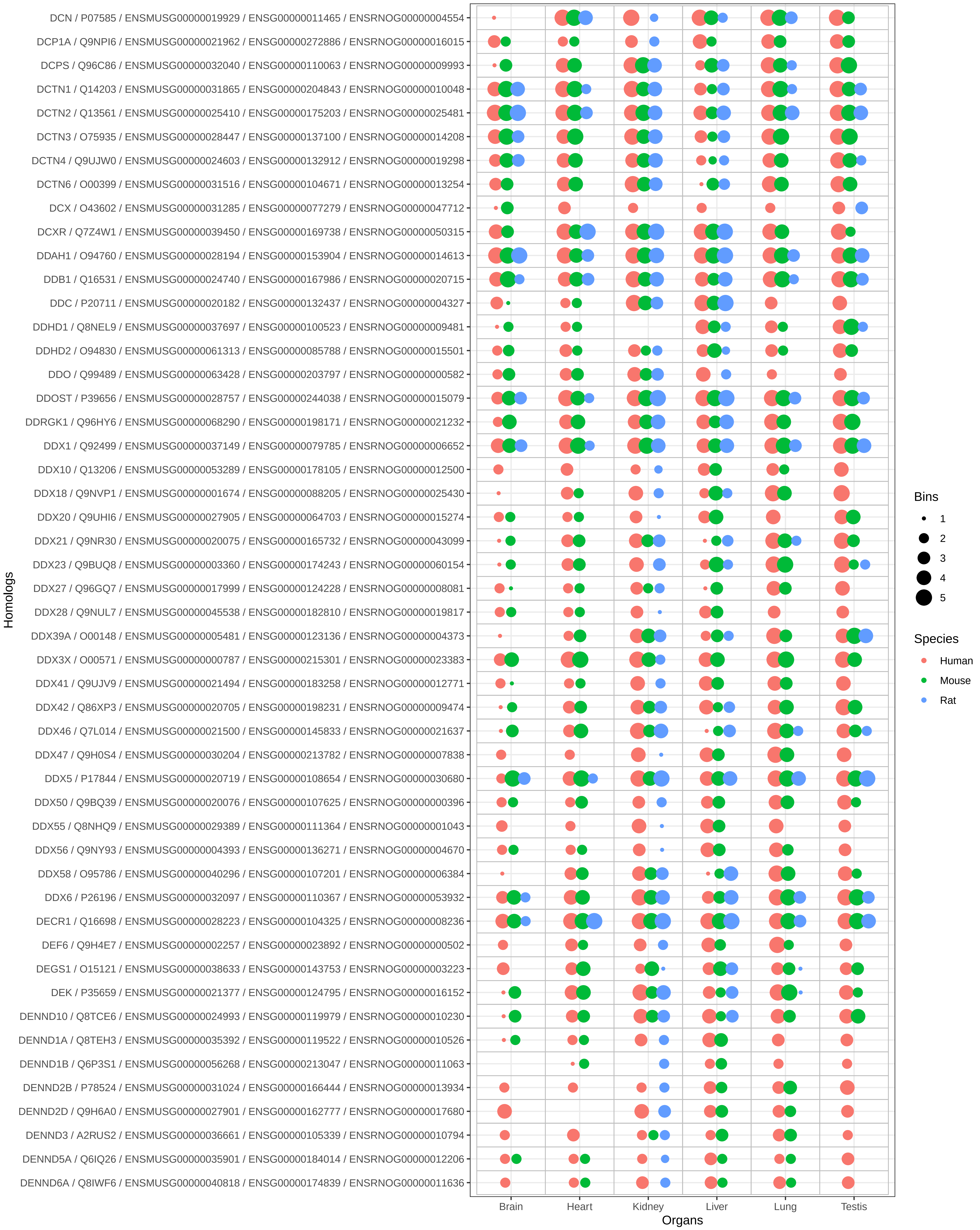

DCN / P07585 / ENSMUSG00000019929 / ENSG00000011465 / ENSRNOG00000004554
DCP1A / Q9NPI6 / ENSMUSG00000021962 / ENSG00000272886 / ENSRNOG00000016015
DCPS / Q96C86 / ENSMUSG00000032040 / ENSG00000110063 / ENSRNOG00000009993
DCTN1 / Q14203 / ENSMUSG00000031865 / ENSG00000204843 / ENSRNOG00000010048
DCTN2 / Q13561 / ENSMUSG00000025410 / ENSG00000175203 / ENSRNOG00000025481
DCTN3 / O75935 / ENSMUSG00000028447 / ENSG00000137100 / ENSRNOG00000014208
DCTN4 / Q9UJW0 / ENSMUSG00000024603 / ENSG00000132912 / ENSRNOG00000019298
DCTN6 / O00399 / ENSMUSG00000031516 / ENSG00000104671 / ENSRNOG00000013254
DCX / O43602 / ENSMUSG00000031285 / ENSG00000077279 / ENSRNOG00000047712
DCXR / Q7Z4W1 / ENSMUSG00000039450 / ENSG00000169738 / ENSRNOG00000050315
DDAH1 / O94760 / ENSMUSG00000028194 / ENSG00000153904 / ENSRNOG00000014613
DDB1 / Q16531 / ENSMUSG00000024740 / ENSG00000167986 / ENSRNOG00000020715
DDC / P20711 / ENSMUSG00000020182 / ENSG00000132437 / ENSRNOG00000004327
DDHD1 / Q8NEL9 / ENSMUSG00000037697 / ENSG00000100523 / ENSRNOG00000009481
DDHD2 / O94830 / ENSMUSG00000061313 / ENSG00000085788 / ENSRNOG00000015501
DDO / Q99489 / ENSMUSG00000063428 / ENSG00000203797 / ENSRNOG00000000582
DDOST / P39656 / ENSMUSG00000028757 / ENSG00000244038 / ENSRNOG00000015079
DDRGK1 / Q96HY6 / ENSMUSG00000068290 / ENSG00000198171 / ENSRNOG00000021232
DDX1 / Q92499 / ENSMUSG00000037149 / ENSG00000079785 / ENSRNOG00000006652
DDX10 / Q13206 / ENSMUSG00000053289 / ENSG00000178105 / ENSRNOG00000012500
DDX18 / Q9NVP1 / ENSMUSG00000001674 / ENSG00000088205 / ENSRNOG00000025430
DDX20 / Q9UHI6 / ENSMUSG00000027905 / ENSG00000064703 / ENSRNOG00000015274
DDX21 / Q9NR30 / ENSMUSG00000020075 / ENSG00000165732 / ENSRNOG00000043099
DDX23 / Q9BUQ8 / ENSMUSG00000003360 / ENSG00000174243 / ENSRNOG00000060154
DDX27 / Q96GQ7 / ENSMUSG00000017999 / ENSG00000124228 / ENSRNOG00000008081
DDX28 / Q9NUL7 / ENSMUSG00000045538 / ENSG00000182810 / ENSRNOG00000019817
DDX39A / O00148 / ENSMUSG00000005481 / ENSG00000123136 / ENSRNOG00000004373
DDX3X / O00571 / ENSMUSG00000000787 / ENSG00000215301 / ENSRNOG00000023383
DDX41 / Q9UJV9 / ENSMUSG00000021494 / ENSG00000183258 / ENSRNOG00000012771
DDX42 / Q86XP3 / ENSMUSG00000020705 / ENSG00000198231 / ENSRNOG00000009474
DDX46 / Q7L014 / ENSMUSG00000021500 / ENSG00000145833 / ENSRNOG00000021637
DDX47 / Q9H0S4 / ENSMUSG00000030204 / ENSG00000213782 / ENSRNOG00000007838
DDX5 / P17844 / ENSMUSG00000020719 / ENSG00000108654 / ENSRNOG00000030680
DDX50 / Q9BQ39 / ENSMUSG00000020076 / ENSG00000107625 / ENSRNOG00000000396
DDX55 / Q8NHQ9 / ENSMUSG00000029389 / ENSG00000111364 / ENSRNOG00000001043
DDX56 / Q9NY93 / ENSMUSG00000004393 / ENSG00000136271 / ENSRNOG00000004670
DDX58 / O95786 / ENSMUSG00000040296 / ENSG00000107201 / ENSRNOG00000006384
DDX6 / P26196 / ENSMUSG00000032097 / ENSG00000110367 / ENSRNOG00000053932
DECR1 / Q16698 / ENSMUSG00000028223 / ENSG00000104325 / ENSRNOG00000008236
DEF6 / Q9H4E7 / ENSMUSG00000002257 / ENSG00000023892 / ENSRNOG00000000502
DEGS1 / O15121 / ENSMUSG00000038633 / ENSG00000143753 / ENSRNOG00000003223
DEK / P35659 / ENSMUSG00000021377 / ENSG00000124795 / ENSRNOG00000016152
DENND10 / Q8TCE6 / ENSMUSG00000024993 / ENSG00000119979 / ENSRNOG00000010230
DENND1A / Q8TEH3 / ENSMUSG00000035392 / ENSG00000119522 / ENSRNOG00000010526
DENND1B / Q6P3S1 / ENSMUSG00000056268 / ENSG00000213047 / ENSRNOG00000011063
DENND2B / P78524 / ENSMUSG00000031024 / ENSG00000166444 / ENSRNOG00000013934
DENND2D / Q9H6A0 / ENSMUSG00000027901 / ENSG00000162777 / ENSRNOG00000017680
DENND3 / A2RUS2 / ENSMUSG00000036661 / ENSG00000105339 / ENSRNOG00000010794
DENND5A / Q6IQ26 / ENSMUSG00000035901 / ENSG00000184014 / ENSRNOG00000012206
DENND6A / Q8IWF6 / ENSMUSG00000040818 / ENSG00000174839 / ENSRNOG00000011636
Homologs
Brain
Heart
Kidney
Liver
Lung
Testis
Organs
Bins
1
2
3
4
5
Species
Human
Mouse
Rat

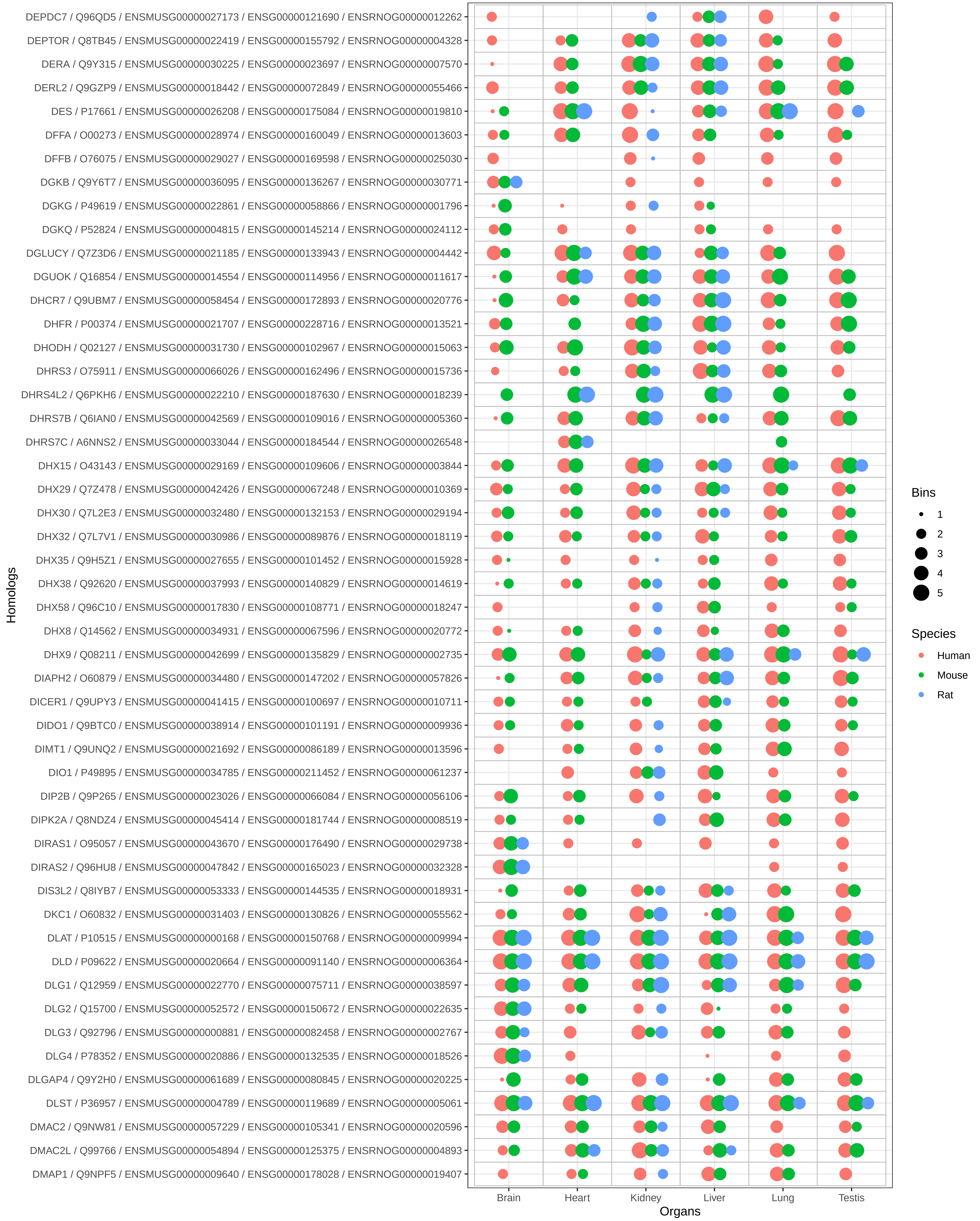
DEPDC7 / Q96QD5 / ENSMUSG00000027173 / ENSG00000121690 / ENSRNOG00000012262
DEPTOR / Q8TB45 / ENSMUSG00000022419 / ENSG00000155792 / ENSRNOG00000004328
DERA / Q9Y315 / ENSMUSG00000030225 / ENSG00000023697 / ENSRNOG00000007570
DERL2 / Q9GZP9 / ENSMUSG00000018442 / ENSG00000072849 / ENSRNOG00000055466
DES / P17661 / ENSMUSG00000026208 / ENSG00000175084 / ENSRNOG00000019810
DFFA / O00273 / ENSMUSG00000028974 / ENSG00000160049 / ENSRNOG00000013603
DFFB / O76075 / ENSMUSG00000029027 / ENSG00000169598 / ENSRNOG00000025030
DGKB / Q9Y6T7 / ENSMUSG00000036095 / ENSG00000136267 / ENSRNOG00000030771
DGKG / P49619 / ENSMUSG00000022861 / ENSG00000058866 / ENSRNOG00000001796
DGKQ / P52824 / ENSMUSG00000004815 / ENSG00000145214 / ENSRNOG00000024112
DGLUCY / Q7Z3D6 / ENSMUSG00000021185 / ENSG00000133943 / ENSRNOG00000004442
DGUOK / Q16854 / ENSMUSG00000014554 / ENSG00000114956 / ENSRNOG00000011617
DHCR7 / Q9UBM7 / ENSMUSG00000058454 / ENSG00000172893 / ENSRNOG00000020776
DHFR / P00374 / ENSMUSG00000021707 / ENSG00000228716 / ENSRNOG00000013521
DHODH / Q02127 / ENSMUSG00000031730 / ENSG00000102967 / ENSRNOG00000015063
DHRS3 / O75911 / ENSMUSG00000066026 / ENSG00000162496 / ENSRNOG00000015736
DHRS4L2 / Q6PKH6 / ENSMUSG00000022210 / ENSG00000187630 / ENSRNOG00000018239
DHRS7B / Q6IAN0 / ENSMUSG00000042569 / ENSG00000109016 / ENSRNOG00000005360
DHRS7C / A6NNS2 / ENSMUSG00000033044 / ENSG00000184544 / ENSRNOG00000026548
DHX15 / O43143 / ENSMUSG00000029169 / ENSG00000109606 / ENSRNOG00000003844
DHX29 / Q7Z478 / ENSMUSG00000042426 / ENSG00000067248 / ENSRNOG00000010369
DHX30 / Q7L2E3 / ENSMUSG00000032480 / ENSG00000132153 / ENSRNOG00000029194
DHX32 / Q7L7V1 / ENSMUSG00000030986 / ENSG00000089876 / ENSRNOG00000018119
DHX35 / Q9H5Z1 / ENSMUSG00000027655 / ENSG00000101452 / ENSRNOG00000015928
DHX38 / Q92620 / ENSMUSG00000037993 / ENSG00000140829 / ENSRNOG00000014619
DHX58 / Q96C10 / ENSMUSG00000017830 / ENSG00000108771 / ENSRNOG00000018247
DHX8 / Q14562 / ENSMUSG00000034931 / ENSG00000067596 / ENSRNOG00000020772
DHX9 / Q08211 / ENSMUSG00000042699 / ENSG00000135829 / ENSRNOG00000002735
DIAPH2 / O60879 / ENSMUSG00000034480 / ENSG00000147202 / ENSRNOG00000057826
DICER1 / Q9UPY3 / ENSMUSG00000041415 / ENSG00000100697 / ENSRNOG00000010711
DIDO1 / Q9BTC0 / ENSMUSG00000038914 / ENSG00000101191 / ENSRNOG00000009936
DIMT1 / Q9UNQ2 / ENSMUSG00000021692 / ENSG00000086189 / ENSRNOG00000013596
DIO1 / P49895 / ENSMUSG00000034785 / ENSG00000211452 / ENSRNOG00000061237
DIP2B / Q9P265 / ENSMUSG00000023026 / ENSG00000066084 / ENSRNOG00000056106
DIPK2A / Q8NDZ4 / ENSMUSG00000045414 / ENSG00000181744 / ENSRNOG00000008519
DIRAS1 / O95057 / ENSMUSG00000043670 / ENSG00000176490 / ENSRNOG00000029738
DIRAS2 / Q96HU8 / ENSMUSG00000047842 / ENSG00000165023 / ENSRNOG00000032328
DIS3L2 / Q8IYB7 / ENSMUSG00000053333 / ENSG00000144535 / ENSRNOG00000018931
DKC1 / O60832 / ENSMUSG00000031403 / ENSG00000130826 / ENSRNOG00000055562
DLAT / P10515 / ENSMUSG00000000168 / ENSG00000150768 / ENSRNOG00000009994
DLD / P09622 / ENSMUSG00000020664 / ENSG00000091140 / ENSRNOG00000006364
DLG1 / Q12959 / ENSMUSG00000022770 / ENSG00000075711 / ENSRNOG00000038597
DLG2 / Q15700 / ENSMUSG00000052572 / ENSG00000150672 / ENSRNOG00000022635
DLG3 / Q92796 / ENSMUSG00000000881 / ENSG00000082458 / ENSRNOG00000002767
DLG4 / P78352 / ENSMUSG00000020886 / ENSG00000132535 / ENSRNOG00000018526
DLGAP4 / Q9Y2H0 / ENSMUSG00000061689 / ENSG00000080845 / ENSRNOG00000020225
DLST / P36957 / ENSMUSG00000004789 / ENSG00000119689 / ENSRNOG00000005061
DMAC2 / Q9NW81 / ENSMUSG00000057229 / ENSG00000105341 / ENSRNOG00000020596
DMAC2L / Q99766 / ENSMUSG00000054894 / ENSG00000125375 / ENSRNOG00000004893
DMAP1 / Q9NPF5 / ENSMUSG00000009640 / ENSG00000178028 / ENSRNOG00000019407
Homologs
Brain
Heart
Kidney
Liver
Lung
Testis
Organs
Bins
1
2
3
4
5
Species
Human
Mouse
Rat

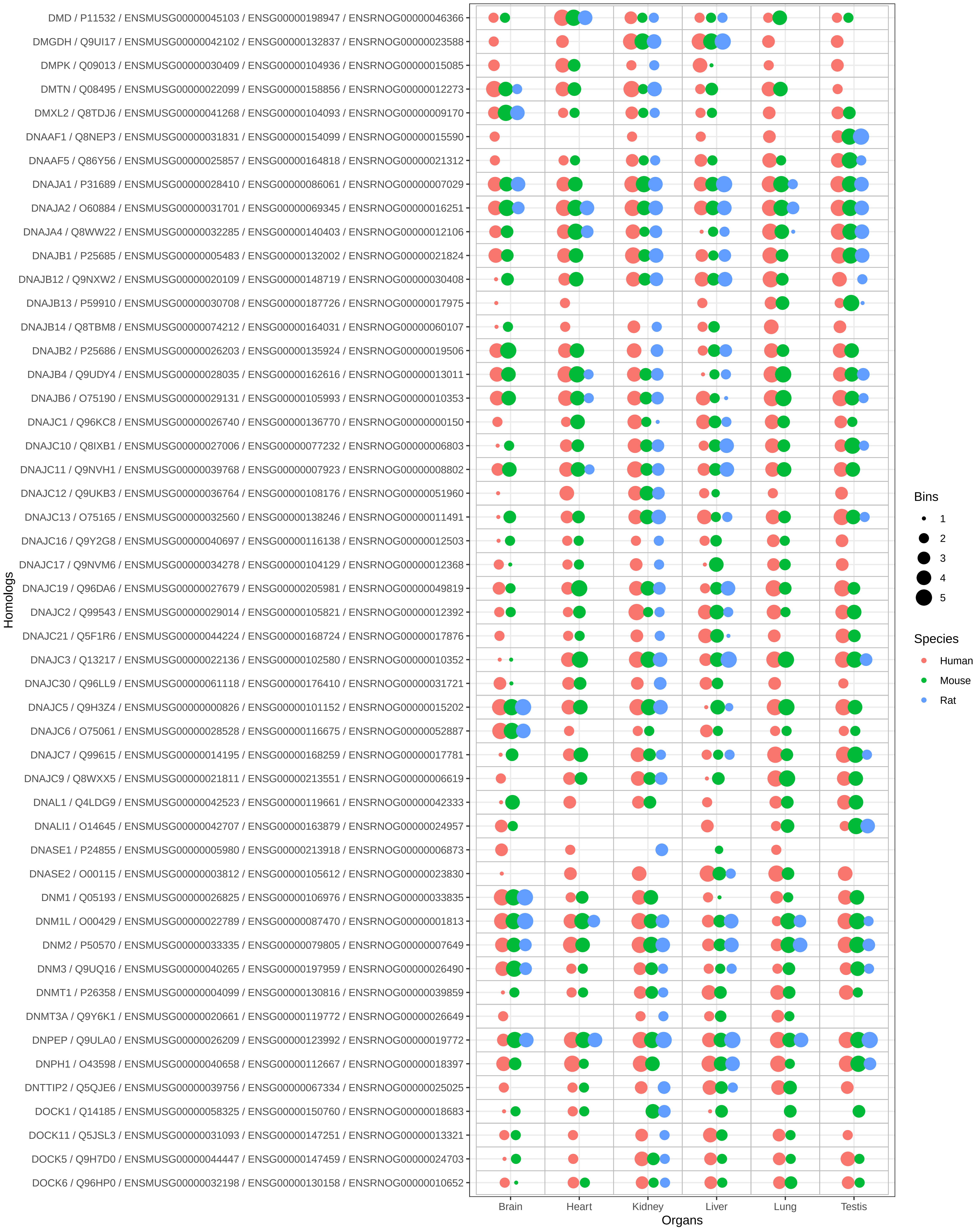
DMD / P11532 / ENSMUSG00000045103 / ENSG00000198947 / ENSRNOG00000046366
DMGDH / Q9UI17 / ENSMUSG00000042102 / ENSG00000132837 / ENSRNOG00000023588
DMPK / Q09013 / ENSMUSG00000030409 / ENSG00000104936 / ENSRNOG00000015085
DMTN / Q08495 / ENSMUSG00000022099 / ENSG00000158856 / ENSRNOG00000012273
DMXL2 / Q8TDJ6 / ENSMUSG00000041268 / ENSG00000104093 / ENSRNOG00000009170
DNAAF1 / Q8NEP3 / ENSMUSG00000031831 / ENSG00000154099 / ENSRNOG00000015590
DNAAF5 / Q86Y56 / ENSMUSG00000025857 / ENSG00000164818 / ENSRNOG00000021312
DNAJA1 / P31689 / ENSMUSG00000028410 / ENSG00000086061 / ENSRNOG00000007029
DNAJA2 / O60884 / ENSMUSG00000031701 / ENSG00000069345 / ENSRNOG00000016251
DNAJA4 / Q8WW22 / ENSMUSG00000032285 / ENSG00000140403 / ENSRNOG00000012106
DNAJB1 / P25685 / ENSMUSG00000005483 / ENSG00000132002 / ENSRNOG00000021824
DNAJB12 / Q9NXW2 / ENSMUSG00000020109 / ENSG00000148719 / ENSRNOG00000030408
DNAJB13 / P59910 / ENSMUSG00000030708 / ENSG00000187726 / ENSRNOG00000017975
DNAJB14 / Q8TBM8 / ENSMUSG00000074212 / ENSG00000164031 / ENSRNOG00000060107
DNAJB2 / P25686 / ENSMUSG00000026203 / ENSG00000135924 / ENSRNOG00000019506
DNAJB4 / Q9UDY4 / ENSMUSG00000028035 / ENSG00000162616 / ENSRNOG00000013011
DNAJB6 / O75190 / ENSMUSG00000029131 / ENSG00000105993 / ENSRNOG00000010353
DNAJC1 / Q96KC8 / ENSMUSG00000026740 / ENSG00000136770 / ENSRNOG00000000150
DNAJC10 / Q8IXB1 / ENSMUSG00000027006 / ENSG00000077232 / ENSRNOG00000006803
DNAJC11 / Q9NVH1 / ENSMUSG00000039768 / ENSG00000007923 / ENSRNOG00000008802
DNAJC12 / Q9UKB3 / ENSMUSG00000036764 / ENSG00000108176 / ENSRNOG00000051960
DNAJC13 / O75165 / ENSMUSG00000032560 / ENSG00000138246 / ENSRNOG00000011491
DNAJC16 / Q9Y2G8 / ENSMUSG00000040697 / ENSG00000116138 / ENSRNOG00000012503
DNAJC17 / Q9NVM6 / ENSMUSG00000034278 / ENSG00000104129 / ENSRNOG00000012368
DNAJC19 / Q96DA6 / ENSMUSG00000027679 / ENSG00000205981 / ENSRNOG00000049819
DNAJC2 / Q99543 / ENSMUSG00000029014 / ENSG00000105821 / ENSRNOG00000012392
DNAJC21 / Q5F1R6 / ENSMUSG00000044224 / ENSG00000168724 / ENSRNOG00000017876
DNAJC3 / Q13217 / ENSMUSG00000022136 / ENSG00000102580 / ENSRNOG00000010352
DNAJC30 / Q96LL9 / ENSMUSG00000061118 / ENSG00000176410 / ENSRNOG00000031721
DNAJC5 / Q9H3Z4 / ENSMUSG00000000826 / ENSG00000101152 / ENSRNOG00000015202
DNAJC6 / O75061 / ENSMUSG00000028528 / ENSG00000116675 / ENSRNOG00000052887
DNAJC7 / Q99615 / ENSMUSG00000014195 / ENSG00000168259 / ENSRNOG00000017781
DNAJC9 / Q8WXX5 / ENSMUSG00000021811 / ENSG00000213551 / ENSRNOG00000006619
DNAL1 / Q4LDG9 / ENSMUSG00000042523 / ENSG00000119661 / ENSRNOG00000042333
DNALI1 / O14645 / ENSMUSG00000042707 / ENSG00000163879 / ENSRNOG00000024957
DNASE1 / P24855 / ENSMUSG00000005980 / ENSG00000213918 / ENSRNOG00000006873
DNASE2 / O00115 / ENSMUSG00000003812 / ENSG00000105612 / ENSRNOG00000023830
DNM1 / Q05193 / ENSMUSG00000026825 / ENSG00000106976 / ENSRNOG00000033835
DNM1L / O00429 / ENSMUSG00000022789 / ENSG00000087470 / ENSRNOG00000001813
DNM2 / P50570 / ENSMUSG00000033335 / ENSG00000079805 / ENSRNOG00000007649
DNM3 / Q9UQ16 / ENSMUSG00000040265 / ENSG00000197959 / ENSRNOG00000026490
DNMT1 / P26358 / ENSMUSG00000004099 / ENSG00000130816 / ENSRNOG00000039859
DNMT3A / Q9Y6K1 / ENSMUSG00000020661 / ENSG00000119772 / ENSRNOG00000026649
DNPEP / Q9ULA0 / ENSMUSG00000026209 / ENSG00000123992 / ENSRNOG00000019772
DNPH1 / O43598 / ENSMUSG00000040658 / ENSG00000112667 / ENSRNOG00000018397
DNTTIP2 / Q5QJE6 / ENSMUSG00000039756 / ENSG00000067334 / ENSRNOG00000025025
DOCK1 / Q14185 / ENSMUSG00000058325 / ENSG00000150760 / ENSRNOG00000018683
DOCK11 / Q5JSL3 / ENSMUSG00000031093 / ENSG00000147251 / ENSRNOG00000013321
DOCK5 / Q9H7D0 / ENSMUSG00000044447 / ENSG00000147459 / ENSRNOG00000024703
DOCK6 / Q96HP0 / ENSMUSG00000032198 / ENSG00000130158 / ENSRNOG00000010652
Homologs
Brain
Heart
Kidney
Liver
Lung
Testis
Organs
Bins
1
2
3
4
5
Species
Human
Mouse
Rat

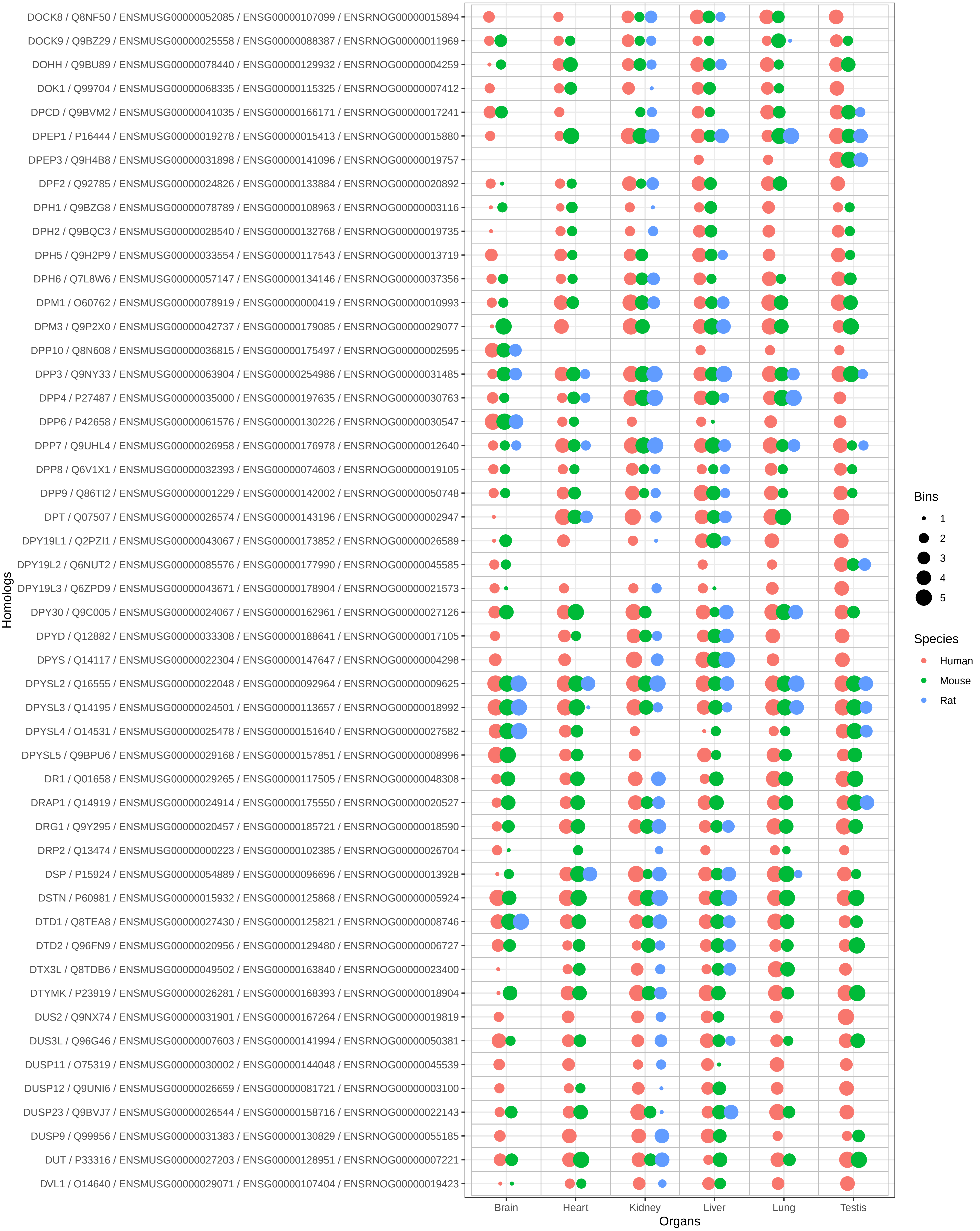
DOCK8 / Q8NF50 / ENSMUSG00000052085 / ENSG00000107099 / ENSRNOG00000015894
DOCK9 / Q9BZ29 / ENSMUSG00000025558 / ENSG00000088387 / ENSRNOG00000011969
DOHH / Q9BU89 / ENSMUSG00000078440 / ENSG00000129932 / ENSRNOG00000004259
DOK1 / Q99704 / ENSMUSG00000068335 / ENSG00000115325 / ENSRNOG00000007412
DPCD / Q9BVM2 / ENSMUSG00000041035 / ENSG00000166171 / ENSRNOG00000017241
DPEP1 / P16444 / ENSMUSG00000019278 / ENSG00000015413 / ENSRNOG00000015880
DPEP3 / Q9H4B8 / ENSMUSG00000031898 / ENSG00000141096 / ENSRNOG00000019757
DPF2 / Q92785 / ENSMUSG00000024826 / ENSG00000133884 / ENSRNOG00000020892
DPH1 / Q9BZG8 / ENSMUSG00000078789 / ENSG00000108963 / ENSRNOG00000003116
DPH2 / Q9BQC3 / ENSMUSG00000028540 / ENSG00000132768 / ENSRNOG00000019735
DPH5 / Q9H2P9 / ENSMUSG00000033554 / ENSG00000117543 / ENSRNOG00000013719
DPH6 / Q7L8W6 / ENSMUSG00000057147 / ENSG00000134146 / ENSRNOG00000037356
DPM1 / O60762 / ENSMUSG00000078919 / ENSG00000000419 / ENSRNOG00000010993
DPM3 / Q9P2X0 / ENSMUSG00000042737 / ENSG00000179085 / ENSRNOG00000029077
DPP10 / Q8N608 / ENSMUSG00000036815 / ENSG00000175497 / ENSRNOG00000002595
DPP3 / Q9NY33 / ENSMUSG00000063904 / ENSG00000254986 / ENSRNOG00000031485
DPP4 / P27487 / ENSMUSG00000035000 / ENSG00000197635 / ENSRNOG00000030763
DPP6 / P42658 / ENSMUSG00000061576 / ENSG00000130226 / ENSRNOG00000030547
DPP7 / Q9UHL4 / ENSMUSG00000026958 / ENSG00000176978 / ENSRNOG00000012640
DPP8 / Q6V1X1 / ENSMUSG00000032393 / ENSG00000074603 / ENSRNOG00000019105
DPP9 / Q86TI2 / ENSMUSG00000001229 / ENSG00000142002 / ENSRNOG00000050748
DPT / Q07507 / ENSMUSG00000026574 / ENSG00000143196 / ENSRNOG00000002947
DPY19L1 / Q2PZI1 / ENSMUSG00000043067 / ENSG00000173852 / ENSRNOG00000026589
DPY19L2 / Q6NUT2 / ENSMUSG00000085576 / ENSG00000177990 / ENSRNOG00000045585
DPY19L3 / Q6ZPD9 / ENSMUSG00000043671 / ENSG00000178904 / ENSRNOG00000021573
DPY30 / Q9C005 / ENSMUSG00000024067 / ENSG00000162961 / ENSRNOG00000027126
DPYD / Q12882 / ENSMUSG00000033308 / ENSG00000188641 / ENSRNOG00000017105
DPYS / Q14117 / ENSMUSG00000022304 / ENSG00000147647 / ENSRNOG00000004298
DPYSL2 / Q16555 / ENSMUSG00000022048 / ENSG00000092964 / ENSRNOG00000009625
DPYSL3 / Q14195 / ENSMUSG00000024501 / ENSG00000113657 / ENSRNOG00000018992
DPYSL4 / O14531 / ENSMUSG00000025478 / ENSG00000151640 / ENSRNOG00000027582
DPYSL5 / Q9BPU6 / ENSMUSG00000029168 / ENSG00000157851 / ENSRNOG00000008996
DR1 / Q01658 / ENSMUSG00000029265 / ENSG00000117505 / ENSRNOG00000048308
DRAP1 / Q14919 / ENSMUSG00000024914 / ENSG00000175550 / ENSRNOG00000020527
DRG1 / Q9Y295 / ENSMUSG00000020457 / ENSG00000185721 / ENSRNOG00000018590
DRP2 / Q13474 / ENSMUSG00000000223 / ENSG00000102385 / ENSRNOG00000026704
DSP / P15924 / ENSMUSG00000054889 / ENSG00000096696 / ENSRNOG00000013928
DSTN / P60981 / ENSMUSG00000015932 / ENSG00000125868 / ENSRNOG00000005924
DTD1 / Q8TEA8 / ENSMUSG00000027430 / ENSG00000125821 / ENSRNOG00000008746
DTD2 / Q96FN9 / ENSMUSG00000020956 / ENSG00000129480 / ENSRNOG00000006727
DTX3L / Q8TDB6 / ENSMUSG00000049502 / ENSG00000163840 / ENSRNOG00000023400
DTYMK / P23919 / ENSMUSG00000026281 / ENSG00000168393 / ENSRNOG00000018904
DUS2 / Q9NX74 / ENSMUSG00000031901 / ENSG00000167264 / ENSRNOG00000019819
DUS3L / Q96G46 / ENSMUSG00000007603 / ENSG00000141994 / ENSRNOG00000050381
DUSP11 / O75319 / ENSMUSG00000030002 / ENSG00000144048 / ENSRNOG00000045539
DUSP12 / Q9UNI6 / ENSMUSG00000026659 / ENSG00000081721 / ENSRNOG00000003100
DUSP23 / Q9BVJ7 / ENSMUSG00000026544 / ENSG00000158716 / ENSRNOG00000022143
DUSP9 / Q99956 / ENSMUSG00000031383 / ENSG00000130829 / ENSRNOG00000055185
DUT / P33316 / ENSMUSG00000027203 / ENSG00000128951 / ENSRNOG00000007221
DVL1 / O14640 / ENSMUSG00000029071 / ENSG00000107404 / ENSRNOG00000019423
Homologs
Brain
Heart
Kidney
Liver
Lung
Testis
Organs
Bins
1
2
3
4
5
Species
Human
Mouse
Rat

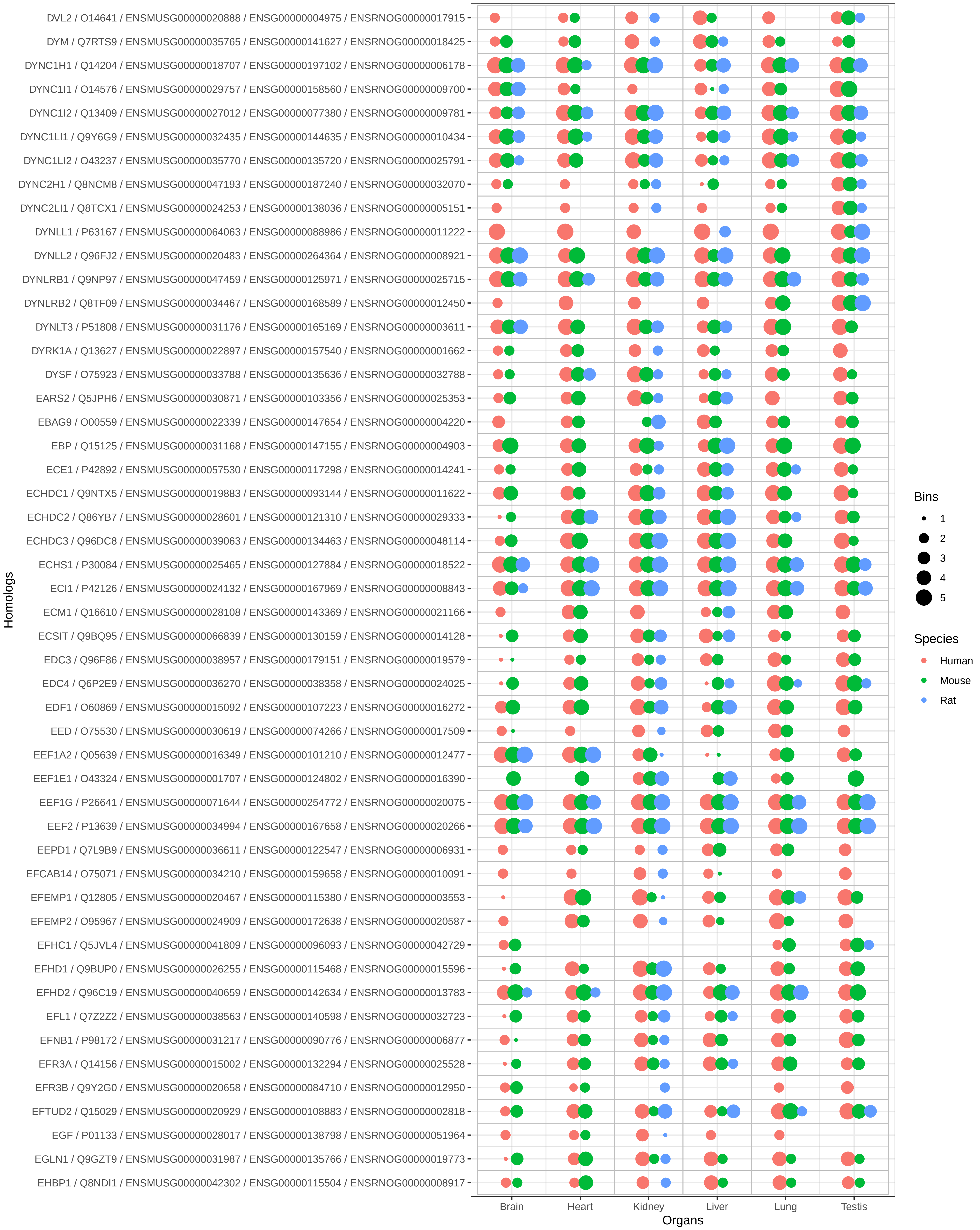

DVL2 / O14641 / ENSMUSG00000020888 / ENSG00000004975 / ENSRNOG00000017915
DYM / Q7RTS9 / ENSMUSG00000035765 / ENSG00000141627 / ENSRNOG00000018425
DYNC1H1 / Q14204 / ENSMUSG00000018707 / ENSG00000197102 / ENSRNOG00000006178
DYNC1I1 / O14576 / ENSMUSG00000029757 / ENSG00000158560 / ENSRNOG00000009700
DYNC1I2 / Q13409 / ENSMUSG00000027012 / ENSG00000077380 / ENSRNOG00000009781
DYNC1LI1 / Q9Y6G9 / ENSMUSG00000032435 / ENSG00000144635 / ENSRNOG00000010434
DYNC1LI2 / O43237 / ENSMUSG00000035770 / ENSG00000135720 / ENSRNOG00000025791
DYNC2H1 / Q8NCM8 / ENSMUSG00000047193 / ENSG00000187240 / ENSRNOG00000032070
DYNC2LI1 / Q8TCX1 / ENSMUSG00000024253 / ENSG00000138036 / ENSRNOG00000005151
DYNLL1 / P63167 / ENSMUSG00000064063 / ENSG00000088986 / ENSRNOG00000011222
DYNLL2 / Q96FJ2 / ENSMUSG00000020483 / ENSG00000264364 / ENSRNOG00000008921
DYNLRB1 / Q9NP97 / ENSMUSG00000047459 / ENSG00000125971 / ENSRNOG00000025715
DYNLRB2 / Q8TF09 / ENSMUSG00000034467 / ENSG00000168589 / ENSRNOG00000012450
DYNLT3 / P51808 / ENSMUSG00000031176 / ENSG00000165169 / ENSRNOG00000003611
DYRK1A / Q13627 / ENSMUSG00000022897 / ENSG00000157540 / ENSRNOG00000001662
DYSF / O75923 / ENSMUSG00000033788 / ENSG00000135636 / ENSRNOG00000032788
EARS2 / Q5JPH6 / ENSMUSG00000030871 / ENSG00000103356 / ENSRNOG00000025353
EBAG9 / O00559 / ENSMUSG00000022339 / ENSG00000147654 / ENSRNOG00000004220
EBP / Q15125 / ENSMUSG00000031168 / ENSG00000147155 / ENSRNOG00000004903
ECE1 / P42892 / ENSMUSG00000057530 / ENSG00000117298 / ENSRNOG00000014241
ECHDC1 / Q9NTX5 / ENSMUSG00000019883 / ENSG00000093144 / ENSRNOG00000011622
ECHDC2 / Q86YB7 / ENSMUSG00000028601 / ENSG00000121310 / ENSRNOG00000029333
ECHDC3 / Q96DC8 / ENSMUSG00000039063 / ENSG00000134463 / ENSRNOG00000048114
ECHS1 / P30084 / ENSMUSG00000025465 / ENSG00000127884 / ENSRNOG00000018522
ECI1 / P42126 / ENSMUSG00000024132 / ENSG00000167969 / ENSRNOG00000008843
ECM1 / Q16610 / ENSMUSG00000028108 / ENSG00000143369 / ENSRNOG00000021166
ECSIT / Q9BQ95 / ENSMUSG00000066839 / ENSG00000130159 / ENSRNOG00000014128
EDC3 / Q96F86 / ENSMUSG00000038957 / ENSG00000179151 / ENSRNOG00000019579
EDC4 / Q6P2E9 / ENSMUSG00000036270 / ENSG00000038358 / ENSRNOG00000024025
EDF1 / O60869 / ENSMUSG00000015092 / ENSG00000107223 / ENSRNOG00000016272
EED / O75530 / ENSMUSG00000030619 / ENSG00000074266 / ENSRNOG00000017509
EEF1A2 / Q05639 / ENSMUSG00000016349 / ENSG00000101210 / ENSRNOG00000012477
EEF1E1 / O43324 / ENSMUSG00000001707 / ENSG00000124802 / ENSRNOG00000016390
EEF1G / P26641 / ENSMUSG00000071644 / ENSG00000254772 / ENSRNOG00000020075
EEF2 / P13639 / ENSMUSG00000034994 / ENSG00000167658 / ENSRNOG00000020266
EEPD1 / Q7L9B9 / ENSMUSG00000036611 / ENSG00000122547 / ENSRNOG00000006931
EFCAB14 / O75071 / ENSMUSG00000034210 / ENSG00000159658 / ENSRNOG00000010091
EFEMP1 / Q12805 / ENSMUSG00000020467 / ENSG00000115380 / ENSRNOG00000003553
EFEMP2 / O95967 / ENSMUSG00000024909 / ENSG00000172638 / ENSRNOG00000020587
EFHC1 / Q5JVL4 / ENSMUSG00000041809 / ENSG00000096093 / ENSRNOG00000042729
EFHD1 / Q9BUP0 / ENSMUSG00000026255 / ENSG00000115468 / ENSRNOG00000015596
EFHD2 / Q96C19 / ENSMUSG00000040659 / ENSG00000142634 / ENSRNOG00000013783
EFL1 / Q7Z2Z2 / ENSMUSG00000038563 / ENSG00000140598 / ENSRNOG00000032723
EFNB1 / P98172 / ENSMUSG00000031217 / ENSG00000090776 / ENSRNOG00000006877
EFR3A / Q14156 / ENSMUSG00000015002 / ENSG00000132294 / ENSRNOG00000025528
EFR3B / Q9Y2G0 / ENSMUSG00000020658 / ENSG00000084710 / ENSRNOG00000012950
EFTUD2 / Q15029 / ENSMUSG00000020929 / ENSG00000108883 / ENSRNOG00000002818
EGF / P01133 / ENSMUSG00000028017 / ENSG00000138798 / ENSRNOG00000051964
EGLN1 / Q9GZT9 / ENSMUSG00000031987 / ENSG00000135766 / ENSRNOG00000019773
EHBP1 / Q8NDI1 / ENSMUSG00000042302 / ENSG00000115504 / ENSRNOG00000008917
Homologs
Brain
Heart
Kidney
Liver
Lung
Testis
Organs
Bins
1
2
3
4
5
Species
Human
Mouse
Rat

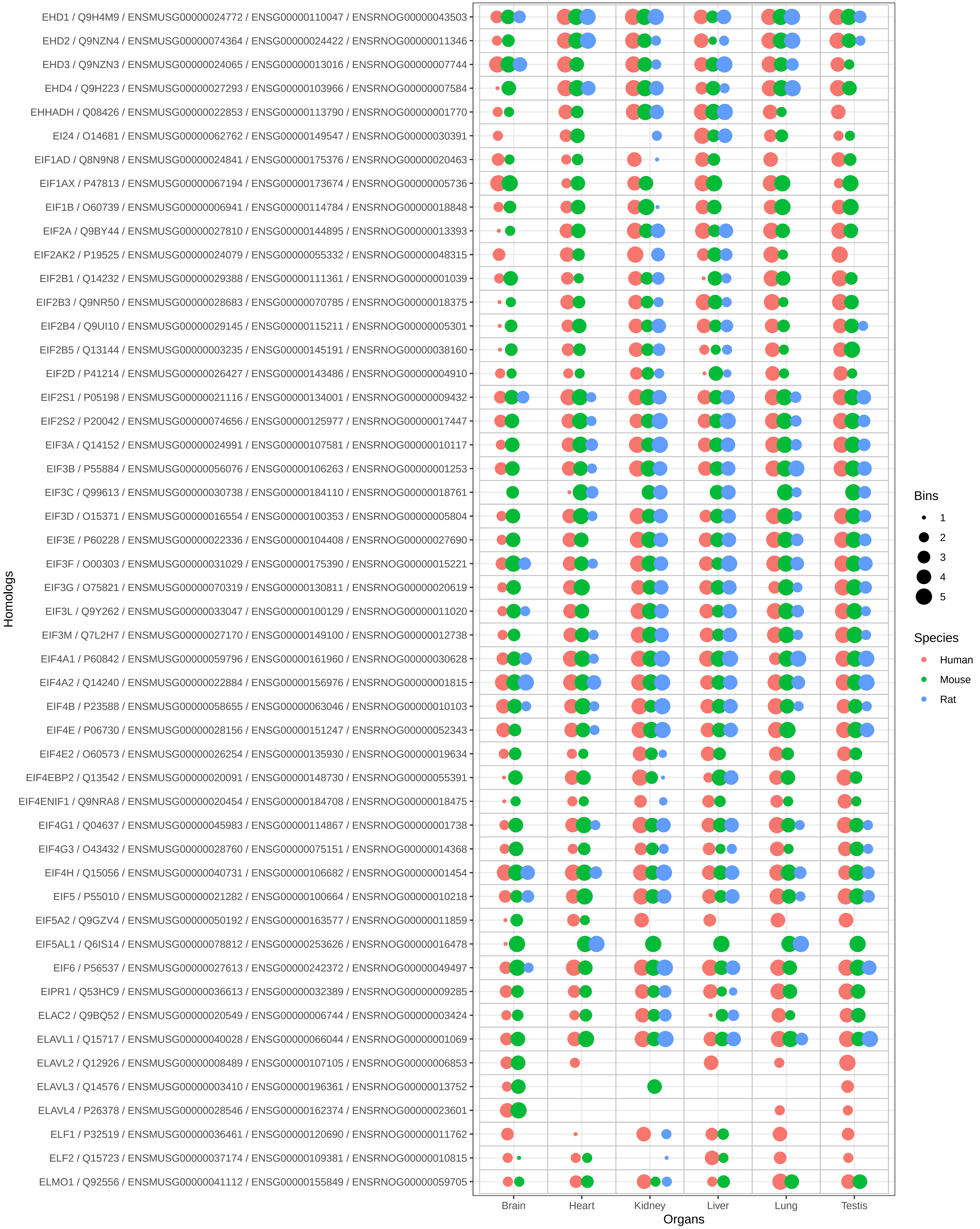
EHD1 / Q9H4M9 / ENSMUSG00000024772 / ENSG00000110047 / ENSRNOG00000043503
EHD2 / Q9NZN4 / ENSMUSG00000074364 / ENSG00000024422 / ENSRNOG00000011346
EHD3 / Q9NZN3 / ENSMUSG00000024065 / ENSG00000013016 / ENSRNOG00000007744
EHD4 / Q9H223 / ENSMUSG00000027293 / ENSG00000103966 / ENSRNOG00000007584
EHHADH / Q08426 / ENSMUSG00000022853 / ENSG00000113790 / ENSRNOG00000001770
EI24 / O14681 / ENSMUSG00000062762 / ENSG00000149547 / ENSRNOG00000030391
EIF1AD / Q8N9N8 / ENSMUSG00000024841 / ENSG00000175376 / ENSRNOG00000020463
EIF1AX / P47813 / ENSMUSG00000067194 / ENSG00000173674 / ENSRNOG00000005736
EIF1B / O60739 / ENSMUSG00000006941 / ENSG00000114784 / ENSRNOG00000018848
EIF2A / Q9BY44 / ENSMUSG00000027810 / ENSG00000144895 / ENSRNOG00000013393
EIF2AK2 / P19525 / ENSMUSG00000024079 / ENSG00000055332 / ENSRNOG00000048315
EIF2B1 / Q14232 / ENSMUSG00000029388 / ENSG00000111361 / ENSRNOG00000001039
EIF2B3 / Q9NR50 / ENSMUSG00000028683 / ENSG00000070785 / ENSRNOG00000018375
EIF2B4 / Q9UI10 / ENSMUSG00000029145 / ENSG00000115211 / ENSRNOG00000005301
EIF2B5 / Q13144 / ENSMUSG00000003235 / ENSG00000145191 / ENSRNOG00000038160
EIF2D / P41214 / ENSMUSG00000026427 / ENSG00000143486 / ENSRNOG00000004910
EIF2S1 / P05198 / ENSMUSG00000021116 / ENSG00000134001 / ENSRNOG00000009432
EIF2S2 / P20042 / ENSMUSG00000074656 / ENSG00000125977 / ENSRNOG00000017447
EIF3A / Q14152 / ENSMUSG00000024991 / ENSG00000107581 / ENSRNOG00000010117
EIF3B / P55884 / ENSMUSG00000056076 / ENSG00000106263 / ENSRNOG00000001253
EIF3C / Q99613 / ENSMUSG00000030738 / ENSG00000184110 / ENSRNOG00000018761
EIF3D / O15371 / ENSMUSG00000016554 / ENSG00000100353 / ENSRNOG00000005804
EIF3E / P60228 / ENSMUSG00000022336 / ENSG00000104408 / ENSRNOG00000027690
EIF3F / O00303 / ENSMUSG00000031029 / ENSG00000175390 / ENSRNOG00000015221
EIF3G / O75821 / ENSMUSG00000070319 / ENSG00000130811 / ENSRNOG00000020619
EIF3L / Q9Y262 / ENSMUSG00000033047 / ENSG00000100129 / ENSRNOG00000011020
EIF3M / Q7L2H7 / ENSMUSG00000027170 / ENSG00000149100 / ENSRNOG00000012738
EIF4A1 / P60842 / ENSMUSG00000059796 / ENSG00000161960 / ENSRNOG00000030628
EIF4A2 / Q14240 / ENSMUSG00000022884 / ENSG00000156976 / ENSRNOG00000001815
EIF4B / P23588 / ENSMUSG00000058655 / ENSG00000063046 / ENSRNOG00000010103
EIF4E / P06730 / ENSMUSG00000028156 / ENSG00000151247 / ENSRNOG00000052343
EIF4E2 / O60573 / ENSMUSG00000026254 / ENSG00000135930 / ENSRNOG00000019634
EIF4EBP2 / Q13542 / ENSMUSG00000020091 / ENSG00000148730 / ENSRNOG00000055391
EIF4ENIF1 / Q9NRA8 / ENSMUSG00000020454 / ENSG00000184708 / ENSRNOG00000018475
EIF4G1 / Q04637 / ENSMUSG00000045983 / ENSG00000114867 / ENSRNOG00000001738
EIF4G3 / O43432 / ENSMUSG00000028760 / ENSG00000075151 / ENSRNOG00000014368
EIF4H / Q15056 / ENSMUSG00000040731 / ENSG00000106682 / ENSRNOG00000001454
EIF5 / P55010 / ENSMUSG00000021282 / ENSG00000100664 / ENSRNOG00000010218
EIF5A2 / Q9GZV4 / ENSMUSG00000050192 / ENSG00000163577 / ENSRNOG00000011859
EIF5AL1 / Q6IS14 / ENSMUSG00000078812 / ENSG00000253626 / ENSRNOG00000016478
EIF6 / P56537 / ENSMUSG00000027613 / ENSG00000242372 / ENSRNOG00000049497
EIPR1 / Q53HC9 / ENSMUSG00000036613 / ENSG00000032389 / ENSRNOG00000009285
ELAC2 / Q9BQ52 / ENSMUSG00000020549 / ENSG00000006744 / ENSRNOG00000003424
ELAVL1 / Q15717 / ENSMUSG00000040028 / ENSG00000066044 / ENSRNOG00000001069
ELAVL2 / Q12926 / ENSMUSG00000008489 / ENSG00000107105 / ENSRNOG00000006853
ELAVL3 / Q14576 / ENSMUSG00000003410 / ENSG00000196361 / ENSRNOG00000013752
ELAVL4 / P26378 / ENSMUSG00000028546 / ENSG00000162374 / ENSRNOG00000023601
ELF1 / P32519 / ENSMUSG00000036461 / ENSG00000120690 / ENSRNOG00000011762
ELF2 / Q15723 / ENSMUSG00000037174 / ENSG00000109381 / ENSRNOG00000010815
ELMO1 / Q92556 / ENSMUSG00000041112 / ENSG00000155849 / ENSRNOG00000059705
Homologs
Brain
Heart
Kidney
Liver
Lung
Testis
Organs
Bins
1
2
3
4
5
Species
Human
Mouse
Rat

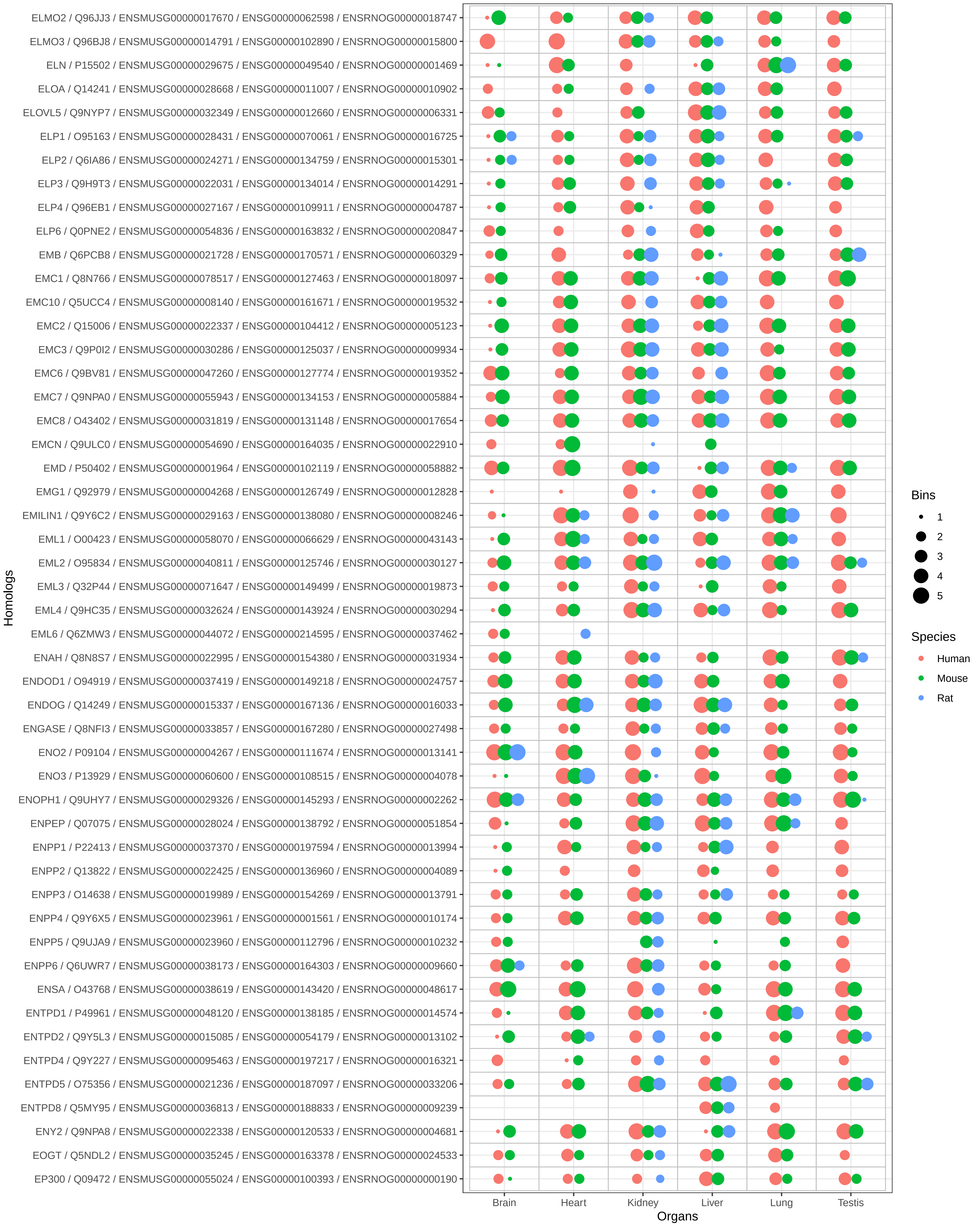
ELMO2 / Q96JJ3 / ENSMUSG00000017670 / ENSG00000062598 / ENSRNOG00000018747
ELMO3 / Q96BJ8 / ENSMUSG00000014791 / ENSG00000102890 / ENSRNOG00000015800
ELN / P15502 / ENSMUSG00000029675 / ENSG00000049540 / ENSRNOG00000001469
ELOA / Q14241 / ENSMUSG00000028668 / ENSG00000011007 / ENSRNOG00000010902
ELOVL5 / Q9NYP7 / ENSMUSG00000032349 / ENSG00000012660 / ENSRNOG00000006331
ELP1 / O95163 / ENSMUSG00000028431 / ENSG00000070061 / ENSRNOG00000016725
ELP2 / Q6IA86 / ENSMUSG00000024271 / ENSG00000134759 / ENSRNOG00000015301
ELP3 / Q9H9T3 / ENSMUSG00000022031 / ENSG00000134014 / ENSRNOG00000014291
ELP4 / Q96EB1 / ENSMUSG00000027167 / ENSG00000109911 / ENSRNOG00000004787
ELP6 / Q0PNE2 / ENSMUSG00000054836 / ENSG00000163832 / ENSRNOG00000020847
EMB / Q6PCB8 / ENSMUSG00000021728 / ENSG00000170571 / ENSRNOG00000060329
EMC1 / Q8N766 / ENSMUSG00000078517 / ENSG00000127463 / ENSRNOG00000018097
EMC10 / Q5UCC4 / ENSMUSG00000008140 / ENSG00000161671 / ENSRNOG00000019532
EMC2 / Q15006 / ENSMUSG00000022337 / ENSG00000104412 / ENSRNOG00000005123
EMC3 / Q9P0I2 / ENSMUSG00000030286 / ENSG00000125037 / ENSRNOG00000009934
EMC6 / Q9BV81 / ENSMUSG00000047260 / ENSG00000127774 / ENSRNOG00000019352
EMC7 / Q9NPA0 / ENSMUSG00000055943 / ENSG00000134153 / ENSRNOG00000005884
EMC8 / O43402 / ENSMUSG00000031819 / ENSG00000131148 / ENSRNOG00000017654
EMCN / Q9ULC0 / ENSMUSG00000054690 / ENSG00000164035 / ENSRNOG00000022910
EMD / P50402 / ENSMUSG00000001964 / ENSG00000102119 / ENSRNOG00000058882
EMG1 / Q92979 / ENSMUSG00000004268 / ENSG00000126749 / ENSRNOG00000012828
EMILIN1 / Q9Y6C2 / ENSMUSG00000029163 / ENSG00000138080 / ENSRNOG00000008246
EML1 / O00423 / ENSMUSG00000058070 / ENSG00000066629 / ENSRNOG00000043143
EML2 / O95834 / ENSMUSG00000040811 / ENSG00000125746 / ENSRNOG00000030127
EML3 / Q32P44 / ENSMUSG00000071647 / ENSG00000149499 / ENSRNOG00000019873
EML4 / Q9HC35 / ENSMUSG00000032624 / ENSG00000143924 / ENSRNOG00000030294
EML6 / Q6ZMW3 / ENSMUSG00000044072 / ENSG00000214595 / ENSRNOG00000037462
ENAH / Q8N8S7 / ENSMUSG00000022995 / ENSG00000154380 / ENSRNOG00000031934
ENDOD1 / O94919 / ENSMUSG00000037419 / ENSG00000149218 / ENSRNOG00000024757
ENDOG / Q14249 / ENSMUSG00000015337 / ENSG00000167136 / ENSRNOG00000016033
ENGASE / Q8NFI3 / ENSMUSG00000033857 / ENSG00000167280 / ENSRNOG00000027498
ENO2 / P09104 / ENSMUSG00000004267 / ENSG00000111674 / ENSRNOG00000013141
ENO3 / P13929 / ENSMUSG00000060600 / ENSG00000108515 / ENSRNOG00000004078
ENOPH1 / Q9UHY7 / ENSMUSG00000029326 / ENSG00000145293 / ENSRNOG00000002262
ENPEP / Q07075 / ENSMUSG00000028024 / ENSG00000138792 / ENSRNOG00000051854
ENPP1 / P22413 / ENSMUSG00000037370 / ENSG00000197594 / ENSRNOG00000013994
ENPP2 / Q13822 / ENSMUSG00000022425 / ENSG00000136960 / ENSRNOG00000004089
ENPP3 / O14638 / ENSMUSG00000019989 / ENSG00000154269 / ENSRNOG00000013791
ENPP4 / Q9Y6X5 / ENSMUSG00000023961 / ENSG00000001561 / ENSRNOG00000010174
ENPP5 / Q9UJA9 / ENSMUSG00000023960 / ENSG00000112796 / ENSRNOG00000010232
ENPP6 / Q6UWR7 / ENSMUSG00000038173 / ENSG00000164303 / ENSRNOG00000009660
ENSA / O43768 / ENSMUSG00000038619 / ENSG00000143420 / ENSRNOG00000048617
ENTPD1 / P49961 / ENSMUSG00000048120 / ENSG00000138185 / ENSRNOG00000014574
ENTPD2 / Q9Y5L3 / ENSMUSG00000015085 / ENSG00000054179 / ENSRNOG00000013102
ENTPD4 / Q9Y227 / ENSMUSG00000095463 / ENSG00000197217 / ENSRNOG00000016321
ENTPD5 / O75356 / ENSMUSG00000021236 / ENSG00000187097 / ENSRNOG00000033206
ENTPD8 / Q5MY95 / ENSMUSG00000036813 / ENSG00000188833 / ENSRNOG00000009239
ENY2 / Q9NPA8 / ENSMUSG00000022338 / ENSG00000120533 / ENSRNOG00000004681
EOGT / Q5NDL2 / ENSMUSG00000035245 / ENSG00000163378 / ENSRNOG00000024533
EP300 / Q09472 / ENSMUSG00000055024 / ENSG00000100393 / ENSRNOG00000000190
Homologs
Brain
Heart
Kidney
Liver
Lung
Testis
Organs
Bins
1
2
3
4
5
Species
Human
Mouse
Rat

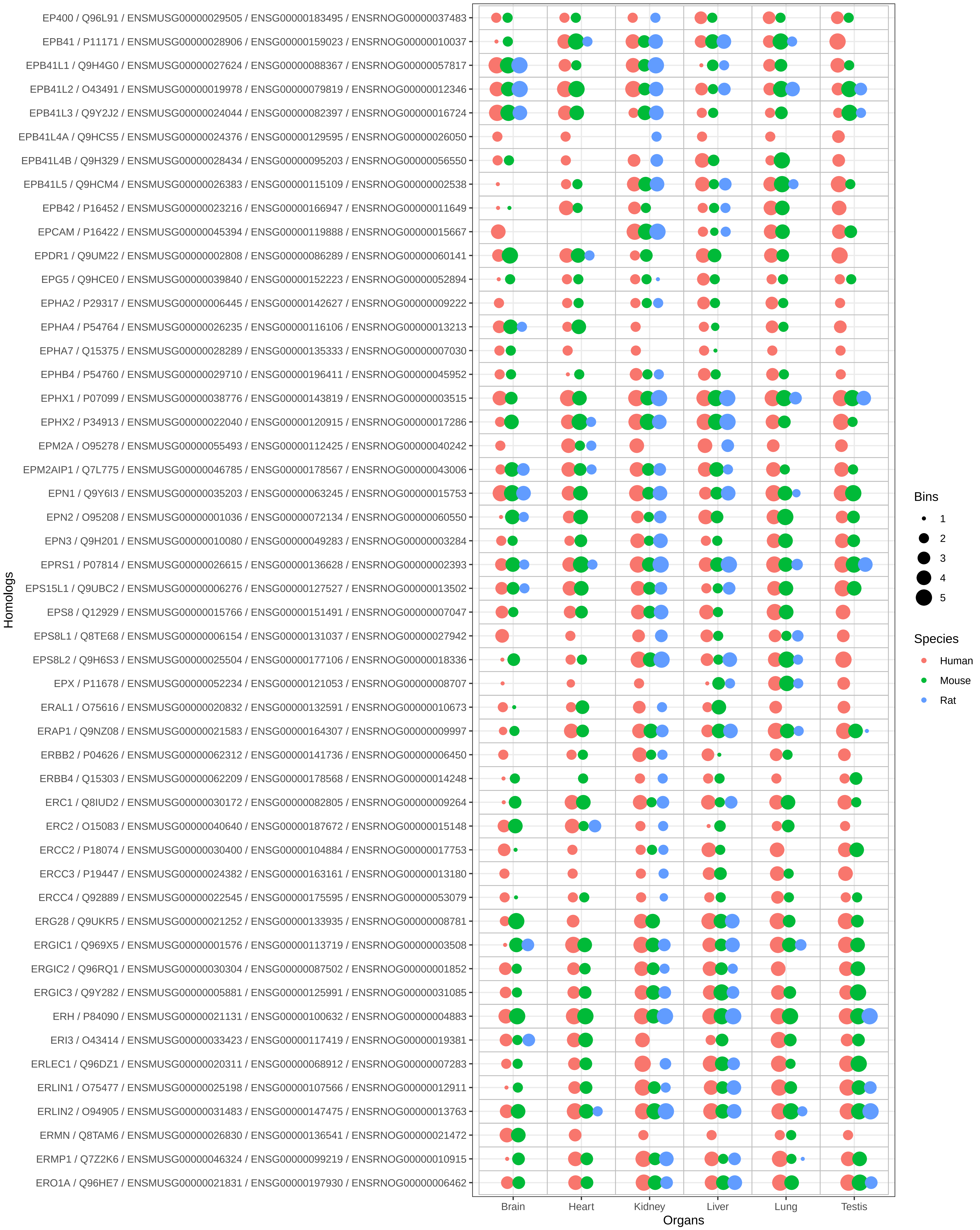
EP400 / Q96L91 / ENSMUSG00000029505 / ENSG00000183495 / ENSRNOG00000037483
EPB41 / P11171 / ENSMUSG00000028906 / ENSG00000159023 / ENSRNOG00000010037
EPB41L1 / Q9H4G0 / ENSMUSG00000027624 / ENSG00000088367 / ENSRNOG00000057817
EPB41L2 / O43491 / ENSMUSG00000019978 / ENSG00000079819 / ENSRNOG00000012346
EPB41L3 / Q9Y2J2 / ENSMUSG00000024044 / ENSG00000082397 / ENSRNOG00000016724
EPB41L4A / Q9HCS5 / ENSMUSG00000024376 / ENSG00000129595 / ENSRNOG00000026050
EPB41L4B / Q9H329 / ENSMUSG00000028434 / ENSG00000095203 / ENSRNOG00000056550
EPB41L5 / Q9HCM4 / ENSMUSG00000026383 / ENSG00000115109 / ENSRNOG00000002538
EPB42 / P16452 / ENSMUSG00000023216 / ENSG00000166947 / ENSRNOG00000011649
EPCAM / P16422 / ENSMUSG00000045394 / ENSG00000119888 / ENSRNOG00000015667
EPDR1 / Q9UM22 / ENSMUSG00000002808 / ENSG00000086289 / ENSRNOG00000060141
EPG5 / Q9HCE0 / ENSMUSG00000039840 / ENSG00000152223 / ENSRNOG00000052894
EPHA2 / P29317 / ENSMUSG00000006445 / ENSG00000142627 / ENSRNOG00000009222
EPHA4 / P54764 / ENSMUSG00000026235 / ENSG00000116106 / ENSRNOG00000013213
EPHA7 / Q15375 / ENSMUSG00000028289 / ENSG00000135333 / ENSRNOG00000007030
EPHB4 / P54760 / ENSMUSG00000029710 / ENSG00000196411 / ENSRNOG00000045952
EPHX1 / P07099 / ENSMUSG00000038776 / ENSG00000143819 / ENSRNOG00000003515
EPHX2 / P34913 / ENSMUSG00000022040 / ENSG00000120915 / ENSRNOG00000017286
EPM2A / O95278 / ENSMUSG00000055493 / ENSG00000112425 / ENSRNOG00000040242
EPM2AIP1 / Q7L775 / ENSMUSG00000046785 / ENSG00000178567 / ENSRNOG00000043006
EPN1 / Q9Y6I3 / ENSMUSG00000035203 / ENSG00000063245 / ENSRNOG00000015753
EPN2 / O95208 / ENSMUSG00000001036 / ENSG00000072134 / ENSRNOG00000060550
EPN3 / Q9H201 / ENSMUSG00000010080 / ENSG00000049283 / ENSRNOG00000003284
EPRS1 / P07814 / ENSMUSG00000026615 / ENSG00000136628 / ENSRNOG00000002393
EPS15L1 / Q9UBC2 / ENSMUSG00000006276 / ENSG00000127527 / ENSRNOG00000013502
EPS8 / Q12929 / ENSMUSG00000015766 / ENSG00000151491 / ENSRNOG00000007047
EPS8L1 / Q8TE68 / ENSMUSG00000006154 / ENSG00000131037 / ENSRNOG00000027942
EPS8L2 / Q9H6S3 / ENSMUSG00000025504 / ENSG00000177106 / ENSRNOG00000018336
EPX / P11678 / ENSMUSG00000052234 / ENSG00000121053 / ENSRNOG00000008707
ERAL1 / O75616 / ENSMUSG00000020832 / ENSG00000132591 / ENSRNOG00000010673
ERAP1 / Q9NZ08 / ENSMUSG00000021583 / ENSG00000164307 / ENSRNOG00000009997
ERBB2 / P04626 / ENSMUSG00000062312 / ENSG00000141736 / ENSRNOG00000006450
ERBB4 / Q15303 / ENSMUSG00000062209 / ENSG00000178568 / ENSRNOG00000014248
ERC1 / Q8IUD2 / ENSMUSG00000030172 / ENSG00000082805 / ENSRNOG00000009264
ERC2 / O15083 / ENSMUSG00000040640 / ENSG00000187672 / ENSRNOG00000015148
ERCC2 / P18074 / ENSMUSG00000030400 / ENSG00000104884 / ENSRNOG00000017753
ERCC3 / P19447 / ENSMUSG00000024382 / ENSG00000163161 / ENSRNOG00000013180
ERCC4 / Q92889 / ENSMUSG00000022545 / ENSG00000175595 / ENSRNOG00000053079
ERG28 / Q9UKR5 / ENSMUSG00000021252 / ENSG00000133935 / ENSRNOG00000008781
ERGIC1 / Q969X5 / ENSMUSG00000001576 / ENSG00000113719 / ENSRNOG00000003508
ERGIC2 / Q96RQ1 / ENSMUSG00000030304 / ENSG00000087502 / ENSRNOG00000001852
ERGIC3 / Q9Y282 / ENSMUSG00000005881 / ENSG00000125991 / ENSRNOG00000031085
ERH / P84090 / ENSMUSG00000021131 / ENSG00000100632 / ENSRNOG00000004883
ERI3 / O43414 / ENSMUSG00000033423 / ENSG00000117419 / ENSRNOG00000019381
ERLEC1 / Q96DZ1 / ENSMUSG00000020311 / ENSG00000068912 / ENSRNOG00000007283
ERLIN1 / O75477 / ENSMUSG00000025198 / ENSG00000107566 / ENSRNOG00000012911
ERLIN2 / O94905 / ENSMUSG00000031483 / ENSG00000147475 / ENSRNOG00000013763
ERMN / Q8TAM6 / ENSMUSG00000026830 / ENSG00000136541 / ENSRNOG00000021472
ERMP1 / Q7Z2K6 / ENSMUSG00000046324 / ENSG00000099219 / ENSRNOG00000010915
ERO1A / Q96HE7 / ENSMUSG00000021831 / ENSG00000197930 / ENSRNOG00000006462
Homologs
Brain
Heart
Kidney
Liver
Lung
Testis
Organs
Bins
1
2
3
4
5
Species
Human
Mouse
Rat

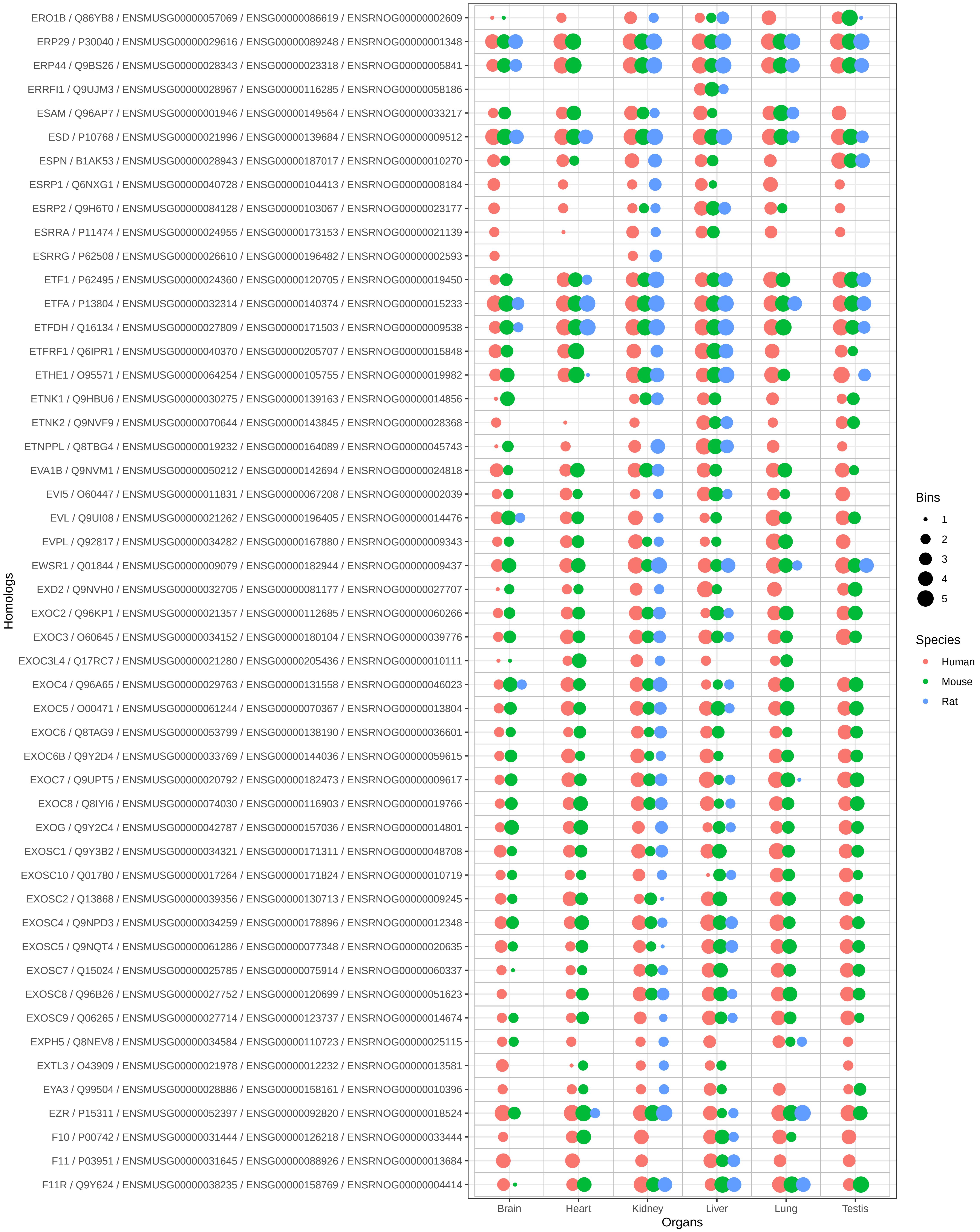
ERO1B / Q86YB8 / ENSMUSG00000057069 / ENSG00000086619 / ENSRNOG00000002609
ERP29 / P30040 / ENSMUSG00000029616 / ENSG00000089248 / ENSRNOG00000001348
ERP44 / Q9BS26 / ENSMUSG00000028343 / ENSG00000023318 / ENSRNOG00000005841
ERRFI1 / Q9UJM3 / ENSMUSG00000028967 / ENSG00000116285 / ENSRNOG00000058186
ESAM / Q96AP7 / ENSMUSG00000001946 / ENSG00000149564 / ENSRNOG00000033217
ESD / P10768 / ENSMUSG00000021996 / ENSG00000139684 / ENSRNOG00000009512
ESPN / B1AK53 / ENSMUSG00000028943 / ENSG00000187017 / ENSRNOG00000010270
ESRP1 / Q6NXG1 / ENSMUSG00000040728 / ENSG00000104413 / ENSRNOG00000008184
ESRP2 / Q9H6T0 / ENSMUSG00000084128 / ENSG00000103067 / ENSRNOG00000023177
ESRRA / P11474 / ENSMUSG00000024955 / ENSG00000173153 / ENSRNOG00000021139
ESRRG / P62508 / ENSMUSG00000026610 / ENSG00000196482 / ENSRNOG00000002593
ETF1 / P62495 / ENSMUSG00000024360 / ENSG00000120705 / ENSRNOG00000019450
ETFA / P13804 / ENSMUSG00000032314 / ENSG00000140374 / ENSRNOG00000015233
ETFDH / Q16134 / ENSMUSG00000027809 / ENSG00000171503 / ENSRNOG00000009538
ETFRF1 / Q6IPR1 / ENSMUSG00000040370 / ENSG00000205707 / ENSRNOG00000015848
ETHE1 / O95571 / ENSMUSG00000064254 / ENSG00000105755 / ENSRNOG00000019982
ETNK1 / Q9HBU6 / ENSMUSG00000030275 / ENSG00000139163 / ENSRNOG00000014856
ETNK2 / Q9NVF9 / ENSMUSG00000070644 / ENSG00000143845 / ENSRNOG00000028368
ETNPPL / Q8TBG4 / ENSMUSG00000019232 / ENSG00000164089 / ENSRNOG00000045743
EVA1B / Q9NVM1 / ENSMUSG00000050212 / ENSG00000142694 / ENSRNOG00000024818
EVI5 / O60447 / ENSMUSG00000011831 / ENSG00000067208 / ENSRNOG00000002039
EVL / Q9UI08 / ENSMUSG00000021262 / ENSG00000196405 / ENSRNOG00000014476
EVPL / Q92817 / ENSMUSG00000034282 / ENSG00000167880 / ENSRNOG00000009343
EWSR1 / Q01844 / ENSMUSG00000009079 / ENSG00000182944 / ENSRNOG00000009437
EXD2 / Q9NVH0 / ENSMUSG00000032705 / ENSG00000081177 / ENSRNOG00000027707
EXOC2 / Q96KP1 / ENSMUSG00000021357 / ENSG00000112685 / ENSRNOG00000060266
EXOC3 / O60645 / ENSMUSG00000034152 / ENSG00000180104 / ENSRNOG00000039776
EXOC3L4 / Q17RC7 / ENSMUSG00000021280 / ENSG00000205436 / ENSRNOG00000010111
EXOC4 / Q96A65 / ENSMUSG00000029763 / ENSG00000131558 / ENSRNOG00000046023
EXOC5 / O00471 / ENSMUSG00000061244 / ENSG00000070367 / ENSRNOG00000013804
EXOC6 / Q8TAG9 / ENSMUSG00000053799 / ENSG00000138190 / ENSRNOG00000036601
EXOC6B / Q9Y2D4 / ENSMUSG00000033769 / ENSG00000144036 / ENSRNOG00000059615
EXOC7 / Q9UPT5 / ENSMUSG00000020792 / ENSG00000182473 / ENSRNOG00000009617
EXOC8 / Q8IYI6 / ENSMUSG00000074030 / ENSG00000116903 / ENSRNOG00000019766
EXOG / Q9Y2C4 / ENSMUSG00000042787 / ENSG00000157036 / ENSRNOG00000014801
EXOSC1 / Q9Y3B2 / ENSMUSG00000034321 / ENSG00000171311 / ENSRNOG00000048708
EXOSC10 / Q01780 / ENSMUSG00000017264 / ENSG00000171824 / ENSRNOG00000010719
EXOSC2 / Q13868 / ENSMUSG00000039356 / ENSG00000130713 / ENSRNOG00000009245
EXOSC4 / Q9NPD3 / ENSMUSG00000034259 / ENSG00000178896 / ENSRNOG00000012348
EXOSC5 / Q9NQT4 / ENSMUSG00000061286 / ENSG00000077348 / ENSRNOG00000020635
EXOSC7 / Q15024 / ENSMUSG00000025785 / ENSG00000075914 / ENSRNOG00000060337
EXOSC8 / Q96B26 / ENSMUSG00000027752 / ENSG00000120699 / ENSRNOG00000051623
EXOSC9 / Q06265 / ENSMUSG00000027714 / ENSG00000123737 / ENSRNOG00000014674
EXPH5 / Q8NEV8 / ENSMUSG00000034584 / ENSG00000110723 / ENSRNOG00000025115
EXTL3 / O43909 / ENSMUSG00000021978 / ENSG00000012232 / ENSRNOG00000013581
EYA3 / Q99504 / ENSMUSG00000028886 / ENSG00000158161 / ENSRNOG00000010396
EZR / P15311 / ENSMUSG00000052397 / ENSG00000092820 / ENSRNOG00000018524
F10 / P00742 / ENSMUSG00000031444 / ENSG00000126218 / ENSRNOG00000033444
F11 / P03951 / ENSMUSG00000031645 / ENSG00000088926 / ENSRNOG00000013684
F11R / Q9Y624 / ENSMUSG00000038235 / ENSG00000158769 / ENSRNOG00000004414
Homologs
Brain
Heart
Kidney
Liver
Lung
Testis
Organs
Bins
1
2
3
4
5
Species
Human
Mouse
Rat

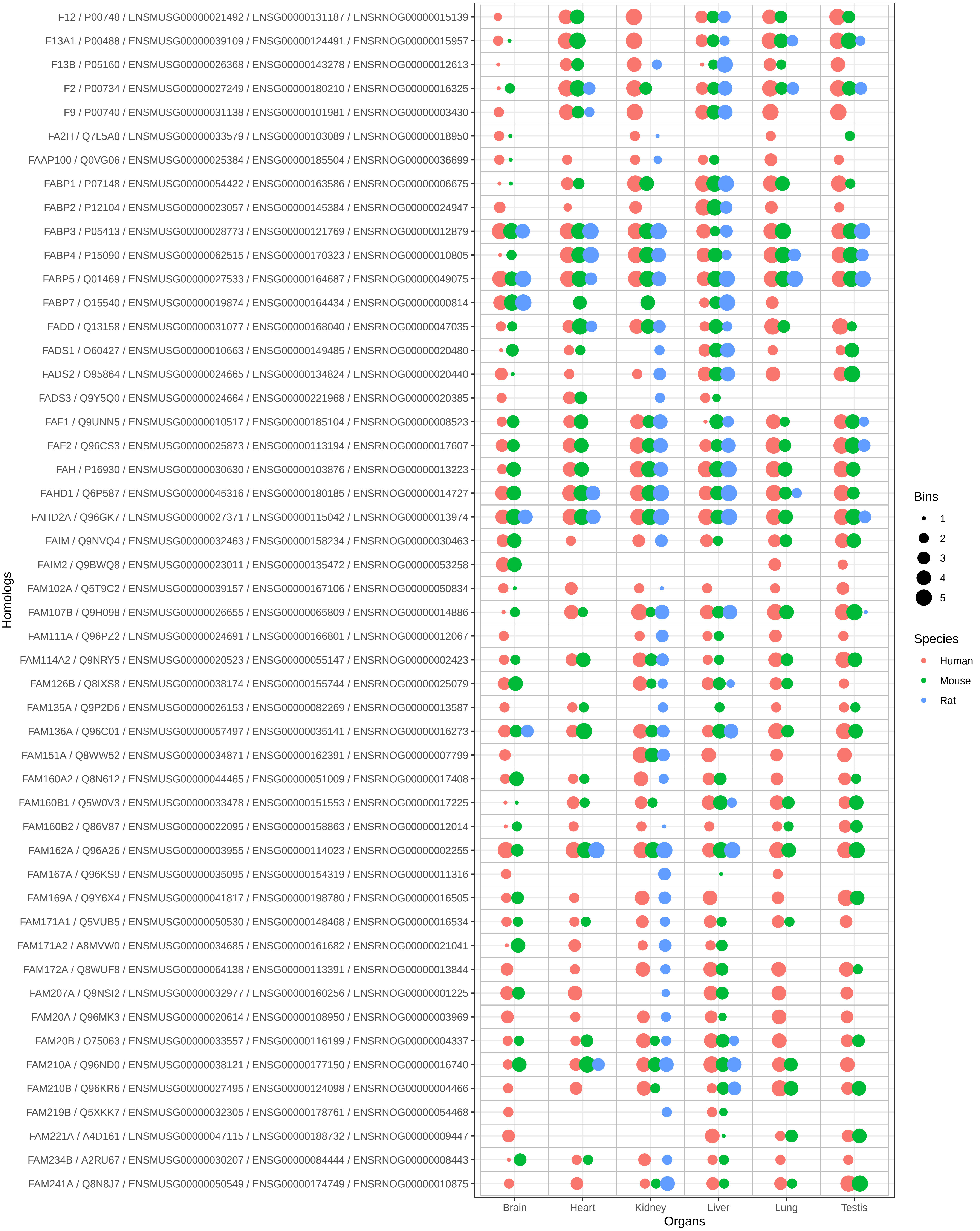

F12 / P00748 / ENSMUSG00000021492 / ENSG00000131187 / ENSRNOG00000015139
F13A1 / P00488 / ENSMUSG00000039109 / ENSG00000124491 / ENSRNOG00000015957
F13B / P05160 / ENSMUSG00000026368 / ENSG00000143278 / ENSRNOG00000012613
F2 / P00734 / ENSMUSG00000027249 / ENSG00000180210 / ENSRNOG00000016325
F9 / P00740 / ENSMUSG00000031138 / ENSG00000101981 / ENSRNOG00000003430
FA2H / Q7L5A8 / ENSMUSG00000033579 / ENSG00000103089 / ENSRNOG00000018950
FAAP100 / Q0VG06 / ENSMUSG00000025384 / ENSG00000185504 / ENSRNOG00000036699
FABP1 / P07148 / ENSMUSG00000054422 / ENSG00000163586 / ENSRNOG00000006675
FABP2 / P12104 / ENSMUSG00000023057 / ENSG00000145384 / ENSRNOG00000024947
FABP3 / P05413 / ENSMUSG00000028773 / ENSG00000121769 / ENSRNOG00000012879
FABP4 / P15090 / ENSMUSG00000062515 / ENSG00000170323 / ENSRNOG00000010805
FABP5 / Q01469 / ENSMUSG00000027533 / ENSG00000164687 / ENSRNOG00000049075
FABP7 / O15540 / ENSMUSG00000019874 / ENSG00000164434 / ENSRNOG00000000814
FADD / Q13158 / ENSMUSG00000031077 / ENSG00000168040 / ENSRNOG00000047035
FADS1 / O60427 / ENSMUSG00000010663 / ENSG00000149485 / ENSRNOG00000020480
FADS2 / O95864 / ENSMUSG00000024665 / ENSG00000134824 / ENSRNOG00000020440
FADS3 / Q9Y5Q0 / ENSMUSG00000024664 / ENSG00000221968 / ENSRNOG00000020385
FAF1 / Q9UNN5 / ENSMUSG00000010517 / ENSG00000185104 / ENSRNOG00000008523
FAF2 / Q96CS3 / ENSMUSG00000025873 / ENSG00000113194 / ENSRNOG00000017607
FAH / P16930 / ENSMUSG00000030630 / ENSG00000103876 / ENSRNOG00000013223
FAHD1 / Q6P587 / ENSMUSG00000045316 / ENSG00000180185 / ENSRNOG00000014727
FAHD2A / Q96GK7 / ENSMUSG00000027371 / ENSG00000115042 / ENSRNOG00000013974
FAIM / Q9NVQ4 / ENSMUSG00000032463 / ENSG00000158234 / ENSRNOG00000030463
FAIM2 / Q9BWQ8 / ENSMUSG00000023011 / ENSG00000135472 / ENSRNOG00000053258
FAM102A / Q5T9C2 / ENSMUSG00000039157 / ENSG00000167106 / ENSRNOG00000050834
FAM107B / Q9H098 / ENSMUSG00000026655 / ENSG00000065809 / ENSRNOG00000014886
FAM111A / Q96PZ2 / ENSMUSG00000024691 / ENSG00000166801 / ENSRNOG00000012067
FAM114A2 / Q9NRY5 / ENSMUSG00000020523 / ENSG00000055147 / ENSRNOG00000002423
FAM126B / Q8IXS8 / ENSMUSG00000038174 / ENSG00000155744 / ENSRNOG00000025079
FAM135A / Q9P2D6 / ENSMUSG00000026153 / ENSG00000082269 / ENSRNOG00000013587
FAM136A / Q96C01 / ENSMUSG00000057497 / ENSG00000035141 / ENSRNOG00000016273
FAM151A / Q8WW52 / ENSMUSG00000034871 / ENSG00000162391 / ENSRNOG00000007799
FAM160A2 / Q8N612 / ENSMUSG00000044465 / ENSG00000051009 / ENSRNOG00000017408
FAM160B1 / Q5W0V3 / ENSMUSG00000033478 / ENSG00000151553 / ENSRNOG00000017225
FAM160B2 / Q86V87 / ENSMUSG00000022095 / ENSG00000158863 / ENSRNOG00000012014
FAM162A / Q96A26 / ENSMUSG00000003955 / ENSG00000114023 / ENSRNOG00000002255
FAM167A / Q96KS9 / ENSMUSG00000035095 / ENSG00000154319 / ENSRNOG00000011316
FAM169A / Q9Y6X4 / ENSMUSG00000041817 / ENSG00000198780 / ENSRNOG00000016505
FAM171A1 / Q5VUB5 / ENSMUSG00000050530 / ENSG00000148468 / ENSRNOG00000016534
FAM171A2 / A8MVW0 / ENSMUSG00000034685 / ENSG00000161682 / ENSRNOG00000021041
FAM172A / Q8WUF8 / ENSMUSG00000064138 / ENSG00000113391 / ENSRNOG00000013844
FAM207A / Q9NSI2 / ENSMUSG00000032977 / ENSG00000160256 / ENSRNOG00000001225
FAM20A / Q96MK3 / ENSMUSG00000020614 / ENSG00000108950 / ENSRNOG00000003969
FAM20B / O75063 / ENSMUSG00000033557 / ENSG00000116199 / ENSRNOG00000004337
FAM210A / Q96ND0 / ENSMUSG00000038121 / ENSG00000177150 / ENSRNOG00000016740
FAM210B / Q96KR6 / ENSMUSG00000027495 / ENSG00000124098 / ENSRNOG00000004466
FAM219B / Q5XKK7 / ENSMUSG00000032305 / ENSG00000178761 / ENSRNOG00000054468
FAM221A / A4D161 / ENSMUSG00000047115 / ENSG00000188732 / ENSRNOG00000009447
FAM234B / A2RU67 / ENSMUSG00000030207 / ENSG00000084444 / ENSRNOG00000008443
FAM241A / Q8N8J7 / ENSMUSG00000050549 / ENSG00000174749 / ENSRNOG00000010875
Homologs
Brain
Heart
Kidney
Liver
Lung
Testis
Organs
Bins
1
2
3
4
5
Species
Human
Mouse
Rat

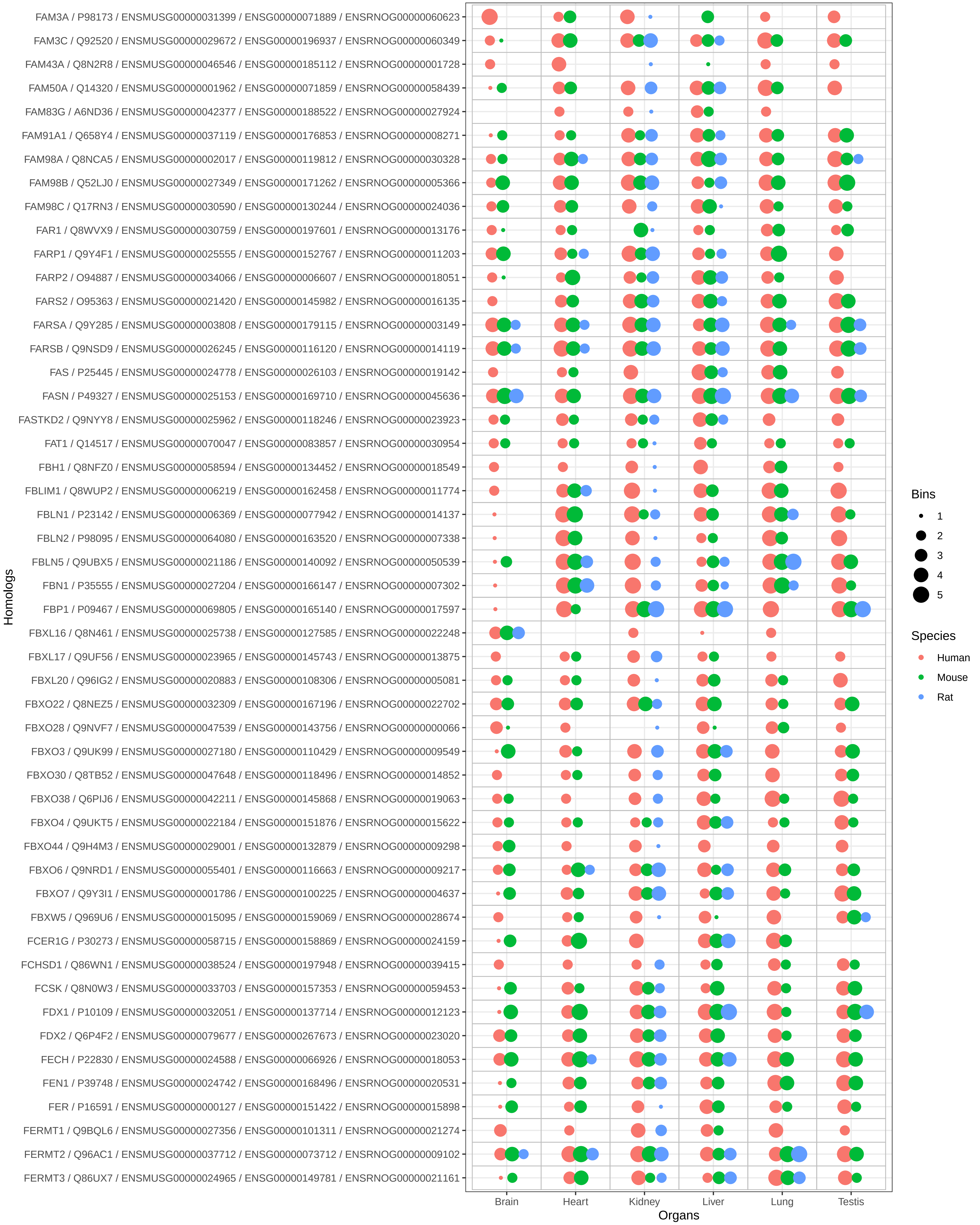
FAM3A / P98173 / ENSMUSG00000031399 / ENSG00000071889 / ENSRNOG00000060623
FAM3C / Q92520 / ENSMUSG00000029672 / ENSG00000196937 / ENSRNOG00000060349
FAM43A / Q8N2R8 / ENSMUSG00000046546 / ENSG00000185112 / ENSRNOG00000001728
FAM50A / Q14320 / ENSMUSG00000001962 / ENSG00000071859 / ENSRNOG00000058439
FAM83G / A6ND36 / ENSMUSG00000042377 / ENSG00000188522 / ENSRNOG00000027924
FAM91A1 / Q658Y4 / ENSMUSG00000037119 / ENSG00000176853 / ENSRNOG00000008271
FAM98A / Q8NCA5 / ENSMUSG00000002017 / ENSG00000119812 / ENSRNOG00000030328
FAM98B / Q52LJ0 / ENSMUSG00000027349 / ENSG00000171262 / ENSRNOG00000005366
FAM98C / Q17RN3 / ENSMUSG00000030590 / ENSG00000130244 / ENSRNOG00000024036
FAR1 / Q8WVX9 / ENSMUSG00000030759 / ENSG00000197601 / ENSRNOG00000013176
FARP1 / Q9Y4F1 / ENSMUSG00000025555 / ENSG00000152767 / ENSRNOG00000011203
FARP2 / O94887 / ENSMUSG00000034066 / ENSG00000006607 / ENSRNOG00000018051
FARS2 / O95363 / ENSMUSG00000021420 / ENSG00000145982 / ENSRNOG00000016135
FARSA / Q9Y285 / ENSMUSG00000003808 / ENSG00000179115 / ENSRNOG00000003149
FARSB / Q9NSD9 / ENSMUSG00000026245 / ENSG00000116120 / ENSRNOG00000014119
FAS / P25445 / ENSMUSG00000024778 / ENSG00000026103 / ENSRNOG00000019142
FASN / P49327 / ENSMUSG00000025153 / ENSG00000169710 / ENSRNOG00000045636
FASTKD2 / Q9NYY8 / ENSMUSG00000025962 / ENSG00000118246 / ENSRNOG00000023923
FAT1 / Q14517 / ENSMUSG00000070047 / ENSG00000083857 / ENSRNOG00000030954
FBH1 / Q8NFZ0 / ENSMUSG00000058594 / ENSG00000134452 / ENSRNOG00000018549
FBLIM1 / Q8WUP2 / ENSMUSG00000006219 / ENSG00000162458 / ENSRNOG00000011774
FBLN1 / P23142 / ENSMUSG00000006369 / ENSG00000077942 / ENSRNOG00000014137
FBLN2 / P98095 / ENSMUSG00000064080 / ENSG00000163520 / ENSRNOG00000007338
FBLN5 / Q9UBX5 / ENSMUSG00000021186 / ENSG00000140092 / ENSRNOG00000050539
FBN1 / P35555 / ENSMUSG00000027204 / ENSG00000166147 / ENSRNOG00000007302
FBP1 / P09467 / ENSMUSG00000069805 / ENSG00000165140 / ENSRNOG00000017597
FBXL16 / Q8N461 / ENSMUSG00000025738 / ENSG00000127585 / ENSRNOG00000022248
FBXL17 / Q9UF56 / ENSMUSG00000023965 / ENSG00000145743 / ENSRNOG00000013875
FBXL20 / Q96IG2 / ENSMUSG00000020883 / ENSG00000108306 / ENSRNOG00000005081
FBXO22 / Q8NEZ5 / ENSMUSG00000032309 / ENSG00000167196 / ENSRNOG00000022702
FBXO28 / Q9NVF7 / ENSMUSG00000047539 / ENSG00000143756 / ENSRNOG00000000066
FBXO3 / Q9UK99 / ENSMUSG00000027180 / ENSG00000110429 / ENSRNOG00000009549
FBXO30 / Q8TB52 / ENSMUSG00000047648 / ENSG00000118496 / ENSRNOG00000014852
FBXO38 / Q6PIJ6 / ENSMUSG00000042211 / ENSG00000145868 / ENSRNOG00000019063
FBXO4 / Q9UKT5 / ENSMUSG00000022184 / ENSG00000151876 / ENSRNOG00000015622
FBXO44 / Q9H4M3 / ENSMUSG00000029001 / ENSG00000132879 / ENSRNOG00000009298
FBXO6 / Q9NRD1 / ENSMUSG00000055401 / ENSG00000116663 / ENSRNOG00000009217
FBXO7 / Q9Y3I1 / ENSMUSG00000001786 / ENSG00000100225 / ENSRNOG00000004637
FBXW5 / Q969U6 / ENSMUSG00000015095 / ENSG00000159069 / ENSRNOG00000028674
FCER1G / P30273 / ENSMUSG00000058715 / ENSG00000158869 / ENSRNOG00000024159
FCHSD1 / Q86WN1 / ENSMUSG00000038524 / ENSG00000197948 / ENSRNOG00000039415
FCSK / Q8N0W3 / ENSMUSG00000033703 / ENSG00000157353 / ENSRNOG00000059453
FDX1 / P10109 / ENSMUSG00000032051 / ENSG00000137714 / ENSRNOG00000012123
FDX2 / Q6P4F2 / ENSMUSG00000079677 / ENSG00000267673 / ENSRNOG00000023020
FECH / P22830 / ENSMUSG00000024588 / ENSG00000066926 / ENSRNOG00000018053
FEN1 / P39748 / ENSMUSG00000024742 / ENSG00000168496 / ENSRNOG00000020531
FER / P16591 / ENSMUSG00000000127 / ENSG00000151422 / ENSRNOG00000015898
FERMT1 / Q9BQL6 / ENSMUSG00000027356 / ENSG00000101311 / ENSRNOG00000021274
FERMT2 / Q96AC1 / ENSMUSG00000037712 / ENSG00000073712 / ENSRNOG00000009102
FERMT3 / Q86UX7 / ENSMUSG00000024965 / ENSG00000149781 / ENSRNOG00000021161
Homologs
Brain
Heart
Kidney
Liver
Lung
Testis
Organs
Bins
1
2
3
4
5
Species
Human
Mouse
Rat

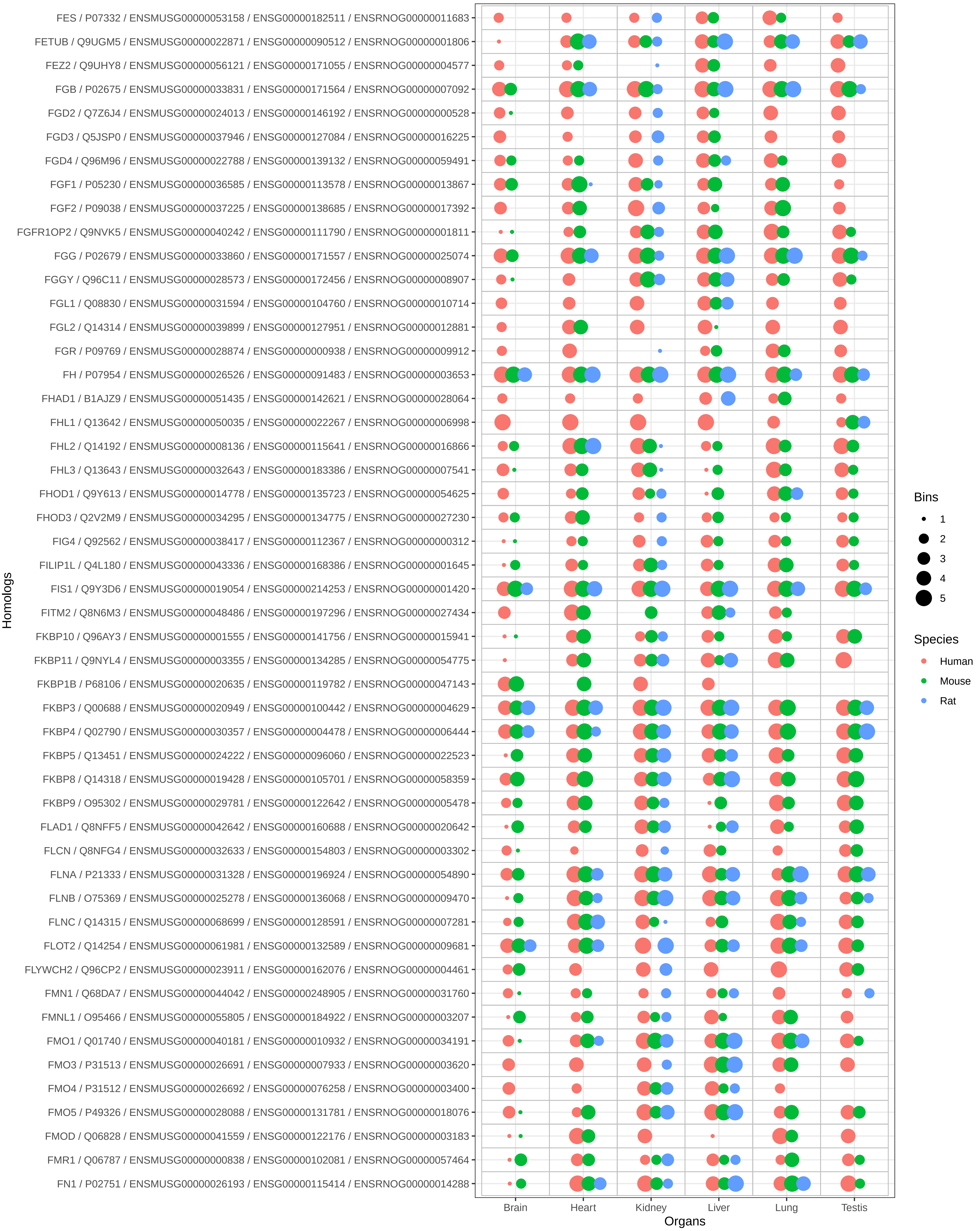
FES / P07332 / ENSMUSG00000053158 / ENSG00000182511 / ENSRNOG00000011683
FETUB / Q9UGM5 / ENSMUSG00000022871 / ENSG00000090512 / ENSRNOG00000001806
FEZ2 / Q9UHY8 / ENSMUSG00000056121 / ENSG00000171055 / ENSRNOG00000004577
FGB / P02675 / ENSMUSG00000033831 / ENSG00000171564 / ENSRNOG00000007092
FGD2 / Q7Z6J4 / ENSMUSG00000024013 / ENSG00000146192 / ENSRNOG00000000528
FGD3 / Q5JSP0 / ENSMUSG00000037946 / ENSG00000127084 / ENSRNOG00000016225
FGD4 / Q96M96 / ENSMUSG00000022788 / ENSG00000139132 / ENSRNOG00000059491
FGF1 / P05230 / ENSMUSG00000036585 / ENSG00000113578 / ENSRNOG00000013867
FGF2 / P09038 / ENSMUSG00000037225 / ENSG00000138685 / ENSRNOG00000017392
FGFR1OP2 / Q9NVK5 / ENSMUSG00000040242 / ENSG00000111790 / ENSRNOG00000001811
FGG / P02679 / ENSMUSG00000033860 / ENSG00000171557 / ENSRNOG00000025074
FGGY / Q96C11 / ENSMUSG00000028573 / ENSG00000172456 / ENSRNOG00000008907
FGL1 / Q08830 / ENSMUSG00000031594 / ENSG00000104760 / ENSRNOG00000010714
FGL2 / Q14314 / ENSMUSG00000039899 / ENSG00000127951 / ENSRNOG00000012881
FGR / P09769 / ENSMUSG00000028874 / ENSG00000000938 / ENSRNOG00000009912
FH / P07954 / ENSMUSG00000026526 / ENSG00000091483 / ENSRNOG00000003653
FHAD1 / B1AJZ9 / ENSMUSG00000051435 / ENSG00000142621 / ENSRNOG00000028064
FHL1 / Q13642 / ENSMUSG00000050035 / ENSG00000022267 / ENSRNOG00000006998
FHL2 / Q14192 / ENSMUSG00000008136 / ENSG00000115641 / ENSRNOG00000016866
FHL3 / Q13643 / ENSMUSG00000032643 / ENSG00000183386 / ENSRNOG00000007541
FHOD1 / Q9Y613 / ENSMUSG00000014778 / ENSG00000135723 / ENSRNOG00000054625
FHOD3 / Q2V2M9 / ENSMUSG00000034295 / ENSG00000134775 / ENSRNOG00000027230
FIG4 / Q92562 / ENSMUSG00000038417 / ENSG00000112367 / ENSRNOG00000000312
FILIP1L / Q4L180 / ENSMUSG00000043336 / ENSG00000168386 / ENSRNOG00000001645
FIS1 / Q9Y3D6 / ENSMUSG00000019054 / ENSG00000214253 / ENSRNOG00000001420
FITM2 / Q8N6M3 / ENSMUSG00000048486 / ENSG00000197296 / ENSRNOG00000027434
FKBP10 / Q96AY3 / ENSMUSG00000001555 / ENSG00000141756 / ENSRNOG00000015941
FKBP11 / Q9NYL4 / ENSMUSG00000003355 / ENSG00000134285 / ENSRNOG00000054775
FKBP1B / P68106 / ENSMUSG00000020635 / ENSG00000119782 / ENSRNOG00000047143
FKBP3 / Q00688 / ENSMUSG00000020949 / ENSG00000100442 / ENSRNOG00000004629
FKBP4 / Q02790 / ENSMUSG00000030357 / ENSG00000004478 / ENSRNOG00000006444
FKBP5 / Q13451 / ENSMUSG00000024222 / ENSG00000096060 / ENSRNOG00000022523
FKBP8 / Q14318 / ENSMUSG00000019428 / ENSG00000105701 / ENSRNOG00000058359
FKBP9 / O95302 / ENSMUSG00000029781 / ENSG00000122642 / ENSRNOG00000005478
FLAD1 / Q8NFF5 / ENSMUSG00000042642 / ENSG00000160688 / ENSRNOG00000020642
FLCN / Q8NFG4 / ENSMUSG00000032633 / ENSG00000154803 / ENSRNOG00000003302
FLNA / P21333 / ENSMUSG00000031328 / ENSG00000196924 / ENSRNOG00000054890
FLNB / O75369 / ENSMUSG00000025278 / ENSG00000136068 / ENSRNOG00000009470
FLNC / Q14315 / ENSMUSG00000068699 / ENSG00000128591 / ENSRNOG00000007281
FLOT2 / Q14254 / ENSMUSG00000061981 / ENSG00000132589 / ENSRNOG00000009681
FLYWCH2 / Q96CP2 / ENSMUSG00000023911 / ENSG00000162076 / ENSRNOG00000004461
FMN1 / Q68DA7 / ENSMUSG00000044042 / ENSG00000248905 / ENSRNOG00000031760
FMNL1 / O95466 / ENSMUSG00000055805 / ENSG00000184922 / ENSRNOG00000003207
FMO1 / Q01740 / ENSMUSG00000040181 / ENSG00000010932 / ENSRNOG00000034191
FMO3 / P31513 / ENSMUSG00000026691 / ENSG00000007933 / ENSRNOG00000003620
FMO4 / P31512 / ENSMUSG00000026692 / ENSG00000076258 / ENSRNOG00000003400
FMO5 / P49326 / ENSMUSG00000028088 / ENSG00000131781 / ENSRNOG00000018076
FMOD / Q06828 / ENSMUSG00000041559 / ENSG00000122176 / ENSRNOG00000003183
FMR1 / Q06787 / ENSMUSG00000000838 / ENSG00000102081 / ENSRNOG00000057464
FN1 / P02751 / ENSMUSG00000026193 / ENSG00000115414 / ENSRNOG00000014288
Homologs
Brain
Heart
Kidney
Liver
Lung
Testis
Organs
Bins
1
2
3
4
5
Species
Human
Mouse
Rat

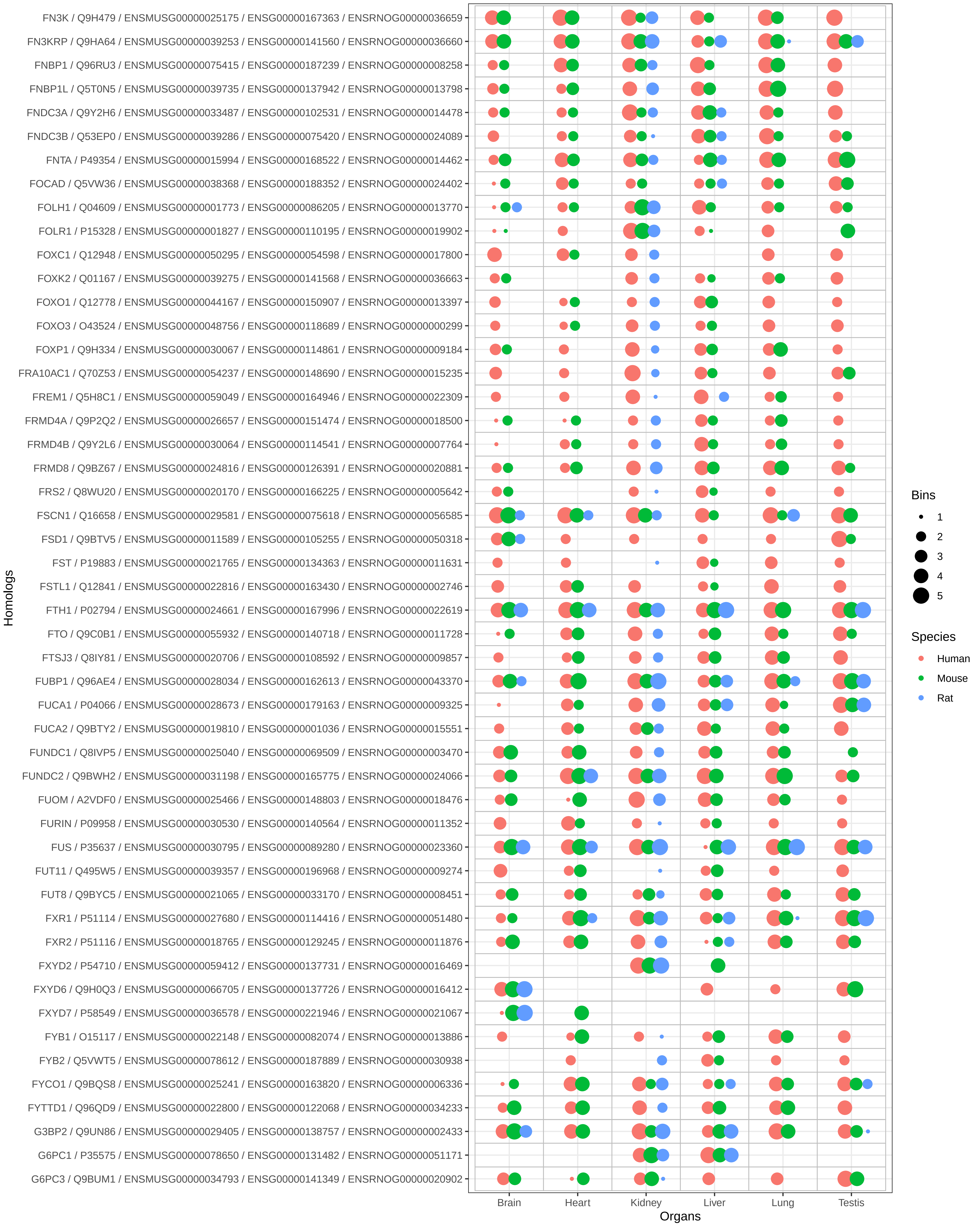

FN3K / Q9H479 / ENSMUSG00000025175 / ENSG00000167363 / ENSRNOG00000036659
FN3KRP / Q9HA64 / ENSMUSG00000039253 / ENSG00000141560 / ENSRNOG00000036660
FNBP1 / Q96RU3 / ENSMUSG00000075415 / ENSG00000187239 / ENSRNOG00000008258
FNBP1L / Q5T0N5 / ENSMUSG00000039735 / ENSG00000137942 / ENSRNOG00000013798
FNDC3A / Q9Y2H6 / ENSMUSG00000033487 / ENSG00000102531 / ENSRNOG00000014478
FNDC3B / Q53EP0 / ENSMUSG00000039286 / ENSG00000075420 / ENSRNOG00000024089
FNTA / P49354 / ENSMUSG00000015994 / ENSG00000168522 / ENSRNOG00000014462
FOCAD / Q5VW36 / ENSMUSG00000038368 / ENSG00000188352 / ENSRNOG00000024402
FOLH1 / Q04609 / ENSMUSG00000001773 / ENSG00000086205 / ENSRNOG00000013770
FOLR1 / P15328 / ENSMUSG00000001827 / ENSG00000110195 / ENSRNOG00000019902
FOXC1 / Q12948 / ENSMUSG00000050295 / ENSG00000054598 / ENSRNOG00000017800
FOXK2 / Q01167 / ENSMUSG00000039275 / ENSG00000141568 / ENSRNOG00000036663
FOXO1 / Q12778 / ENSMUSG00000044167 / ENSG00000150907 / ENSRNOG00000013397
FOXO3 / O43524 / ENSMUSG00000048756 / ENSG00000118689 / ENSRNOG00000000299
FOXP1 / Q9H334 / ENSMUSG00000030067 / ENSG00000114861 / ENSRNOG00000009184
FRA10AC1 / Q70Z53 / ENSMUSG00000054237 / ENSG00000148690 / ENSRNOG00000015235
FREM1 / Q5H8C1 / ENSMUSG00000059049 / ENSG00000164946 / ENSRNOG00000022309
FRMD4A / Q9P2Q2 / ENSMUSG00000026657 / ENSG00000151474 / ENSRNOG00000018500
FRMD4B / Q9Y2L6 / ENSMUSG00000030064 / ENSG00000114541 / ENSRNOG00000007764
FRMD8 / Q9BZ67 / ENSMUSG00000024816 / ENSG00000126391 / ENSRNOG00000020881
FRS2 / Q8WU20 / ENSMUSG00000020170 / ENSG00000166225 / ENSRNOG00000005642
FSCN1 / Q16658 / ENSMUSG00000029581 / ENSG00000075618 / ENSRNOG00000056585
FSD1 / Q9BTV5 / ENSMUSG00000011589 / ENSG00000105255 / ENSRNOG00000050318
FST / P19883 / ENSMUSG00000021765 / ENSG00000134363 / ENSRNOG00000011631
FSTL1 / Q12841 / ENSMUSG00000022816 / ENSG00000163430 / ENSRNOG00000002746
FTH1 / P02794 / ENSMUSG00000024661 / ENSG00000167996 / ENSRNOG00000022619
FTO / Q9C0B1 / ENSMUSG00000055932 / ENSG00000140718 / ENSRNOG00000011728
FTSJ3 / Q8IY81 / ENSMUSG00000020706 / ENSG00000108592 / ENSRNOG00000009857
FUBP1 / Q96AE4 / ENSMUSG00000028034 / ENSG00000162613 / ENSRNOG00000043370
FUCA1 / P04066 / ENSMUSG00000028673 / ENSG00000179163 / ENSRNOG00000009325
FUCA2 / Q9BTY2 / ENSMUSG00000019810 / ENSG00000001036 / ENSRNOG00000015551
FUNDC1 / Q8IVP5 / ENSMUSG00000025040 / ENSG00000069509 / ENSRNOG00000003470
FUNDC2 / Q9BWH2 / ENSMUSG00000031198 / ENSG00000165775 / ENSRNOG00000024066
FUOM / A2VDF0 / ENSMUSG00000025466 / ENSG00000148803 / ENSRNOG00000018476
FURIN / P09958 / ENSMUSG00000030530 / ENSG00000140564 / ENSRNOG00000011352
FUS / P35637 / ENSMUSG00000030795 / ENSG00000089280 / ENSRNOG00000023360
FUT11 / Q495W5 / ENSMUSG00000039357 / ENSG00000196968 / ENSRNOG00000009274
FUT8 / Q9BYC5 / ENSMUSG00000021065 / ENSG00000033170 / ENSRNOG00000008451
FXR1 / P51114 / ENSMUSG00000027680 / ENSG00000114416 / ENSRNOG00000051480
FXR2 / P51116 / ENSMUSG00000018765 / ENSG00000129245 / ENSRNOG00000011876
FXYD2 / P54710 / ENSMUSG00000059412 / ENSG00000137731 / ENSRNOG00000016469
FXYD6 / Q9H0Q3 / ENSMUSG00000066705 / ENSG00000137726 / ENSRNOG00000016412
FXYD7 / P58549 / ENSMUSG00000036578 / ENSG00000221946 / ENSRNOG00000021067
FYB1 / O15117 / ENSMUSG00000022148 / ENSG00000082074 / ENSRNOG00000013886
FYB2 / Q5VWT5 / ENSMUSG00000078612 / ENSG00000187889 / ENSRNOG00000030938
FYCO1 / Q9BQS8 / ENSMUSG00000025241 / ENSG00000163820 / ENSRNOG00000006336
FYTTD1 / Q96QD9 / ENSMUSG00000022800 / ENSG00000122068 / ENSRNOG00000034233
G3BP2 / Q9UN86 / ENSMUSG00000029405 / ENSG00000138757 / ENSRNOG00000002433
G6PC1 / P35575 / ENSMUSG00000078650 / ENSG00000131482 / ENSRNOG00000051171
G6PC3 / Q9BUM1 / ENSMUSG00000034793 / ENSG00000141349 / ENSRNOG00000020902
Homologs
Brain
Heart
Kidney
Liver
Lung
Testis
Organs
Bins
1
2
3
4
5
Species
Human
Mouse
Rat

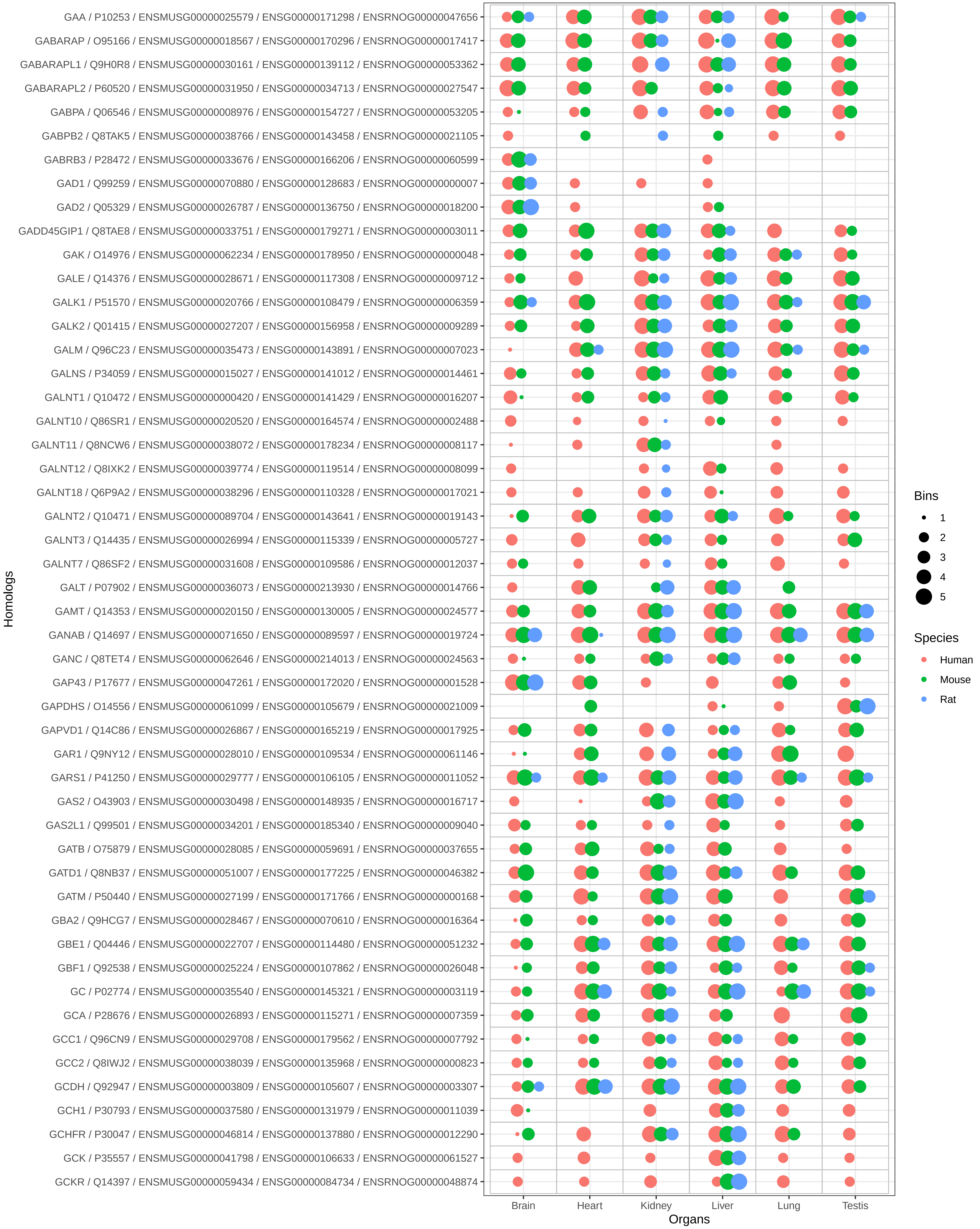
GAA / P10253 / ENSMUSG00000025579 / ENSG00000171298 / ENSRNOG00000047656
GABARAP / O95166 / ENSMUSG00000018567 / ENSG00000170296 / ENSRNOG00000017417
GABARAPL1 / Q9H0R8 / ENSMUSG00000030161 / ENSG00000139112 / ENSRNOG00000053362
GABARAPL2 / P60520 / ENSMUSG00000031950 / ENSG00000034713 / ENSRNOG00000027547
GABPA / Q06546 / ENSMUSG00000008976 / ENSG00000154727 / ENSRNOG00000053205
GABPB2 / Q8TAK5 / ENSMUSG00000038766 / ENSG00000143458 / ENSRNOG00000021105
GABRB3 / P28472 / ENSMUSG00000033676 / ENSG00000166206 / ENSRNOG00000060599
GAD1 / Q99259 / ENSMUSG00000070880 / ENSG00000128683 / ENSRNOG00000000007
GAD2 / Q05329 / ENSMUSG00000026787 / ENSG00000136750 / ENSRNOG00000018200
GADD45GIP1 / Q8TAE8 / ENSMUSG00000033751 / ENSG00000179271 / ENSRNOG00000003011
GAK / O14976 / ENSMUSG00000062234 / ENSG00000178950 / ENSRNOG00000000048
GALE / Q14376 / ENSMUSG00000028671 / ENSG00000117308 / ENSRNOG00000009712
GALK1 / P51570 / ENSMUSG00000020766 / ENSG00000108479 / ENSRNOG00000006359
GALK2 / Q01415 / ENSMUSG00000027207 / ENSG00000156958 / ENSRNOG00000009289
GALM / Q96C23 / ENSMUSG00000035473 / ENSG00000143891 / ENSRNOG00000007023
GALNS / P34059 / ENSMUSG00000015027 / ENSG00000141012 / ENSRNOG00000014461
GALNT1 / Q10472 / ENSMUSG00000000420 / ENSG00000141429 / ENSRNOG00000016207
GALNT10 / Q86SR1 / ENSMUSG00000020520 / ENSG00000164574 / ENSRNOG00000002488
GALNT11 / Q8NCW6 / ENSMUSG00000038072 / ENSG00000178234 / ENSRNOG00000008117
GALNT12 / Q8IXK2 / ENSMUSG00000039774 / ENSG00000119514 / ENSRNOG00000008099
GALNT18 / Q6P9A2 / ENSMUSG00000038296 / ENSG00000110328 / ENSRNOG00000017021
GALNT2 / Q10471 / ENSMUSG00000089704 / ENSG00000143641 / ENSRNOG00000019143
GALNT3 / Q14435 / ENSMUSG00000026994 / ENSG00000115339 / ENSRNOG00000005727
GALNT7 / Q86SF2 / ENSMUSG00000031608 / ENSG00000109586 / ENSRNOG00000012037
GALT / P07902 / ENSMUSG00000036073 / ENSG00000213930 / ENSRNOG00000014766
GAMT / Q14353 / ENSMUSG00000020150 / ENSG00000130005 / ENSRNOG00000024577
GANAB / Q14697 / ENSMUSG00000071650 / ENSG00000089597 / ENSRNOG00000019724
GANC / Q8TET4 / ENSMUSG00000062646 / ENSG00000214013 / ENSRNOG00000024563
GAP43 / P17677 / ENSMUSG00000047261 / ENSG00000172020 / ENSRNOG00000001528
GAPDHS / O14556 / ENSMUSG00000061099 / ENSG00000105679 / ENSRNOG00000021009
GAPVD1 / Q14C86 / ENSMUSG00000026867 / ENSG00000165219 / ENSRNOG00000017925
GAR1 / Q9NY12 / ENSMUSG00000028010 / ENSG00000109534 / ENSRNOG00000061146
GARS1 / P41250 / ENSMUSG00000029777 / ENSG00000106105 / ENSRNOG00000011052
GAS2 / O43903 / ENSMUSG00000030498 / ENSG00000148935 / ENSRNOG00000016717
GAS2L1 / Q99501 / ENSMUSG00000034201 / ENSG00000185340 / ENSRNOG00000009040
GATB / O75879 / ENSMUSG00000028085 / ENSG00000059691 / ENSRNOG00000037655
GATD1 / Q8NB37 / ENSMUSG00000051007 / ENSG00000177225 / ENSRNOG00000046382
GATM / P50440 / ENSMUSG00000027199 / ENSG00000171766 / ENSRNOG00000000168
GBA2 / Q9HCG7 / ENSMUSG00000028467 / ENSG00000070610 / ENSRNOG00000016364
GBE1 / Q04446 / ENSMUSG00000022707 / ENSG00000114480 / ENSRNOG00000051232
GBF1 / Q92538 / ENSMUSG00000025224 / ENSG00000107862 / ENSRNOG00000026048
GC / P02774 / ENSMUSG00000035540 / ENSG00000145321 / ENSRNOG00000003119
GCA / P28676 / ENSMUSG00000026893 / ENSG00000115271 / ENSRNOG00000007359
GCC1 / Q96CN9 / ENSMUSG00000029708 / ENSG00000179562 / ENSRNOG00000007792
GCC2 / Q8IWJ2 / ENSMUSG00000038039 / ENSG00000135968 / ENSRNOG00000000823
GCDH / Q92947 / ENSMUSG00000003809 / ENSG00000105607 / ENSRNOG00000003307
GCH1 / P30793 / ENSMUSG00000037580 / ENSG00000131979 / ENSRNOG00000011039
GCHFR / P30047 / ENSMUSG00000046814 / ENSG00000137880 / ENSRNOG00000012290
GCK / P35557 / ENSMUSG00000041798 / ENSG00000106633 / ENSRNOG00000061527
GCKR / Q14397 / ENSMUSG00000059434 / ENSG00000084734 / ENSRNOG00000048874
Homologs
Brain
Heart
Kidney
Liver
Lung
Testis
Organs
Bins
1
2
3
4
5
Species
Human
Mouse
Rat

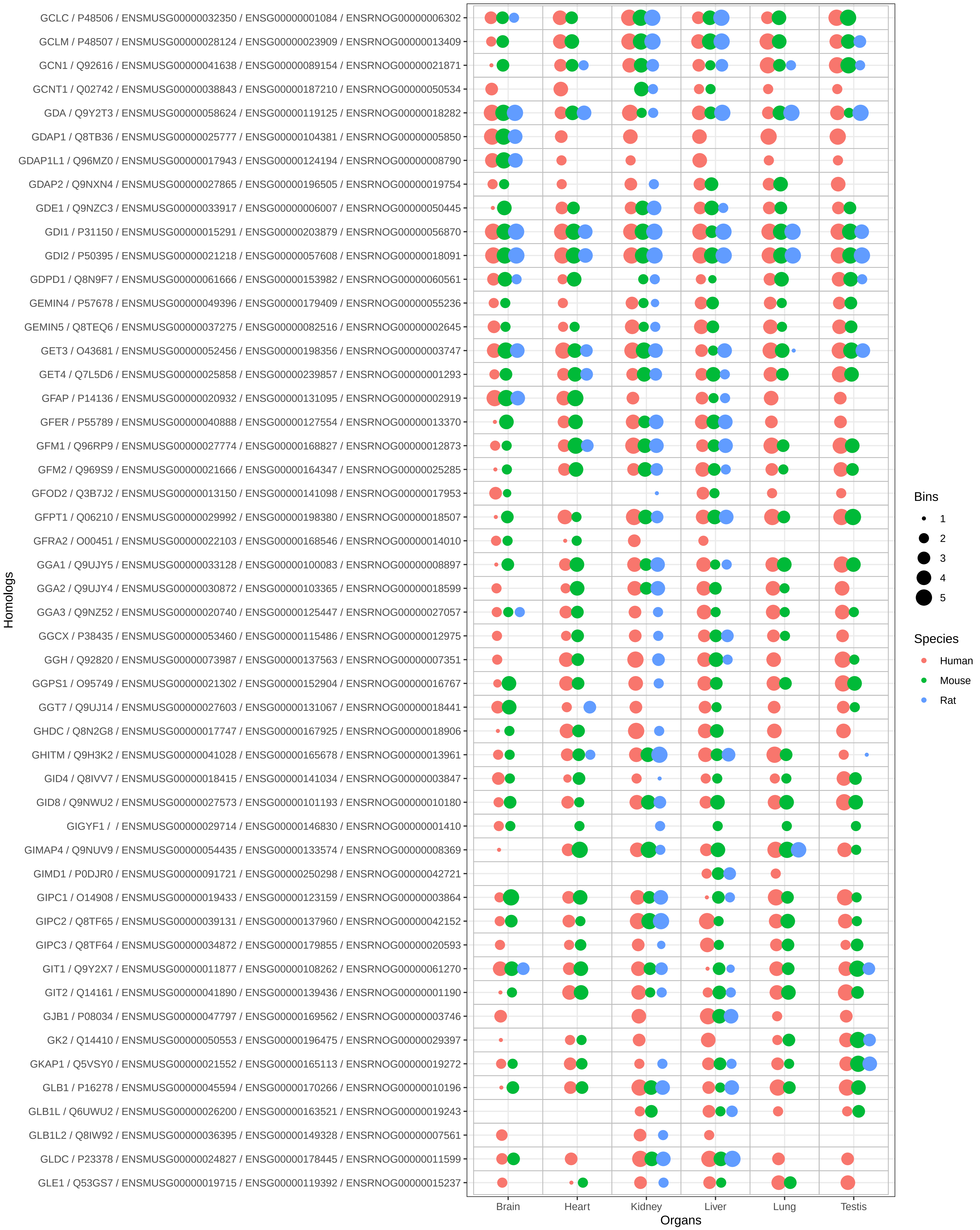
GCLC / P48506 / ENSMUSG00000032350 / ENSG00000001084 / ENSRNOG00000006302
GCLM / P48507 / ENSMUSG00000028124 / ENSG00000023909 / ENSRNOG00000013409
GCN1 / Q92616 / ENSMUSG00000041638 / ENSG00000089154 / ENSRNOG00000021871
GCNT1 / Q02742 / ENSMUSG00000038843 / ENSG00000187210 / ENSRNOG00000050534
GDA / Q9Y2T3 / ENSMUSG00000058624 / ENSG00000119125 / ENSRNOG00000018282
GDAP1 / Q8TB36 / ENSMUSG00000025777 / ENSG00000104381 / ENSRNOG00000005850
GDAP1L1 / Q96MZ0 / ENSMUSG00000017943 / ENSG00000124194 / ENSRNOG00000008790
GDAP2 / Q9NXN4 / ENSMUSG00000027865 / ENSG00000196505 / ENSRNOG00000019754
GDE1 / Q9NZC3 / ENSMUSG00000033917 / ENSG00000006007 / ENSRNOG00000050445
GDI1 / P31150 / ENSMUSG00000015291 / ENSG00000203879 / ENSRNOG00000056870
GDI2 / P50395 / ENSMUSG00000021218 / ENSG00000057608 / ENSRNOG00000018091
GDPD1 / Q8N9F7 / ENSMUSG00000061666 / ENSG00000153982 / ENSRNOG00000060561
GEMIN4 / P57678 / ENSMUSG00000049396 / ENSG00000179409 / ENSRNOG00000055236
GEMIN5 / Q8TEQ6 / ENSMUSG00000037275 / ENSG00000082516 / ENSRNOG00000002645
GET3 / O43681 / ENSMUSG00000052456 / ENSG00000198356 / ENSRNOG00000003747
GET4 / Q7L5D6 / ENSMUSG00000025858 / ENSG00000239857 / ENSRNOG00000001293
GFAP / P14136 / ENSMUSG00000020932 / ENSG00000131095 / ENSRNOG00000002919
GFER / P55789 / ENSMUSG00000040888 / ENSG00000127554 / ENSRNOG00000013370
GFM1 / Q96RP9 / ENSMUSG00000027774 / ENSG00000168827 / ENSRNOG00000012873
GFM2 / Q969S9 / ENSMUSG00000021666 / ENSG00000164347 / ENSRNOG00000025285
GFOD2 / Q3B7J2 / ENSMUSG00000013150 / ENSG00000141098 / ENSRNOG00000017953
GFPT1 / Q06210 / ENSMUSG00000029992 / ENSG00000198380 / ENSRNOG00000018507
GFRA2 / O00451 / ENSMUSG00000022103 / ENSG00000168546 / ENSRNOG00000014010
GGA1 / Q9UJY5 / ENSMUSG00000033128 / ENSG00000100083 / ENSRNOG00000008897
GGA2 / Q9UJY4 / ENSMUSG00000030872 / ENSG00000103365 / ENSRNOG00000018599
GGA3 / Q9NZ52 / ENSMUSG00000020740 / ENSG00000125447 / ENSRNOG00000027057
GGCX / P38435 / ENSMUSG00000053460 / ENSG00000115486 / ENSRNOG00000012975
GGH / Q92820 / ENSMUSG00000073987 / ENSG00000137563 / ENSRNOG00000007351
GGPS1 / O95749 / ENSMUSG00000021302 / ENSG00000152904 / ENSRNOG00000016767
GGT7 / Q9UJ14 / ENSMUSG00000027603 / ENSG00000131067 / ENSRNOG00000018441
GHDC / Q8N2G8 / ENSMUSG00000017747 / ENSG00000167925 / ENSRNOG00000018906
GHITM / Q9H3K2 / ENSMUSG00000041028 / ENSG00000165678 / ENSRNOG00000013961
GID4 / Q8IVV7 / ENSMUSG00000018415 / ENSG00000141034 / ENSRNOG00000003847
GID8 / Q9NWU2 / ENSMUSG00000027573 / ENSG00000101193 / ENSRNOG00000010180
GIGYF1 / / ENSMUSG00000029714 / ENSG00000146830 / ENSRNOG00000001410
GIMAP4 / Q9NUV9 / ENSMUSG00000054435 / ENSG00000133574 / ENSRNOG00000008369
GIMD1 / P0DJR0 / ENSMUSG00000091721 / ENSG00000250298 / ENSRNOG00000042721
GIPC1 / O14908 / ENSMUSG00000019433 / ENSG00000123159 / ENSRNOG00000003864
GIPC2 / Q8TF65 / ENSMUSG00000039131 / ENSG00000137960 / ENSRNOG00000042152
GIPC3 / Q8TF64 / ENSMUSG00000034872 / ENSG00000179855 / ENSRNOG00000020593
GIT1 / Q9Y2X7 / ENSMUSG00000011877 / ENSG00000108262 / ENSRNOG00000061270
GIT2 / Q14161 / ENSMUSG00000041890 / ENSG00000139436 / ENSRNOG00000001190
GJB1 / P08034 / ENSMUSG00000047797 / ENSG00000169562 / ENSRNOG00000003746
GK2 / Q14410 / ENSMUSG00000050553 / ENSG00000196475 / ENSRNOG00000029397
GKAP1 / Q5VSY0 / ENSMUSG00000021552 / ENSG00000165113 / ENSRNOG00000019272
GLB1 / P16278 / ENSMUSG00000045594 / ENSG00000170266 / ENSRNOG00000010196
GLB1L / Q6UWU2 / ENSMUSG00000026200 / ENSG00000163521 / ENSRNOG00000019243
GLB1L2 / Q8IW92 / ENSMUSG00000036395 / ENSG00000149328 / ENSRNOG00000007561
GLDC / P23378 / ENSMUSG00000024827 / ENSG00000178445 / ENSRNOG00000011599
GLE1 / Q53GS7 / ENSMUSG00000019715 / ENSG00000119392 / ENSRNOG00000015237
Homologs
Brain
Heart
Kidney
Liver
Lung
Testis
Organs
Bins
1
2
3
4
5
Species
Human
Mouse
Rat

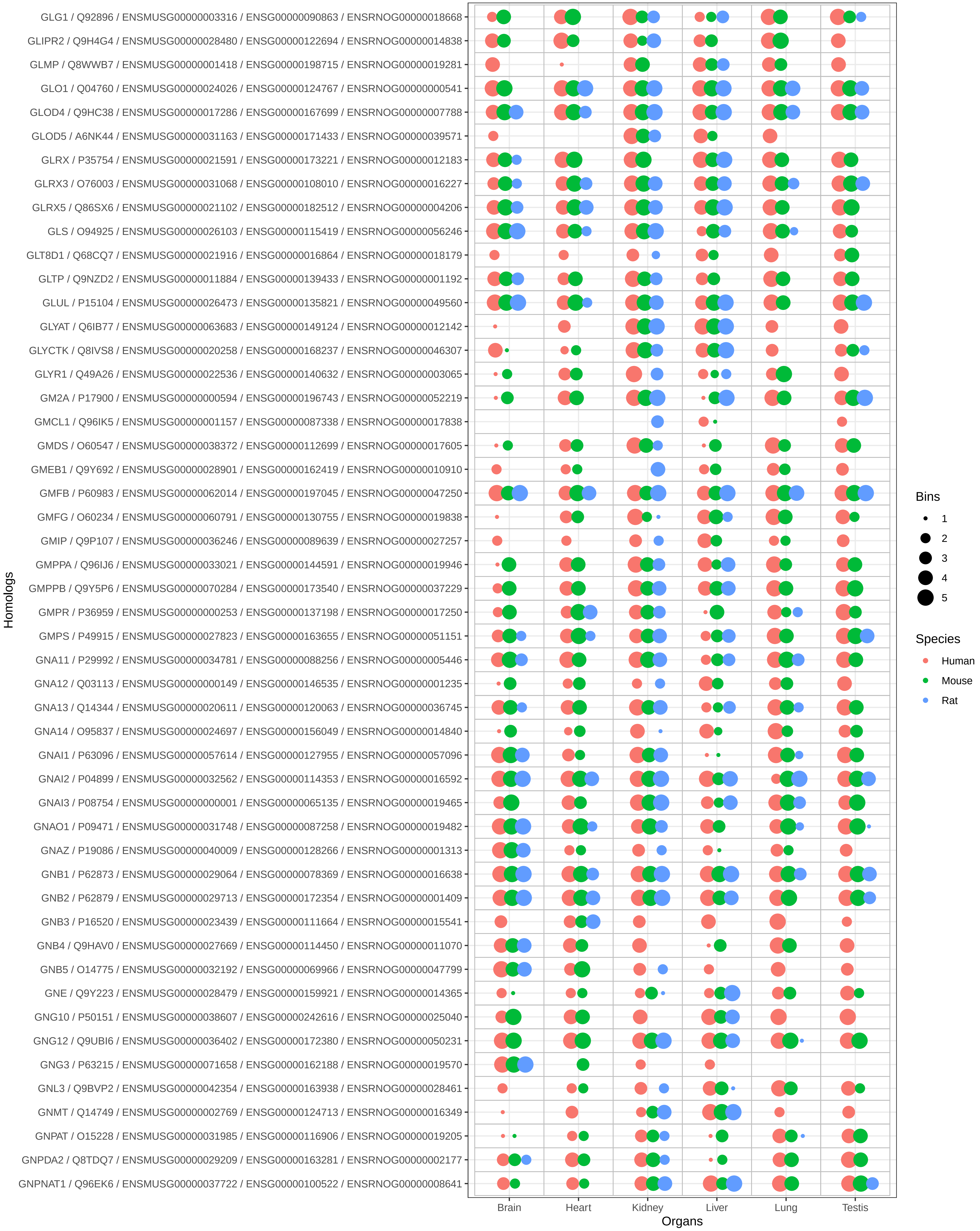

GLG1 / Q92896 / ENSMUSG00000003316 / ENSG00000090863 / ENSRNOG00000018668
GLIPR2 / Q9H4G4 / ENSMUSG00000028480 / ENSG00000122694 / ENSRNOG00000014838
GLMP / Q8WWB7 / ENSMUSG00000001418 / ENSG00000198715 / ENSRNOG00000019281
GLO1 / Q04760 / ENSMUSG00000024026 / ENSG00000124767 / ENSRNOG00000000541
GLOD4 / Q9HC38 / ENSMUSG00000017286 / ENSG00000167699 / ENSRNOG00000007788
GLOD5 / A6NK44 / ENSMUSG00000031163 / ENSG00000171433 / ENSRNOG00000039571
GLRX / P35754 / ENSMUSG00000021591 / ENSG00000173221 / ENSRNOG00000012183
GLRX3 / O76003 / ENSMUSG00000031068 / ENSG00000108010 / ENSRNOG00000016227
GLRX5 / Q86SX6 / ENSMUSG00000021102 / ENSG00000182512 / ENSRNOG00000004206
GLS / O94925 / ENSMUSG00000026103 / ENSG00000115419 / ENSRNOG00000056246
GLT8D1 / Q68CQ7 / ENSMUSG00000021916 / ENSG00000016864 / ENSRNOG00000018179
GLTP / Q9NZD2 / ENSMUSG00000011884 / ENSG00000139433 / ENSRNOG00000001192
GLUL / P15104 / ENSMUSG00000026473 / ENSG00000135821 / ENSRNOG00000049560
GLYAT / Q6IB77 / ENSMUSG00000063683 / ENSG00000149124 / ENSRNOG00000012142
GLYCTK / Q8IVS8 / ENSMUSG00000020258 / ENSG00000168237 / ENSRNOG00000046307
GLYR1 / Q49A26 / ENSMUSG00000022536 / ENSG00000140632 / ENSRNOG00000003065
GM2A / P17900 / ENSMUSG00000000594 / ENSG00000196743 / ENSRNOG00000052219
GMCL1 / Q96IK5 / ENSMUSG00000001157 / ENSG00000087338 / ENSRNOG00000017838
GMDS / O60547 / ENSMUSG00000038372 / ENSG00000112699 / ENSRNOG00000017605
GMEB1 / Q9Y692 / ENSMUSG00000028901 / ENSG00000162419 / ENSRNOG00000010910
GMFB / P60983 / ENSMUSG00000062014 / ENSG00000197045 / ENSRNOG00000047250
GMFG / O60234 / ENSMUSG00000060791 / ENSG00000130755 / ENSRNOG00000019838
GMIP / Q9P107 / ENSMUSG00000036246 / ENSG00000089639 / ENSRNOG00000027257
GMPPA / Q96IJ6 / ENSMUSG00000033021 / ENSG00000144591 / ENSRNOG00000019946
GMPPB / Q9Y5P6 / ENSMUSG00000070284 / ENSG00000173540 / ENSRNOG00000037229
GMPR / P36959 / ENSMUSG00000000253 / ENSG00000137198 / ENSRNOG00000017250
GMPS / P49915 / ENSMUSG00000027823 / ENSG00000163655 / ENSRNOG00000051151
GNA11 / P29992 / ENSMUSG00000034781 / ENSG00000088256 / ENSRNOG00000005446
GNA12 / Q03113 / ENSMUSG00000000149 / ENSG00000146535 / ENSRNOG00000001235
GNA13 / Q14344 / ENSMUSG00000020611 / ENSG00000120063 / ENSRNOG00000036745
GNA14 / O95837 / ENSMUSG00000024697 / ENSG00000156049 / ENSRNOG00000014840
GNAI1 / P63096 / ENSMUSG00000057614 / ENSG00000127955 / ENSRNOG00000057096
GNAI2 / P04899 / ENSMUSG00000032562 / ENSG00000114353 / ENSRNOG00000016592
GNAI3 / P08754 / ENSMUSG00000000001 / ENSG00000065135 / ENSRNOG00000019465
GNAO1 / P09471 / ENSMUSG00000031748 / ENSG00000087258 / ENSRNOG00000019482
GNAZ / P19086 / ENSMUSG00000040009 / ENSG00000128266 / ENSRNOG00000001313
GNB1 / P62873 / ENSMUSG00000029064 / ENSG00000078369 / ENSRNOG00000016638
GNB2 / P62879 / ENSMUSG00000029713 / ENSG00000172354 / ENSRNOG00000001409
GNB3 / P16520 / ENSMUSG00000023439 / ENSG00000111664 / ENSRNOG00000015541
GNB4 / Q9HAV0 / ENSMUSG00000027669 / ENSG00000114450 / ENSRNOG00000011070
GNB5 / O14775 / ENSMUSG00000032192 / ENSG00000069966 / ENSRNOG00000047799
GNE / Q9Y223 / ENSMUSG00000028479 / ENSG00000159921 / ENSRNOG00000014365
GNG10 / P50151 / ENSMUSG00000038607 / ENSG00000242616 / ENSRNOG00000025040
GNG12 / Q9UBI6 / ENSMUSG00000036402 / ENSG00000172380 / ENSRNOG00000050231
GNG3 / P63215 / ENSMUSG00000071658 / ENSG00000162188 / ENSRNOG00000019570
GNL3 / Q9BVP2 / ENSMUSG00000042354 / ENSG00000163938 / ENSRNOG00000028461
GNMT / Q14749 / ENSMUSG00000002769 / ENSG00000124713 / ENSRNOG00000016349
GNPAT / O15228 / ENSMUSG00000031985 / ENSG00000116906 / ENSRNOG00000019205
GNPDA2 / Q8TDQ7 / ENSMUSG00000029209 / ENSG00000163281 / ENSRNOG00000002177
GNPNAT1 / Q96EK6 / ENSMUSG00000037722 / ENSG00000100522 / ENSRNOG00000008641
Homologs
Brain
Heart
Kidney
Liver
Lung
Testis
Organs
Bins
1
2
3
4
5
Species
Human
Mouse
Rat

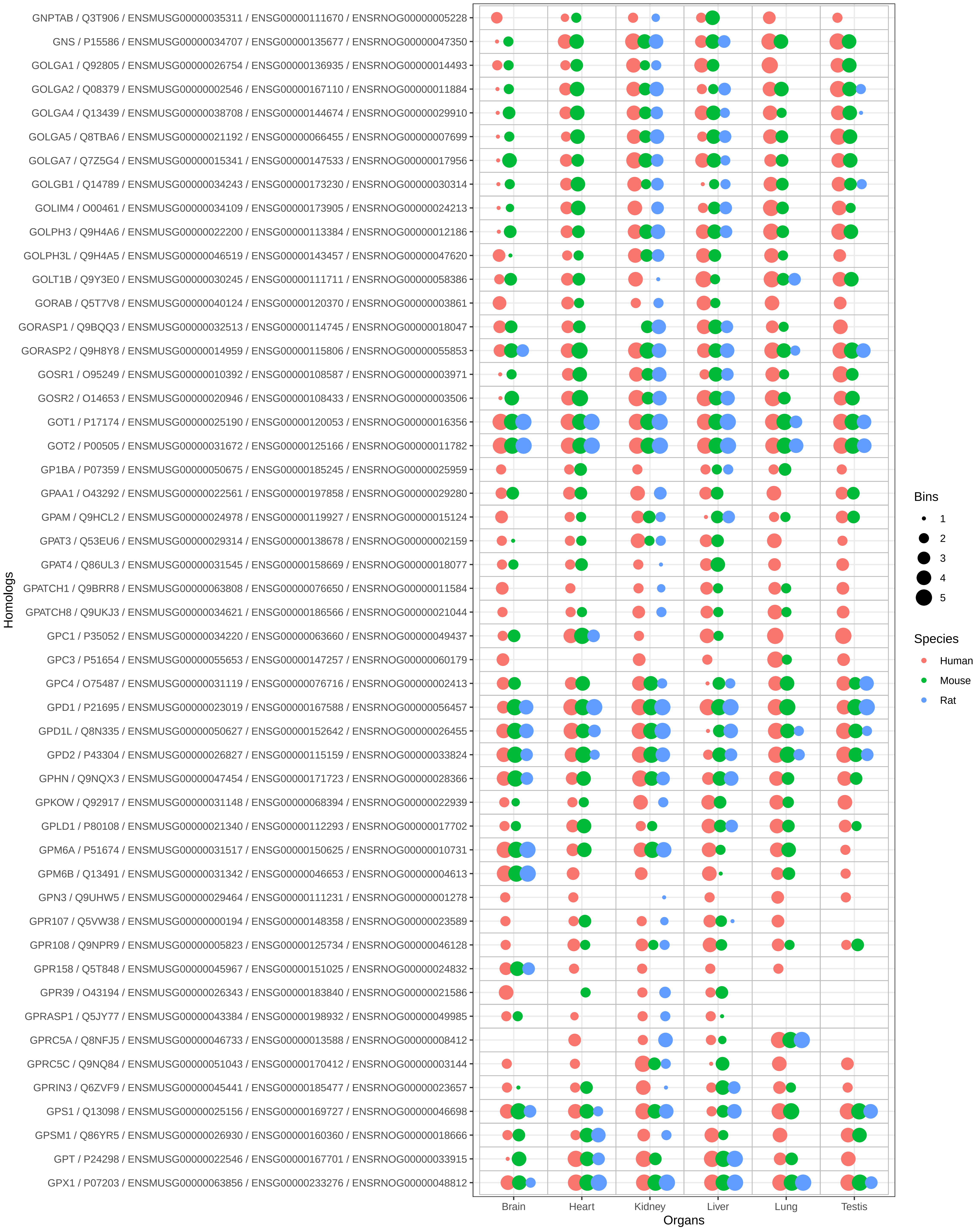
GNPTAB / Q3T906 / ENSMUSG00000035311 / ENSG00000111670 / ENSRNOG00000005228
GNS / P15586 / ENSMUSG00000034707 / ENSG00000135677 / ENSRNOG00000047350
GOLGA1 / Q92805 / ENSMUSG00000026754 / ENSG00000136935 / ENSRNOG00000014493
GOLGA2 / Q08379 / ENSMUSG00000002546 / ENSG00000167110 / ENSRNOG00000011884
GOLGA4 / Q13439 / ENSMUSG00000038708 / ENSG00000144674 / ENSRNOG00000029910
GOLGA5 / Q8TBA6 / ENSMUSG00000021192 / ENSG00000066455 / ENSRNOG00000007699
GOLGA7 / Q7Z5G4 / ENSMUSG00000015341 / ENSG00000147533 / ENSRNOG00000017956
GOLGB1 / Q14789 / ENSMUSG00000034243 / ENSG00000173230 / ENSRNOG00000030314
GOLIM4 / O00461 / ENSMUSG00000034109 / ENSG00000173905 / ENSRNOG00000024213
GOLPH3 / Q9H4A6 / ENSMUSG00000022200 / ENSG00000113384 / ENSRNOG00000012186
GOLPH3L / Q9H4A5 / ENSMUSG00000046519 / ENSG00000143457 / ENSRNOG00000047620
GOLT1B / Q9Y3E0 / ENSMUSG00000030245 / ENSG00000111711 / ENSRNOG00000058386
GORAB / Q5T7V8 / ENSMUSG00000040124 / ENSG00000120370 / ENSRNOG00000003861
GORASP1 / Q9BQQ3 / ENSMUSG00000032513 / ENSG00000114745 / ENSRNOG00000018047
GORASP2 / Q9H8Y8 / ENSMUSG00000014959 / ENSG00000115806 / ENSRNOG00000055853
GOSR1 / O95249 / ENSMUSG00000010392 / ENSG00000108587 / ENSRNOG00000003971
GOSR2 / O14653 / ENSMUSG00000020946 / ENSG00000108433 / ENSRNOG00000003506
GOT1 / P17174 / ENSMUSG00000025190 / ENSG00000120053 / ENSRNOG00000016356
GOT2 / P00505 / ENSMUSG00000031672 / ENSG00000125166 / ENSRNOG00000011782
GP1BA / P07359 / ENSMUSG00000050675 / ENSG00000185245 / ENSRNOG00000025959
GPAA1 / O43292 / ENSMUSG00000022561 / ENSG00000197858 / ENSRNOG00000029280
GPAM / Q9HCL2 / ENSMUSG00000024978 / ENSG00000119927 / ENSRNOG00000015124
GPAT3 / Q53EU6 / ENSMUSG00000029314 / ENSG00000138678 / ENSRNOG00000002159
GPAT4 / Q86UL3 / ENSMUSG00000031545 / ENSG00000158669 / ENSRNOG00000018077
GPATCH1 / Q9BRR8 / ENSMUSG00000063808 / ENSG00000076650 / ENSRNOG00000011584
GPATCH8 / Q9UKJ3 / ENSMUSG00000034621 / ENSG00000186566 / ENSRNOG00000021044
GPC1 / P35052 / ENSMUSG00000034220 / ENSG00000063660 / ENSRNOG00000049437
GPC3 / P51654 / ENSMUSG00000055653 / ENSG00000147257 / ENSRNOG00000060179
GPC4 / O75487 / ENSMUSG00000031119 / ENSG00000076716 / ENSRNOG00000002413
GPD1 / P21695 / ENSMUSG00000023019 / ENSG00000167588 / ENSRNOG00000056457
GPD1L / Q8N335 / ENSMUSG00000050627 / ENSG00000152642 / ENSRNOG00000026455
GPD2 / P43304 / ENSMUSG00000026827 / ENSG00000115159 / ENSRNOG00000033824
GPHN / Q9NQX3 / ENSMUSG00000047454 / ENSG00000171723 / ENSRNOG00000028366
GPKOW / Q92917 / ENSMUSG00000031148 / ENSG00000068394 / ENSRNOG00000022939
GPLD1 / P80108 / ENSMUSG00000021340 / ENSG00000112293 / ENSRNOG00000017702
GPM6A / P51674 / ENSMUSG00000031517 / ENSG00000150625 / ENSRNOG00000010731
GPM6B / Q13491 / ENSMUSG00000031342 / ENSG00000046653 / ENSRNOG00000004613
GPN3 / Q9UHW5 / ENSMUSG00000029464 / ENSG00000111231 / ENSRNOG00000001278
GPR107 / Q5VW38 / ENSMUSG00000000194 / ENSG00000148358 / ENSRNOG00000023589
GPR108 / Q9NPR9 / ENSMUSG00000005823 / ENSG00000125734 / ENSRNOG00000046128
GPR158 / Q5T848 / ENSMUSG00000045967 / ENSG00000151025 / ENSRNOG00000024832
GPR39 / O43194 / ENSMUSG00000026343 / ENSG00000183840 / ENSRNOG00000021586
GPRASP1 / Q5JY77 / ENSMUSG00000043384 / ENSG00000198932 / ENSRNOG00000049985
GPRC5A / Q8NFJ5 / ENSMUSG00000046733 / ENSG00000013588 / ENSRNOG00000008412
GPRC5C / Q9NQ84 / ENSMUSG00000051043 / ENSG00000170412 / ENSRNOG00000003144
GPRIN3 / Q6ZVF9 / ENSMUSG00000045441 / ENSG00000185477 / ENSRNOG00000023657
GPS1 / Q13098 / ENSMUSG00000025156 / ENSG00000169727 / ENSRNOG00000046698
GPSM1 / Q86YR5 / ENSMUSG00000026930 / ENSG00000160360 / ENSRNOG00000018666
GPT / P24298 / ENSMUSG00000022546 / ENSG00000167701 / ENSRNOG00000033915
GPX1 / P07203 / ENSMUSG00000063856 / ENSG00000233276 / ENSRNOG00000048812
Homologs
Brain
Heart
Kidney
Liver
Lung
Testis
Organs
Bins
1
2
3
4
5
Species
Human
Mouse
Rat

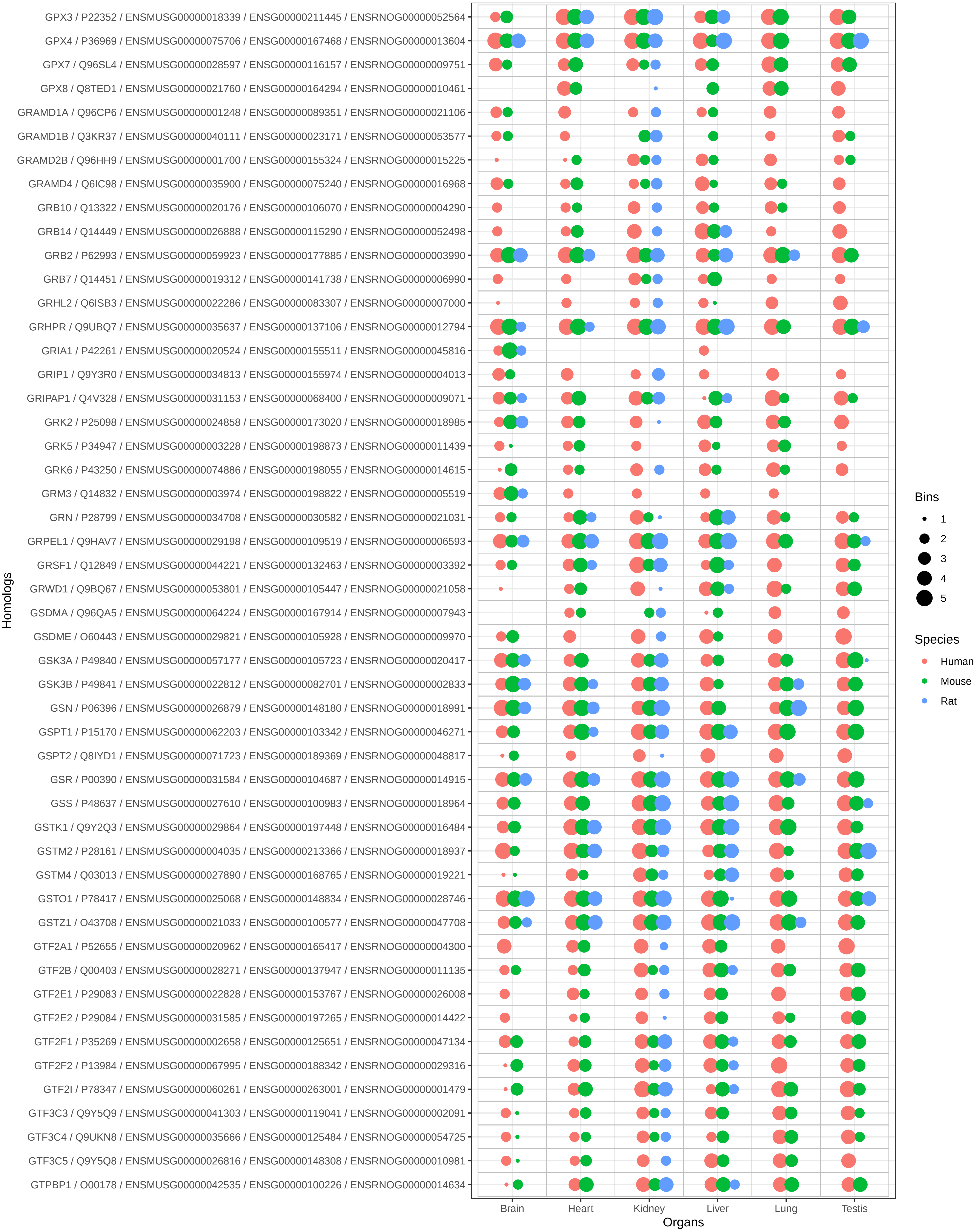

GPX3 / P22352 / ENSMUSG00000018339 / ENSG00000211445 / ENSRNOG00000052564
GPX4 / P36969 / ENSMUSG00000075706 / ENSG00000167468 / ENSRNOG00000013604
GPX7 / Q96SL4 / ENSMUSG00000028597 / ENSG00000116157 / ENSRNOG00000009751
GPX8 / Q8TED1 / ENSMUSG00000021760 / ENSG00000164294 / ENSRNOG00000010461
GRAMD1A / Q96CP6 / ENSMUSG00000001248 / ENSG00000089351 / ENSRNOG00000021106
GRAMD1B / Q3KR37 / ENSMUSG00000040111 / ENSG00000023171 / ENSRNOG00000053577
GRAMD2B / Q96HH9 / ENSMUSG00000001700 / ENSG00000155324 / ENSRNOG00000015225
GRAMD4 / Q6IC98 / ENSMUSG00000035900 / ENSG00000075240 / ENSRNOG00000016968
GRB10 / Q13322 / ENSMUSG00000020176 / ENSG00000106070 / ENSRNOG00000004290
GRB14 / Q14449 / ENSMUSG00000026888 / ENSG00000115290 / ENSRNOG00000052498
GRB2 / P62993 / ENSMUSG00000059923 / ENSG00000177885 / ENSRNOG00000003990
GRB7 / Q14451 / ENSMUSG00000019312 / ENSG00000141738 / ENSRNOG00000006990
GRHL2 / Q6ISB3 / ENSMUSG00000022286 / ENSG00000083307 / ENSRNOG00000007000
GRHPR / Q9UBQ7 / ENSMUSG00000035637 / ENSG00000137106 / ENSRNOG00000012794
GRIA1 / P42261 / ENSMUSG00000020524 / ENSG00000155511 / ENSRNOG00000045816
GRIP1 / Q9Y3R0 / ENSMUSG00000034813 / ENSG00000155974 / ENSRNOG00000004013
GRIPAP1 / Q4V328 / ENSMUSG00000031153 / ENSG00000068400 / ENSRNOG00000009071
GRK2 / P25098 / ENSMUSG00000024858 / ENSG00000173020 / ENSRNOG00000018985
GRK5 / P34947 / ENSMUSG00000003228 / ENSG00000198873 / ENSRNOG00000011439
GRK6 / P43250 / ENSMUSG00000074886 / ENSG00000198055 / ENSRNOG00000014615
GRM3 / Q14832 / ENSMUSG00000003974 / ENSG00000198822 / ENSRNOG00000005519
GRN / P28799 / ENSMUSG00000034708 / ENSG00000030582 / ENSRNOG00000021031
GRPEL1 / Q9HAV7 / ENSMUSG00000029198 / ENSG00000109519 / ENSRNOG00000006593
GRSF1 / Q12849 / ENSMUSG00000044221 / ENSG00000132463 / ENSRNOG00000003392
GRWD1 / Q9BQ67 / ENSMUSG00000053801 / ENSG00000105447 / ENSRNOG00000021058
GSDMA / Q96QA5 / ENSMUSG00000064224 / ENSG00000167914 / ENSRNOG00000007943
GSDME / O60443 / ENSMUSG00000029821 / ENSG00000105928 / ENSRNOG00000009970
GSK3A / P49840 / ENSMUSG00000057177 / ENSG00000105723 / ENSRNOG00000020417
GSK3B / P49841 / ENSMUSG00000022812 / ENSG00000082701 / ENSRNOG00000002833
GSN / P06396 / ENSMUSG00000026879 / ENSG00000148180 / ENSRNOG00000018991
GSPT1 / P15170 / ENSMUSG00000062203 / ENSG00000103342 / ENSRNOG00000046271
GSPT2 / Q8IYD1 / ENSMUSG00000071723 / ENSG00000189369 / ENSRNOG00000048817
GSR / P00390 / ENSMUSG00000031584 / ENSG00000104687 / ENSRNOG00000014915
GSS / P48637 / ENSMUSG00000027610 / ENSG00000100983 / ENSRNOG00000018964
GSTK1 / Q9Y2Q3 / ENSMUSG00000029864 / ENSG00000197448 / ENSRNOG00000016484
GSTM2 / P28161 / ENSMUSG00000004035 / ENSG00000213366 / ENSRNOG00000018937
GSTM4 / Q03013 / ENSMUSG00000027890 / ENSG00000168765 / ENSRNOG00000019221
GSTO1 / P78417 / ENSMUSG00000025068 / ENSG00000148834 / ENSRNOG00000028746
GSTZ1 / O43708 / ENSMUSG00000021033 / ENSG00000100577 / ENSRNOG00000047708
GTF2A1 / P52655 / ENSMUSG00000020962 / ENSG00000165417 / ENSRNOG00000004300
GTF2B / Q00403 / ENSMUSG00000028271 / ENSG00000137947 / ENSRNOG00000011135
GTF2E1 / P29083 / ENSMUSG00000022828 / ENSG00000153767 / ENSRNOG00000026008
GTF2E2 / P29084 / ENSMUSG00000031585 / ENSG00000197265 / ENSRNOG00000014422
GTF2F1 / P35269 / ENSMUSG00000002658 / ENSG00000125651 / ENSRNOG00000047134
GTF2F2 / P13984 / ENSMUSG00000067995 / ENSG00000188342 / ENSRNOG00000029316
GTF2I / P78347 / ENSMUSG00000060261 / ENSG00000263001 / ENSRNOG00000001479
GTF3C3 / Q9Y5Q9 / ENSMUSG00000041303 / ENSG00000119041 / ENSRNOG00000002091
GTF3C4 / Q9UKN8 / ENSMUSG00000035666 / ENSG00000125484 / ENSRNOG00000054725
GTF3C5 / Q9Y5Q8 / ENSMUSG00000026816 / ENSG00000148308 / ENSRNOG00000010981
GTPBP1 / O00178 / ENSMUSG00000042535 / ENSG00000100226 / ENSRNOG00000014634
Homologs
Brain
Heart
Kidney
Liver
Lung
Testis
Organs
Bins
1
2
3
4
5
Species
Human
Mouse
Rat

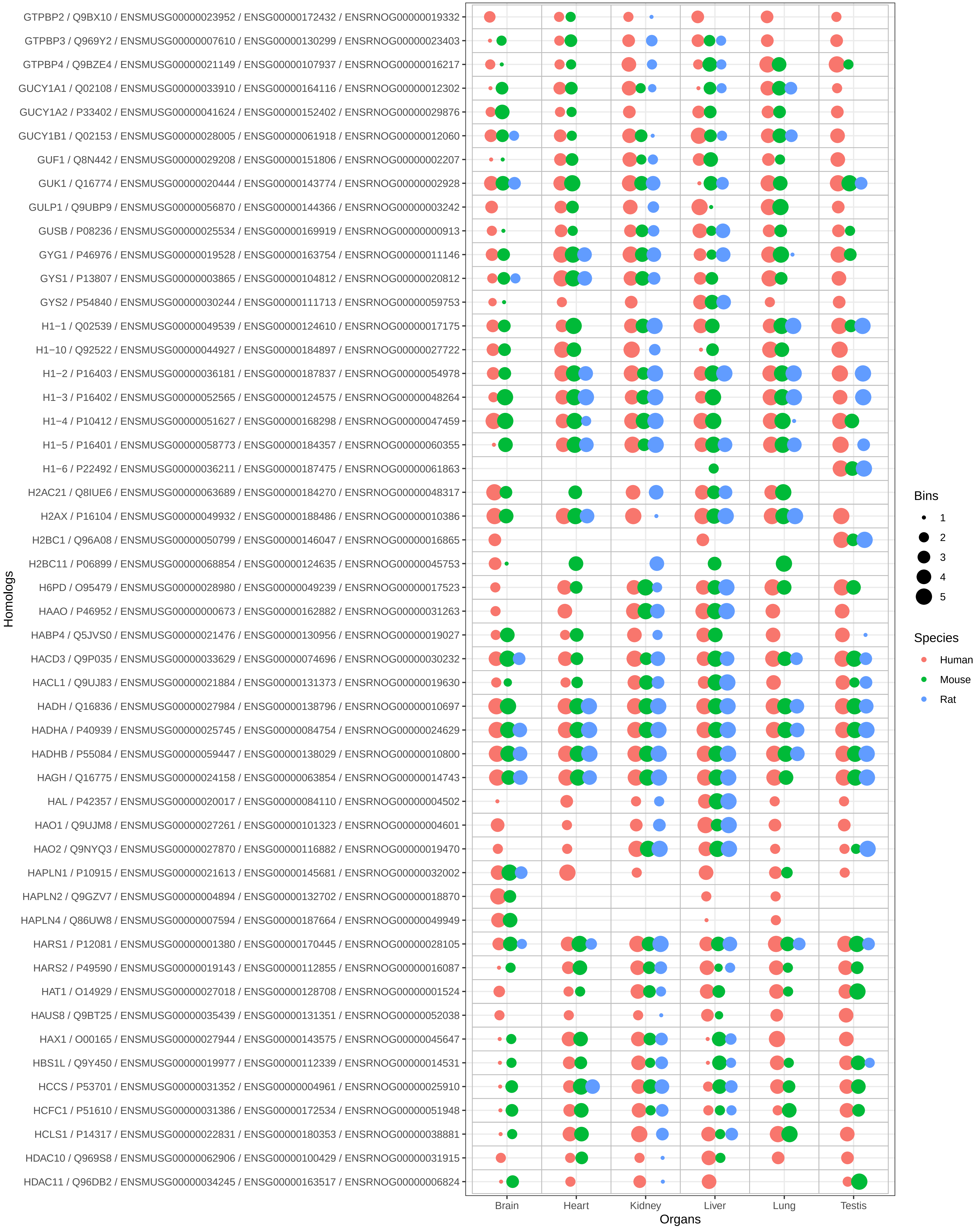
GTPBP2 / Q9BX10 / ENSMUSG00000023952 / ENSG00000172432 / ENSRNOG00000019332
GTPBP3 / Q969Y2 / ENSMUSG00000007610 / ENSG00000130299 / ENSRNOG00000023403
GTPBP4 / Q9BZE4 / ENSMUSG00000021149 / ENSG00000107937 / ENSRNOG00000016217
GUCY1A1 / Q02108 / ENSMUSG00000033910 / ENSG00000164116 / ENSRNOG00000012302
GUCY1A2 / P33402 / ENSMUSG00000041624 / ENSG00000152402 / ENSRNOG00000029876
GUCY1B1 / Q02153 / ENSMUSG00000028005 / ENSG00000061918 / ENSRNOG00000012060
GUF1 / Q8N442 / ENSMUSG00000029208 / ENSG00000151806 / ENSRNOG00000002207
GUK1 / Q16774 / ENSMUSG00000020444 / ENSG00000143774 / ENSRNOG00000002928
GULP1 / Q9UBP9 / ENSMUSG00000056870 / ENSG00000144366 / ENSRNOG00000003242
GUSB / P08236 / ENSMUSG00000025534 / ENSG00000169919 / ENSRNOG00000000913
GYG1 / P46976 / ENSMUSG00000019528 / ENSG00000163754 / ENSRNOG00000011146
GYS1 / P13807 / ENSMUSG00000003865 / ENSG00000104812 / ENSRNOG00000020812
GYS2 / P54840 / ENSMUSG00000030244 / ENSG00000111713 / ENSRNOG00000059753
H1−1 / Q02539 / ENSMUSG00000049539 / ENSG00000124610 / ENSRNOG00000017175
H1−10 / Q92522 / ENSMUSG00000044927 / ENSG00000184897 / ENSRNOG00000027722
H1−2 / P16403 / ENSMUSG00000036181 / ENSG00000187837 / ENSRNOG00000054978
H1−3 / P16402 / ENSMUSG00000052565 / ENSG00000124575 / ENSRNOG00000048264
H1−4 / P10412 / ENSMUSG00000051627 / ENSG00000168298 / ENSRNOG00000047459
H1−5 / P16401 / ENSMUSG00000058773 / ENSG00000184357 / ENSRNOG00000060355
H1−6 / P22492 / ENSMUSG00000036211 / ENSG00000187475 / ENSRNOG00000061863
H2AC21 / Q8IUE6 / ENSMUSG00000063689 / ENSG00000184270 / ENSRNOG00000048317
H2AX / P16104 / ENSMUSG00000049932 / ENSG00000188486 / ENSRNOG00000010386
H2BC1 / Q96A08 / ENSMUSG00000050799 / ENSG00000146047 / ENSRNOG00000016865
H2BC11 / P06899 / ENSMUSG00000068854 / ENSG00000124635 / ENSRNOG00000045753
H6PD / O95479 / ENSMUSG00000028980 / ENSG00000049239 / ENSRNOG00000017523
HAAO / P46952 / ENSMUSG00000000673 / ENSG00000162882 / ENSRNOG00000031263
HABP4 / Q5JVS0 / ENSMUSG00000021476 / ENSG00000130956 / ENSRNOG00000019027
HACD3 / Q9P035 / ENSMUSG00000033629 / ENSG00000074696 / ENSRNOG00000030232
HACL1 / Q9UJ83 / ENSMUSG00000021884 / ENSG00000131373 / ENSRNOG00000019630
HADH / Q16836 / ENSMUSG00000027984 / ENSG00000138796 / ENSRNOG00000010697
HADHA / P40939 / ENSMUSG00000025745 / ENSG00000084754 / ENSRNOG00000024629
HADHB / P55084 / ENSMUSG00000059447 / ENSG00000138029 / ENSRNOG00000010800
HAGH / Q16775 / ENSMUSG00000024158 / ENSG00000063854 / ENSRNOG00000014743
HAL / P42357 / ENSMUSG00000020017 / ENSG00000084110 / ENSRNOG00000004502
HAO1 / Q9UJM8 / ENSMUSG00000027261 / ENSG00000101323 / ENSRNOG00000004601
HAO2 / Q9NYQ3 / ENSMUSG00000027870 / ENSG00000116882 / ENSRNOG00000019470
HAPLN1 / P10915 / ENSMUSG00000021613 / ENSG00000145681 / ENSRNOG00000032002
HAPLN2 / Q9GZV7 / ENSMUSG00000004894 / ENSG00000132702 / ENSRNOG00000018870
HAPLN4 / Q86UW8 / ENSMUSG00000007594 / ENSG00000187664 / ENSRNOG00000049949
HARS1 / P12081 / ENSMUSG00000001380 / ENSG00000170445 / ENSRNOG00000028105
HARS2 / P49590 / ENSMUSG00000019143 / ENSG00000112855 / ENSRNOG00000016087
HAT1 / O14929 / ENSMUSG00000027018 / ENSG00000128708 / ENSRNOG00000001524
HAUS8 / Q9BT25 / ENSMUSG00000035439 / ENSG00000131351 / ENSRNOG00000052038
HAX1 / O00165 / ENSMUSG00000027944 / ENSG00000143575 / ENSRNOG00000045647
HBS1L / Q9Y450 / ENSMUSG00000019977 / ENSG00000112339 / ENSRNOG00000014531
HCCS / P53701 / ENSMUSG00000031352 / ENSG00000004961 / ENSRNOG00000025910
HCFC1 / P51610 / ENSMUSG00000031386 / ENSG00000172534 / ENSRNOG00000051948
HCLS1 / P14317 / ENSMUSG00000022831 / ENSG00000180353 / ENSRNOG00000038881
HDAC10 / Q969S8 / ENSMUSG00000062906 / ENSG00000100429 / ENSRNOG00000031915
HDAC11 / Q96DB2 / ENSMUSG00000034245 / ENSG00000163517 / ENSRNOG00000006824
Homologs
Brain
Heart
Kidney
Liver
Lung
Testis
Organs
Bins
1
2
3
4
5
Species
Human
Mouse
Rat

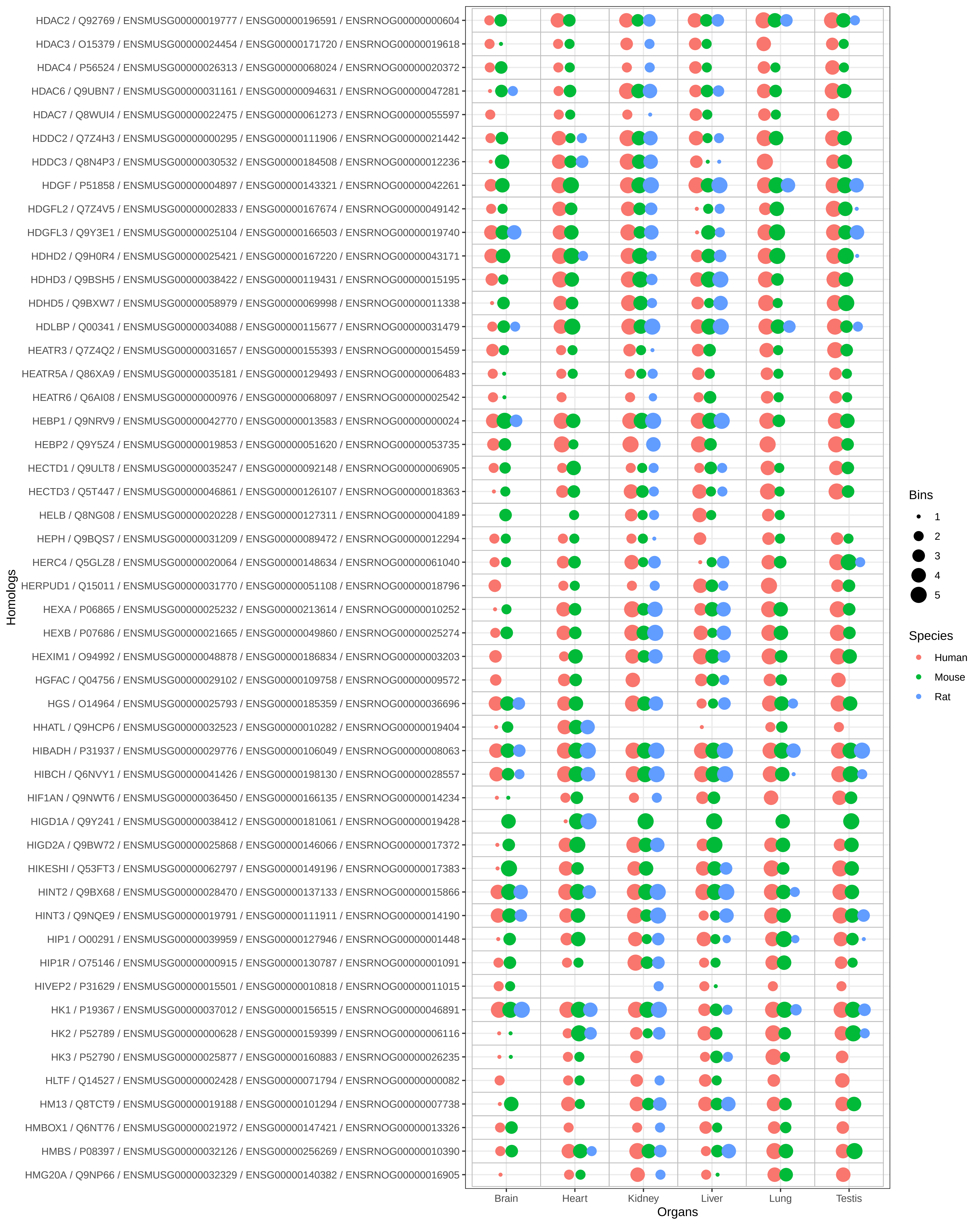
HDAC2 / Q92769 / ENSMUSG00000019777 / ENSG00000196591 / ENSRNOG00000000604
HDAC3 / O15379 / ENSMUSG00000024454 / ENSG00000171720 / ENSRNOG00000019618
HDAC4 / P56524 / ENSMUSG00000026313 / ENSG00000068024 / ENSRNOG00000020372
HDAC6 / Q9UBN7 / ENSMUSG00000031161 / ENSG00000094631 / ENSRNOG00000047281
HDAC7 / Q8WUI4 / ENSMUSG00000022475 / ENSG00000061273 / ENSRNOG00000055597
HDDC2 / Q7Z4H3 / ENSMUSG00000000295 / ENSG00000111906 / ENSRNOG00000021442
HDDC3 / Q8N4P3 / ENSMUSG00000030532 / ENSG00000184508 / ENSRNOG00000012236
HDGF / P51858 / ENSMUSG00000004897 / ENSG00000143321 / ENSRNOG00000042261
HDGFL2 / Q7Z4V5 / ENSMUSG00000002833 / ENSG00000167674 / ENSRNOG00000049142
HDGFL3 / Q9Y3E1 / ENSMUSG00000025104 / ENSG00000166503 / ENSRNOG00000019740
HDHD2 / Q9H0R4 / ENSMUSG00000025421 / ENSG00000167220 / ENSRNOG00000043171
HDHD3 / Q9BSH5 / ENSMUSG00000038422 / ENSG00000119431 / ENSRNOG00000015195
HDHD5 / Q9BXW7 / ENSMUSG00000058979 / ENSG00000069998 / ENSRNOG00000011338
HDLBP / Q00341 / ENSMUSG00000034088 / ENSG00000115677 / ENSRNOG00000031479
HEATR3 / Q7Z4Q2 / ENSMUSG00000031657 / ENSG00000155393 / ENSRNOG00000015459
HEATR5A / Q86XA9 / ENSMUSG00000035181 / ENSG00000129493 / ENSRNOG00000006483
HEATR6 / Q6AI08 / ENSMUSG00000000976 / ENSG00000068097 / ENSRNOG00000002542
HEBP1 / Q9NRV9 / ENSMUSG00000042770 / ENSG00000013583 / ENSRNOG00000000024
HEBP2 / Q9Y5Z4 / ENSMUSG00000019853 / ENSG00000051620 / ENSRNOG00000053735
HECTD1 / Q9ULT8 / ENSMUSG00000035247 / ENSG00000092148 / ENSRNOG00000006905
HECTD3 / Q5T447 / ENSMUSG00000046861 / ENSG00000126107 / ENSRNOG00000018363
HELB / Q8NG08 / ENSMUSG00000020228 / ENSG00000127311 / ENSRNOG00000004189
HEPH / Q9BQS7 / ENSMUSG00000031209 / ENSG00000089472 / ENSRNOG00000012294
HERC4 / Q5GLZ8 / ENSMUSG00000020064 / ENSG00000148634 / ENSRNOG00000061040
HERPUD1 / Q15011 / ENSMUSG00000031770 / ENSG00000051108 / ENSRNOG00000018796
HEXA / P06865 / ENSMUSG00000025232 / ENSG00000213614 / ENSRNOG00000010252
HEXB / P07686 / ENSMUSG00000021665 / ENSG00000049860 / ENSRNOG00000025274
HEXIM1 / O94992 / ENSMUSG00000048878 / ENSG00000186834 / ENSRNOG00000003203
HGFAC / Q04756 / ENSMUSG00000029102 / ENSG00000109758 / ENSRNOG00000009572
HGS / O14964 / ENSMUSG00000025793 / ENSG00000185359 / ENSRNOG00000036696
HHATL / Q9HCP6 / ENSMUSG00000032523 / ENSG00000010282 / ENSRNOG00000019404
HIBADH / P31937 / ENSMUSG00000029776 / ENSG00000106049 / ENSRNOG00000008063
HIBCH / Q6NVY1 / ENSMUSG00000041426 / ENSG00000198130 / ENSRNOG00000028557
HIF1AN / Q9NWT6 / ENSMUSG00000036450 / ENSG00000166135 / ENSRNOG00000014234
HIGD1A / Q9Y241 / ENSMUSG00000038412 / ENSG00000181061 / ENSRNOG00000019428
HIGD2A / Q9BW72 / ENSMUSG00000025868 / ENSG00000146066 / ENSRNOG00000017372
HIKESHI / Q53FT3 / ENSMUSG00000062797 / ENSG00000149196 / ENSRNOG00000017383
HINT2 / Q9BX68 / ENSMUSG00000028470 / ENSG00000137133 / ENSRNOG00000015866
HINT3 / Q9NQE9 / ENSMUSG00000019791 / ENSG00000111911 / ENSRNOG00000014190
HIP1 / O00291 / ENSMUSG00000039959 / ENSG00000127946 / ENSRNOG00000001448
HIP1R / O75146 / ENSMUSG00000000915 / ENSG00000130787 / ENSRNOG00000001091
HIVEP2 / P31629 / ENSMUSG00000015501 / ENSG00000010818 / ENSRNOG00000011015
HK1 / P19367 / ENSMUSG00000037012 / ENSG00000156515 / ENSRNOG00000046891
HK2 / P52789 / ENSMUSG00000000628 / ENSG00000159399 / ENSRNOG00000006116
HK3 / P52790 / ENSMUSG00000025877 / ENSG00000160883 / ENSRNOG00000026235
HLTF / Q14527 / ENSMUSG00000002428 / ENSG00000071794 / ENSRNOG00000000082
HM13 / Q8TCT9 / ENSMUSG00000019188 / ENSG00000101294 / ENSRNOG00000007738
HMBOX1 / Q6NT76 / ENSMUSG00000021972 / ENSG00000147421 / ENSRNOG00000013326
HMBS / P08397 / ENSMUSG00000032126 / ENSG00000256269 / ENSRNOG00000010390
HMG20A / Q9NP66 / ENSMUSG00000032329 / ENSG00000140382 / ENSRNOG00000016905
Homologs
Brain
Heart
Kidney
Liver
Lung
Testis
Organs
Bins
1
2
3
4
5
Species
Human
Mouse
Rat

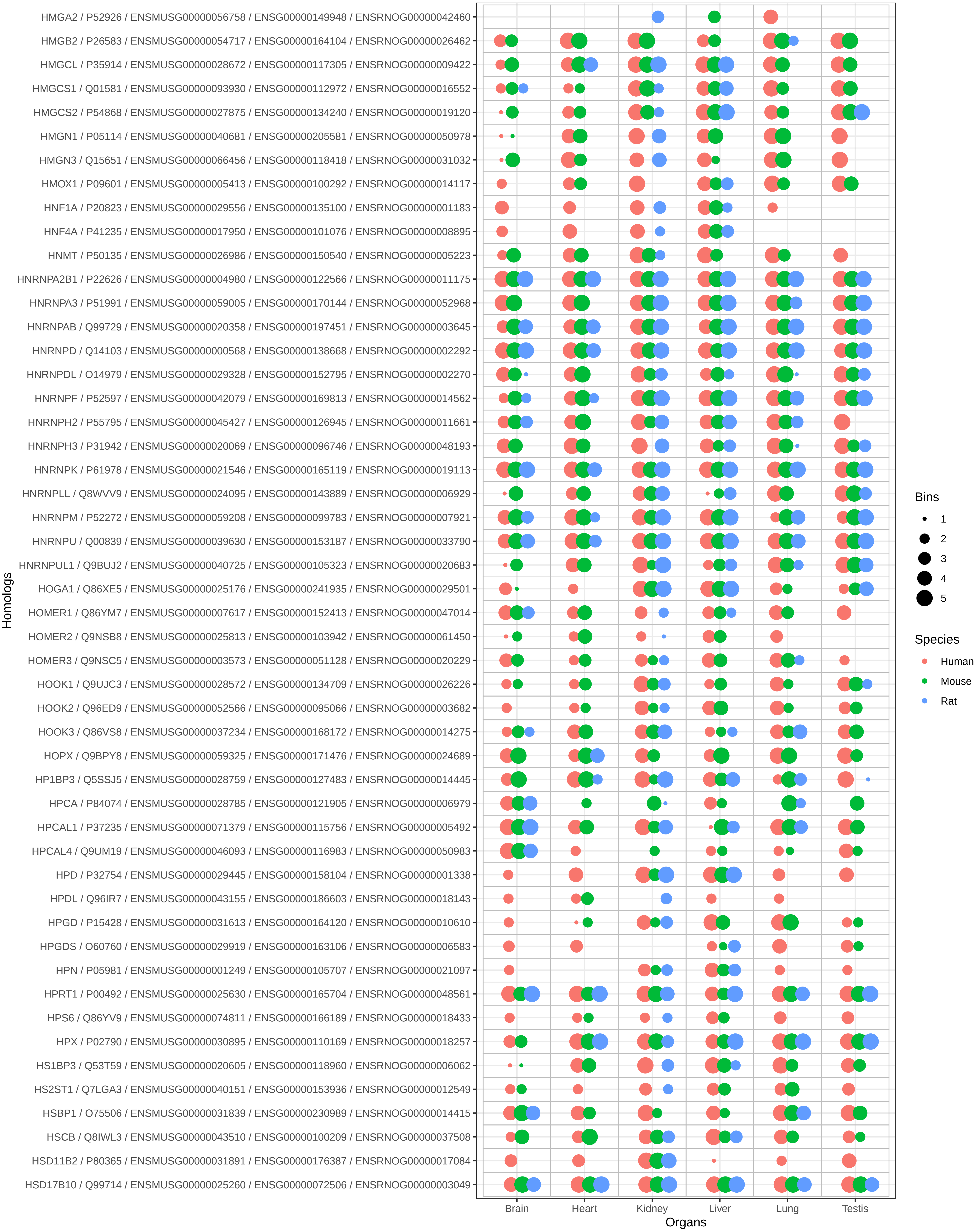
HMGA2 / P52926 / ENSMUSG00000056758 / ENSG00000149948 / ENSRNOG00000042460
HMGB2 / P26583 / ENSMUSG00000054717 / ENSG00000164104 / ENSRNOG00000026462
HMGCL / P35914 / ENSMUSG00000028672 / ENSG00000117305 / ENSRNOG00000009422
HMGCS1 / Q01581 / ENSMUSG00000093930 / ENSG00000112972 / ENSRNOG00000016552
HMGCS2 / P54868 / ENSMUSG00000027875 / ENSG00000134240 / ENSRNOG00000019120
HMGN1 / P05114 / ENSMUSG00000040681 / ENSG00000205581 / ENSRNOG00000050978
HMGN3 / Q15651 / ENSMUSG00000066456 / ENSG00000118418 / ENSRNOG00000031032
HMOX1 / P09601 / ENSMUSG00000005413 / ENSG00000100292 / ENSRNOG00000014117
HNF1A / P20823 / ENSMUSG00000029556 / ENSG00000135100 / ENSRNOG00000001183
HNF4A / P41235 / ENSMUSG00000017950 / ENSG00000101076 / ENSRNOG00000008895
HNMT / P50135 / ENSMUSG00000026986 / ENSG00000150540 / ENSRNOG00000005223
HNRNPA2B1 / P22626 / ENSMUSG00000004980 / ENSG00000122566 / ENSRNOG00000011175
HNRNPA3 / P51991 / ENSMUSG00000059005 / ENSG00000170144 / ENSRNOG00000052968
HNRNPAB / Q99729 / ENSMUSG00000020358 / ENSG00000197451 / ENSRNOG00000003645
HNRNPD / Q14103 / ENSMUSG00000000568 / ENSG00000138668 / ENSRNOG00000002292
HNRNPDL / O14979 / ENSMUSG00000029328 / ENSG00000152795 / ENSRNOG00000002270
HNRNPF / P52597 / ENSMUSG00000042079 / ENSG00000169813 / ENSRNOG00000014562
HNRNPH2 / P55795 / ENSMUSG00000045427 / ENSG00000126945 / ENSRNOG00000011661
HNRNPH3 / P31942 / ENSMUSG00000020069 / ENSG00000096746 / ENSRNOG00000048193
HNRNPK / P61978 / ENSMUSG00000021546 / ENSG00000165119 / ENSRNOG00000019113
HNRNPLL / Q8WVV9 / ENSMUSG00000024095 / ENSG00000143889 / ENSRNOG00000006929
HNRNPM / P52272 / ENSMUSG00000059208 / ENSG00000099783 / ENSRNOG00000007921
HNRNPU / Q00839 / ENSMUSG00000039630 / ENSG00000153187 / ENSRNOG00000033790
HNRNPUL1 / Q9BUJ2 / ENSMUSG00000040725 / ENSG00000105323 / ENSRNOG00000020683
HOGA1 / Q86XE5 / ENSMUSG00000025176 / ENSG00000241935 / ENSRNOG00000029501
HOMER1 / Q86YM7 / ENSMUSG00000007617 / ENSG00000152413 / ENSRNOG00000047014
HOMER2 / Q9NSB8 / ENSMUSG00000025813 / ENSG00000103942 / ENSRNOG00000061450
HOMER3 / Q9NSC5 / ENSMUSG00000003573 / ENSG00000051128 / ENSRNOG00000020229
HOOK1 / Q9UJC3 / ENSMUSG00000028572 / ENSG00000134709 / ENSRNOG00000026226
HOOK2 / Q96ED9 / ENSMUSG00000052566 / ENSG00000095066 / ENSRNOG00000003682
HOOK3 / Q86VS8 / ENSMUSG00000037234 / ENSG00000168172 / ENSRNOG00000014275
HOPX / Q9BPY8 / ENSMUSG00000059325 / ENSG00000171476 / ENSRNOG00000024689
HP1BP3 / Q5SSJ5 / ENSMUSG00000028759 / ENSG00000127483 / ENSRNOG00000014445
HPCA / P84074 / ENSMUSG00000028785 / ENSG00000121905 / ENSRNOG00000006979
HPCAL1 / P37235 / ENSMUSG00000071379 / ENSG00000115756 / ENSRNOG00000005492
HPCAL4 / Q9UM19 / ENSMUSG00000046093 / ENSG00000116983 / ENSRNOG00000050983
HPD / P32754 / ENSMUSG00000029445 / ENSG00000158104 / ENSRNOG00000001338
HPDL / Q96IR7 / ENSMUSG00000043155 / ENSG00000186603 / ENSRNOG00000018143
HPGD / P15428 / ENSMUSG00000031613 / ENSG00000164120 / ENSRNOG00000010610
HPGDS / O60760 / ENSMUSG00000029919 / ENSG00000163106 / ENSRNOG00000006583
HPN / P05981 / ENSMUSG00000001249 / ENSG00000105707 / ENSRNOG00000021097
HPRT1 / P00492 / ENSMUSG00000025630 / ENSG00000165704 / ENSRNOG00000048561
HPS6 / Q86YV9 / ENSMUSG00000074811 / ENSG00000166189 / ENSRNOG00000018433
HPX / P02790 / ENSMUSG00000030895 / ENSG00000110169 / ENSRNOG00000018257
HS1BP3 / Q53T59 / ENSMUSG00000020605 / ENSG00000118960 / ENSRNOG00000006062
HS2ST1 / Q7LGA3 / ENSMUSG00000040151 / ENSG00000153936 / ENSRNOG00000012549
HSBP1 / O75506 / ENSMUSG00000031839 / ENSG00000230989 / ENSRNOG00000014415
HSCB / Q8IWL3 / ENSMUSG00000043510 / ENSG00000100209 / ENSRNOG00000037508
HSD11B2 / P80365 / ENSMUSG00000031891 / ENSG00000176387 / ENSRNOG00000017084
HSD17B10 / Q99714 / ENSMUSG00000025260 / ENSG00000072506 / ENSRNOG00000003049
Homologs
Brain
Heart
Kidney
Liver
Lung
Testis
Organs
Bins
1
2
3
4
5
Species
Human
Mouse
Rat

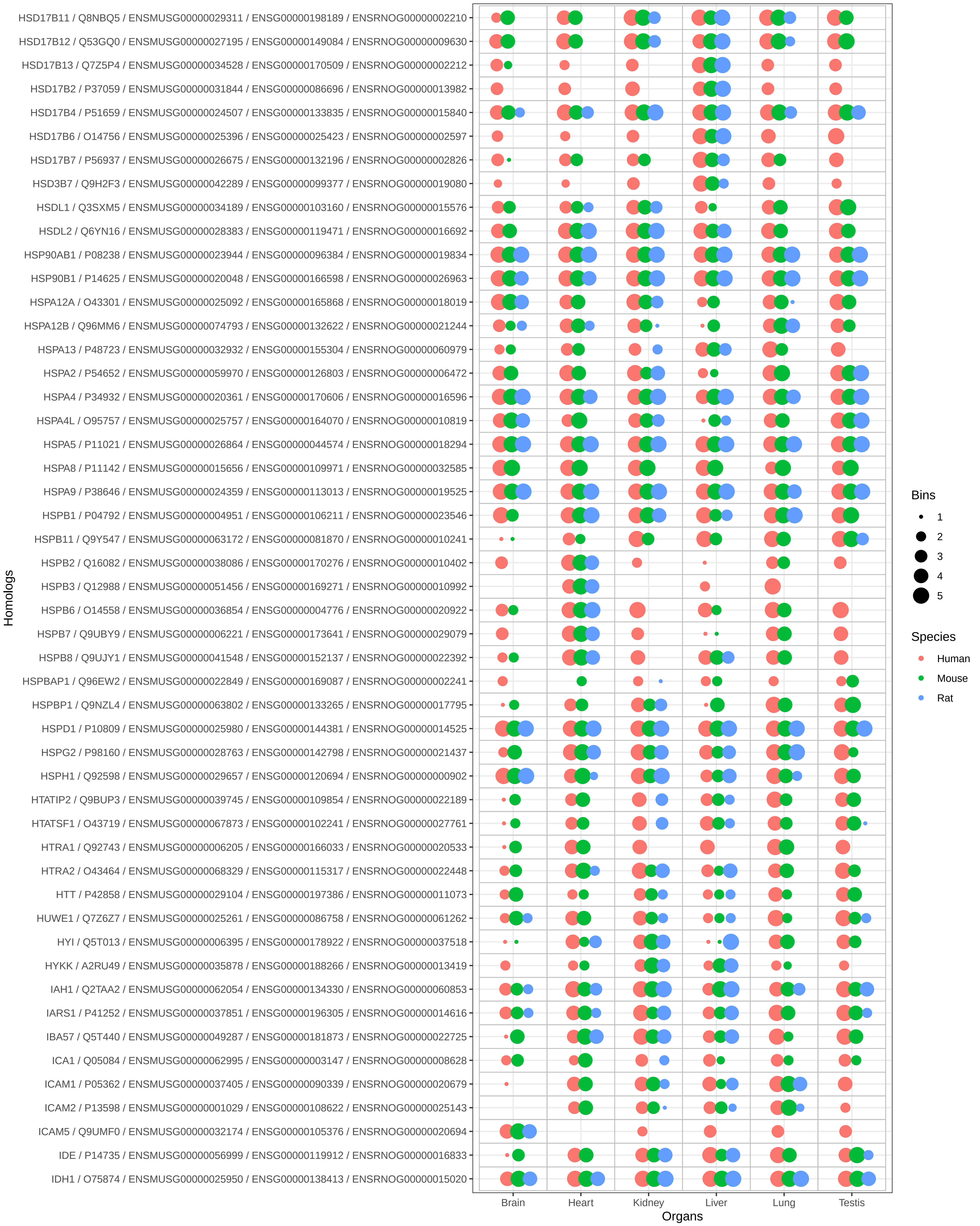

HSD17B11 / Q8NBQ5 / ENSMUSG00000029311 / ENSG00000198189 / ENSRNOG00000002210
HSD17B12 / Q53GQ0 / ENSMUSG00000027195 / ENSG00000149084 / ENSRNOG00000009630
HSD17B13 / Q7Z5P4 / ENSMUSG00000034528 / ENSG00000170509 / ENSRNOG00000002212
HSD17B2 / P37059 / ENSMUSG00000031844 / ENSG00000086696 / ENSRNOG00000013982
HSD17B4 / P51659 / ENSMUSG00000024507 / ENSG00000133835 / ENSRNOG00000015840
HSD17B6 / O14756 / ENSMUSG00000025396 / ENSG00000025423 / ENSRNOG00000002597
HSD17B7 / P56937 / ENSMUSG00000026675 / ENSG00000132196 / ENSRNOG00000002826
HSD3B7 / Q9H2F3 / ENSMUSG00000042289 / ENSG00000099377 / ENSRNOG00000019080
HSDL1 / Q3SXM5 / ENSMUSG00000034189 / ENSG00000103160 / ENSRNOG00000015576
HSDL2 / Q6YN16 / ENSMUSG00000028383 / ENSG00000119471 / ENSRNOG00000016692
HSP90AB1 / P08238 / ENSMUSG00000023944 / ENSG00000096384 / ENSRNOG00000019834
HSP90B1 / P14625 / ENSMUSG00000020048 / ENSG00000166598 / ENSRNOG00000026963
HSPA12A / O43301 / ENSMUSG00000025092 / ENSG00000165868 / ENSRNOG00000018019
HSPA12B / Q96MM6 / ENSMUSG00000074793 / ENSG00000132622 / ENSRNOG00000021244
HSPA13 / P48723 / ENSMUSG00000032932 / ENSG00000155304 / ENSRNOG00000060979
HSPA2 / P54652 / ENSMUSG00000059970 / ENSG00000126803 / ENSRNOG00000006472
HSPA4 / P34932 / ENSMUSG00000020361 / ENSG00000170606 / ENSRNOG00000016596
HSPA4L / O95757 / ENSMUSG00000025757 / ENSG00000164070 / ENSRNOG00000010819
HSPA5 / P11021 / ENSMUSG00000026864 / ENSG00000044574 / ENSRNOG00000018294
HSPA8 / P11142 / ENSMUSG00000015656 / ENSG00000109971 / ENSRNOG00000032585
HSPA9 / P38646 / ENSMUSG00000024359 / ENSG00000113013 / ENSRNOG00000019525
HSPB1 / P04792 / ENSMUSG00000004951 / ENSG00000106211 / ENSRNOG00000023546
HSPB11 / Q9Y547 / ENSMUSG00000063172 / ENSG00000081870 / ENSRNOG00000010241
HSPB2 / Q16082 / ENSMUSG00000038086 / ENSG00000170276 / ENSRNOG00000010402
HSPB3 / Q12988 / ENSMUSG00000051456 / ENSG00000169271 / ENSRNOG00000010992
HSPB6 / O14558 / ENSMUSG00000036854 / ENSG00000004776 / ENSRNOG00000020922
HSPB7 / Q9UBY9 / ENSMUSG00000006221 / ENSG00000173641 / ENSRNOG00000029079
HSPB8 / Q9UJY1 / ENSMUSG00000041548 / ENSG00000152137 / ENSRNOG00000022392
HSPBAP1 / Q96EW2 / ENSMUSG00000022849 / ENSG00000169087 / ENSRNOG00000002241
HSPBP1 / Q9NZL4 / ENSMUSG00000063802 / ENSG00000133265 / ENSRNOG00000017795
HSPD1 / P10809 / ENSMUSG00000025980 / ENSG00000144381 / ENSRNOG00000014525
HSPG2 / P98160 / ENSMUSG00000028763 / ENSG00000142798 / ENSRNOG00000021437
HSPH1 / Q92598 / ENSMUSG00000029657 / ENSG00000120694 / ENSRNOG00000000902
HTATIP2 / Q9BUP3 / ENSMUSG00000039745 / ENSG00000109854 / ENSRNOG00000022189
HTATSF1 / O43719 / ENSMUSG00000067873 / ENSG00000102241 / ENSRNOG00000027761
HTRA1 / Q92743 / ENSMUSG00000006205 / ENSG00000166033 / ENSRNOG00000020533
HTRA2 / O43464 / ENSMUSG00000068329 / ENSG00000115317 / ENSRNOG00000022448
HTT / P42858 / ENSMUSG00000029104 / ENSG00000197386 / ENSRNOG00000011073
HUWE1 / Q7Z6Z7 / ENSMUSG00000025261 / ENSG00000086758 / ENSRNOG00000061262
HYI / Q5T013 / ENSMUSG00000006395 / ENSG00000178922 / ENSRNOG00000037518
HYKK / A2RU49 / ENSMUSG00000035878 / ENSG00000188266 / ENSRNOG00000013419
IAH1 / Q2TAA2 / ENSMUSG00000062054 / ENSG00000134330 / ENSRNOG00000060853
IARS1 / P41252 / ENSMUSG00000037851 / ENSG00000196305 / ENSRNOG00000014616
IBA57 / Q5T440 / ENSMUSG00000049287 / ENSG00000181873 / ENSRNOG00000022725
ICA1 / Q05084 / ENSMUSG00000062995 / ENSG00000003147 / ENSRNOG00000008628
ICAM1 / P05362 / ENSMUSG00000037405 / ENSG00000090339 / ENSRNOG00000020679
ICAM2 / P13598 / ENSMUSG00000001029 / ENSG00000108622 / ENSRNOG00000025143
ICAM5 / Q9UMF0 / ENSMUSG00000032174 / ENSG00000105376 / ENSRNOG00000020694
IDE / P14735 / ENSMUSG00000056999 / ENSG00000119912 / ENSRNOG00000016833
IDH1 / O75874 / ENSMUSG00000025950 / ENSG00000138413 / ENSRNOG00000015020
Homologs
Brain
Heart
Kidney
Liver
Lung
Testis
Organs
Bins
1
2
3
4
5
Species
Human
Mouse
Rat

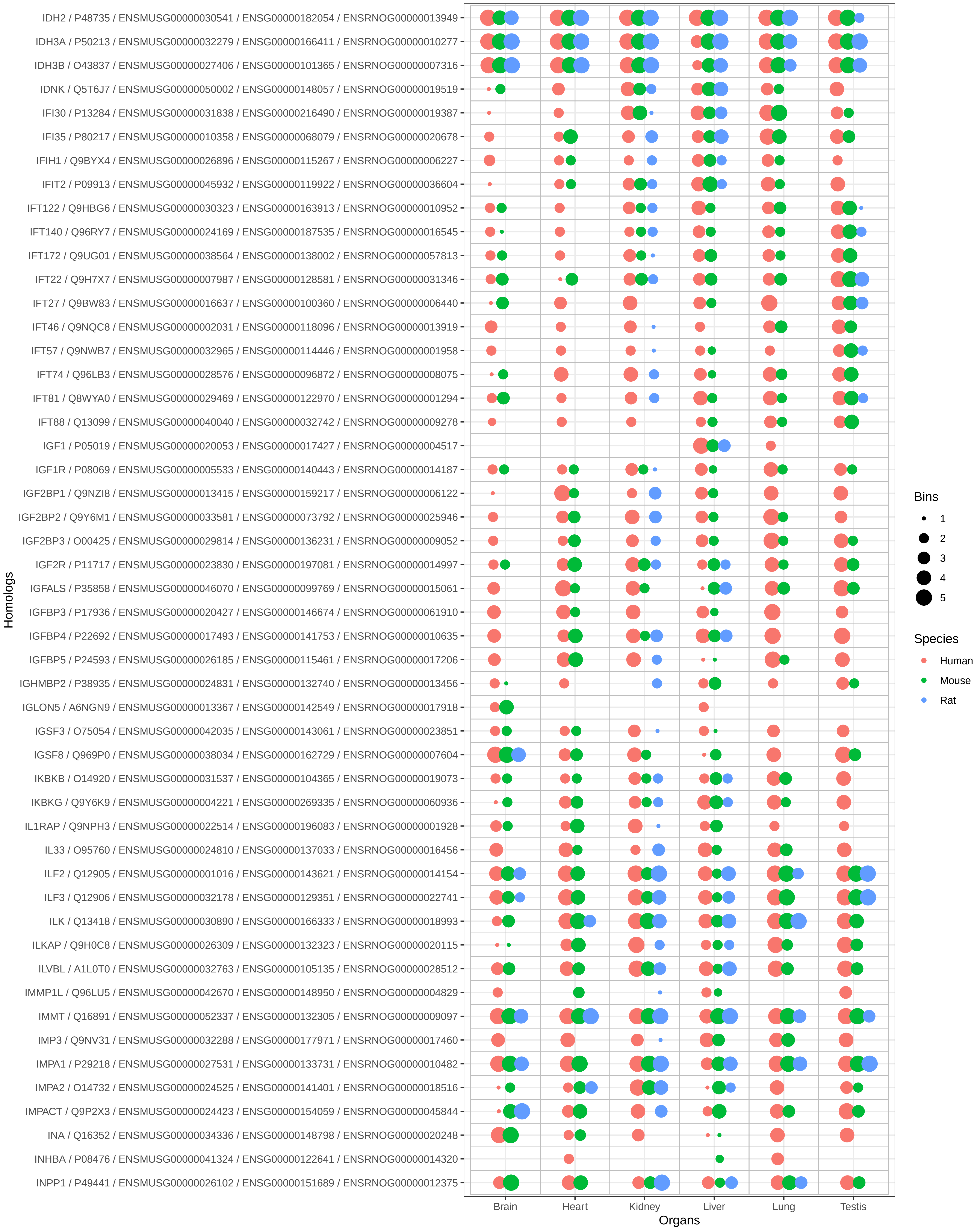
IDH2 / P48735 / ENSMUSG00000030541 / ENSG00000182054 / ENSRNOG00000013949
IDH3A / P50213 / ENSMUSG00000032279 / ENSG00000166411 / ENSRNOG00000010277
IDH3B / O43837 / ENSMUSG00000027406 / ENSG00000101365 / ENSRNOG00000007316
IDNK / Q5T6J7 / ENSMUSG00000050002 / ENSG00000148057 / ENSRNOG00000019519
IFI30 / P13284 / ENSMUSG00000031838 / ENSG00000216490 / ENSRNOG00000019387
IFI35 / P80217 / ENSMUSG00000010358 / ENSG00000068079 / ENSRNOG00000020678
IFIH1 / Q9BYX4 / ENSMUSG00000026896 / ENSG00000115267 / ENSRNOG00000006227
IFIT2 / P09913 / ENSMUSG00000045932 / ENSG00000119922 / ENSRNOG00000036604
IFT122 / Q9HBG6 / ENSMUSG00000030323 / ENSG00000163913 / ENSRNOG00000010952
IFT140 / Q96RY7 / ENSMUSG00000024169 / ENSG00000187535 / ENSRNOG00000016545
IFT172 / Q9UG01 / ENSMUSG00000038564 / ENSG00000138002 / ENSRNOG00000057813
IFT22 / Q9H7X7 / ENSMUSG00000007987 / ENSG00000128581 / ENSRNOG00000031346
IFT27 / Q9BW83 / ENSMUSG00000016637 / ENSG00000100360 / ENSRNOG00000006440
IFT46 / Q9NQC8 / ENSMUSG00000002031 / ENSG00000118096 / ENSRNOG00000013919
IFT57 / Q9NWB7 / ENSMUSG00000032965 / ENSG00000114446 / ENSRNOG00000001958
IFT74 / Q96LB3 / ENSMUSG00000028576 / ENSG00000096872 / ENSRNOG00000008075
IFT81 / Q8WYA0 / ENSMUSG00000029469 / ENSG00000122970 / ENSRNOG00000001294
IFT88 / Q13099 / ENSMUSG00000040040 / ENSG00000032742 / ENSRNOG00000009278
IGF1 / P05019 / ENSMUSG00000020053 / ENSG00000017427 / ENSRNOG00000004517
IGF1R / P08069 / ENSMUSG00000005533 / ENSG00000140443 / ENSRNOG00000014187
IGF2BP1 / Q9NZI8 / ENSMUSG00000013415 / ENSG00000159217 / ENSRNOG00000006122
IGF2BP2 / Q9Y6M1 / ENSMUSG00000033581 / ENSG00000073792 / ENSRNOG00000025946
IGF2BP3 / O00425 / ENSMUSG00000029814 / ENSG00000136231 / ENSRNOG00000009052
IGF2R / P11717 / ENSMUSG00000023830 / ENSG00000197081 / ENSRNOG00000014997
IGFALS / P35858 / ENSMUSG00000046070 / ENSG00000099769 / ENSRNOG00000015061
IGFBP3 / P17936 / ENSMUSG00000020427 / ENSG00000146674 / ENSRNOG00000061910
IGFBP4 / P22692 / ENSMUSG00000017493 / ENSG00000141753 / ENSRNOG00000010635
IGFBP5 / P24593 / ENSMUSG00000026185 / ENSG00000115461 / ENSRNOG00000017206
IGHMBP2 / P38935 / ENSMUSG00000024831 / ENSG00000132740 / ENSRNOG00000013456
IGLON5 / A6NGN9 / ENSMUSG00000013367 / ENSG00000142549 / ENSRNOG00000017918
IGSF3 / O75054 / ENSMUSG00000042035 / ENSG00000143061 / ENSRNOG00000023851
IGSF8 / Q969P0 / ENSMUSG00000038034 / ENSG00000162729 / ENSRNOG00000007604
IKBKB / O14920 / ENSMUSG00000031537 / ENSG00000104365 / ENSRNOG00000019073
IKBKG / Q9Y6K9 / ENSMUSG00000004221 / ENSG00000269335 / ENSRNOG00000060936
IL1RAP / Q9NPH3 / ENSMUSG00000022514 / ENSG00000196083 / ENSRNOG00000001928
IL33 / O95760 / ENSMUSG00000024810 / ENSG00000137033 / ENSRNOG00000016456
ILF2 / Q12905 / ENSMUSG00000001016 / ENSG00000143621 / ENSRNOG00000014154
ILF3 / Q12906 / ENSMUSG00000032178 / ENSG00000129351 / ENSRNOG00000022741
ILK / Q13418 / ENSMUSG00000030890 / ENSG00000166333 / ENSRNOG00000018993
ILKAP / Q9H0C8 / ENSMUSG00000026309 / ENSG00000132323 / ENSRNOG00000020115
ILVBL / A1L0T0 / ENSMUSG00000032763 / ENSG00000105135 / ENSRNOG00000028512
IMMP1L / Q96LU5 / ENSMUSG00000042670 / ENSG00000148950 / ENSRNOG00000004829
IMMT / Q16891 / ENSMUSG00000052337 / ENSG00000132305 / ENSRNOG00000009097
IMP3 / Q9NV31 / ENSMUSG00000032288 / ENSG00000177971 / ENSRNOG00000017460
IMPA1 / P29218 / ENSMUSG00000027531 / ENSG00000133731 / ENSRNOG00000010482
IMPA2 / O14732 / ENSMUSG00000024525 / ENSG00000141401 / ENSRNOG00000018516
IMPACT / Q9P2X3 / ENSMUSG00000024423 / ENSG00000154059 / ENSRNOG00000045844
INA / Q16352 / ENSMUSG00000034336 / ENSG00000148798 / ENSRNOG00000020248
INHBA / P08476 / ENSMUSG00000041324 / ENSG00000122641 / ENSRNOG00000014320
INPP1 / P49441 / ENSMUSG00000026102 / ENSG00000151689 / ENSRNOG00000012375
Homologs
Brain
Heart
Kidney
Liver
Lung
Testis
Organs
Bins
1
2
3
4
5
Species
Human
Mouse
Rat

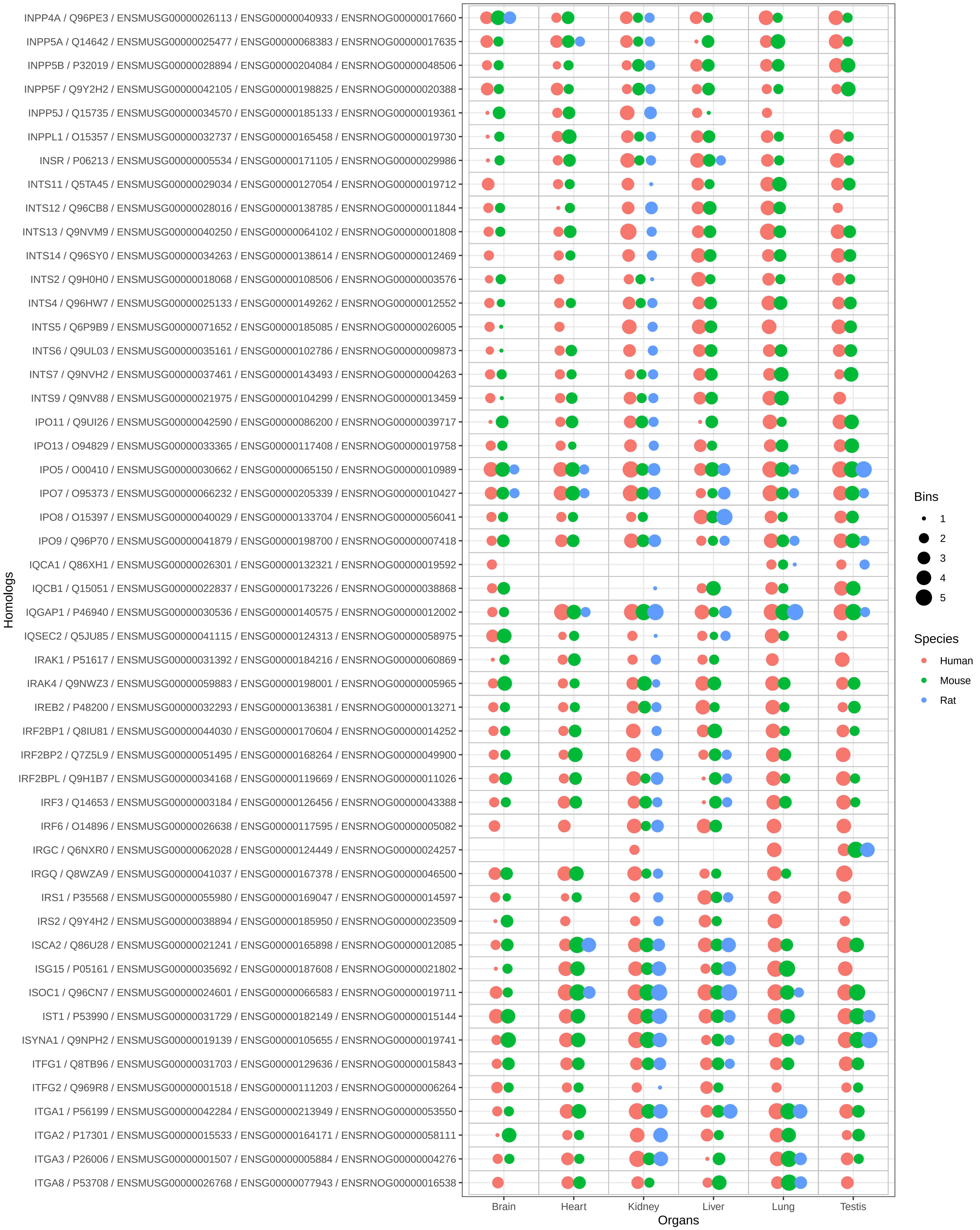
INPP4A / Q96PE3 / ENSMUSG00000026113 / ENSG00000040933 / ENSRNOG00000017660
INPP5A / Q14642 / ENSMUSG00000025477 / ENSG00000068383 / ENSRNOG00000017635
INPP5B / P32019 / ENSMUSG00000028894 / ENSG00000204084 / ENSRNOG00000048506
INPP5F / Q9Y2H2 / ENSMUSG00000042105 / ENSG00000198825 / ENSRNOG00000020388
INPP5J / Q15735 / ENSMUSG00000034570 / ENSG00000185133 / ENSRNOG00000019361
INPPL1 / O15357 / ENSMUSG00000032737 / ENSG00000165458 / ENSRNOG00000019730
INSR / P06213 / ENSMUSG00000005534 / ENSG00000171105 / ENSRNOG00000029986
INTS11 / Q5TA45 / ENSMUSG00000029034 / ENSG00000127054 / ENSRNOG00000019712
INTS12 / Q96CB8 / ENSMUSG00000028016 / ENSG00000138785 / ENSRNOG00000011844
INTS13 / Q9NVM9 / ENSMUSG00000040250 / ENSG00000064102 / ENSRNOG00000001808
INTS14 / Q96SY0 / ENSMUSG00000034263 / ENSG00000138614 / ENSRNOG00000012469
INTS2 / Q9H0H0 / ENSMUSG00000018068 / ENSG00000108506 / ENSRNOG00000003576
INTS4 / Q96HW7 / ENSMUSG00000025133 / ENSG00000149262 / ENSRNOG00000012552
INTS5 / Q6P9B9 / ENSMUSG00000071652 / ENSG00000185085 / ENSRNOG00000026005
INTS6 / Q9UL03 / ENSMUSG00000035161 / ENSG00000102786 / ENSRNOG00000009873
INTS7 / Q9NVH2 / ENSMUSG00000037461 / ENSG00000143493 / ENSRNOG00000004263
INTS9 / Q9NV88 / ENSMUSG00000021975 / ENSG00000104299 / ENSRNOG00000013459
IPO11 / Q9UI26 / ENSMUSG00000042590 / ENSG00000086200 / ENSRNOG00000039717
IPO13 / O94829 / ENSMUSG00000033365 / ENSG00000117408 / ENSRNOG00000019758
IPO5 / O00410 / ENSMUSG00000030662 / ENSG00000065150 / ENSRNOG00000010989
IPO7 / O95373 / ENSMUSG00000066232 / ENSG00000205339 / ENSRNOG00000010427
IPO8 / O15397 / ENSMUSG00000040029 / ENSG00000133704 / ENSRNOG00000056041
IPO9 / Q96P70 / ENSMUSG00000041879 / ENSG00000198700 / ENSRNOG00000007418
IQCA1 / Q86XH1 / ENSMUSG00000026301 / ENSG00000132321 / ENSRNOG00000019592
IQCB1 / Q15051 / ENSMUSG00000022837 / ENSG00000173226 / ENSRNOG00000038868
IQGAP1 / P46940 / ENSMUSG00000030536 / ENSG00000140575 / ENSRNOG00000012002
IQSEC2 / Q5JU85 / ENSMUSG00000041115 / ENSG00000124313 / ENSRNOG00000058975
IRAK1 / P51617 / ENSMUSG00000031392 / ENSG00000184216 / ENSRNOG00000060869
IRAK4 / Q9NWZ3 / ENSMUSG00000059883 / ENSG00000198001 / ENSRNOG00000005965
IREB2 / P48200 / ENSMUSG00000032293 / ENSG00000136381 / ENSRNOG00000013271
IRF2BP1 / Q8IU81 / ENSMUSG00000044030 / ENSG00000170604 / ENSRNOG00000014252
IRF2BP2 / Q7Z5L9 / ENSMUSG00000051495 / ENSG00000168264 / ENSRNOG00000049900
IRF2BPL / Q9H1B7 / ENSMUSG00000034168 / ENSG00000119669 / ENSRNOG00000011026
IRF3 / Q14653 / ENSMUSG00000003184 / ENSG00000126456 / ENSRNOG00000043388
IRF6 / O14896 / ENSMUSG00000026638 / ENSG00000117595 / ENSRNOG00000005082
IRGC / Q6NXR0 / ENSMUSG00000062028 / ENSG00000124449 / ENSRNOG00000024257
IRGQ / Q8WZA9 / ENSMUSG00000041037 / ENSG00000167378 / ENSRNOG00000046500
IRS1 / P35568 / ENSMUSG00000055980 / ENSG00000169047 / ENSRNOG00000014597
IRS2 / Q9Y4H2 / ENSMUSG00000038894 / ENSG00000185950 / ENSRNOG00000023509
ISCA2 / Q86U28 / ENSMUSG00000021241 / ENSG00000165898 / ENSRNOG00000012085
ISG15 / P05161 / ENSMUSG00000035692 / ENSG00000187608 / ENSRNOG00000021802
ISOC1 / Q96CN7 / ENSMUSG00000024601 / ENSG00000066583 / ENSRNOG00000019711
IST1 / P53990 / ENSMUSG00000031729 / ENSG00000182149 / ENSRNOG00000015144
ISYNA1 / Q9NPH2 / ENSMUSG00000019139 / ENSG00000105655 / ENSRNOG00000019741
ITFG1 / Q8TB96 / ENSMUSG00000031703 / ENSG00000129636 / ENSRNOG00000015843
ITFG2 / Q969R8 / ENSMUSG00000001518 / ENSG00000111203 / ENSRNOG00000006264
ITGA1 / P56199 / ENSMUSG00000042284 / ENSG00000213949 / ENSRNOG00000053550
ITGA2 / P17301 / ENSMUSG00000015533 / ENSG00000164171 / ENSRNOG00000058111
ITGA3 / P26006 / ENSMUSG00000001507 / ENSG00000005884 / ENSRNOG00000004276
ITGA8 / P53708 / ENSMUSG00000026768 / ENSG00000077943 / ENSRNOG00000016538
Homologs
Brain
Heart
Kidney
Liver
Lung
Testis
Organs
Bins
1
2
3
4
5
Species
Human
Mouse
Rat

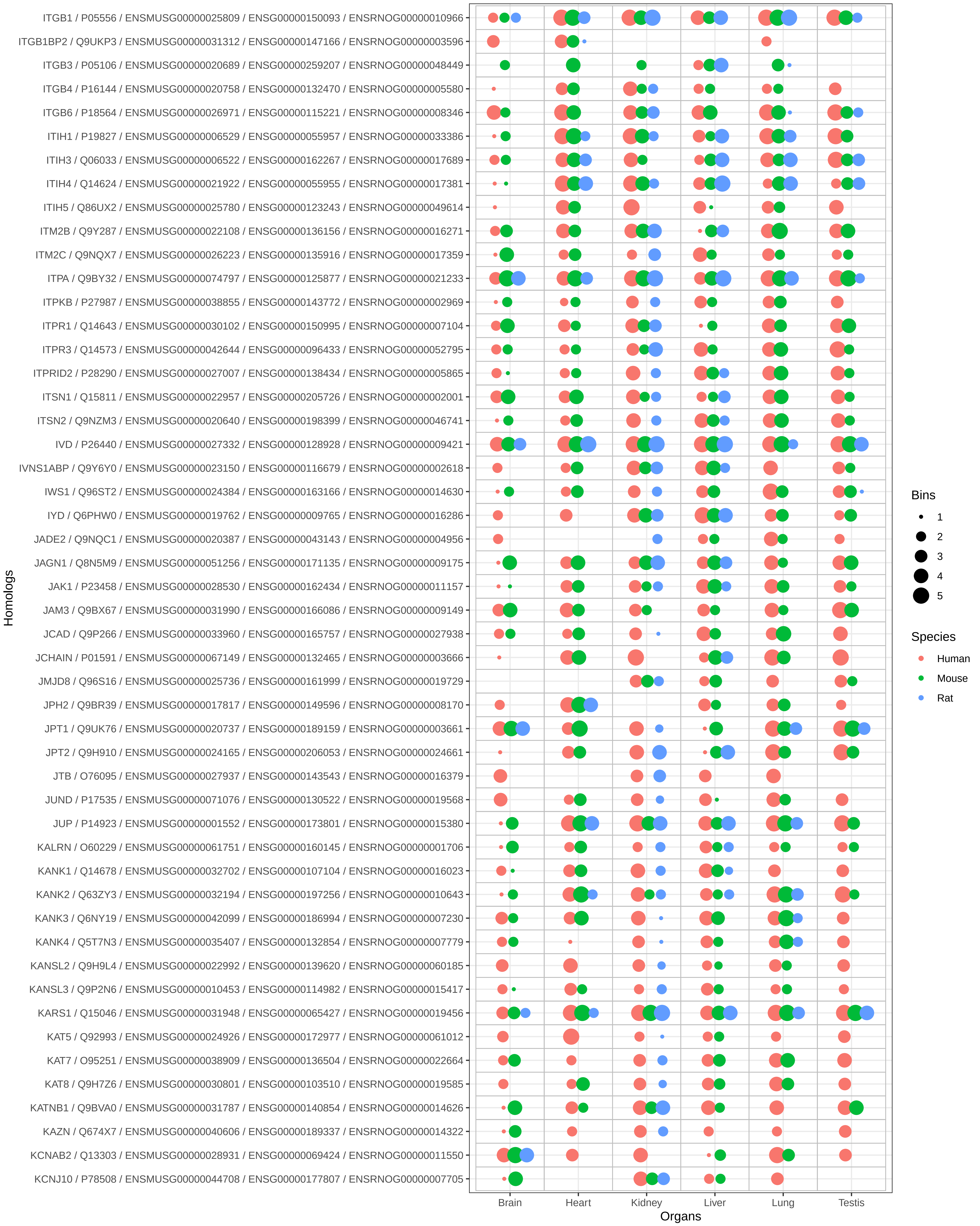
ITGB1 / P05556 / ENSMUSG00000025809 / ENSG00000150093 / ENSRNOG00000010966
ITGB1BP2 / Q9UKP3 / ENSMUSG00000031312 / ENSG00000147166 / ENSRNOG00000003596
ITGB3 / P05106 / ENSMUSG00000020689 / ENSG00000259207 / ENSRNOG00000048449
ITGB4 / P16144 / ENSMUSG00000020758 / ENSG00000132470 / ENSRNOG00000005580
ITGB6 / P18564 / ENSMUSG00000026971 / ENSG00000115221 / ENSRNOG00000008346
ITIH1 / P19827 / ENSMUSG00000006529 / ENSG00000055957 / ENSRNOG00000033386
ITIH3 / Q06033 / ENSMUSG00000006522 / ENSG00000162267 / ENSRNOG00000017689
ITIH4 / Q14624 / ENSMUSG00000021922 / ENSG00000055955 / ENSRNOG00000017381
ITIH5 / Q86UX2 / ENSMUSG00000025780 / ENSG00000123243 / ENSRNOG00000049614
ITM2B / Q9Y287 / ENSMUSG00000022108 / ENSG00000136156 / ENSRNOG00000016271
ITM2C / Q9NQX7 / ENSMUSG00000026223 / ENSG00000135916 / ENSRNOG00000017359
ITPA / Q9BY32 / ENSMUSG00000074797 / ENSG00000125877 / ENSRNOG00000021233
ITPKB / P27987 / ENSMUSG00000038855 / ENSG00000143772 / ENSRNOG00000002969
ITPR1 / Q14643 / ENSMUSG00000030102 / ENSG00000150995 / ENSRNOG00000007104
ITPR3 / Q14573 / ENSMUSG00000042644 / ENSG00000096433 / ENSRNOG00000052795
ITPRID2 / P28290 / ENSMUSG00000027007 / ENSG00000138434 / ENSRNOG00000005865
ITSN1 / Q15811 / ENSMUSG00000022957 / ENSG00000205726 / ENSRNOG00000002001
ITSN2 / Q9NZM3 / ENSMUSG00000020640 / ENSG00000198399 / ENSRNOG00000046741
IVD / P26440 / ENSMUSG00000027332 / ENSG00000128928 / ENSRNOG00000009421
IVNS1ABP / Q9Y6Y0 / ENSMUSG00000023150 / ENSG00000116679 / ENSRNOG00000002618
IWS1 / Q96ST2 / ENSMUSG00000024384 / ENSG00000163166 / ENSRNOG00000014630
IYD / Q6PHW0 / ENSMUSG00000019762 / ENSG00000009765 / ENSRNOG00000016286
JADE2 / Q9NQC1 / ENSMUSG00000020387 / ENSG00000043143 / ENSRNOG00000004956
JAGN1 / Q8N5M9 / ENSMUSG00000051256 / ENSG00000171135 / ENSRNOG00000009175
JAK1 / P23458 / ENSMUSG00000028530 / ENSG00000162434 / ENSRNOG00000011157
JAM3 / Q9BX67 / ENSMUSG00000031990 / ENSG00000166086 / ENSRNOG00000009149
JCAD / Q9P266 / ENSMUSG00000033960 / ENSG00000165757 / ENSRNOG00000027938
JCHAIN / P01591 / ENSMUSG00000067149 / ENSG00000132465 / ENSRNOG00000003666
JMJD8 / Q96S16 / ENSMUSG00000025736 / ENSG00000161999 / ENSRNOG00000019729
JPH2 / Q9BR39 / ENSMUSG00000017817 / ENSG00000149596 / ENSRNOG00000008170
JPT1 / Q9UK76 / ENSMUSG00000020737 / ENSG00000189159 / ENSRNOG00000003661
JPT2 / Q9H910 / ENSMUSG00000024165 / ENSG00000206053 / ENSRNOG00000024661
JTB / O76095 / ENSMUSG00000027937 / ENSG00000143543 / ENSRNOG00000016379
JUND / P17535 / ENSMUSG00000071076 / ENSG00000130522 / ENSRNOG00000019568
JUP / P14923 / ENSMUSG00000001552 / ENSG00000173801 / ENSRNOG00000015380
KALRN / O60229 / ENSMUSG00000061751 / ENSG00000160145 / ENSRNOG00000001706
KANK1 / Q14678 / ENSMUSG00000032702 / ENSG00000107104 / ENSRNOG00000016023
KANK2 / Q63ZY3 / ENSMUSG00000032194 / ENSG00000197256 / ENSRNOG00000010643
KANK3 / Q6NY19 / ENSMUSG00000042099 / ENSG00000186994 / ENSRNOG00000007230
KANK4 / Q5T7N3 / ENSMUSG00000035407 / ENSG00000132854 / ENSRNOG00000007779
KANSL2 / Q9H9L4 / ENSMUSG00000022992 / ENSG00000139620 / ENSRNOG00000060185
KANSL3 / Q9P2N6 / ENSMUSG00000010453 / ENSG00000114982 / ENSRNOG00000015417
KARS1 / Q15046 / ENSMUSG00000031948 / ENSG00000065427 / ENSRNOG00000019456
KAT5 / Q92993 / ENSMUSG00000024926 / ENSG00000172977 / ENSRNOG00000061012
KAT7 / O95251 / ENSMUSG00000038909 / ENSG00000136504 / ENSRNOG00000022664
KAT8 / Q9H7Z6 / ENSMUSG00000030801 / ENSG00000103510 / ENSRNOG00000019585
KATNB1 / Q9BVA0 / ENSMUSG00000031787 / ENSG00000140854 / ENSRNOG00000014626
KAZN / Q674X7 / ENSMUSG00000040606 / ENSG00000189337 / ENSRNOG00000014322
KCNAB2 / Q13303 / ENSMUSG00000028931 / ENSG00000069424 / ENSRNOG00000011550
KCNJ10 / P78508 / ENSMUSG00000044708 / ENSG00000177807 / ENSRNOG00000007705
Homologs
Brain
Heart
Kidney
Liver
Lung
Testis
Organs
Bins
1
2
3
4
5
Species
Human
Mouse
Rat

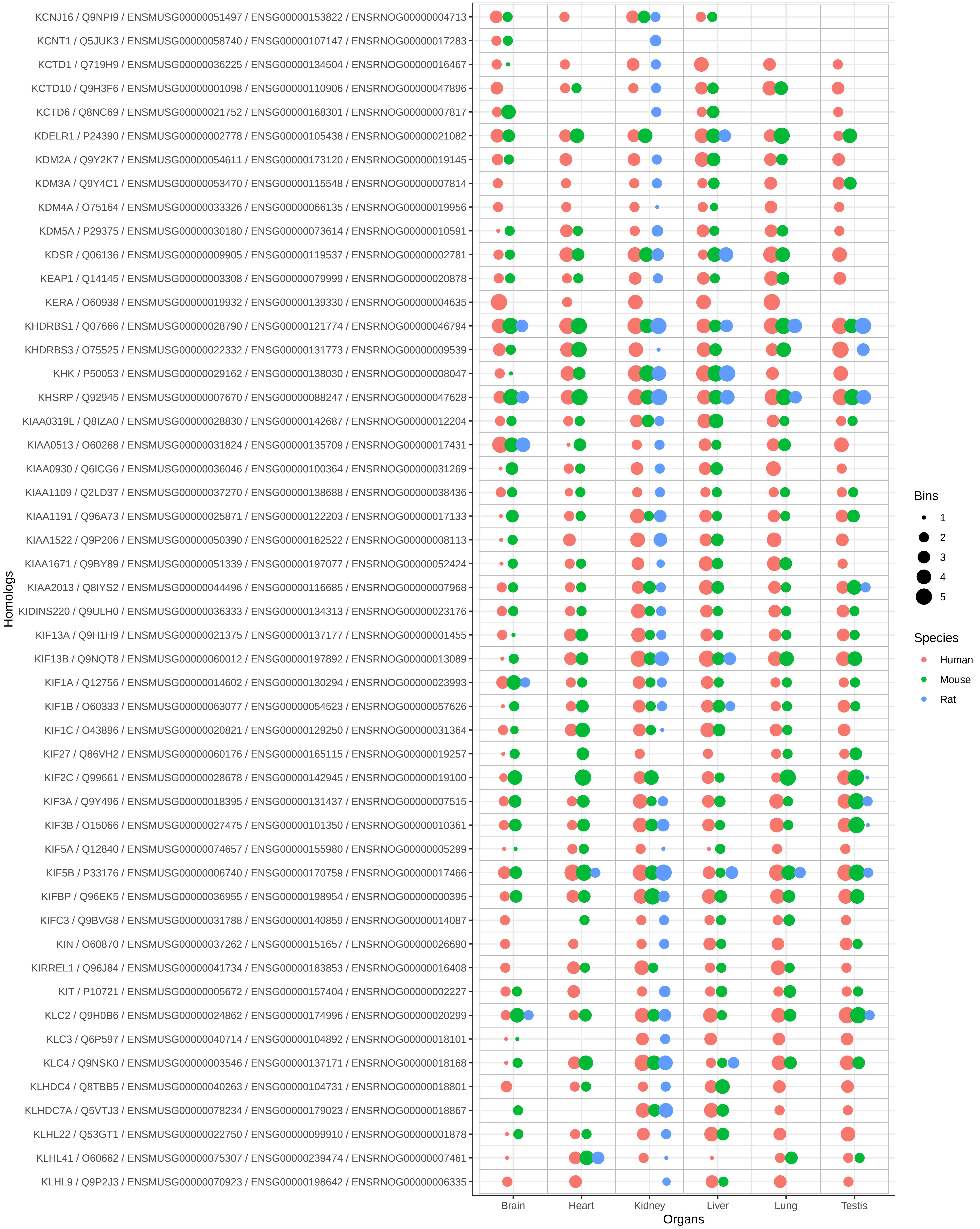
KCNJ16 / Q9NPI9 / ENSMUSG00000051497 / ENSG00000153822 / ENSRNOG00000004713
KCNT1 / Q5JUK3 / ENSMUSG00000058740 / ENSG00000107147 / ENSRNOG00000017283
KCTD1 / Q719H9 / ENSMUSG00000036225 / ENSG00000134504 / ENSRNOG00000016467
KCTD10 / Q9H3F6 / ENSMUSG00000001098 / ENSG00000110906 / ENSRNOG00000047896
KCTD6 / Q8NC69 / ENSMUSG00000021752 / ENSG00000168301 / ENSRNOG00000007817
KDELR1 / P24390 / ENSMUSG00000002778 / ENSG00000105438 / ENSRNOG00000021082
KDM2A / Q9Y2K7 / ENSMUSG00000054611 / ENSG00000173120 / ENSRNOG00000019145
KDM3A / Q9Y4C1 / ENSMUSG00000053470 / ENSG00000115548 / ENSRNOG00000007814
KDM4A / O75164 / ENSMUSG00000033326 / ENSG00000066135 / ENSRNOG00000019956
KDM5A / P29375 / ENSMUSG00000030180 / ENSG00000073614 / ENSRNOG00000010591
KDSR / Q06136 / ENSMUSG00000009905 / ENSG00000119537 / ENSRNOG00000002781
KEAP1 / Q14145 / ENSMUSG00000003308 / ENSG00000079999 / ENSRNOG00000020878
KERA / O60938 / ENSMUSG00000019932 / ENSG00000139330 / ENSRNOG00000004635
KHDRBS1 / Q07666 / ENSMUSG00000028790 / ENSG00000121774 / ENSRNOG00000046794
KHDRBS3 / O75525 / ENSMUSG00000022332 / ENSG00000131773 / ENSRNOG00000009539
KHK / P50053 / ENSMUSG00000029162 / ENSG00000138030 / ENSRNOG00000008047
KHSRP / Q92945 / ENSMUSG00000007670 / ENSG00000088247 / ENSRNOG00000047628
KIAA0319L / Q8IZA0 / ENSMUSG00000028830 / ENSG00000142687 / ENSRNOG00000012204
KIAA0513 / O60268 / ENSMUSG00000031824 / ENSG00000135709 / ENSRNOG00000017431
KIAA0930 / Q6ICG6 / ENSMUSG00000036046 / ENSG00000100364 / ENSRNOG00000031269
KIAA1109 / Q2LD37 / ENSMUSG00000037270 / ENSG00000138688 / ENSRNOG00000038436
KIAA1191 / Q96A73 / ENSMUSG00000025871 / ENSG00000122203 / ENSRNOG00000017133
KIAA1522 / Q9P206 / ENSMUSG00000050390 / ENSG00000162522 / ENSRNOG00000008113
KIAA1671 / Q9BY89 / ENSMUSG00000051339 / ENSG00000197077 / ENSRNOG00000052424
KIAA2013 / Q8IYS2 / ENSMUSG00000044496 / ENSG00000116685 / ENSRNOG00000007968
KIDINS220 / Q9ULH0 / ENSMUSG00000036333 / ENSG00000134313 / ENSRNOG00000023176
KIF13A / Q9H1H9 / ENSMUSG00000021375 / ENSG00000137177 / ENSRNOG00000001455
KIF13B / Q9NQT8 / ENSMUSG00000060012 / ENSG00000197892 / ENSRNOG00000013089
KIF1A / Q12756 / ENSMUSG00000014602 / ENSG00000130294 / ENSRNOG00000023993
KIF1B / O60333 / ENSMUSG00000063077 / ENSG00000054523 / ENSRNOG00000057626
KIF1C / O43896 / ENSMUSG00000020821 / ENSG00000129250 / ENSRNOG00000031364
KIF27 / Q86VH2 / ENSMUSG00000060176 / ENSG00000165115 / ENSRNOG00000019257
KIF2C / Q99661 / ENSMUSG00000028678 / ENSG00000142945 / ENSRNOG00000019100
KIF3A / Q9Y496 / ENSMUSG00000018395 / ENSG00000131437 / ENSRNOG00000007515
KIF3B / O15066 / ENSMUSG00000027475 / ENSG00000101350 / ENSRNOG00000010361
KIF5A / Q12840 / ENSMUSG00000074657 / ENSG00000155980 / ENSRNOG00000005299
KIF5B / P33176 / ENSMUSG00000006740 / ENSG00000170759 / ENSRNOG00000017466
KIFBP / Q96EK5 / ENSMUSG00000036955 / ENSG00000198954 / ENSRNOG00000000395
KIFC3 / Q9BVG8 / ENSMUSG00000031788 / ENSG00000140859 / ENSRNOG00000014087
KIN / O60870 / ENSMUSG00000037262 / ENSG00000151657 / ENSRNOG00000026690
KIRREL1 / Q96J84 / ENSMUSG00000041734 / ENSG00000183853 / ENSRNOG00000016408
KIT / P10721 / ENSMUSG00000005672 / ENSG00000157404 / ENSRNOG00000002227
KLC2 / Q9H0B6 / ENSMUSG00000024862 / ENSG00000174996 / ENSRNOG00000020299
KLC3 / Q6P597 / ENSMUSG00000040714 / ENSG00000104892 / ENSRNOG00000018101
KLC4 / Q9NSK0 / ENSMUSG00000003546 / ENSG00000137171 / ENSRNOG00000018168
KLHDC4 / Q8TBB5 / ENSMUSG00000040263 / ENSG00000104731 / ENSRNOG00000018801
KLHDC7A / Q5VTJ3 / ENSMUSG00000078234 / ENSG00000179023 / ENSRNOG00000018867
KLHL22 / Q53GT1 / ENSMUSG00000022750 / ENSG00000099910 / ENSRNOG00000001878
KLHL41 / O60662 / ENSMUSG00000075307 / ENSG00000239474 / ENSRNOG00000007461
KLHL9 / Q9P2J3 / ENSMUSG00000070923 / ENSG00000198642 / ENSRNOG00000006335
Homologs
Brain
Heart
Kidney
Liver
Lung
Testis
Organs
Bins
1
2
3
4
5
Species
Human
Mouse
Rat

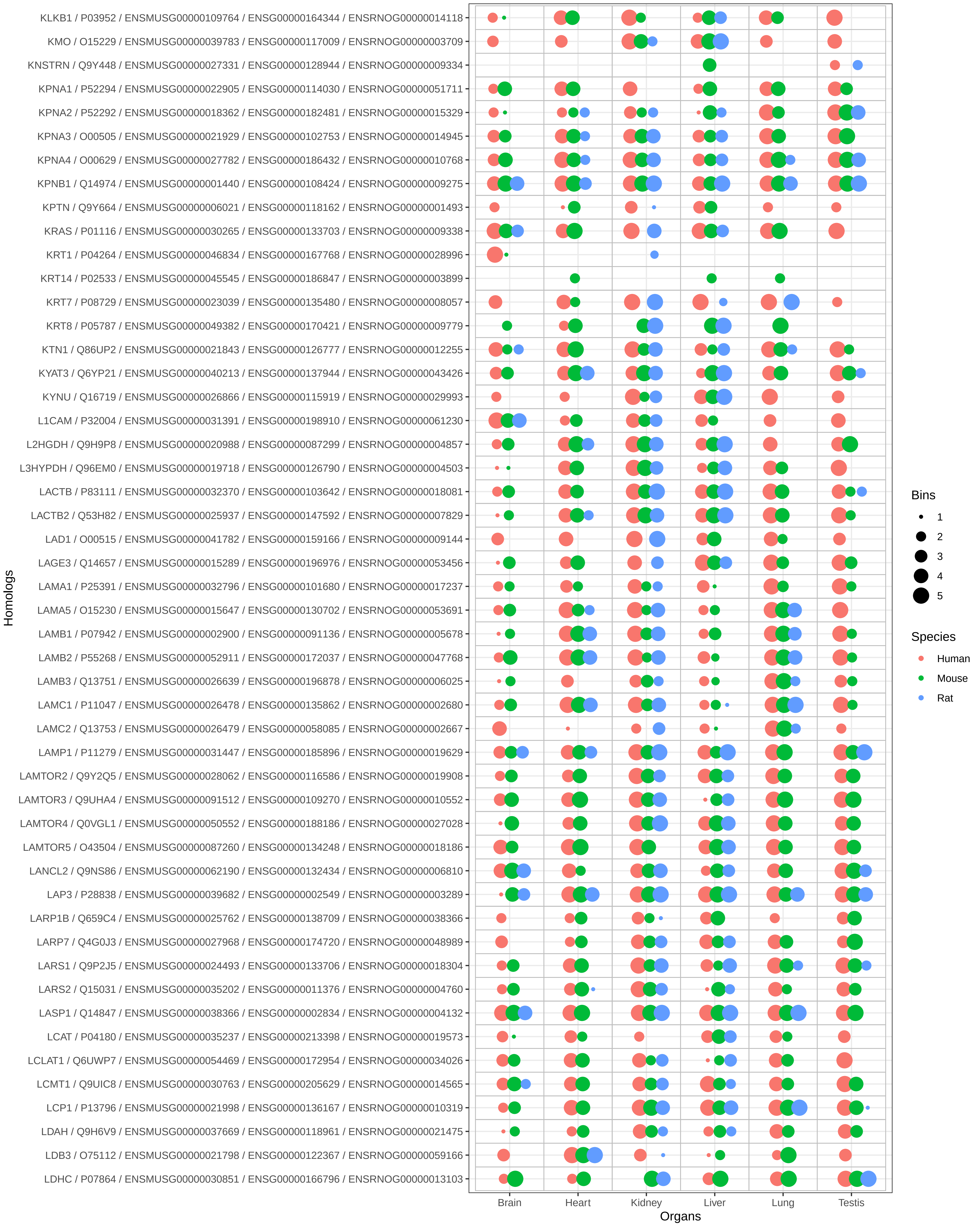
KLKB1 / P03952 / ENSMUSG00000109764 / ENSG00000164344 / ENSRNOG00000014118
KMO / O15229 / ENSMUSG00000039783 / ENSG00000117009 / ENSRNOG00000003709
KNSTRN / Q9Y448 / ENSMUSG00000027331 / ENSG00000128944 / ENSRNOG00000009334
KPNA1 / P52294 / ENSMUSG00000022905 / ENSG00000114030 / ENSRNOG00000051711
KPNA2 / P52292 / ENSMUSG00000018362 / ENSG00000182481 / ENSRNOG00000015329
KPNA3 / O00505 / ENSMUSG00000021929 / ENSG00000102753 / ENSRNOG00000014945
KPNA4 / O00629 / ENSMUSG00000027782 / ENSG00000186432 / ENSRNOG00000010768
KPNB1 / Q14974 / ENSMUSG00000001440 / ENSG00000108424 / ENSRNOG00000009275
KPTN / Q9Y664 / ENSMUSG00000006021 / ENSG00000118162 / ENSRNOG00000001493
KRAS / P01116 / ENSMUSG00000030265 / ENSG00000133703 / ENSRNOG00000009338
KRT1 / P04264 / ENSMUSG00000046834 / ENSG00000167768 / ENSRNOG00000028996
KRT14 / P02533 / ENSMUSG00000045545 / ENSG00000186847 / ENSRNOG00000003899
KRT7 / P08729 / ENSMUSG00000023039 / ENSG00000135480 / ENSRNOG00000008057
KRT8 / P05787 / ENSMUSG00000049382 / ENSG00000170421 / ENSRNOG00000009779
KTN1 / Q86UP2 / ENSMUSG00000021843 / ENSG00000126777 / ENSRNOG00000012255
KYAT3 / Q6YP21 / ENSMUSG00000040213 / ENSG00000137944 / ENSRNOG00000043426
KYNU / Q16719 / ENSMUSG00000026866 / ENSG00000115919 / ENSRNOG00000029993
L1CAM / P32004 / ENSMUSG00000031391 / ENSG00000198910 / ENSRNOG00000061230
L2HGDH / Q9H9P8 / ENSMUSG00000020988 / ENSG00000087299 / ENSRNOG00000004857
L3HYPDH / Q96EM0 / ENSMUSG00000019718 / ENSG00000126790 / ENSRNOG00000004503
LACTB / P83111 / ENSMUSG00000032370 / ENSG00000103642 / ENSRNOG00000018081
LACTB2 / Q53H82 / ENSMUSG00000025937 / ENSG00000147592 / ENSRNOG00000007829
LAD1 / O00515 / ENSMUSG00000041782 / ENSG00000159166 / ENSRNOG00000009144
LAGE3 / Q14657 / ENSMUSG00000015289 / ENSG00000196976 / ENSRNOG00000053456
LAMA1 / P25391 / ENSMUSG00000032796 / ENSG00000101680 / ENSRNOG00000017237
LAMA5 / O15230 / ENSMUSG00000015647 / ENSG00000130702 / ENSRNOG00000053691
LAMB1 / P07942 / ENSMUSG00000002900 / ENSG00000091136 / ENSRNOG00000005678
LAMB2 / P55268 / ENSMUSG00000052911 / ENSG00000172037 / ENSRNOG00000047768
LAMB3 / Q13751 / ENSMUSG00000026639 / ENSG00000196878 / ENSRNOG00000006025
LAMC1 / P11047 / ENSMUSG00000026478 / ENSG00000135862 / ENSRNOG00000002680
LAMC2 / Q13753 / ENSMUSG00000026479 / ENSG00000058085 / ENSRNOG00000002667
LAMP1 / P11279 / ENSMUSG00000031447 / ENSG00000185896 / ENSRNOG00000019629
LAMTOR2 / Q9Y2Q5 / ENSMUSG00000028062 / ENSG00000116586 / ENSRNOG00000019908
LAMTOR3 / Q9UHA4 / ENSMUSG00000091512 / ENSG00000109270 / ENSRNOG00000010552
LAMTOR4 / Q0VGL1 / ENSMUSG00000050552 / ENSG00000188186 / ENSRNOG00000027028
LAMTOR5 / O43504 / ENSMUSG00000087260 / ENSG00000134248 / ENSRNOG00000018186
LANCL2 / Q9NS86 / ENSMUSG00000062190 / ENSG00000132434 / ENSRNOG00000006810
LAP3 / P28838 / ENSMUSG00000039682 / ENSG00000002549 / ENSRNOG00000003289
LARP1B / Q659C4 / ENSMUSG00000025762 / ENSG00000138709 / ENSRNOG00000038366
LARP7 / Q4G0J3 / ENSMUSG00000027968 / ENSG00000174720 / ENSRNOG00000048989
LARS1 / Q9P2J5 / ENSMUSG00000024493 / ENSG00000133706 / ENSRNOG00000018304
LARS2 / Q15031 / ENSMUSG00000035202 / ENSG00000011376 / ENSRNOG00000004760
LASP1 / Q14847 / ENSMUSG00000038366 / ENSG00000002834 / ENSRNOG00000004132
LCAT / P04180 / ENSMUSG00000035237 / ENSG00000213398 / ENSRNOG00000019573
LCLAT1 / Q6UWP7 / ENSMUSG00000054469 / ENSG00000172954 / ENSRNOG00000034026
LCMT1 / Q9UIC8 / ENSMUSG00000030763 / ENSG00000205629 / ENSRNOG00000014565
LCP1 / P13796 / ENSMUSG00000021998 / ENSG00000136167 / ENSRNOG00000010319
LDAH / Q9H6V9 / ENSMUSG00000037669 / ENSG00000118961 / ENSRNOG00000021475
LDB3 / O75112 / ENSMUSG00000021798 / ENSG00000122367 / ENSRNOG00000059166
LDHC / P07864 / ENSMUSG00000030851 / ENSG00000166796 / ENSRNOG00000013103
Homologs
Brain
Heart
Kidney
Liver
Lung
Testis
Organs
Bins
1
2
3
4
5
Species
Human
Mouse
Rat

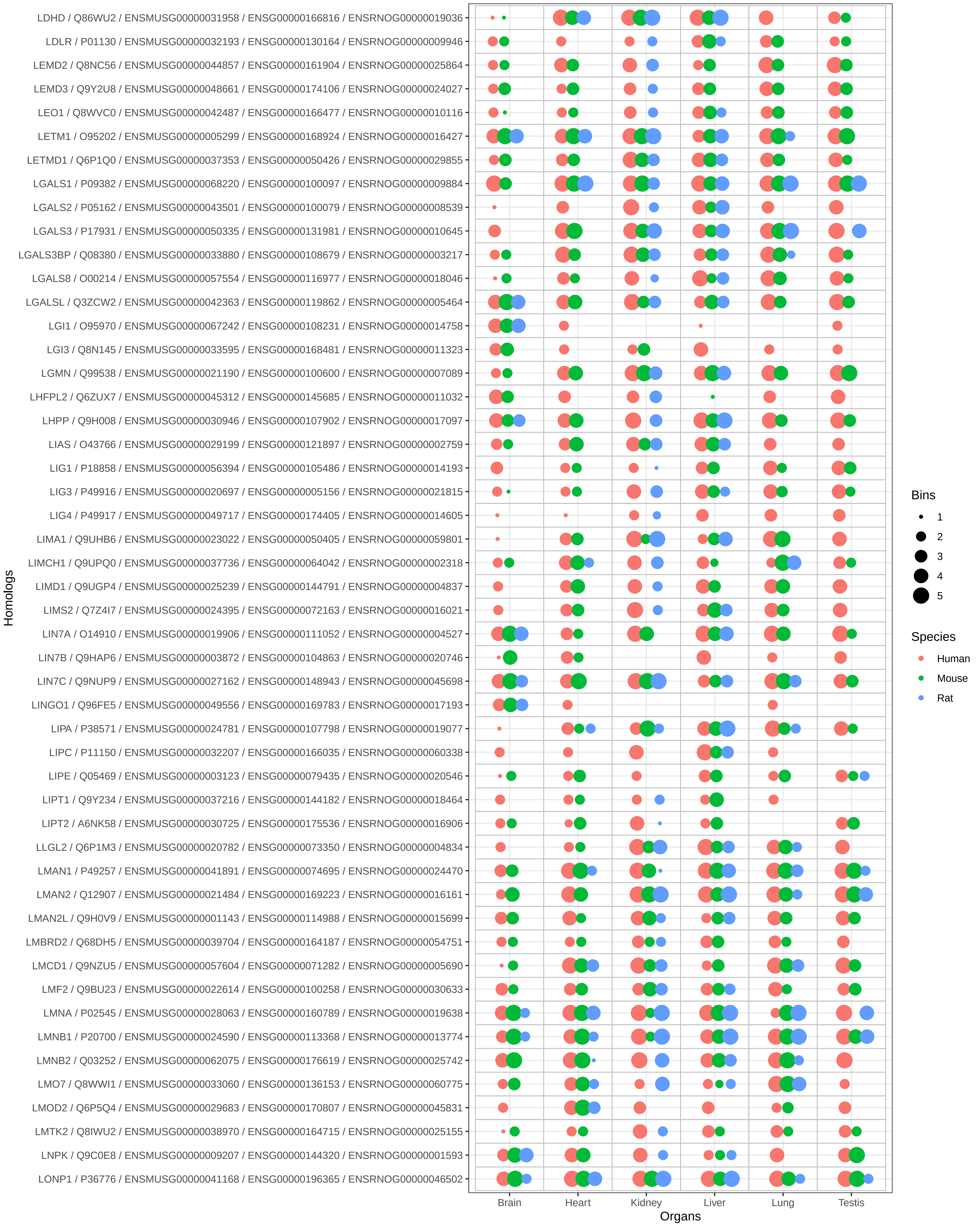

LDHD / Q86WU2 / ENSMUSG00000031958 / ENSG00000166816 / ENSRNOG00000019036
LDLR / P01130 / ENSMUSG00000032193 / ENSG00000130164 / ENSRNOG00000009946
LEMD2 / Q8NC56 / ENSMUSG00000044857 / ENSG00000161904 / ENSRNOG00000025864
LEMD3 / Q9Y2U8 / ENSMUSG00000048661 / ENSG00000174106 / ENSRNOG00000024027
LEO1 / Q8WVC0 / ENSMUSG00000042487 / ENSG00000166477 / ENSRNOG00000010116
LETM1 / O95202 / ENSMUSG00000005299 / ENSG00000168924 / ENSRNOG00000016427
LETMD1 / Q6P1Q0 / ENSMUSG00000037353 / ENSG00000050426 / ENSRNOG00000029855
LGALS1 / P09382 / ENSMUSG00000068220 / ENSG00000100097 / ENSRNOG00000009884
LGALS2 / P05162 / ENSMUSG00000043501 / ENSG00000100079 / ENSRNOG00000008539
LGALS3 / P17931 / ENSMUSG00000050335 / ENSG00000131981 / ENSRNOG00000010645
LGALS3BP / Q08380 / ENSMUSG00000033880 / ENSG00000108679 / ENSRNOG00000003217
LGALS8 / O00214 / ENSMUSG00000057554 / ENSG00000116977 / ENSRNOG00000018046
LGALSL / Q3ZCW2 / ENSMUSG00000042363 / ENSG00000119862 / ENSRNOG00000005464
LGI1 / O95970 / ENSMUSG00000067242 / ENSG00000108231 / ENSRNOG00000014758
LGI3 / Q8N145 / ENSMUSG00000033595 / ENSG00000168481 / ENSRNOG00000011323
LGMN / Q99538 / ENSMUSG00000021190 / ENSG00000100600 / ENSRNOG00000007089
LHFPL2 / Q6ZUX7 / ENSMUSG00000045312 / ENSG00000145685 / ENSRNOG00000011032
LHPP / Q9H008 / ENSMUSG00000030946 / ENSG00000107902 / ENSRNOG00000017097
LIAS / O43766 / ENSMUSG00000029199 / ENSG00000121897 / ENSRNOG00000002759
LIG1 / P18858 / ENSMUSG00000056394 / ENSG00000105486 / ENSRNOG00000014193
LIG3 / P49916 / ENSMUSG00000020697 / ENSG00000005156 / ENSRNOG00000021815
LIG4 / P49917 / ENSMUSG00000049717 / ENSG00000174405 / ENSRNOG00000014605
LIMA1 / Q9UHB6 / ENSMUSG00000023022 / ENSG00000050405 / ENSRNOG00000059801
LIMCH1 / Q9UPQ0 / ENSMUSG00000037736 / ENSG00000064042 / ENSRNOG00000002318
LIMD1 / Q9UGP4 / ENSMUSG00000025239 / ENSG00000144791 / ENSRNOG00000004837
LIMS2 / Q7Z4I7 / ENSMUSG00000024395 / ENSG00000072163 / ENSRNOG00000016021
LIN7A / O14910 / ENSMUSG00000019906 / ENSG00000111052 / ENSRNOG00000004527
LIN7B / Q9HAP6 / ENSMUSG00000003872 / ENSG00000104863 / ENSRNOG00000020746
LIN7C / Q9NUP9 / ENSMUSG00000027162 / ENSG00000148943 / ENSRNOG00000045698
LINGO1 / Q96FE5 / ENSMUSG00000049556 / ENSG00000169783 / ENSRNOG00000017193
LIPA / P38571 / ENSMUSG00000024781 / ENSG00000107798 / ENSRNOG00000019077
LIPC / P11150 / ENSMUSG00000032207 / ENSG00000166035 / ENSRNOG00000060338
LIPE / Q05469 / ENSMUSG00000003123 / ENSG00000079435 / ENSRNOG00000020546
LIPT1 / Q9Y234 / ENSMUSG00000037216 / ENSG00000144182 / ENSRNOG00000018464
LIPT2 / A6NK58 / ENSMUSG00000030725 / ENSG00000175536 / ENSRNOG00000016906
LLGL2 / Q6P1M3 / ENSMUSG00000020782 / ENSG00000073350 / ENSRNOG00000004834
LMAN1 / P49257 / ENSMUSG00000041891 / ENSG00000074695 / ENSRNOG00000024470
LMAN2 / Q12907 / ENSMUSG00000021484 / ENSG00000169223 / ENSRNOG00000016161
LMAN2L / Q9H0V9 / ENSMUSG00000001143 / ENSG00000114988 / ENSRNOG00000015699
LMBRD2 / Q68DH5 / ENSMUSG00000039704 / ENSG00000164187 / ENSRNOG00000054751
LMCD1 / Q9NZU5 / ENSMUSG00000057604 / ENSG00000071282 / ENSRNOG00000005690
LMF2 / Q9BU23 / ENSMUSG00000022614 / ENSG00000100258 / ENSRNOG00000030633
LMNA / P02545 / ENSMUSG00000028063 / ENSG00000160789 / ENSRNOG00000019638
LMNB1 / P20700 / ENSMUSG00000024590 / ENSG00000113368 / ENSRNOG00000013774
LMNB2 / Q03252 / ENSMUSG00000062075 / ENSG00000176619 / ENSRNOG00000025742
LMO7 / Q8WWI1 / ENSMUSG00000033060 / ENSG00000136153 / ENSRNOG00000060775
LMOD2 / Q6P5Q4 / ENSMUSG00000029683 / ENSG00000170807 / ENSRNOG00000045831
LMTK2 / Q8IWU2 / ENSMUSG00000038970 / ENSG00000164715 / ENSRNOG00000025155
LNPK / Q9C0E8 / ENSMUSG00000009207 / ENSG00000144320 / ENSRNOG00000001593
LONP1 / P36776 / ENSMUSG00000041168 / ENSG00000196365 / ENSRNOG00000046502
Homologs
Brain
Heart
Kidney
Liver
Lung
Testis
Organs
Bins
1
2
3
4
5
Species
Human
Mouse
Rat

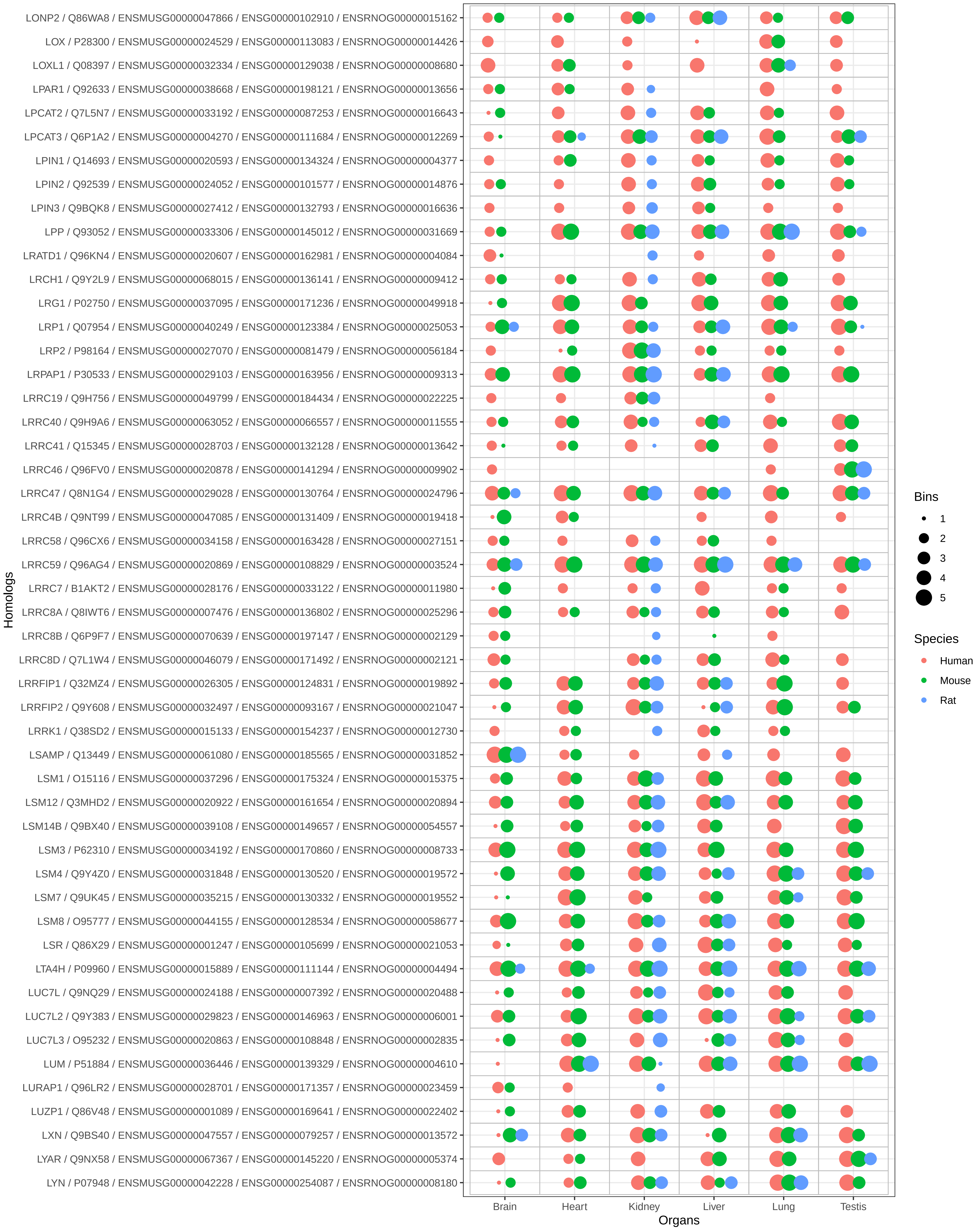
LONP2 / Q86WA8 / ENSMUSG00000047866 / ENSG00000102910 / ENSRNOG00000015162
LOX / P28300 / ENSMUSG00000024529 / ENSG00000113083 / ENSRNOG00000014426
LOXL1 / Q08397 / ENSMUSG00000032334 / ENSG00000129038 / ENSRNOG00000008680
LPAR1 / Q92633 / ENSMUSG00000038668 / ENSG00000198121 / ENSRNOG00000013656
LPCAT2 / Q7L5N7 / ENSMUSG00000033192 / ENSG00000087253 / ENSRNOG00000016643
LPCAT3 / Q6P1A2 / ENSMUSG00000004270 / ENSG00000111684 / ENSRNOG00000012269
LPIN1 / Q14693 / ENSMUSG00000020593 / ENSG00000134324 / ENSRNOG00000004377
LPIN2 / Q92539 / ENSMUSG00000024052 / ENSG00000101577 / ENSRNOG00000014876
LPIN3 / Q9BQK8 / ENSMUSG00000027412 / ENSG00000132793 / ENSRNOG00000016636
LPP / Q93052 / ENSMUSG00000033306 / ENSG00000145012 / ENSRNOG00000031669
LRATD1 / Q96KN4 / ENSMUSG00000020607 / ENSG00000162981 / ENSRNOG00000004084
LRCH1 / Q9Y2L9 / ENSMUSG00000068015 / ENSG00000136141 / ENSRNOG00000009412
LRG1 / P02750 / ENSMUSG00000037095 / ENSG00000171236 / ENSRNOG00000049918
LRP1 / Q07954 / ENSMUSG00000040249 / ENSG00000123384 / ENSRNOG00000025053
LRP2 / P98164 / ENSMUSG00000027070 / ENSG00000081479 / ENSRNOG00000056184
LRPAP1 / P30533 / ENSMUSG00000029103 / ENSG00000163956 / ENSRNOG00000009313
LRRC19 / Q9H756 / ENSMUSG00000049799 / ENSG00000184434 / ENSRNOG00000022225
LRRC40 / Q9H9A6 / ENSMUSG00000063052 / ENSG00000066557 / ENSRNOG00000011555
LRRC41 / Q15345 / ENSMUSG00000028703 / ENSG00000132128 / ENSRNOG00000013642
LRRC46 / Q96FV0 / ENSMUSG00000020878 / ENSG00000141294 / ENSRNOG00000009902
LRRC47 / Q8N1G4 / ENSMUSG00000029028 / ENSG00000130764 / ENSRNOG00000024796
LRRC4B / Q9NT99 / ENSMUSG00000047085 / ENSG00000131409 / ENSRNOG00000019418
LRRC58 / Q96CX6 / ENSMUSG00000034158 / ENSG00000163428 / ENSRNOG00000027151
LRRC59 / Q96AG4 / ENSMUSG00000020869 / ENSG00000108829 / ENSRNOG00000003524
LRRC7 / B1AKT2 / ENSMUSG00000028176 / ENSG00000033122 / ENSRNOG00000011980
LRRC8A / Q8IWT6 / ENSMUSG00000007476 / ENSG00000136802 / ENSRNOG00000025296
LRRC8B / Q6P9F7 / ENSMUSG00000070639 / ENSG00000197147 / ENSRNOG00000002129
LRRC8D / Q7L1W4 / ENSMUSG00000046079 / ENSG00000171492 / ENSRNOG00000002121
LRRFIP1 / Q32MZ4 / ENSMUSG00000026305 / ENSG00000124831 / ENSRNOG00000019892
LRRFIP2 / Q9Y608 / ENSMUSG00000032497 / ENSG00000093167 / ENSRNOG00000021047
LRRK1 / Q38SD2 / ENSMUSG00000015133 / ENSG00000154237 / ENSRNOG00000012730
LSAMP / Q13449 / ENSMUSG00000061080 / ENSG00000185565 / ENSRNOG00000031852
LSM1 / O15116 / ENSMUSG00000037296 / ENSG00000175324 / ENSRNOG00000015375
LSM12 / Q3MHD2 / ENSMUSG00000020922 / ENSG00000161654 / ENSRNOG00000020894
LSM14B / Q9BX40 / ENSMUSG00000039108 / ENSG00000149657 / ENSRNOG00000054557
LSM3 / P62310 / ENSMUSG00000034192 / ENSG00000170860 / ENSRNOG00000008733
LSM4 / Q9Y4Z0 / ENSMUSG00000031848 / ENSG00000130520 / ENSRNOG00000019572
LSM7 / Q9UK45 / ENSMUSG00000035215 / ENSG00000130332 / ENSRNOG00000019552
LSM8 / O95777 / ENSMUSG00000044155 / ENSG00000128534 / ENSRNOG00000058677
LSR / Q86X29 / ENSMUSG00000001247 / ENSG00000105699 / ENSRNOG00000021053
LTA4H / P09960 / ENSMUSG00000015889 / ENSG00000111144 / ENSRNOG00000004494
LUC7L / Q9NQ29 / ENSMUSG00000024188 / ENSG00000007392 / ENSRNOG00000020488
LUC7L2 / Q9Y383 / ENSMUSG00000029823 / ENSG00000146963 / ENSRNOG00000006001
LUC7L3 / O95232 / ENSMUSG00000020863 / ENSG00000108848 / ENSRNOG00000002835
LUM / P51884 / ENSMUSG00000036446 / ENSG00000139329 / ENSRNOG00000004610
LURAP1 / Q96LR2 / ENSMUSG00000028701 / ENSG00000171357 / ENSRNOG00000023459
LUZP1 / Q86V48 / ENSMUSG00000001089 / ENSG00000169641 / ENSRNOG00000022402
LXN / Q9BS40 / ENSMUSG00000047557 / ENSG00000079257 / ENSRNOG00000013572
LYAR / Q9NX58 / ENSMUSG00000067367 / ENSG00000145220 / ENSRNOG00000005374
LYN / P07948 / ENSMUSG00000042228 / ENSG00000254087 / ENSRNOG00000008180
Homologs
Brain
Heart
Kidney
Liver
Lung
Testis
Organs
Bins
1
2
3
4
5
Species
Human
Mouse
Rat

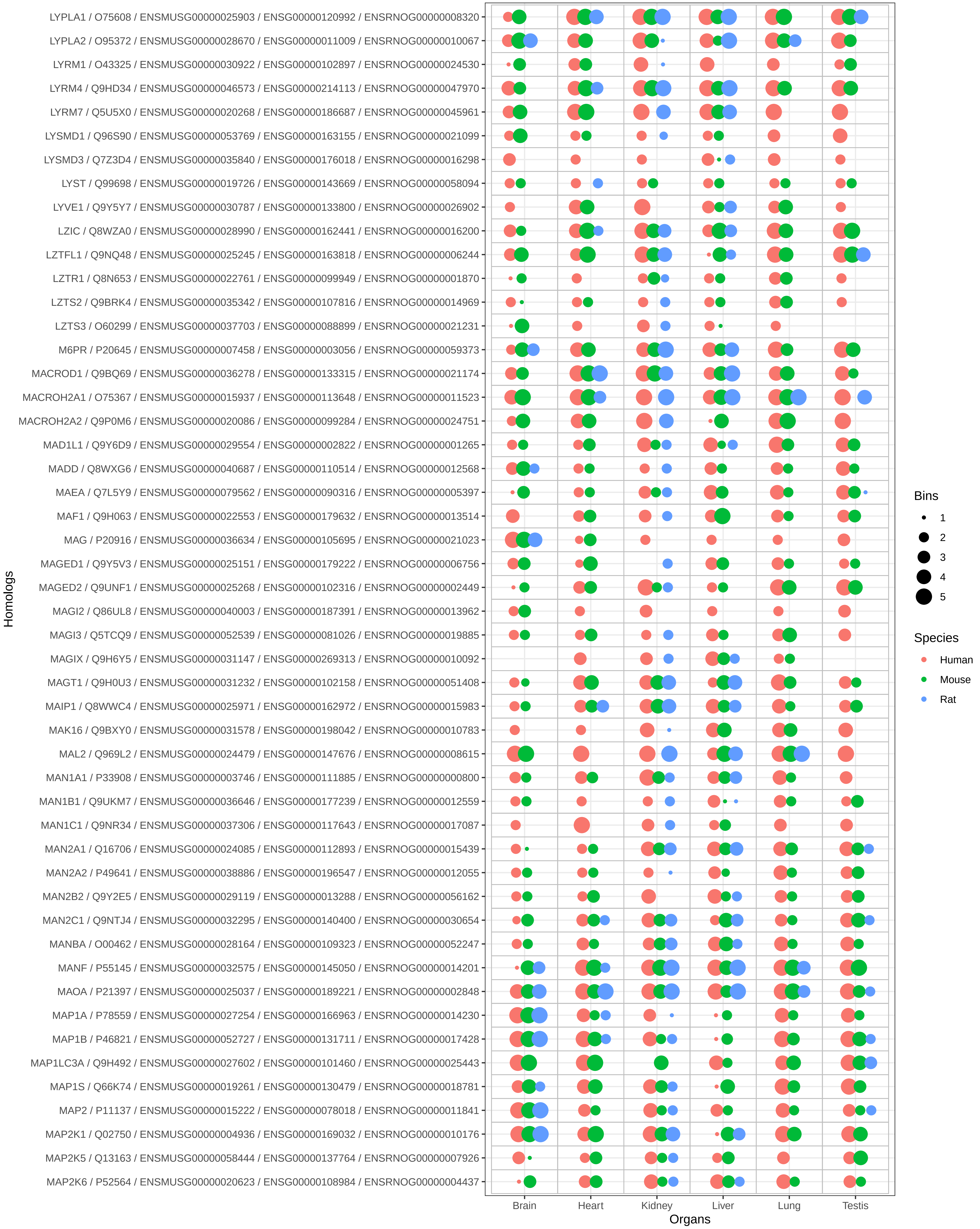
LYPLA1 / O75608 / ENSMUSG00000025903 / ENSG00000120992 / ENSRNOG00000008320
LYPLA2 / O95372 / ENSMUSG00000028670 / ENSG00000011009 / ENSRNOG00000010067
LYRM1 / O43325 / ENSMUSG00000030922 / ENSG00000102897 / ENSRNOG00000024530
LYRM4 / Q9HD34 / ENSMUSG00000046573 / ENSG00000214113 / ENSRNOG00000047970
LYRM7 / Q5U5X0 / ENSMUSG00000020268 / ENSG00000186687 / ENSRNOG00000045961
LYSMD1 / Q96S90 / ENSMUSG00000053769 / ENSG00000163155 / ENSRNOG00000021099
LYSMD3 / Q7Z3D4 / ENSMUSG00000035840 / ENSG00000176018 / ENSRNOG00000016298
LYST / Q99698 / ENSMUSG00000019726 / ENSG00000143669 / ENSRNOG00000058094
LYVE1 / Q9Y5Y7 / ENSMUSG00000030787 / ENSG00000133800 / ENSRNOG00000026902
LZIC / Q8WZA0 / ENSMUSG00000028990 / ENSG00000162441 / ENSRNOG00000016200
LZTFL1 / Q9NQ48 / ENSMUSG00000025245 / ENSG00000163818 / ENSRNOG00000006244
LZTR1 / Q8N653 / ENSMUSG00000022761 / ENSG00000099949 / ENSRNOG00000001870
LZTS2 / Q9BRK4 / ENSMUSG00000035342 / ENSG00000107816 / ENSRNOG00000014969
LZTS3 / O60299 / ENSMUSG00000037703 / ENSG00000088899 / ENSRNOG00000021231
M6PR / P20645 / ENSMUSG00000007458 / ENSG00000003056 / ENSRNOG00000059373
MACROD1 / Q9BQ69 / ENSMUSG00000036278 / ENSG00000133315 / ENSRNOG00000021174
MACROH2A1 / O75367 / ENSMUSG00000015937 / ENSG00000113648 / ENSRNOG00000011523
MACROH2A2 / Q9P0M6 / ENSMUSG00000020086 / ENSG00000099284 / ENSRNOG00000024751
MAD1L1 / Q9Y6D9 / ENSMUSG00000029554 / ENSG00000002822 / ENSRNOG00000001265
MADD / Q8WXG6 / ENSMUSG00000040687 / ENSG00000110514 / ENSRNOG00000012568
MAEA / Q7L5Y9 / ENSMUSG00000079562 / ENSG00000090316 / ENSRNOG00000005397
MAF1 / Q9H063 / ENSMUSG00000022553 / ENSG00000179632 / ENSRNOG00000013514
MAG / P20916 / ENSMUSG00000036634 / ENSG00000105695 / ENSRNOG00000021023
MAGED1 / Q9Y5V3 / ENSMUSG00000025151 / ENSG00000179222 / ENSRNOG00000006756
MAGED2 / Q9UNF1 / ENSMUSG00000025268 / ENSG00000102316 / ENSRNOG00000002449
MAGI2 / Q86UL8 / ENSMUSG00000040003 / ENSG00000187391 / ENSRNOG00000013962
MAGI3 / Q5TCQ9 / ENSMUSG00000052539 / ENSG00000081026 / ENSRNOG00000019885
MAGIX / Q9H6Y5 / ENSMUSG00000031147 / ENSG00000269313 / ENSRNOG00000010092
MAGT1 / Q9H0U3 / ENSMUSG00000031232 / ENSG00000102158 / ENSRNOG00000051408
MAIP1 / Q8WWC4 / ENSMUSG00000025971 / ENSG00000162972 / ENSRNOG00000015983
MAK16 / Q9BXY0 / ENSMUSG00000031578 / ENSG00000198042 / ENSRNOG00000010783
MAL2 / Q969L2 / ENSMUSG00000024479 / ENSG00000147676 / ENSRNOG00000008615
MAN1A1 / P33908 / ENSMUSG00000003746 / ENSG00000111885 / ENSRNOG00000000800
MAN1B1 / Q9UKM7 / ENSMUSG00000036646 / ENSG00000177239 / ENSRNOG00000012559
MAN1C1 / Q9NR34 / ENSMUSG00000037306 / ENSG00000117643 / ENSRNOG00000017087
MAN2A1 / Q16706 / ENSMUSG00000024085 / ENSG00000112893 / ENSRNOG00000015439
MAN2A2 / P49641 / ENSMUSG00000038886 / ENSG00000196547 / ENSRNOG00000012055
MAN2B2 / Q9Y2E5 / ENSMUSG00000029119 / ENSG00000013288 / ENSRNOG00000056162
MAN2C1 / Q9NTJ4 / ENSMUSG00000032295 / ENSG00000140400 / ENSRNOG00000030654
MANBA / O00462 / ENSMUSG00000028164 / ENSG00000109323 / ENSRNOG00000052247
MANF / P55145 / ENSMUSG00000032575 / ENSG00000145050 / ENSRNOG00000014201
MAOA / P21397 / ENSMUSG00000025037 / ENSG00000189221 / ENSRNOG00000002848
MAP1A / P78559 / ENSMUSG00000027254 / ENSG00000166963 / ENSRNOG00000014230
MAP1B / P46821 / ENSMUSG00000052727 / ENSG00000131711 / ENSRNOG00000017428
MAP1LC3A / Q9H492 / ENSMUSG00000027602 / ENSG00000101460 / ENSRNOG00000025443
MAP1S / Q66K74 / ENSMUSG00000019261 / ENSG00000130479 / ENSRNOG00000018781
MAP2 / P11137 / ENSMUSG00000015222 / ENSG00000078018 / ENSRNOG00000011841
MAP2K1 / Q02750 / ENSMUSG00000004936 / ENSG00000169032 / ENSRNOG00000010176
MAP2K5 / Q13163 / ENSMUSG00000058444 / ENSG00000137764 / ENSRNOG00000007926
MAP2K6 / P52564 / ENSMUSG00000020623 / ENSG00000108984 / ENSRNOG00000004437
Homologs
Brain
Heart
Kidney
Liver
Lung
Testis
Organs
Bins
1
2
3
4
5
Species
Human
Mouse
Rat

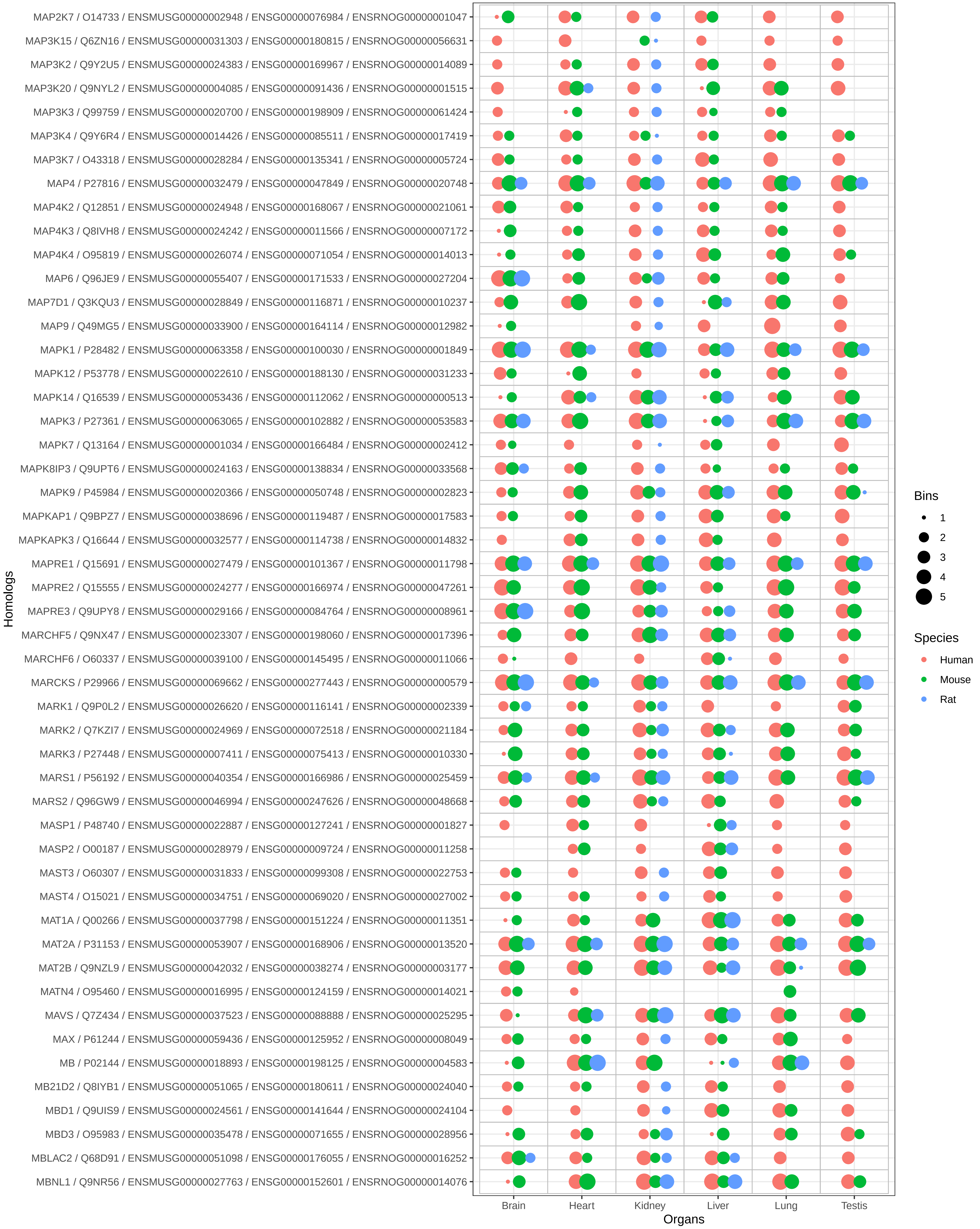
MAP2K7 / O14733 / ENSMUSG00000002948 / ENSG00000076984 / ENSRNOG00000001047
MAP3K15 / Q6ZN16 / ENSMUSG00000031303 / ENSG00000180815 / ENSRNOG00000056631
MAP3K2 / Q9Y2U5 / ENSMUSG00000024383 / ENSG00000169967 / ENSRNOG00000014089
MAP3K20 / Q9NYL2 / ENSMUSG00000004085 / ENSG00000091436 / ENSRNOG00000001515
MAP3K3 / Q99759 / ENSMUSG00000020700 / ENSG00000198909 / ENSRNOG00000061424
MAP3K4 / Q9Y6R4 / ENSMUSG00000014426 / ENSG00000085511 / ENSRNOG00000017419
MAP3K7 / O43318 / ENSMUSG00000028284 / ENSG00000135341 / ENSRNOG00000005724
MAP4 / P27816 / ENSMUSG00000032479 / ENSG00000047849 / ENSRNOG00000020748
MAP4K2 / Q12851 / ENSMUSG00000024948 / ENSG00000168067 / ENSRNOG00000021061
MAP4K3 / Q8IVH8 / ENSMUSG00000024242 / ENSG00000011566 / ENSRNOG00000007172
MAP4K4 / O95819 / ENSMUSG00000026074 / ENSG00000071054 / ENSRNOG00000014013
MAP6 / Q96JE9 / ENSMUSG00000055407 / ENSG00000171533 / ENSRNOG00000027204
MAP7D1 / Q3KQU3 / ENSMUSG00000028849 / ENSG00000116871 / ENSRNOG00000010237
MAP9 / Q49MG5 / ENSMUSG00000033900 / ENSG00000164114 / ENSRNOG00000012982
MAPK1 / P28482 / ENSMUSG00000063358 / ENSG00000100030 / ENSRNOG00000001849
MAPK12 / P53778 / ENSMUSG00000022610 / ENSG00000188130 / ENSRNOG00000031233
MAPK14 / Q16539 / ENSMUSG00000053436 / ENSG00000112062 / ENSRNOG00000000513
MAPK3 / P27361 / ENSMUSG00000063065 / ENSG00000102882 / ENSRNOG00000053583
MAPK7 / Q13164 / ENSMUSG00000001034 / ENSG00000166484 / ENSRNOG00000002412
MAPK8IP3 / Q9UPT6 / ENSMUSG00000024163 / ENSG00000138834 / ENSRNOG00000033568
MAPK9 / P45984 / ENSMUSG00000020366 / ENSG00000050748 / ENSRNOG00000002823
MAPKAP1 / Q9BPZ7 / ENSMUSG00000038696 / ENSG00000119487 / ENSRNOG00000017583
MAPKAPK3 / Q16644 / ENSMUSG00000032577 / ENSG00000114738 / ENSRNOG00000014832
MAPRE1 / Q15691 / ENSMUSG00000027479 / ENSG00000101367 / ENSRNOG00000011798
MAPRE2 / Q15555 / ENSMUSG00000024277 / ENSG00000166974 / ENSRNOG00000047261
MAPRE3 / Q9UPY8 / ENSMUSG00000029166 / ENSG00000084764 / ENSRNOG00000008961
MARCHF5 / Q9NX47 / ENSMUSG00000023307 / ENSG00000198060 / ENSRNOG00000017396
MARCHF6 / O60337 / ENSMUSG00000039100 / ENSG00000145495 / ENSRNOG00000011066
MARCKS / P29966 / ENSMUSG00000069662 / ENSG00000277443 / ENSRNOG00000000579
MARK1 / Q9P0L2 / ENSMUSG00000026620 / ENSG00000116141 / ENSRNOG00000002339
MARK2 / Q7KZI7 / ENSMUSG00000024969 / ENSG00000072518 / ENSRNOG00000021184
MARK3 / P27448 / ENSMUSG00000007411 / ENSG00000075413 / ENSRNOG00000010330
MARS1 / P56192 / ENSMUSG00000040354 / ENSG00000166986 / ENSRNOG00000025459
MARS2 / Q96GW9 / ENSMUSG00000046994 / ENSG00000247626 / ENSRNOG00000048668
MASP1 / P48740 / ENSMUSG00000022887 / ENSG00000127241 / ENSRNOG00000001827
MASP2 / O00187 / ENSMUSG00000028979 / ENSG00000009724 / ENSRNOG00000011258
MAST3 / O60307 / ENSMUSG00000031833 / ENSG00000099308 / ENSRNOG00000022753
MAST4 / O15021 / ENSMUSG00000034751 / ENSG00000069020 / ENSRNOG00000027002
MAT1A / Q00266 / ENSMUSG00000037798 / ENSG00000151224 / ENSRNOG00000011351
MAT2A / P31153 / ENSMUSG00000053907 / ENSG00000168906 / ENSRNOG00000013520
MAT2B / Q9NZL9 / ENSMUSG00000042032 / ENSG00000038274 / ENSRNOG00000003177
MATN4 / O95460 / ENSMUSG00000016995 / ENSG00000124159 / ENSRNOG00000014021
MAVS / Q7Z434 / ENSMUSG00000037523 / ENSG00000088888 / ENSRNOG00000025295
MAX / P61244 / ENSMUSG00000059436 / ENSG00000125952 / ENSRNOG00000008049
MB / P02144 / ENSMUSG00000018893 / ENSG00000198125 / ENSRNOG00000004583
MB21D2 / Q8IYB1 / ENSMUSG00000051065 / ENSG00000180611 / ENSRNOG00000024040
MBD1 / Q9UIS9 / ENSMUSG00000024561 / ENSG00000141644 / ENSRNOG00000024104
MBD3 / O95983 / ENSMUSG00000035478 / ENSG00000071655 / ENSRNOG00000028956
MBLAC2 / Q68D91 / ENSMUSG00000051098 / ENSG00000176055 / ENSRNOG00000016252
MBNL1 / Q9NR56 / ENSMUSG00000027763 / ENSG00000152601 / ENSRNOG00000014076
Homologs
Brain
Heart
Kidney
Liver
Lung
Testis
Organs
Bins
1
2
3
4
5
Species
Human
Mouse
Rat

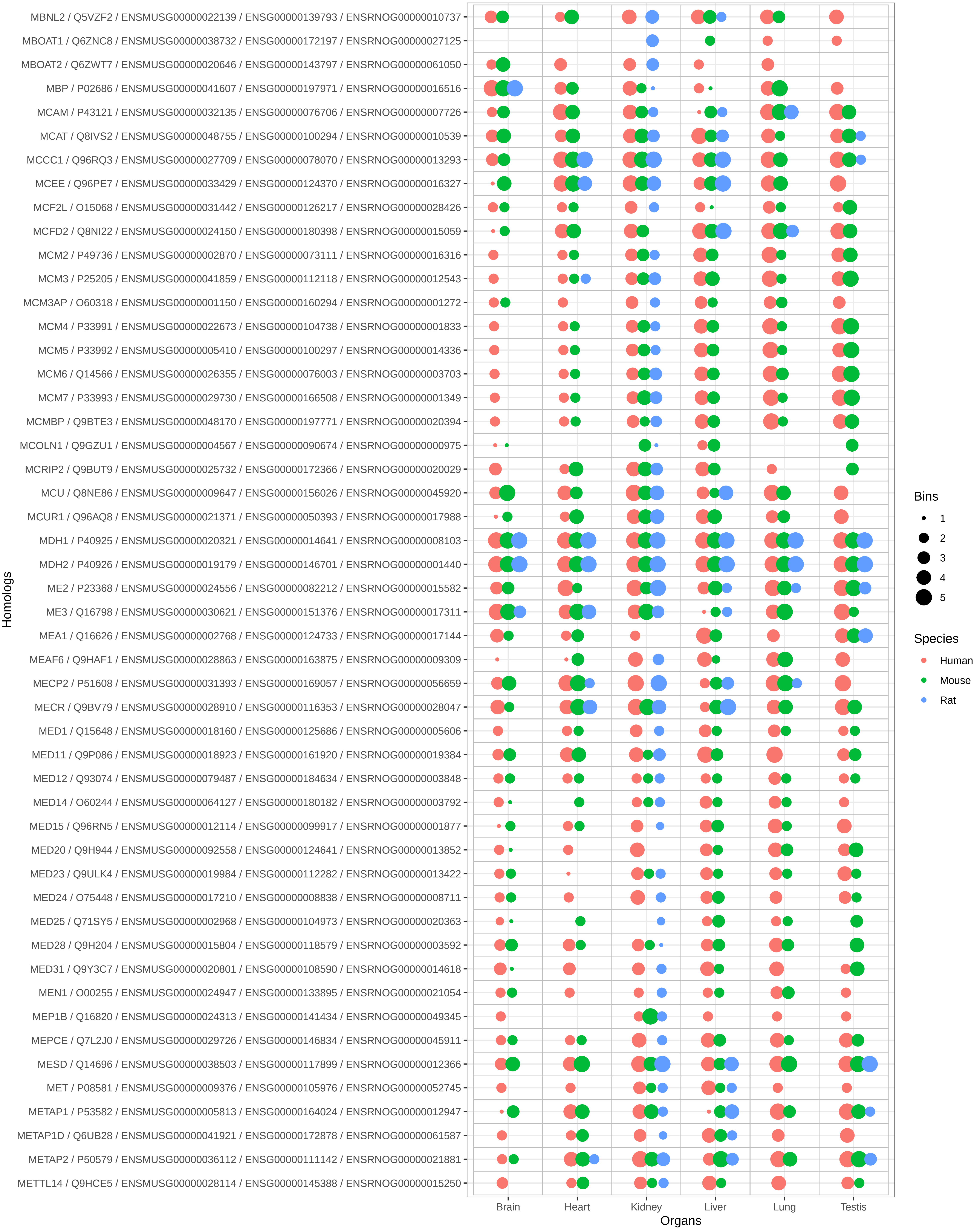
MBNL2 / Q5VZF2 / ENSMUSG00000022139 / ENSG00000139793 / ENSRNOG00000010737
MBOAT1 / Q6ZNC8 / ENSMUSG00000038732 / ENSG00000172197 / ENSRNOG00000027125
MBOAT2 / Q6ZWT7 / ENSMUSG00000020646 / ENSG00000143797 / ENSRNOG00000061050
MBP / P02686 / ENSMUSG00000041607 / ENSG00000197971 / ENSRNOG00000016516
MCAM / P43121 / ENSMUSG00000032135 / ENSG00000076706 / ENSRNOG00000007726
MCAT / Q8IVS2 / ENSMUSG00000048755 / ENSG00000100294 / ENSRNOG00000010539
MCCC1 / Q96RQ3 / ENSMUSG00000027709 / ENSG00000078070 / ENSRNOG00000013293
MCEE / Q96PE7 / ENSMUSG00000033429 / ENSG00000124370 / ENSRNOG00000016327
MCF2L / O15068 / ENSMUSG00000031442 / ENSG00000126217 / ENSRNOG00000028426
MCFD2 / Q8NI22 / ENSMUSG00000024150 / ENSG00000180398 / ENSRNOG00000015059
MCM2 / P49736 / ENSMUSG00000002870 / ENSG00000073111 / ENSRNOG00000016316
MCM3 / P25205 / ENSMUSG00000041859 / ENSG00000112118 / ENSRNOG00000012543
MCM3AP / O60318 / ENSMUSG00000001150 / ENSG00000160294 / ENSRNOG00000001272
MCM4 / P33991 / ENSMUSG00000022673 / ENSG00000104738 / ENSRNOG00000001833
MCM5 / P33992 / ENSMUSG00000005410 / ENSG00000100297 / ENSRNOG00000014336
MCM6 / Q14566 / ENSMUSG00000026355 / ENSG00000076003 / ENSRNOG00000003703
MCM7 / P33993 / ENSMUSG00000029730 / ENSG00000166508 / ENSRNOG00000001349
MCMBP / Q9BTE3 / ENSMUSG00000048170 / ENSG00000197771 / ENSRNOG00000020394
MCOLN1 / Q9GZU1 / ENSMUSG00000004567 / ENSG00000090674 / ENSRNOG00000000975
MCRIP2 / Q9BUT9 / ENSMUSG00000025732 / ENSG00000172366 / ENSRNOG00000020029
MCU / Q8NE86 / ENSMUSG00000009647 / ENSG00000156026 / ENSRNOG00000045920
MCUR1 / Q96AQ8 / ENSMUSG00000021371 / ENSG00000050393 / ENSRNOG00000017988
MDH1 / P40925 / ENSMUSG00000020321 / ENSG00000014641 / ENSRNOG00000008103
MDH2 / P40926 / ENSMUSG00000019179 / ENSG00000146701 / ENSRNOG00000001440
ME2 / P23368 / ENSMUSG00000024556 / ENSG00000082212 / ENSRNOG00000015582
ME3 / Q16798 / ENSMUSG00000030621 / ENSG00000151376 / ENSRNOG00000017311
MEA1 / Q16626 / ENSMUSG00000002768 / ENSG00000124733 / ENSRNOG00000017144
MEAF6 / Q9HAF1 / ENSMUSG00000028863 / ENSG00000163875 / ENSRNOG00000009309
MECP2 / P51608 / ENSMUSG00000031393 / ENSG00000169057 / ENSRNOG00000056659
MECR / Q9BV79 / ENSMUSG00000028910 / ENSG00000116353 / ENSRNOG00000028047
MED1 / Q15648 / ENSMUSG00000018160 / ENSG00000125686 / ENSRNOG00000005606
MED11 / Q9P086 / ENSMUSG00000018923 / ENSG00000161920 / ENSRNOG00000019384
MED12 / Q93074 / ENSMUSG00000079487 / ENSG00000184634 / ENSRNOG00000003848
MED14 / O60244 / ENSMUSG00000064127 / ENSG00000180182 / ENSRNOG00000003792
MED15 / Q96RN5 / ENSMUSG00000012114 / ENSG00000099917 / ENSRNOG00000001877
MED20 / Q9H944 / ENSMUSG00000092558 / ENSG00000124641 / ENSRNOG00000013852
MED23 / Q9ULK4 / ENSMUSG00000019984 / ENSG00000112282 / ENSRNOG00000013422
MED24 / O75448 / ENSMUSG00000017210 / ENSG00000008838 / ENSRNOG00000008711
MED25 / Q71SY5 / ENSMUSG00000002968 / ENSG00000104973 / ENSRNOG00000020363
MED28 / Q9H204 / ENSMUSG00000015804 / ENSG00000118579 / ENSRNOG00000003592
MED31 / Q9Y3C7 / ENSMUSG00000020801 / ENSG00000108590 / ENSRNOG00000014618
MEN1 / O00255 / ENSMUSG00000024947 / ENSG00000133895 / ENSRNOG00000021054
MEP1B / Q16820 / ENSMUSG00000024313 / ENSG00000141434 / ENSRNOG00000049345
MEPCE / Q7L2J0 / ENSMUSG00000029726 / ENSG00000146834 / ENSRNOG00000045911
MESD / Q14696 / ENSMUSG00000038503 / ENSG00000117899 / ENSRNOG00000012366
MET / P08581 / ENSMUSG00000009376 / ENSG00000105976 / ENSRNOG00000052745
METAP1 / P53582 / ENSMUSG00000005813 / ENSG00000164024 / ENSRNOG00000012947
METAP1D / Q6UB28 / ENSMUSG00000041921 / ENSG00000172878 / ENSRNOG00000061587
METAP2 / P50579 / ENSMUSG00000036112 / ENSG00000111142 / ENSRNOG00000021881
METTL14 / Q9HCE5 / ENSMUSG00000028114 / ENSG00000145388 / ENSRNOG00000015250
Homologs
Brain
Heart
Kidney
Liver
Lung
Testis
Organs
Bins
1
2
3
4
5
Species
Human
Mouse
Rat

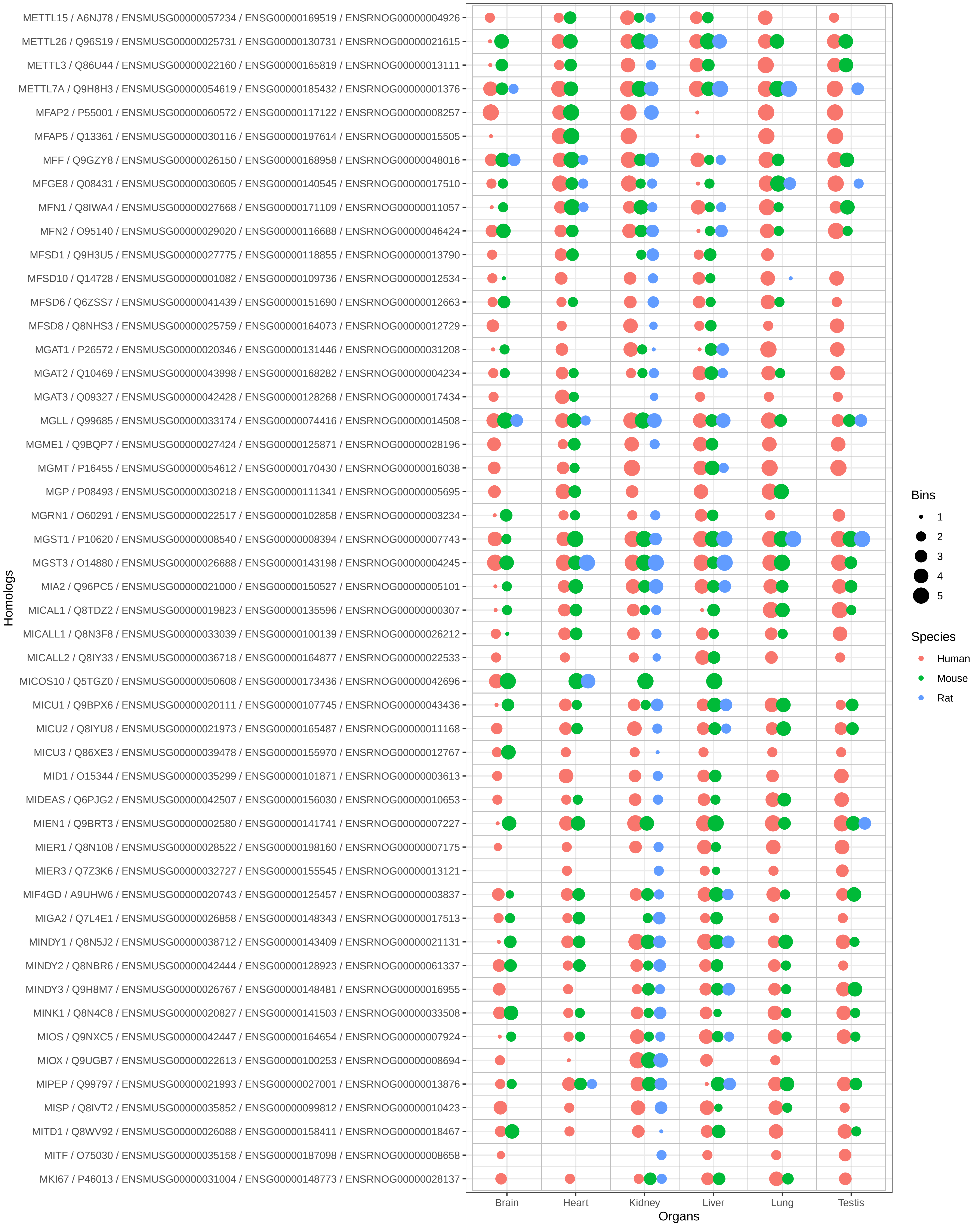
METTL15 / A6NJ78 / ENSMUSG00000057234 / ENSG00000169519 / ENSRNOG00000004926
METTL26 / Q96S19 / ENSMUSG00000025731 / ENSG00000130731 / ENSRNOG00000021615
METTL3 / Q86U44 / ENSMUSG00000022160 / ENSG00000165819 / ENSRNOG00000013111
METTL7A / Q9H8H3 / ENSMUSG00000054619 / ENSG00000185432 / ENSRNOG00000001376
MFAP2 / P55001 / ENSMUSG00000060572 / ENSG00000117122 / ENSRNOG00000008257
MFAP5 / Q13361 / ENSMUSG00000030116 / ENSG00000197614 / ENSRNOG00000015505
MFF / Q9GZY8 / ENSMUSG00000026150 / ENSG00000168958 / ENSRNOG00000048016
MFGE8 / Q08431 / ENSMUSG00000030605 / ENSG00000140545 / ENSRNOG00000017510
MFN1 / Q8IWA4 / ENSMUSG00000027668 / ENSG00000171109 / ENSRNOG00000011057
MFN2 / O95140 / ENSMUSG00000029020 / ENSG00000116688 / ENSRNOG00000046424
MFSD1 / Q9H3U5 / ENSMUSG00000027775 / ENSG00000118855 / ENSRNOG00000013790
MFSD10 / Q14728 / ENSMUSG00000001082 / ENSG00000109736 / ENSRNOG00000012534
MFSD6 / Q6ZSS7 / ENSMUSG00000041439 / ENSG00000151690 / ENSRNOG00000012663
MFSD8 / Q8NHS3 / ENSMUSG00000025759 / ENSG00000164073 / ENSRNOG00000012729
MGAT1 / P26572 / ENSMUSG00000020346 / ENSG00000131446 / ENSRNOG00000031208
MGAT2 / Q10469 / ENSMUSG00000043998 / ENSG00000168282 / ENSRNOG00000004234
MGAT3 / Q09327 / ENSMUSG00000042428 / ENSG00000128268 / ENSRNOG00000017434
MGLL / Q99685 / ENSMUSG00000033174 / ENSG00000074416 / ENSRNOG00000014508
MGME1 / Q9BQP7 / ENSMUSG00000027424 / ENSG00000125871 / ENSRNOG00000028196
MGMT / P16455 / ENSMUSG00000054612 / ENSG00000170430 / ENSRNOG00000016038
MGP / P08493 / ENSMUSG00000030218 / ENSG00000111341 / ENSRNOG00000005695
MGRN1 / O60291 / ENSMUSG00000022517 / ENSG00000102858 / ENSRNOG00000003234
MGST1 / P10620 / ENSMUSG00000008540 / ENSG00000008394 / ENSRNOG00000007743
MGST3 / O14880 / ENSMUSG00000026688 / ENSG00000143198 / ENSRNOG00000004245
MIA2 / Q96PC5 / ENSMUSG00000021000 / ENSG00000150527 / ENSRNOG00000005101
MICAL1 / Q8TDZ2 / ENSMUSG00000019823 / ENSG00000135596 / ENSRNOG00000000307
MICALL1 / Q8N3F8 / ENSMUSG00000033039 / ENSG00000100139 / ENSRNOG00000026212
MICALL2 / Q8IY33 / ENSMUSG00000036718 / ENSG00000164877 / ENSRNOG00000022533
MICOS10 / Q5TGZ0 / ENSMUSG00000050608 / ENSG00000173436 / ENSRNOG00000042696
MICU1 / Q9BPX6 / ENSMUSG00000020111 / ENSG00000107745 / ENSRNOG00000043436
MICU2 / Q8IYU8 / ENSMUSG00000021973 / ENSG00000165487 / ENSRNOG00000011168
MICU3 / Q86XE3 / ENSMUSG00000039478 / ENSG00000155970 / ENSRNOG00000012767
MID1 / O15344 / ENSMUSG00000035299 / ENSG00000101871 / ENSRNOG00000003613
MIDEAS / Q6PJG2 / ENSMUSG00000042507 / ENSG00000156030 / ENSRNOG00000010653
MIEN1 / Q9BRT3 / ENSMUSG00000002580 / ENSG00000141741 / ENSRNOG00000007227
MIER1 / Q8N108 / ENSMUSG00000028522 / ENSG00000198160 / ENSRNOG00000007175
MIER3 / Q7Z3K6 / ENSMUSG00000032727 / ENSG00000155545 / ENSRNOG00000013121
MIF4GD / A9UHW6 / ENSMUSG00000020743 / ENSG00000125457 / ENSRNOG00000003837
MIGA2 / Q7L4E1 / ENSMUSG00000026858 / ENSG00000148343 / ENSRNOG00000017513
MINDY1 / Q8N5J2 / ENSMUSG00000038712 / ENSG00000143409 / ENSRNOG00000021131
MINDY2 / Q8NBR6 / ENSMUSG00000042444 / ENSG00000128923 / ENSRNOG00000061337
MINDY3 / Q9H8M7 / ENSMUSG00000026767 / ENSG00000148481 / ENSRNOG00000016955
MINK1 / Q8N4C8 / ENSMUSG00000020827 / ENSG00000141503 / ENSRNOG00000033508
MIOS / Q9NXC5 / ENSMUSG00000042447 / ENSG00000164654 / ENSRNOG00000007924
MIOX / Q9UGB7 / ENSMUSG00000022613 / ENSG00000100253 / ENSRNOG00000008694
MIPEP / Q99797 / ENSMUSG00000021993 / ENSG00000027001 / ENSRNOG00000013876
MISP / Q8IVT2 / ENSMUSG00000035852 / ENSG00000099812 / ENSRNOG00000010423
MITD1 / Q8WV92 / ENSMUSG00000026088 / ENSG00000158411 / ENSRNOG00000018467
MITF / O75030 / ENSMUSG00000035158 / ENSG00000187098 / ENSRNOG00000008658
MKI67 / P46013 / ENSMUSG00000031004 / ENSG00000148773 / ENSRNOG00000028137
Homologs
Brain
Heart
Kidney
Liver
Lung
Testis
Organs
Bins
1
2
3
4
5
Species
Human
Mouse
Rat

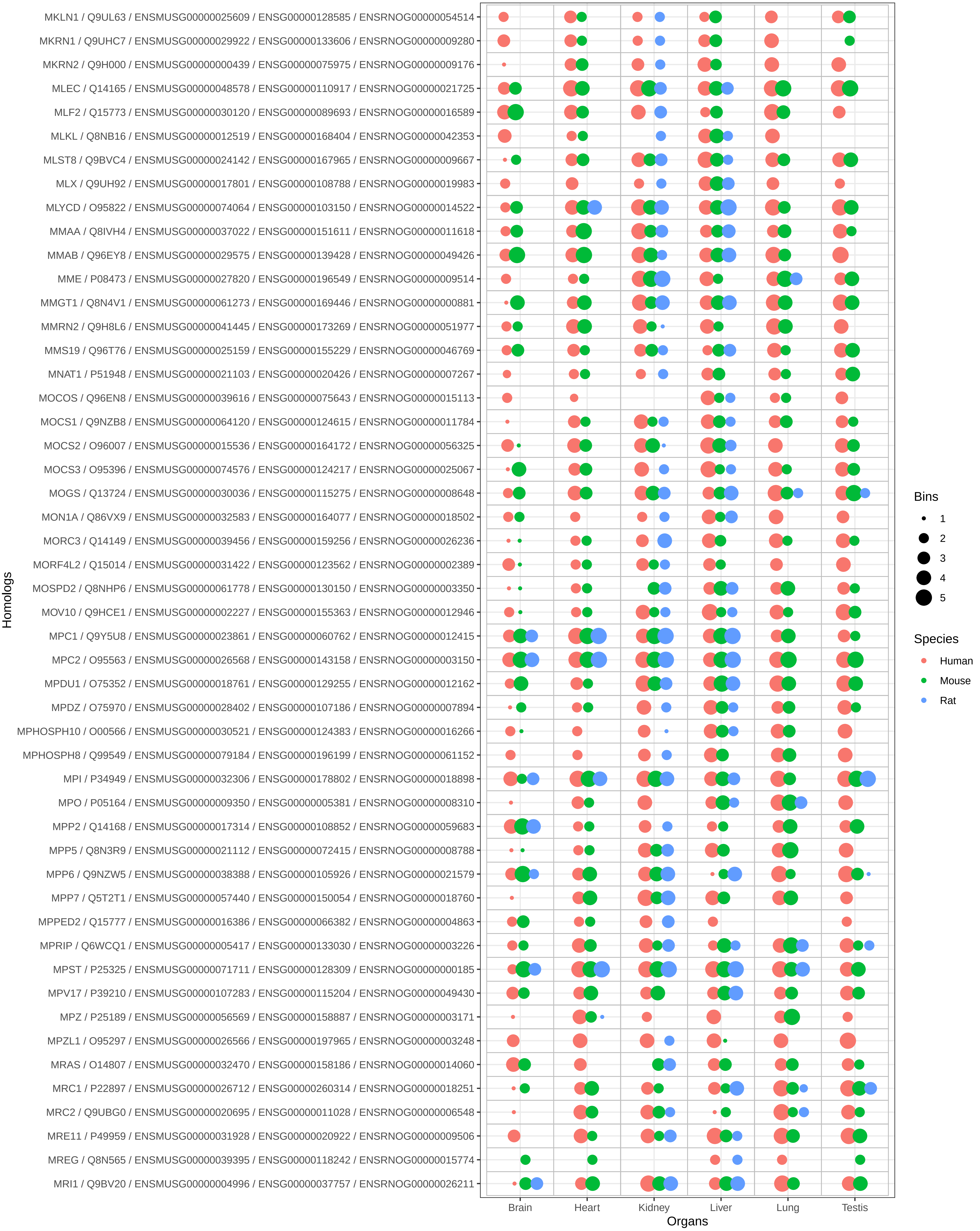
MKLN1 / Q9UL63 / ENSMUSG00000025609 / ENSG00000128585 / ENSRNOG00000054514
MKRN1 / Q9UHC7 / ENSMUSG00000029922 / ENSG00000133606 / ENSRNOG00000009280
MKRN2 / Q9H000 / ENSMUSG00000000439 / ENSG00000075975 / ENSRNOG00000009176
MLEC / Q14165 / ENSMUSG00000048578 / ENSG00000110917 / ENSRNOG00000021725
MLF2 / Q15773 / ENSMUSG00000030120 / ENSG00000089693 / ENSRNOG00000016589
MLKL / Q8NB16 / ENSMUSG00000012519 / ENSG00000168404 / ENSRNOG00000042353
MLST8 / Q9BVC4 / ENSMUSG00000024142 / ENSG00000167965 / ENSRNOG00000009667
MLX / Q9UH92 / ENSMUSG00000017801 / ENSG00000108788 / ENSRNOG00000019983
MLYCD / O95822 / ENSMUSG00000074064 / ENSG00000103150 / ENSRNOG00000014522
MMAA / Q8IVH4 / ENSMUSG00000037022 / ENSG00000151611 / ENSRNOG00000011618
MMAB / Q96EY8 / ENSMUSG00000029575 / ENSG00000139428 / ENSRNOG00000049426
MME / P08473 / ENSMUSG00000027820 / ENSG00000196549 / ENSRNOG00000009514
MMGT1 / Q8N4V1 / ENSMUSG00000061273 / ENSG00000169446 / ENSRNOG00000000881
MMRN2 / Q9H8L6 / ENSMUSG00000041445 / ENSG00000173269 / ENSRNOG00000051977
MMS19 / Q96T76 / ENSMUSG00000025159 / ENSG00000155229 / ENSRNOG00000046769
MNAT1 / P51948 / ENSMUSG00000021103 / ENSG00000020426 / ENSRNOG00000007267
MOCOS / Q96EN8 / ENSMUSG00000039616 / ENSG00000075643 / ENSRNOG00000015113
MOCS1 / Q9NZB8 / ENSMUSG00000064120 / ENSG00000124615 / ENSRNOG00000011784
MOCS2 / O96007 / ENSMUSG00000015536 / ENSG00000164172 / ENSRNOG00000056325
MOCS3 / O95396 / ENSMUSG00000074576 / ENSG00000124217 / ENSRNOG00000025067
MOGS / Q13724 / ENSMUSG00000030036 / ENSG00000115275 / ENSRNOG00000008648
MON1A / Q86VX9 / ENSMUSG00000032583 / ENSG00000164077 / ENSRNOG00000018502
MORC3 / Q14149 / ENSMUSG00000039456 / ENSG00000159256 / ENSRNOG00000026236
MORF4L2 / Q15014 / ENSMUSG00000031422 / ENSG00000123562 / ENSRNOG00000002389
MOSPD2 / Q8NHP6 / ENSMUSG00000061778 / ENSG00000130150 / ENSRNOG00000003350
MOV10 / Q9HCE1 / ENSMUSG00000002227 / ENSG00000155363 / ENSRNOG00000012946
MPC1 / Q9Y5U8 / ENSMUSG00000023861 / ENSG00000060762 / ENSRNOG00000012415
MPC2 / O95563 / ENSMUSG00000026568 / ENSG00000143158 / ENSRNOG00000003150
MPDU1 / O75352 / ENSMUSG00000018761 / ENSG00000129255 / ENSRNOG00000012162
MPDZ / O75970 / ENSMUSG00000028402 / ENSG00000107186 / ENSRNOG00000007894
MPHOSPH10 / O00566 / ENSMUSG00000030521 / ENSG00000124383 / ENSRNOG00000016266
MPHOSPH8 / Q99549 / ENSMUSG00000079184 / ENSG00000196199 / ENSRNOG00000061152
MPI / P34949 / ENSMUSG00000032306 / ENSG00000178802 / ENSRNOG00000018898
MPO / P05164 / ENSMUSG00000009350 / ENSG00000005381 / ENSRNOG00000008310
MPP2 / Q14168 / ENSMUSG00000017314 / ENSG00000108852 / ENSRNOG00000059683
MPP5 / Q8N3R9 / ENSMUSG00000021112 / ENSG00000072415 / ENSRNOG00000008788
MPP6 / Q9NZW5 / ENSMUSG00000038388 / ENSG00000105926 / ENSRNOG00000021579
MPP7 / Q5T2T1 / ENSMUSG00000057440 / ENSG00000150054 / ENSRNOG00000018760
MPPED2 / Q15777 / ENSMUSG00000016386 / ENSG00000066382 / ENSRNOG00000004863
MPRIP / Q6WCQ1 / ENSMUSG00000005417 / ENSG00000133030 / ENSRNOG00000003226
MPST / P25325 / ENSMUSG00000071711 / ENSG00000128309 / ENSRNOG00000000185
MPV17 / P39210 / ENSMUSG00000107283 / ENSG00000115204 / ENSRNOG00000049430
MPZ / P25189 / ENSMUSG00000056569 / ENSG00000158887 / ENSRNOG00000003171
MPZL1 / O95297 / ENSMUSG00000026566 / ENSG00000197965 / ENSRNOG00000003248
MRAS / O14807 / ENSMUSG00000032470 / ENSG00000158186 / ENSRNOG00000014060
MRC1 / P22897 / ENSMUSG00000026712 / ENSG00000260314 / ENSRNOG00000018251
MRC2 / Q9UBG0 / ENSMUSG00000020695 / ENSG00000011028 / ENSRNOG00000006548
MRE11 / P49959 / ENSMUSG00000031928 / ENSG00000020922 / ENSRNOG00000009506
MREG / Q8N565 / ENSMUSG00000039395 / ENSG00000118242 / ENSRNOG00000015774
MRI1 / Q9BV20 / ENSMUSG00000004996 / ENSG00000037757 / ENSRNOG00000026211
Homologs
Brain
Heart
Kidney
Liver
Lung
Testis
Organs
Bins
1
2
3
4
5
Species
Human
Mouse
Rat

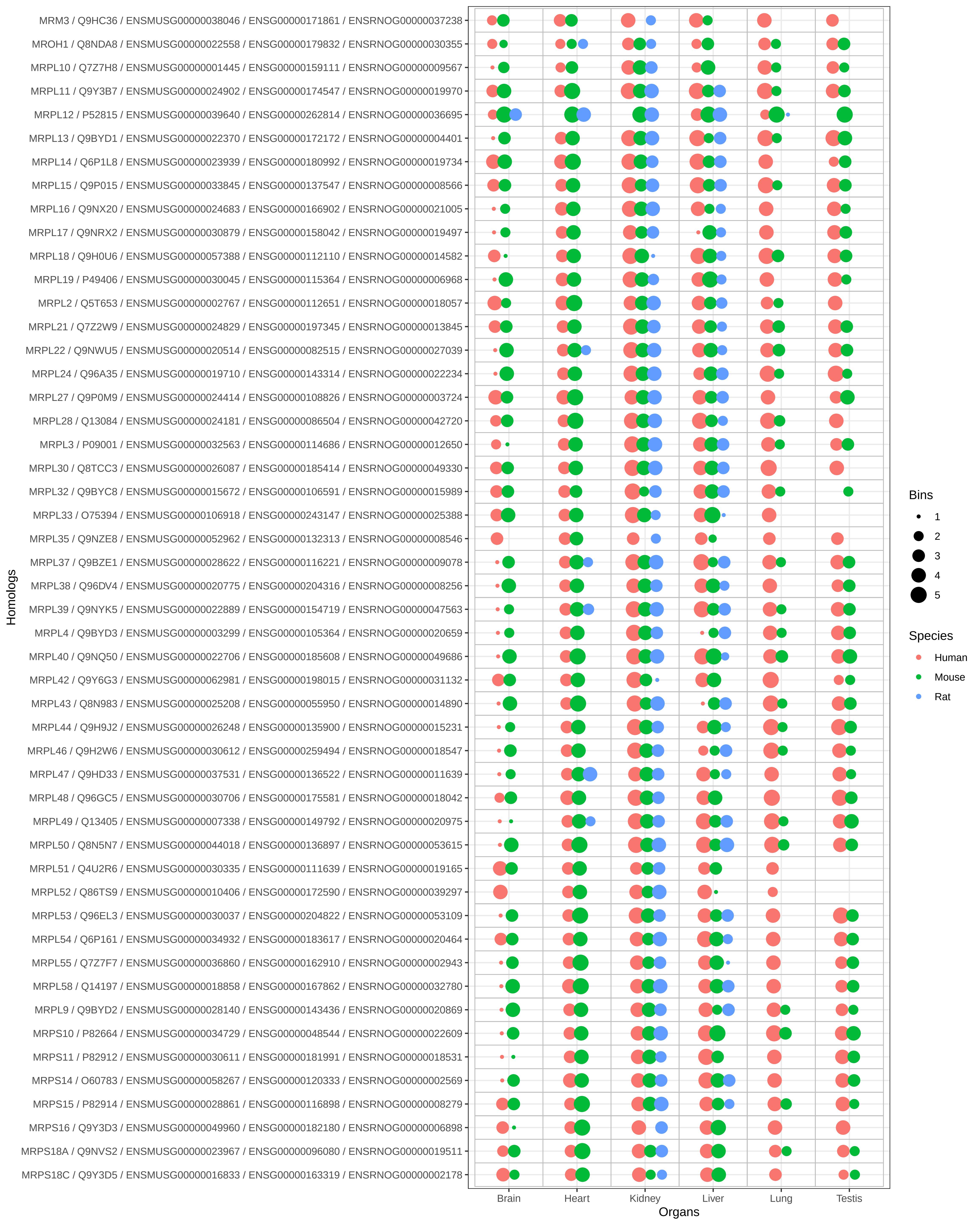
MRM3 / Q9HC36 / ENSMUSG00000038046 / ENSG00000171861 / ENSRNOG00000037238
MROH1 / Q8NDA8 / ENSMUSG00000022558 / ENSG00000179832 / ENSRNOG00000030355
MRPL10 / Q7Z7H8 / ENSMUSG00000001445 / ENSG00000159111 / ENSRNOG00000009567
MRPL11 / Q9Y3B7 / ENSMUSG00000024902 / ENSG00000174547 / ENSRNOG00000019970
MRPL12 / P52815 / ENSMUSG00000039640 / ENSG00000262814 / ENSRNOG00000036695
MRPL13 / Q9BYD1 / ENSMUSG00000022370 / ENSG00000172172 / ENSRNOG00000004401
MRPL14 / Q6P1L8 / ENSMUSG00000023939 / ENSG00000180992 / ENSRNOG00000019734
MRPL15 / Q9P015 / ENSMUSG00000033845 / ENSG00000137547 / ENSRNOG00000008566
MRPL16 / Q9NX20 / ENSMUSG00000024683 / ENSG00000166902 / ENSRNOG00000021005
MRPL17 / Q9NRX2 / ENSMUSG00000030879 / ENSG00000158042 / ENSRNOG00000019497
MRPL18 / Q9H0U6 / ENSMUSG00000057388 / ENSG00000112110 / ENSRNOG00000014582
MRPL19 / P49406 / ENSMUSG00000030045 / ENSG00000115364 / ENSRNOG00000006968
MRPL2 / Q5T653 / ENSMUSG00000002767 / ENSG00000112651 / ENSRNOG00000018057
MRPL21 / Q7Z2W9 / ENSMUSG00000024829 / ENSG00000197345 / ENSRNOG00000013845
MRPL22 / Q9NWU5 / ENSMUSG00000020514 / ENSG00000082515 / ENSRNOG00000027039
MRPL24 / Q96A35 / ENSMUSG00000019710 / ENSG00000143314 / ENSRNOG00000022234
MRPL27 / Q9P0M9 / ENSMUSG00000024414 / ENSG00000108826 / ENSRNOG00000003724
MRPL28 / Q13084 / ENSMUSG00000024181 / ENSG00000086504 / ENSRNOG00000042720
MRPL3 / P09001 / ENSMUSG00000032563 / ENSG00000114686 / ENSRNOG00000012650
MRPL30 / Q8TCC3 / ENSMUSG00000026087 / ENSG00000185414 / ENSRNOG00000049330
MRPL32 / Q9BYC8 / ENSMUSG00000015672 / ENSG00000106591 / ENSRNOG00000015989
MRPL33 / O75394 / ENSMUSG00000106918 / ENSG00000243147 / ENSRNOG00000025388
MRPL35 / Q9NZE8 / ENSMUSG00000052962 / ENSG00000132313 / ENSRNOG00000008546
MRPL37 / Q9BZE1 / ENSMUSG00000028622 / ENSG00000116221 / ENSRNOG00000009078
MRPL38 / Q96DV4 / ENSMUSG00000020775 / ENSG00000204316 / ENSRNOG00000008256
MRPL39 / Q9NYK5 / ENSMUSG00000022889 / ENSG00000154719 / ENSRNOG00000047563
MRPL4 / Q9BYD3 / ENSMUSG00000003299 / ENSG00000105364 / ENSRNOG00000020659
MRPL40 / Q9NQ50 / ENSMUSG00000022706 / ENSG00000185608 / ENSRNOG00000049686
MRPL42 / Q9Y6G3 / ENSMUSG00000062981 / ENSG00000198015 / ENSRNOG00000031132
MRPL43 / Q8N983 / ENSMUSG00000025208 / ENSG00000055950 / ENSRNOG00000014890
MRPL44 / Q9H9J2 / ENSMUSG00000026248 / ENSG00000135900 / ENSRNOG00000015231
MRPL46 / Q9H2W6 / ENSMUSG00000030612 / ENSG00000259494 / ENSRNOG00000018547
MRPL47 / Q9HD33 / ENSMUSG00000037531 / ENSG00000136522 / ENSRNOG00000011639
MRPL48 / Q96GC5 / ENSMUSG00000030706 / ENSG00000175581 / ENSRNOG00000018042
MRPL49 / Q13405 / ENSMUSG00000007338 / ENSG00000149792 / ENSRNOG00000020975
MRPL50 / Q8N5N7 / ENSMUSG00000044018 / ENSG00000136897 / ENSRNOG00000053615
MRPL51 / Q4U2R6 / ENSMUSG00000030335 / ENSG00000111639 / ENSRNOG00000019165
MRPL52 / Q86TS9 / ENSMUSG00000010406 / ENSG00000172590 / ENSRNOG00000039297
MRPL53 / Q96EL3 / ENSMUSG00000030037 / ENSG00000204822 / ENSRNOG00000053109
MRPL54 / Q6P161 / ENSMUSG00000034932 / ENSG00000183617 / ENSRNOG00000020464
MRPL55 / Q7Z7F7 / ENSMUSG00000036860 / ENSG00000162910 / ENSRNOG00000002943
MRPL58 / Q14197 / ENSMUSG00000018858 / ENSG00000167862 / ENSRNOG00000032780
MRPL9 / Q9BYD2 / ENSMUSG00000028140 / ENSG00000143436 / ENSRNOG00000020869
MRPS10 / P82664 / ENSMUSG00000034729 / ENSG00000048544 / ENSRNOG00000022609
MRPS11 / P82912 / ENSMUSG00000030611 / ENSG00000181991 / ENSRNOG00000018531
MRPS14 / O60783 / ENSMUSG00000058267 / ENSG00000120333 / ENSRNOG00000002569
MRPS15 / P82914 / ENSMUSG00000028861 / ENSG00000116898 / ENSRNOG00000008279
MRPS16 / Q9Y3D3 / ENSMUSG00000049960 / ENSG00000182180 / ENSRNOG00000006898
MRPS18A / Q9NVS2 / ENSMUSG00000023967 / ENSG00000096080 / ENSRNOG00000019511
MRPS18C / Q9Y3D5 / ENSMUSG00000016833 / ENSG00000163319 / ENSRNOG00000002178
Homologs
Brain
Heart
Kidney
Liver
Lung
Testis
Organs
Bins
1
2
3
4
5
Species
Human
Mouse
Rat

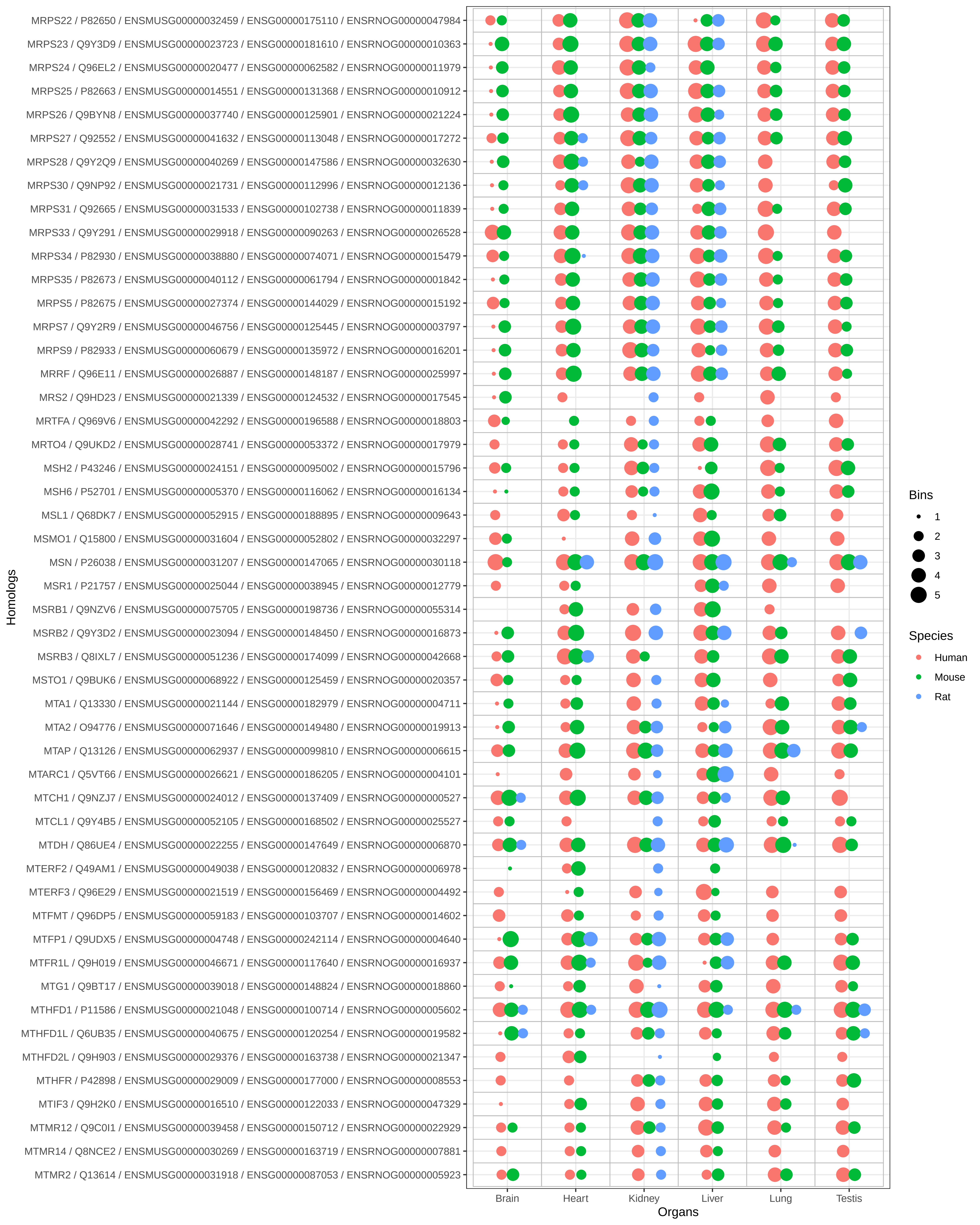

MRPS22 / P82650 / ENSMUSG00000032459 / ENSG00000175110 / ENSRNOG00000047984
MRPS23 / Q9Y3D9 / ENSMUSG00000023723 / ENSG00000181610 / ENSRNOG00000010363
MRPS24 / Q96EL2 / ENSMUSG00000020477 / ENSG00000062582 / ENSRNOG00000011979
MRPS25 / P82663 / ENSMUSG00000014551 / ENSG00000131368 / ENSRNOG00000010912
MRPS26 / Q9BYN8 / ENSMUSG00000037740 / ENSG00000125901 / ENSRNOG00000021224
MRPS27 / Q92552 / ENSMUSG00000041632 / ENSG00000113048 / ENSRNOG00000017272
MRPS28 / Q9Y2Q9 / ENSMUSG00000040269 / ENSG00000147586 / ENSRNOG00000032630
MRPS30 / Q9NP92 / ENSMUSG00000021731 / ENSG00000112996 / ENSRNOG00000012136
MRPS31 / Q92665 / ENSMUSG00000031533 / ENSG00000102738 / ENSRNOG00000011839
MRPS33 / Q9Y291 / ENSMUSG00000029918 / ENSG00000090263 / ENSRNOG00000026528
MRPS34 / P82930 / ENSMUSG00000038880 / ENSG00000074071 / ENSRNOG00000015479
MRPS35 / P82673 / ENSMUSG00000040112 / ENSG00000061794 / ENSRNOG00000001842
MRPS5 / P82675 / ENSMUSG00000027374 / ENSG00000144029 / ENSRNOG00000015192
MRPS7 / Q9Y2R9 / ENSMUSG00000046756 / ENSG00000125445 / ENSRNOG00000003797
MRPS9 / P82933 / ENSMUSG00000060679 / ENSG00000135972 / ENSRNOG00000016201
MRRF / Q96E11 / ENSMUSG00000026887 / ENSG00000148187 / ENSRNOG00000025997
MRS2 / Q9HD23 / ENSMUSG00000021339 / ENSG00000124532 / ENSRNOG00000017545
MRTFA / Q969V6 / ENSMUSG00000042292 / ENSG00000196588 / ENSRNOG00000018803
MRTO4 / Q9UKD2 / ENSMUSG00000028741 / ENSG00000053372 / ENSRNOG00000017979
MSH2 / P43246 / ENSMUSG00000024151 / ENSG00000095002 / ENSRNOG00000015796
MSH6 / P52701 / ENSMUSG00000005370 / ENSG00000116062 / ENSRNOG00000016134
MSL1 / Q68DK7 / ENSMUSG00000052915 / ENSG00000188895 / ENSRNOG00000009643
MSMO1 / Q15800 / ENSMUSG00000031604 / ENSG00000052802 / ENSRNOG00000032297
MSN / P26038 / ENSMUSG00000031207 / ENSG00000147065 / ENSRNOG00000030118
MSR1 / P21757 / ENSMUSG00000025044 / ENSG00000038945 / ENSRNOG00000012779
MSRB1 / Q9NZV6 / ENSMUSG00000075705 / ENSG00000198736 / ENSRNOG00000055314
MSRB2 / Q9Y3D2 / ENSMUSG00000023094 / ENSG00000148450 / ENSRNOG00000016873
MSRB3 / Q8IXL7 / ENSMUSG00000051236 / ENSG00000174099 / ENSRNOG00000042668
MSTO1 / Q9BUK6 / ENSMUSG00000068922 / ENSG00000125459 / ENSRNOG00000020357
MTA1 / Q13330 / ENSMUSG00000021144 / ENSG00000182979 / ENSRNOG00000004711
MTA2 / O94776 / ENSMUSG00000071646 / ENSG00000149480 / ENSRNOG00000019913
MTAP / Q13126 / ENSMUSG00000062937 / ENSG00000099810 / ENSRNOG00000006615
MTARC1 / Q5VT66 / ENSMUSG00000026621 / ENSG00000186205 / ENSRNOG00000004101
MTCH1 / Q9NZJ7 / ENSMUSG00000024012 / ENSG00000137409 / ENSRNOG00000000527
MTCL1 / Q9Y4B5 / ENSMUSG00000052105 / ENSG00000168502 / ENSRNOG00000025527
MTDH / Q86UE4 / ENSMUSG00000022255 / ENSG00000147649 / ENSRNOG00000006870
MTERF2 / Q49AM1 / ENSMUSG00000049038 / ENSG00000120832 / ENSRNOG00000006978
MTERF3 / Q96E29 / ENSMUSG00000021519 / ENSG00000156469 / ENSRNOG00000004492
MTFMT / Q96DP5 / ENSMUSG00000059183 / ENSG00000103707 / ENSRNOG00000014602
MTFP1 / Q9UDX5 / ENSMUSG00000004748 / ENSG00000242114 / ENSRNOG00000004640
MTFR1L / Q9H019 / ENSMUSG00000046671 / ENSG00000117640 / ENSRNOG00000016937
MTG1 / Q9BT17 / ENSMUSG00000039018 / ENSG00000148824 / ENSRNOG00000018860
MTHFD1 / P11586 / ENSMUSG00000021048 / ENSG00000100714 / ENSRNOG00000005602
MTHFD1L / Q6UB35 / ENSMUSG00000040675 / ENSG00000120254 / ENSRNOG00000019582
MTHFD2L / Q9H903 / ENSMUSG00000029376 / ENSG00000163738 / ENSRNOG00000021347
MTHFR / P42898 / ENSMUSG00000029009 / ENSG00000177000 / ENSRNOG00000008553
MTIF3 / Q9H2K0 / ENSMUSG00000016510 / ENSG00000122033 / ENSRNOG00000047329
MTMR12 / Q9C0I1 / ENSMUSG00000039458 / ENSG00000150712 / ENSRNOG00000022929
MTMR14 / Q8NCE2 / ENSMUSG00000030269 / ENSG00000163719 / ENSRNOG00000007881
MTMR2 / Q13614 / ENSMUSG00000031918 / ENSG00000087053 / ENSRNOG00000005923
Homologs
Brain
Heart
Kidney
Liver
Lung
Testis
Organs
Bins
1
2
3
4
5
Species
Human
Mouse
Rat

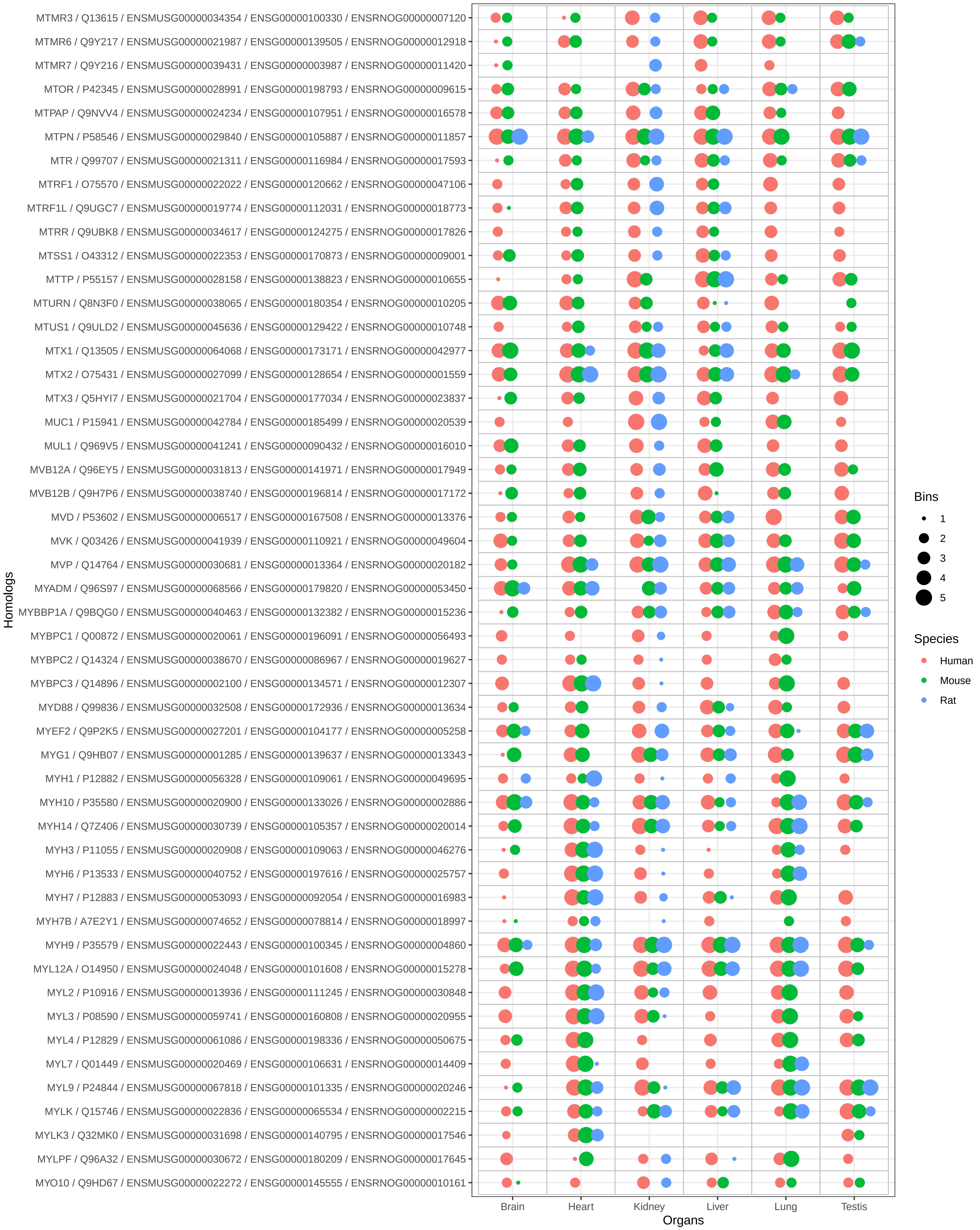
MTMR3 / Q13615 / ENSMUSG00000034354 / ENSG00000100330 / ENSRNOG00000007120
MTMR6 / Q9Y217 / ENSMUSG00000021987 / ENSG00000139505 / ENSRNOG00000012918
MTMR7 / Q9Y216 / ENSMUSG00000039431 / ENSG00000003987 / ENSRNOG00000011420
MTOR / P42345 / ENSMUSG00000028991 / ENSG00000198793 / ENSRNOG00000009615
MTPAP / Q9NVV4 / ENSMUSG00000024234 / ENSG00000107951 / ENSRNOG00000016578
MTPN / P58546 / ENSMUSG00000029840 / ENSG00000105887 / ENSRNOG00000011857
MTR / Q99707 / ENSMUSG00000021311 / ENSG00000116984 / ENSRNOG00000017593
MTRF1 / O75570 / ENSMUSG00000022022 / ENSG00000120662 / ENSRNOG00000047106
MTRF1L / Q9UGC7 / ENSMUSG00000019774 / ENSG00000112031 / ENSRNOG00000018773
MTRR / Q9UBK8 / ENSMUSG00000034617 / ENSG00000124275 / ENSRNOG00000017826
MTSS1 / O43312 / ENSMUSG00000022353 / ENSG00000170873 / ENSRNOG00000009001
MTTP / P55157 / ENSMUSG00000028158 / ENSG00000138823 / ENSRNOG00000010655
MTURN / Q8N3F0 / ENSMUSG00000038065 / ENSG00000180354 / ENSRNOG00000010205
MTUS1 / Q9ULD2 / ENSMUSG00000045636 / ENSG00000129422 / ENSRNOG00000010748
MTX1 / Q13505 / ENSMUSG00000064068 / ENSG00000173171 / ENSRNOG00000042977
MTX2 / O75431 / ENSMUSG00000027099 / ENSG00000128654 / ENSRNOG00000001559
MTX3 / Q5HYI7 / ENSMUSG00000021704 / ENSG00000177034 / ENSRNOG00000023837
MUC1 / P15941 / ENSMUSG00000042784 / ENSG00000185499 / ENSRNOG00000020539
MUL1 / Q969V5 / ENSMUSG00000041241 / ENSG00000090432 / ENSRNOG00000016010
MVB12A / Q96EY5 / ENSMUSG00000031813 / ENSG00000141971 / ENSRNOG00000017949
MVB12B / Q9H7P6 / ENSMUSG00000038740 / ENSG00000196814 / ENSRNOG00000017172
MVD / P53602 / ENSMUSG00000006517 / ENSG00000167508 / ENSRNOG00000013376
MVK / Q03426 / ENSMUSG00000041939 / ENSG00000110921 / ENSRNOG00000049604
MVP / Q14764 / ENSMUSG00000030681 / ENSG00000013364 / ENSRNOG00000020182
MYADM / Q96S97 / ENSMUSG00000068566 / ENSG00000179820 / ENSRNOG00000053450
MYBBP1A / Q9BQG0 / ENSMUSG00000040463 / ENSG00000132382 / ENSRNOG00000015236
MYBPC1 / Q00872 / ENSMUSG00000020061 / ENSG00000196091 / ENSRNOG00000056493
MYBPC2 / Q14324 / ENSMUSG00000038670 / ENSG00000086967 / ENSRNOG00000019627
MYBPC3 / Q14896 / ENSMUSG00000002100 / ENSG00000134571 / ENSRNOG00000012307
MYD88 / Q99836 / ENSMUSG00000032508 / ENSG00000172936 / ENSRNOG00000013634
MYEF2 / Q9P2K5 / ENSMUSG00000027201 / ENSG00000104177 / ENSRNOG00000005258
MYG1 / Q9HB07 / ENSMUSG00000001285 / ENSG00000139637 / ENSRNOG00000013343
MYH1 / P12882 / ENSMUSG00000056328 / ENSG00000109061 / ENSRNOG00000049695
MYH10 / P35580 / ENSMUSG00000020900 / ENSG00000133026 / ENSRNOG00000002886
MYH14 / Q7Z406 / ENSMUSG00000030739 / ENSG00000105357 / ENSRNOG00000020014
MYH3 / P11055 / ENSMUSG00000020908 / ENSG00000109063 / ENSRNOG00000046276
MYH6 / P13533 / ENSMUSG00000040752 / ENSG00000197616 / ENSRNOG00000025757
MYH7 / P12883 / ENSMUSG00000053093 / ENSG00000092054 / ENSRNOG00000016983
MYH7B / A7E2Y1 / ENSMUSG00000074652 / ENSG00000078814 / ENSRNOG00000018997
MYH9 / P35579 / ENSMUSG00000022443 / ENSG00000100345 / ENSRNOG00000004860
MYL12A / O14950 / ENSMUSG00000024048 / ENSG00000101608 / ENSRNOG00000015278
MYL2 / P10916 / ENSMUSG00000013936 / ENSG00000111245 / ENSRNOG00000030848
MYL3 / P08590 / ENSMUSG00000059741 / ENSG00000160808 / ENSRNOG00000020955
MYL4 / P12829 / ENSMUSG00000061086 / ENSG00000198336 / ENSRNOG00000050675
MYL7 / Q01449 / ENSMUSG00000020469 / ENSG00000106631 / ENSRNOG00000014409
MYL9 / P24844 / ENSMUSG00000067818 / ENSG00000101335 / ENSRNOG00000020246
MYLK / Q15746 / ENSMUSG00000022836 / ENSG00000065534 / ENSRNOG00000002215
MYLK3 / Q32MK0 / ENSMUSG00000031698 / ENSG00000140795 / ENSRNOG00000017546
MYLPF / Q96A32 / ENSMUSG00000030672 / ENSG00000180209 / ENSRNOG00000017645
MYO10 / Q9HD67 / ENSMUSG00000022272 / ENSG00000145555 / ENSRNOG00000010161
Homologs
Brain
Heart
Kidney
Liver
Lung
Testis
Organs
Bins
1
2
3
4
5
Species
Human
Mouse
Rat

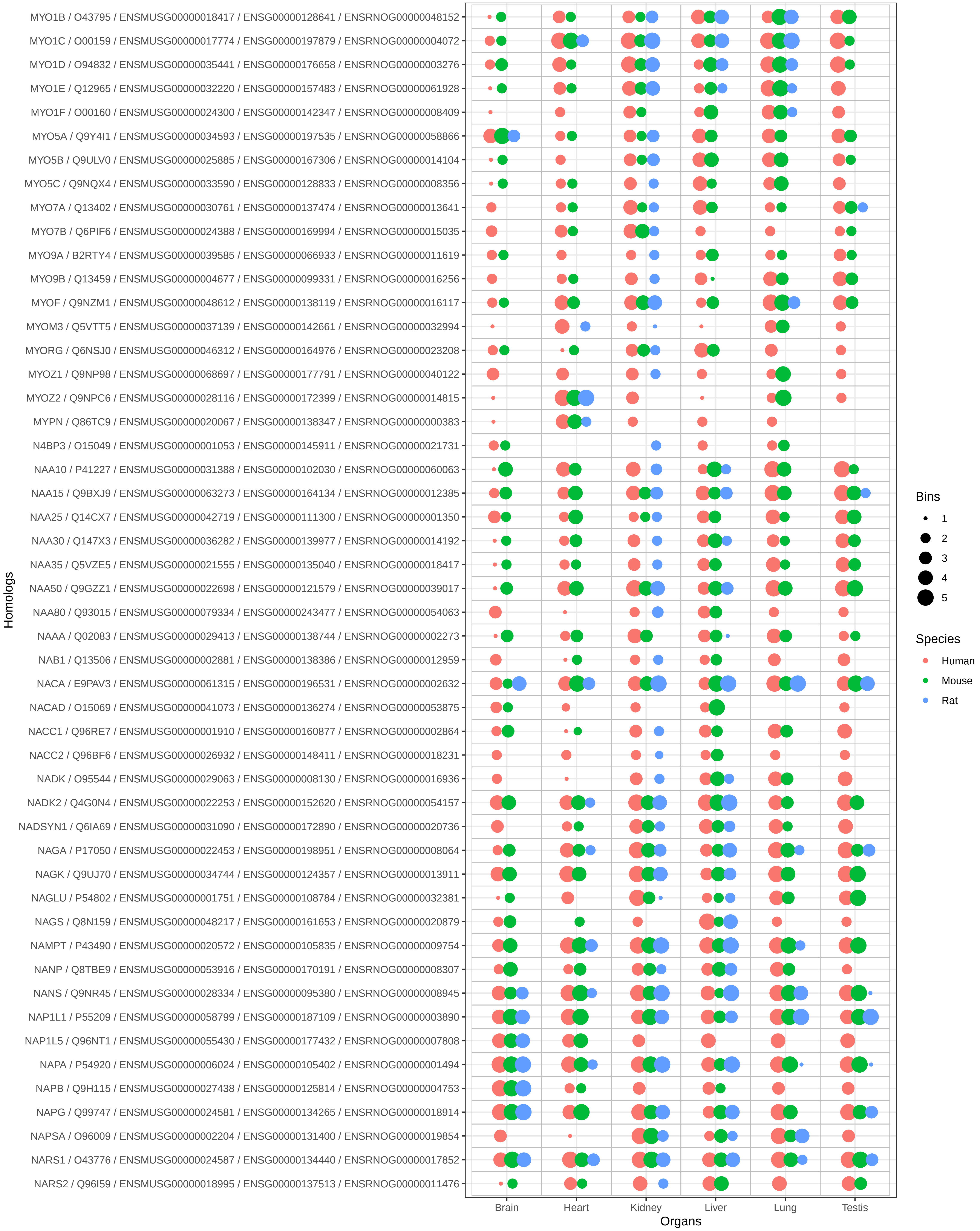
MYO1B / O43795 / ENSMUSG00000018417 / ENSG00000128641 / ENSRNOG00000048152
MYO1C / O00159 / ENSMUSG00000017774 / ENSG00000197879 / ENSRNOG00000004072
MYO1D / O94832 / ENSMUSG00000035441 / ENSG00000176658 / ENSRNOG00000003276
MYO1E / Q12965 / ENSMUSG00000032220 / ENSG00000157483 / ENSRNOG00000061928
MYO1F / O00160 / ENSMUSG00000024300 / ENSG00000142347 / ENSRNOG00000008409
MYO5A / Q9Y4I1 / ENSMUSG00000034593 / ENSG00000197535 / ENSRNOG00000058866
MYO5B / Q9ULV0 / ENSMUSG00000025885 / ENSG00000167306 / ENSRNOG00000014104
MYO5C / Q9NQX4 / ENSMUSG00000033590 / ENSG00000128833 / ENSRNOG00000008356
MYO7A / Q13402 / ENSMUSG00000030761 / ENSG00000137474 / ENSRNOG00000013641
MYO7B / Q6PIF6 / ENSMUSG00000024388 / ENSG00000169994 / ENSRNOG00000015035
MYO9A / B2RTY4 / ENSMUSG00000039585 / ENSG00000066933 / ENSRNOG00000011619
MYO9B / Q13459 / ENSMUSG00000004677 / ENSG00000099331 / ENSRNOG00000016256
MYOF / Q9NZM1 / ENSMUSG00000048612 / ENSG00000138119 / ENSRNOG00000016117
MYOM3 / Q5VTT5 / ENSMUSG00000037139 / ENSG00000142661 / ENSRNOG00000032994
MYORG / Q6NSJ0 / ENSMUSG00000046312 / ENSG00000164976 / ENSRNOG00000023208
MYOZ1 / Q9NP98 / ENSMUSG00000068697 / ENSG00000177791 / ENSRNOG00000040122
MYOZ2 / Q9NPC6 / ENSMUSG00000028116 / ENSG00000172399 / ENSRNOG00000014815
MYPN / Q86TC9 / ENSMUSG00000020067 / ENSG00000138347 / ENSRNOG00000000383
N4BP3 / O15049 / ENSMUSG00000001053 / ENSG00000145911 / ENSRNOG00000021731
NAA10 / P41227 / ENSMUSG00000031388 / ENSG00000102030 / ENSRNOG00000060063
NAA15 / Q9BXJ9 / ENSMUSG00000063273 / ENSG00000164134 / ENSRNOG00000012385
NAA25 / Q14CX7 / ENSMUSG00000042719 / ENSG00000111300 / ENSRNOG00000001350
NAA30 / Q147X3 / ENSMUSG00000036282 / ENSG00000139977 / ENSRNOG00000014192
NAA35 / Q5VZE5 / ENSMUSG00000021555 / ENSG00000135040 / ENSRNOG00000018417
NAA50 / Q9GZZ1 / ENSMUSG00000022698 / ENSG00000121579 / ENSRNOG00000039017
NAA80 / Q93015 / ENSMUSG00000079334 / ENSG00000243477 / ENSRNOG00000054063
NAAA / Q02083 / ENSMUSG00000029413 / ENSG00000138744 / ENSRNOG00000002273
NAB1 / Q13506 / ENSMUSG00000002881 / ENSG00000138386 / ENSRNOG00000012959
NACA / E9PAV3 / ENSMUSG00000061315 / ENSG00000196531 / ENSRNOG00000002632
NACAD / O15069 / ENSMUSG00000041073 / ENSG00000136274 / ENSRNOG00000053875
NACC1 / Q96RE7 / ENSMUSG00000001910 / ENSG00000160877 / ENSRNOG00000002864
NACC2 / Q96BF6 / ENSMUSG00000026932 / ENSG00000148411 / ENSRNOG00000018231
NADK / O95544 / ENSMUSG00000029063 / ENSG00000008130 / ENSRNOG00000016936
NADK2 / Q4G0N4 / ENSMUSG00000022253 / ENSG00000152620 / ENSRNOG00000054157
NADSYN1 / Q6IA69 / ENSMUSG00000031090 / ENSG00000172890 / ENSRNOG00000020736
NAGA / P17050 / ENSMUSG00000022453 / ENSG00000198951 / ENSRNOG00000008064
NAGK / Q9UJ70 / ENSMUSG00000034744 / ENSG00000124357 / ENSRNOG00000013911
NAGLU / P54802 / ENSMUSG00000001751 / ENSG00000108784 / ENSRNOG00000032381
NAGS / Q8N159 / ENSMUSG00000048217 / ENSG00000161653 / ENSRNOG00000020879
NAMPT / P43490 / ENSMUSG00000020572 / ENSG00000105835 / ENSRNOG00000009754
NANP / Q8TBE9 / ENSMUSG00000053916 / ENSG00000170191 / ENSRNOG00000008307
NANS / Q9NR45 / ENSMUSG00000028334 / ENSG00000095380 / ENSRNOG00000008945
NAP1L1 / P55209 / ENSMUSG00000058799 / ENSG00000187109 / ENSRNOG00000003890
NAP1L5 / Q96NT1 / ENSMUSG00000055430 / ENSG00000177432 / ENSRNOG00000007808
NAPA / P54920 / ENSMUSG00000006024 / ENSG00000105402 / ENSRNOG00000001494
NAPB / Q9H115 / ENSMUSG00000027438 / ENSG00000125814 / ENSRNOG00000004753
NAPG / Q99747 / ENSMUSG00000024581 / ENSG00000134265 / ENSRNOG00000018914
NAPSA / O96009 / ENSMUSG00000002204 / ENSG00000131400 / ENSRNOG00000019854
NARS1 / O43776 / ENSMUSG00000024587 / ENSG00000134440 / ENSRNOG00000017852
NARS2 / Q96I59 / ENSMUSG00000018995 / ENSG00000137513 / ENSRNOG00000011476
Homologs
Brain
Heart
Kidney
Liver
Lung
Testis
Organs
Bins
1
2
3
4
5
Species
Human
Mouse
Rat

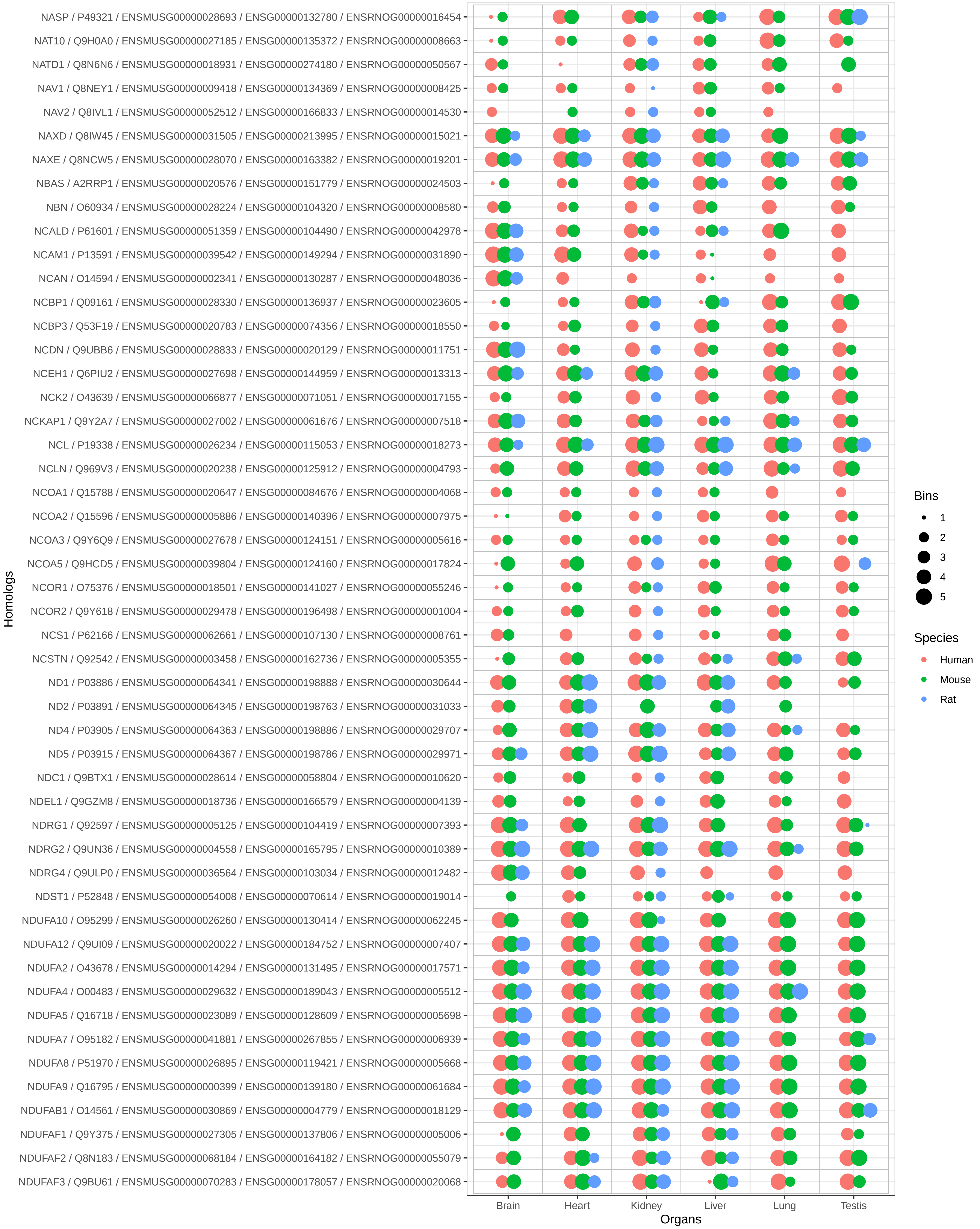
NASP / P49321 / ENSMUSG00000028693 / ENSG00000132780 / ENSRNOG00000016454
NAT10 / Q9H0A0 / ENSMUSG00000027185 / ENSG00000135372 / ENSRNOG00000008663
NATD1 / Q8N6N6 / ENSMUSG00000018931 / ENSG00000274180 / ENSRNOG00000050567
NAV1 / Q8NEY1 / ENSMUSG00000009418 / ENSG00000134369 / ENSRNOG00000008425
NAV2 / Q8IVL1 / ENSMUSG00000052512 / ENSG00000166833 / ENSRNOG00000014530
NAXD / Q8IW45 / ENSMUSG00000031505 / ENSG00000213995 / ENSRNOG00000015021
NAXE / Q8NCW5 / ENSMUSG00000028070 / ENSG00000163382 / ENSRNOG00000019201
NBAS / A2RRP1 / ENSMUSG00000020576 / ENSG00000151779 / ENSRNOG00000024503
NBN / O60934 / ENSMUSG00000028224 / ENSG00000104320 / ENSRNOG00000008580
NCALD / P61601 / ENSMUSG00000051359 / ENSG00000104490 / ENSRNOG00000042978
NCAM1 / P13591 / ENSMUSG00000039542 / ENSG00000149294 / ENSRNOG00000031890
NCAN / O14594 / ENSMUSG00000002341 / ENSG00000130287 / ENSRNOG00000048036
NCBP1 / Q09161 / ENSMUSG00000028330 / ENSG00000136937 / ENSRNOG00000023605
NCBP3 / Q53F19 / ENSMUSG00000020783 / ENSG00000074356 / ENSRNOG00000018550
NCDN / Q9UBB6 / ENSMUSG00000028833 / ENSG00000020129 / ENSRNOG00000011751
NCEH1 / Q6PIU2 / ENSMUSG00000027698 / ENSG00000144959 / ENSRNOG00000013313
NCK2 / O43639 / ENSMUSG00000066877 / ENSG00000071051 / ENSRNOG00000017155
NCKAP1 / Q9Y2A7 / ENSMUSG00000027002 / ENSG00000061676 / ENSRNOG00000007518
NCL / P19338 / ENSMUSG00000026234 / ENSG00000115053 / ENSRNOG00000018273
NCLN / Q969V3 / ENSMUSG00000020238 / ENSG00000125912 / ENSRNOG00000004793
NCOA1 / Q15788 / ENSMUSG00000020647 / ENSG00000084676 / ENSRNOG00000004068
NCOA2 / Q15596 / ENSMUSG00000005886 / ENSG00000140396 / ENSRNOG00000007975
NCOA3 / Q9Y6Q9 / ENSMUSG00000027678 / ENSG00000124151 / ENSRNOG00000005616
NCOA5 / Q9HCD5 / ENSMUSG00000039804 / ENSG00000124160 / ENSRNOG00000017824
NCOR1 / O75376 / ENSMUSG00000018501 / ENSG00000141027 / ENSRNOG00000055246
NCOR2 / Q9Y618 / ENSMUSG00000029478 / ENSG00000196498 / ENSRNOG00000001004
NCS1 / P62166 / ENSMUSG00000062661 / ENSG00000107130 / ENSRNOG00000008761
NCSTN / Q92542 / ENSMUSG00000003458 / ENSG00000162736 / ENSRNOG00000005355
ND1 / P03886 / ENSMUSG00000064341 / ENSG00000198888 / ENSRNOG00000030644
ND2 / P03891 / ENSMUSG00000064345 / ENSG00000198763 / ENSRNOG00000031033
ND4 / P03905 / ENSMUSG00000064363 / ENSG00000198886 / ENSRNOG00000029707
ND5 / P03915 / ENSMUSG00000064367 / ENSG00000198786 / ENSRNOG00000029971
NDC1 / Q9BTX1 / ENSMUSG00000028614 / ENSG00000058804 / ENSRNOG00000010620
NDEL1 / Q9GZM8 / ENSMUSG00000018736 / ENSG00000166579 / ENSRNOG00000004139
NDRG1 / Q92597 / ENSMUSG00000005125 / ENSG00000104419 / ENSRNOG00000007393
NDRG2 / Q9UN36 / ENSMUSG00000004558 / ENSG00000165795 / ENSRNOG00000010389
NDRG4 / Q9ULP0 / ENSMUSG00000036564 / ENSG00000103034 / ENSRNOG00000012482
NDST1 / P52848 / ENSMUSG00000054008 / ENSG00000070614 / ENSRNOG00000019014
NDUFA10 / O95299 / ENSMUSG00000026260 / ENSG00000130414 / ENSRNOG00000062245
NDUFA12 / Q9UI09 / ENSMUSG00000020022 / ENSG00000184752 / ENSRNOG00000007407
NDUFA2 / O43678 / ENSMUSG00000014294 / ENSG00000131495 / ENSRNOG00000017571
NDUFA4 / O00483 / ENSMUSG00000029632 / ENSG00000189043 / ENSRNOG00000005512
NDUFA5 / Q16718 / ENSMUSG00000023089 / ENSG00000128609 / ENSRNOG00000005698
NDUFA7 / O95182 / ENSMUSG00000041881 / ENSG00000267855 / ENSRNOG00000006939
NDUFA8 / P51970 / ENSMUSG00000026895 / ENSG00000119421 / ENSRNOG00000005668
NDUFA9 / Q16795 / ENSMUSG00000000399 / ENSG00000139180 / ENSRNOG00000061684
NDUFAB1 / O14561 / ENSMUSG00000030869 / ENSG00000004779 / ENSRNOG00000018129
NDUFAF1 / Q9Y375 / ENSMUSG00000027305 / ENSG00000137806 / ENSRNOG00000005006
NDUFAF2 / Q8N183 / ENSMUSG00000068184 / ENSG00000164182 / ENSRNOG00000055079
NDUFAF3 / Q9BU61 / ENSMUSG00000070283 / ENSG00000178057 / ENSRNOG00000020068
Homologs
Brain
Heart
Kidney
Liver
Lung
Testis
Organs
Bins
1
2
3
4
5
Species
Human
Mouse
Rat

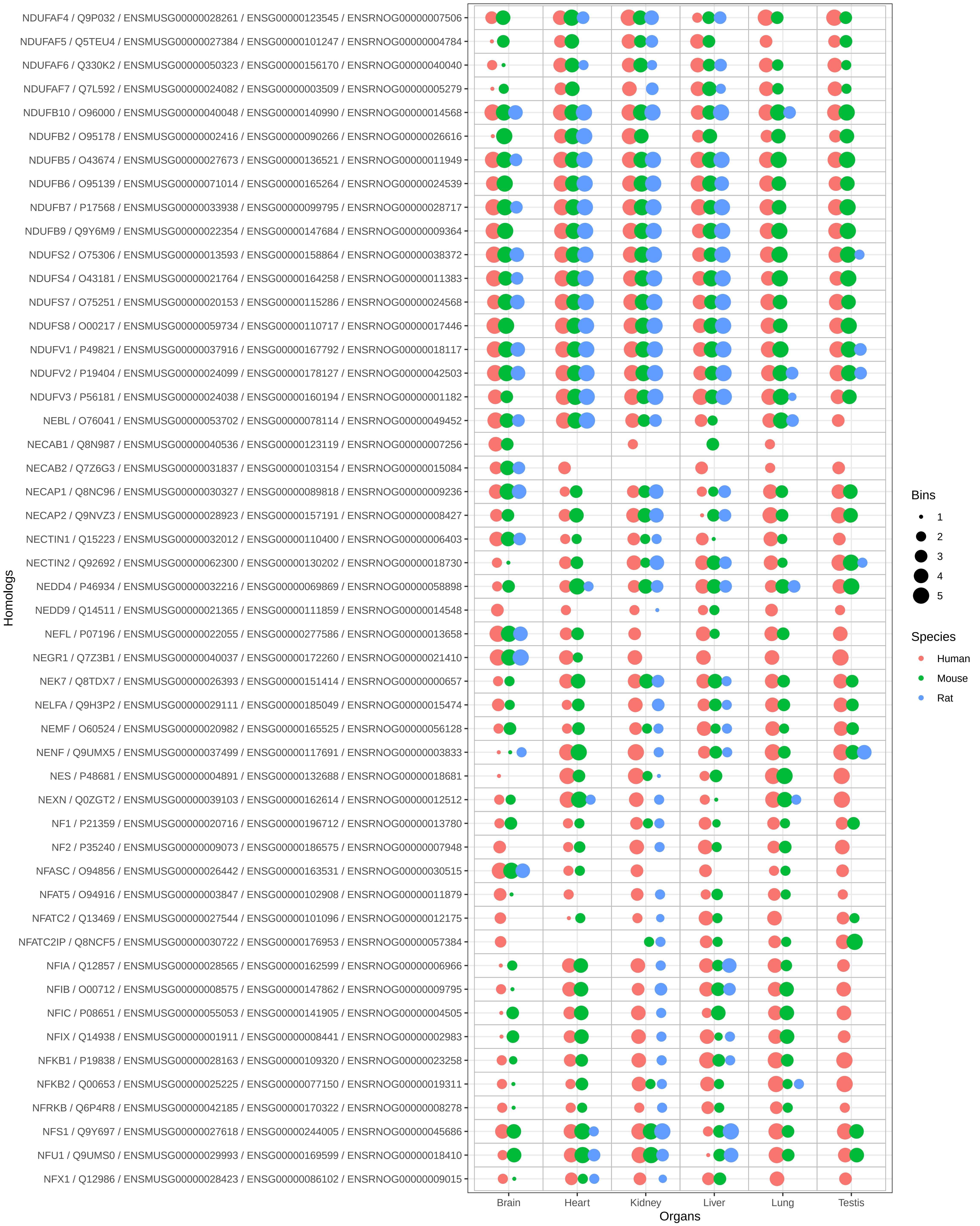
NDUFAF4 / Q9P032 / ENSMUSG00000028261 / ENSG00000123545 / ENSRNOG00000007506
NDUFAF5 / Q5TEU4 / ENSMUSG00000027384 / ENSG00000101247 / ENSRNOG00000004784
NDUFAF6 / Q330K2 / ENSMUSG00000050323 / ENSG00000156170 / ENSRNOG00000040040
NDUFAF7 / Q7L592 / ENSMUSG00000024082 / ENSG00000003509 / ENSRNOG00000005279
NDUFB10 / O96000 / ENSMUSG00000040048 / ENSG00000140990 / ENSRNOG00000014568
NDUFB2 / O95178 / ENSMUSG00000002416 / ENSG00000090266 / ENSRNOG00000026616
NDUFB5 / O43674 / ENSMUSG00000027673 / ENSG00000136521 / ENSRNOG00000011949
NDUFB6 / O95139 / ENSMUSG00000071014 / ENSG00000165264 / ENSRNOG00000024539
NDUFB7 / P17568 / ENSMUSG00000033938 / ENSG00000099795 / ENSRNOG00000028717
NDUFB9 / Q9Y6M9 / ENSMUSG00000022354 / ENSG00000147684 / ENSRNOG00000009364
NDUFS2 / O75306 / ENSMUSG00000013593 / ENSG00000158864 / ENSRNOG00000038372
NDUFS4 / O43181 / ENSMUSG00000021764 / ENSG00000164258 / ENSRNOG00000011383
NDUFS7 / O75251 / ENSMUSG00000020153 / ENSG00000115286 / ENSRNOG00000024568
NDUFS8 / O00217 / ENSMUSG00000059734 / ENSG00000110717 / ENSRNOG00000017446
NDUFV1 / P49821 / ENSMUSG00000037916 / ENSG00000167792 / ENSRNOG00000018117
NDUFV2 / P19404 / ENSMUSG00000024099 / ENSG00000178127 / ENSRNOG00000042503
NDUFV3 / P56181 / ENSMUSG00000024038 / ENSG00000160194 / ENSRNOG00000001182
NEBL / O76041 / ENSMUSG00000053702 / ENSG00000078114 / ENSRNOG00000049452
NECAB1 / Q8N987 / ENSMUSG00000040536 / ENSG00000123119 / ENSRNOG00000007256
NECAB2 / Q7Z6G3 / ENSMUSG00000031837 / ENSG00000103154 / ENSRNOG00000015084
NECAP1 / Q8NC96 / ENSMUSG00000030327 / ENSG00000089818 / ENSRNOG00000009236
NECAP2 / Q9NVZ3 / ENSMUSG00000028923 / ENSG00000157191 / ENSRNOG00000008427
NECTIN1 / Q15223 / ENSMUSG00000032012 / ENSG00000110400 / ENSRNOG00000006403
NECTIN2 / Q92692 / ENSMUSG00000062300 / ENSG00000130202 / ENSRNOG00000018730
NEDD4 / P46934 / ENSMUSG00000032216 / ENSG00000069869 / ENSRNOG00000058898
NEDD9 / Q14511 / ENSMUSG00000021365 / ENSG00000111859 / ENSRNOG00000014548
NEFL / P07196 / ENSMUSG00000022055 / ENSG00000277586 / ENSRNOG00000013658
NEGR1 / Q7Z3B1 / ENSMUSG00000040037 / ENSG00000172260 / ENSRNOG00000021410
NEK7 / Q8TDX7 / ENSMUSG00000026393 / ENSG00000151414 / ENSRNOG00000000657
NELFA / Q9H3P2 / ENSMUSG00000029111 / ENSG00000185049 / ENSRNOG00000015474
NEMF / O60524 / ENSMUSG00000020982 / ENSG00000165525 / ENSRNOG00000056128
NENF / Q9UMX5 / ENSMUSG00000037499 / ENSG00000117691 / ENSRNOG00000003833
NES / P48681 / ENSMUSG00000004891 / ENSG00000132688 / ENSRNOG00000018681
NEXN / Q0ZGT2 / ENSMUSG00000039103 / ENSG00000162614 / ENSRNOG00000012512
NF1 / P21359 / ENSMUSG00000020716 / ENSG00000196712 / ENSRNOG00000013780
NF2 / P35240 / ENSMUSG00000009073 / ENSG00000186575 / ENSRNOG00000007948
NFASC / O94856 / ENSMUSG00000026442 / ENSG00000163531 / ENSRNOG00000030515
NFAT5 / O94916 / ENSMUSG00000003847 / ENSG00000102908 / ENSRNOG00000011879
NFATC2 / Q13469 / ENSMUSG00000027544 / ENSG00000101096 / ENSRNOG00000012175
NFATC2IP / Q8NCF5 / ENSMUSG00000030722 / ENSG00000176953 / ENSRNOG00000057384
NFIA / Q12857 / ENSMUSG00000028565 / ENSG00000162599 / ENSRNOG00000006966
NFIB / O00712 / ENSMUSG00000008575 / ENSG00000147862 / ENSRNOG00000009795
NFIC / P08651 / ENSMUSG00000055053 / ENSG00000141905 / ENSRNOG00000004505
NFIX / Q14938 / ENSMUSG00000001911 / ENSG00000008441 / ENSRNOG00000002983
NFKB1 / P19838 / ENSMUSG00000028163 / ENSG00000109320 / ENSRNOG00000023258
NFKB2 / Q00653 / ENSMUSG00000025225 / ENSG00000077150 / ENSRNOG00000019311
NFRKB / Q6P4R8 / ENSMUSG00000042185 / ENSG00000170322 / ENSRNOG00000008278
NFS1 / Q9Y697 / ENSMUSG00000027618 / ENSG00000244005 / ENSRNOG00000045686
NFU1 / Q9UMS0 / ENSMUSG00000029993 / ENSG00000169599 / ENSRNOG00000018410
NFX1 / Q12986 / ENSMUSG00000028423 / ENSG00000086102 / ENSRNOG00000009015
Homologs
Brain
Heart
Kidney
Liver
Lung
Testis
Organs
Bins
1
2
3
4
5
Species
Human
Mouse
Rat

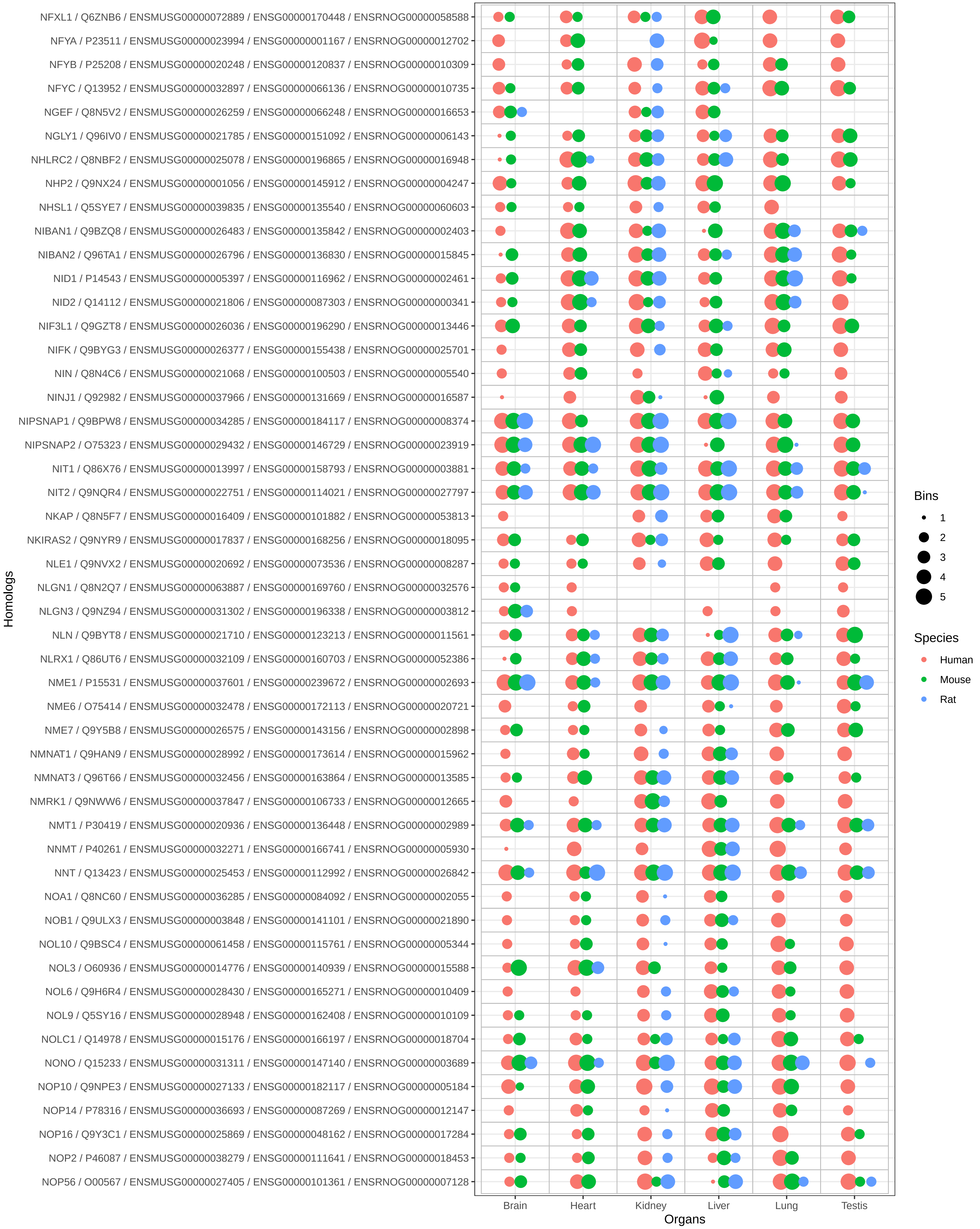

NFXL1 / Q6ZNB6 / ENSMUSG00000072889 / ENSG00000170448 / ENSRNOG00000058588
NFYA / P23511 / ENSMUSG00000023994 / ENSG00000001167 / ENSRNOG00000012702
NFYB / P25208 / ENSMUSG00000020248 / ENSG00000120837 / ENSRNOG00000010309
NFYC / Q13952 / ENSMUSG00000032897 / ENSG00000066136 / ENSRNOG00000010735
NGEF / Q8N5V2 / ENSMUSG00000026259 / ENSG00000066248 / ENSRNOG00000016653
NGLY1 / Q96IV0 / ENSMUSG00000021785 / ENSG00000151092 / ENSRNOG00000006143
NHLRC2 / Q8NBF2 / ENSMUSG00000025078 / ENSG00000196865 / ENSRNOG00000016948
NHP2 / Q9NX24 / ENSMUSG00000001056 / ENSG00000145912 / ENSRNOG00000004247
NHSL1 / Q5SYE7 / ENSMUSG00000039835 / ENSG00000135540 / ENSRNOG00000060603
NIBAN1 / Q9BZQ8 / ENSMUSG00000026483 / ENSG00000135842 / ENSRNOG00000002403
NIBAN2 / Q96TA1 / ENSMUSG00000026796 / ENSG00000136830 / ENSRNOG00000015845
NID1 / P14543 / ENSMUSG00000005397 / ENSG00000116962 / ENSRNOG00000002461
NID2 / Q14112 / ENSMUSG00000021806 / ENSG00000087303 / ENSRNOG00000000341
NIF3L1 / Q9GZT8 / ENSMUSG00000026036 / ENSG00000196290 / ENSRNOG00000013446
NIFK / Q9BYG3 / ENSMUSG00000026377 / ENSG00000155438 / ENSRNOG00000025701
NIN / Q8N4C6 / ENSMUSG00000021068 / ENSG00000100503 / ENSRNOG00000005540
NINJ1 / Q92982 / ENSMUSG00000037966 / ENSG00000131669 / ENSRNOG00000016587
NIPSNAP1 / Q9BPW8 / ENSMUSG00000034285 / ENSG00000184117 / ENSRNOG00000008374
NIPSNAP2 / O75323 / ENSMUSG00000029432 / ENSG00000146729 / ENSRNOG00000023919
NIT1 / Q86X76 / ENSMUSG00000013997 / ENSG00000158793 / ENSRNOG00000003881
NIT2 / Q9NQR4 / ENSMUSG00000022751 / ENSG00000114021 / ENSRNOG00000027797
NKAP / Q8N5F7 / ENSMUSG00000016409 / ENSG00000101882 / ENSRNOG00000053813
NKIRAS2 / Q9NYR9 / ENSMUSG00000017837 / ENSG00000168256 / ENSRNOG00000018095
NLE1 / Q9NVX2 / ENSMUSG00000020692 / ENSG00000073536 / ENSRNOG00000008287
NLGN1 / Q8N2Q7 / ENSMUSG00000063887 / ENSG00000169760 / ENSRNOG00000032576
NLGN3 / Q9NZ94 / ENSMUSG00000031302 / ENSG00000196338 / ENSRNOG00000003812
NLN / Q9BYT8 / ENSMUSG00000021710 / ENSG00000123213 / ENSRNOG00000011561
NLRX1 / Q86UT6 / ENSMUSG00000032109 / ENSG00000160703 / ENSRNOG00000052386
NME1 / P15531 / ENSMUSG00000037601 / ENSG00000239672 / ENSRNOG00000002693
NME6 / O75414 / ENSMUSG00000032478 / ENSG00000172113 / ENSRNOG00000020721
NME7 / Q9Y5B8 / ENSMUSG00000026575 / ENSG00000143156 / ENSRNOG00000002898
NMNAT1 / Q9HAN9 / ENSMUSG00000028992 / ENSG00000173614 / ENSRNOG00000015962
NMNAT3 / Q96T66 / ENSMUSG00000032456 / ENSG00000163864 / ENSRNOG00000013585
NMRK1 / Q9NWW6 / ENSMUSG00000037847 / ENSG00000106733 / ENSRNOG00000012665
NMT1 / P30419 / ENSMUSG00000020936 / ENSG00000136448 / ENSRNOG00000002989
NNMT / P40261 / ENSMUSG00000032271 / ENSG00000166741 / ENSRNOG00000005930
NNT / Q13423 / ENSMUSG00000025453 / ENSG00000112992 / ENSRNOG00000026842
NOA1 / Q8NC60 / ENSMUSG00000036285 / ENSG00000084092 / ENSRNOG00000002055
NOB1 / Q9ULX3 / ENSMUSG00000003848 / ENSG00000141101 / ENSRNOG00000021890
NOL10 / Q9BSC4 / ENSMUSG00000061458 / ENSG00000115761 / ENSRNOG00000005344
NOL3 / O60936 / ENSMUSG00000014776 / ENSG00000140939 / ENSRNOG00000015588
NOL6 / Q9H6R4 / ENSMUSG00000028430 / ENSG00000165271 / ENSRNOG00000010409
NOL9 / Q5SY16 / ENSMUSG00000028948 / ENSG00000162408 / ENSRNOG00000010109
NOLC1 / Q14978 / ENSMUSG00000015176 / ENSG00000166197 / ENSRNOG00000018704
NONO / Q15233 / ENSMUSG00000031311 / ENSG00000147140 / ENSRNOG00000003689
NOP10 / Q9NPE3 / ENSMUSG00000027133 / ENSG00000182117 / ENSRNOG00000005184
NOP14 / P78316 / ENSMUSG00000036693 / ENSG00000087269 / ENSRNOG00000012147
NOP16 / Q9Y3C1 / ENSMUSG00000025869 / ENSG00000048162 / ENSRNOG00000017284
NOP2 / P46087 / ENSMUSG00000038279 / ENSG00000111641 / ENSRNOG00000018453
NOP56 / O00567 / ENSMUSG00000027405 / ENSG00000101361 / ENSRNOG00000007128
Homologs
Brain
Heart
Kidney
Liver
Lung
Testis
Organs
Bins
1
2
3
4
5
Species
Human
Mouse
Rat

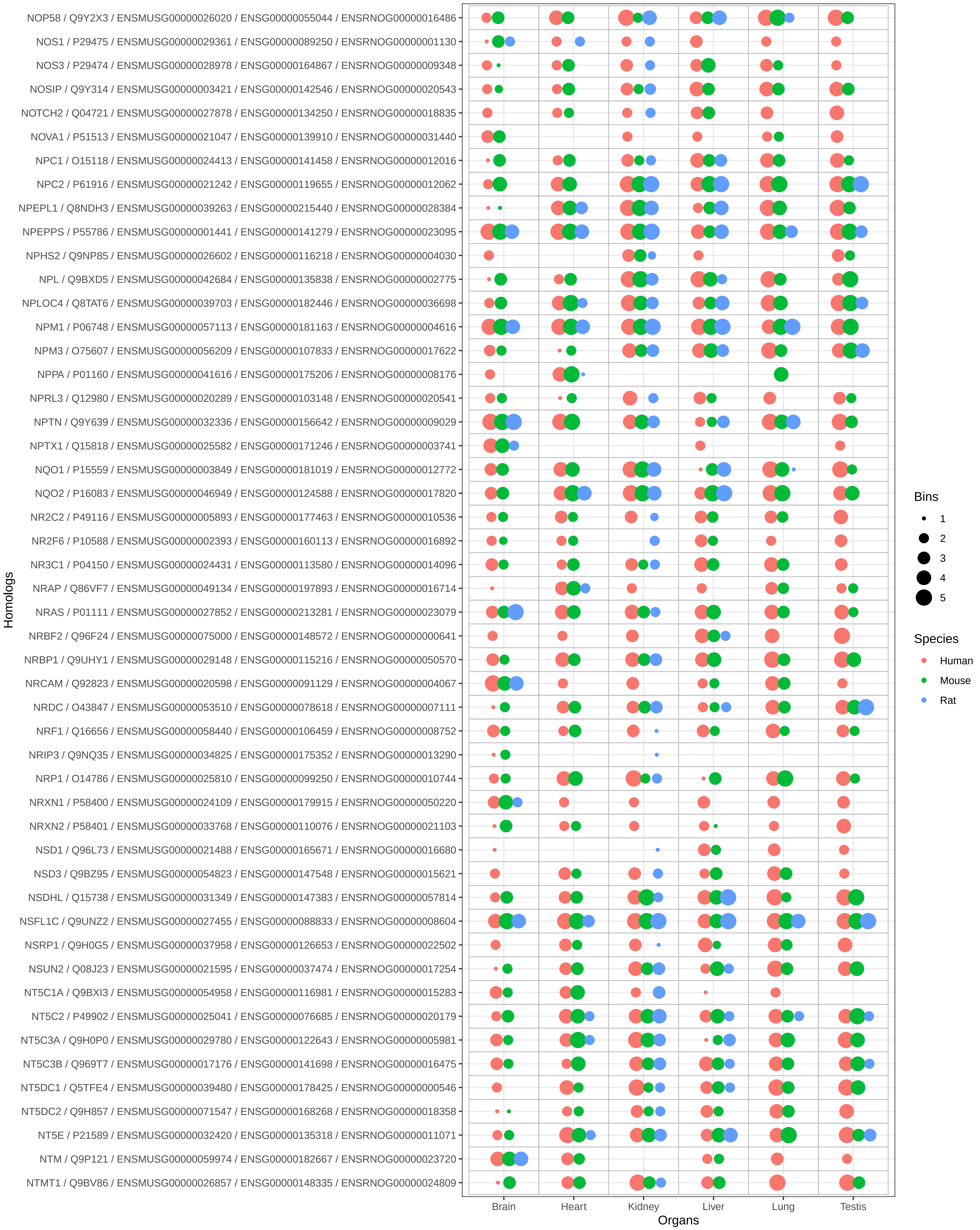
NOP58 / Q9Y2X3 / ENSMUSG00000026020 / ENSG00000055044 / ENSRNOG00000016486
NOS1 / P29475 / ENSMUSG00000029361 / ENSG00000089250 / ENSRNOG00000001130
NOS3 / P29474 / ENSMUSG00000028978 / ENSG00000164867 / ENSRNOG00000009348
NOSIP / Q9Y314 / ENSMUSG00000003421 / ENSG00000142546 / ENSRNOG00000020543
NOTCH2 / Q04721 / ENSMUSG00000027878 / ENSG00000134250 / ENSRNOG00000018835
NOVA1 / P51513 / ENSMUSG00000021047 / ENSG00000139910 / ENSRNOG00000031440
NPC1 / O15118 / ENSMUSG00000024413 / ENSG00000141458 / ENSRNOG00000012016
NPC2 / P61916 / ENSMUSG00000021242 / ENSG00000119655 / ENSRNOG00000012062
NPEPL1 / Q8NDH3 / ENSMUSG00000039263 / ENSG00000215440 / ENSRNOG00000028384
NPEPPS / P55786 / ENSMUSG00000001441 / ENSG00000141279 / ENSRNOG00000023095
NPHS2 / Q9NP85 / ENSMUSG00000026602 / ENSG00000116218 / ENSRNOG00000004030
NPL / Q9BXD5 / ENSMUSG00000042684 / ENSG00000135838 / ENSRNOG00000002775
NPLOC4 / Q8TAT6 / ENSMUSG00000039703 / ENSG00000182446 / ENSRNOG00000036698
NPM1 / P06748 / ENSMUSG00000057113 / ENSG00000181163 / ENSRNOG00000004616
NPM3 / O75607 / ENSMUSG00000056209 / ENSG00000107833 / ENSRNOG00000017622
NPPA / P01160 / ENSMUSG00000041616 / ENSG00000175206 / ENSRNOG00000008176
NPRL3 / Q12980 / ENSMUSG00000020289 / ENSG00000103148 / ENSRNOG00000020541
NPTN / Q9Y639 / ENSMUSG00000032336 / ENSG00000156642 / ENSRNOG00000009029
NPTX1 / Q15818 / ENSMUSG00000025582 / ENSG00000171246 / ENSRNOG00000003741
NQO1 / P15559 / ENSMUSG00000003849 / ENSG00000181019 / ENSRNOG00000012772
NQO2 / P16083 / ENSMUSG00000046949 / ENSG00000124588 / ENSRNOG00000017820
NR2C2 / P49116 / ENSMUSG00000005893 / ENSG00000177463 / ENSRNOG00000010536
NR2F6 / P10588 / ENSMUSG00000002393 / ENSG00000160113 / ENSRNOG00000016892
NR3C1 / P04150 / ENSMUSG00000024431 / ENSG00000113580 / ENSRNOG00000014096
NRAP / Q86VF7 / ENSMUSG00000049134 / ENSG00000197893 / ENSRNOG00000016714
NRAS / P01111 / ENSMUSG00000027852 / ENSG00000213281 / ENSRNOG00000023079
NRBF2 / Q96F24 / ENSMUSG00000075000 / ENSG00000148572 / ENSRNOG00000000641
NRBP1 / Q9UHY1 / ENSMUSG00000029148 / ENSG00000115216 / ENSRNOG00000050570
NRCAM / Q92823 / ENSMUSG00000020598 / ENSG00000091129 / ENSRNOG00000004067
NRDC / O43847 / ENSMUSG00000053510 / ENSG00000078618 / ENSRNOG00000007111
NRF1 / Q16656 / ENSMUSG00000058440 / ENSG00000106459 / ENSRNOG00000008752
NRIP3 / Q9NQ35 / ENSMUSG00000034825 / ENSG00000175352 / ENSRNOG00000013290
NRP1 / O14786 / ENSMUSG00000025810 / ENSG00000099250 / ENSRNOG00000010744
NRXN1 / P58400 / ENSMUSG00000024109 / ENSG00000179915 / ENSRNOG00000050220
NRXN2 / P58401 / ENSMUSG00000033768 / ENSG00000110076 / ENSRNOG00000021103
NSD1 / Q96L73 / ENSMUSG00000021488 / ENSG00000165671 / ENSRNOG00000016680
NSD3 / Q9BZ95 / ENSMUSG00000054823 / ENSG00000147548 / ENSRNOG00000015621
NSDHL / Q15738 / ENSMUSG00000031349 / ENSG00000147383 / ENSRNOG00000057814
NSFL1C / Q9UNZ2 / ENSMUSG00000027455 / ENSG00000088833 / ENSRNOG00000008604
NSRP1 / Q9H0G5 / ENSMUSG00000037958 / ENSG00000126653 / ENSRNOG00000022502
NSUN2 / Q08J23 / ENSMUSG00000021595 / ENSG00000037474 / ENSRNOG00000017254
NT5C1A / Q9BXI3 / ENSMUSG00000054958 / ENSG00000116981 / ENSRNOG00000015283
NT5C2 / P49902 / ENSMUSG00000025041 / ENSG00000076685 / ENSRNOG00000020179
NT5C3A / Q9H0P0 / ENSMUSG00000029780 / ENSG00000122643 / ENSRNOG00000005981
NT5C3B / Q969T7 / ENSMUSG00000017176 / ENSG00000141698 / ENSRNOG00000016475
NT5DC1 / Q5TFE4 / ENSMUSG00000039480 / ENSG00000178425 / ENSRNOG00000000546
NT5DC2 / Q9H857 / ENSMUSG00000071547 / ENSG00000168268 / ENSRNOG00000018358
NT5E / P21589 / ENSMUSG00000032420 / ENSG00000135318 / ENSRNOG00000011071
NTM / Q9P121 / ENSMUSG00000059974 / ENSG00000182667 / ENSRNOG00000023720
NTMT1 / Q9BV86 / ENSMUSG00000026857 / ENSG00000148335 / ENSRNOG00000024809
Homologs
Brain
Heart
Kidney
Liver
Lung
Testis
Organs
Bins
1
2
3
4
5
Species
Human
Mouse
Rat

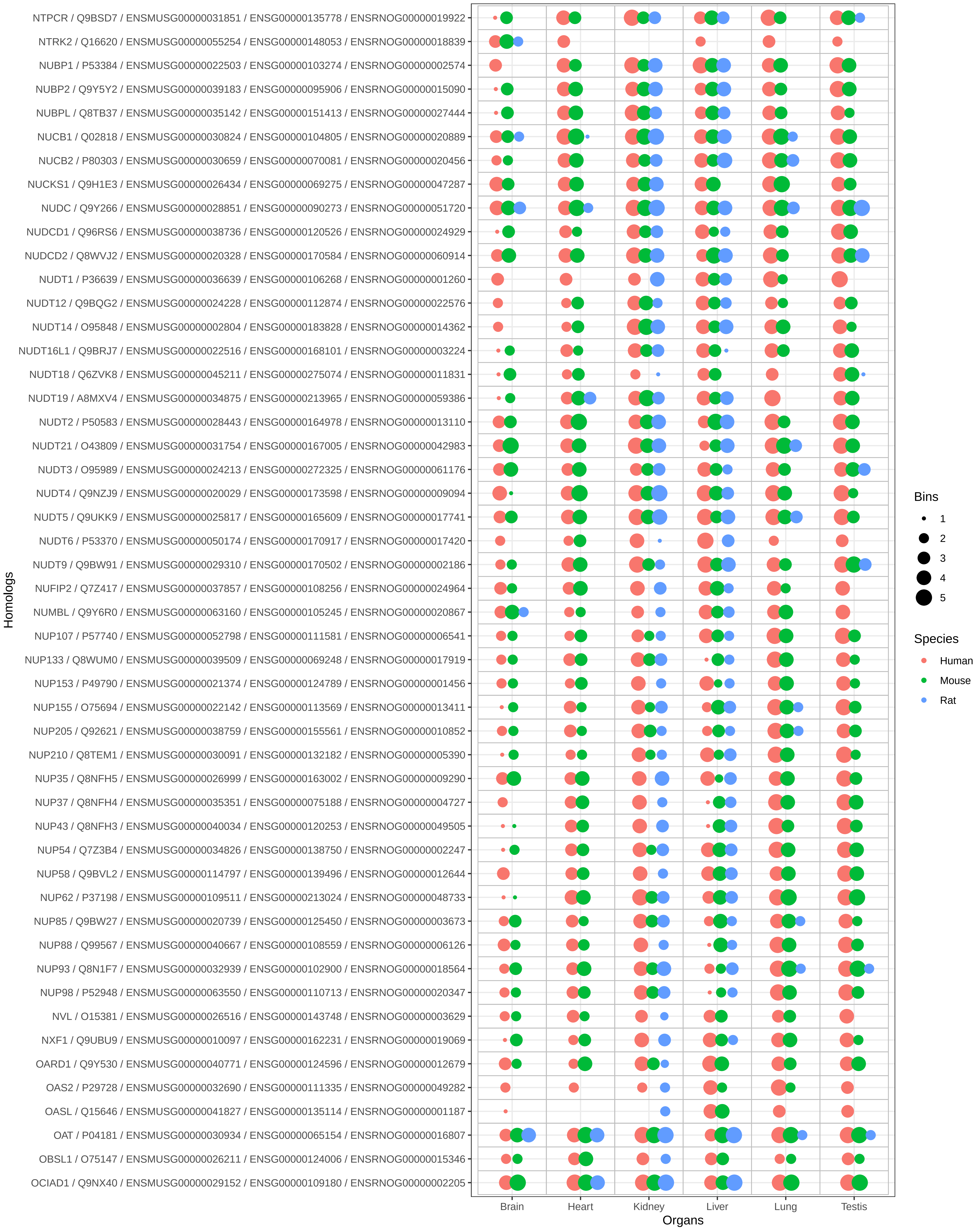

NTPCR / Q9BSD7 / ENSMUSG00000031851 / ENSG00000135778 / ENSRNOG00000019922
NTRK2 / Q16620 / ENSMUSG00000055254 / ENSG00000148053 / ENSRNOG00000018839
NUBP1 / P53384 / ENSMUSG00000022503 / ENSG00000103274 / ENSRNOG00000002574
NUBP2 / Q9Y5Y2 / ENSMUSG00000039183 / ENSG00000095906 / ENSRNOG00000015090
NUBPL / Q8TB37 / ENSMUSG00000035142 / ENSG00000151413 / ENSRNOG00000027444
NUCB1 / Q02818 / ENSMUSG00000030824 / ENSG00000104805 / ENSRNOG00000020889
NUCB2 / P80303 / ENSMUSG00000030659 / ENSG00000070081 / ENSRNOG00000020456
NUCKS1 / Q9H1E3 / ENSMUSG00000026434 / ENSG00000069275 / ENSRNOG00000047287
NUDC / Q9Y266 / ENSMUSG00000028851 / ENSG00000090273 / ENSRNOG00000051720
NUDCD1 / Q96RS6 / ENSMUSG00000038736 / ENSG00000120526 / ENSRNOG00000024929
NUDCD2 / Q8WVJ2 / ENSMUSG00000020328 / ENSG00000170584 / ENSRNOG00000060914
NUDT1 / P36639 / ENSMUSG00000036639 / ENSG00000106268 / ENSRNOG00000001260
NUDT12 / Q9BQG2 / ENSMUSG00000024228 / ENSG00000112874 / ENSRNOG00000022576
NUDT14 / O95848 / ENSMUSG00000002804 / ENSG00000183828 / ENSRNOG00000014362
NUDT16L1 / Q9BRJ7 / ENSMUSG00000022516 / ENSG00000168101 / ENSRNOG00000003224
NUDT18 / Q6ZVK8 / ENSMUSG00000045211 / ENSG00000275074 / ENSRNOG00000011831
NUDT19 / A8MXV4 / ENSMUSG00000034875 / ENSG00000213965 / ENSRNOG00000059386
NUDT2 / P50583 / ENSMUSG00000028443 / ENSG00000164978 / ENSRNOG00000013110
NUDT21 / O43809 / ENSMUSG00000031754 / ENSG00000167005 / ENSRNOG00000042983
NUDT3 / O95989 / ENSMUSG00000024213 / ENSG00000272325 / ENSRNOG00000061176
NUDT4 / Q9NZJ9 / ENSMUSG00000020029 / ENSG00000173598 / ENSRNOG00000009094
NUDT5 / Q9UKK9 / ENSMUSG00000025817 / ENSG00000165609 / ENSRNOG00000017741
NUDT6 / P53370 / ENSMUSG00000050174 / ENSG00000170917 / ENSRNOG00000017420
NUDT9 / Q9BW91 / ENSMUSG00000029310 / ENSG00000170502 / ENSRNOG00000002186
NUFIP2 / Q7Z417 / ENSMUSG00000037857 / ENSG00000108256 / ENSRNOG00000024964
NUMBL / Q9Y6R0 / ENSMUSG00000063160 / ENSG00000105245 / ENSRNOG00000020867
NUP107 / P57740 / ENSMUSG00000052798 / ENSG00000111581 / ENSRNOG00000006541
NUP133 / Q8WUM0 / ENSMUSG00000039509 / ENSG00000069248 / ENSRNOG00000017919
NUP153 / P49790 / ENSMUSG00000021374 / ENSG00000124789 / ENSRNOG00000001456
NUP155 / O75694 / ENSMUSG00000022142 / ENSG00000113569 / ENSRNOG00000013411
NUP205 / Q92621 / ENSMUSG00000038759 / ENSG00000155561 / ENSRNOG00000010852
NUP210 / Q8TEM1 / ENSMUSG00000030091 / ENSG00000132182 / ENSRNOG00000005390
NUP35 / Q8NFH5 / ENSMUSG00000026999 / ENSG00000163002 / ENSRNOG00000009290
NUP37 / Q8NFH4 / ENSMUSG00000035351 / ENSG00000075188 / ENSRNOG00000004727
NUP43 / Q8NFH3 / ENSMUSG00000040034 / ENSG00000120253 / ENSRNOG00000049505
NUP54 / Q7Z3B4 / ENSMUSG00000034826 / ENSG00000138750 / ENSRNOG00000002247
NUP58 / Q9BVL2 / ENSMUSG00000114797 / ENSG00000139496 / ENSRNOG00000012644
NUP62 / P37198 / ENSMUSG00000109511 / ENSG00000213024 / ENSRNOG00000048733
NUP85 / Q9BW27 / ENSMUSG00000020739 / ENSG00000125450 / ENSRNOG00000003673
NUP88 / Q99567 / ENSMUSG00000040667 / ENSG00000108559 / ENSRNOG00000006126
NUP93 / Q8N1F7 / ENSMUSG00000032939 / ENSG00000102900 / ENSRNOG00000018564
NUP98 / P52948 / ENSMUSG00000063550 / ENSG00000110713 / ENSRNOG00000020347
NVL / O15381 / ENSMUSG00000026516 / ENSG00000143748 / ENSRNOG00000003629
NXF1 / Q9UBU9 / ENSMUSG00000010097 / ENSG00000162231 / ENSRNOG00000019069
OARD1 / Q9Y530 / ENSMUSG00000040771 / ENSG00000124596 / ENSRNOG00000012679
OAS2 / P29728 / ENSMUSG00000032690 / ENSG00000111335 / ENSRNOG00000049282
OASL / Q15646 / ENSMUSG00000041827 / ENSG00000135114 / ENSRNOG00000001187
OAT / P04181 / ENSMUSG00000030934 / ENSG00000065154 / ENSRNOG00000016807
OBSL1 / O75147 / ENSMUSG00000026211 / ENSG00000124006 / ENSRNOG00000015346
OCIAD1 / Q9NX40 / ENSMUSG00000029152 / ENSG00000109180 / ENSRNOG00000002205
Homologs
Brain
Heart
Kidney
Liver
Lung
Testis
Organs
Bins
1
2
3
4
5
Species
Human
Mouse
Rat

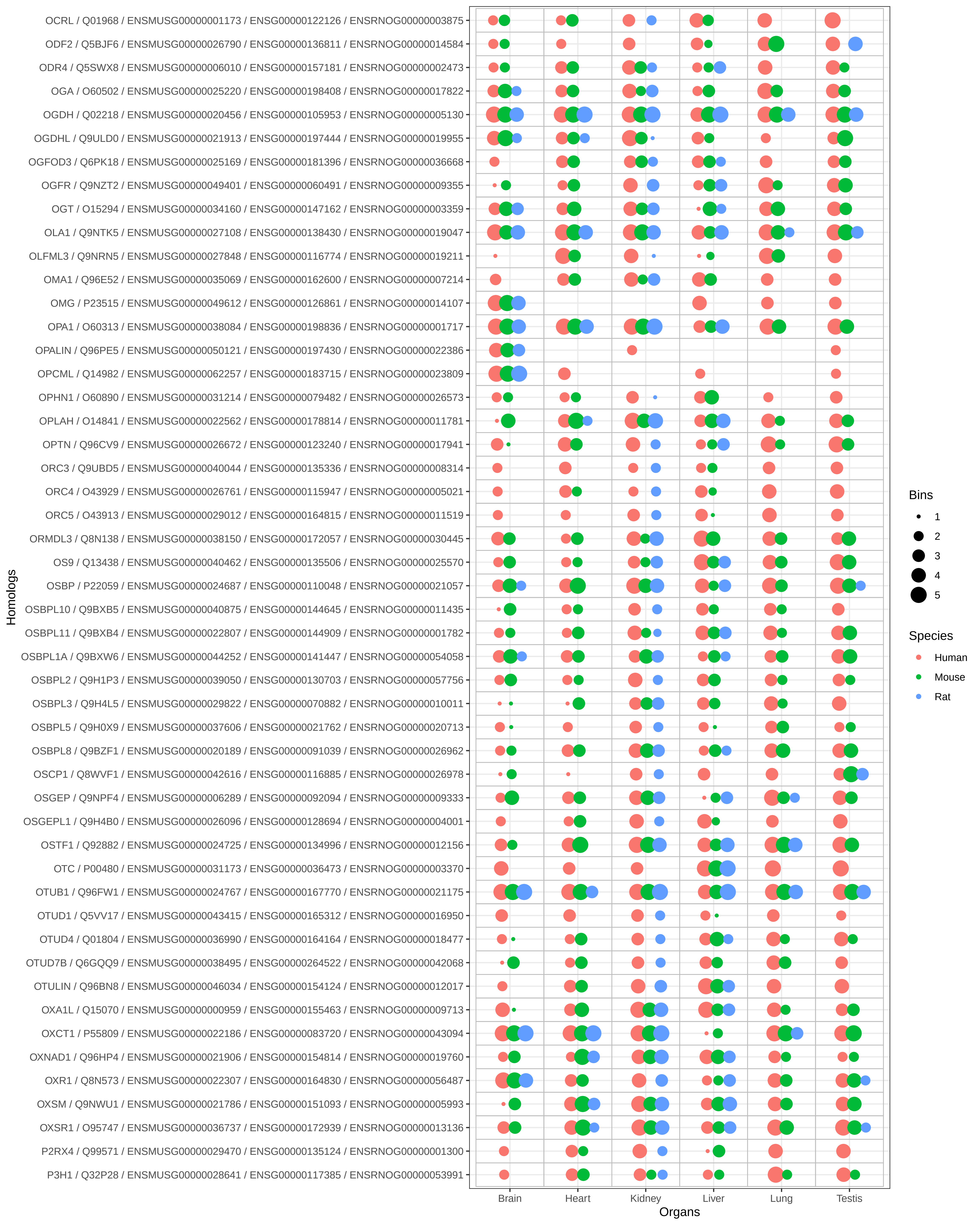

OCRL / Q01968 / ENSMUSG00000001173 / ENSG00000122126 / ENSRNOG00000003875
ODF2 / Q5BJF6 / ENSMUSG00000026790 / ENSG00000136811 / ENSRNOG00000014584
ODR4 / Q5SWX8 / ENSMUSG00000006010 / ENSG00000157181 / ENSRNOG00000002473
OGA / O60502 / ENSMUSG00000025220 / ENSG00000198408 / ENSRNOG00000017822
OGDH / Q02218 / ENSMUSG00000020456 / ENSG00000105953 / ENSRNOG00000005130
OGDHL / Q9ULD0 / ENSMUSG00000021913 / ENSG00000197444 / ENSRNOG00000019955
OGFOD3 / Q6PK18 / ENSMUSG00000025169 / ENSG00000181396 / ENSRNOG00000036668
OGFR / Q9NZT2 / ENSMUSG00000049401 / ENSG00000060491 / ENSRNOG00000009355
OGT / O15294 / ENSMUSG00000034160 / ENSG00000147162 / ENSRNOG00000003359
OLA1 / Q9NTK5 / ENSMUSG00000027108 / ENSG00000138430 / ENSRNOG00000019047
OLFML3 / Q9NRN5 / ENSMUSG00000027848 / ENSG00000116774 / ENSRNOG00000019211
OMA1 / Q96E52 / ENSMUSG00000035069 / ENSG00000162600 / ENSRNOG00000007214
OMG / P23515 / ENSMUSG00000049612 / ENSG00000126861 / ENSRNOG00000014107
OPA1 / O60313 / ENSMUSG00000038084 / ENSG00000198836 / ENSRNOG00000001717
OPALIN / Q96PE5 / ENSMUSG00000050121 / ENSG00000197430 / ENSRNOG00000022386
OPCML / Q14982 / ENSMUSG00000062257 / ENSG00000183715 / ENSRNOG00000023809
OPHN1 / O60890 / ENSMUSG00000031214 / ENSG00000079482 / ENSRNOG00000026573
OPLAH / O14841 / ENSMUSG00000022562 / ENSG00000178814 / ENSRNOG00000011781
OPTN / Q96CV9 / ENSMUSG00000026672 / ENSG00000123240 / ENSRNOG00000017941
ORC3 / Q9UBD5 / ENSMUSG00000040044 / ENSG00000135336 / ENSRNOG00000008314
ORC4 / O43929 / ENSMUSG00000026761 / ENSG00000115947 / ENSRNOG00000005021
ORC5 / O43913 / ENSMUSG00000029012 / ENSG00000164815 / ENSRNOG00000011519
ORMDL3 / Q8N138 / ENSMUSG00000038150 / ENSG00000172057 / ENSRNOG00000030445
OS9 / Q13438 / ENSMUSG00000040462 / ENSG00000135506 / ENSRNOG00000025570
OSBP / P22059 / ENSMUSG00000024687 / ENSG00000110048 / ENSRNOG00000021057
OSBPL10 / Q9BXB5 / ENSMUSG00000040875 / ENSG00000144645 / ENSRNOG00000011435
OSBPL11 / Q9BXB4 / ENSMUSG00000022807 / ENSG00000144909 / ENSRNOG00000001782
OSBPL1A / Q9BXW6 / ENSMUSG00000044252 / ENSG00000141447 / ENSRNOG00000054058
OSBPL2 / Q9H1P3 / ENSMUSG00000039050 / ENSG00000130703 / ENSRNOG00000057756
OSBPL3 / Q9H4L5 / ENSMUSG00000029822 / ENSG00000070882 / ENSRNOG00000010011
OSBPL5 / Q9H0X9 / ENSMUSG00000037606 / ENSG00000021762 / ENSRNOG00000020713
OSBPL8 / Q9BZF1 / ENSMUSG00000020189 / ENSG00000091039 / ENSRNOG00000026962
OSCP1 / Q8WVF1 / ENSMUSG00000042616 / ENSG00000116885 / ENSRNOG00000026978
OSGEP / Q9NPF4 / ENSMUSG00000006289 / ENSG00000092094 / ENSRNOG00000009333
OSGEPL1 / Q9H4B0 / ENSMUSG00000026096 / ENSG00000128694 / ENSRNOG00000004001
OSTF1 / Q92882 / ENSMUSG00000024725 / ENSG00000134996 / ENSRNOG00000012156
OTC / P00480 / ENSMUSG00000031173 / ENSG00000036473 / ENSRNOG00000003370
OTUB1 / Q96FW1 / ENSMUSG00000024767 / ENSG00000167770 / ENSRNOG00000021175
OTUD1 / Q5VV17 / ENSMUSG00000043415 / ENSG00000165312 / ENSRNOG00000016950
OTUD4 / Q01804 / ENSMUSG00000036990 / ENSG00000164164 / ENSRNOG00000018477
OTUD7B / Q6GQQ9 / ENSMUSG00000038495 / ENSG00000264522 / ENSRNOG00000042068
OTULIN / Q96BN8 / ENSMUSG00000046034 / ENSG00000154124 / ENSRNOG00000012017
OXA1L / Q15070 / ENSMUSG00000000959 / ENSG00000155463 / ENSRNOG00000009713
OXCT1 / P55809 / ENSMUSG00000022186 / ENSG00000083720 / ENSRNOG00000043094
OXNAD1 / Q96HP4 / ENSMUSG00000021906 / ENSG00000154814 / ENSRNOG00000019760
OXR1 / Q8N573 / ENSMUSG00000022307 / ENSG00000164830 / ENSRNOG00000056487
OXSM / Q9NWU1 / ENSMUSG00000021786 / ENSG00000151093 / ENSRNOG00000005993
OXSR1 / O95747 / ENSMUSG00000036737 / ENSG00000172939 / ENSRNOG00000013136
P2RX4 / Q99571 / ENSMUSG00000029470 / ENSG00000135124 / ENSRNOG00000001300
P3H1 / Q32P28 / ENSMUSG00000028641 / ENSG00000117385 / ENSRNOG00000053991
Homologs
Brain
Heart
Kidney
Liver
Lung
Testis
Organs
Bins
1
2
3
4
5
Species
Human
Mouse
Rat

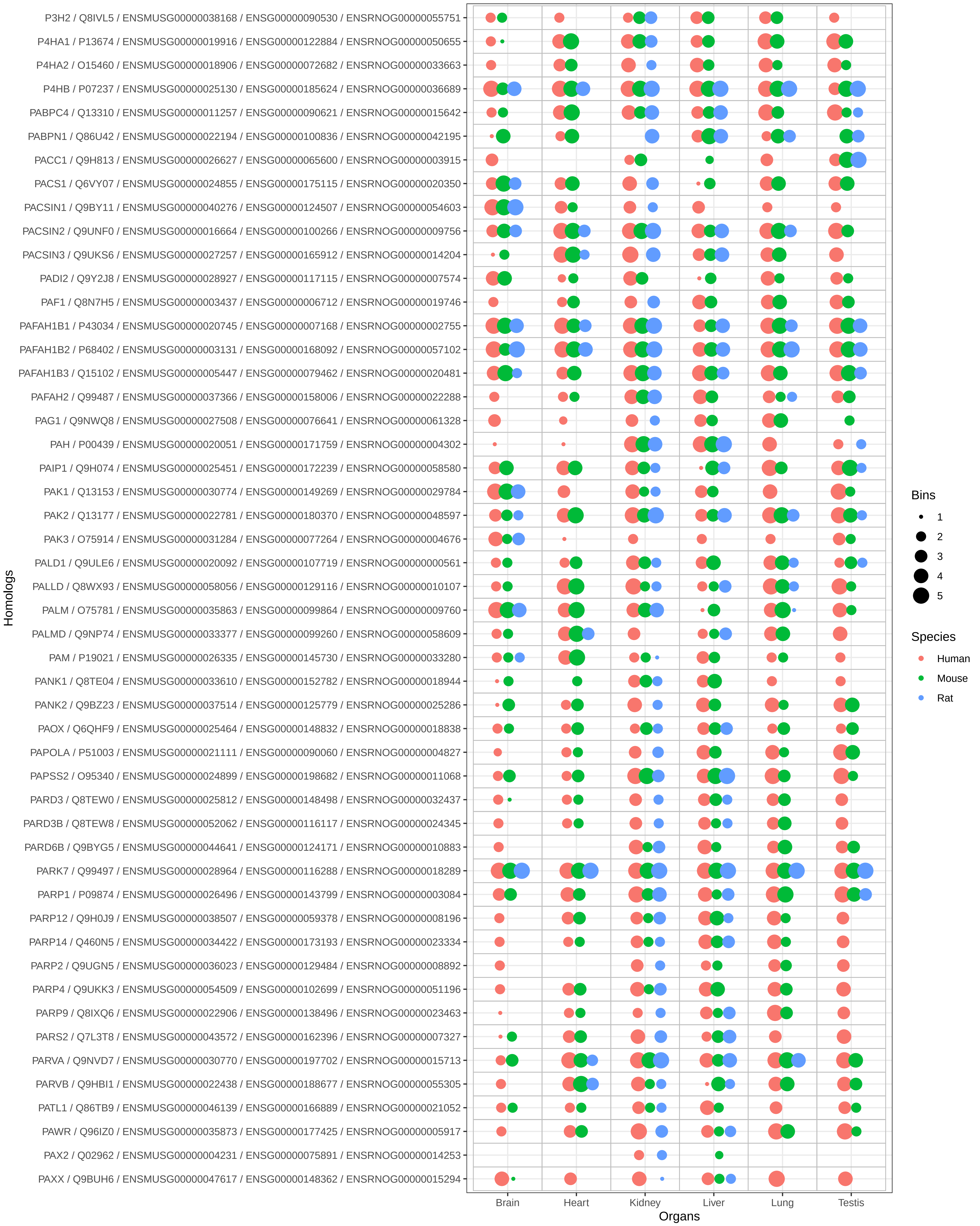
P3H2 / Q8IVL5 / ENSMUSG00000038168 / ENSG00000090530 / ENSRNOG00000055751
P4HA1 / P13674 / ENSMUSG00000019916 / ENSG00000122884 / ENSRNOG00000050655
P4HA2 / O15460 / ENSMUSG00000018906 / ENSG00000072682 / ENSRNOG00000033663
P4HB / P07237 / ENSMUSG00000025130 / ENSG00000185624 / ENSRNOG00000036689
PABPC4 / Q13310 / ENSMUSG00000011257 / ENSG00000090621 / ENSRNOG00000015642
PABPN1 / Q86U42 / ENSMUSG00000022194 / ENSG00000100836 / ENSRNOG00000042195
PACC1 / Q9H813 / ENSMUSG00000026627 / ENSG00000065600 / ENSRNOG00000003915
PACS1 / Q6VY07 / ENSMUSG00000024855 / ENSG00000175115 / ENSRNOG00000020350
PACSIN1 / Q9BY11 / ENSMUSG00000040276 / ENSG00000124507 / ENSRNOG00000054603
PACSIN2 / Q9UNF0 / ENSMUSG00000016664 / ENSG00000100266 / ENSRNOG00000009756
PACSIN3 / Q9UKS6 / ENSMUSG00000027257 / ENSG00000165912 / ENSRNOG00000014204
PADI2 / Q9Y2J8 / ENSMUSG00000028927 / ENSG00000117115 / ENSRNOG00000007574
PAF1 / Q8N7H5 / ENSMUSG00000003437 / ENSG00000006712 / ENSRNOG00000019746
PAFAH1B1 / P43034 / ENSMUSG00000020745 / ENSG00000007168 / ENSRNOG00000002755
PAFAH1B2 / P68402 / ENSMUSG00000003131 / ENSG00000168092 / ENSRNOG00000057102
PAFAH1B3 / Q15102 / ENSMUSG00000005447 / ENSG00000079462 / ENSRNOG00000020481
PAFAH2 / Q99487 / ENSMUSG00000037366 / ENSG00000158006 / ENSRNOG00000022288
PAG1 / Q9NWQ8 / ENSMUSG00000027508 / ENSG00000076641 / ENSRNOG00000061328
PAH / P00439 / ENSMUSG00000020051 / ENSG00000171759 / ENSRNOG00000004302
PAIP1 / Q9H074 / ENSMUSG00000025451 / ENSG00000172239 / ENSRNOG00000058580
PAK1 / Q13153 / ENSMUSG00000030774 / ENSG00000149269 / ENSRNOG00000029784
PAK2 / Q13177 / ENSMUSG00000022781 / ENSG00000180370 / ENSRNOG00000048597
PAK3 / O75914 / ENSMUSG00000031284 / ENSG00000077264 / ENSRNOG00000004676
PALD1 / Q9ULE6 / ENSMUSG00000020092 / ENSG00000107719 / ENSRNOG00000000561
PALLD / Q8WX93 / ENSMUSG00000058056 / ENSG00000129116 / ENSRNOG00000010107
PALM / O75781 / ENSMUSG00000035863 / ENSG00000099864 / ENSRNOG00000009760
PALMD / Q9NP74 / ENSMUSG00000033377 / ENSG00000099260 / ENSRNOG00000058609
PAM / P19021 / ENSMUSG00000026335 / ENSG00000145730 / ENSRNOG00000033280
PANK1 / Q8TE04 / ENSMUSG00000033610 / ENSG00000152782 / ENSRNOG00000018944
PANK2 / Q9BZ23 / ENSMUSG00000037514 / ENSG00000125779 / ENSRNOG00000025286
PAOX / Q6QHF9 / ENSMUSG00000025464 / ENSG00000148832 / ENSRNOG00000018838
PAPOLA / P51003 / ENSMUSG00000021111 / ENSG00000090060 / ENSRNOG00000004827
PAPSS2 / O95340 / ENSMUSG00000024899 / ENSG00000198682 / ENSRNOG00000011068
PARD3 / Q8TEW0 / ENSMUSG00000025812 / ENSG00000148498 / ENSRNOG00000032437
PARD3B / Q8TEW8 / ENSMUSG00000052062 / ENSG00000116117 / ENSRNOG00000024345
PARD6B / Q9BYG5 / ENSMUSG00000044641 / ENSG00000124171 / ENSRNOG00000010883
PARK7 / Q99497 / ENSMUSG00000028964 / ENSG00000116288 / ENSRNOG00000018289
PARP1 / P09874 / ENSMUSG00000026496 / ENSG00000143799 / ENSRNOG00000003084
PARP12 / Q9H0J9 / ENSMUSG00000038507 / ENSG00000059378 / ENSRNOG00000008196
PARP14 / Q460N5 / ENSMUSG00000034422 / ENSG00000173193 / ENSRNOG00000023334
PARP2 / Q9UGN5 / ENSMUSG00000036023 / ENSG00000129484 / ENSRNOG00000008892
PARP4 / Q9UKK3 / ENSMUSG00000054509 / ENSG00000102699 / ENSRNOG00000051196
PARP9 / Q8IXQ6 / ENSMUSG00000022906 / ENSG00000138496 / ENSRNOG00000023463
PARS2 / Q7L3T8 / ENSMUSG00000043572 / ENSG00000162396 / ENSRNOG00000007327
PARVA / Q9NVD7 / ENSMUSG00000030770 / ENSG00000197702 / ENSRNOG00000015713
PARVB / Q9HBI1 / ENSMUSG00000022438 / ENSG00000188677 / ENSRNOG00000055305
PATL1 / Q86TB9 / ENSMUSG00000046139 / ENSG00000166889 / ENSRNOG00000021052
PAWR / Q96IZ0 / ENSMUSG00000035873 / ENSG00000177425 / ENSRNOG00000005917
PAX2 / Q02962 / ENSMUSG00000004231 / ENSG00000075891 / ENSRNOG00000014253
PAXX / Q9BUH6 / ENSMUSG00000047617 / ENSG00000148362 / ENSRNOG00000015294
Homologs
Brain
Heart
Kidney
Liver
Lung
Testis
Organs
Bins
1
2
3
4
5
Species
Human
Mouse
Rat

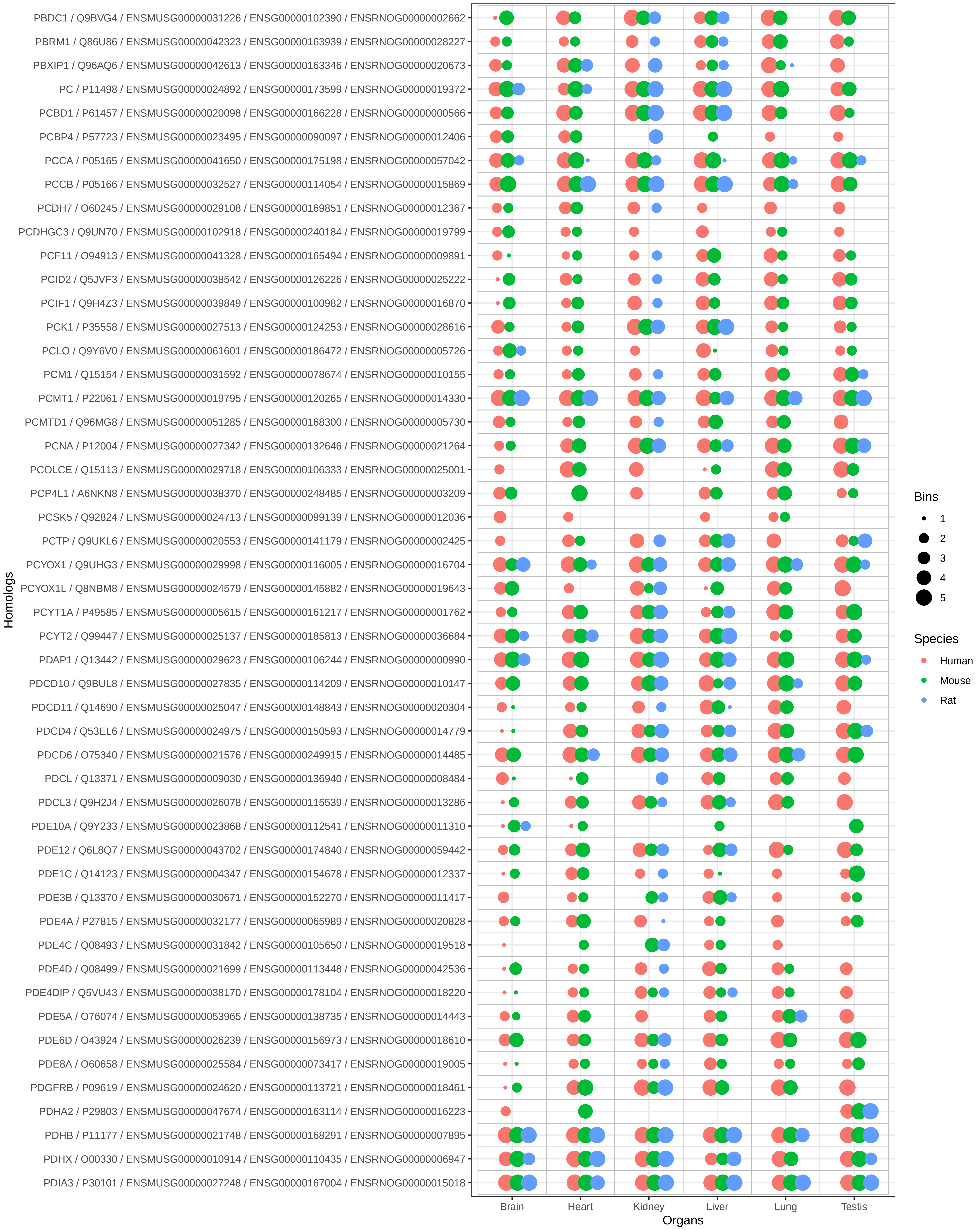
PBDC1 / Q9BVG4 / ENSMUSG00000031226 / ENSG00000102390 / ENSRNOG00000002662
PBRM1 / Q86U86 / ENSMUSG00000042323 / ENSG00000163939 / ENSRNOG00000028227
PBXIP1 / Q96AQ6 / ENSMUSG00000042613 / ENSG00000163346 / ENSRNOG00000020673
PC / P11498 / ENSMUSG00000024892 / ENSG00000173599 / ENSRNOG00000019372
PCBD1 / P61457 / ENSMUSG00000020098 / ENSG00000166228 / ENSRNOG00000000566
PCBP4 / P57723 / ENSMUSG00000023495 / ENSG00000090097 / ENSRNOG00000012406
PCCA / P05165 / ENSMUSG00000041650 / ENSG00000175198 / ENSRNOG00000057042
PCCB / P05166 / ENSMUSG00000032527 / ENSG00000114054 / ENSRNOG00000015869
PCDH7 / O60245 / ENSMUSG00000029108 / ENSG00000169851 / ENSRNOG00000012367
PCDHGC3 / Q9UN70 / ENSMUSG00000102918 / ENSG00000240184 / ENSRNOG00000019799
PCF11 / O94913 / ENSMUSG00000041328 / ENSG00000165494 / ENSRNOG00000009891
PCID2 / Q5JVF3 / ENSMUSG00000038542 / ENSG00000126226 / ENSRNOG00000025222
PCIF1 / Q9H4Z3 / ENSMUSG00000039849 / ENSG00000100982 / ENSRNOG00000016870
PCK1 / P35558 / ENSMUSG00000027513 / ENSG00000124253 / ENSRNOG00000028616
PCLO / Q9Y6V0 / ENSMUSG00000061601 / ENSG00000186472 / ENSRNOG00000005726
PCM1 / Q15154 / ENSMUSG00000031592 / ENSG00000078674 / ENSRNOG00000010155
PCMT1 / P22061 / ENSMUSG00000019795 / ENSG00000120265 / ENSRNOG00000014330
PCMTD1 / Q96MG8 / ENSMUSG00000051285 / ENSG00000168300 / ENSRNOG00000005730
PCNA / P12004 / ENSMUSG00000027342 / ENSG00000132646 / ENSRNOG00000021264
PCOLCE / Q15113 / ENSMUSG00000029718 / ENSG00000106333 / ENSRNOG00000025001
PCP4L1 / A6NKN8 / ENSMUSG00000038370 / ENSG00000248485 / ENSRNOG00000003209
PCSK5 / Q92824 / ENSMUSG00000024713 / ENSG00000099139 / ENSRNOG00000012036
PCTP / Q9UKL6 / ENSMUSG00000020553 / ENSG00000141179 / ENSRNOG00000002425
PCYOX1 / Q9UHG3 / ENSMUSG00000029998 / ENSG00000116005 / ENSRNOG00000016704
PCYOX1L / Q8NBM8 / ENSMUSG00000024579 / ENSG00000145882 / ENSRNOG00000019643
PCYT1A / P49585 / ENSMUSG00000005615 / ENSG00000161217 / ENSRNOG00000001762
PCYT2 / Q99447 / ENSMUSG00000025137 / ENSG00000185813 / ENSRNOG00000036684
PDAP1 / Q13442 / ENSMUSG00000029623 / ENSG00000106244 / ENSRNOG00000000990
PDCD10 / Q9BUL8 / ENSMUSG00000027835 / ENSG00000114209 / ENSRNOG00000010147
PDCD11 / Q14690 / ENSMUSG00000025047 / ENSG00000148843 / ENSRNOG00000020304
PDCD4 / Q53EL6 / ENSMUSG00000024975 / ENSG00000150593 / ENSRNOG00000014779
PDCD6 / O75340 / ENSMUSG00000021576 / ENSG00000249915 / ENSRNOG00000014485
PDCL / Q13371 / ENSMUSG00000009030 / ENSG00000136940 / ENSRNOG00000008484
PDCL3 / Q9H2J4 / ENSMUSG00000026078 / ENSG00000115539 / ENSRNOG00000013286
PDE10A / Q9Y233 / ENSMUSG00000023868 / ENSG00000112541 / ENSRNOG00000011310
PDE12 / Q6L8Q7 / ENSMUSG00000043702 / ENSG00000174840 / ENSRNOG00000059442
PDE1C / Q14123 / ENSMUSG00000004347 / ENSG00000154678 / ENSRNOG00000012337
PDE3B / Q13370 / ENSMUSG00000030671 / ENSG00000152270 / ENSRNOG00000011417
PDE4A / P27815 / ENSMUSG00000032177 / ENSG00000065989 / ENSRNOG00000020828
PDE4C / Q08493 / ENSMUSG00000031842 / ENSG00000105650 / ENSRNOG00000019518
PDE4D / Q08499 / ENSMUSG00000021699 / ENSG00000113448 / ENSRNOG00000042536
PDE4DIP / Q5VU43 / ENSMUSG00000038170 / ENSG00000178104 / ENSRNOG00000018220
PDE5A / O76074 / ENSMUSG00000053965 / ENSG00000138735 / ENSRNOG00000014443
PDE6D / O43924 / ENSMUSG00000026239 / ENSG00000156973 / ENSRNOG00000018610
PDE8A / O60658 / ENSMUSG00000025584 / ENSG00000073417 / ENSRNOG00000019005
PDGFRB / P09619 / ENSMUSG00000024620 / ENSG00000113721 / ENSRNOG00000018461
PDHA2 / P29803 / ENSMUSG00000047674 / ENSG00000163114 / ENSRNOG00000016223
PDHB / P11177 / ENSMUSG00000021748 / ENSG00000168291 / ENSRNOG00000007895
PDHX / O00330 / ENSMUSG00000010914 / ENSG00000110435 / ENSRNOG00000006947
PDIA3 / P30101 / ENSMUSG00000027248 / ENSG00000167004 / ENSRNOG00000015018
Homologs
Brain
Heart
Kidney
Liver
Lung
Testis
Organs
Bins
1
2
3
4
5
Species
Human
Mouse
Rat

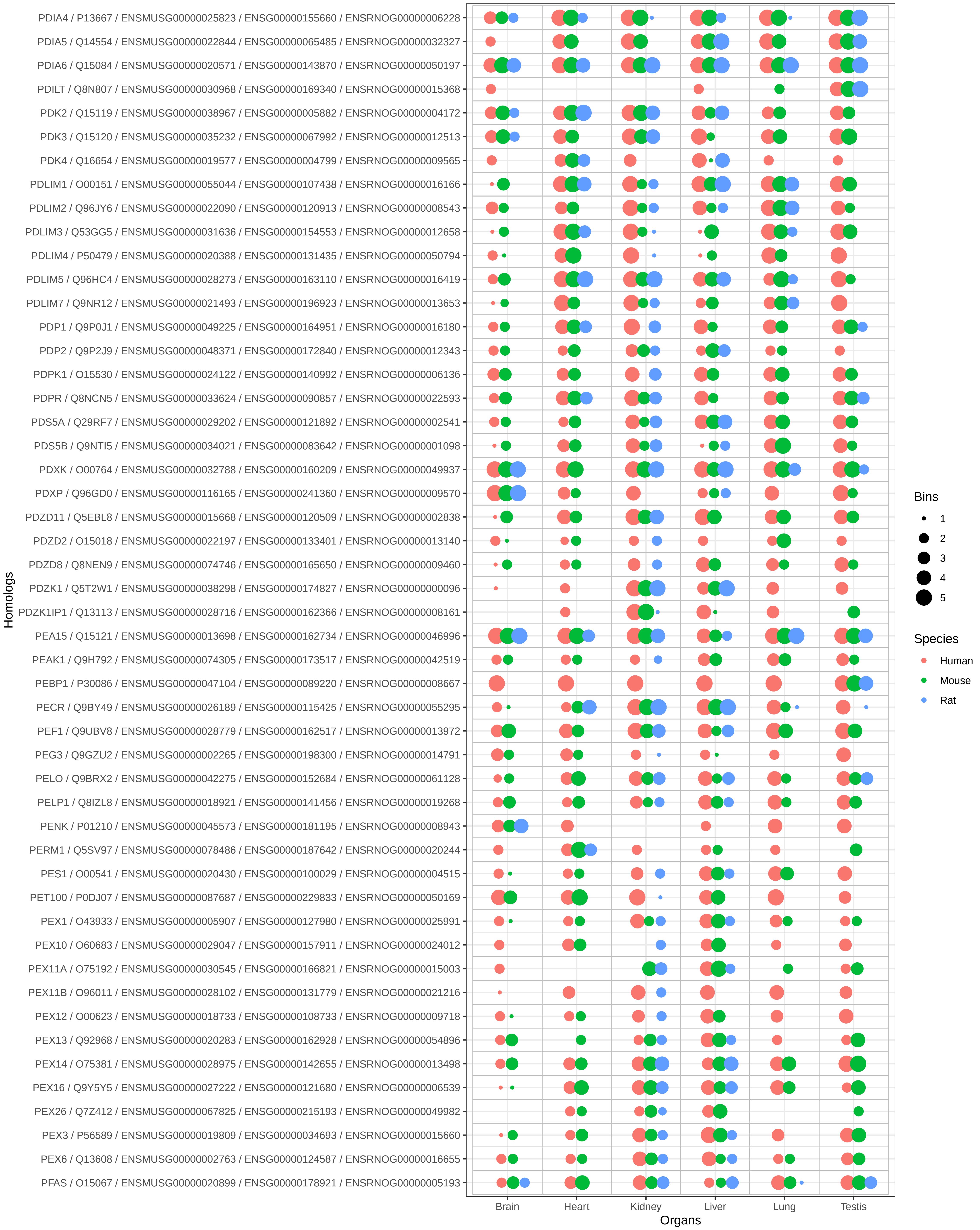
PDIA4 / P13667 / ENSMUSG00000025823 / ENSG00000155660 / ENSRNOG00000006228
PDIA5 / Q14554 / ENSMUSG00000022844 / ENSG00000065485 / ENSRNOG00000032327
PDIA6 / Q15084 / ENSMUSG00000020571 / ENSG00000143870 / ENSRNOG00000050197
PDILT / Q8N807 / ENSMUSG00000030968 / ENSG00000169340 / ENSRNOG00000015368
PDK2 / Q15119 / ENSMUSG00000038967 / ENSG00000005882 / ENSRNOG00000004172
PDK3 / Q15120 / ENSMUSG00000035232 / ENSG00000067992 / ENSRNOG00000012513
PDK4 / Q16654 / ENSMUSG00000019577 / ENSG00000004799 / ENSRNOG00000009565
PDLIM1 / O00151 / ENSMUSG00000055044 / ENSG00000107438 / ENSRNOG00000016166
PDLIM2 / Q96JY6 / ENSMUSG00000022090 / ENSG00000120913 / ENSRNOG00000008543
PDLIM3 / Q53GG5 / ENSMUSG00000031636 / ENSG00000154553 / ENSRNOG00000012658
PDLIM4 / P50479 / ENSMUSG00000020388 / ENSG00000131435 / ENSRNOG00000050794
PDLIM5 / Q96HC4 / ENSMUSG00000028273 / ENSG00000163110 / ENSRNOG00000016419
PDLIM7 / Q9NR12 / ENSMUSG00000021493 / ENSG00000196923 / ENSRNOG00000013653
PDP1 / Q9P0J1 / ENSMUSG00000049225 / ENSG00000164951 / ENSRNOG00000016180
PDP2 / Q9P2J9 / ENSMUSG00000048371 / ENSG00000172840 / ENSRNOG00000012343
PDPK1 / O15530 / ENSMUSG00000024122 / ENSG00000140992 / ENSRNOG00000006136
PDPR / Q8NCN5 / ENSMUSG00000033624 / ENSG00000090857 / ENSRNOG00000022593
PDS5A / Q29RF7 / ENSMUSG00000029202 / ENSG00000121892 / ENSRNOG00000002541
PDS5B / Q9NTI5 / ENSMUSG00000034021 / ENSG00000083642 / ENSRNOG00000001098
PDXK / O00764 / ENSMUSG00000032788 / ENSG00000160209 / ENSRNOG00000049937
PDXP / Q96GD0 / ENSMUSG00000116165 / ENSG00000241360 / ENSRNOG00000009570
PDZD11 / Q5EBL8 / ENSMUSG00000015668 / ENSG00000120509 / ENSRNOG00000002838
PDZD2 / O15018 / ENSMUSG00000022197 / ENSG00000133401 / ENSRNOG00000013140
PDZD8 / Q8NEN9 / ENSMUSG00000074746 / ENSG00000165650 / ENSRNOG00000009460
PDZK1 / Q5T2W1 / ENSMUSG00000038298 / ENSG00000174827 / ENSRNOG00000000096
PDZK1IP1 / Q13113 / ENSMUSG00000028716 / ENSG00000162366 / ENSRNOG00000008161
PEA15 / Q15121 / ENSMUSG00000013698 / ENSG00000162734 / ENSRNOG00000046996
PEAK1 / Q9H792 / ENSMUSG00000074305 / ENSG00000173517 / ENSRNOG00000042519
PEBP1 / P30086 / ENSMUSG00000047104 / ENSG00000089220 / ENSRNOG00000008667
PECR / Q9BY49 / ENSMUSG00000026189 / ENSG00000115425 / ENSRNOG00000055295
PEF1 / Q9UBV8 / ENSMUSG00000028779 / ENSG00000162517 / ENSRNOG00000013972
PEG3 / Q9GZU2 / ENSMUSG00000002265 / ENSG00000198300 / ENSRNOG00000014791
PELO / Q9BRX2 / ENSMUSG00000042275 / ENSG00000152684 / ENSRNOG00000061128
PELP1 / Q8IZL8 / ENSMUSG00000018921 / ENSG00000141456 / ENSRNOG00000019268
PENK / P01210 / ENSMUSG00000045573 / ENSG00000181195 / ENSRNOG00000008943
PERM1 / Q5SV97 / ENSMUSG00000078486 / ENSG00000187642 / ENSRNOG00000020244
PES1 / O00541 / ENSMUSG00000020430 / ENSG00000100029 / ENSRNOG00000004515
PET100 / P0DJ07 / ENSMUSG00000087687 / ENSG00000229833 / ENSRNOG00000050169
PEX1 / O43933 / ENSMUSG00000005907 / ENSG00000127980 / ENSRNOG00000025991
PEX10 / O60683 / ENSMUSG00000029047 / ENSG00000157911 / ENSRNOG00000024012
PEX11A / O75192 / ENSMUSG00000030545 / ENSG00000166821 / ENSRNOG00000015003
PEX11B / O96011 / ENSMUSG00000028102 / ENSG00000131779 / ENSRNOG00000021216
PEX12 / O00623 / ENSMUSG00000018733 / ENSG00000108733 / ENSRNOG00000009718
PEX13 / Q92968 / ENSMUSG00000020283 / ENSG00000162928 / ENSRNOG00000054896
PEX14 / O75381 / ENSMUSG00000028975 / ENSG00000142655 / ENSRNOG00000013498
PEX16 / Q9Y5Y5 / ENSMUSG00000027222 / ENSG00000121680 / ENSRNOG00000006539
PEX26 / Q7Z412 / ENSMUSG00000067825 / ENSG00000215193 / ENSRNOG00000049982
PEX3 / P56589 / ENSMUSG00000019809 / ENSG00000034693 / ENSRNOG00000015660
PEX6 / Q13608 / ENSMUSG00000002763 / ENSG00000124587 / ENSRNOG00000016655
PFAS / O15067 / ENSMUSG00000020899 / ENSG00000178921 / ENSRNOG00000005193
Homologs
Brain
Heart
Kidney
Liver
Lung
Testis
Organs
Bins
1
2
3
4
5
Species
Human
Mouse
Rat

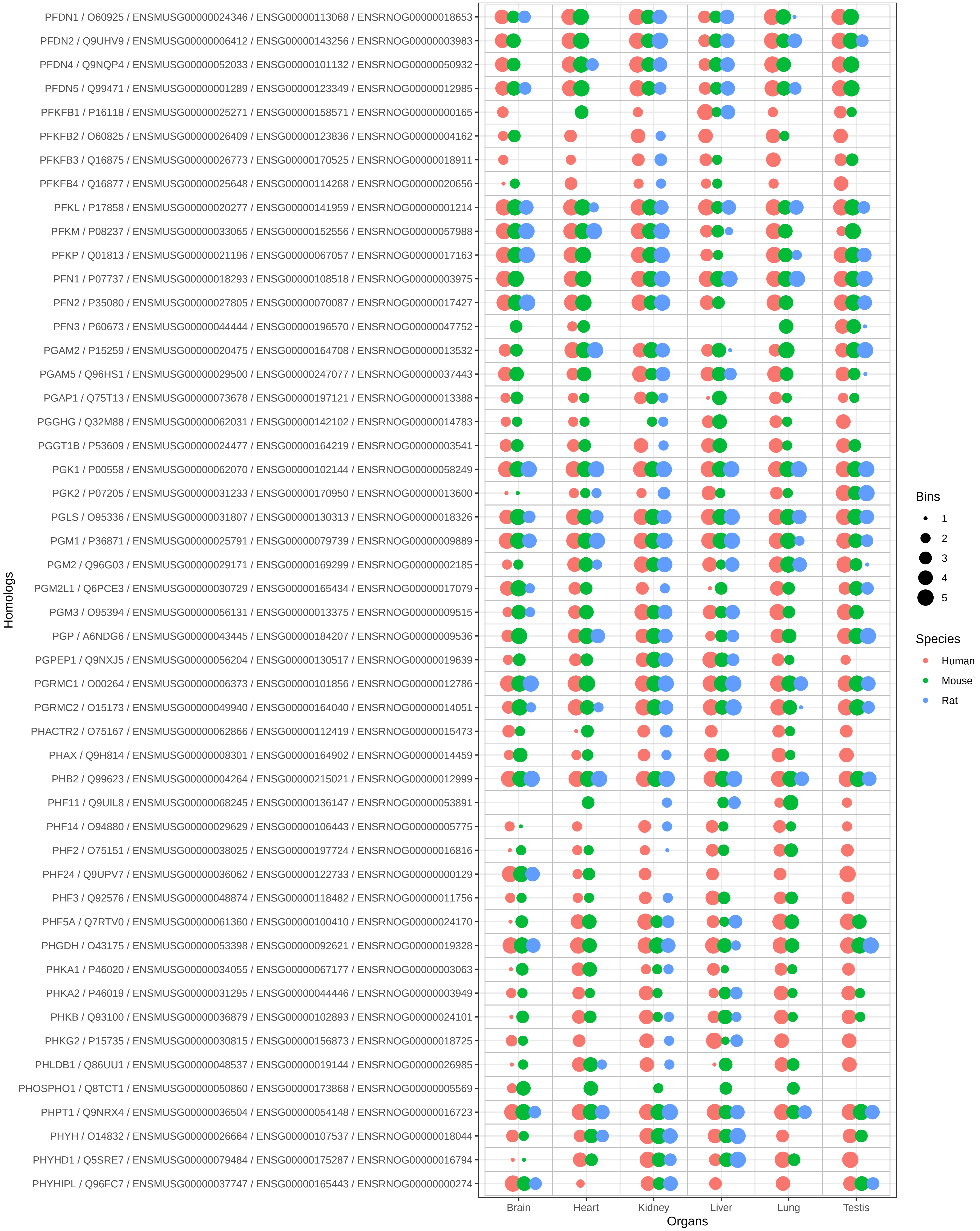
PFDN1 / O60925 / ENSMUSG00000024346 / ENSG00000113068 / ENSRNOG00000018653
PFDN2 / Q9UHV9 / ENSMUSG00000006412 / ENSG00000143256 / ENSRNOG00000003983
PFDN4 / Q9NQP4 / ENSMUSG00000052033 / ENSG00000101132 / ENSRNOG00000050932
PFDN5 / Q99471 / ENSMUSG00000001289 / ENSG00000123349 / ENSRNOG00000012985
PFKFB1 / P16118 / ENSMUSG00000025271 / ENSG00000158571 / ENSRNOG00000000165
PFKFB2 / O60825 / ENSMUSG00000026409 / ENSG00000123836 / ENSRNOG00000004162
PFKFB3 / Q16875 / ENSMUSG00000026773 / ENSG00000170525 / ENSRNOG00000018911
PFKFB4 / Q16877 / ENSMUSG00000025648 / ENSG00000114268 / ENSRNOG00000020656
PFKL / P17858 / ENSMUSG00000020277 / ENSG00000141959 / ENSRNOG00000001214
PFKM / P08237 / ENSMUSG00000033065 / ENSG00000152556 / ENSRNOG00000057988
PFKP / Q01813 / ENSMUSG00000021196 / ENSG00000067057 / ENSRNOG00000017163
PFN1 / P07737 / ENSMUSG00000018293 / ENSG00000108518 / ENSRNOG00000003975
PFN2 / P35080 / ENSMUSG00000027805 / ENSG00000070087 / ENSRNOG00000017427
PFN3 / P60673 / ENSMUSG00000044444 / ENSG00000196570 / ENSRNOG00000047752
PGAM2 / P15259 / ENSMUSG00000020475 / ENSG00000164708 / ENSRNOG00000013532
PGAM5 / Q96HS1 / ENSMUSG00000029500 / ENSG00000247077 / ENSRNOG00000037443
PGAP1 / Q75T13 / ENSMUSG00000073678 / ENSG00000197121 / ENSRNOG00000013388
PGGHG / Q32M88 / ENSMUSG00000062031 / ENSG00000142102 / ENSRNOG00000014783
PGGT1B / P53609 / ENSMUSG00000024477 / ENSG00000164219 / ENSRNOG00000003541
PGK1 / P00558 / ENSMUSG00000062070 / ENSG00000102144 / ENSRNOG00000058249
PGK2 / P07205 / ENSMUSG00000031233 / ENSG00000170950 / ENSRNOG00000013600
PGLS / O95336 / ENSMUSG00000031807 / ENSG00000130313 / ENSRNOG00000018326
PGM1 / P36871 / ENSMUSG00000025791 / ENSG00000079739 / ENSRNOG00000009889
PGM2 / Q96G03 / ENSMUSG00000029171 / ENSG00000169299 / ENSRNOG00000002185
PGM2L1 / Q6PCE3 / ENSMUSG00000030729 / ENSG00000165434 / ENSRNOG00000017079
PGM3 / O95394 / ENSMUSG00000056131 / ENSG00000013375 / ENSRNOG00000009515
PGP / A6NDG6 / ENSMUSG00000043445 / ENSG00000184207 / ENSRNOG00000009536
PGPEP1 / Q9NXJ5 / ENSMUSG00000056204 / ENSG00000130517 / ENSRNOG00000019639
PGRMC1 / O00264 / ENSMUSG00000006373 / ENSG00000101856 / ENSRNOG00000012786
PGRMC2 / O15173 / ENSMUSG00000049940 / ENSG00000164040 / ENSRNOG00000014051
PHACTR2 / O75167 / ENSMUSG00000062866 / ENSG00000112419 / ENSRNOG00000015473
PHAX / Q9H814 / ENSMUSG00000008301 / ENSG00000164902 / ENSRNOG00000014459
PHB2 / Q99623 / ENSMUSG00000004264 / ENSG00000215021 / ENSRNOG00000012999
PHF11 / Q9UIL8 / ENSMUSG00000068245 / ENSG00000136147 / ENSRNOG00000053891
PHF14 / O94880 / ENSMUSG00000029629 / ENSG00000106443 / ENSRNOG00000005775
PHF2 / O75151 / ENSMUSG00000038025 / ENSG00000197724 / ENSRNOG00000016816
PHF24 / Q9UPV7 / ENSMUSG00000036062 / ENSG00000122733 / ENSRNOG00000000129
PHF3 / Q92576 / ENSMUSG00000048874 / ENSG00000118482 / ENSRNOG00000011756
PHF5A / Q7RTV0 / ENSMUSG00000061360 / ENSG00000100410 / ENSRNOG00000024170
PHGDH / O43175 / ENSMUSG00000053398 / ENSG00000092621 / ENSRNOG00000019328
PHKA1 / P46020 / ENSMUSG00000034055 / ENSG00000067177 / ENSRNOG00000003063
PHKA2 / P46019 / ENSMUSG00000031295 / ENSG00000044446 / ENSRNOG00000003949
PHKB / Q93100 / ENSMUSG00000036879 / ENSG00000102893 / ENSRNOG00000024101
PHKG2 / P15735 / ENSMUSG00000030815 / ENSG00000156873 / ENSRNOG00000018725
PHLDB1 / Q86UU1 / ENSMUSG00000048537 / ENSG00000019144 / ENSRNOG00000026985
PHOSPHO1 / Q8TCT1 / ENSMUSG00000050860 / ENSG00000173868 / ENSRNOG00000005569
PHPT1 / Q9NRX4 / ENSMUSG00000036504 / ENSG00000054148 / ENSRNOG00000016723
PHYH / O14832 / ENSMUSG00000026664 / ENSG00000107537 / ENSRNOG00000018044
PHYHD1 / Q5SRE7 / ENSMUSG00000079484 / ENSG00000175287 / ENSRNOG00000016794
PHYHIPL / Q96FC7 / ENSMUSG00000037747 / ENSG00000165443 / ENSRNOG00000000274
Homologs
Brain
Heart
Kidney
Liver
Lung
Testis
Organs
Bins
1
2
3
4
5
Species
Human
Mouse
Rat

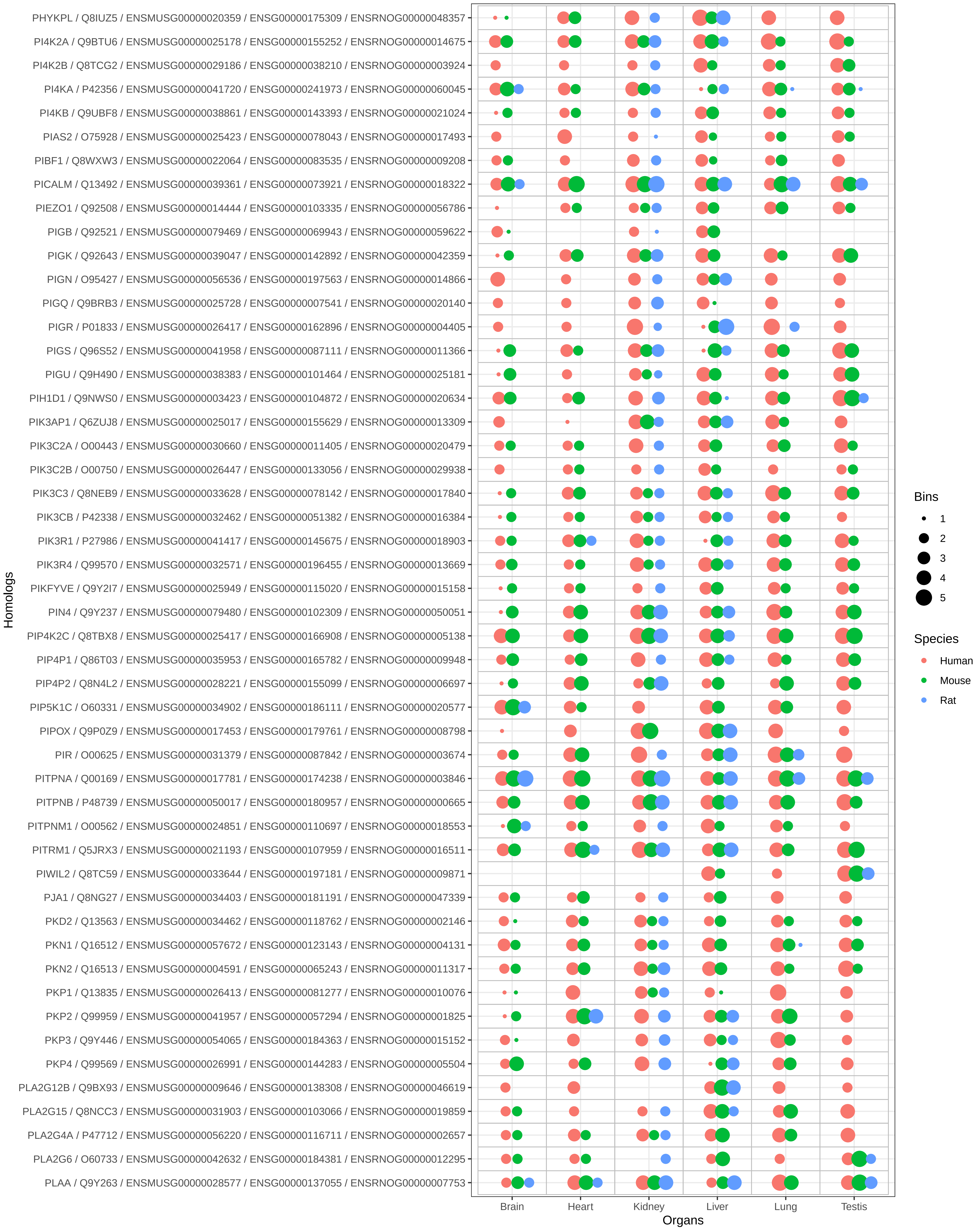
PHYKPL / Q8IUZ5 / ENSMUSG00000020359 / ENSG00000175309 / ENSRNOG00000048357
PI4K2A / Q9BTU6 / ENSMUSG00000025178 / ENSG00000155252 / ENSRNOG00000014675
PI4K2B / Q8TCG2 / ENSMUSG00000029186 / ENSG00000038210 / ENSRNOG00000003924
PI4KA / P42356 / ENSMUSG00000041720 / ENSG00000241973 / ENSRNOG00000060045
PI4KB / Q9UBF8 / ENSMUSG00000038861 / ENSG00000143393 / ENSRNOG00000021024
PIAS2 / O75928 / ENSMUSG00000025423 / ENSG00000078043 / ENSRNOG00000017493
PIBF1 / Q8WXW3 / ENSMUSG00000022064 / ENSG00000083535 / ENSRNOG00000009208
PICALM / Q13492 / ENSMUSG00000039361 / ENSG00000073921 / ENSRNOG00000018322
PIEZO1 / Q92508 / ENSMUSG00000014444 / ENSG00000103335 / ENSRNOG00000056786
PIGB / Q92521 / ENSMUSG00000079469 / ENSG00000069943 / ENSRNOG00000059622
PIGK / Q92643 / ENSMUSG00000039047 / ENSG00000142892 / ENSRNOG00000042359
PIGN / O95427 / ENSMUSG00000056536 / ENSG00000197563 / ENSRNOG00000014866
PIGQ / Q9BRB3 / ENSMUSG00000025728 / ENSG00000007541 / ENSRNOG00000020140
PIGR / P01833 / ENSMUSG00000026417 / ENSG00000162896 / ENSRNOG00000004405
PIGS / Q96S52 / ENSMUSG00000041958 / ENSG00000087111 / ENSRNOG00000011366
PIGU / Q9H490 / ENSMUSG00000038383 / ENSG00000101464 / ENSRNOG00000025181
PIH1D1 / Q9NWS0 / ENSMUSG00000003423 / ENSG00000104872 / ENSRNOG00000020634
PIK3AP1 / Q6ZUJ8 / ENSMUSG00000025017 / ENSG00000155629 / ENSRNOG00000013309
PIK3C2A / O00443 / ENSMUSG00000030660 / ENSG00000011405 / ENSRNOG00000020479
PIK3C2B / O00750 / ENSMUSG00000026447 / ENSG00000133056 / ENSRNOG00000029938
PIK3C3 / Q8NEB9 / ENSMUSG00000033628 / ENSG00000078142 / ENSRNOG00000017840
PIK3CB / P42338 / ENSMUSG00000032462 / ENSG00000051382 / ENSRNOG00000016384
PIK3R1 / P27986 / ENSMUSG00000041417 / ENSG00000145675 / ENSRNOG00000018903
PIK3R4 / Q99570 / ENSMUSG00000032571 / ENSG00000196455 / ENSRNOG00000013669
PIKFYVE / Q9Y2I7 / ENSMUSG00000025949 / ENSG00000115020 / ENSRNOG00000015158
PIN4 / Q9Y237 / ENSMUSG00000079480 / ENSG00000102309 / ENSRNOG00000050051
PIP4K2C / Q8TBX8 / ENSMUSG00000025417 / ENSG00000166908 / ENSRNOG00000005138
PIP4P1 / Q86T03 / ENSMUSG00000035953 / ENSG00000165782 / ENSRNOG00000009948
PIP4P2 / Q8N4L2 / ENSMUSG00000028221 / ENSG00000155099 / ENSRNOG00000006697
PIP5K1C / O60331 / ENSMUSG00000034902 / ENSG00000186111 / ENSRNOG00000020577
PIPOX / Q9P0Z9 / ENSMUSG00000017453 / ENSG00000179761 / ENSRNOG00000008798
PIR / O00625 / ENSMUSG00000031379 / ENSG00000087842 / ENSRNOG00000003674
PITPNA / Q00169 / ENSMUSG00000017781 / ENSG00000174238 / ENSRNOG00000003846
PITPNB / P48739 / ENSMUSG00000050017 / ENSG00000180957 / ENSRNOG00000000665
PITPNM1 / O00562 / ENSMUSG00000024851 / ENSG00000110697 / ENSRNOG00000018553
PITRM1 / Q5JRX3 / ENSMUSG00000021193 / ENSG00000107959 / ENSRNOG00000016511
PIWIL2 / Q8TC59 / ENSMUSG00000033644 / ENSG00000197181 / ENSRNOG00000009871
PJA1 / Q8NG27 / ENSMUSG00000034403 / ENSG00000181191 / ENSRNOG00000047339
PKD2 / Q13563 / ENSMUSG00000034462 / ENSG00000118762 / ENSRNOG00000002146
PKN1 / Q16512 / ENSMUSG00000057672 / ENSG00000123143 / ENSRNOG00000004131
PKN2 / Q16513 / ENSMUSG00000004591 / ENSG00000065243 / ENSRNOG00000011317
PKP1 / Q13835 / ENSMUSG00000026413 / ENSG00000081277 / ENSRNOG00000010076
PKP2 / Q99959 / ENSMUSG00000041957 / ENSG00000057294 / ENSRNOG00000001825
PKP3 / Q9Y446 / ENSMUSG00000054065 / ENSG00000184363 / ENSRNOG00000015152
PKP4 / Q99569 / ENSMUSG00000026991 / ENSG00000144283 / ENSRNOG00000005504
PLA2G12B / Q9BX93 / ENSMUSG00000009646 / ENSG00000138308 / ENSRNOG00000046619
PLA2G15 / Q8NCC3 / ENSMUSG00000031903 / ENSG00000103066 / ENSRNOG00000019859
PLA2G4A / P47712 / ENSMUSG00000056220 / ENSG00000116711 / ENSRNOG00000002657
PLA2G6 / O60733 / ENSMUSG00000042632 / ENSG00000184381 / ENSRNOG00000012295
PLAA / Q9Y263 / ENSMUSG00000028577 / ENSG00000137055 / ENSRNOG00000007753
Homologs
Brain
Heart
Kidney
Liver
Lung
Testis
Organs
Bins
1
2
3
4
5
Species
Human
Mouse
Rat

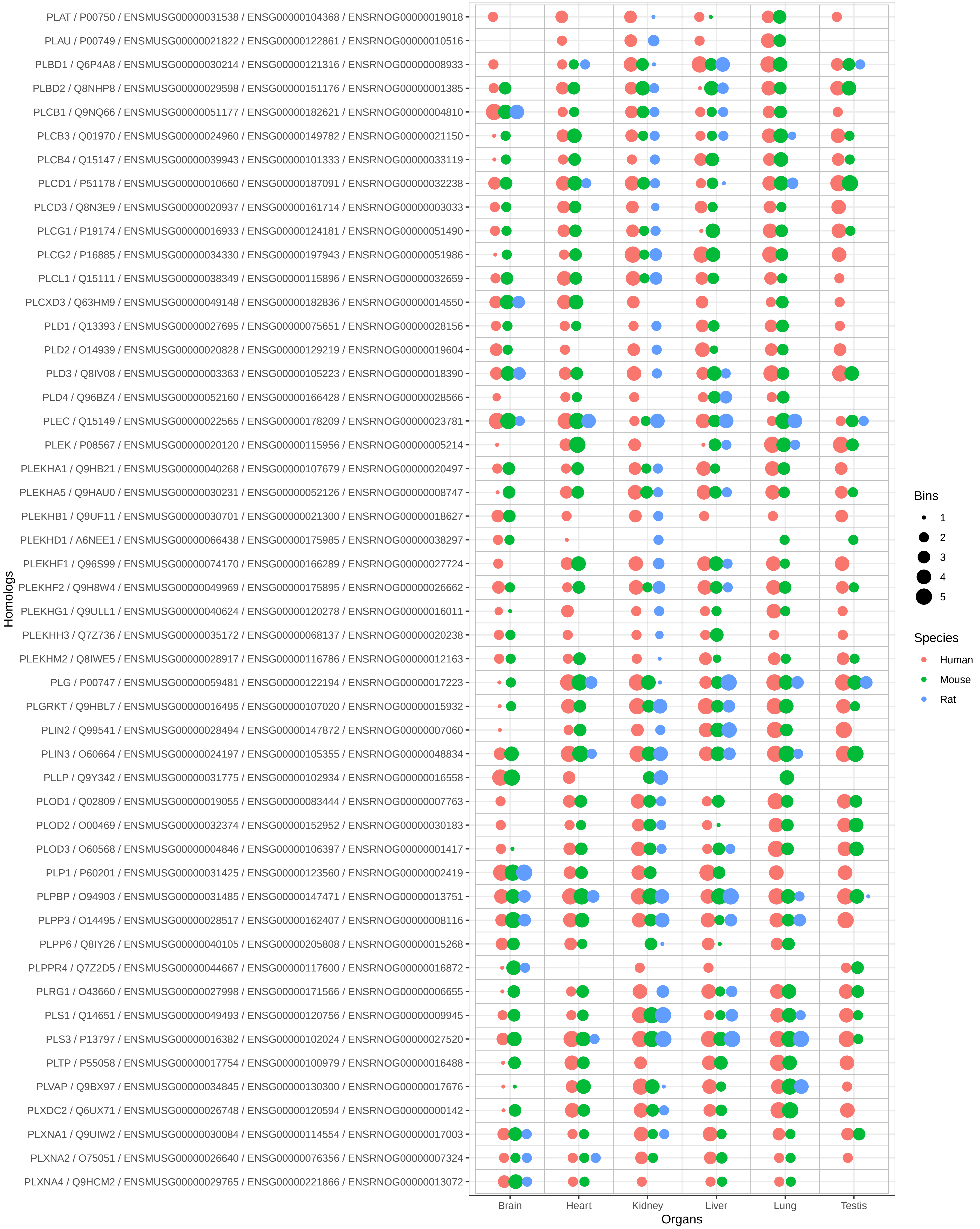
PLAT / P00750 / ENSMUSG00000031538 / ENSG00000104368 / ENSRNOG00000019018
PLAU / P00749 / ENSMUSG00000021822 / ENSG00000122861 / ENSRNOG00000010516
PLBD1 / Q6P4A8 / ENSMUSG00000030214 / ENSG00000121316 / ENSRNOG00000008933
PLBD2 / Q8NHP8 / ENSMUSG00000029598 / ENSG00000151176 / ENSRNOG00000001385
PLCB1 / Q9NQ66 / ENSMUSG00000051177 / ENSG00000182621 / ENSRNOG00000004810
PLCB3 / Q01970 / ENSMUSG00000024960 / ENSG00000149782 / ENSRNOG00000021150
PLCB4 / Q15147 / ENSMUSG00000039943 / ENSG00000101333 / ENSRNOG00000033119
PLCD1 / P51178 / ENSMUSG00000010660 / ENSG00000187091 / ENSRNOG00000032238
PLCD3 / Q8N3E9 / ENSMUSG00000020937 / ENSG00000161714 / ENSRNOG00000003033
PLCG1 / P19174 / ENSMUSG00000016933 / ENSG00000124181 / ENSRNOG00000051490
PLCG2 / P16885 / ENSMUSG00000034330 / ENSG00000197943 / ENSRNOG00000051986
PLCL1 / Q15111 / ENSMUSG00000038349 / ENSG00000115896 / ENSRNOG00000032659
PLCXD3 / Q63HM9 / ENSMUSG00000049148 / ENSG00000182836 / ENSRNOG00000014550
PLD1 / Q13393 / ENSMUSG00000027695 / ENSG00000075651 / ENSRNOG00000028156
PLD2 / O14939 / ENSMUSG00000020828 / ENSG00000129219 / ENSRNOG00000019604
PLD3 / Q8IV08 / ENSMUSG00000003363 / ENSG00000105223 / ENSRNOG00000018390
PLD4 / Q96BZ4 / ENSMUSG00000052160 / ENSG00000166428 / ENSRNOG00000028566
PLEC / Q15149 / ENSMUSG00000022565 / ENSG00000178209 / ENSRNOG00000023781
PLEK / P08567 / ENSMUSG00000020120 / ENSG00000115956 / ENSRNOG00000005214
PLEKHA1 / Q9HB21 / ENSMUSG00000040268 / ENSG00000107679 / ENSRNOG00000020497
PLEKHA5 / Q9HAU0 / ENSMUSG00000030231 / ENSG00000052126 / ENSRNOG00000008747
PLEKHB1 / Q9UF11 / ENSMUSG00000030701 / ENSG00000021300 / ENSRNOG00000018627
PLEKHD1 / A6NEE1 / ENSMUSG00000066438 / ENSG00000175985 / ENSRNOG00000038297
PLEKHF1 / Q96S99 / ENSMUSG00000074170 / ENSG00000166289 / ENSRNOG00000027724
PLEKHF2 / Q9H8W4 / ENSMUSG00000049969 / ENSG00000175895 / ENSRNOG00000026662
PLEKHG1 / Q9ULL1 / ENSMUSG00000040624 / ENSG00000120278 / ENSRNOG00000016011
PLEKHH3 / Q7Z736 / ENSMUSG00000035172 / ENSG00000068137 / ENSRNOG00000020238
PLEKHM2 / Q8IWE5 / ENSMUSG00000028917 / ENSG00000116786 / ENSRNOG00000012163
PLG / P00747 / ENSMUSG00000059481 / ENSG00000122194 / ENSRNOG00000017223
PLGRKT / Q9HBL7 / ENSMUSG00000016495 / ENSG00000107020 / ENSRNOG00000015932
PLIN2 / Q99541 / ENSMUSG00000028494 / ENSG00000147872 / ENSRNOG00000007060
PLIN3 / O60664 / ENSMUSG00000024197 / ENSG00000105355 / ENSRNOG00000048834
PLLP / Q9Y342 / ENSMUSG00000031775 / ENSG00000102934 / ENSRNOG00000016558
PLOD1 / Q02809 / ENSMUSG00000019055 / ENSG00000083444 / ENSRNOG00000007763
PLOD2 / O00469 / ENSMUSG00000032374 / ENSG00000152952 / ENSRNOG00000030183
PLOD3 / O60568 / ENSMUSG00000004846 / ENSG00000106397 / ENSRNOG00000001417
PLP1 / P60201 / ENSMUSG00000031425 / ENSG00000123560 / ENSRNOG00000002419
PLPBP / O94903 / ENSMUSG00000031485 / ENSG00000147471 / ENSRNOG00000013751
PLPP3 / O14495 / ENSMUSG00000028517 / ENSG00000162407 / ENSRNOG00000008116
PLPP6 / Q8IY26 / ENSMUSG00000040105 / ENSG00000205808 / ENSRNOG00000015268
PLPPR4 / Q7Z2D5 / ENSMUSG00000044667 / ENSG00000117600 / ENSRNOG00000016872
PLRG1 / O43660 / ENSMUSG00000027998 / ENSG00000171566 / ENSRNOG00000006655
PLS1 / Q14651 / ENSMUSG00000049493 / ENSG00000120756 / ENSRNOG00000009945
PLS3 / P13797 / ENSMUSG00000016382 / ENSG00000102024 / ENSRNOG00000027520
PLTP / P55058 / ENSMUSG00000017754 / ENSG00000100979 / ENSRNOG00000016488
PLVAP / Q9BX97 / ENSMUSG00000034845 / ENSG00000130300 / ENSRNOG00000017676
PLXDC2 / Q6UX71 / ENSMUSG00000026748 / ENSG00000120594 / ENSRNOG00000000142
PLXNA1 / Q9UIW2 / ENSMUSG00000030084 / ENSG00000114554 / ENSRNOG00000017003
PLXNA2 / O75051 / ENSMUSG00000026640 / ENSG00000076356 / ENSRNOG00000007324
PLXNA4 / Q9HCM2 / ENSMUSG00000029765 / ENSG00000221866 / ENSRNOG00000013072
Homologs
Brain
Heart
Kidney
Liver
Lung
Testis
Organs
Bins
1
2
3
4
5
Species
Human
Mouse
Rat

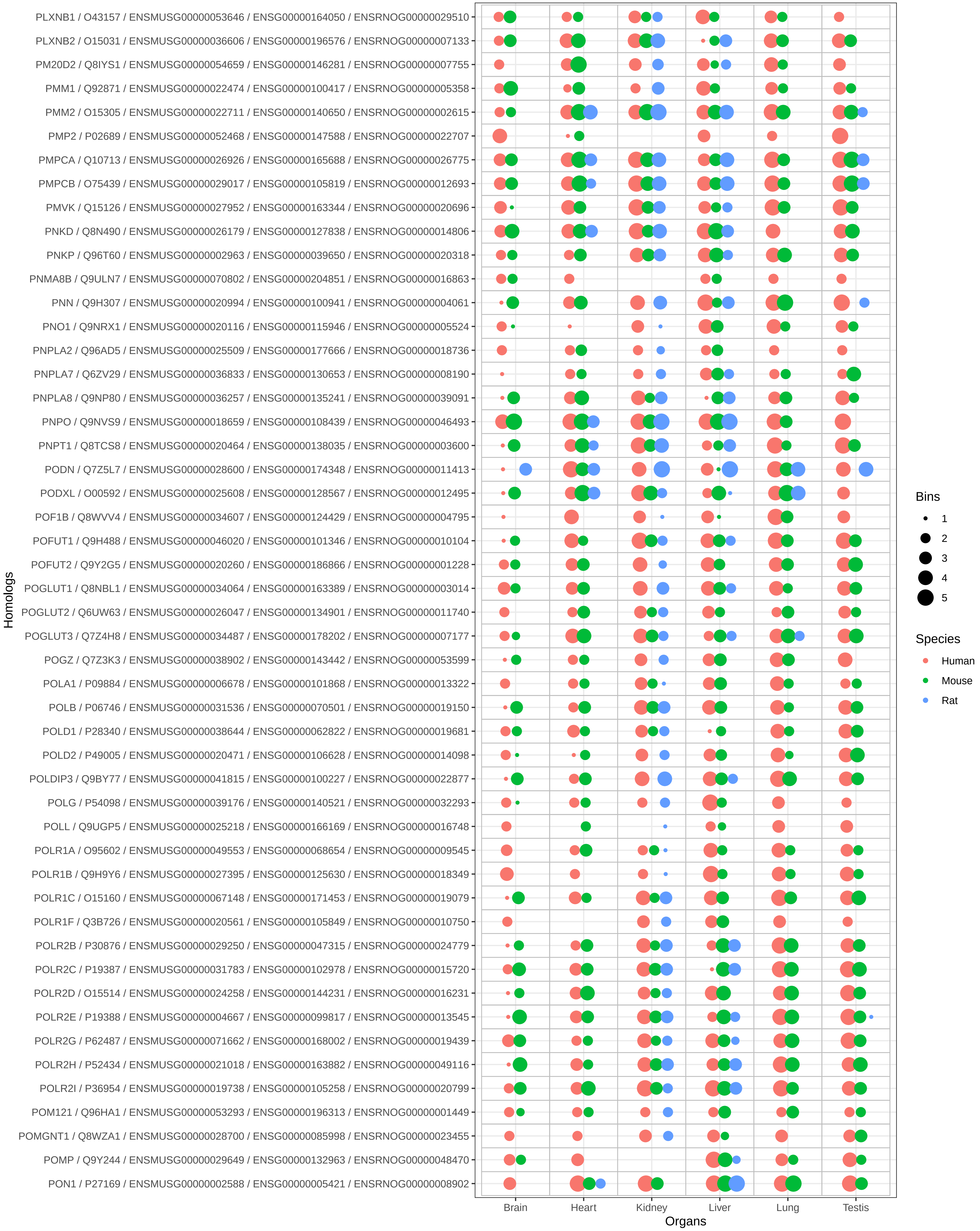
PLXNB1 / O43157 / ENSMUSG00000053646 / ENSG00000164050 / ENSRNOG00000029510
PLXNB2 / O15031 / ENSMUSG00000036606 / ENSG00000196576 / ENSRNOG00000007133
PM20D2 / Q8IYS1 / ENSMUSG00000054659 / ENSG00000146281 / ENSRNOG00000007755
PMM1 / Q92871 / ENSMUSG00000022474 / ENSG00000100417 / ENSRNOG00000005358
PMM2 / O15305 / ENSMUSG00000022711 / ENSG00000140650 / ENSRNOG00000002615
PMP2 / P02689 / ENSMUSG00000052468 / ENSG00000147588 / ENSRNOG00000022707
PMPCA / Q10713 / ENSMUSG00000026926 / ENSG00000165688 / ENSRNOG00000026775
PMPCB / O75439 / ENSMUSG00000029017 / ENSG00000105819 / ENSRNOG00000012693
PMVK / Q15126 / ENSMUSG00000027952 / ENSG00000163344 / ENSRNOG00000020696
PNKD / Q8N490 / ENSMUSG00000026179 / ENSG00000127838 / ENSRNOG00000014806
PNKP / Q96T60 / ENSMUSG00000002963 / ENSG00000039650 / ENSRNOG00000020318
PNMA8B / Q9ULN7 / ENSMUSG00000070802 / ENSG00000204851 / ENSRNOG00000016863
PNN / Q9H307 / ENSMUSG00000020994 / ENSG00000100941 / ENSRNOG00000004061
PNO1 / Q9NRX1 / ENSMUSG00000020116 / ENSG00000115946 / ENSRNOG00000005524
PNPLA2 / Q96AD5 / ENSMUSG00000025509 / ENSG00000177666 / ENSRNOG00000018736
PNPLA7 / Q6ZV29 / ENSMUSG00000036833 / ENSG00000130653 / ENSRNOG00000008190
PNPLA8 / Q9NP80 / ENSMUSG00000036257 / ENSG00000135241 / ENSRNOG00000039091
PNPO / Q9NVS9 / ENSMUSG00000018659 / ENSG00000108439 / ENSRNOG00000046493
PNPT1 / Q8TCS8 / ENSMUSG00000020464 / ENSG00000138035 / ENSRNOG00000003600
PODN / Q7Z5L7 / ENSMUSG00000028600 / ENSG00000174348 / ENSRNOG00000011413
PODXL / O00592 / ENSMUSG00000025608 / ENSG00000128567 / ENSRNOG00000012495
POF1B / Q8WVV4 / ENSMUSG00000034607 / ENSG00000124429 / ENSRNOG00000004795
POFUT1 / Q9H488 / ENSMUSG00000046020 / ENSG00000101346 / ENSRNOG00000010104
POFUT2 / Q9Y2G5 / ENSMUSG00000020260 / ENSG00000186866 / ENSRNOG00000001228
POGLUT1 / Q8NBL1 / ENSMUSG00000034064 / ENSG00000163389 / ENSRNOG00000003014
POGLUT2 / Q6UW63 / ENSMUSG00000026047 / ENSG00000134901 / ENSRNOG00000011740
POGLUT3 / Q7Z4H8 / ENSMUSG00000034487 / ENSG00000178202 / ENSRNOG00000007177
POGZ / Q7Z3K3 / ENSMUSG00000038902 / ENSG00000143442 / ENSRNOG00000053599
POLA1 / P09884 / ENSMUSG00000006678 / ENSG00000101868 / ENSRNOG00000013322
POLB / P06746 / ENSMUSG00000031536 / ENSG00000070501 / ENSRNOG00000019150
POLD1 / P28340 / ENSMUSG00000038644 / ENSG00000062822 / ENSRNOG00000019681
POLD2 / P49005 / ENSMUSG00000020471 / ENSG00000106628 / ENSRNOG00000014098
POLDIP3 / Q9BY77 / ENSMUSG00000041815 / ENSG00000100227 / ENSRNOG00000022877
POLG / P54098 / ENSMUSG00000039176 / ENSG00000140521 / ENSRNOG00000032293
POLL / Q9UGP5 / ENSMUSG00000025218 / ENSG00000166169 / ENSRNOG00000016748
POLR1A / O95602 / ENSMUSG00000049553 / ENSG00000068654 / ENSRNOG00000009545
POLR1B / Q9H9Y6 / ENSMUSG00000027395 / ENSG00000125630 / ENSRNOG00000018349
POLR1C / O15160 / ENSMUSG00000067148 / ENSG00000171453 / ENSRNOG00000019079
POLR1F / Q3B726 / ENSMUSG00000020561 / ENSG00000105849 / ENSRNOG00000010750
POLR2B / P30876 / ENSMUSG00000029250 / ENSG00000047315 / ENSRNOG00000024779
POLR2C / P19387 / ENSMUSG00000031783 / ENSG00000102978 / ENSRNOG00000015720
POLR2D / O15514 / ENSMUSG00000024258 / ENSG00000144231 / ENSRNOG00000016231
POLR2E / P19388 / ENSMUSG00000004667 / ENSG00000099817 / ENSRNOG00000013545
POLR2G / P62487 / ENSMUSG00000071662 / ENSG00000168002 / ENSRNOG00000019439
POLR2H / P52434 / ENSMUSG00000021018 / ENSG00000163882 / ENSRNOG00000049116
POLR2I / P36954 / ENSMUSG00000019738 / ENSG00000105258 / ENSRNOG00000020799
POM121 / Q96HA1 / ENSMUSG00000053293 / ENSG00000196313 / ENSRNOG00000001449
POMGNT1 / Q8WZA1 / ENSMUSG00000028700 / ENSG00000085998 / ENSRNOG00000023455
POMP / Q9Y244 / ENSMUSG00000029649 / ENSG00000132963 / ENSRNOG00000048470
PON1 / P27169 / ENSMUSG00000002588 / ENSG00000005421 / ENSRNOG00000008902
Homologs
Brain
Heart
Kidney
Liver
Lung
Testis
Organs
Bins
1
2
3
4
5
Species
Human
Mouse
Rat

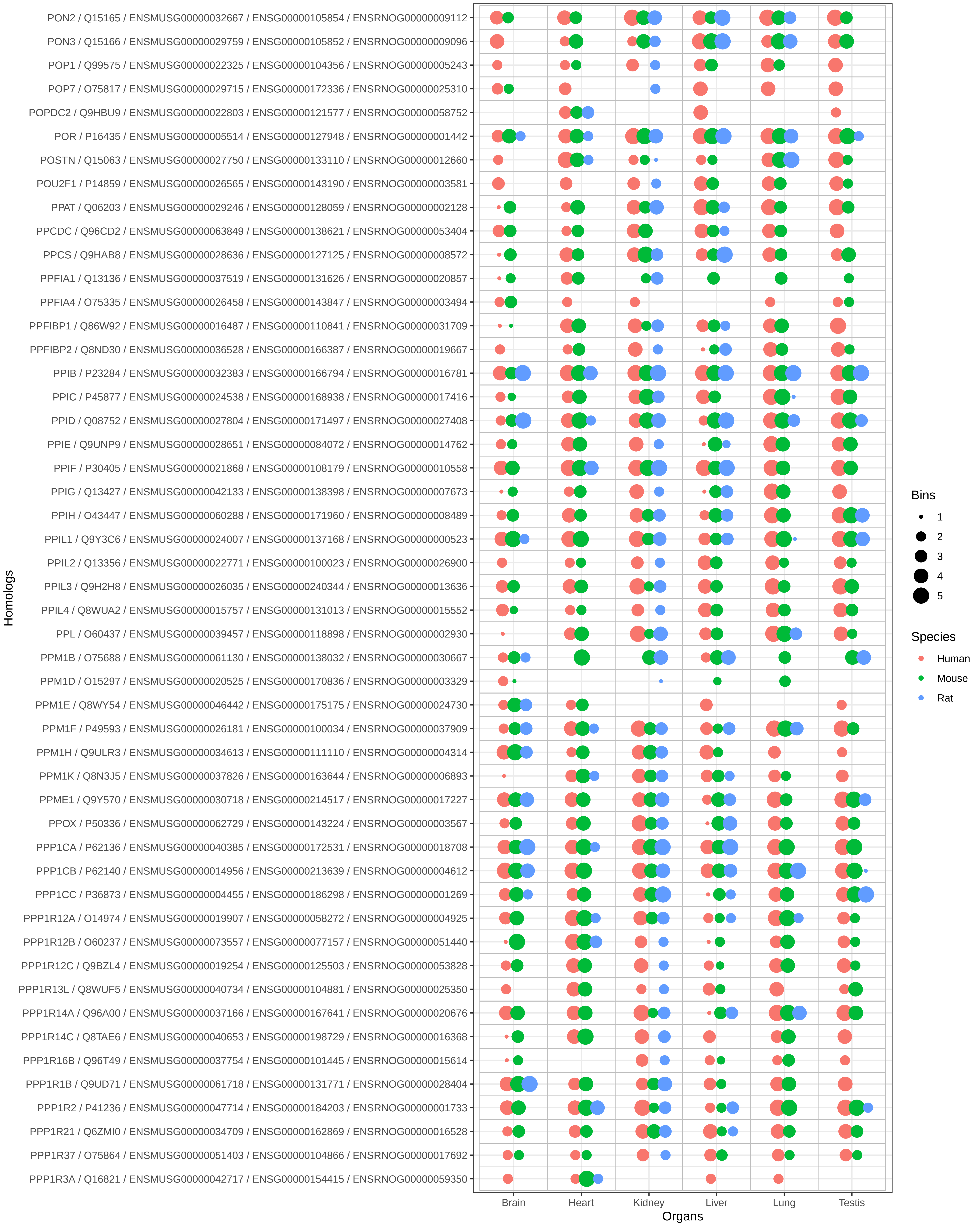
PON2 / Q15165 / ENSMUSG00000032667 / ENSG00000105854 / ENSRNOG00000009112
PON3 / Q15166 / ENSMUSG00000029759 / ENSG00000105852 / ENSRNOG00000009096
POP1 / Q99575 / ENSMUSG00000022325 / ENSG00000104356 / ENSRNOG00000005243
POP7 / O75817 / ENSMUSG00000029715 / ENSG00000172336 / ENSRNOG00000025310
POPDC2 / Q9HBU9 / ENSMUSG00000022803 / ENSG00000121577 / ENSRNOG00000058752
POR / P16435 / ENSMUSG00000005514 / ENSG00000127948 / ENSRNOG00000001442
POSTN / Q15063 / ENSMUSG00000027750 / ENSG00000133110 / ENSRNOG00000012660
POU2F1 / P14859 / ENSMUSG00000026565 / ENSG00000143190 / ENSRNOG00000003581
PPAT / Q06203 / ENSMUSG00000029246 / ENSG00000128059 / ENSRNOG00000002128
PPCDC / Q96CD2 / ENSMUSG00000063849 / ENSG00000138621 / ENSRNOG00000053404
PPCS / Q9HAB8 / ENSMUSG00000028636 / ENSG00000127125 / ENSRNOG00000008572
PPFIA1 / Q13136 / ENSMUSG00000037519 / ENSG00000131626 / ENSRNOG00000020857
PPFIA4 / O75335 / ENSMUSG00000026458 / ENSG00000143847 / ENSRNOG00000003494
PPFIBP1 / Q86W92 / ENSMUSG00000016487 / ENSG00000110841 / ENSRNOG00000031709
PPFIBP2 / Q8ND30 / ENSMUSG00000036528 / ENSG00000166387 / ENSRNOG00000019667
PPIB / P23284 / ENSMUSG00000032383 / ENSG00000166794 / ENSRNOG00000016781
PPIC / P45877 / ENSMUSG00000024538 / ENSG00000168938 / ENSRNOG00000017416
PPID / Q08752 / ENSMUSG00000027804 / ENSG00000171497 / ENSRNOG00000027408
PPIE / Q9UNP9 / ENSMUSG00000028651 / ENSG00000084072 / ENSRNOG00000014762
PPIF / P30405 / ENSMUSG00000021868 / ENSG00000108179 / ENSRNOG00000010558
PPIG / Q13427 / ENSMUSG00000042133 / ENSG00000138398 / ENSRNOG00000007673
PPIH / O43447 / ENSMUSG00000060288 / ENSG00000171960 / ENSRNOG00000008489
PPIL1 / Q9Y3C6 / ENSMUSG00000024007 / ENSG00000137168 / ENSRNOG00000000523
PPIL2 / Q13356 / ENSMUSG00000022771 / ENSG00000100023 / ENSRNOG00000026900
PPIL3 / Q9H2H8 / ENSMUSG00000026035 / ENSG00000240344 / ENSRNOG00000013636
PPIL4 / Q8WUA2 / ENSMUSG00000015757 / ENSG00000131013 / ENSRNOG00000015552
PPL / O60437 / ENSMUSG00000039457 / ENSG00000118898 / ENSRNOG00000002930
PPM1B / O75688 / ENSMUSG00000061130 / ENSG00000138032 / ENSRNOG00000030667
PPM1D / O15297 / ENSMUSG00000020525 / ENSG00000170836 / ENSRNOG00000003329
PPM1E / Q8WY54 / ENSMUSG00000046442 / ENSG00000175175 / ENSRNOG00000024730
PPM1F / P49593 / ENSMUSG00000026181 / ENSG00000100034 / ENSRNOG00000037909
PPM1H / Q9ULR3 / ENSMUSG00000034613 / ENSG00000111110 / ENSRNOG00000004314
PPM1K / Q8N3J5 / ENSMUSG00000037826 / ENSG00000163644 / ENSRNOG00000006893
PPME1 / Q9Y570 / ENSMUSG00000030718 / ENSG00000214517 / ENSRNOG00000017227
PPOX / P50336 / ENSMUSG00000062729 / ENSG00000143224 / ENSRNOG00000003567
PPP1CA / P62136 / ENSMUSG00000040385 / ENSG00000172531 / ENSRNOG00000018708
PPP1CB / P62140 / ENSMUSG00000014956 / ENSG00000213639 / ENSRNOG00000004612
PPP1CC / P36873 / ENSMUSG00000004455 / ENSG00000186298 / ENSRNOG00000001269
PPP1R12A / O14974 / ENSMUSG00000019907 / ENSG00000058272 / ENSRNOG00000004925
PPP1R12B / O60237 / ENSMUSG00000073557 / ENSG00000077157 / ENSRNOG00000051440
PPP1R12C / Q9BZL4 / ENSMUSG00000019254 / ENSG00000125503 / ENSRNOG00000053828
PPP1R13L / Q8WUF5 / ENSMUSG00000040734 / ENSG00000104881 / ENSRNOG00000025350
PPP1R14A / Q96A00 / ENSMUSG00000037166 / ENSG00000167641 / ENSRNOG00000020676
PPP1R14C / Q8TAE6 / ENSMUSG00000040653 / ENSG00000198729 / ENSRNOG00000016368
PPP1R16B / Q96T49 / ENSMUSG00000037754 / ENSG00000101445 / ENSRNOG00000015614
PPP1R1B / Q9UD71 / ENSMUSG00000061718 / ENSG00000131771 / ENSRNOG00000028404
PPP1R2 / P41236 / ENSMUSG00000047714 / ENSG00000184203 / ENSRNOG00000001733
PPP1R21 / Q6ZMI0 / ENSMUSG00000034709 / ENSG00000162869 / ENSRNOG00000016528
PPP1R37 / O75864 / ENSMUSG00000051403 / ENSG00000104866 / ENSRNOG00000017692
PPP1R3A / Q16821 / ENSMUSG00000042717 / ENSG00000154415 / ENSRNOG00000059350
Homologs
Brain
Heart
Kidney
Liver
Lung
Testis
Organs
Bins
1
2
3
4
5
Species
Human
Mouse
Rat

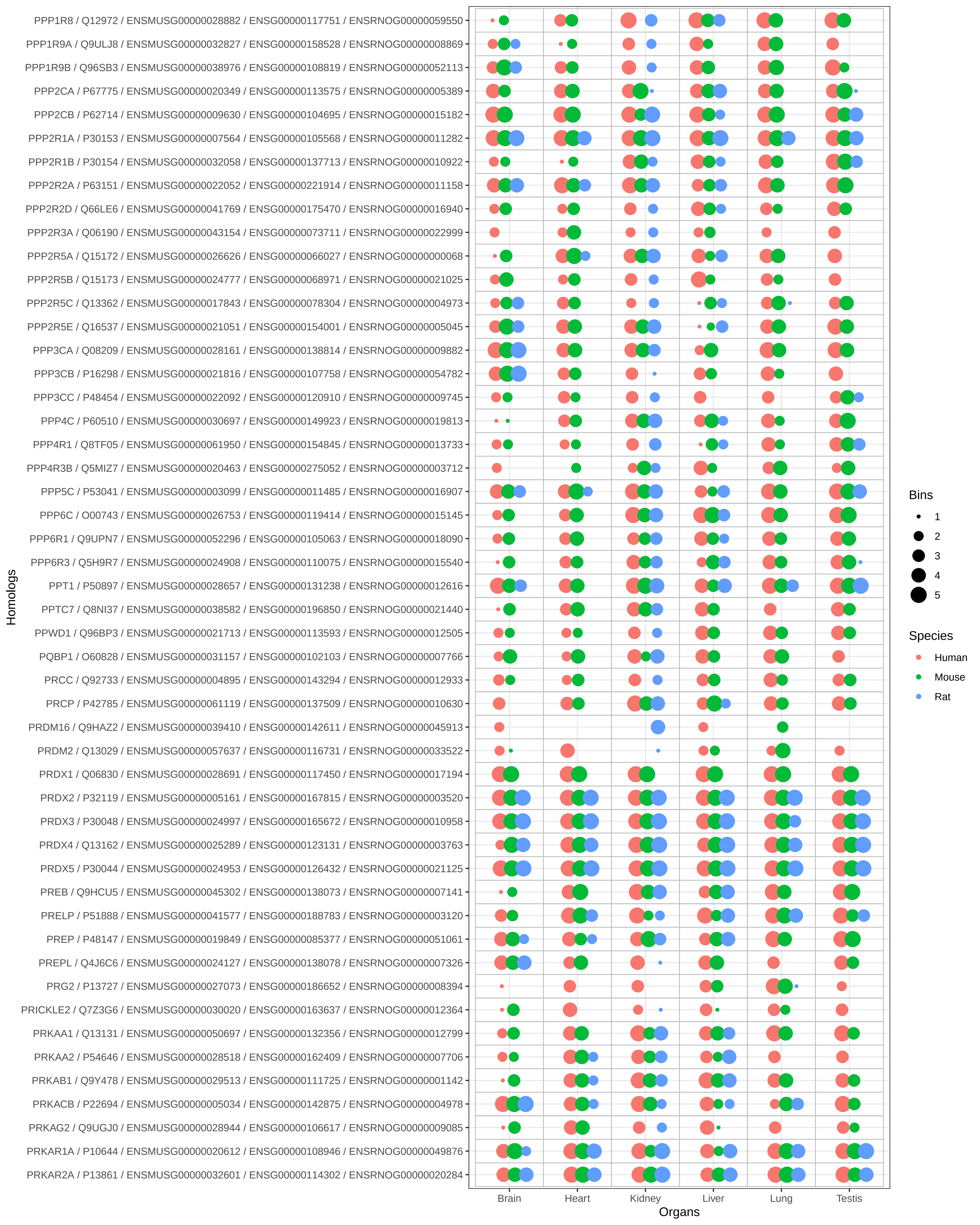
PPP1R8 / Q12972 / ENSMUSG00000028882 / ENSG00000117751 / ENSRNOG00000059550
PPP1R9A / Q9ULJ8 / ENSMUSG00000032827 / ENSG00000158528 / ENSRNOG00000008869
PPP1R9B / Q96SB3 / ENSMUSG00000038976 / ENSG00000108819 / ENSRNOG00000052113
PPP2CA / P67775 / ENSMUSG00000020349 / ENSG00000113575 / ENSRNOG00000005389
PPP2CB / P62714 / ENSMUSG00000009630 / ENSG00000104695 / ENSRNOG00000015182
PPP2R1A / P30153 / ENSMUSG00000007564 / ENSG00000105568 / ENSRNOG00000011282
PPP2R1B / P30154 / ENSMUSG00000032058 / ENSG00000137713 / ENSRNOG00000010922
PPP2R2A / P63151 / ENSMUSG00000022052 / ENSG00000221914 / ENSRNOG00000011158
PPP2R2D / Q66LE6 / ENSMUSG00000041769 / ENSG00000175470 / ENSRNOG00000016940
PPP2R3A / Q06190 / ENSMUSG00000043154 / ENSG00000073711 / ENSRNOG00000022999
PPP2R5A / Q15172 / ENSMUSG00000026626 / ENSG00000066027 / ENSRNOG00000000068
PPP2R5B / Q15173 / ENSMUSG00000024777 / ENSG00000068971 / ENSRNOG00000021025
PPP2R5C / Q13362 / ENSMUSG00000017843 / ENSG00000078304 / ENSRNOG00000004973
PPP2R5E / Q16537 / ENSMUSG00000021051 / ENSG00000154001 / ENSRNOG00000005045
PPP3CA / Q08209 / ENSMUSG00000028161 / ENSG00000138814 / ENSRNOG00000009882
PPP3CB / P16298 / ENSMUSG00000021816 / ENSG00000107758 / ENSRNOG00000054782
PPP3CC / P48454 / ENSMUSG00000022092 / ENSG00000120910 / ENSRNOG00000009745
PPP4C / P60510 / ENSMUSG00000030697 / ENSG00000149923 / ENSRNOG00000019813
PPP4R1 / Q8TF05 / ENSMUSG00000061950 / ENSG00000154845 / ENSRNOG00000013733
PPP4R3B / Q5MIZ7 / ENSMUSG00000020463 / ENSG00000275052 / ENSRNOG00000003712
PPP5C / P53041 / ENSMUSG00000003099 / ENSG00000011485 / ENSRNOG00000016907
PPP6C / O00743 / ENSMUSG00000026753 / ENSG00000119414 / ENSRNOG00000015145
PPP6R1 / Q9UPN7 / ENSMUSG00000052296 / ENSG00000105063 / ENSRNOG00000018090
PPP6R3 / Q5H9R7 / ENSMUSG00000024908 / ENSG00000110075 / ENSRNOG00000015540
PPT1 / P50897 / ENSMUSG00000028657 / ENSG00000131238 / ENSRNOG00000012616
PPTC7 / Q8NI37 / ENSMUSG00000038582 / ENSG00000196850 / ENSRNOG00000021440
PPWD1 / Q96BP3 / ENSMUSG00000021713 / ENSG00000113593 / ENSRNOG00000012505
PQBP1 / O60828 / ENSMUSG00000031157 / ENSG00000102103 / ENSRNOG00000007766
PRCC / Q92733 / ENSMUSG00000004895 / ENSG00000143294 / ENSRNOG00000012933
PRCP / P42785 / ENSMUSG00000061119 / ENSG00000137509 / ENSRNOG00000010630
PRDM16 / Q9HAZ2 / ENSMUSG00000039410 / ENSG00000142611 / ENSRNOG00000045913
PRDM2 / Q13029 / ENSMUSG00000057637 / ENSG00000116731 / ENSRNOG00000033522
PRDX1 / Q06830 / ENSMUSG00000028691 / ENSG00000117450 / ENSRNOG00000017194
PRDX2 / P32119 / ENSMUSG00000005161 / ENSG00000167815 / ENSRNOG00000003520
PRDX3 / P30048 / ENSMUSG00000024997 / ENSG00000165672 / ENSRNOG00000010958
PRDX4 / Q13162 / ENSMUSG00000025289 / ENSG00000123131 / ENSRNOG00000003763
PRDX5 / P30044 / ENSMUSG00000024953 / ENSG00000126432 / ENSRNOG00000021125
PREB / Q9HCU5 / ENSMUSG00000045302 / ENSG00000138073 / ENSRNOG00000007141
PRELP / P51888 / ENSMUSG00000041577 / ENSG00000188783 / ENSRNOG00000003120
PREP / P48147 / ENSMUSG00000019849 / ENSG00000085377 / ENSRNOG00000051061
PREPL / Q4J6C6 / ENSMUSG00000024127 / ENSG00000138078 / ENSRNOG00000007326
PRG2 / P13727 / ENSMUSG00000027073 / ENSG00000186652 / ENSRNOG00000008394
PRICKLE2 / Q7Z3G6 / ENSMUSG00000030020 / ENSG00000163637 / ENSRNOG00000012364
PRKAA1 / Q13131 / ENSMUSG00000050697 / ENSG00000132356 / ENSRNOG00000012799
PRKAA2 / P54646 / ENSMUSG00000028518 / ENSG00000162409 / ENSRNOG00000007706
PRKAB1 / Q9Y478 / ENSMUSG00000029513 / ENSG00000111725 / ENSRNOG00000001142
PRKACB / P22694 / ENSMUSG00000005034 / ENSG00000142875 / ENSRNOG00000004978
PRKAG2 / Q9UGJ0 / ENSMUSG00000028944 / ENSG00000106617 / ENSRNOG00000009085
PRKAR1A / P10644 / ENSMUSG00000020612 / ENSG00000108946 / ENSRNOG00000049876
PRKAR2A / P13861 / ENSMUSG00000032601 / ENSG00000114302 / ENSRNOG00000020284
Homologs
Brain
Heart
Kidney
Liver
Lung
Testis
Organs
Bins
1
2
3
4
5
Species
Human
Mouse
Rat

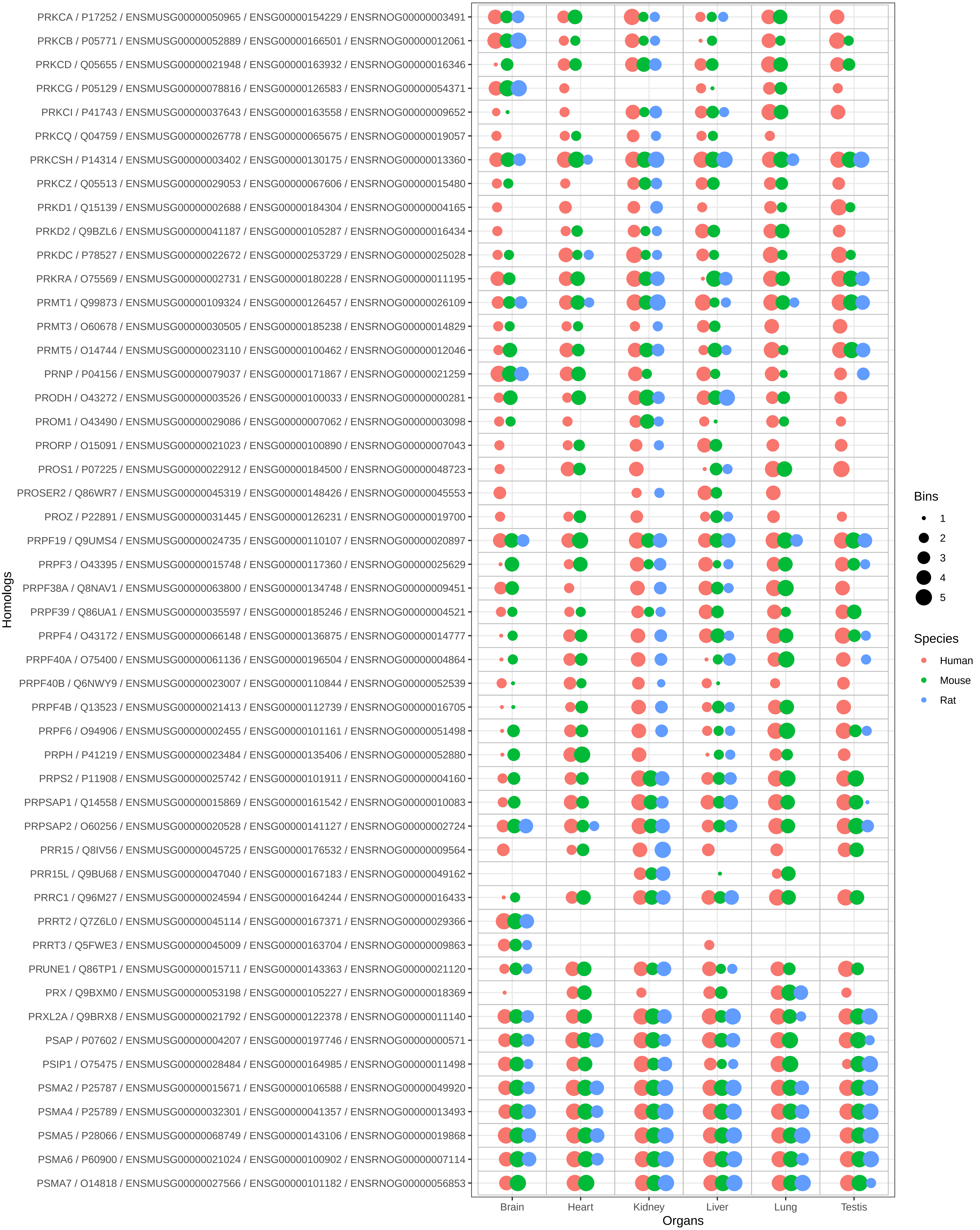
PRKCA / P17252 / ENSMUSG00000050965 / ENSG00000154229 / ENSRNOG00000003491
PRKCB / P05771 / ENSMUSG00000052889 / ENSG00000166501 / ENSRNOG00000012061
PRKCD / Q05655 / ENSMUSG00000021948 / ENSG00000163932 / ENSRNOG00000016346
PRKCG / P05129 / ENSMUSG00000078816 / ENSG00000126583 / ENSRNOG00000054371
PRKCI / P41743 / ENSMUSG00000037643 / ENSG00000163558 / ENSRNOG00000009652
PRKCQ / Q04759 / ENSMUSG00000026778 / ENSG00000065675 / ENSRNOG00000019057
PRKCSH / P14314 / ENSMUSG00000003402 / ENSG00000130175 / ENSRNOG00000013360
PRKCZ / Q05513 / ENSMUSG00000029053 / ENSG00000067606 / ENSRNOG00000015480
PRKD1 / Q15139 / ENSMUSG00000002688 / ENSG00000184304 / ENSRNOG00000004165
PRKD2 / Q9BZL6 / ENSMUSG00000041187 / ENSG00000105287 / ENSRNOG00000016434
PRKDC / P78527 / ENSMUSG00000022672 / ENSG00000253729 / ENSRNOG00000025028
PRKRA / O75569 / ENSMUSG00000002731 / ENSG00000180228 / ENSRNOG00000011195
PRMT1 / Q99873 / ENSMUSG00000109324 / ENSG00000126457 / ENSRNOG00000026109
PRMT3 / O60678 / ENSMUSG00000030505 / ENSG00000185238 / ENSRNOG00000014829
PRMT5 / O14744 / ENSMUSG00000023110 / ENSG00000100462 / ENSRNOG00000012046
PRNP / P04156 / ENSMUSG00000079037 / ENSG00000171867 / ENSRNOG00000021259
PRODH / O43272 / ENSMUSG00000003526 / ENSG00000100033 / ENSRNOG00000000281
PROM1 / O43490 / ENSMUSG00000029086 / ENSG00000007062 / ENSRNOG00000003098
PRORP / O15091 / ENSMUSG00000021023 / ENSG00000100890 / ENSRNOG00000007043
PROS1 / P07225 / ENSMUSG00000022912 / ENSG00000184500 / ENSRNOG00000048723
PROSER2 / Q86WR7 / ENSMUSG00000045319 / ENSG00000148426 / ENSRNOG00000045553
PROZ / P22891 / ENSMUSG00000031445 / ENSG00000126231 / ENSRNOG00000019700
PRPF19 / Q9UMS4 / ENSMUSG00000024735 / ENSG00000110107 / ENSRNOG00000020897
PRPF3 / O43395 / ENSMUSG00000015748 / ENSG00000117360 / ENSRNOG00000025629
PRPF38A / Q8NAV1 / ENSMUSG00000063800 / ENSG00000134748 / ENSRNOG00000009451
PRPF39 / Q86UA1 / ENSMUSG00000035597 / ENSG00000185246 / ENSRNOG00000004521
PRPF4 / O43172 / ENSMUSG00000066148 / ENSG00000136875 / ENSRNOG00000014777
PRPF40A / O75400 / ENSMUSG00000061136 / ENSG00000196504 / ENSRNOG00000004864
PRPF40B / Q6NWY9 / ENSMUSG00000023007 / ENSG00000110844 / ENSRNOG00000052539
PRPF4B / Q13523 / ENSMUSG00000021413 / ENSG00000112739 / ENSRNOG00000016705
PRPF6 / O94906 / ENSMUSG00000002455 / ENSG00000101161 / ENSRNOG00000051498
PRPH / P41219 / ENSMUSG00000023484 / ENSG00000135406 / ENSRNOG00000052880
PRPS2 / P11908 / ENSMUSG00000025742 / ENSG00000101911 / ENSRNOG00000004160
PRPSAP1 / Q14558 / ENSMUSG00000015869 / ENSG00000161542 / ENSRNOG00000010083
PRPSAP2 / O60256 / ENSMUSG00000020528 / ENSG00000141127 / ENSRNOG00000002724
PRR15 / Q8IV56 / ENSMUSG00000045725 / ENSG00000176532 / ENSRNOG00000009564
PRR15L / Q9BU68 / ENSMUSG00000047040 / ENSG00000167183 / ENSRNOG00000049162
PRRC1 / Q96M27 / ENSMUSG00000024594 / ENSG00000164244 / ENSRNOG00000016433
PRRT2 / Q7Z6L0 / ENSMUSG00000045114 / ENSG00000167371 / ENSRNOG00000029366
PRRT3 / Q5FWE3 / ENSMUSG00000045009 / ENSG00000163704 / ENSRNOG00000009863
PRUNE1 / Q86TP1 / ENSMUSG00000015711 / ENSG00000143363 / ENSRNOG00000021120
PRX / Q9BXM0 / ENSMUSG00000053198 / ENSG00000105227 / ENSRNOG00000018369
PRXL2A / Q9BRX8 / ENSMUSG00000021792 / ENSG00000122378 / ENSRNOG00000011140
PSAP / P07602 / ENSMUSG00000004207 / ENSG00000197746 / ENSRNOG00000000571
PSIP1 / O75475 / ENSMUSG00000028484 / ENSG00000164985 / ENSRNOG00000011498
PSMA2 / P25787 / ENSMUSG00000015671 / ENSG00000106588 / ENSRNOG00000049920
PSMA4 / P25789 / ENSMUSG00000032301 / ENSG00000041357 / ENSRNOG00000013493
PSMA5 / P28066 / ENSMUSG00000068749 / ENSG00000143106 / ENSRNOG00000019868
PSMA6 / P60900 / ENSMUSG00000021024 / ENSG00000100902 / ENSRNOG00000007114
PSMA7 / O14818 / ENSMUSG00000027566 / ENSG00000101182 / ENSRNOG00000056853
Homologs
Brain
Heart
Kidney
Liver
Lung
Testis
Organs
Bins
1
2
3
4
5
Species
Human
Mouse
Rat

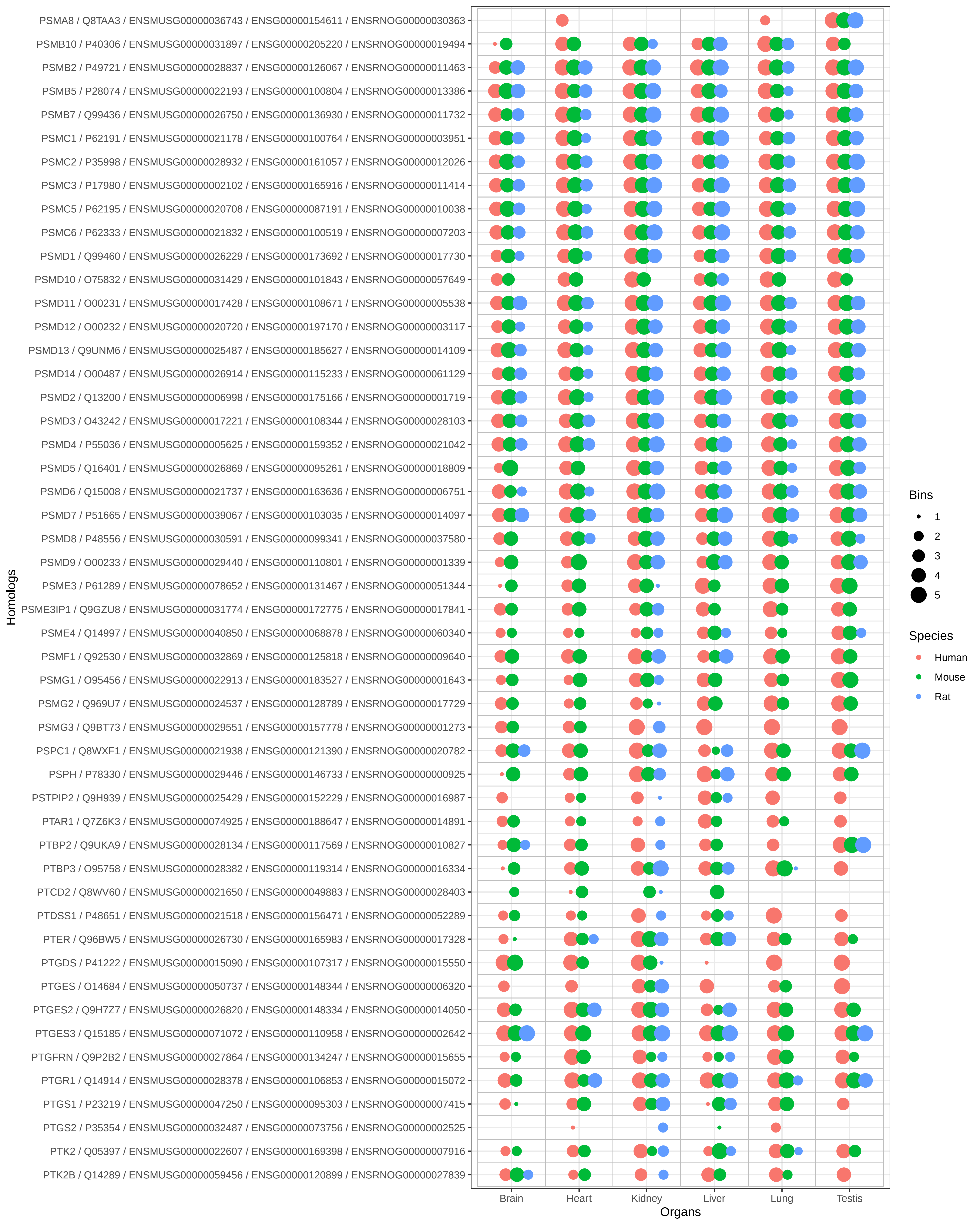
PSMA8 / Q8TAA3 / ENSMUSG00000036743 / ENSG00000154611 / ENSRNOG00000030363
PSMB10 / P40306 / ENSMUSG00000031897 / ENSG00000205220 / ENSRNOG00000019494
PSMB2 / P49721 / ENSMUSG00000028837 / ENSG00000126067 / ENSRNOG00000011463
PSMB5 / P28074 / ENSMUSG00000022193 / ENSG00000100804 / ENSRNOG00000013386
PSMB7 / Q99436 / ENSMUSG00000026750 / ENSG00000136930 / ENSRNOG00000011732
PSMC1 / P62191 / ENSMUSG00000021178 / ENSG00000100764 / ENSRNOG00000003951
PSMC2 / P35998 / ENSMUSG00000028932 / ENSG00000161057 / ENSRNOG00000012026
PSMC3 / P17980 / ENSMUSG00000002102 / ENSG00000165916 / ENSRNOG00000011414
PSMC5 / P62195 / ENSMUSG00000020708 / ENSG00000087191 / ENSRNOG00000010038
PSMC6 / P62333 / ENSMUSG00000021832 / ENSG00000100519 / ENSRNOG00000007203
PSMD1 / Q99460 / ENSMUSG00000026229 / ENSG00000173692 / ENSRNOG00000017730
PSMD10 / O75832 / ENSMUSG00000031429 / ENSG00000101843 / ENSRNOG00000057649
PSMD11 / O00231 / ENSMUSG00000017428 / ENSG00000108671 / ENSRNOG00000005538
PSMD12 / O00232 / ENSMUSG00000020720 / ENSG00000197170 / ENSRNOG00000003117
PSMD13 / Q9UNM6 / ENSMUSG00000025487 / ENSG00000185627 / ENSRNOG00000014109
PSMD14 / O00487 / ENSMUSG00000026914 / ENSG00000115233 / ENSRNOG00000061129
PSMD2 / Q13200 / ENSMUSG00000006998 / ENSG00000175166 / ENSRNOG00000001719
PSMD3 / O43242 / ENSMUSG00000017221 / ENSG00000108344 / ENSRNOG00000028103
PSMD4 / P55036 / ENSMUSG00000005625 / ENSG00000159352 / ENSRNOG00000021042
PSMD5 / Q16401 / ENSMUSG00000026869 / ENSG00000095261 / ENSRNOG00000018809
PSMD6 / Q15008 / ENSMUSG00000021737 / ENSG00000163636 / ENSRNOG00000006751
PSMD7 / P51665 / ENSMUSG00000039067 / ENSG00000103035 / ENSRNOG00000014097
PSMD8 / P48556 / ENSMUSG00000030591 / ENSG00000099341 / ENSRNOG00000037580
PSMD9 / O00233 / ENSMUSG00000029440 / ENSG00000110801 / ENSRNOG00000001339
PSME3 / P61289 / ENSMUSG00000078652 / ENSG00000131467 / ENSRNOG00000051344
PSME3IP1 / Q9GZU8 / ENSMUSG00000031774 / ENSG00000172775 / ENSRNOG00000017841
PSME4 / Q14997 / ENSMUSG00000040850 / ENSG00000068878 / ENSRNOG00000060340
PSMF1 / Q92530 / ENSMUSG00000032869 / ENSG00000125818 / ENSRNOG00000009640
PSMG1 / O95456 / ENSMUSG00000022913 / ENSG00000183527 / ENSRNOG00000001643
PSMG2 / Q969U7 / ENSMUSG00000024537 / ENSG00000128789 / ENSRNOG00000017729
PSMG3 / Q9BT73 / ENSMUSG00000029551 / ENSG00000157778 / ENSRNOG00000001273
PSPC1 / Q8WXF1 / ENSMUSG00000021938 / ENSG00000121390 / ENSRNOG00000020782
PSPH / P78330 / ENSMUSG00000029446 / ENSG00000146733 / ENSRNOG00000000925
PSTPIP2 / Q9H939 / ENSMUSG00000025429 / ENSG00000152229 / ENSRNOG00000016987
PTAR1 / Q7Z6K3 / ENSMUSG00000074925 / ENSG00000188647 / ENSRNOG00000014891
PTBP2 / Q9UKA9 / ENSMUSG00000028134 / ENSG00000117569 / ENSRNOG00000010827
PTBP3 / O95758 / ENSMUSG00000028382 / ENSG00000119314 / ENSRNOG00000016334
PTCD2 / Q8WV60 / ENSMUSG00000021650 / ENSG00000049883 / ENSRNOG00000028403
PTDSS1 / P48651 / ENSMUSG00000021518 / ENSG00000156471 / ENSRNOG00000052289
PTER / Q96BW5 / ENSMUSG00000026730 / ENSG00000165983 / ENSRNOG00000017328
PTGDS / P41222 / ENSMUSG00000015090 / ENSG00000107317 / ENSRNOG00000015550
PTGES / O14684 / ENSMUSG00000050737 / ENSG00000148344 / ENSRNOG00000006320
PTGES2 / Q9H7Z7 / ENSMUSG00000026820 / ENSG00000148334 / ENSRNOG00000014050
PTGES3 / Q15185 / ENSMUSG00000071072 / ENSG00000110958 / ENSRNOG00000002642
PTGFRN / Q9P2B2 / ENSMUSG00000027864 / ENSG00000134247 / ENSRNOG00000015655
PTGR1 / Q14914 / ENSMUSG00000028378 / ENSG00000106853 / ENSRNOG00000015072
PTGS1 / P23219 / ENSMUSG00000047250 / ENSG00000095303 / ENSRNOG00000007415
PTGS2 / P35354 / ENSMUSG00000032487 / ENSG00000073756 / ENSRNOG00000002525
PTK2 / Q05397 / ENSMUSG00000022607 / ENSG00000169398 / ENSRNOG00000007916
PTK2B / Q14289 / ENSMUSG00000059456 / ENSG00000120899 / ENSRNOG00000027839
Homologs
Brain
Heart
Kidney
Liver
Lung
Testis
Organs
Bins
1
2
3
4
5
Species
Human
Mouse
Rat

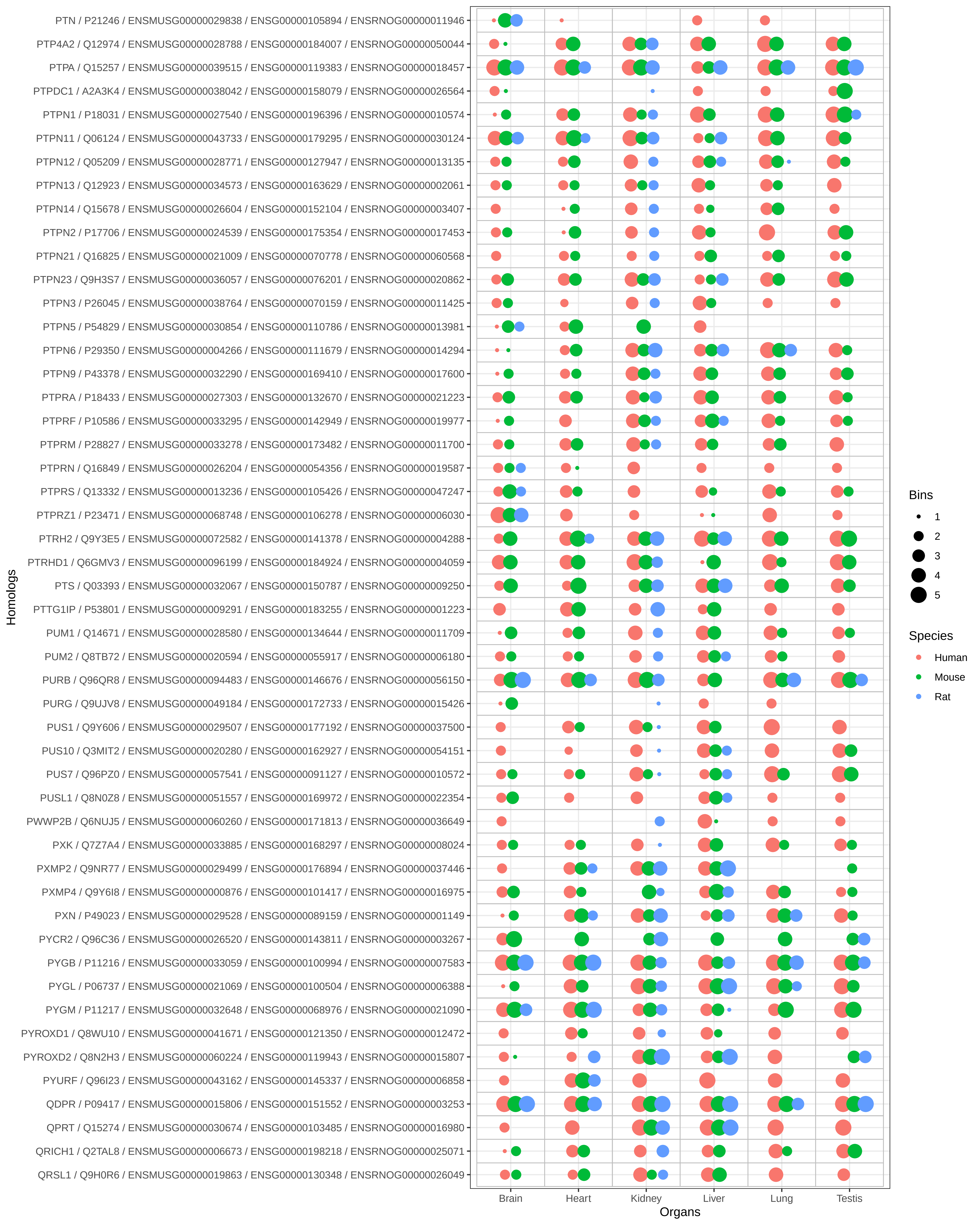
PTN / P21246 / ENSMUSG00000029838 / ENSG00000105894 / ENSRNOG00000011946
PTP4A2 / Q12974 / ENSMUSG00000028788 / ENSG00000184007 / ENSRNOG00000050044
PTPA / Q15257 / ENSMUSG00000039515 / ENSG00000119383 / ENSRNOG00000018457
PTPDC1 / A2A3K4 / ENSMUSG00000038042 / ENSG00000158079 / ENSRNOG00000026564
PTPN1 / P18031 / ENSMUSG00000027540 / ENSG00000196396 / ENSRNOG00000010574
PTPN11 / Q06124 / ENSMUSG00000043733 / ENSG00000179295 / ENSRNOG00000030124
PTPN12 / Q05209 / ENSMUSG00000028771 / ENSG00000127947 / ENSRNOG00000013135
PTPN13 / Q12923 / ENSMUSG00000034573 / ENSG00000163629 / ENSRNOG00000002061
PTPN14 / Q15678 / ENSMUSG00000026604 / ENSG00000152104 / ENSRNOG00000003407
PTPN2 / P17706 / ENSMUSG00000024539 / ENSG00000175354 / ENSRNOG00000017453
PTPN21 / Q16825 / ENSMUSG00000021009 / ENSG00000070778 / ENSRNOG00000060568
PTPN23 / Q9H3S7 / ENSMUSG00000036057 / ENSG00000076201 / ENSRNOG00000020862
PTPN3 / P26045 / ENSMUSG00000038764 / ENSG00000070159 / ENSRNOG00000011425
PTPN5 / P54829 / ENSMUSG00000030854 / ENSG00000110786 / ENSRNOG00000013981
PTPN6 / P29350 / ENSMUSG00000004266 / ENSG00000111679 / ENSRNOG00000014294
PTPN9 / P43378 / ENSMUSG00000032290 / ENSG00000169410 / ENSRNOG00000017600
PTPRA / P18433 / ENSMUSG00000027303 / ENSG00000132670 / ENSRNOG00000021223
PTPRF / P10586 / ENSMUSG00000033295 / ENSG00000142949 / ENSRNOG00000019977
PTPRM / P28827 / ENSMUSG00000033278 / ENSG00000173482 / ENSRNOG00000011700
PTPRN / Q16849 / ENSMUSG00000026204 / ENSG00000054356 / ENSRNOG00000019587
PTPRS / Q13332 / ENSMUSG00000013236 / ENSG00000105426 / ENSRNOG00000047247
PTPRZ1 / P23471 / ENSMUSG00000068748 / ENSG00000106278 / ENSRNOG00000006030
PTRH2 / Q9Y3E5 / ENSMUSG00000072582 / ENSG00000141378 / ENSRNOG00000004288
PTRHD1 / Q6GMV3 / ENSMUSG00000096199 / ENSG00000184924 / ENSRNOG00000004059
PTS / Q03393 / ENSMUSG00000032067 / ENSG00000150787 / ENSRNOG00000009250
PTTG1IP / P53801 / ENSMUSG00000009291 / ENSG00000183255 / ENSRNOG00000001223
PUM1 / Q14671 / ENSMUSG00000028580 / ENSG00000134644 / ENSRNOG00000011709
PUM2 / Q8TB72 / ENSMUSG00000020594 / ENSG00000055917 / ENSRNOG00000006180
PURB / Q96QR8 / ENSMUSG00000094483 / ENSG00000146676 / ENSRNOG00000056150
PURG / Q9UJV8 / ENSMUSG00000049184 / ENSG00000172733 / ENSRNOG00000015426
PUS1 / Q9Y606 / ENSMUSG00000029507 / ENSG00000177192 / ENSRNOG00000037500
PUS10 / Q3MIT2 / ENSMUSG00000020280 / ENSG00000162927 / ENSRNOG00000054151
PUS7 / Q96PZ0 / ENSMUSG00000057541 / ENSG00000091127 / ENSRNOG00000010572
PUSL1 / Q8N0Z8 / ENSMUSG00000051557 / ENSG00000169972 / ENSRNOG00000022354
PWWP2B / Q6NUJ5 / ENSMUSG00000060260 / ENSG00000171813 / ENSRNOG00000036649
PXK / Q7Z7A4 / ENSMUSG00000033885 / ENSG00000168297 / ENSRNOG00000008024
PXMP2 / Q9NR77 / ENSMUSG00000029499 / ENSG00000176894 / ENSRNOG00000037446
PXMP4 / Q9Y6I8 / ENSMUSG00000000876 / ENSG00000101417 / ENSRNOG00000016975
PXN / P49023 / ENSMUSG00000029528 / ENSG00000089159 / ENSRNOG00000001149
PYCR2 / Q96C36 / ENSMUSG00000026520 / ENSG00000143811 / ENSRNOG00000003267
PYGB / P11216 / ENSMUSG00000033059 / ENSG00000100994 / ENSRNOG00000007583
PYGL / P06737 / ENSMUSG00000021069 / ENSG00000100504 / ENSRNOG00000006388
PYGM / P11217 / ENSMUSG00000032648 / ENSG00000068976 / ENSRNOG00000021090
PYROXD1 / Q8WU10 / ENSMUSG00000041671 / ENSG00000121350 / ENSRNOG00000012472
PYROXD2 / Q8N2H3 / ENSMUSG00000060224 / ENSG00000119943 / ENSRNOG00000015807
PYURF / Q96I23 / ENSMUSG00000043162 / ENSG00000145337 / ENSRNOG00000006858
QDPR / P09417 / ENSMUSG00000015806 / ENSG00000151552 / ENSRNOG00000003253
QPRT / Q15274 / ENSMUSG00000030674 / ENSG00000103485 / ENSRNOG00000016980
QRICH1 / Q2TAL8 / ENSMUSG00000006673 / ENSG00000198218 / ENSRNOG00000025071
QRSL1 / Q9H0R6 / ENSMUSG00000019863 / ENSG00000130348 / ENSRNOG00000026049
Homologs
Brain
Heart
Kidney
Liver
Lung
Testis
Organs
Bins
1
2
3
4
5
Species
Human
Mouse
Rat

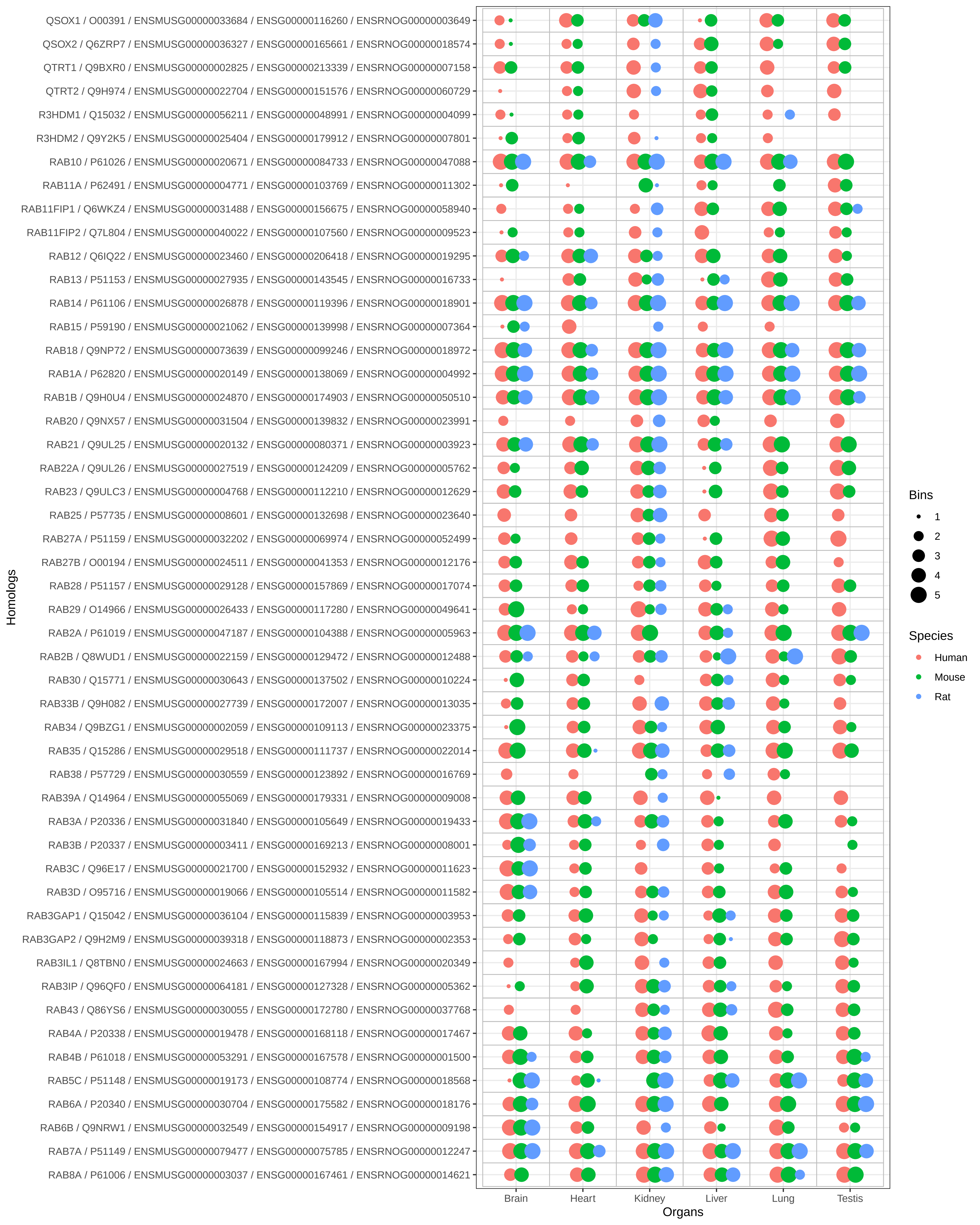

QSOX1 / O00391 / ENSMUSG00000033684 / ENSG00000116260 / ENSRNOG00000003649
QSOX2 / Q6ZRP7 / ENSMUSG00000036327 / ENSG00000165661 / ENSRNOG00000018574
QTRT1 / Q9BXR0 / ENSMUSG00000002825 / ENSG00000213339 / ENSRNOG00000007158
QTRT2 / Q9H974 / ENSMUSG00000022704 / ENSG00000151576 / ENSRNOG00000060729
R3HDM1 / Q15032 / ENSMUSG00000056211 / ENSG00000048991 / ENSRNOG00000004099
R3HDM2 / Q9Y2K5 / ENSMUSG00000025404 / ENSG00000179912 / ENSRNOG00000007801
RAB10 / P61026 / ENSMUSG00000020671 / ENSG00000084733 / ENSRNOG00000047088
RAB11A / P62491 / ENSMUSG00000004771 / ENSG00000103769 / ENSRNOG00000011302
RAB11FIP1 / Q6WKZ4 / ENSMUSG00000031488 / ENSG00000156675 / ENSRNOG00000058940
RAB11FIP2 / Q7L804 / ENSMUSG00000040022 / ENSG00000107560 / ENSRNOG00000009523
RAB12 / Q6IQ22 / ENSMUSG00000023460 / ENSG00000206418 / ENSRNOG00000019295
RAB13 / P51153 / ENSMUSG00000027935 / ENSG00000143545 / ENSRNOG00000016733
RAB14 / P61106 / ENSMUSG00000026878 / ENSG00000119396 / ENSRNOG00000018901
RAB15 / P59190 / ENSMUSG00000021062 / ENSG00000139998 / ENSRNOG00000007364
RAB18 / Q9NP72 / ENSMUSG00000073639 / ENSG00000099246 / ENSRNOG00000018972
RAB1A / P62820 / ENSMUSG00000020149 / ENSG00000138069 / ENSRNOG00000004992
RAB1B / Q9H0U4 / ENSMUSG00000024870 / ENSG00000174903 / ENSRNOG00000050510
RAB20 / Q9NX57 / ENSMUSG00000031504 / ENSG00000139832 / ENSRNOG00000023991
RAB21 / Q9UL25 / ENSMUSG00000020132 / ENSG00000080371 / ENSRNOG00000003923
RAB22A / Q9UL26 / ENSMUSG00000027519 / ENSG00000124209 / ENSRNOG00000005762
RAB23 / Q9ULC3 / ENSMUSG00000004768 / ENSG00000112210 / ENSRNOG00000012629
RAB25 / P57735 / ENSMUSG00000008601 / ENSG00000132698 / ENSRNOG00000023640
RAB27A / P51159 / ENSMUSG00000032202 / ENSG00000069974 / ENSRNOG00000052499
RAB27B / O00194 / ENSMUSG00000024511 / ENSG00000041353 / ENSRNOG00000012176
RAB28 / P51157 / ENSMUSG00000029128 / ENSG00000157869 / ENSRNOG00000017074
RAB29 / O14966 / ENSMUSG00000026433 / ENSG00000117280 / ENSRNOG00000049641
RAB2A / P61019 / ENSMUSG00000047187 / ENSG00000104388 / ENSRNOG00000005963
RAB2B / Q8WUD1 / ENSMUSG00000022159 / ENSG00000129472 / ENSRNOG00000012488
RAB30 / Q15771 / ENSMUSG00000030643 / ENSG00000137502 / ENSRNOG00000010224
RAB33B / Q9H082 / ENSMUSG00000027739 / ENSG00000172007 / ENSRNOG00000013035
RAB34 / Q9BZG1 / ENSMUSG00000002059 / ENSG00000109113 / ENSRNOG00000023375
RAB35 / Q15286 / ENSMUSG00000029518 / ENSG00000111737 / ENSRNOG00000022014
RAB38 / P57729 / ENSMUSG00000030559 / ENSG00000123892 / ENSRNOG00000016769
RAB39A / Q14964 / ENSMUSG00000055069 / ENSG00000179331 / ENSRNOG00000009008
RAB3A / P20336 / ENSMUSG00000031840 / ENSG00000105649 / ENSRNOG00000019433
RAB3B / P20337 / ENSMUSG00000003411 / ENSG00000169213 / ENSRNOG00000008001
RAB3C / Q96E17 / ENSMUSG00000021700 / ENSG00000152932 / ENSRNOG00000011623
RAB3D / O95716 / ENSMUSG00000019066 / ENSG00000105514 / ENSRNOG00000011582
RAB3GAP1 / Q15042 / ENSMUSG00000036104 / ENSG00000115839 / ENSRNOG00000003953
RAB3GAP2 / Q9H2M9 / ENSMUSG00000039318 / ENSG00000118873 / ENSRNOG00000002353
RAB3IL1 / Q8TBN0 / ENSMUSG00000024663 / ENSG00000167994 / ENSRNOG00000020349
RAB3IP / Q96QF0 / ENSMUSG00000064181 / ENSG00000127328 / ENSRNOG00000005362
RAB43 / Q86YS6 / ENSMUSG00000030055 / ENSG00000172780 / ENSRNOG00000037768
RAB4A / P20338 / ENSMUSG00000019478 / ENSG00000168118 / ENSRNOG00000017467
RAB4B / P61018 / ENSMUSG00000053291 / ENSG00000167578 / ENSRNOG00000001500
RAB5C / P51148 / ENSMUSG00000019173 / ENSG00000108774 / ENSRNOG00000018568
RAB6A / P20340 / ENSMUSG00000030704 / ENSG00000175582 / ENSRNOG00000018176
RAB6B / Q9NRW1 / ENSMUSG00000032549 / ENSG00000154917 / ENSRNOG00000009198
RAB7A / P51149 / ENSMUSG00000079477 / ENSG00000075785 / ENSRNOG00000012247
RAB8A / P61006 / ENSMUSG00000003037 / ENSG00000167461 / ENSRNOG00000014621
Homologs
Brain
Heart
Kidney
Liver
Lung
Testis
Organs
Bins
1
2
3
4
5
Species
Human
Mouse
Rat

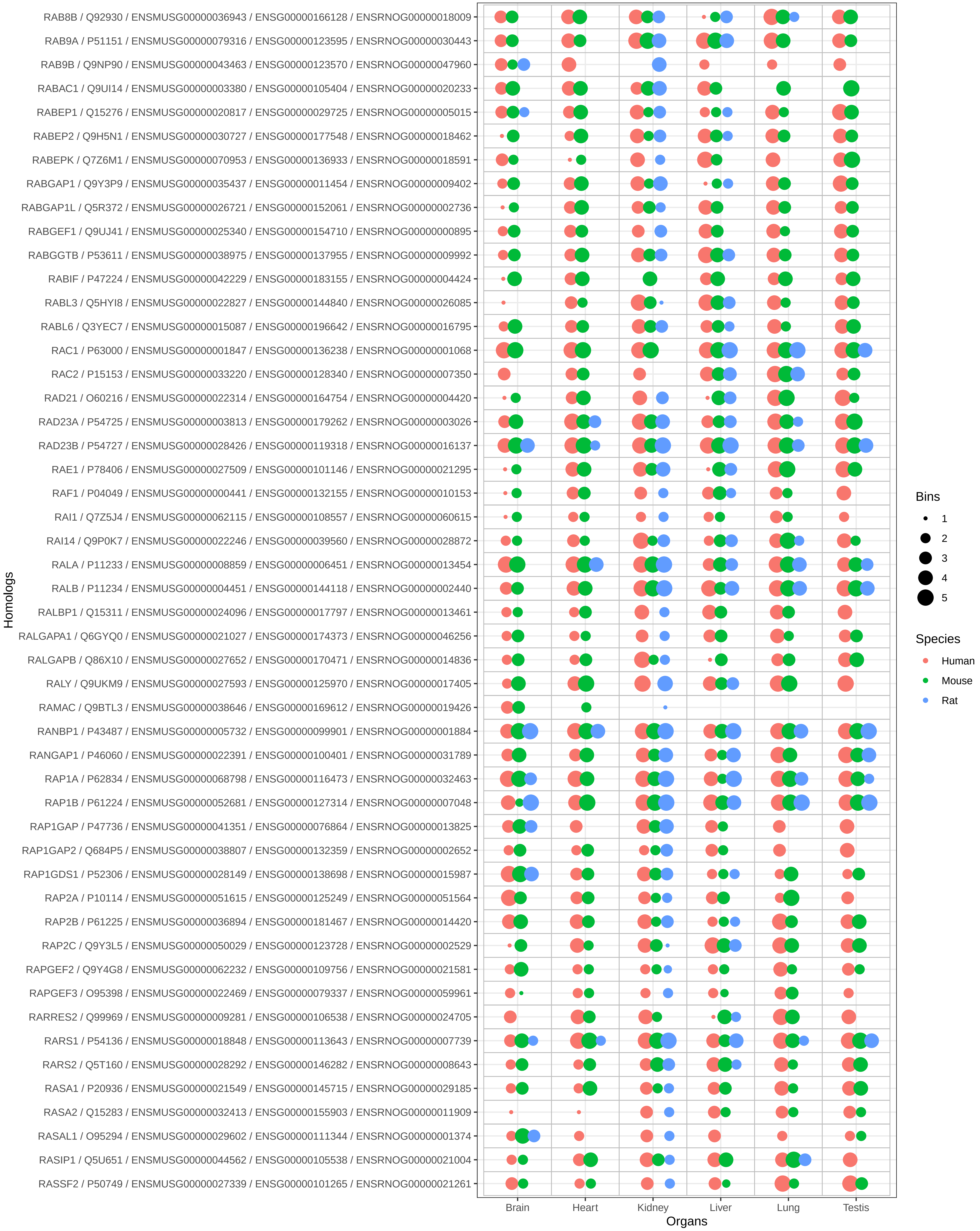
RAB8B / Q92930 / ENSMUSG00000036943 / ENSG00000166128 / ENSRNOG00000018009
RAB9A / P51151 / ENSMUSG00000079316 / ENSG00000123595 / ENSRNOG00000030443
RAB9B / Q9NP90 / ENSMUSG00000043463 / ENSG00000123570 / ENSRNOG00000047960
RABAC1 / Q9UI14 / ENSMUSG00000003380 / ENSG00000105404 / ENSRNOG00000020233
RABEP1 / Q15276 / ENSMUSG00000020817 / ENSG00000029725 / ENSRNOG00000005015
RABEP2 / Q9H5N1 / ENSMUSG00000030727 / ENSG00000177548 / ENSRNOG00000018462
RABEPK / Q7Z6M1 / ENSMUSG00000070953 / ENSG00000136933 / ENSRNOG00000018591
RABGAP1 / Q9Y3P9 / ENSMUSG00000035437 / ENSG00000011454 / ENSRNOG00000009402
RABGAP1L / Q5R372 / ENSMUSG00000026721 / ENSG00000152061 / ENSRNOG00000002736
RABGEF1 / Q9UJ41 / ENSMUSG00000025340 / ENSG00000154710 / ENSRNOG00000000895
RABGGTB / P53611 / ENSMUSG00000038975 / ENSG00000137955 / ENSRNOG00000009992
RABIF / P47224 / ENSMUSG00000042229 / ENSG00000183155 / ENSRNOG00000004424
RABL3 / Q5HYI8 / ENSMUSG00000022827 / ENSG00000144840 / ENSRNOG00000026085
RABL6 / Q3YEC7 / ENSMUSG00000015087 / ENSG00000196642 / ENSRNOG00000016795
RAC1 / P63000 / ENSMUSG00000001847 / ENSG00000136238 / ENSRNOG00000001068
RAC2 / P15153 / ENSMUSG00000033220 / ENSG00000128340 / ENSRNOG00000007350
RAD21 / O60216 / ENSMUSG00000022314 / ENSG00000164754 / ENSRNOG00000004420
RAD23A / P54725 / ENSMUSG00000003813 / ENSG00000179262 / ENSRNOG00000003026
RAD23B / P54727 / ENSMUSG00000028426 / ENSG00000119318 / ENSRNOG00000016137
RAE1 / P78406 / ENSMUSG00000027509 / ENSG00000101146 / ENSRNOG00000021295
RAF1 / P04049 / ENSMUSG00000000441 / ENSG00000132155 / ENSRNOG00000010153
RAI1 / Q7Z5J4 / ENSMUSG00000062115 / ENSG00000108557 / ENSRNOG00000060615
RAI14 / Q9P0K7 / ENSMUSG00000022246 / ENSG00000039560 / ENSRNOG00000028872
RALA / P11233 / ENSMUSG00000008859 / ENSG00000006451 / ENSRNOG00000013454
RALB / P11234 / ENSMUSG00000004451 / ENSG00000144118 / ENSRNOG00000002440
RALBP1 / Q15311 / ENSMUSG00000024096 / ENSG00000017797 / ENSRNOG00000013461
RALGAPA1 / Q6GYQ0 / ENSMUSG00000021027 / ENSG00000174373 / ENSRNOG00000046256
RALGAPB / Q86X10 / ENSMUSG00000027652 / ENSG00000170471 / ENSRNOG00000014836
RALY / Q9UKM9 / ENSMUSG00000027593 / ENSG00000125970 / ENSRNOG00000017405
RAMAC / Q9BTL3 / ENSMUSG00000038646 / ENSG00000169612 / ENSRNOG00000019426
RANBP1 / P43487 / ENSMUSG00000005732 / ENSG00000099901 / ENSRNOG00000001884
RANGAP1 / P46060 / ENSMUSG00000022391 / ENSG00000100401 / ENSRNOG00000031789
RAP1A / P62834 / ENSMUSG00000068798 / ENSG00000116473 / ENSRNOG00000032463
RAP1B / P61224 / ENSMUSG00000052681 / ENSG00000127314 / ENSRNOG00000007048
RAP1GAP / P47736 / ENSMUSG00000041351 / ENSG00000076864 / ENSRNOG00000013825
RAP1GAP2 / Q684P5 / ENSMUSG00000038807 / ENSG00000132359 / ENSRNOG00000002652
RAP1GDS1 / P52306 / ENSMUSG00000028149 / ENSG00000138698 / ENSRNOG00000015987
RAP2A / P10114 / ENSMUSG00000051615 / ENSG00000125249 / ENSRNOG00000051564
RAP2B / P61225 / ENSMUSG00000036894 / ENSG00000181467 / ENSRNOG00000014420
RAP2C / Q9Y3L5 / ENSMUSG00000050029 / ENSG00000123728 / ENSRNOG00000002529
RAPGEF2 / Q9Y4G8 / ENSMUSG00000062232 / ENSG00000109756 / ENSRNOG00000021581
RAPGEF3 / O95398 / ENSMUSG00000022469 / ENSG00000079337 / ENSRNOG00000059961
RARRES2 / Q99969 / ENSMUSG00000009281 / ENSG00000106538 / ENSRNOG00000024705
RARS1 / P54136 / ENSMUSG00000018848 / ENSG00000113643 / ENSRNOG00000007739
RARS2 / Q5T160 / ENSMUSG00000028292 / ENSG00000146282 / ENSRNOG00000008643
RASA1 / P20936 / ENSMUSG00000021549 / ENSG00000145715 / ENSRNOG00000029185
RASA2 / Q15283 / ENSMUSG00000032413 / ENSG00000155903 / ENSRNOG00000011909
RASAL1 / O95294 / ENSMUSG00000029602 / ENSG00000111344 / ENSRNOG00000001374
RASIP1 / Q5U651 / ENSMUSG00000044562 / ENSG00000105538 / ENSRNOG00000021004
RASSF2 / P50749 / ENSMUSG00000027339 / ENSG00000101265 / ENSRNOG00000021261
Homologs
Brain
Heart
Kidney
Liver
Lung
Testis
Organs
Bins
1
2
3
4
5
Species
Human
Mouse
Rat

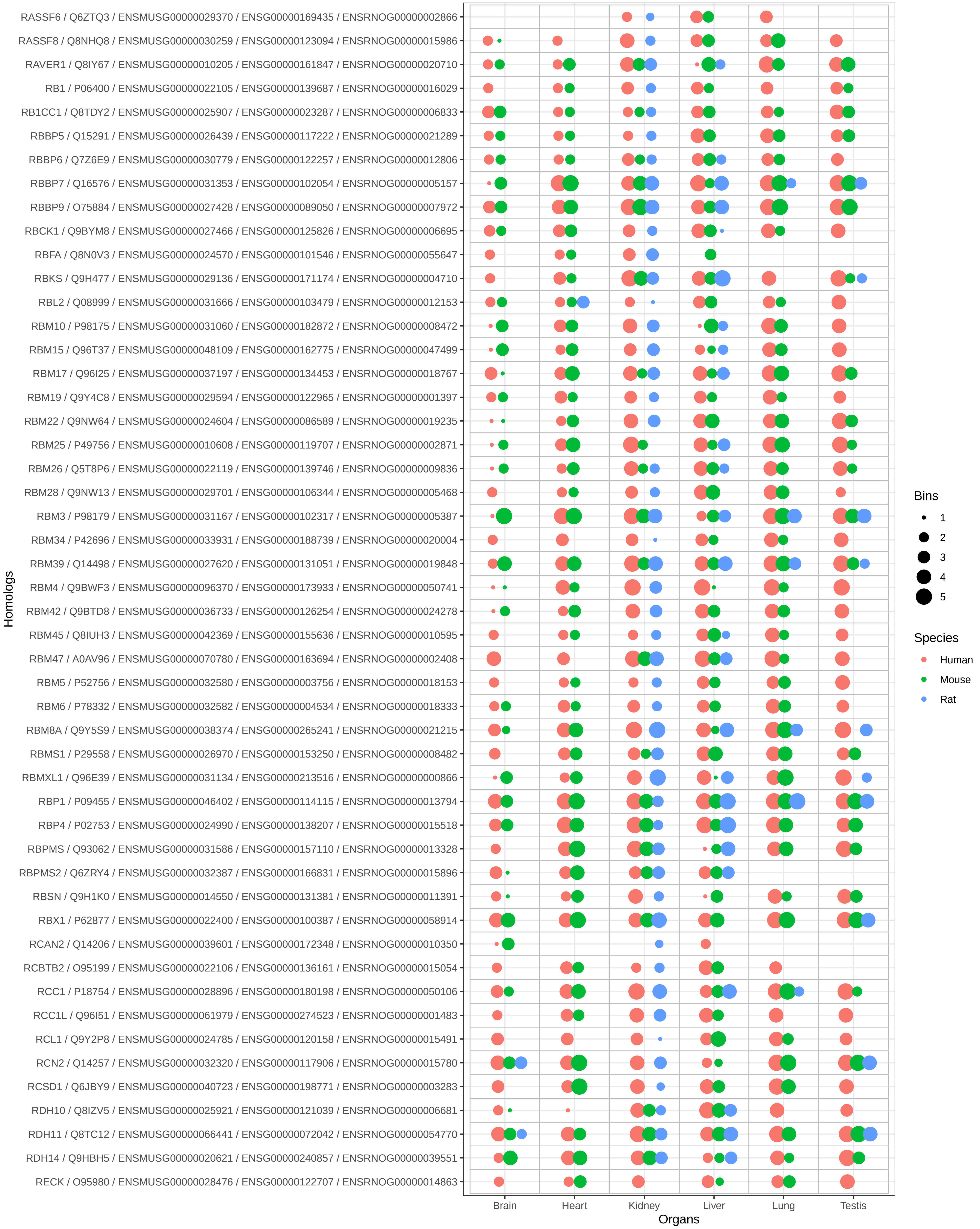

RASSF6 / Q6ZTQ3 / ENSMUSG00000029370 / ENSG00000169435 / ENSRNOG00000002866
RASSF8 / Q8NHQ8 / ENSMUSG00000030259 / ENSG00000123094 / ENSRNOG00000015986
RAVER1 / Q8IY67 / ENSMUSG00000010205 / ENSG00000161847 / ENSRNOG00000020710
RB1 / P06400 / ENSMUSG00000022105 / ENSG00000139687 / ENSRNOG00000016029
RB1CC1 / Q8TDY2 / ENSMUSG00000025907 / ENSG00000023287 / ENSRNOG00000006833
RBBP5 / Q15291 / ENSMUSG00000026439 / ENSG00000117222 / ENSRNOG00000021289
RBBP6 / Q7Z6E9 / ENSMUSG00000030779 / ENSG00000122257 / ENSRNOG00000012806
RBBP7 / Q16576 / ENSMUSG00000031353 / ENSG00000102054 / ENSRNOG00000005157
RBBP9 / O75884 / ENSMUSG00000027428 / ENSG00000089050 / ENSRNOG00000007972
RBCK1 / Q9BYM8 / ENSMUSG00000027466 / ENSG00000125826 / ENSRNOG00000006695
RBFA / Q8N0V3 / ENSMUSG00000024570 / ENSG00000101546 / ENSRNOG00000055647
RBKS / Q9H477 / ENSMUSG00000029136 / ENSG00000171174 / ENSRNOG00000004710
RBL2 / Q08999 / ENSMUSG00000031666 / ENSG00000103479 / ENSRNOG00000012153
RBM10 / P98175 / ENSMUSG00000031060 / ENSG00000182872 / ENSRNOG00000008472
RBM15 / Q96T37 / ENSMUSG00000048109 / ENSG00000162775 / ENSRNOG00000047499
RBM17 / Q96I25 / ENSMUSG00000037197 / ENSG00000134453 / ENSRNOG00000018767
RBM19 / Q9Y4C8 / ENSMUSG00000029594 / ENSG00000122965 / ENSRNOG00000001397
RBM22 / Q9NW64 / ENSMUSG00000024604 / ENSG00000086589 / ENSRNOG00000019235
RBM25 / P49756 / ENSMUSG00000010608 / ENSG00000119707 / ENSRNOG00000002871
RBM26 / Q5T8P6 / ENSMUSG00000022119 / ENSG00000139746 / ENSRNOG00000009836
RBM28 / Q9NW13 / ENSMUSG00000029701 / ENSG00000106344 / ENSRNOG00000005468
RBM3 / P98179 / ENSMUSG00000031167 / ENSG00000102317 / ENSRNOG00000005387
RBM34 / P42696 / ENSMUSG00000033931 / ENSG00000188739 / ENSRNOG00000020004
RBM39 / Q14498 / ENSMUSG00000027620 / ENSG00000131051 / ENSRNOG00000019848
RBM4 / Q9BWF3 / ENSMUSG00000096370 / ENSG00000173933 / ENSRNOG00000050741
RBM42 / Q9BTD8 / ENSMUSG00000036733 / ENSG00000126254 / ENSRNOG00000024278
RBM45 / Q8IUH3 / ENSMUSG00000042369 / ENSG00000155636 / ENSRNOG00000010595
RBM47 / A0AV96 / ENSMUSG00000070780 / ENSG00000163694 / ENSRNOG00000002408
RBM5 / P52756 / ENSMUSG00000032580 / ENSG00000003756 / ENSRNOG00000018153
RBM6 / P78332 / ENSMUSG00000032582 / ENSG00000004534 / ENSRNOG00000018333
RBM8A / Q9Y5S9 / ENSMUSG00000038374 / ENSG00000265241 / ENSRNOG00000021215
RBMS1 / P29558 / ENSMUSG00000026970 / ENSG00000153250 / ENSRNOG00000008482
RBMXL1 / Q96E39 / ENSMUSG00000031134 / ENSG00000213516 / ENSRNOG00000000866
RBP1 / P09455 / ENSMUSG00000046402 / ENSG00000114115 / ENSRNOG00000013794
RBP4 / P02753 / ENSMUSG00000024990 / ENSG00000138207 / ENSRNOG00000015518
RBPMS / Q93062 / ENSMUSG00000031586 / ENSG00000157110 / ENSRNOG00000013328
RBPMS2 / Q6ZRY4 / ENSMUSG00000032387 / ENSG00000166831 / ENSRNOG00000015896
RBSN / Q9H1K0 / ENSMUSG00000014550 / ENSG00000131381 / ENSRNOG00000011391
RBX1 / P62877 / ENSMUSG00000022400 / ENSG00000100387 / ENSRNOG00000058914
RCAN2 / Q14206 / ENSMUSG00000039601 / ENSG00000172348 / ENSRNOG00000010350
RCBTB2 / O95199 / ENSMUSG00000022106 / ENSG00000136161 / ENSRNOG00000015054
RCC1 / P18754 / ENSMUSG00000028896 / ENSG00000180198 / ENSRNOG00000050106
RCC1L / Q96I51 / ENSMUSG00000061979 / ENSG00000274523 / ENSRNOG00000001483
RCL1 / Q9Y2P8 / ENSMUSG00000024785 / ENSG00000120158 / ENSRNOG00000015491
RCN2 / Q14257 / ENSMUSG00000032320 / ENSG00000117906 / ENSRNOG00000015780
RCSD1 / Q6JBY9 / ENSMUSG00000040723 / ENSG00000198771 / ENSRNOG00000003283
RDH10 / Q8IZV5 / ENSMUSG00000025921 / ENSG00000121039 / ENSRNOG00000006681
RDH11 / Q8TC12 / ENSMUSG00000066441 / ENSG00000072042 / ENSRNOG00000054770
RDH14 / Q9HBH5 / ENSMUSG00000020621 / ENSG00000240857 / ENSRNOG00000039551
RECK / O95980 / ENSMUSG00000028476 / ENSG00000122707 / ENSRNOG00000014863
Homologs
Brain
Heart
Kidney
Liver
Lung
Testis
Organs
Bins
1
2
3
4
5
Species
Human
Mouse
Rat

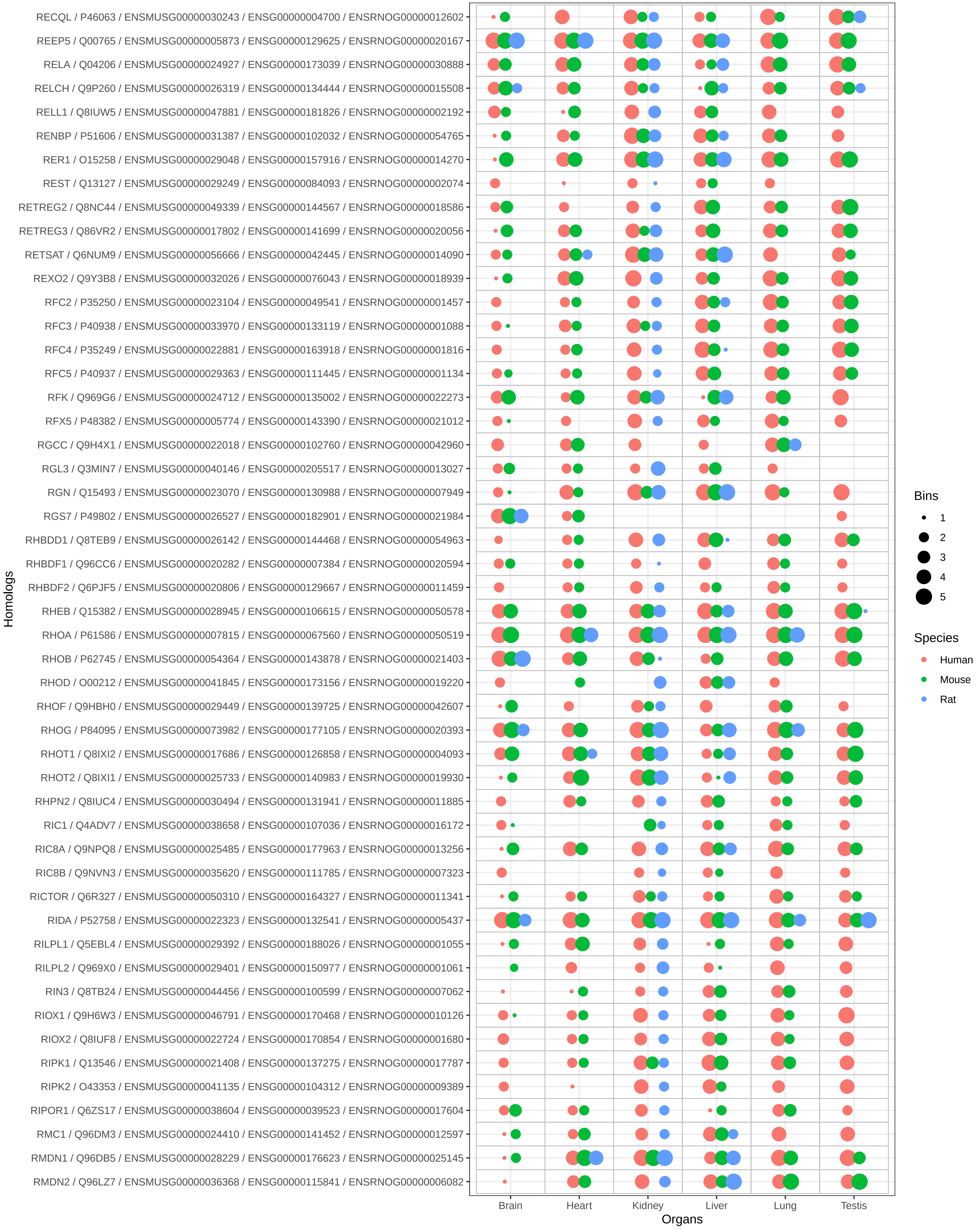
RECQL / P46063 / ENSMUSG00000030243 / ENSG00000004700 / ENSRNOG00000012602
REEP5 / Q00765 / ENSMUSG00000005873 / ENSG00000129625 / ENSRNOG00000020167
RELA / Q04206 / ENSMUSG00000024927 / ENSG00000173039 / ENSRNOG00000030888
RELCH / Q9P260 / ENSMUSG00000026319 / ENSG00000134444 / ENSRNOG00000015508
RELL1 / Q8IUW5 / ENSMUSG00000047881 / ENSG00000181826 / ENSRNOG00000002192
RENBP / P51606 / ENSMUSG00000031387 / ENSG00000102032 / ENSRNOG00000054765
RER1 / O15258 / ENSMUSG00000029048 / ENSG00000157916 / ENSRNOG00000014270
REST / Q13127 / ENSMUSG00000029249 / ENSG00000084093 / ENSRNOG00000002074
RETREG2 / Q8NC44 / ENSMUSG00000049339 / ENSG00000144567 / ENSRNOG00000018586
RETREG3 / Q86VR2 / ENSMUSG00000017802 / ENSG00000141699 / ENSRNOG00000020056
RETSAT / Q6NUM9 / ENSMUSG00000056666 / ENSG00000042445 / ENSRNOG00000014090
REXO2 / Q9Y3B8 / ENSMUSG00000032026 / ENSG00000076043 / ENSRNOG00000018939
RFC2 / P35250 / ENSMUSG00000023104 / ENSG00000049541 / ENSRNOG00000001457
RFC3 / P40938 / ENSMUSG00000033970 / ENSG00000133119 / ENSRNOG00000001088
RFC4 / P35249 / ENSMUSG00000022881 / ENSG00000163918 / ENSRNOG00000001816
RFC5 / P40937 / ENSMUSG00000029363 / ENSG00000111445 / ENSRNOG00000001134
RFK / Q969G6 / ENSMUSG00000024712 / ENSG00000135002 / ENSRNOG00000022273
RFX5 / P48382 / ENSMUSG00000005774 / ENSG00000143390 / ENSRNOG00000021012
RGCC / Q9H4X1 / ENSMUSG00000022018 / ENSG00000102760 / ENSRNOG00000042960
RGL3 / Q3MIN7 / ENSMUSG00000040146 / ENSG00000205517 / ENSRNOG00000013027
RGN / Q15493 / ENSMUSG00000023070 / ENSG00000130988 / ENSRNOG00000007949
RGS7 / P49802 / ENSMUSG00000026527 / ENSG00000182901 / ENSRNOG00000021984
RHBDD1 / Q8TEB9 / ENSMUSG00000026142 / ENSG00000144468 / ENSRNOG00000054963
RHBDF1 / Q96CC6 / ENSMUSG00000020282 / ENSG00000007384 / ENSRNOG00000020594
RHBDF2 / Q6PJF5 / ENSMUSG00000020806 / ENSG00000129667 / ENSRNOG00000011459
RHEB / Q15382 / ENSMUSG00000028945 / ENSG00000106615 / ENSRNOG00000050578
RHOA / P61586 / ENSMUSG00000007815 / ENSG00000067560 / ENSRNOG00000050519
RHOB / P62745 / ENSMUSG00000054364 / ENSG00000143878 / ENSRNOG00000021403
RHOD / O00212 / ENSMUSG00000041845 / ENSG00000173156 / ENSRNOG00000019220
RHOF / Q9HBH0 / ENSMUSG00000029449 / ENSG00000139725 / ENSRNOG00000042607
RHOG / P84095 / ENSMUSG00000073982 / ENSG00000177105 / ENSRNOG00000020393
RHOT1 / Q8IXI2 / ENSMUSG00000017686 / ENSG00000126858 / ENSRNOG00000004093
RHOT2 / Q8IXI1 / ENSMUSG00000025733 / ENSG00000140983 / ENSRNOG00000019930
RHPN2 / Q8IUC4 / ENSMUSG00000030494 / ENSG00000131941 / ENSRNOG00000011885
RIC1 / Q4ADV7 / ENSMUSG00000038658 / ENSG00000107036 / ENSRNOG00000016172
RIC8A / Q9NPQ8 / ENSMUSG00000025485 / ENSG00000177963 / ENSRNOG00000013256
RIC8B / Q9NVN3 / ENSMUSG00000035620 / ENSG00000111785 / ENSRNOG00000007323
RICTOR / Q6R327 / ENSMUSG00000050310 / ENSG00000164327 / ENSRNOG00000011341
RIDA / P52758 / ENSMUSG00000022323 / ENSG00000132541 / ENSRNOG00000005437
RILPL1 / Q5EBL4 / ENSMUSG00000029392 / ENSG00000188026 / ENSRNOG00000001055
RILPL2 / Q969X0 / ENSMUSG00000029401 / ENSG00000150977 / ENSRNOG00000001061
RIN3 / Q8TB24 / ENSMUSG00000044456 / ENSG00000100599 / ENSRNOG00000007062
RIOX1 / Q9H6W3 / ENSMUSG00000046791 / ENSG00000170468 / ENSRNOG00000010126
RIOX2 / Q8IUF8 / ENSMUSG00000022724 / ENSG00000170854 / ENSRNOG00000001680
RIPK1 / Q13546 / ENSMUSG00000021408 / ENSG00000137275 / ENSRNOG00000017787
RIPK2 / O43353 / ENSMUSG00000041135 / ENSG00000104312 / ENSRNOG00000009389
RIPOR1 / Q6ZS17 / ENSMUSG00000038604 / ENSG00000039523 / ENSRNOG00000017604
RMC1 / Q96DM3 / ENSMUSG00000024410 / ENSG00000141452 / ENSRNOG00000012597
RMDN1 / Q96DB5 / ENSMUSG00000028229 / ENSG00000176623 / ENSRNOG00000025145
RMDN2 / Q96LZ7 / ENSMUSG00000036368 / ENSG00000115841 / ENSRNOG00000006082
Homologs
Brain
Heart
Kidney
Liver
Lung
Testis
Organs
Bins
1
2
3
4
5
Species
Human
Mouse
Rat

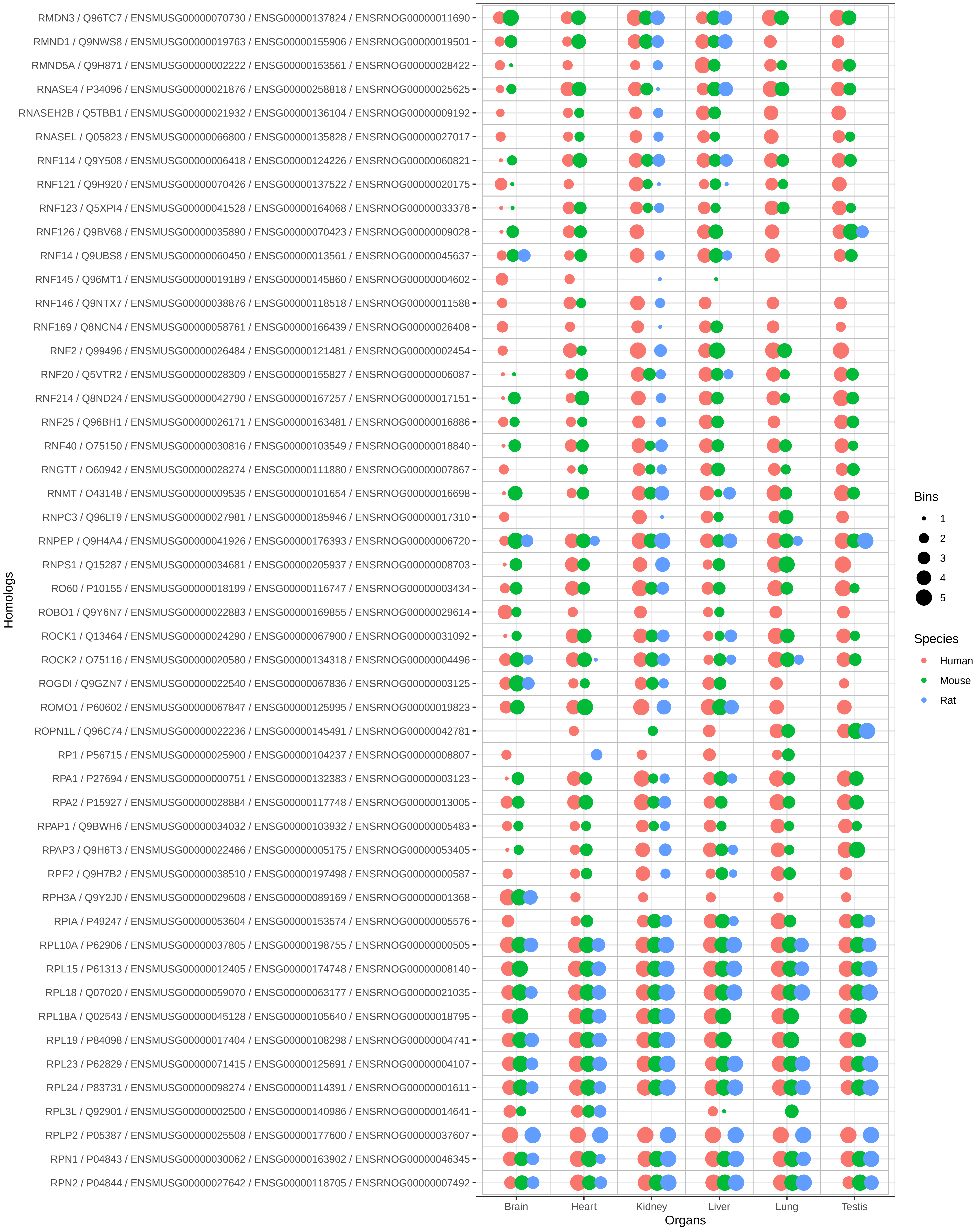
RMDN3 / Q96TC7 / ENSMUSG00000070730 / ENSG00000137824 / ENSRNOG00000011690
RMND1 / Q9NWS8 / ENSMUSG00000019763 / ENSG00000155906 / ENSRNOG00000019501
RMND5A / Q9H871 / ENSMUSG00000002222 / ENSG00000153561 / ENSRNOG00000028422
RNASE4 / P34096 / ENSMUSG00000021876 / ENSG00000258818 / ENSRNOG00000025625
RNASEH2B / Q5TBB1 / ENSMUSG00000021932 / ENSG00000136104 / ENSRNOG00000009192
RNASEL / Q05823 / ENSMUSG00000066800 / ENSG00000135828 / ENSRNOG00000027017
RNF114 / Q9Y508 / ENSMUSG00000006418 / ENSG00000124226 / ENSRNOG00000060821
RNF121 / Q9H920 / ENSMUSG00000070426 / ENSG00000137522 / ENSRNOG00000020175
RNF123 / Q5XPI4 / ENSMUSG00000041528 / ENSG00000164068 / ENSRNOG00000033378
RNF126 / Q9BV68 / ENSMUSG00000035890 / ENSG00000070423 / ENSRNOG00000009028
RNF14 / Q9UBS8 / ENSMUSG00000060450 / ENSG00000013561 / ENSRNOG00000045637
RNF145 / Q96MT1 / ENSMUSG00000019189 / ENSG00000145860 / ENSRNOG00000004602
RNF146 / Q9NTX7 / ENSMUSG00000038876 / ENSG00000118518 / ENSRNOG00000011588
RNF169 / Q8NCN4 / ENSMUSG00000058761 / ENSG00000166439 / ENSRNOG00000026408
RNF2 / Q99496 / ENSMUSG00000026484 / ENSG00000121481 / ENSRNOG00000002454
RNF20 / Q5VTR2 / ENSMUSG00000028309 / ENSG00000155827 / ENSRNOG00000006087
RNF214 / Q8ND24 / ENSMUSG00000042790 / ENSG00000167257 / ENSRNOG00000017151
RNF25 / Q96BH1 / ENSMUSG00000026171 / ENSG00000163481 / ENSRNOG00000016886
RNF40 / O75150 / ENSMUSG00000030816 / ENSG00000103549 / ENSRNOG00000018840
RNGTT / O60942 / ENSMUSG00000028274 / ENSG00000111880 / ENSRNOG00000007867
RNMT / O43148 / ENSMUSG00000009535 / ENSG00000101654 / ENSRNOG00000016698
RNPC3 / Q96LT9 / ENSMUSG00000027981 / ENSG00000185946 / ENSRNOG00000017310
RNPEP / Q9H4A4 / ENSMUSG00000041926 / ENSG00000176393 / ENSRNOG00000006720
RNPS1 / Q15287 / ENSMUSG00000034681 / ENSG00000205937 / ENSRNOG00000008703
RO60 / P10155 / ENSMUSG00000018199 / ENSG00000116747 / ENSRNOG00000003434
ROBO1 / Q9Y6N7 / ENSMUSG00000022883 / ENSG00000169855 / ENSRNOG00000029614
ROCK1 / Q13464 / ENSMUSG00000024290 / ENSG00000067900 / ENSRNOG00000031092
ROCK2 / O75116 / ENSMUSG00000020580 / ENSG00000134318 / ENSRNOG00000004496
ROGDI / Q9GZN7 / ENSMUSG00000022540 / ENSG00000067836 / ENSRNOG00000003125
ROMO1 / P60602 / ENSMUSG00000067847 / ENSG00000125995 / ENSRNOG00000019823
ROPN1L / Q96C74 / ENSMUSG00000022236 / ENSG00000145491 / ENSRNOG00000042781
RP1 / P56715 / ENSMUSG00000025900 / ENSG00000104237 / ENSRNOG00000008807
RPA1 / P27694 / ENSMUSG00000000751 / ENSG00000132383 / ENSRNOG00000003123
RPA2 / P15927 / ENSMUSG00000028884 / ENSG00000117748 / ENSRNOG00000013005
RPAP1 / Q9BWH6 / ENSMUSG00000034032 / ENSG00000103932 / ENSRNOG00000005483
RPAP3 / Q9H6T3 / ENSMUSG00000022466 / ENSG00000005175 / ENSRNOG00000053405
RPF2 / Q9H7B2 / ENSMUSG00000038510 / ENSG00000197498 / ENSRNOG00000000587
RPH3A / Q9Y2J0 / ENSMUSG00000029608 / ENSG00000089169 / ENSRNOG00000001368
RPIA / P49247 / ENSMUSG00000053604 / ENSG00000153574 / ENSRNOG00000005576
RPL10A / P62906 / ENSMUSG00000037805 / ENSG00000198755 / ENSRNOG00000000505
RPL15 / P61313 / ENSMUSG00000012405 / ENSG00000174748 / ENSRNOG00000008140
RPL18 / Q07020 / ENSMUSG00000059070 / ENSG00000063177 / ENSRNOG00000021035
RPL18A / Q02543 / ENSMUSG00000045128 / ENSG00000105640 / ENSRNOG00000018795
RPL19 / P84098 / ENSMUSG00000017404 / ENSG00000108298 / ENSRNOG00000004741
RPL23 / P62829 / ENSMUSG00000071415 / ENSG00000125691 / ENSRNOG00000004107
RPL24 / P83731 / ENSMUSG00000098274 / ENSG00000114391 / ENSRNOG00000001611
RPL3L / Q92901 / ENSMUSG00000002500 / ENSG00000140986 / ENSRNOG00000014641
RPLP2 / P05387 / ENSMUSG00000025508 / ENSG00000177600 / ENSRNOG00000037607
RPN1 / P04843 / ENSMUSG00000030062 / ENSG00000163902 / ENSRNOG00000046345
RPN2 / P04844 / ENSMUSG00000027642 / ENSG00000118705 / ENSRNOG00000007492
Homologs
Brain
Heart
Kidney
Liver
Lung
Testis
Organs
Bins
1
2
3
4
5
Species
Human
Mouse
Rat

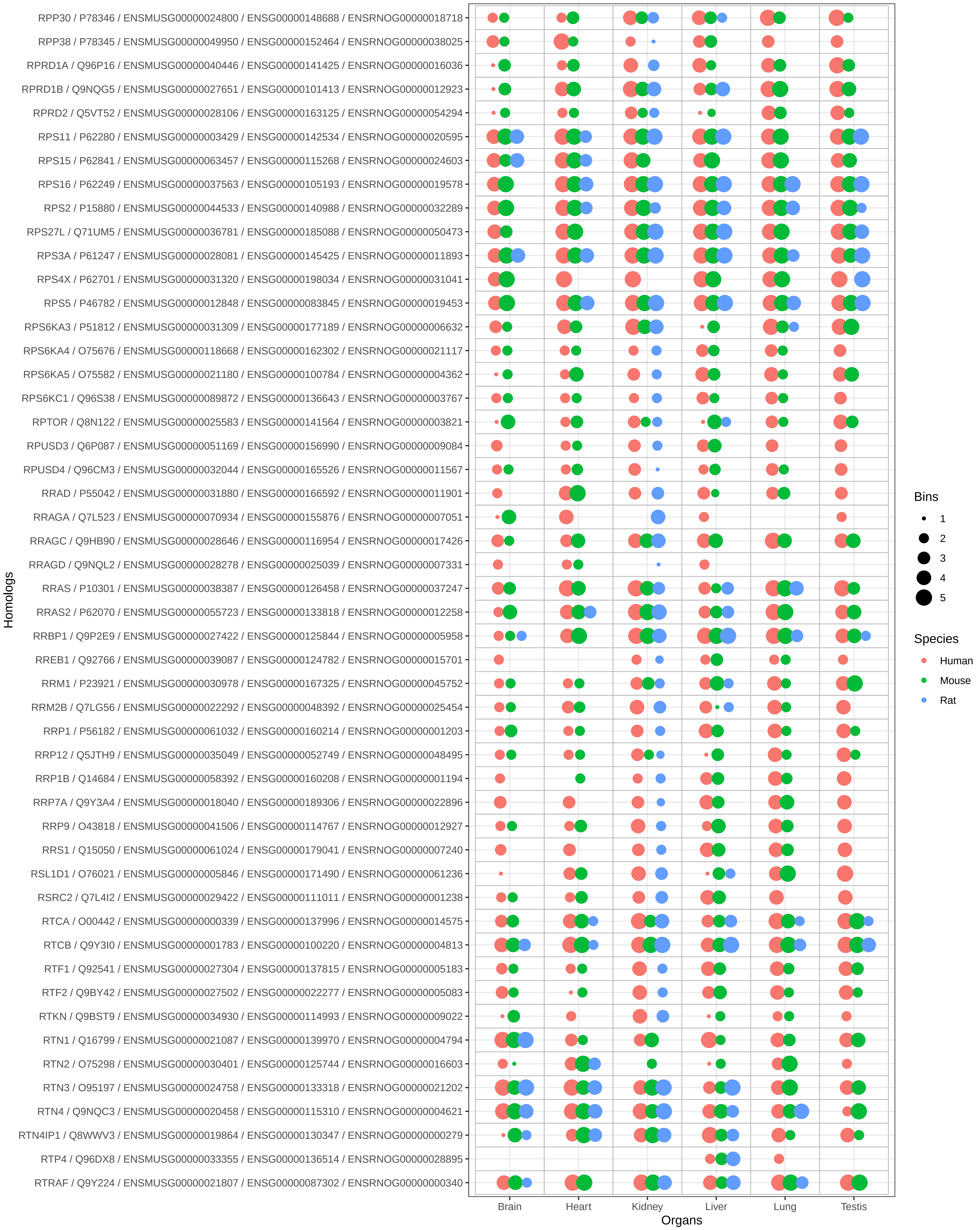
RPP30 / P78346 / ENSMUSG00000024800 / ENSG00000148688 / ENSRNOG00000018718
RPP38 / P78345 / ENSMUSG00000049950 / ENSG00000152464 / ENSRNOG00000038025
RPRD1A / Q96P16 / ENSMUSG00000040446 / ENSG00000141425 / ENSRNOG00000016036
RPRD1B / Q9NQG5 / ENSMUSG00000027651 / ENSG00000101413 / ENSRNOG00000012923
RPRD2 / Q5VT52 / ENSMUSG00000028106 / ENSG00000163125 / ENSRNOG00000054294
RPS11 / P62280 / ENSMUSG00000003429 / ENSG00000142534 / ENSRNOG00000020595
RPS15 / P62841 / ENSMUSG00000063457 / ENSG00000115268 / ENSRNOG00000024603
RPS16 / P62249 / ENSMUSG00000037563 / ENSG00000105193 / ENSRNOG00000019578
RPS2 / P15880 / ENSMUSG00000044533 / ENSG00000140988 / ENSRNOG00000032289
RPS27L / Q71UM5 / ENSMUSG00000036781 / ENSG00000185088 / ENSRNOG00000050473
RPS3A / P61247 / ENSMUSG00000028081 / ENSG00000145425 / ENSRNOG00000011893
RPS4X / P62701 / ENSMUSG00000031320 / ENSG00000198034 / ENSRNOG00000031041
RPS5 / P46782 / ENSMUSG00000012848 / ENSG00000083845 / ENSRNOG00000019453
RPS6KA3 / P51812 / ENSMUSG00000031309 / ENSG00000177189 / ENSRNOG00000006632
RPS6KA4 / O75676 / ENSMUSG00000024952 / ENSG00000162302 / ENSRNOG00000021117
RPS6KA5 / O75582 / ENSMUSG00000021180 / ENSG00000100784 / ENSRNOG00000004362
RPS6KC1 / Q96S38 / ENSMUSG00000089872 / ENSG00000136643 / ENSRNOG00000003767
RPTOR / Q8N122 / ENSMUSG00000025583 / ENSG00000141564 / ENSRNOG00000003821
RPUSD3 / Q6P087 / ENSMUSG00000051169 / ENSG00000156990 / ENSRNOG00000009084
RPUSD4 / Q96CM3 / ENSMUSG00000032044 / ENSG00000165526 / ENSRNOG00000011567
RRAD / P55042 / ENSMUSG00000031880 / ENSG00000166592 / ENSRNOG00000011901
RRAGA / Q7L523 / ENSMUSG00000070934 / ENSG00000155876 / ENSRNOG00000007051
RRAGC / Q9HB90 / ENSMUSG00000028646 / ENSG00000116954 / ENSRNOG00000017426
RRAGD / Q9NQL2 / ENSMUSG00000028278 / ENSG00000025039 / ENSRNOG00000007331
RRAS / P10301 / ENSMUSG00000038387 / ENSG00000126458 / ENSRNOG00000037247
RRAS2 / P62070 / ENSMUSG00000055723 / ENSG00000133818 / ENSRNOG00000012258
RRBP1 / Q9P2E9 / ENSMUSG00000027422 / ENSG00000125844 / ENSRNOG00000005958
RREB1 / Q92766 / ENSMUSG00000039087 / ENSG00000124782 / ENSRNOG00000015701
RRM1 / P23921 / ENSMUSG00000030978 / ENSG00000167325 / ENSRNOG00000045752
RRM2B / Q7LG56 / ENSMUSG00000022292 / ENSG00000048392 / ENSRNOG00000025454
RRP1 / P56182 / ENSMUSG00000061032 / ENSG00000160214 / ENSRNOG00000001203
RRP12 / Q5JTH9 / ENSMUSG00000035049 / ENSG00000052749 / ENSRNOG00000048495
RRP1B / Q14684 / ENSMUSG00000058392 / ENSG00000160208 / ENSRNOG00000001194
RRP7A / Q9Y3A4 / ENSMUSG00000018040 / ENSG00000189306 / ENSRNOG00000022896
RRP9 / O43818 / ENSMUSG00000041506 / ENSG00000114767 / ENSRNOG00000012927
RRS1 / Q15050 / ENSMUSG00000061024 / ENSG00000179041 / ENSRNOG00000007240
RSL1D1 / O76021 / ENSMUSG00000005846 / ENSG00000171490 / ENSRNOG00000061236
RSRC2 / Q7L4I2 / ENSMUSG00000029422 / ENSG00000111011 / ENSRNOG00000001238
RTCA / O00442 / ENSMUSG00000000339 / ENSG00000137996 / ENSRNOG00000014575
RTCB / Q9Y3I0 / ENSMUSG00000001783 / ENSG00000100220 / ENSRNOG00000004813
RTF1 / Q92541 / ENSMUSG00000027304 / ENSG00000137815 / ENSRNOG00000005183
RTF2 / Q9BY42 / ENSMUSG00000027502 / ENSG00000022277 / ENSRNOG00000005083
RTKN / Q9BST9 / ENSMUSG00000034930 / ENSG00000114993 / ENSRNOG00000009022
RTN1 / Q16799 / ENSMUSG00000021087 / ENSG00000139970 / ENSRNOG00000004794
RTN2 / O75298 / ENSMUSG00000030401 / ENSG00000125744 / ENSRNOG00000016603
RTN3 / O95197 / ENSMUSG00000024758 / ENSG00000133318 / ENSRNOG00000021202
RTN4 / Q9NQC3 / ENSMUSG00000020458 / ENSG00000115310 / ENSRNOG00000004621
RTN4IP1 / Q8WWV3 / ENSMUSG00000019864 / ENSG00000130347 / ENSRNOG00000000279
RTP4 / Q96DX8 / ENSMUSG00000033355 / ENSG00000136514 / ENSRNOG00000028895
RTRAF / Q9Y224 / ENSMUSG00000021807 / ENSG00000087302 / ENSRNOG00000000340
Homologs
Brain
Heart
Kidney
Liver
Lung
Testis
Organs
Bins
1
2
3
4
5
Species
Human
Mouse
Rat

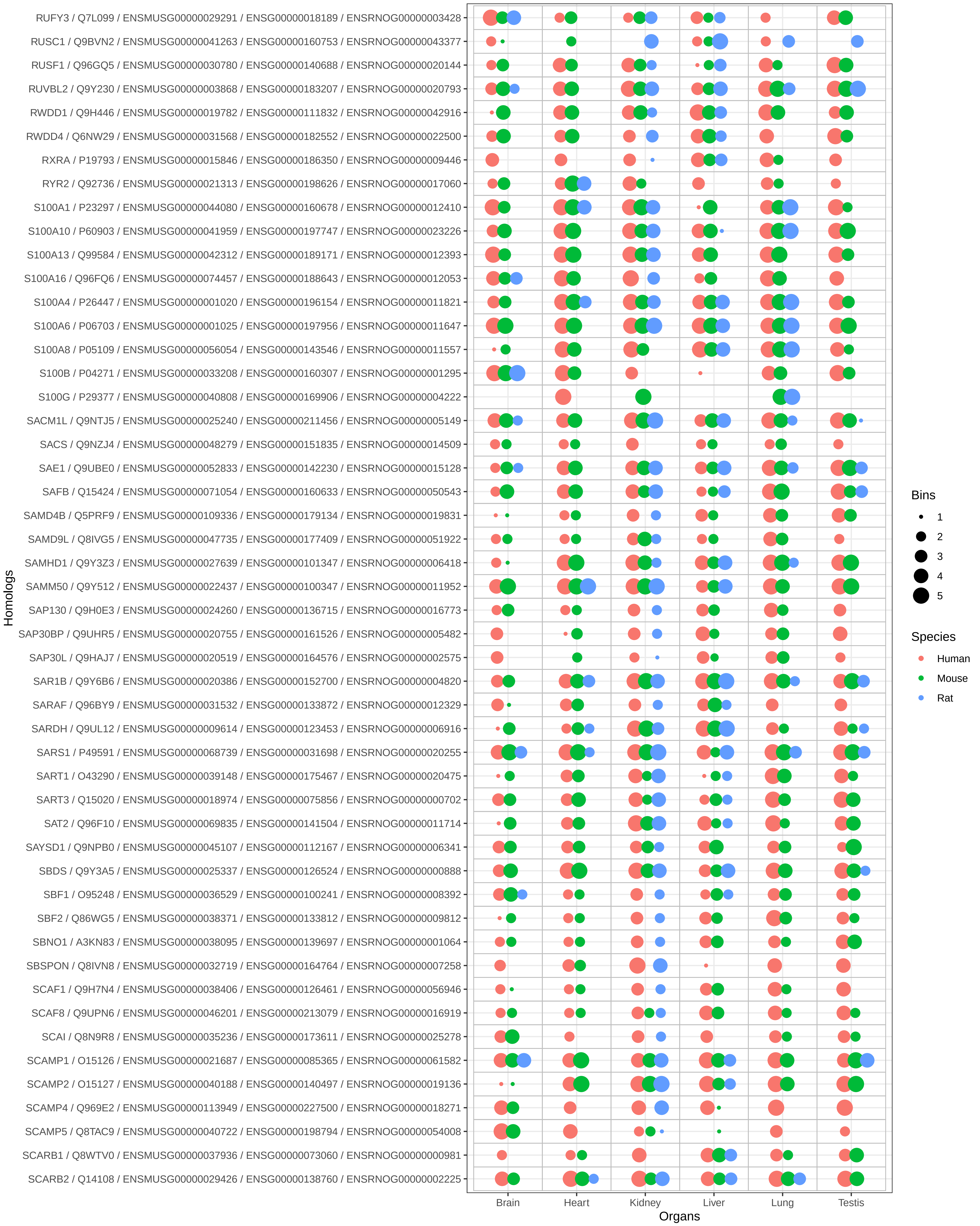
RUFY3 / Q7L099 / ENSMUSG00000029291 / ENSG00000018189 / ENSRNOG00000003428
RUSC1 / Q9BVN2 / ENSMUSG00000041263 / ENSG00000160753 / ENSRNOG00000043377
RUSF1 / Q96GQ5 / ENSMUSG00000030780 / ENSG00000140688 / ENSRNOG00000020144
RUVBL2 / Q9Y230 / ENSMUSG00000003868 / ENSG00000183207 / ENSRNOG00000020793
RWDD1 / Q9H446 / ENSMUSG00000019782 / ENSG00000111832 / ENSRNOG00000042916
RWDD4 / Q6NW29 / ENSMUSG00000031568 / ENSG00000182552 / ENSRNOG00000022500
RXRA / P19793 / ENSMUSG00000015846 / ENSG00000186350 / ENSRNOG00000009446
RYR2 / Q92736 / ENSMUSG00000021313 / ENSG00000198626 / ENSRNOG00000017060
S100A1 / P23297 / ENSMUSG00000044080 / ENSG00000160678 / ENSRNOG00000012410
S100A10 / P60903 / ENSMUSG00000041959 / ENSG00000197747 / ENSRNOG00000023226
S100A13 / Q99584 / ENSMUSG00000042312 / ENSG00000189171 / ENSRNOG00000012393
S100A16 / Q96FQ6 / ENSMUSG00000074457 / ENSG00000188643 / ENSRNOG00000012053
S100A4 / P26447 / ENSMUSG00000001020 / ENSG00000196154 / ENSRNOG00000011821
S100A6 / P06703 / ENSMUSG00000001025 / ENSG00000197956 / ENSRNOG00000011647
S100A8 / P05109 / ENSMUSG00000056054 / ENSG00000143546 / ENSRNOG00000011557
S100B / P04271 / ENSMUSG00000033208 / ENSG00000160307 / ENSRNOG00000001295
S100G / P29377 / ENSMUSG00000040808 / ENSG00000169906 / ENSRNOG00000004222
SACM1L / Q9NTJ5 / ENSMUSG00000025240 / ENSG00000211456 / ENSRNOG00000005149
SACS / Q9NZJ4 / ENSMUSG00000048279 / ENSG00000151835 / ENSRNOG00000014509
SAE1 / Q9UBE0 / ENSMUSG00000052833 / ENSG00000142230 / ENSRNOG00000015128
SAFB / Q15424 / ENSMUSG00000071054 / ENSG00000160633 / ENSRNOG00000050543
SAMD4B / Q5PRF9 / ENSMUSG00000109336 / ENSG00000179134 / ENSRNOG00000019831
SAMD9L / Q8IVG5 / ENSMUSG00000047735 / ENSG00000177409 / ENSRNOG00000051922
SAMHD1 / Q9Y3Z3 / ENSMUSG00000027639 / ENSG00000101347 / ENSRNOG00000006418
SAMM50 / Q9Y512 / ENSMUSG00000022437 / ENSG00000100347 / ENSRNOG00000011952
SAP130 / Q9H0E3 / ENSMUSG00000024260 / ENSG00000136715 / ENSRNOG00000016773
SAP30BP / Q9UHR5 / ENSMUSG00000020755 / ENSG00000161526 / ENSRNOG00000005482
SAP30L / Q9HAJ7 / ENSMUSG00000020519 / ENSG00000164576 / ENSRNOG00000002575
SAR1B / Q9Y6B6 / ENSMUSG00000020386 / ENSG00000152700 / ENSRNOG00000004820
SARAF / Q96BY9 / ENSMUSG00000031532 / ENSG00000133872 / ENSRNOG00000012329
SARDH / Q9UL12 / ENSMUSG00000009614 / ENSG00000123453 / ENSRNOG00000006916
SARS1 / P49591 / ENSMUSG00000068739 / ENSG00000031698 / ENSRNOG00000020255
SART1 / O43290 / ENSMUSG00000039148 / ENSG00000175467 / ENSRNOG00000020475
SART3 / Q15020 / ENSMUSG00000018974 / ENSG00000075856 / ENSRNOG00000000702
SAT2 / Q96F10 / ENSMUSG00000069835 / ENSG00000141504 / ENSRNOG00000011714
SAYSD1 / Q9NPB0 / ENSMUSG00000045107 / ENSG00000112167 / ENSRNOG00000006341
SBDS / Q9Y3A5 / ENSMUSG00000025337 / ENSG00000126524 / ENSRNOG00000000888
SBF1 / O95248 / ENSMUSG00000036529 / ENSG00000100241 / ENSRNOG00000008392
SBF2 / Q86WG5 / ENSMUSG00000038371 / ENSG00000133812 / ENSRNOG00000009812
SBNO1 / A3KN83 / ENSMUSG00000038095 / ENSG00000139697 / ENSRNOG00000001064
SBSPON / Q8IVN8 / ENSMUSG00000032719 / ENSG00000164764 / ENSRNOG00000007258
SCAF1 / Q9H7N4 / ENSMUSG00000038406 / ENSG00000126461 / ENSRNOG00000056946
SCAF8 / Q9UPN6 / ENSMUSG00000046201 / ENSG00000213079 / ENSRNOG00000016919
SCAI / Q8N9R8 / ENSMUSG00000035236 / ENSG00000173611 / ENSRNOG00000025278
SCAMP1 / O15126 / ENSMUSG00000021687 / ENSG00000085365 / ENSRNOG00000061582
SCAMP2 / O15127 / ENSMUSG00000040188 / ENSG00000140497 / ENSRNOG00000019136
SCAMP4 / Q969E2 / ENSMUSG00000113949 / ENSG00000227500 / ENSRNOG00000018271
SCAMP5 / Q8TAC9 / ENSMUSG00000040722 / ENSG00000198794 / ENSRNOG00000054008
SCARB1 / Q8WTV0 / ENSMUSG00000037936 / ENSG00000073060 / ENSRNOG00000000981
SCARB2 / Q14108 / ENSMUSG00000029426 / ENSG00000138760 / ENSRNOG00000002225
Homologs
Brain
Heart
Kidney
Liver
Lung
Testis
Organs
Bins
1
2
3
4
5
Species
Human
Mouse
Rat

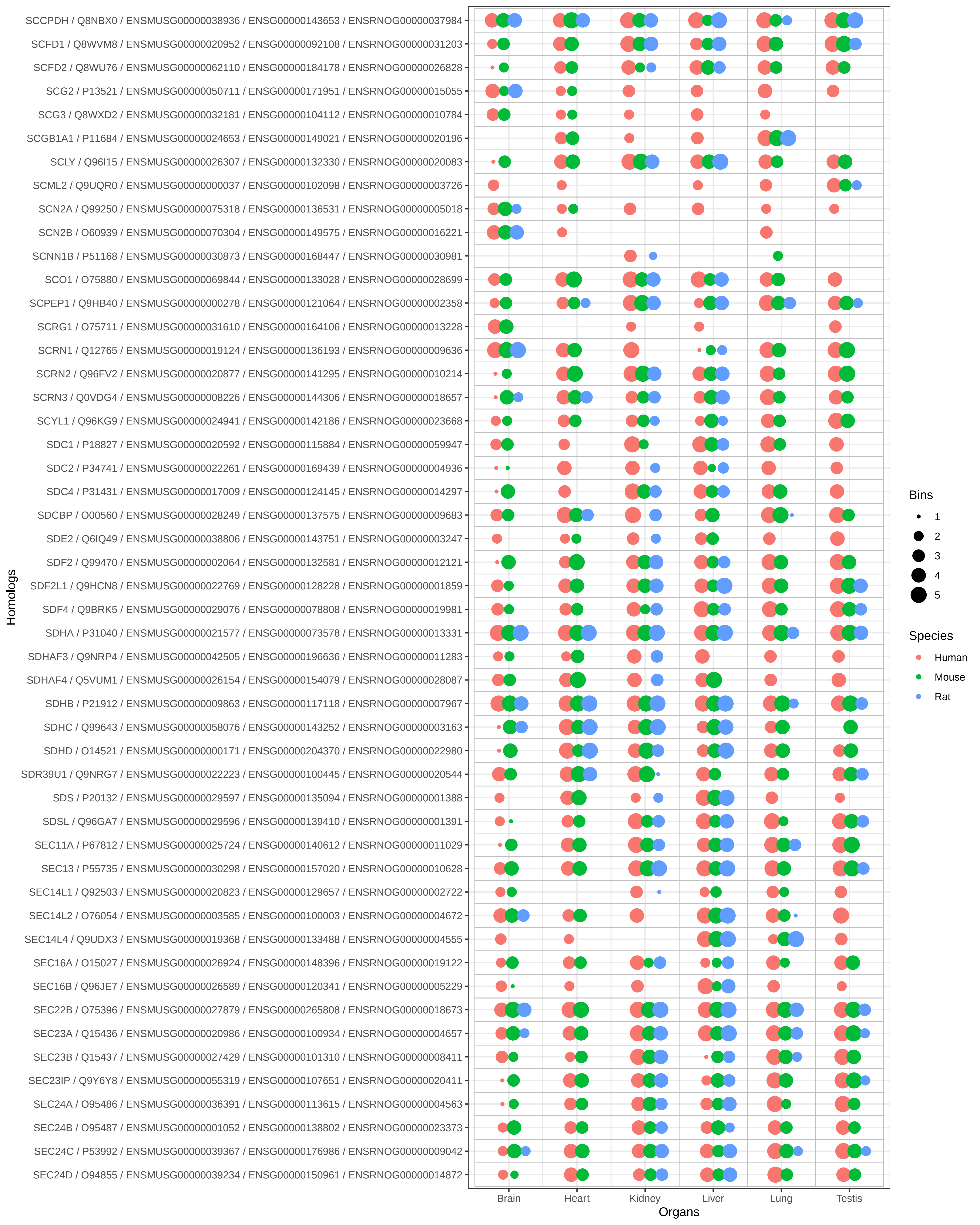
SCCPDH / Q8NBX0 / ENSMUSG00000038936 / ENSG00000143653 / ENSRNOG00000037984
SCFD1 / Q8WVM8 / ENSMUSG00000020952 / ENSG00000092108 / ENSRNOG00000031203
SCFD2 / Q8WU76 / ENSMUSG00000062110 / ENSG00000184178 / ENSRNOG00000026828
SCG2 / P13521 / ENSMUSG00000050711 / ENSG00000171951 / ENSRNOG00000015055
SCG3 / Q8WXD2 / ENSMUSG00000032181 / ENSG00000104112 / ENSRNOG00000010784
SCGB1A1 / P11684 / ENSMUSG00000024653 / ENSG00000149021 / ENSRNOG00000020196
SCLY / Q96I15 / ENSMUSG00000026307 / ENSG00000132330 / ENSRNOG00000020083
SCML2 / Q9UQR0 / ENSMUSG00000000037 / ENSG00000102098 / ENSRNOG00000003726
SCN2A / Q99250 / ENSMUSG00000075318 / ENSG00000136531 / ENSRNOG00000005018
SCN2B / O60939 / ENSMUSG00000070304 / ENSG00000149575 / ENSRNOG00000016221
SCNN1B / P51168 / ENSMUSG00000030873 / ENSG00000168447 / ENSRNOG00000030981
SCO1 / O75880 / ENSMUSG00000069844 / ENSG00000133028 / ENSRNOG00000028699
SCPEP1 / Q9HB40 / ENSMUSG00000000278 / ENSG00000121064 / ENSRNOG00000002358
SCRG1 / O75711 / ENSMUSG00000031610 / ENSG00000164106 / ENSRNOG00000013228
SCRN1 / Q12765 / ENSMUSG00000019124 / ENSG00000136193 / ENSRNOG00000009636
SCRN2 / Q96FV2 / ENSMUSG00000020877 / ENSG00000141295 / ENSRNOG00000010214
SCRN3 / Q0VDG4 / ENSMUSG00000008226 / ENSG00000144306 / ENSRNOG00000018657
SCYL1 / Q96KG9 / ENSMUSG00000024941 / ENSG00000142186 / ENSRNOG00000023668
SDC1 / P18827 / ENSMUSG00000020592 / ENSG00000115884 / ENSRNOG00000059947
SDC2 / P34741 / ENSMUSG00000022261 / ENSG00000169439 / ENSRNOG00000004936
SDC4 / P31431 / ENSMUSG00000017009 / ENSG00000124145 / ENSRNOG00000014297
SDCBP / O00560 / ENSMUSG00000028249 / ENSG00000137575 / ENSRNOG00000009683
SDE2 / Q6IQ49 / ENSMUSG00000038806 / ENSG00000143751 / ENSRNOG00000003247
SDF2 / Q99470 / ENSMUSG00000002064 / ENSG00000132581 / ENSRNOG00000012121
SDF2L1 / Q9HCN8 / ENSMUSG00000022769 / ENSG00000128228 / ENSRNOG00000001859
SDF4 / Q9BRK5 / ENSMUSG00000029076 / ENSG00000078808 / ENSRNOG00000019981
SDHA / P31040 / ENSMUSG00000021577 / ENSG00000073578 / ENSRNOG00000013331
SDHAF3 / Q9NRP4 / ENSMUSG00000042505 / ENSG00000196636 / ENSRNOG00000011283
SDHAF4 / Q5VUM1 / ENSMUSG00000026154 / ENSG00000154079 / ENSRNOG00000028087
SDHB / P21912 / ENSMUSG00000009863 / ENSG00000117118 / ENSRNOG00000007967
SDHC / Q99643 / ENSMUSG00000058076 / ENSG00000143252 / ENSRNOG00000003163
SDHD / O14521 / ENSMUSG00000000171 / ENSG00000204370 / ENSRNOG00000022980
SDR39U1 / Q9NRG7 / ENSMUSG00000022223 / ENSG00000100445 / ENSRNOG00000020544
SDS / P20132 / ENSMUSG00000029597 / ENSG00000135094 / ENSRNOG00000001388
SDSL / Q96GA7 / ENSMUSG00000029596 / ENSG00000139410 / ENSRNOG00000001391
SEC11A / P67812 / ENSMUSG00000025724 / ENSG00000140612 / ENSRNOG00000011029
SEC13 / P55735 / ENSMUSG00000030298 / ENSG00000157020 / ENSRNOG00000010628
SEC14L1 / Q92503 / ENSMUSG00000020823 / ENSG00000129657 / ENSRNOG00000002722
SEC14L2 / O76054 / ENSMUSG00000003585 / ENSG00000100003 / ENSRNOG00000004672
SEC14L4 / Q9UDX3 / ENSMUSG00000019368 / ENSG00000133488 / ENSRNOG00000004555
SEC16A / O15027 / ENSMUSG00000026924 / ENSG00000148396 / ENSRNOG00000019122
SEC16B / Q96JE7 / ENSMUSG00000026589 / ENSG00000120341 / ENSRNOG00000005229
SEC22B / O75396 / ENSMUSG00000027879 / ENSG00000265808 / ENSRNOG00000018673
SEC23A / Q15436 / ENSMUSG00000020986 / ENSG00000100934 / ENSRNOG00000004657
SEC23B / Q15437 / ENSMUSG00000027429 / ENSG00000101310 / ENSRNOG00000008411
SEC23IP / Q9Y6Y8 / ENSMUSG00000055319 / ENSG00000107651 / ENSRNOG00000020411
SEC24A / O95486 / ENSMUSG00000036391 / ENSG00000113615 / ENSRNOG00000004563
SEC24B / O95487 / ENSMUSG00000001052 / ENSG00000138802 / ENSRNOG00000023373
SEC24C / P53992 / ENSMUSG00000039367 / ENSG00000176986 / ENSRNOG00000009042
SEC24D / O94855 / ENSMUSG00000039234 / ENSG00000150961 / ENSRNOG00000014872
Homologs
Brain
Heart
Kidney
Liver
Lung
Testis
Organs
Bins
1
2
3
4
5
Species
Human
Mouse
Rat

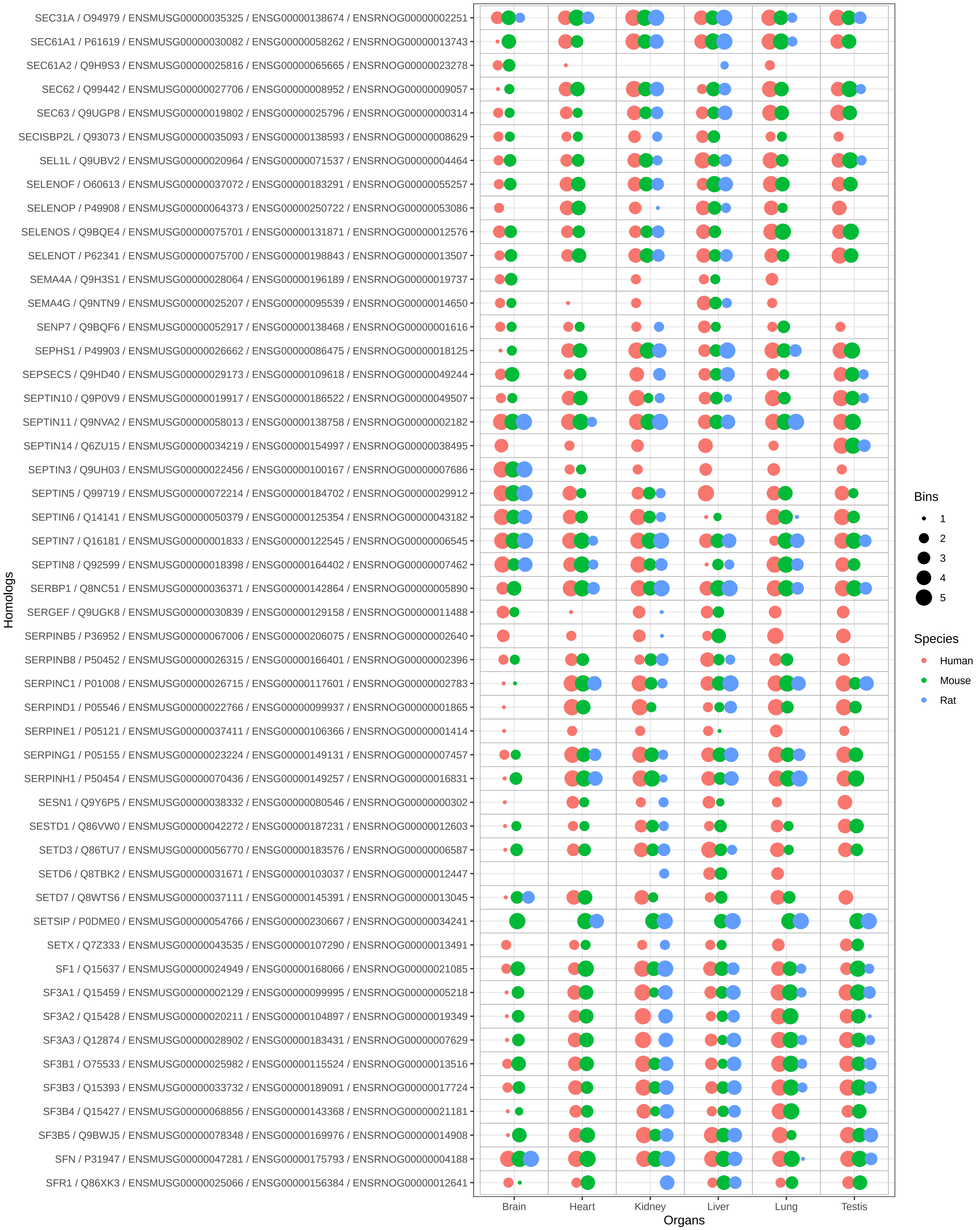
SEC31A / O94979 / ENSMUSG00000035325 / ENSG00000138674 / ENSRNOG00000002251
SEC61A1 / P61619 / ENSMUSG00000030082 / ENSG00000058262 / ENSRNOG00000013743
SEC61A2 / Q9H9S3 / ENSMUSG00000025816 / ENSG00000065665 / ENSRNOG00000023278
SEC62 / Q99442 / ENSMUSG00000027706 / ENSG00000008952 / ENSRNOG00000009057
SEC63 / Q9UGP8 / ENSMUSG00000019802 / ENSG00000025796 / ENSRNOG00000000314
SECISBP2L / Q93073 / ENSMUSG00000035093 / ENSG00000138593 / ENSRNOG00000008629
SEL1L / Q9UBV2 / ENSMUSG00000020964 / ENSG00000071537 / ENSRNOG00000004464
SELENOF / O60613 / ENSMUSG00000037072 / ENSG00000183291 / ENSRNOG00000055257
SELENOP / P49908 / ENSMUSG00000064373 / ENSG00000250722 / ENSRNOG00000053086
SELENOS / Q9BQE4 / ENSMUSG00000075701 / ENSG00000131871 / ENSRNOG00000012576
SELENOT / P62341 / ENSMUSG00000075700 / ENSG00000198843 / ENSRNOG00000013507
SEMA4A / Q9H3S1 / ENSMUSG00000028064 / ENSG00000196189 / ENSRNOG00000019737
SEMA4G / Q9NTN9 / ENSMUSG00000025207 / ENSG00000095539 / ENSRNOG00000014650
SENP7 / Q9BQF6 / ENSMUSG00000052917 / ENSG00000138468 / ENSRNOG00000001616
SEPHS1 / P49903 / ENSMUSG00000026662 / ENSG00000086475 / ENSRNOG00000018125
SEPSECS / Q9HD40 / ENSMUSG00000029173 / ENSG00000109618 / ENSRNOG00000049244
SEPTIN10 / Q9P0V9 / ENSMUSG00000019917 / ENSG00000186522 / ENSRNOG00000049507
SEPTIN11 / Q9NVA2 / ENSMUSG00000058013 / ENSG00000138758 / ENSRNOG00000002182
SEPTIN14 / Q6ZU15 / ENSMUSG00000034219 / ENSG00000154997 / ENSRNOG00000038495
SEPTIN3 / Q9UH03 / ENSMUSG00000022456 / ENSG00000100167 / ENSRNOG00000007686
SEPTIN5 / Q99719 / ENSMUSG00000072214 / ENSG00000184702 / ENSRNOG00000029912
SEPTIN6 / Q14141 / ENSMUSG00000050379 / ENSG00000125354 / ENSRNOG00000043182
SEPTIN7 / Q16181 / ENSMUSG00000001833 / ENSG00000122545 / ENSRNOG00000006545
SEPTIN8 / Q92599 / ENSMUSG00000018398 / ENSG00000164402 / ENSRNOG00000007462
SERBP1 / Q8NC51 / ENSMUSG00000036371 / ENSG00000142864 / ENSRNOG00000005890
SERGEF / Q9UGK8 / ENSMUSG00000030839 / ENSG00000129158 / ENSRNOG00000011488
SERPINB5 / P36952 / ENSMUSG00000067006 / ENSG00000206075 / ENSRNOG00000002640
SERPINB8 / P50452 / ENSMUSG00000026315 / ENSG00000166401 / ENSRNOG00000002396
SERPINC1 / P01008 / ENSMUSG00000026715 / ENSG00000117601 / ENSRNOG00000002783
SERPIND1 / P05546 / ENSMUSG00000022766 / ENSG00000099937 / ENSRNOG00000001865
SERPINE1 / P05121 / ENSMUSG00000037411 / ENSG00000106366 / ENSRNOG00000001414
SERPING1 / P05155 / ENSMUSG00000023224 / ENSG00000149131 / ENSRNOG00000007457
SERPINH1 / P50454 / ENSMUSG00000070436 / ENSG00000149257 / ENSRNOG00000016831
SESN1 / Q9Y6P5 / ENSMUSG00000038332 / ENSG00000080546 / ENSRNOG00000000302
SESTD1 / Q86VW0 / ENSMUSG00000042272 / ENSG00000187231 / ENSRNOG00000012603
SETD3 / Q86TU7 / ENSMUSG00000056770 / ENSG00000183576 / ENSRNOG00000006587
SETD6 / Q8TBK2 / ENSMUSG00000031671 / ENSG00000103037 / ENSRNOG00000012447
SETD7 / Q8WTS6 / ENSMUSG00000037111 / ENSG00000145391 / ENSRNOG00000013045
SETSIP / P0DME0 / ENSMUSG00000054766 / ENSG00000230667 / ENSRNOG00000034241
SETX / Q7Z333 / ENSMUSG00000043535 / ENSG00000107290 / ENSRNOG00000013491
SF1 / Q15637 / ENSMUSG00000024949 / ENSG00000168066 / ENSRNOG00000021085
SF3A1 / Q15459 / ENSMUSG00000002129 / ENSG00000099995 / ENSRNOG00000005218
SF3A2 / Q15428 / ENSMUSG00000020211 / ENSG00000104897 / ENSRNOG00000019349
SF3A3 / Q12874 / ENSMUSG00000028902 / ENSG00000183431 / ENSRNOG00000007629
SF3B1 / O75533 / ENSMUSG00000025982 / ENSG00000115524 / ENSRNOG00000013516
SF3B3 / Q15393 / ENSMUSG00000033732 / ENSG00000189091 / ENSRNOG00000017724
SF3B4 / Q15427 / ENSMUSG00000068856 / ENSG00000143368 / ENSRNOG00000021181
SF3B5 / Q9BWJ5 / ENSMUSG00000078348 / ENSG00000169976 / ENSRNOG00000014908
SFN / P31947 / ENSMUSG00000047281 / ENSG00000175793 / ENSRNOG00000004188
SFR1 / Q86XK3 / ENSMUSG00000025066 / ENSG00000156384 / ENSRNOG00000012641
Homologs
Brain
Heart
Kidney
Liver
Lung
Testis
Organs
Bins
1
2
3
4
5
Species
Human
Mouse
Rat

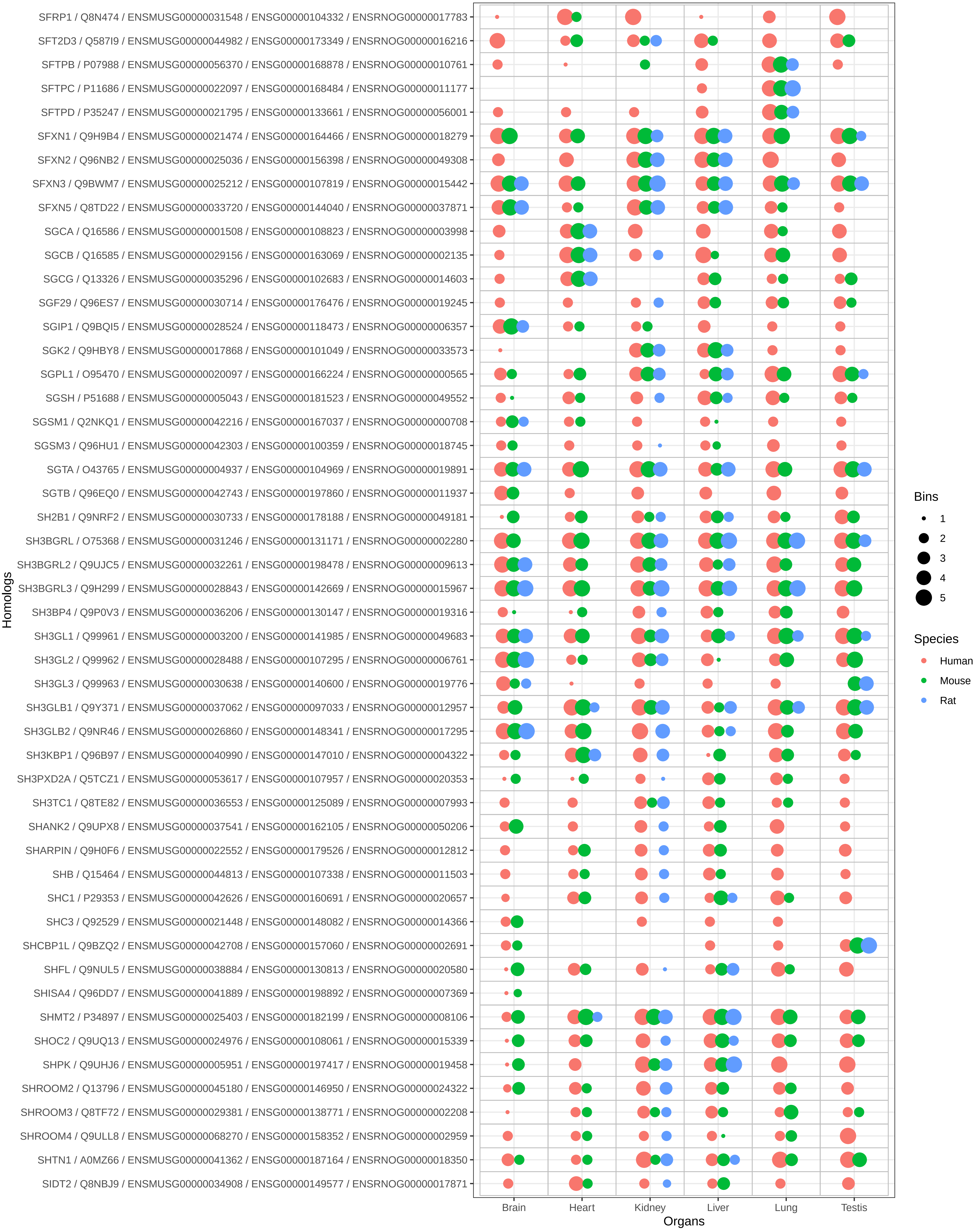
SFRP1 / Q8N474 / ENSMUSG00000031548 / ENSG00000104332 / ENSRNOG00000017783
SFT2D3 / Q587I9 / ENSMUSG00000044982 / ENSG00000173349 / ENSRNOG00000016216
SFTPB / P07988 / ENSMUSG00000056370 / ENSG00000168878 / ENSRNOG00000010761
SFTPC / P11686 / ENSMUSG00000022097 / ENSG00000168484 / ENSRNOG00000011177
SFTPD / P35247 / ENSMUSG00000021795 / ENSG00000133661 / ENSRNOG00000056001
SFXN1 / Q9H9B4 / ENSMUSG00000021474 / ENSG00000164466 / ENSRNOG00000018279
SFXN2 / Q96NB2 / ENSMUSG00000025036 / ENSG00000156398 / ENSRNOG00000049308
SFXN3 / Q9BWM7 / ENSMUSG00000025212 / ENSG00000107819 / ENSRNOG00000015442
SFXN5 / Q8TD22 / ENSMUSG00000033720 / ENSG00000144040 / ENSRNOG00000037871
SGCA / Q16586 / ENSMUSG00000001508 / ENSG00000108823 / ENSRNOG00000003998
SGCB / Q16585 / ENSMUSG00000029156 / ENSG00000163069 / ENSRNOG00000002135
SGCG / Q13326 / ENSMUSG00000035296 / ENSG00000102683 / ENSRNOG00000014603
SGF29 / Q96ES7 / ENSMUSG00000030714 / ENSG00000176476 / ENSRNOG00000019245
SGIP1 / Q9BQI5 / ENSMUSG00000028524 / ENSG00000118473 / ENSRNOG00000006357
SGK2 / Q9HBY8 / ENSMUSG00000017868 / ENSG00000101049 / ENSRNOG00000033573
SGPL1 / O95470 / ENSMUSG00000020097 / ENSG00000166224 / ENSRNOG00000000565
SGSH / P51688 / ENSMUSG00000005043 / ENSG00000181523 / ENSRNOG00000049552
SGSM1 / Q2NKQ1 / ENSMUSG00000042216 / ENSG00000167037 / ENSRNOG00000000708
SGSM3 / Q96HU1 / ENSMUSG00000042303 / ENSG00000100359 / ENSRNOG00000018745
SGTA / O43765 / ENSMUSG00000004937 / ENSG00000104969 / ENSRNOG00000019891
SGTB / Q96EQ0 / ENSMUSG00000042743 / ENSG00000197860 / ENSRNOG00000011937
SH2B1 / Q9NRF2 / ENSMUSG00000030733 / ENSG00000178188 / ENSRNOG00000049181
SH3BGRL / O75368 / ENSMUSG00000031246 / ENSG00000131171 / ENSRNOG00000002280
SH3BGRL2 / Q9UJC5 / ENSMUSG00000032261 / ENSG00000198478 / ENSRNOG00000009613
SH3BGRL3 / Q9H299 / ENSMUSG00000028843 / ENSG00000142669 / ENSRNOG00000015967
SH3BP4 / Q9P0V3 / ENSMUSG00000036206 / ENSG00000130147 / ENSRNOG00000019316
SH3GL1 / Q99961 / ENSMUSG00000003200 / ENSG00000141985 / ENSRNOG00000049683
SH3GL2 / Q99962 / ENSMUSG00000028488 / ENSG00000107295 / ENSRNOG00000006761
SH3GL3 / Q99963 / ENSMUSG00000030638 / ENSG00000140600 / ENSRNOG00000019776
SH3GLB1 / Q9Y371 / ENSMUSG00000037062 / ENSG00000097033 / ENSRNOG00000012957
SH3GLB2 / Q9NR46 / ENSMUSG00000026860 / ENSG00000148341 / ENSRNOG00000017295
SH3KBP1 / Q96B97 / ENSMUSG00000040990 / ENSG00000147010 / ENSRNOG00000004322
SH3PXD2A / Q5TCZ1 / ENSMUSG00000053617 / ENSG00000107957 / ENSRNOG00000020353
SH3TC1 / Q8TE82 / ENSMUSG00000036553 / ENSG00000125089 / ENSRNOG00000007993
SHANK2 / Q9UPX8 / ENSMUSG00000037541 / ENSG00000162105 / ENSRNOG00000050206
SHARPIN / Q9H0F6 / ENSMUSG00000022552 / ENSG00000179526 / ENSRNOG00000012812
SHB / Q15464 / ENSMUSG00000044813 / ENSG00000107338 / ENSRNOG00000011503
SHC1 / P29353 / ENSMUSG00000042626 / ENSG00000160691 / ENSRNOG00000020657
SHC3 / Q92529 / ENSMUSG00000021448 / ENSG00000148082 / ENSRNOG00000014366
SHCBP1L / Q9BZQ2 / ENSMUSG00000042708 / ENSG00000157060 / ENSRNOG00000002691
SHFL / Q9NUL5 / ENSMUSG00000038884 / ENSG00000130813 / ENSRNOG00000020580
SHISA4 / Q96DD7 / ENSMUSG00000041889 / ENSG00000198892 / ENSRNOG00000007369
SHMT2 / P34897 / ENSMUSG00000025403 / ENSG00000182199 / ENSRNOG00000008106
SHOC2 / Q9UQ13 / ENSMUSG00000024976 / ENSG00000108061 / ENSRNOG00000015339
SHPK / Q9UHJ6 / ENSMUSG00000005951 / ENSG00000197417 / ENSRNOG00000019458
SHROOM2 / Q13796 / ENSMUSG00000045180 / ENSG00000146950 / ENSRNOG00000024322
SHROOM3 / Q8TF72 / ENSMUSG00000029381 / ENSG00000138771 / ENSRNOG00000002208
SHROOM4 / Q9ULL8 / ENSMUSG00000068270 / ENSG00000158352 / ENSRNOG00000002959
SHTN1 / A0MZ66 / ENSMUSG00000041362 / ENSG00000187164 / ENSRNOG00000018350
SIDT2 / Q8NBJ9 / ENSMUSG00000034908 / ENSG00000149577 / ENSRNOG00000017871
Homologs
Brain
Heart
Kidney
Liver
Lung
Testis
Organs
Bins
1
2
3
4
5
Species
Human
Mouse
Rat

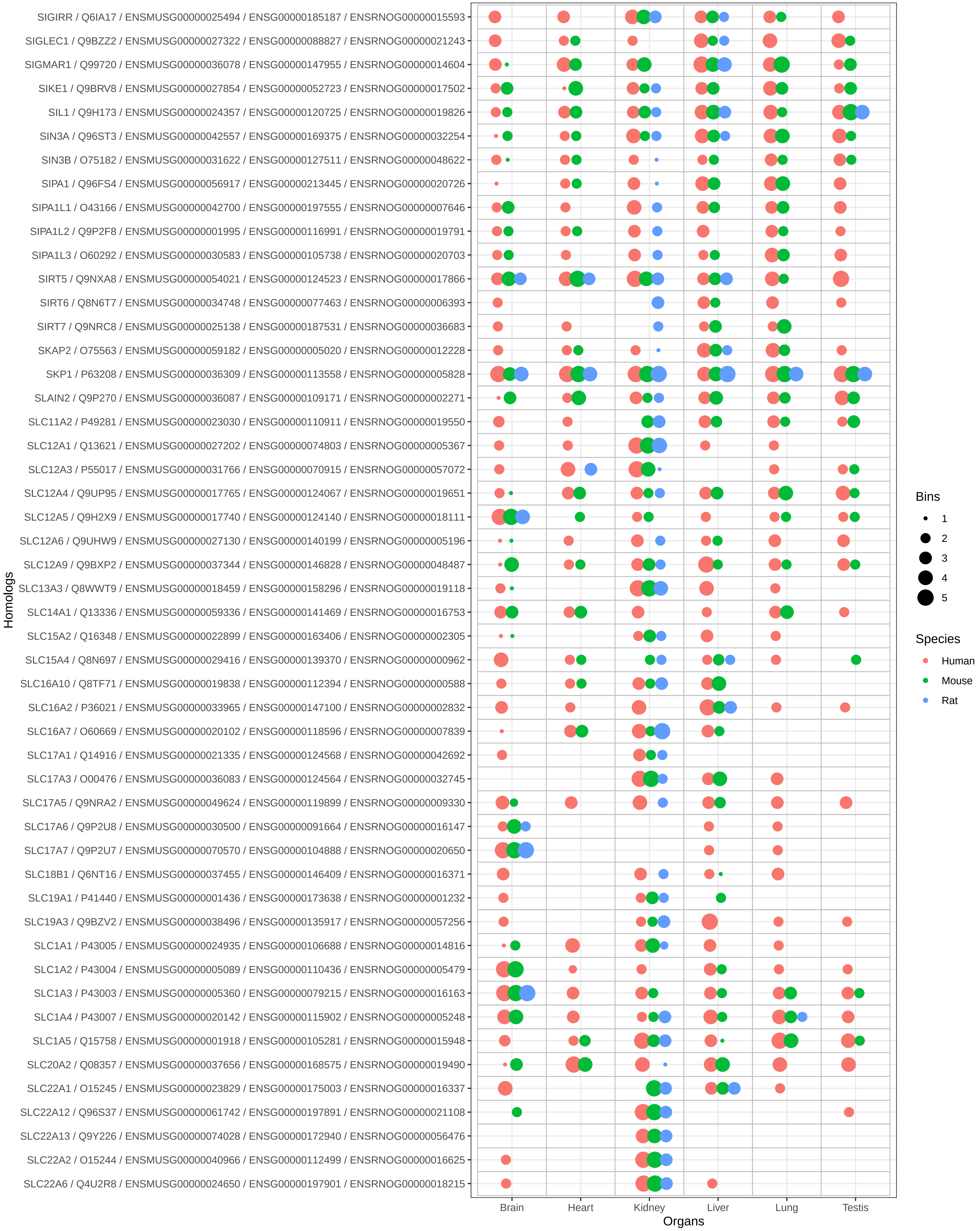
SIGIRR / Q6IA17 / ENSMUSG00000025494 / ENSG00000185187 / ENSRNOG00000015593
SIGLEC1 / Q9BZZ2 / ENSMUSG00000027322 / ENSG00000088827 / ENSRNOG00000021243
SIGMAR1 / Q99720 / ENSMUSG00000036078 / ENSG00000147955 / ENSRNOG00000014604
SIKE1 / Q9BRV8 / ENSMUSG00000027854 / ENSG00000052723 / ENSRNOG00000017502
SIL1 / Q9H173 / ENSMUSG00000024357 / ENSG00000120725 / ENSRNOG00000019826
SIN3A / Q96ST3 / ENSMUSG00000042557 / ENSG00000169375 / ENSRNOG00000032254
SIN3B / O75182 / ENSMUSG00000031622 / ENSG00000127511 / ENSRNOG00000048622
SIPA1 / Q96FS4 / ENSMUSG00000056917 / ENSG00000213445 / ENSRNOG00000020726
SIPA1L1 / O43166 / ENSMUSG00000042700 / ENSG00000197555 / ENSRNOG00000007646
SIPA1L2 / Q9P2F8 / ENSMUSG00000001995 / ENSG00000116991 / ENSRNOG00000019791
SIPA1L3 / O60292 / ENSMUSG00000030583 / ENSG00000105738 / ENSRNOG00000020703
SIRT5 / Q9NXA8 / ENSMUSG00000054021 / ENSG00000124523 / ENSRNOG00000017866
SIRT6 / Q8N6T7 / ENSMUSG00000034748 / ENSG00000077463 / ENSRNOG00000006393
SIRT7 / Q9NRC8 / ENSMUSG00000025138 / ENSG00000187531 / ENSRNOG00000036683
SKAP2 / O75563 / ENSMUSG00000059182 / ENSG00000005020 / ENSRNOG00000012228
SKP1 / P63208 / ENSMUSG00000036309 / ENSG00000113558 / ENSRNOG00000005828
SLAIN2 / Q9P270 / ENSMUSG00000036087 / ENSG00000109171 / ENSRNOG00000002271
SLC11A2 / P49281 / ENSMUSG00000023030 / ENSG00000110911 / ENSRNOG00000019550
SLC12A1 / Q13621 / ENSMUSG00000027202 / ENSG00000074803 / ENSRNOG00000005367
SLC12A3 / P55017 / ENSMUSG00000031766 / ENSG00000070915 / ENSRNOG00000057072
SLC12A4 / Q9UP95 / ENSMUSG00000017765 / ENSG00000124067 / ENSRNOG00000019651
SLC12A5 / Q9H2X9 / ENSMUSG00000017740 / ENSG00000124140 / ENSRNOG00000018111
SLC12A6 / Q9UHW9 / ENSMUSG00000027130 / ENSG00000140199 / ENSRNOG00000005196
SLC12A9 / Q9BXP2 / ENSMUSG00000037344 / ENSG00000146828 / ENSRNOG00000048487
SLC13A3 / Q8WWT9 / ENSMUSG00000018459 / ENSG00000158296 / ENSRNOG00000019118
SLC14A1 / Q13336 / ENSMUSG00000059336 / ENSG00000141469 / ENSRNOG00000016753
SLC15A2 / Q16348 / ENSMUSG00000022899 / ENSG00000163406 / ENSRNOG00000002305
SLC15A4 / Q8N697 / ENSMUSG00000029416 / ENSG00000139370 / ENSRNOG00000000962
SLC16A10 / Q8TF71 / ENSMUSG00000019838 / ENSG00000112394 / ENSRNOG00000000588
SLC16A2 / P36021 / ENSMUSG00000033965 / ENSG00000147100 / ENSRNOG00000002832
SLC16A7 / O60669 / ENSMUSG00000020102 / ENSG00000118596 / ENSRNOG00000007839
SLC17A1 / Q14916 / ENSMUSG00000021335 / ENSG00000124568 / ENSRNOG00000042692
SLC17A3 / O00476 / ENSMUSG00000036083 / ENSG00000124564 / ENSRNOG00000032745
SLC17A5 / Q9NRA2 / ENSMUSG00000049624 / ENSG00000119899 / ENSRNOG00000009330
SLC17A6 / Q9P2U8 / ENSMUSG00000030500 / ENSG00000091664 / ENSRNOG00000016147
SLC17A7 / Q9P2U7 / ENSMUSG00000070570 / ENSG00000104888 / ENSRNOG00000020650
SLC18B1 / Q6NT16 / ENSMUSG00000037455 / ENSG00000146409 / ENSRNOG00000016371
SLC19A1 / P41440 / ENSMUSG00000001436 / ENSG00000173638 / ENSRNOG00000001232
SLC19A3 / Q9BZV2 / ENSMUSG00000038496 / ENSG00000135917 / ENSRNOG00000057256
SLC1A1 / P43005 / ENSMUSG00000024935 / ENSG00000106688 / ENSRNOG00000014816
SLC1A2 / P43004 / ENSMUSG00000005089 / ENSG00000110436 / ENSRNOG00000005479
SLC1A3 / P43003 / ENSMUSG00000005360 / ENSG00000079215 / ENSRNOG00000016163
SLC1A4 / P43007 / ENSMUSG00000020142 / ENSG00000115902 / ENSRNOG00000005248
SLC1A5 / Q15758 / ENSMUSG00000001918 / ENSG00000105281 / ENSRNOG00000015948
SLC20A2 / Q08357 / ENSMUSG00000037656 / ENSG00000168575 / ENSRNOG00000019490
SLC22A1 / O15245 / ENSMUSG00000023829 / ENSG00000175003 / ENSRNOG00000016337
SLC22A12 / Q96S37 / ENSMUSG00000061742 / ENSG00000197891 / ENSRNOG00000021108
SLC22A13 / Q9Y226 / ENSMUSG00000074028 / ENSG00000172940 / ENSRNOG00000056476
SLC22A2 / O15244 / ENSMUSG00000040966 / ENSG00000112499 / ENSRNOG00000016625
SLC22A6 / Q4U2R8 / ENSMUSG00000024650 / ENSG00000197901 / ENSRNOG00000018215
Homologs
Brain
Heart
Kidney
Liver
Lung
Testis
Organs
Bins
1
2
3
4
5
Species
Human
Mouse
Rat

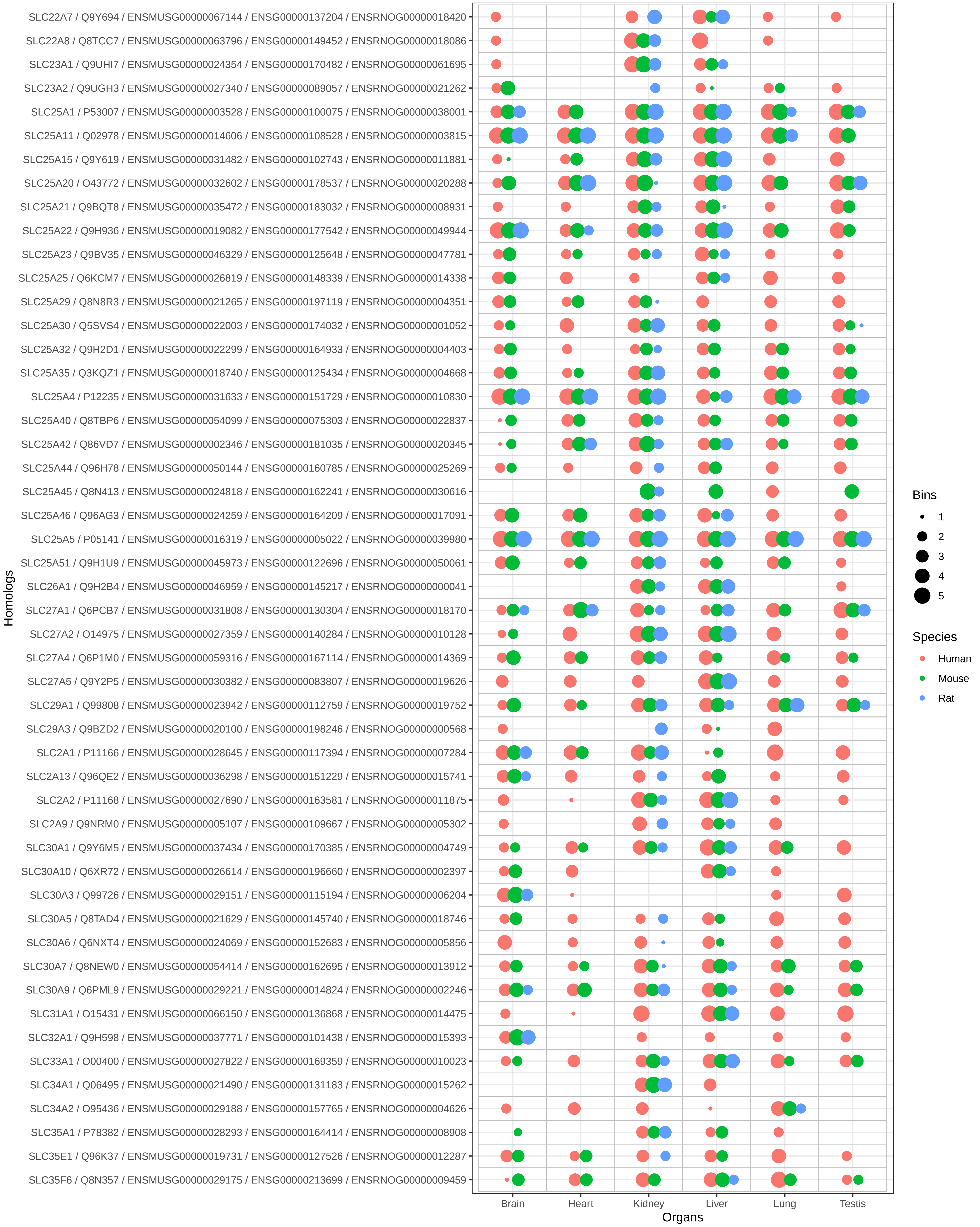
SLC22A7 / Q9Y694 / ENSMUSG00000067144 / ENSG00000137204 / ENSRNOG00000018420
SLC22A8 / Q8TCC7 / ENSMUSG00000063796 / ENSG00000149452 / ENSRNOG00000018086
SLC23A1 / Q9UHI7 / ENSMUSG00000024354 / ENSG00000170482 / ENSRNOG00000061695
SLC23A2 / Q9UGH3 / ENSMUSG00000027340 / ENSG00000089057 / ENSRNOG00000021262
SLC25A1 / P53007 / ENSMUSG00000003528 / ENSG00000100075 / ENSRNOG00000038001
SLC25A11 / Q02978 / ENSMUSG00000014606 / ENSG00000108528 / ENSRNOG00000003815
SLC25A15 / Q9Y619 / ENSMUSG00000031482 / ENSG00000102743 / ENSRNOG00000011881
SLC25A20 / O43772 / ENSMUSG00000032602 / ENSG00000178537 / ENSRNOG00000020288
SLC25A21 / Q9BQT8 / ENSMUSG00000035472 / ENSG00000183032 / ENSRNOG00000008931
SLC25A22 / Q9H936 / ENSMUSG00000019082 / ENSG00000177542 / ENSRNOG00000049944
SLC25A23 / Q9BV35 / ENSMUSG00000046329 / ENSG00000125648 / ENSRNOG00000047781
SLC25A25 / Q6KCM7 / ENSMUSG00000026819 / ENSG00000148339 / ENSRNOG00000014338
SLC25A29 / Q8N8R3 / ENSMUSG00000021265 / ENSG00000197119 / ENSRNOG00000004351
SLC25A30 / Q5SVS4 / ENSMUSG00000022003 / ENSG00000174032 / ENSRNOG00000001052
SLC25A32 / Q9H2D1 / ENSMUSG00000022299 / ENSG00000164933 / ENSRNOG00000004403
SLC25A35 / Q3KQZ1 / ENSMUSG00000018740 / ENSG00000125434 / ENSRNOG00000004668
SLC25A4 / P12235 / ENSMUSG00000031633 / ENSG00000151729 / ENSRNOG00000010830
SLC25A40 / Q8TBP6 / ENSMUSG00000054099 / ENSG00000075303 / ENSRNOG00000022837
SLC25A42 / Q86VD7 / ENSMUSG00000002346 / ENSG00000181035 / ENSRNOG00000020345
SLC25A44 / Q96H78 / ENSMUSG00000050144 / ENSG00000160785 / ENSRNOG00000025269
SLC25A45 / Q8N413 / ENSMUSG00000024818 / ENSG00000162241 / ENSRNOG00000030616
SLC25A46 / Q96AG3 / ENSMUSG00000024259 / ENSG00000164209 / ENSRNOG00000017091
SLC25A5 / P05141 / ENSMUSG00000016319 / ENSG00000005022 / ENSRNOG00000039980
SLC25A51 / Q9H1U9 / ENSMUSG00000045973 / ENSG00000122696 / ENSRNOG00000050061
SLC26A1 / Q9H2B4 / ENSMUSG00000046959 / ENSG00000145217 / ENSRNOG00000000041
SLC27A1 / Q6PCB7 / ENSMUSG00000031808 / ENSG00000130304 / ENSRNOG00000018170
SLC27A2 / O14975 / ENSMUSG00000027359 / ENSG00000140284 / ENSRNOG00000010128
SLC27A4 / Q6P1M0 / ENSMUSG00000059316 / ENSG00000167114 / ENSRNOG00000014369
SLC27A5 / Q9Y2P5 / ENSMUSG00000030382 / ENSG00000083807 / ENSRNOG00000019626
SLC29A1 / Q99808 / ENSMUSG00000023942 / ENSG00000112759 / ENSRNOG00000019752
SLC29A3 / Q9BZD2 / ENSMUSG00000020100 / ENSG00000198246 / ENSRNOG00000000568
SLC2A1 / P11166 / ENSMUSG00000028645 / ENSG00000117394 / ENSRNOG00000007284
SLC2A13 / Q96QE2 / ENSMUSG00000036298 / ENSG00000151229 / ENSRNOG00000015741
SLC2A2 / P11168 / ENSMUSG00000027690 / ENSG00000163581 / ENSRNOG00000011875
SLC2A9 / Q9NRM0 / ENSMUSG00000005107 / ENSG00000109667 / ENSRNOG00000005302
SLC30A1 / Q9Y6M5 / ENSMUSG00000037434 / ENSG00000170385 / ENSRNOG00000004749
SLC30A10 / Q6XR72 / ENSMUSG00000026614 / ENSG00000196660 / ENSRNOG00000002397
SLC30A3 / Q99726 / ENSMUSG00000029151 / ENSG00000115194 / ENSRNOG00000006204
SLC30A5 / Q8TAD4 / ENSMUSG00000021629 / ENSG00000145740 / ENSRNOG00000018746
SLC30A6 / Q6NXT4 / ENSMUSG00000024069 / ENSG00000152683 / ENSRNOG00000005856
SLC30A7 / Q8NEW0 / ENSMUSG00000054414 / ENSG00000162695 / ENSRNOG00000013912
SLC30A9 / Q6PML9 / ENSMUSG00000029221 / ENSG00000014824 / ENSRNOG00000002246
SLC31A1 / O15431 / ENSMUSG00000066150 / ENSG00000136868 / ENSRNOG00000014475
SLC32A1 / Q9H598 / ENSMUSG00000037771 / ENSG00000101438 / ENSRNOG00000015393
SLC33A1 / O00400 / ENSMUSG00000027822 / ENSG00000169359 / ENSRNOG00000010023
SLC34A1 / Q06495 / ENSMUSG00000021490 / ENSG00000131183 / ENSRNOG00000015262
SLC34A2 / O95436 / ENSMUSG00000029188 / ENSG00000157765 / ENSRNOG00000004626
SLC35A1 / P78382 / ENSMUSG00000028293 / ENSG00000164414 / ENSRNOG00000008908
SLC35E1 / Q96K37 / ENSMUSG00000019731 / ENSG00000127526 / ENSRNOG00000012287
SLC35F6 / Q8N357 / ENSMUSG00000029175 / ENSG00000213699 / ENSRNOG00000009459
Homologs
Brain
Heart
Kidney
Liver
Lung
Testis
Organs
Bins
1
2
3
4
5
Species
Human
Mouse
Rat

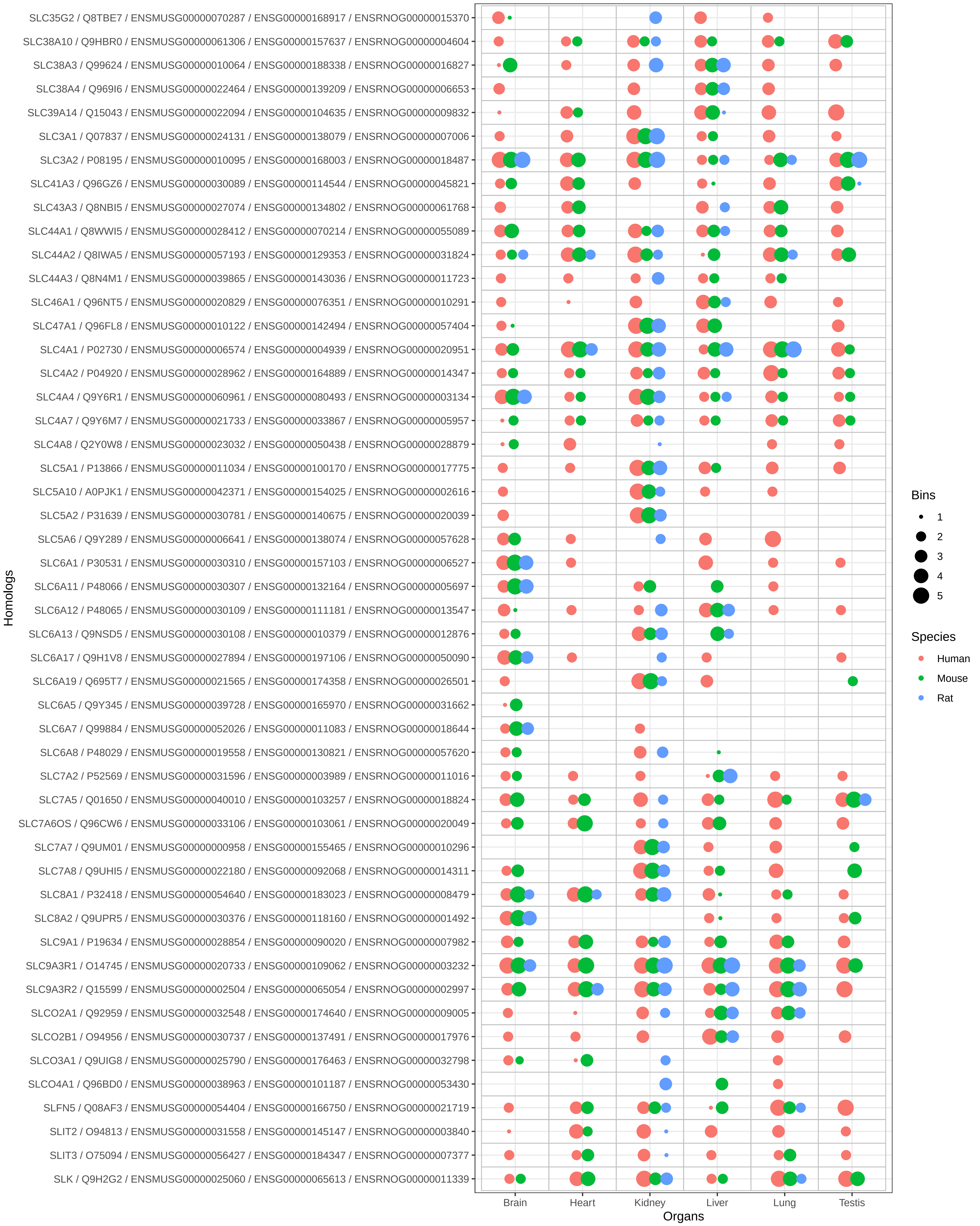

SLC35G2 / Q8TBE7 / ENSMUSG00000070287 / ENSG00000168917 / ENSRNOG00000015370
SLC38A10 / Q9HBR0 / ENSMUSG00000061306 / ENSG00000157637 / ENSRNOG00000004604
SLC38A3 / Q99624 / ENSMUSG00000010064 / ENSG00000188338 / ENSRNOG00000016827
SLC38A4 / Q969I6 / ENSMUSG00000022464 / ENSG00000139209 / ENSRNOG00000006653
SLC39A14 / Q15043 / ENSMUSG00000022094 / ENSG00000104635 / ENSRNOG00000009832
SLC3A1 / Q07837 / ENSMUSG00000024131 / ENSG00000138079 / ENSRNOG00000007006
SLC3A2 / P08195 / ENSMUSG00000010095 / ENSG00000168003 / ENSRNOG00000018487
SLC41A3 / Q96GZ6 / ENSMUSG00000030089 / ENSG00000114544 / ENSRNOG00000045821
SLC43A3 / Q8NBI5 / ENSMUSG00000027074 / ENSG00000134802 / ENSRNOG00000061768
SLC44A1 / Q8WWI5 / ENSMUSG00000028412 / ENSG00000070214 / ENSRNOG00000055089
SLC44A2 / Q8IWA5 / ENSMUSG00000057193 / ENSG00000129353 / ENSRNOG00000031824
SLC44A3 / Q8N4M1 / ENSMUSG00000039865 / ENSG00000143036 / ENSRNOG00000011723
SLC46A1 / Q96NT5 / ENSMUSG00000020829 / ENSG00000076351 / ENSRNOG00000010291
SLC47A1 / Q96FL8 / ENSMUSG00000010122 / ENSG00000142494 / ENSRNOG00000057404
SLC4A1 / P02730 / ENSMUSG00000006574 / ENSG00000004939 / ENSRNOG00000020951
SLC4A2 / P04920 / ENSMUSG00000028962 / ENSG00000164889 / ENSRNOG00000014347
SLC4A4 / Q9Y6R1 / ENSMUSG00000060961 / ENSG00000080493 / ENSRNOG00000003134
SLC4A7 / Q9Y6M7 / ENSMUSG00000021733 / ENSG00000033867 / ENSRNOG00000005957
SLC4A8 / Q2Y0W8 / ENSMUSG00000023032 / ENSG00000050438 / ENSRNOG00000028879
SLC5A1 / P13866 / ENSMUSG00000011034 / ENSG00000100170 / ENSRNOG00000017775
SLC5A10 / A0PJK1 / ENSMUSG00000042371 / ENSG00000154025 / ENSRNOG00000002616
SLC5A2 / P31639 / ENSMUSG00000030781 / ENSG00000140675 / ENSRNOG00000020039
SLC5A6 / Q9Y289 / ENSMUSG00000006641 / ENSG00000138074 / ENSRNOG00000057628
SLC6A1 / P30531 / ENSMUSG00000030310 / ENSG00000157103 / ENSRNOG00000006527
SLC6A11 / P48066 / ENSMUSG00000030307 / ENSG00000132164 / ENSRNOG00000005697
SLC6A12 / P48065 / ENSMUSG00000030109 / ENSG00000111181 / ENSRNOG00000013547
SLC6A13 / Q9NSD5 / ENSMUSG00000030108 / ENSG00000010379 / ENSRNOG00000012876
SLC6A17 / Q9H1V8 / ENSMUSG00000027894 / ENSG00000197106 / ENSRNOG00000050090
SLC6A19 / Q695T7 / ENSMUSG00000021565 / ENSG00000174358 / ENSRNOG00000026501
SLC6A5 / Q9Y345 / ENSMUSG00000039728 / ENSG00000165970 / ENSRNOG00000031662
SLC6A7 / Q99884 / ENSMUSG00000052026 / ENSG00000011083 / ENSRNOG00000018644
SLC6A8 / P48029 / ENSMUSG00000019558 / ENSG00000130821 / ENSRNOG00000057620
SLC7A2 / P52569 / ENSMUSG00000031596 / ENSG00000003989 / ENSRNOG00000011016
SLC7A5 / Q01650 / ENSMUSG00000040010 / ENSG00000103257 / ENSRNOG00000018824
SLC7A6OS / Q96CW6 / ENSMUSG00000033106 / ENSG00000103061 / ENSRNOG00000020049
SLC7A7 / Q9UM01 / ENSMUSG00000000958 / ENSG00000155465 / ENSRNOG00000010296
SLC7A8 / Q9UHI5 / ENSMUSG00000022180 / ENSG00000092068 / ENSRNOG00000014311
SLC8A1 / P32418 / ENSMUSG00000054640 / ENSG00000183023 / ENSRNOG00000008479
SLC8A2 / Q9UPR5 / ENSMUSG00000030376 / ENSG00000118160 / ENSRNOG00000001492
SLC9A1 / P19634 / ENSMUSG00000028854 / ENSG00000090020 / ENSRNOG00000007982
SLC9A3R1 / O14745 / ENSMUSG00000020733 / ENSG00000109062 / ENSRNOG00000003232
SLC9A3R2 / Q15599 / ENSMUSG00000002504 / ENSG00000065054 / ENSRNOG00000002997
SLCO2A1 / Q92959 / ENSMUSG00000032548 / ENSG00000174640 / ENSRNOG00000009005
SLCO2B1 / O94956 / ENSMUSG00000030737 / ENSG00000137491 / ENSRNOG00000017976
SLCO3A1 / Q9UIG8 / ENSMUSG00000025790 / ENSG00000176463 / ENSRNOG00000032798
SLCO4A1 / Q96BD0 / ENSMUSG00000038963 / ENSG00000101187 / ENSRNOG00000053430
SLFN5 / Q08AF3 / ENSMUSG00000054404 / ENSG00000166750 / ENSRNOG00000021719
SLIT2 / O94813 / ENSMUSG00000031558 / ENSG00000145147 / ENSRNOG00000003840
SLIT3 / O75094 / ENSMUSG00000056427 / ENSG00000184347 / ENSRNOG00000007377
SLK / Q9H2G2 / ENSMUSG00000025060 / ENSG00000065613 / ENSRNOG00000011339
Homologs
Brain
Heart
Kidney
Liver
Lung
Testis
Organs
Bins
1
2
3
4
5
Species
Human
Mouse
Rat

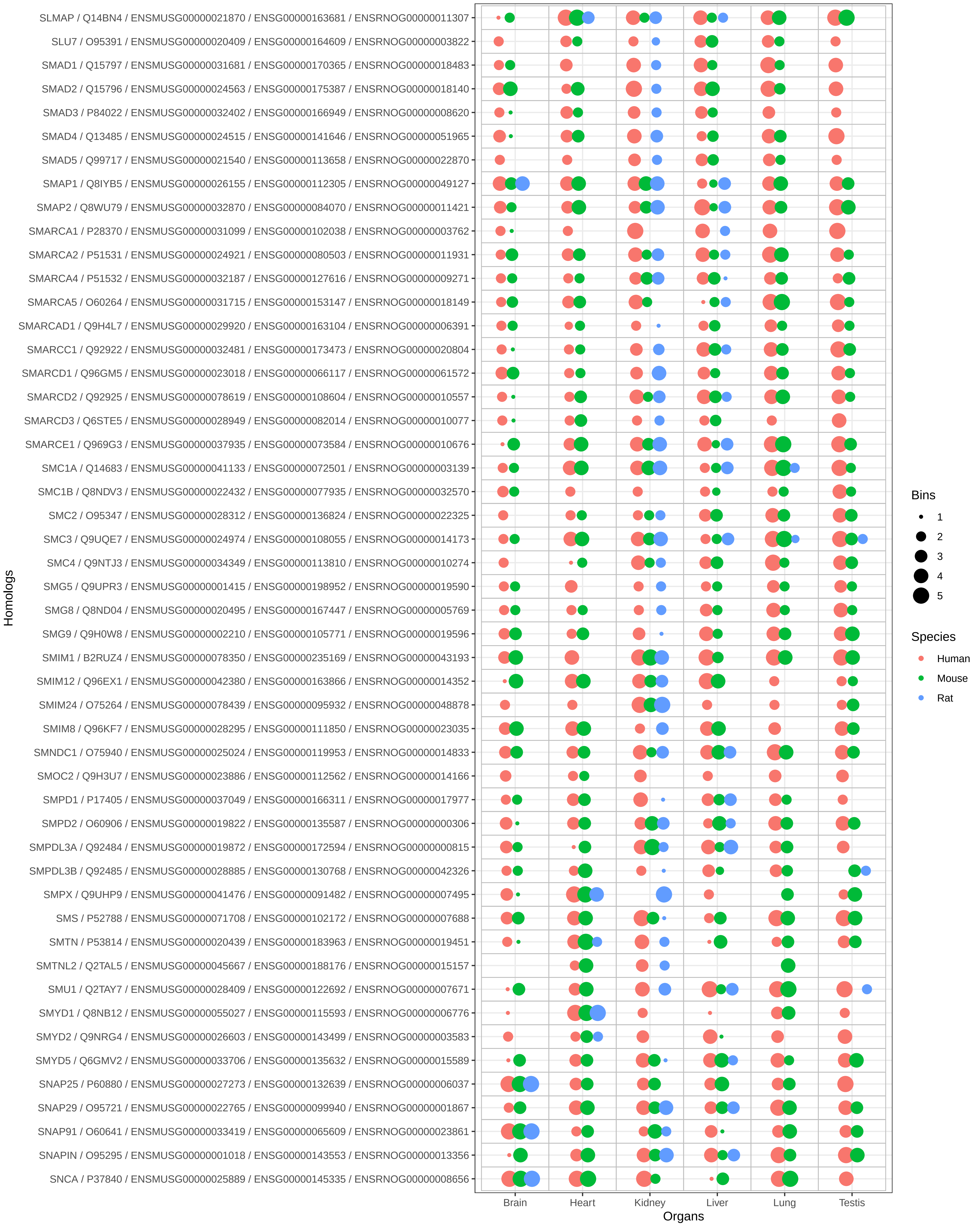
SLMAP / Q14BN4 / ENSMUSG00000021870 / ENSG00000163681 / ENSRNOG00000011307
SLU7 / O95391 / ENSMUSG00000020409 / ENSG00000164609 / ENSRNOG00000003822
SMAD1 / Q15797 / ENSMUSG00000031681 / ENSG00000170365 / ENSRNOG00000018483
SMAD2 / Q15796 / ENSMUSG00000024563 / ENSG00000175387 / ENSRNOG00000018140
SMAD3 / P84022 / ENSMUSG00000032402 / ENSG00000166949 / ENSRNOG00000008620
SMAD4 / Q13485 / ENSMUSG00000024515 / ENSG00000141646 / ENSRNOG00000051965
SMAD5 / Q99717 / ENSMUSG00000021540 / ENSG00000113658 / ENSRNOG00000022870
SMAP1 / Q8IYB5 / ENSMUSG00000026155 / ENSG00000112305 / ENSRNOG00000049127
SMAP2 / Q8WU79 / ENSMUSG00000032870 / ENSG00000084070 / ENSRNOG00000011421
SMARCA1 / P28370 / ENSMUSG00000031099 / ENSG00000102038 / ENSRNOG00000003762
SMARCA2 / P51531 / ENSMUSG00000024921 / ENSG00000080503 / ENSRNOG00000011931
SMARCA4 / P51532 / ENSMUSG00000032187 / ENSG00000127616 / ENSRNOG00000009271
SMARCA5 / O60264 / ENSMUSG00000031715 / ENSG00000153147 / ENSRNOG00000018149
SMARCAD1 / Q9H4L7 / ENSMUSG00000029920 / ENSG00000163104 / ENSRNOG00000006391
SMARCC1 / Q92922 / ENSMUSG00000032481 / ENSG00000173473 / ENSRNOG00000020804
SMARCD1 / Q96GM5 / ENSMUSG00000023018 / ENSG00000066117 / ENSRNOG00000061572
SMARCD2 / Q92925 / ENSMUSG00000078619 / ENSG00000108604 / ENSRNOG00000010557
SMARCD3 / Q6STE5 / ENSMUSG00000028949 / ENSG00000082014 / ENSRNOG00000010077
SMARCE1 / Q969G3 / ENSMUSG00000037935 / ENSG00000073584 / ENSRNOG00000010676
SMC1A / Q14683 / ENSMUSG00000041133 / ENSG00000072501 / ENSRNOG00000003139
SMC1B / Q8NDV3 / ENSMUSG00000022432 / ENSG00000077935 / ENSRNOG00000032570
SMC2 / O95347 / ENSMUSG00000028312 / ENSG00000136824 / ENSRNOG00000022325
SMC3 / Q9UQE7 / ENSMUSG00000024974 / ENSG00000108055 / ENSRNOG00000014173
SMC4 / Q9NTJ3 / ENSMUSG00000034349 / ENSG00000113810 / ENSRNOG00000010274
SMG5 / Q9UPR3 / ENSMUSG00000001415 / ENSG00000198952 / ENSRNOG00000019590
SMG8 / Q8ND04 / ENSMUSG00000020495 / ENSG00000167447 / ENSRNOG00000005769
SMG9 / Q9H0W8 / ENSMUSG00000002210 / ENSG00000105771 / ENSRNOG00000019596
SMIM1 / B2RUZ4 / ENSMUSG00000078350 / ENSG00000235169 / ENSRNOG00000043193
SMIM12 / Q96EX1 / ENSMUSG00000042380 / ENSG00000163866 / ENSRNOG00000014352
SMIM24 / O75264 / ENSMUSG00000078439 / ENSG00000095932 / ENSRNOG00000048878
SMIM8 / Q96KF7 / ENSMUSG00000028295 / ENSG00000111850 / ENSRNOG00000023035
SMNDC1 / O75940 / ENSMUSG00000025024 / ENSG00000119953 / ENSRNOG00000014833
SMOC2 / Q9H3U7 / ENSMUSG00000023886 / ENSG00000112562 / ENSRNOG00000014166
SMPD1 / P17405 / ENSMUSG00000037049 / ENSG00000166311 / ENSRNOG00000017977
SMPD2 / O60906 / ENSMUSG00000019822 / ENSG00000135587 / ENSRNOG00000000306
SMPDL3A / Q92484 / ENSMUSG00000019872 / ENSG00000172594 / ENSRNOG00000000815
SMPDL3B / Q92485 / ENSMUSG00000028885 / ENSG00000130768 / ENSRNOG00000042326
SMPX / Q9UHP9 / ENSMUSG00000041476 / ENSG00000091482 / ENSRNOG00000007495
SMS / P52788 / ENSMUSG00000071708 / ENSG00000102172 / ENSRNOG00000007688
SMTN / P53814 / ENSMUSG00000020439 / ENSG00000183963 / ENSRNOG00000019451
SMTNL2 / Q2TAL5 / ENSMUSG00000045667 / ENSG00000188176 / ENSRNOG00000015157
SMU1 / Q2TAY7 / ENSMUSG00000028409 / ENSG00000122692 / ENSRNOG00000007671
SMYD1 / Q8NB12 / ENSMUSG00000055027 / ENSG00000115593 / ENSRNOG00000006776
SMYD2 / Q9NRG4 / ENSMUSG00000026603 / ENSG00000143499 / ENSRNOG00000003583
SMYD5 / Q6GMV2 / ENSMUSG00000033706 / ENSG00000135632 / ENSRNOG00000015589
SNAP25 / P60880 / ENSMUSG00000027273 / ENSG00000132639 / ENSRNOG00000006037
SNAP29 / O95721 / ENSMUSG00000022765 / ENSG00000099940 / ENSRNOG00000001867
SNAP91 / O60641 / ENSMUSG00000033419 / ENSG00000065609 / ENSRNOG00000023861
SNAPIN / O95295 / ENSMUSG00000001018 / ENSG00000143553 / ENSRNOG00000013356
SNCA / P37840 / ENSMUSG00000025889 / ENSG00000145335 / ENSRNOG00000008656
Homologs
Brain
Heart
Kidney
Liver
Lung
Testis
Organs
Bins
1
2
3
4
5
Species
Human
Mouse
Rat

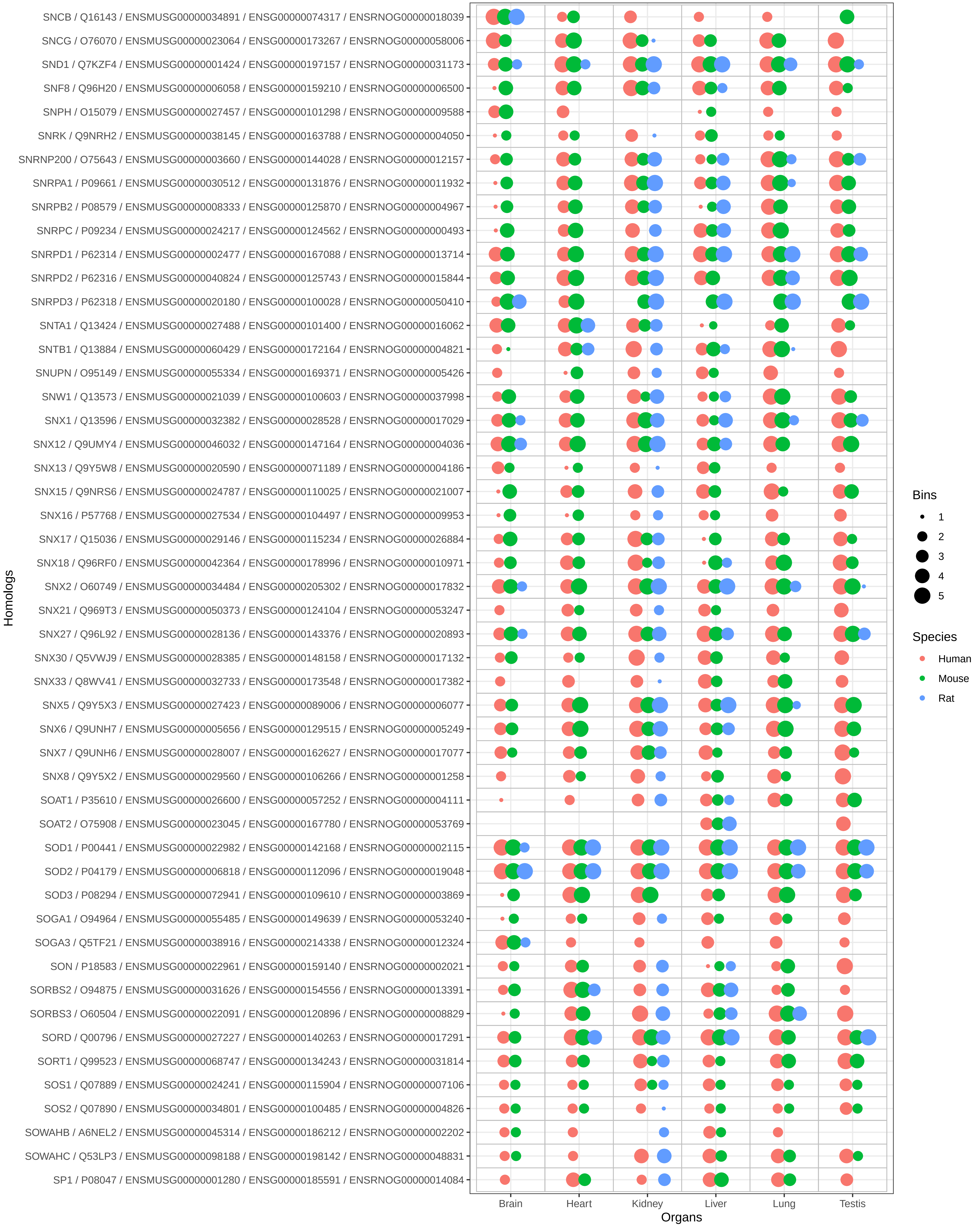
SNCB / Q16143 / ENSMUSG00000034891 / ENSG00000074317 / ENSRNOG00000018039
SNCG / O76070 / ENSMUSG00000023064 / ENSG00000173267 / ENSRNOG00000058006
SND1 / Q7KZF4 / ENSMUSG00000001424 / ENSG00000197157 / ENSRNOG00000031173
SNF8 / Q96H20 / ENSMUSG00000006058 / ENSG00000159210 / ENSRNOG00000006500
SNPH / O15079 / ENSMUSG00000027457 / ENSG00000101298 / ENSRNOG00000009588
SNRK / Q9NRH2 / ENSMUSG00000038145 / ENSG00000163788 / ENSRNOG00000004050
SNRNP200 / O75643 / ENSMUSG00000003660 / ENSG00000144028 / ENSRNOG00000012157
SNRPA1 / P09661 / ENSMUSG00000030512 / ENSG00000131876 / ENSRNOG00000011932
SNRPB2 / P08579 / ENSMUSG00000008333 / ENSG00000125870 / ENSRNOG00000004967
SNRPC / P09234 / ENSMUSG00000024217 / ENSG00000124562 / ENSRNOG00000000493
SNRPD1 / P62314 / ENSMUSG00000002477 / ENSG00000167088 / ENSRNOG00000013714
SNRPD2 / P62316 / ENSMUSG00000040824 / ENSG00000125743 / ENSRNOG00000015844
SNRPD3 / P62318 / ENSMUSG00000020180 / ENSG00000100028 / ENSRNOG00000050410
SNTA1 / Q13424 / ENSMUSG00000027488 / ENSG00000101400 / ENSRNOG00000016062
SNTB1 / Q13884 / ENSMUSG00000060429 / ENSG00000172164 / ENSRNOG00000004821
SNUPN / O95149 / ENSMUSG00000055334 / ENSG00000169371 / ENSRNOG00000005426
SNW1 / Q13573 / ENSMUSG00000021039 / ENSG00000100603 / ENSRNOG00000037998
SNX1 / Q13596 / ENSMUSG00000032382 / ENSG00000028528 / ENSRNOG00000017029
SNX12 / Q9UMY4 / ENSMUSG00000046032 / ENSG00000147164 / ENSRNOG00000004036
SNX13 / Q9Y5W8 / ENSMUSG00000020590 / ENSG00000071189 / ENSRNOG00000004186
SNX15 / Q9NRS6 / ENSMUSG00000024787 / ENSG00000110025 / ENSRNOG00000021007
SNX16 / P57768 / ENSMUSG00000027534 / ENSG00000104497 / ENSRNOG00000009953
SNX17 / Q15036 / ENSMUSG00000029146 / ENSG00000115234 / ENSRNOG00000026884
SNX18 / Q96RF0 / ENSMUSG00000042364 / ENSG00000178996 / ENSRNOG00000010971
SNX2 / O60749 / ENSMUSG00000034484 / ENSG00000205302 / ENSRNOG00000017832
SNX21 / Q969T3 / ENSMUSG00000050373 / ENSG00000124104 / ENSRNOG00000053247
SNX27 / Q96L92 / ENSMUSG00000028136 / ENSG00000143376 / ENSRNOG00000020893
SNX30 / Q5VWJ9 / ENSMUSG00000028385 / ENSG00000148158 / ENSRNOG00000017132
SNX33 / Q8WV41 / ENSMUSG00000032733 / ENSG00000173548 / ENSRNOG00000017382
SNX5 / Q9Y5X3 / ENSMUSG00000027423 / ENSG00000089006 / ENSRNOG00000006077
SNX6 / Q9UNH7 / ENSMUSG00000005656 / ENSG00000129515 / ENSRNOG00000005249
SNX7 / Q9UNH6 / ENSMUSG00000028007 / ENSG00000162627 / ENSRNOG00000017077
SNX8 / Q9Y5X2 / ENSMUSG00000029560 / ENSG00000106266 / ENSRNOG00000001258
SOAT1 / P35610 / ENSMUSG00000026600 / ENSG00000057252 / ENSRNOG00000004111
SOAT2 / O75908 / ENSMUSG00000023045 / ENSG00000167780 / ENSRNOG00000053769
SOD1 / P00441 / ENSMUSG00000022982 / ENSG00000142168 / ENSRNOG00000002115
SOD2 / P04179 / ENSMUSG00000006818 / ENSG00000112096 / ENSRNOG00000019048
SOD3 / P08294 / ENSMUSG00000072941 / ENSG00000109610 / ENSRNOG00000003869
SOGA1 / O94964 / ENSMUSG00000055485 / ENSG00000149639 / ENSRNOG00000053240
SOGA3 / Q5TF21 / ENSMUSG00000038916 / ENSG00000214338 / ENSRNOG00000012324
SON / P18583 / ENSMUSG00000022961 / ENSG00000159140 / ENSRNOG00000002021
SORBS2 / O94875 / ENSMUSG00000031626 / ENSG00000154556 / ENSRNOG00000013391
SORBS3 / O60504 / ENSMUSG00000022091 / ENSG00000120896 / ENSRNOG00000008829
SORD / Q00796 / ENSMUSG00000027227 / ENSG00000140263 / ENSRNOG00000017291
SORT1 / Q99523 / ENSMUSG00000068747 / ENSG00000134243 / ENSRNOG00000031814
SOS1 / Q07889 / ENSMUSG00000024241 / ENSG00000115904 / ENSRNOG00000007106
SOS2 / Q07890 / ENSMUSG00000034801 / ENSG00000100485 / ENSRNOG00000004826
SOWAHB / A6NEL2 / ENSMUSG00000045314 / ENSG00000186212 / ENSRNOG00000002202
SOWAHC / Q53LP3 / ENSMUSG00000098188 / ENSG00000198142 / ENSRNOG00000048831
SP1 / P08047 / ENSMUSG00000001280 / ENSG00000185591 / ENSRNOG00000014084
Homologs
Brain
Heart
Kidney
Liver
Lung
Testis
Organs
Bins
1
2
3
4
5
Species
Human
Mouse
Rat

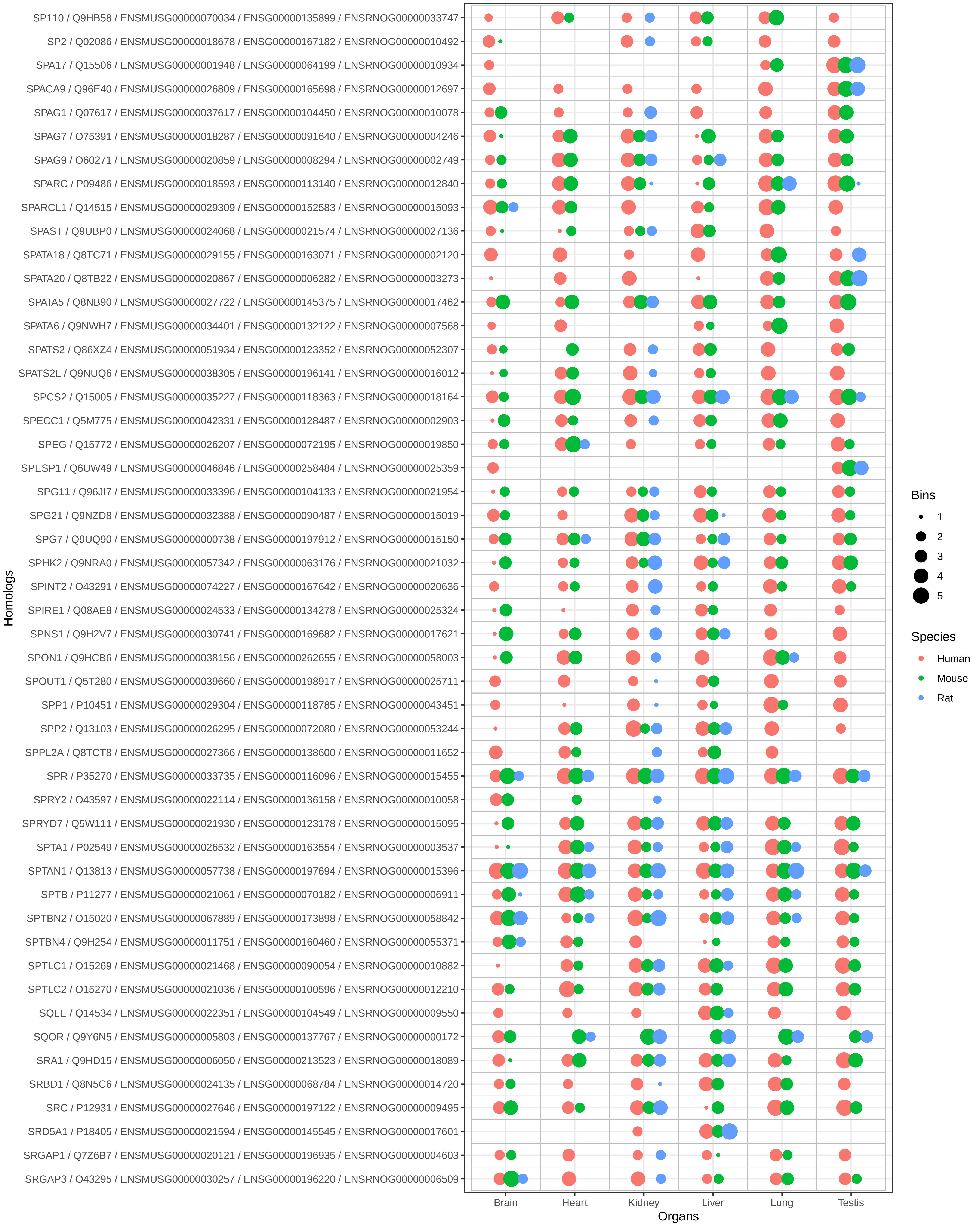
SP110 / Q9HB58 / ENSMUSG00000070034 / ENSG00000135899 / ENSRNOG00000033747
SP2 / Q02086 / ENSMUSG00000018678 / ENSG00000167182 / ENSRNOG00000010492
SPA17 / Q15506 / ENSMUSG00000001948 / ENSG00000064199 / ENSRNOG00000010934
SPACA9 / Q96E40 / ENSMUSG00000026809 / ENSG00000165698 / ENSRNOG00000012697
SPAG1 / Q07617 / ENSMUSG00000037617 / ENSG00000104450 / ENSRNOG00000010078
SPAG7 / O75391 / ENSMUSG00000018287 / ENSG00000091640 / ENSRNOG00000004246
SPAG9 / O60271 / ENSMUSG00000020859 / ENSG00000008294 / ENSRNOG00000002749
SPARC / P09486 / ENSMUSG00000018593 / ENSG00000113140 / ENSRNOG00000012840
SPARCL1 / Q14515 / ENSMUSG00000029309 / ENSG00000152583 / ENSRNOG00000015093
SPAST / Q9UBP0 / ENSMUSG00000024068 / ENSG00000021574 / ENSRNOG00000027136
SPATA18 / Q8TC71 / ENSMUSG00000029155 / ENSG00000163071 / ENSRNOG00000002120
SPATA20 / Q8TB22 / ENSMUSG00000020867 / ENSG00000006282 / ENSRNOG00000003273
SPATA5 / Q8NB90 / ENSMUSG00000027722 / ENSG00000145375 / ENSRNOG00000017462
SPATA6 / Q9NWH7 / ENSMUSG00000034401 / ENSG00000132122 / ENSRNOG00000007568
SPATS2 / Q86XZ4 / ENSMUSG00000051934 / ENSG00000123352 / ENSRNOG00000052307
SPATS2L / Q9NUQ6 / ENSMUSG00000038305 / ENSG00000196141 / ENSRNOG00000016012
SPCS2 / Q15005 / ENSMUSG00000035227 / ENSG00000118363 / ENSRNOG00000018164
SPECC1 / Q5M775 / ENSMUSG00000042331 / ENSG00000128487 / ENSRNOG00000002903
SPEG / Q15772 / ENSMUSG00000026207 / ENSG00000072195 / ENSRNOG00000019850
SPESP1 / Q6UW49 / ENSMUSG00000046846 / ENSG00000258484 / ENSRNOG00000025359
SPG11 / Q96JI7 / ENSMUSG00000033396 / ENSG00000104133 / ENSRNOG00000021954
SPG21 / Q9NZD8 / ENSMUSG00000032388 / ENSG00000090487 / ENSRNOG00000015019
SPG7 / Q9UQ90 / ENSMUSG00000000738 / ENSG00000197912 / ENSRNOG00000015150
SPHK2 / Q9NRA0 / ENSMUSG00000057342 / ENSG00000063176 / ENSRNOG00000021032
SPINT2 / O43291 / ENSMUSG00000074227 / ENSG00000167642 / ENSRNOG00000020636
SPIRE1 / Q08AE8 / ENSMUSG00000024533 / ENSG00000134278 / ENSRNOG00000025324
SPNS1 / Q9H2V7 / ENSMUSG00000030741 / ENSG00000169682 / ENSRNOG00000017621
SPON1 / Q9HCB6 / ENSMUSG00000038156 / ENSG00000262655 / ENSRNOG00000058003
SPOUT1 / Q5T280 / ENSMUSG00000039660 / ENSG00000198917 / ENSRNOG00000025711
SPP1 / P10451 / ENSMUSG00000029304 / ENSG00000118785 / ENSRNOG00000043451
SPP2 / Q13103 / ENSMUSG00000026295 / ENSG00000072080 / ENSRNOG00000053244
SPPL2A / Q8TCT8 / ENSMUSG00000027366 / ENSG00000138600 / ENSRNOG00000011652
SPR / P35270 / ENSMUSG00000033735 / ENSG00000116096 / ENSRNOG00000015455
SPRY2 / O43597 / ENSMUSG00000022114 / ENSG00000136158 / ENSRNOG00000010058
SPRYD7 / Q5W111 / ENSMUSG00000021930 / ENSG00000123178 / ENSRNOG00000015095
SPTA1 / P02549 / ENSMUSG00000026532 / ENSG00000163554 / ENSRNOG00000003537
SPTAN1 / Q13813 / ENSMUSG00000057738 / ENSG00000197694 / ENSRNOG00000015396
SPTB / P11277 / ENSMUSG00000021061 / ENSG00000070182 / ENSRNOG00000006911
SPTBN2 / O15020 / ENSMUSG00000067889 / ENSG00000173898 / ENSRNOG00000058842
SPTBN4 / Q9H254 / ENSMUSG00000011751 / ENSG00000160460 / ENSRNOG00000055371
SPTLC1 / O15269 / ENSMUSG00000021468 / ENSG00000090054 / ENSRNOG00000010882
SPTLC2 / O15270 / ENSMUSG00000021036 / ENSG00000100596 / ENSRNOG00000012210
SQLE / Q14534 / ENSMUSG00000022351 / ENSG00000104549 / ENSRNOG00000009550
SQOR / Q9Y6N5 / ENSMUSG00000005803 / ENSG00000137767 / ENSRNOG00000000172
SRA1 / Q9HD15 / ENSMUSG00000006050 / ENSG00000213523 / ENSRNOG00000018089
SRBD1 / Q8N5C6 / ENSMUSG00000024135 / ENSG00000068784 / ENSRNOG00000014720
SRC / P12931 / ENSMUSG00000027646 / ENSG00000197122 / ENSRNOG00000009495
SRD5A1 / P18405 / ENSMUSG00000021594 / ENSG00000145545 / ENSRNOG00000017601
SRGAP1 / Q7Z6B7 / ENSMUSG00000020121 / ENSG00000196935 / ENSRNOG00000004603
SRGAP3 / O43295 / ENSMUSG00000030257 / ENSG00000196220 / ENSRNOG00000006509
Homologs
Brain
Heart
Kidney
Liver
Lung
Testis
Organs
Bins
1
2
3
4
5
Species
Human
Mouse
Rat

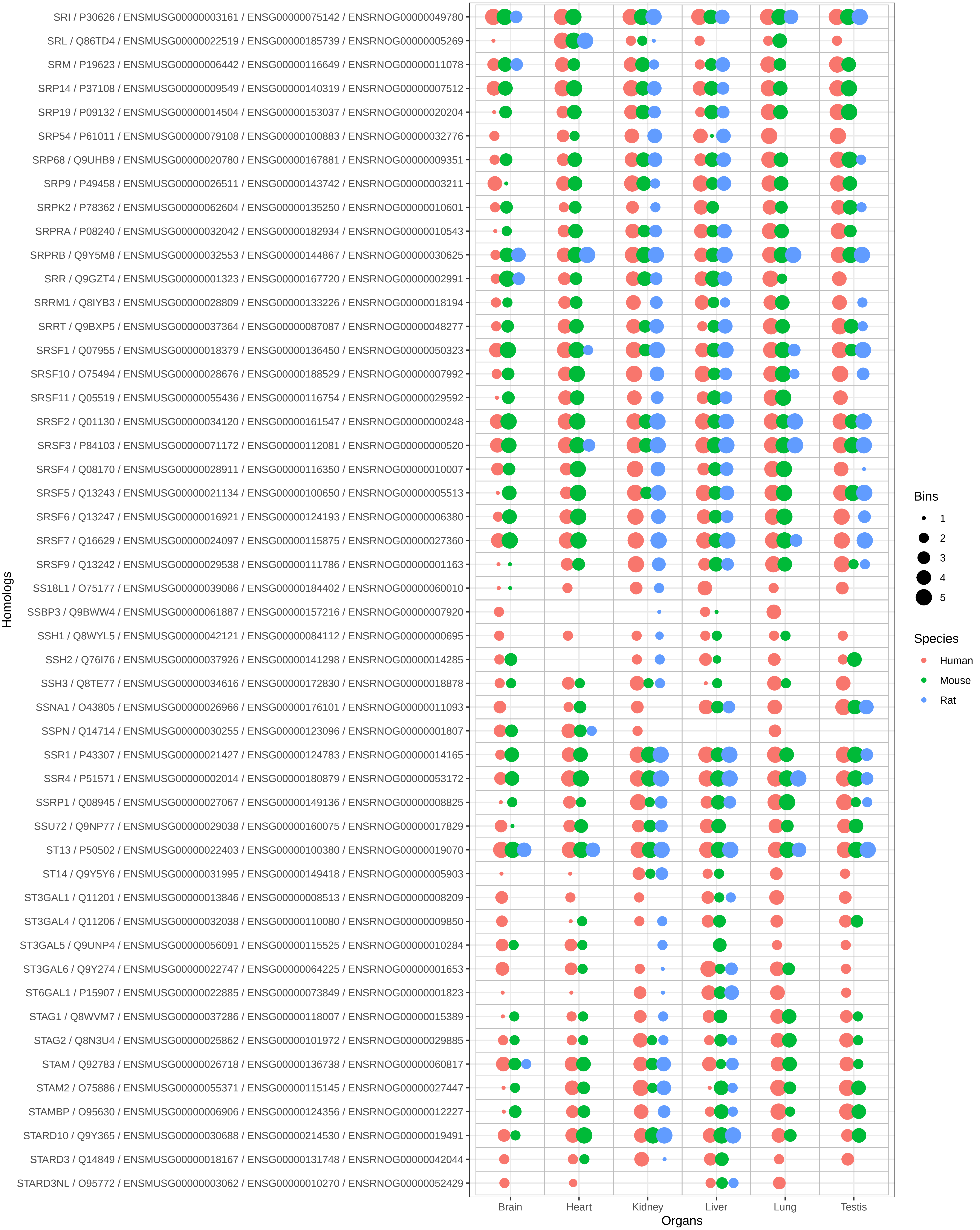
SRI / P30626 / ENSMUSG00000003161 / ENSG00000075142 / ENSRNOG00000049780
SRL / Q86TD4 / ENSMUSG00000022519 / ENSG00000185739 / ENSRNOG00000005269
SRM / P19623 / ENSMUSG00000006442 / ENSG00000116649 / ENSRNOG00000011078
SRP14 / P37108 / ENSMUSG00000009549 / ENSG00000140319 / ENSRNOG00000007512
SRP19 / P09132 / ENSMUSG00000014504 / ENSG00000153037 / ENSRNOG00000020204
SRP54 / P61011 / ENSMUSG00000079108 / ENSG00000100883 / ENSRNOG00000032776
SRP68 / Q9UHB9 / ENSMUSG00000020780 / ENSG00000167881 / ENSRNOG00000009351
SRP9 / P49458 / ENSMUSG00000026511 / ENSG00000143742 / ENSRNOG00000003211
SRPK2 / P78362 / ENSMUSG00000062604 / ENSG00000135250 / ENSRNOG00000010601
SRPRA / P08240 / ENSMUSG00000032042 / ENSG00000182934 / ENSRNOG00000010543
SRPRB / Q9Y5M8 / ENSMUSG00000032553 / ENSG00000144867 / ENSRNOG00000030625
SRR / Q9GZT4 / ENSMUSG00000001323 / ENSG00000167720 / ENSRNOG00000002991
SRRM1 / Q8IYB3 / ENSMUSG00000028809 / ENSG00000133226 / ENSRNOG00000018194
SRRT / Q9BXP5 / ENSMUSG00000037364 / ENSG00000087087 / ENSRNOG00000048277
SRSF1 / Q07955 / ENSMUSG00000018379 / ENSG00000136450 / ENSRNOG00000050323
SRSF10 / O75494 / ENSMUSG00000028676 / ENSG00000188529 / ENSRNOG00000007992
SRSF11 / Q05519 / ENSMUSG00000055436 / ENSG00000116754 / ENSRNOG00000029592
SRSF2 / Q01130 / ENSMUSG00000034120 / ENSG00000161547 / ENSRNOG00000000248
SRSF3 / P84103 / ENSMUSG00000071172 / ENSG00000112081 / ENSRNOG00000000520
SRSF4 / Q08170 / ENSMUSG00000028911 / ENSG00000116350 / ENSRNOG00000010007
SRSF5 / Q13243 / ENSMUSG00000021134 / ENSG00000100650 / ENSRNOG00000005513
SRSF6 / Q13247 / ENSMUSG00000016921 / ENSG00000124193 / ENSRNOG00000006380
SRSF7 / Q16629 / ENSMUSG00000024097 / ENSG00000115875 / ENSRNOG00000027360
SRSF9 / Q13242 / ENSMUSG00000029538 / ENSG00000111786 / ENSRNOG00000001163
SS18L1 / O75177 / ENSMUSG00000039086 / ENSG00000184402 / ENSRNOG00000060010
SSBP3 / Q9BWW4 / ENSMUSG00000061887 / ENSG00000157216 / ENSRNOG00000007920
SSH1 / Q8WYL5 / ENSMUSG00000042121 / ENSG00000084112 / ENSRNOG00000000695
SSH2 / Q76I76 / ENSMUSG00000037926 / ENSG00000141298 / ENSRNOG00000014285
SSH3 / Q8TE77 / ENSMUSG00000034616 / ENSG00000172830 / ENSRNOG00000018878
SSNA1 / O43805 / ENSMUSG00000026966 / ENSG00000176101 / ENSRNOG00000011093
SSPN / Q14714 / ENSMUSG00000030255 / ENSG00000123096 / ENSRNOG00000001807
SSR1 / P43307 / ENSMUSG00000021427 / ENSG00000124783 / ENSRNOG00000014165
SSR4 / P51571 / ENSMUSG00000002014 / ENSG00000180879 / ENSRNOG00000053172
SSRP1 / Q08945 / ENSMUSG00000027067 / ENSG00000149136 / ENSRNOG00000008825
SSU72 / Q9NP77 / ENSMUSG00000029038 / ENSG00000160075 / ENSRNOG00000017829
ST13 / P50502 / ENSMUSG00000022403 / ENSG00000100380 / ENSRNOG00000019070
ST14 / Q9Y5Y6 / ENSMUSG00000031995 / ENSG00000149418 / ENSRNOG00000005903
ST3GAL1 / Q11201 / ENSMUSG00000013846 / ENSG00000008513 / ENSRNOG00000008209
ST3GAL4 / Q11206 / ENSMUSG00000032038 / ENSG00000110080 / ENSRNOG00000009850
ST3GAL5 / Q9UNP4 / ENSMUSG00000056091 / ENSG00000115525 / ENSRNOG00000010284
ST3GAL6 / Q9Y274 / ENSMUSG00000022747 / ENSG00000064225 / ENSRNOG00000001653
ST6GAL1 / P15907 / ENSMUSG00000022885 / ENSG00000073849 / ENSRNOG00000001823
STAG1 / Q8WVM7 / ENSMUSG00000037286 / ENSG00000118007 / ENSRNOG00000015389
STAG2 / Q8N3U4 / ENSMUSG00000025862 / ENSG00000101972 / ENSRNOG00000029885
STAM / Q92783 / ENSMUSG00000026718 / ENSG00000136738 / ENSRNOG00000060817
STAM2 / O75886 / ENSMUSG00000055371 / ENSG00000115145 / ENSRNOG00000027447
STAMBP / O95630 / ENSMUSG00000006906 / ENSG00000124356 / ENSRNOG00000012227
STARD10 / Q9Y365 / ENSMUSG00000030688 / ENSG00000214530 / ENSRNOG00000019491
STARD3 / Q14849 / ENSMUSG00000018167 / ENSG00000131748 / ENSRNOG00000042044
STARD3NL / O95772 / ENSMUSG00000003062 / ENSG00000010270 / ENSRNOG00000052429
Homologs
Brain
Heart
Kidney
Liver
Lung
Testis
Organs
Bins
1
2
3
4
5
Species
Human
Mouse
Rat

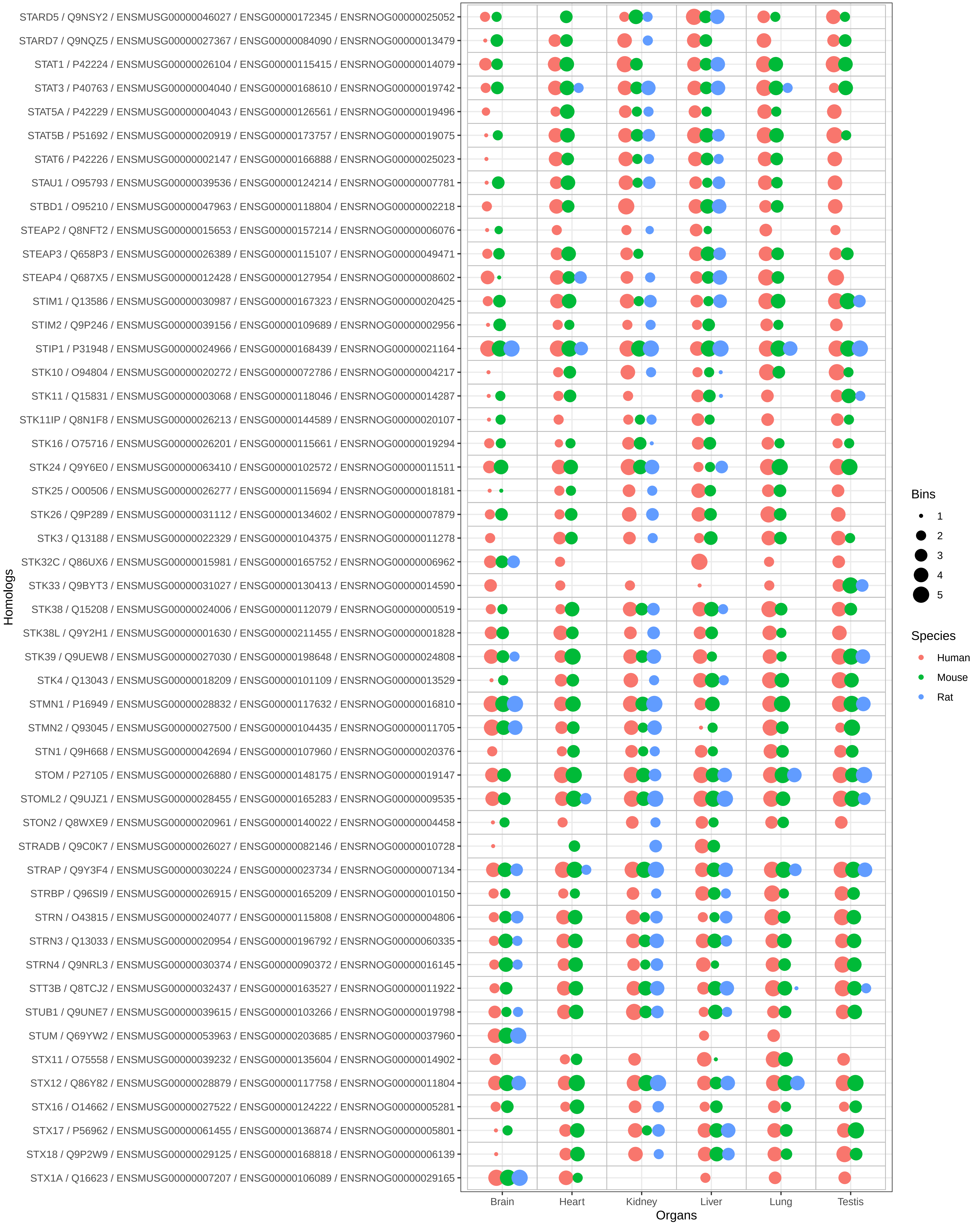
STARD5 / Q9NSY2 / ENSMUSG00000046027 / ENSG00000172345 / ENSRNOG00000025052
STARD7 / Q9NQZ5 / ENSMUSG00000027367 / ENSG00000084090 / ENSRNOG00000013479
STAT1 / P42224 / ENSMUSG00000026104 / ENSG00000115415 / ENSRNOG00000014079
STAT3 / P40763 / ENSMUSG00000004040 / ENSG00000168610 / ENSRNOG00000019742
STAT5A / P42229 / ENSMUSG00000004043 / ENSG00000126561 / ENSRNOG00000019496
STAT5B / P51692 / ENSMUSG00000020919 / ENSG00000173757 / ENSRNOG00000019075
STAT6 / P42226 / ENSMUSG00000002147 / ENSG00000166888 / ENSRNOG00000025023
STAU1 / O95793 / ENSMUSG00000039536 / ENSG00000124214 / ENSRNOG00000007781
STBD1 / O95210 / ENSMUSG00000047963 / ENSG00000118804 / ENSRNOG00000002218
STEAP2 / Q8NFT2 / ENSMUSG00000015653 / ENSG00000157214 / ENSRNOG00000006076
STEAP3 / Q658P3 / ENSMUSG00000026389 / ENSG00000115107 / ENSRNOG00000049471
STEAP4 / Q687X5 / ENSMUSG00000012428 / ENSG00000127954 / ENSRNOG00000008602
STIM1 / Q13586 / ENSMUSG00000030987 / ENSG00000167323 / ENSRNOG00000020425
STIM2 / Q9P246 / ENSMUSG00000039156 / ENSG00000109689 / ENSRNOG00000002956
STIP1 / P31948 / ENSMUSG00000024966 / ENSG00000168439 / ENSRNOG00000021164
STK10 / O94804 / ENSMUSG00000020272 / ENSG00000072786 / ENSRNOG00000004217
STK11 / Q15831 / ENSMUSG00000003068 / ENSG00000118046 / ENSRNOG00000014287
STK11IP / Q8N1F8 / ENSMUSG00000026213 / ENSG00000144589 / ENSRNOG00000020107
STK16 / O75716 / ENSMUSG00000026201 / ENSG00000115661 / ENSRNOG00000019294
STK24 / Q9Y6E0 / ENSMUSG00000063410 / ENSG00000102572 / ENSRNOG00000011511
STK25 / O00506 / ENSMUSG00000026277 / ENSG00000115694 / ENSRNOG00000018181
STK26 / Q9P289 / ENSMUSG00000031112 / ENSG00000134602 / ENSRNOG00000007879
STK3 / Q13188 / ENSMUSG00000022329 / ENSG00000104375 / ENSRNOG00000011278
STK32C / Q86UX6 / ENSMUSG00000015981 / ENSG00000165752 / ENSRNOG00000006962
STK33 / Q9BYT3 / ENSMUSG00000031027 / ENSG00000130413 / ENSRNOG00000014590
STK38 / Q15208 / ENSMUSG00000024006 / ENSG00000112079 / ENSRNOG00000000519
STK38L / Q9Y2H1 / ENSMUSG00000001630 / ENSG00000211455 / ENSRNOG00000001828
STK39 / Q9UEW8 / ENSMUSG00000027030 / ENSG00000198648 / ENSRNOG00000024808
STK4 / Q13043 / ENSMUSG00000018209 / ENSG00000101109 / ENSRNOG00000013529
STMN1 / P16949 / ENSMUSG00000028832 / ENSG00000117632 / ENSRNOG00000016810
STMN2 / Q93045 / ENSMUSG00000027500 / ENSG00000104435 / ENSRNOG00000011705
STN1 / Q9H668 / ENSMUSG00000042694 / ENSG00000107960 / ENSRNOG00000020376
STOM / P27105 / ENSMUSG00000026880 / ENSG00000148175 / ENSRNOG00000019147
STOML2 / Q9UJZ1 / ENSMUSG00000028455 / ENSG00000165283 / ENSRNOG00000009535
STON2 / Q8WXE9 / ENSMUSG00000020961 / ENSG00000140022 / ENSRNOG00000004458
STRADB / Q9C0K7 / ENSMUSG00000026027 / ENSG00000082146 / ENSRNOG00000010728
STRAP / Q9Y3F4 / ENSMUSG00000030224 / ENSG00000023734 / ENSRNOG00000007134
STRBP / Q96SI9 / ENSMUSG00000026915 / ENSG00000165209 / ENSRNOG00000010150
STRN / O43815 / ENSMUSG00000024077 / ENSG00000115808 / ENSRNOG00000004806
STRN3 / Q13033 / ENSMUSG00000020954 / ENSG00000196792 / ENSRNOG00000060335
STRN4 / Q9NRL3 / ENSMUSG00000030374 / ENSG00000090372 / ENSRNOG00000016145
STT3B / Q8TCJ2 / ENSMUSG00000032437 / ENSG00000163527 / ENSRNOG00000011922
STUB1 / Q9UNE7 / ENSMUSG00000039615 / ENSG00000103266 / ENSRNOG00000019798
STUM / Q69YW2 / ENSMUSG00000053963 / ENSG00000203685 / ENSRNOG00000037960
STX11 / O75558 / ENSMUSG00000039232 / ENSG00000135604 / ENSRNOG00000014902
STX12 / Q86Y82 / ENSMUSG00000028879 / ENSG00000117758 / ENSRNOG00000011804
STX16 / O14662 / ENSMUSG00000027522 / ENSG00000124222 / ENSRNOG00000005281
STX17 / P56962 / ENSMUSG00000061455 / ENSG00000136874 / ENSRNOG00000005801
STX18 / Q9P2W9 / ENSMUSG00000029125 / ENSG00000168818 / ENSRNOG00000006139
STX1A / Q16623 / ENSMUSG00000007207 / ENSG00000106089 / ENSRNOG00000029165
Homologs
Brain
Heart
Kidney
Liver
Lung
Testis
Organs
Bins
1
2
3
4
5
Species
Human
Mouse
Rat

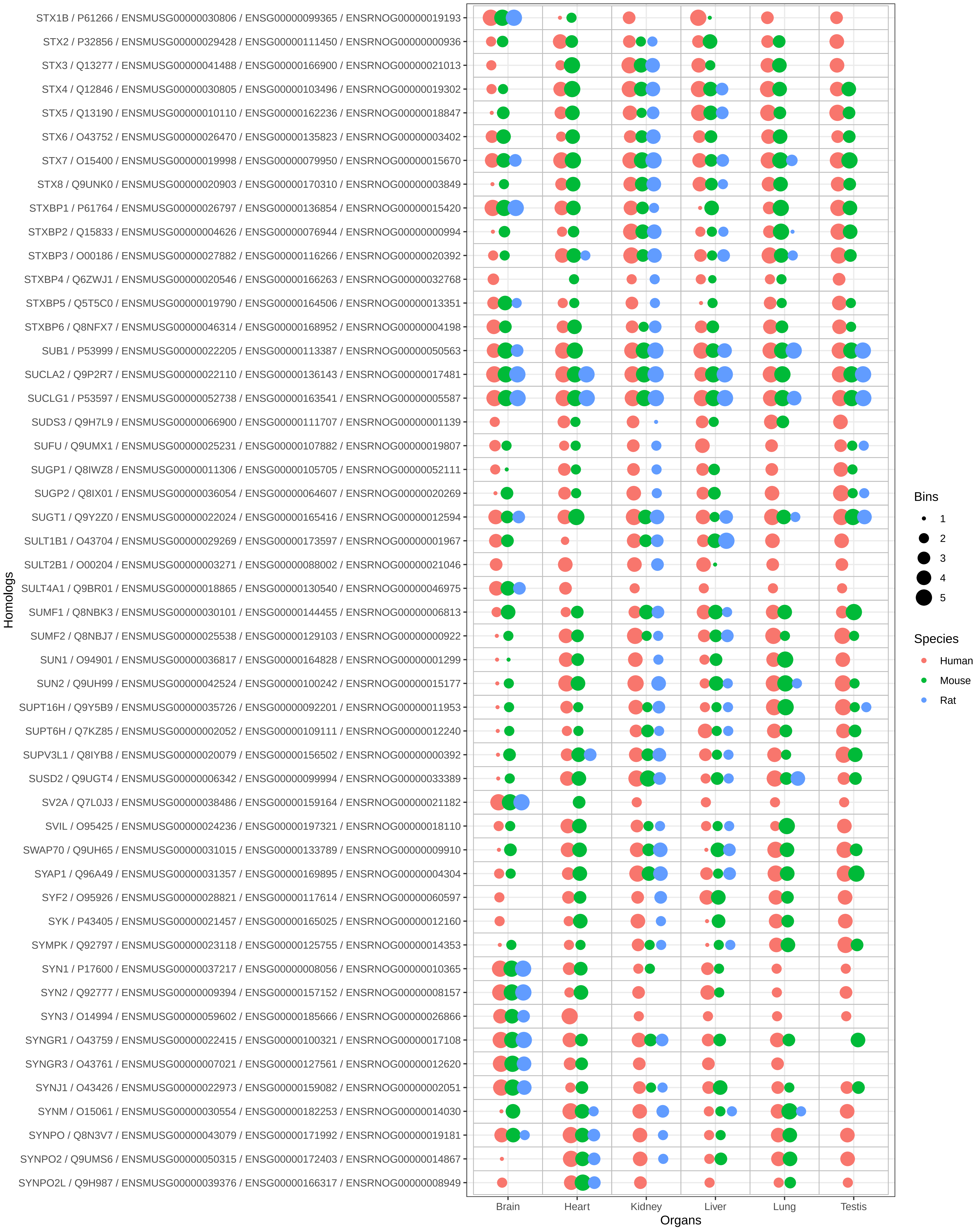

STX1B / P61266 / ENSMUSG00000030806 / ENSG00000099365 / ENSRNOG00000019193
STX2 / P32856 / ENSMUSG00000029428 / ENSG00000111450 / ENSRNOG00000000936
STX3 / Q13277 / ENSMUSG00000041488 / ENSG00000166900 / ENSRNOG00000021013
STX4 / Q12846 / ENSMUSG00000030805 / ENSG00000103496 / ENSRNOG00000019302
STX5 / Q13190 / ENSMUSG00000010110 / ENSG00000162236 / ENSRNOG00000018847
STX6 / O43752 / ENSMUSG00000026470 / ENSG00000135823 / ENSRNOG00000003402
STX7 / O15400 / ENSMUSG00000019998 / ENSG00000079950 / ENSRNOG00000015670
STX8 / Q9UNK0 / ENSMUSG00000020903 / ENSG00000170310 / ENSRNOG00000003849
STXBP1 / P61764 / ENSMUSG00000026797 / ENSG00000136854 / ENSRNOG00000015420
STXBP2 / Q15833 / ENSMUSG00000004626 / ENSG00000076944 / ENSRNOG00000000994
STXBP3 / O00186 / ENSMUSG00000027882 / ENSG00000116266 / ENSRNOG00000020392
STXBP4 / Q6ZWJ1 / ENSMUSG00000020546 / ENSG00000166263 / ENSRNOG00000032768
STXBP5 / Q5T5C0 / ENSMUSG00000019790 / ENSG00000164506 / ENSRNOG00000013351
STXBP6 / Q8NFX7 / ENSMUSG00000046314 / ENSG00000168952 / ENSRNOG00000004198
SUB1 / P53999 / ENSMUSG00000022205 / ENSG00000113387 / ENSRNOG00000050563
SUCLA2 / Q9P2R7 / ENSMUSG00000022110 / ENSG00000136143 / ENSRNOG00000017481
SUCLG1 / P53597 / ENSMUSG00000052738 / ENSG00000163541 / ENSRNOG00000005587
SUDS3 / Q9H7L9 / ENSMUSG00000066900 / ENSG00000111707 / ENSRNOG00000001139
SUFU / Q9UMX1 / ENSMUSG00000025231 / ENSG00000107882 / ENSRNOG00000019807
SUGP1 / Q8IWZ8 / ENSMUSG00000011306 / ENSG00000105705 / ENSRNOG00000052111
SUGP2 / Q8IX01 / ENSMUSG00000036054 / ENSG00000064607 / ENSRNOG00000020269
SUGT1 / Q9Y2Z0 / ENSMUSG00000022024 / ENSG00000165416 / ENSRNOG00000012594
SULT1B1 / O43704 / ENSMUSG00000029269 / ENSG00000173597 / ENSRNOG00000001967
SULT2B1 / O00204 / ENSMUSG00000003271 / ENSG00000088002 / ENSRNOG00000021046
SULT4A1 / Q9BR01 / ENSMUSG00000018865 / ENSG00000130540 / ENSRNOG00000046975
SUMF1 / Q8NBK3 / ENSMUSG00000030101 / ENSG00000144455 / ENSRNOG00000006813
SUMF2 / Q8NBJ7 / ENSMUSG00000025538 / ENSG00000129103 / ENSRNOG00000000922
SUN1 / O94901 / ENSMUSG00000036817 / ENSG00000164828 / ENSRNOG00000001299
SUN2 / Q9UH99 / ENSMUSG00000042524 / ENSG00000100242 / ENSRNOG00000015177
SUPT16H / Q9Y5B9 / ENSMUSG00000035726 / ENSG00000092201 / ENSRNOG00000011953
SUPT6H / Q7KZ85 / ENSMUSG00000002052 / ENSG00000109111 / ENSRNOG00000012240
SUPV3L1 / Q8IYB8 / ENSMUSG00000020079 / ENSG00000156502 / ENSRNOG00000000392
SUSD2 / Q9UGT4 / ENSMUSG00000006342 / ENSG00000099994 / ENSRNOG00000033389
SV2A / Q7L0J3 / ENSMUSG00000038486 / ENSG00000159164 / ENSRNOG00000021182
SVIL / O95425 / ENSMUSG00000024236 / ENSG00000197321 / ENSRNOG00000018110
SWAP70 / Q9UH65 / ENSMUSG00000031015 / ENSG00000133789 / ENSRNOG00000009910
SYAP1 / Q96A49 / ENSMUSG00000031357 / ENSG00000169895 / ENSRNOG00000004304
SYF2 / O95926 / ENSMUSG00000028821 / ENSG00000117614 / ENSRNOG00000060597
SYK / P43405 / ENSMUSG00000021457 / ENSG00000165025 / ENSRNOG00000012160
SYMPK / Q92797 / ENSMUSG00000023118 / ENSG00000125755 / ENSRNOG00000014353
SYN1 / P17600 / ENSMUSG00000037217 / ENSG00000008056 / ENSRNOG00000010365
SYN2 / Q92777 / ENSMUSG00000009394 / ENSG00000157152 / ENSRNOG00000008157
SYN3 / O14994 / ENSMUSG00000059602 / ENSG00000185666 / ENSRNOG00000026866
SYNGR1 / O43759 / ENSMUSG00000022415 / ENSG00000100321 / ENSRNOG00000017108
SYNGR3 / O43761 / ENSMUSG00000007021 / ENSG00000127561 / ENSRNOG00000012620
SYNJ1 / O43426 / ENSMUSG00000022973 / ENSG00000159082 / ENSRNOG00000002051
SYNM / O15061 / ENSMUSG00000030554 / ENSG00000182253 / ENSRNOG00000014030
SYNPO / Q8N3V7 / ENSMUSG00000043079 / ENSG00000171992 / ENSRNOG00000019181
SYNPO2 / Q9UMS6 / ENSMUSG00000050315 / ENSG00000172403 / ENSRNOG00000014867
SYNPO2L / Q9H987 / ENSMUSG00000039376 / ENSG00000166317 / ENSRNOG00000008949
Homologs
Brain
Heart
Kidney
Liver
Lung
Testis
Organs
Bins
1
2
3
4
5
Species
Human
Mouse
Rat

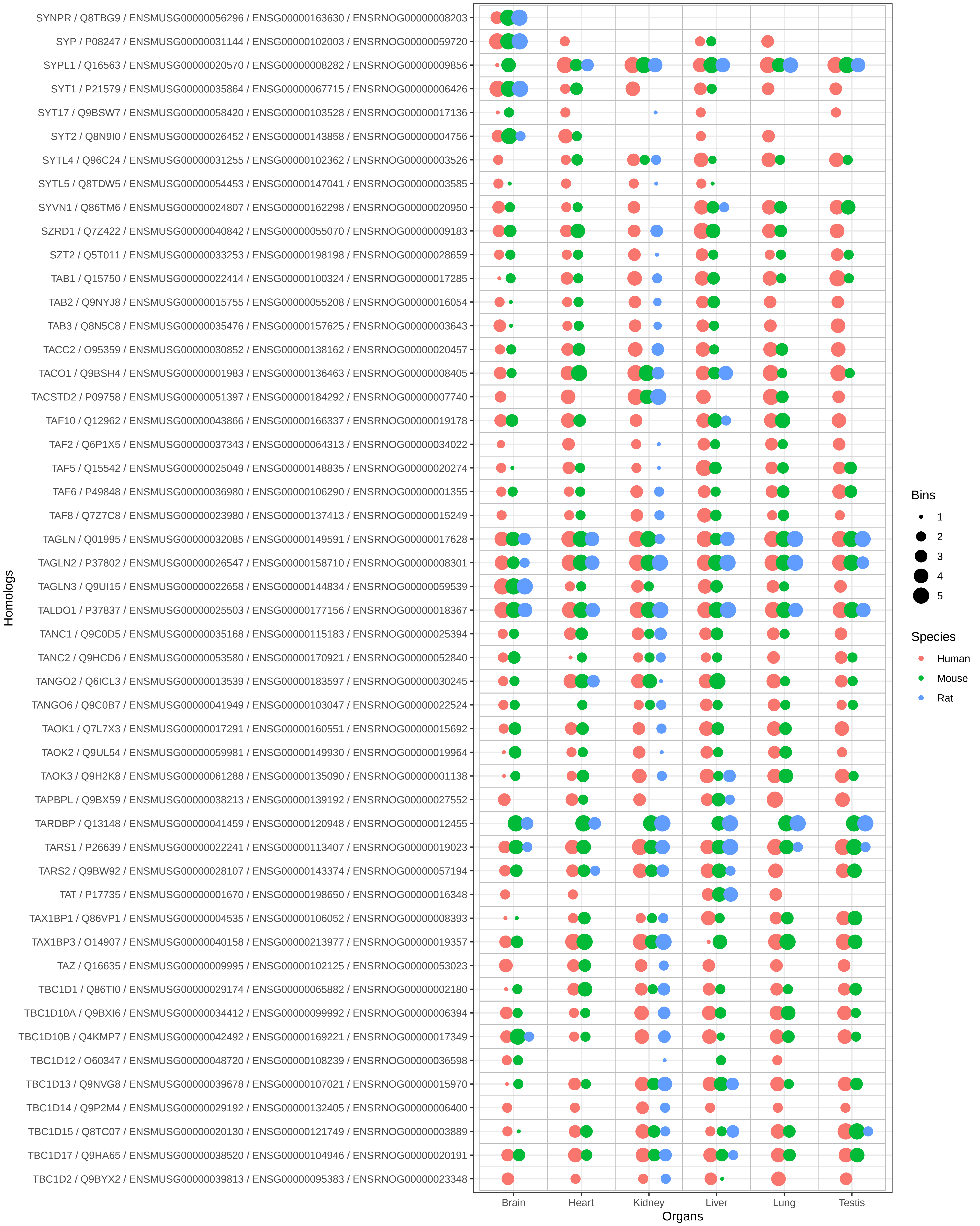
SYNPR / Q8TBG9 / ENSMUSG00000056296 / ENSG00000163630 / ENSRNOG00000008203
SYP / P08247 / ENSMUSG00000031144 / ENSG00000102003 / ENSRNOG00000059720
SYPL1 / Q16563 / ENSMUSG00000020570 / ENSG00000008282 / ENSRNOG00000009856
SYT1 / P21579 / ENSMUSG00000035864 / ENSG00000067715 / ENSRNOG00000006426
SYT17 / Q9BSW7 / ENSMUSG00000058420 / ENSG00000103528 / ENSRNOG00000017136
SYT2 / Q8N9I0 / ENSMUSG00000026452 / ENSG00000143858 / ENSRNOG00000004756
SYTL4 / Q96C24 / ENSMUSG00000031255 / ENSG00000102362 / ENSRNOG00000003526
SYTL5 / Q8TDW5 / ENSMUSG00000054453 / ENSG00000147041 / ENSRNOG00000003585
SYVN1 / Q86TM6 / ENSMUSG00000024807 / ENSG00000162298 / ENSRNOG00000020950
SZRD1 / Q7Z422 / ENSMUSG00000040842 / ENSG00000055070 / ENSRNOG00000009183
SZT2 / Q5T011 / ENSMUSG00000033253 / ENSG00000198198 / ENSRNOG00000028659
TAB1 / Q15750 / ENSMUSG00000022414 / ENSG00000100324 / ENSRNOG00000017285
TAB2 / Q9NYJ8 / ENSMUSG00000015755 / ENSG00000055208 / ENSRNOG00000016054
TAB3 / Q8N5C8 / ENSMUSG00000035476 / ENSG00000157625 / ENSRNOG00000003643
TACC2 / O95359 / ENSMUSG00000030852 / ENSG00000138162 / ENSRNOG00000020457
TACO1 / Q9BSH4 / ENSMUSG00000001983 / ENSG00000136463 / ENSRNOG00000008405
TACSTD2 / P09758 / ENSMUSG00000051397 / ENSG00000184292 / ENSRNOG00000007740
TAF10 / Q12962 / ENSMUSG00000043866 / ENSG00000166337 / ENSRNOG00000019178
TAF2 / Q6P1X5 / ENSMUSG00000037343 / ENSG00000064313 / ENSRNOG00000034022
TAF5 / Q15542 / ENSMUSG00000025049 / ENSG00000148835 / ENSRNOG00000020274
TAF6 / P49848 / ENSMUSG00000036980 / ENSG00000106290 / ENSRNOG00000001355
TAF8 / Q7Z7C8 / ENSMUSG00000023980 / ENSG00000137413 / ENSRNOG00000015249
TAGLN / Q01995 / ENSMUSG00000032085 / ENSG00000149591 / ENSRNOG00000017628
TAGLN2 / P37802 / ENSMUSG00000026547 / ENSG00000158710 / ENSRNOG00000008301
TAGLN3 / Q9UI15 / ENSMUSG00000022658 / ENSG00000144834 / ENSRNOG00000059539
TALDO1 / P37837 / ENSMUSG00000025503 / ENSG00000177156 / ENSRNOG00000018367
TANC1 / Q9C0D5 / ENSMUSG00000035168 / ENSG00000115183 / ENSRNOG00000025394
TANC2 / Q9HCD6 / ENSMUSG00000053580 / ENSG00000170921 / ENSRNOG00000052840
TANGO2 / Q6ICL3 / ENSMUSG00000013539 / ENSG00000183597 / ENSRNOG00000030245
TANGO6 / Q9C0B7 / ENSMUSG00000041949 / ENSG00000103047 / ENSRNOG00000022524
TAOK1 / Q7L7X3 / ENSMUSG00000017291 / ENSG00000160551 / ENSRNOG00000015692
TAOK2 / Q9UL54 / ENSMUSG00000059981 / ENSG00000149930 / ENSRNOG00000019964
TAOK3 / Q9H2K8 / ENSMUSG00000061288 / ENSG00000135090 / ENSRNOG00000001138
TAPBPL / Q9BX59 / ENSMUSG00000038213 / ENSG00000139192 / ENSRNOG00000027552
TARDBP / Q13148 / ENSMUSG00000041459 / ENSG00000120948 / ENSRNOG00000012455
TARS1 / P26639 / ENSMUSG00000022241 / ENSG00000113407 / ENSRNOG00000019023
TARS2 / Q9BW92 / ENSMUSG00000028107 / ENSG00000143374 / ENSRNOG00000057194
TAT / P17735 / ENSMUSG00000001670 / ENSG00000198650 / ENSRNOG00000016348
TAX1BP1 / Q86VP1 / ENSMUSG00000004535 / ENSG00000106052 / ENSRNOG00000008393
TAX1BP3 / O14907 / ENSMUSG00000040158 / ENSG00000213977 / ENSRNOG00000019357
TAZ / Q16635 / ENSMUSG00000009995 / ENSG00000102125 / ENSRNOG00000053023
TBC1D1 / Q86TI0 / ENSMUSG00000029174 / ENSG00000065882 / ENSRNOG00000002180
TBC1D10A / Q9BXI6 / ENSMUSG00000034412 / ENSG00000099992 / ENSRNOG00000006394
TBC1D10B / Q4KMP7 / ENSMUSG00000042492 / ENSG00000169221 / ENSRNOG00000017349
TBC1D12 / O60347 / ENSMUSG00000048720 / ENSG00000108239 / ENSRNOG00000036598
TBC1D13 / Q9NVG8 / ENSMUSG00000039678 / ENSG00000107021 / ENSRNOG00000015970
TBC1D14 / Q9P2M4 / ENSMUSG00000029192 / ENSG00000132405 / ENSRNOG00000006400
TBC1D15 / Q8TC07 / ENSMUSG00000020130 / ENSG00000121749 / ENSRNOG00000003889
TBC1D17 / Q9HA65 / ENSMUSG00000038520 / ENSG00000104946 / ENSRNOG00000020191
TBC1D2 / Q9BYX2 / ENSMUSG00000039813 / ENSG00000095383 / ENSRNOG00000023348
Homologs
Brain
Heart
Kidney
Liver
Lung
Testis
Organs
Bins
1
2
3
4
5
Species
Human
Mouse
Rat

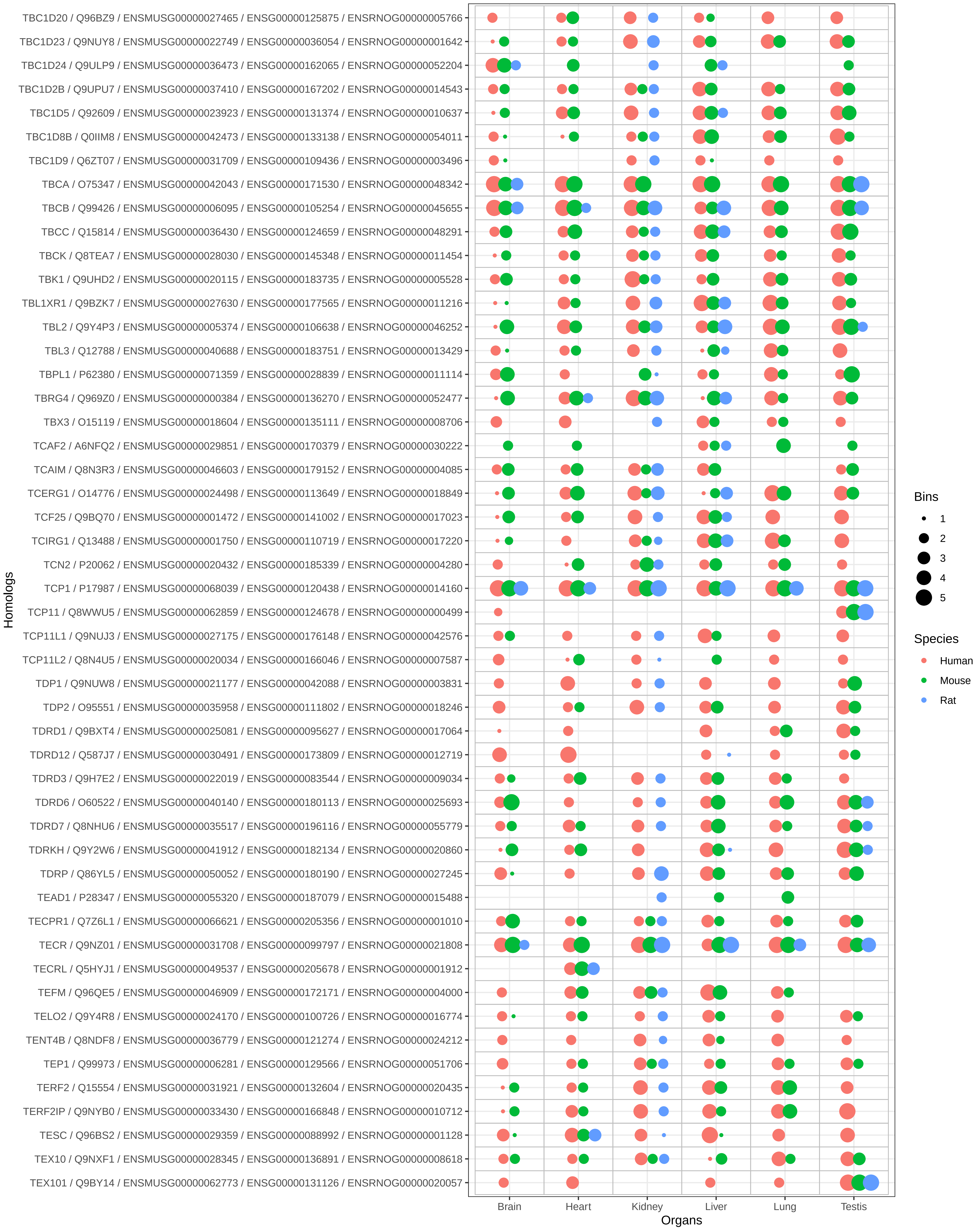
TBC1D20 / Q96BZ9 / ENSMUSG00000027465 / ENSG00000125875 / ENSRNOG00000005766
TBC1D23 / Q9NUY8 / ENSMUSG00000022749 / ENSG00000036054 / ENSRNOG00000001642
TBC1D24 / Q9ULP9 / ENSMUSG00000036473 / ENSG00000162065 / ENSRNOG00000052204
TBC1D2B / Q9UPU7 / ENSMUSG00000037410 / ENSG00000167202 / ENSRNOG00000014543
TBC1D5 / Q92609 / ENSMUSG00000023923 / ENSG00000131374 / ENSRNOG00000010637
TBC1D8B / Q0IIM8 / ENSMUSG00000042473 / ENSG00000133138 / ENSRNOG00000054011
TBC1D9 / Q6ZT07 / ENSMUSG00000031709 / ENSG00000109436 / ENSRNOG00000003496
TBCA / O75347 / ENSMUSG00000042043 / ENSG00000171530 / ENSRNOG00000048342
TBCB / Q99426 / ENSMUSG00000006095 / ENSG00000105254 / ENSRNOG00000045655
TBCC / Q15814 / ENSMUSG00000036430 / ENSG00000124659 / ENSRNOG00000048291
TBCK / Q8TEA7 / ENSMUSG00000028030 / ENSG00000145348 / ENSRNOG00000011454
TBK1 / Q9UHD2 / ENSMUSG00000020115 / ENSG00000183735 / ENSRNOG00000005528
TBL1XR1 / Q9BZK7 / ENSMUSG00000027630 / ENSG00000177565 / ENSRNOG00000011216
TBL2 / Q9Y4P3 / ENSMUSG00000005374 / ENSG00000106638 / ENSRNOG00000046252
TBL3 / Q12788 / ENSMUSG00000040688 / ENSG00000183751 / ENSRNOG00000013429
TBPL1 / P62380 / ENSMUSG00000071359 / ENSG00000028839 / ENSRNOG00000011114
TBRG4 / Q969Z0 / ENSMUSG00000000384 / ENSG00000136270 / ENSRNOG00000052477
TBX3 / O15119 / ENSMUSG00000018604 / ENSG00000135111 / ENSRNOG00000008706
TCAF2 / A6NFQ2 / ENSMUSG00000029851 / ENSG00000170379 / ENSRNOG00000030222
TCAIM / Q8N3R3 / ENSMUSG00000046603 / ENSG00000179152 / ENSRNOG00000004085
TCERG1 / O14776 / ENSMUSG00000024498 / ENSG00000113649 / ENSRNOG00000018849
TCF25 / Q9BQ70 / ENSMUSG00000001472 / ENSG00000141002 / ENSRNOG00000017023
TCIRG1 / Q13488 / ENSMUSG00000001750 / ENSG00000110719 / ENSRNOG00000017220
TCN2 / P20062 / ENSMUSG00000020432 / ENSG00000185339 / ENSRNOG00000004280
TCP1 / P17987 / ENSMUSG00000068039 / ENSG00000120438 / ENSRNOG00000014160
TCP11 / Q8WWU5 / ENSMUSG00000062859 / ENSG00000124678 / ENSRNOG00000000499
TCP11L1 / Q9NUJ3 / ENSMUSG00000027175 / ENSG00000176148 / ENSRNOG00000042576
TCP11L2 / Q8N4U5 / ENSMUSG00000020034 / ENSG00000166046 / ENSRNOG00000007587
TDP1 / Q9NUW8 / ENSMUSG00000021177 / ENSG00000042088 / ENSRNOG00000003831
TDP2 / O95551 / ENSMUSG00000035958 / ENSG00000111802 / ENSRNOG00000018246
TDRD1 / Q9BXT4 / ENSMUSG00000025081 / ENSG00000095627 / ENSRNOG00000017064
TDRD12 / Q587J7 / ENSMUSG00000030491 / ENSG00000173809 / ENSRNOG00000012719
TDRD3 / Q9H7E2 / ENSMUSG00000022019 / ENSG00000083544 / ENSRNOG00000009034
TDRD6 / O60522 / ENSMUSG00000040140 / ENSG00000180113 / ENSRNOG00000025693
TDRD7 / Q8NHU6 / ENSMUSG00000035517 / ENSG00000196116 / ENSRNOG00000055779
TDRKH / Q9Y2W6 / ENSMUSG00000041912 / ENSG00000182134 / ENSRNOG00000020860
TDRP / Q86YL5 / ENSMUSG00000050052 / ENSG00000180190 / ENSRNOG00000027245
TEAD1 / P28347 / ENSMUSG00000055320 / ENSG00000187079 / ENSRNOG00000015488
TECPR1 / Q7Z6L1 / ENSMUSG00000066621 / ENSG00000205356 / ENSRNOG00000001010
TECR / Q9NZ01 / ENSMUSG00000031708 / ENSG00000099797 / ENSRNOG00000021808
TECRL / Q5HYJ1 / ENSMUSG00000049537 / ENSG00000205678 / ENSRNOG00000001912
TEFM / Q96QE5 / ENSMUSG00000046909 / ENSG00000172171 / ENSRNOG00000004000
TELO2 / Q9Y4R8 / ENSMUSG00000024170 / ENSG00000100726 / ENSRNOG00000016774
TENT4B / Q8NDF8 / ENSMUSG00000036779 / ENSG00000121274 / ENSRNOG00000024212
TEP1 / Q99973 / ENSMUSG00000006281 / ENSG00000129566 / ENSRNOG00000051706
TERF2 / Q15554 / ENSMUSG00000031921 / ENSG00000132604 / ENSRNOG00000020435
TERF2IP / Q9NYB0 / ENSMUSG00000033430 / ENSG00000166848 / ENSRNOG00000010712
TESC / Q96BS2 / ENSMUSG00000029359 / ENSG00000088992 / ENSRNOG00000001128
TEX10 / Q9NXF1 / ENSMUSG00000028345 / ENSG00000136891 / ENSRNOG00000008618
TEX101 / Q9BY14 / ENSMUSG00000062773 / ENSG00000131126 / ENSRNOG00000020057
Homologs
Brain
Heart
Kidney
Liver
Lung
Testis
Organs
Bins
1
2
3
4
5
Species
Human
Mouse
Rat

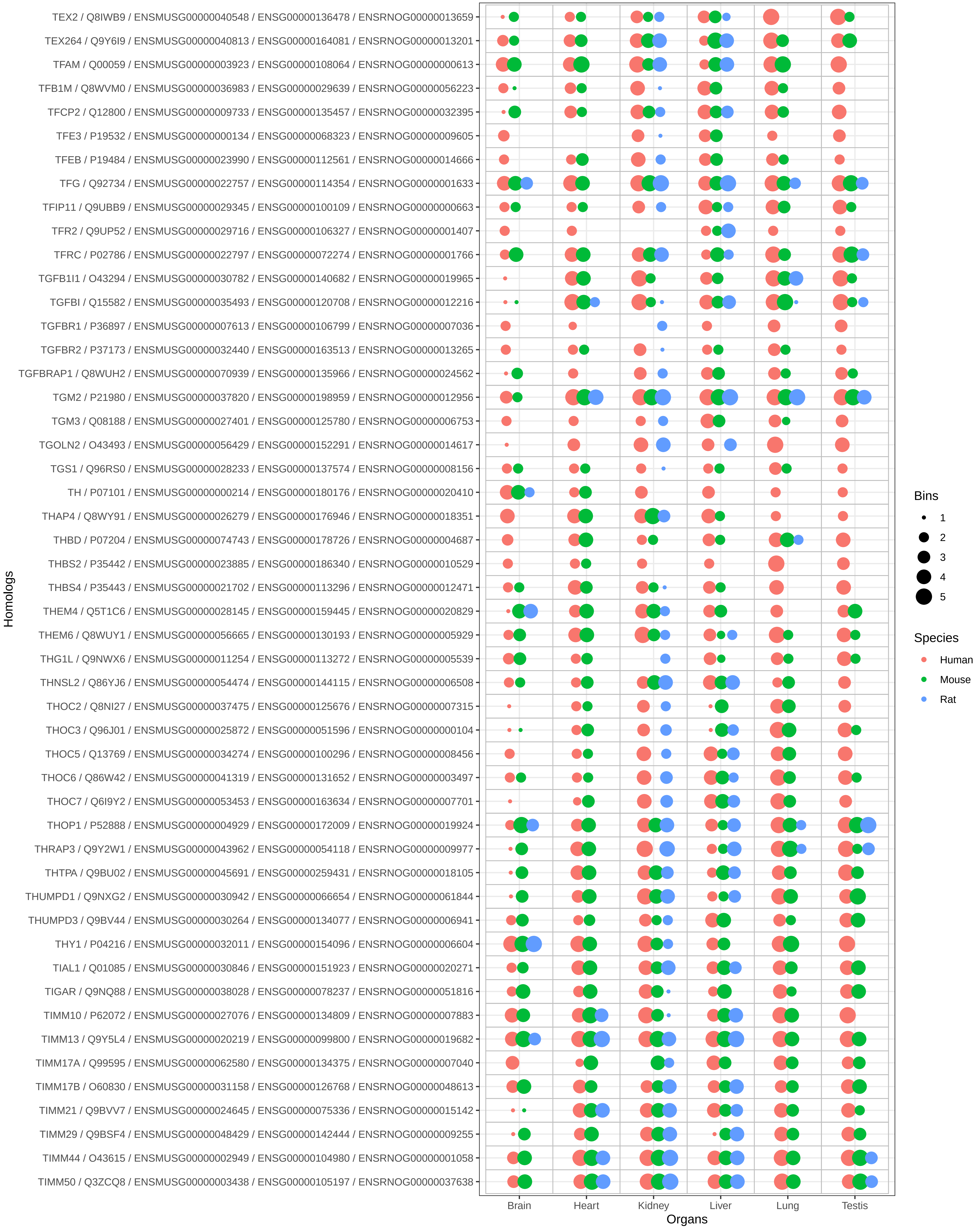

TEX2 / Q8IWB9 / ENSMUSG00000040548 / ENSG00000136478 / ENSRNOG00000013659
TEX264 / Q9Y6I9 / ENSMUSG00000040813 / ENSG00000164081 / ENSRNOG00000013201
TFAM / Q00059 / ENSMUSG00000003923 / ENSG00000108064 / ENSRNOG00000000613
TFB1M / Q8WVM0 / ENSMUSG00000036983 / ENSG00000029639 / ENSRNOG00000056223
TFCP2 / Q12800 / ENSMUSG00000009733 / ENSG00000135457 / ENSRNOG00000032395
TFE3 / P19532 / ENSMUSG00000000134 / ENSG00000068323 / ENSRNOG00000009605
TFEB / P19484 / ENSMUSG00000023990 / ENSG00000112561 / ENSRNOG00000014666
TFG / Q92734 / ENSMUSG00000022757 / ENSG00000114354 / ENSRNOG00000001633
TFIP11 / Q9UBB9 / ENSMUSG00000029345 / ENSG00000100109 / ENSRNOG00000000663
TFR2 / Q9UP52 / ENSMUSG00000029716 / ENSG00000106327 / ENSRNOG00000001407
TFRC / P02786 / ENSMUSG00000022797 / ENSG00000072274 / ENSRNOG00000001766
TGFB1I1 / O43294 / ENSMUSG00000030782 / ENSG00000140682 / ENSRNOG00000019965
TGFBI / Q15582 / ENSMUSG00000035493 / ENSG00000120708 / ENSRNOG00000012216
TGFBR1 / P36897 / ENSMUSG00000007613 / ENSG00000106799 / ENSRNOG00000007036
TGFBR2 / P37173 / ENSMUSG00000032440 / ENSG00000163513 / ENSRNOG00000013265
TGFBRAP1 / Q8WUH2 / ENSMUSG00000070939 / ENSG00000135966 / ENSRNOG00000024562
TGM2 / P21980 / ENSMUSG00000037820 / ENSG00000198959 / ENSRNOG00000012956
TGM3 / Q08188 / ENSMUSG00000027401 / ENSG00000125780 / ENSRNOG00000006753
TGOLN2 / O43493 / ENSMUSG00000056429 / ENSG00000152291 / ENSRNOG00000014617
TGS1 / Q96RS0 / ENSMUSG00000028233 / ENSG00000137574 / ENSRNOG00000008156
TH / P07101 / ENSMUSG00000000214 / ENSG00000180176 / ENSRNOG00000020410
THAP4 / Q8WY91 / ENSMUSG00000026279 / ENSG00000176946 / ENSRNOG00000018351
THBD / P07204 / ENSMUSG00000074743 / ENSG00000178726 / ENSRNOG00000004687
THBS2 / P35442 / ENSMUSG00000023885 / ENSG00000186340 / ENSRNOG00000010529
THBS4 / P35443 / ENSMUSG00000021702 / ENSG00000113296 / ENSRNOG00000012471
THEM4 / Q5T1C6 / ENSMUSG00000028145 / ENSG00000159445 / ENSRNOG00000020829
THEM6 / Q8WUY1 / ENSMUSG00000056665 / ENSG00000130193 / ENSRNOG00000005929
THG1L / Q9NWX6 / ENSMUSG00000011254 / ENSG00000113272 / ENSRNOG00000005539
THNSL2 / Q86YJ6 / ENSMUSG00000054474 / ENSG00000144115 / ENSRNOG00000006508
THOC2 / Q8NI27 / ENSMUSG00000037475 / ENSG00000125676 / ENSRNOG00000007315
THOC3 / Q96J01 / ENSMUSG00000025872 / ENSG00000051596 / ENSRNOG00000000104
THOC5 / Q13769 / ENSMUSG00000034274 / ENSG00000100296 / ENSRNOG00000008456
THOC6 / Q86W42 / ENSMUSG00000041319 / ENSG00000131652 / ENSRNOG00000003497
THOC7 / Q6I9Y2 / ENSMUSG00000053453 / ENSG00000163634 / ENSRNOG00000007701
THOP1 / P52888 / ENSMUSG00000004929 / ENSG00000172009 / ENSRNOG00000019924
THRAP3 / Q9Y2W1 / ENSMUSG00000043962 / ENSG00000054118 / ENSRNOG00000009977
THTPA / Q9BU02 / ENSMUSG00000045691 / ENSG00000259431 / ENSRNOG00000018105
THUMPD1 / Q9NXG2 / ENSMUSG00000030942 / ENSG00000066654 / ENSRNOG00000061844
THUMPD3 / Q9BV44 / ENSMUSG00000030264 / ENSG00000134077 / ENSRNOG00000006941
THY1 / P04216 / ENSMUSG00000032011 / ENSG00000154096 / ENSRNOG00000006604
TIAL1 / Q01085 / ENSMUSG00000030846 / ENSG00000151923 / ENSRNOG00000020271
TIGAR / Q9NQ88 / ENSMUSG00000038028 / ENSG00000078237 / ENSRNOG00000051816
TIMM10 / P62072 / ENSMUSG00000027076 / ENSG00000134809 / ENSRNOG00000007883
TIMM13 / Q9Y5L4 / ENSMUSG00000020219 / ENSG00000099800 / ENSRNOG00000019682
TIMM17A / Q99595 / ENSMUSG00000062580 / ENSG00000134375 / ENSRNOG00000007040
TIMM17B / O60830 / ENSMUSG00000031158 / ENSG00000126768 / ENSRNOG00000048613
TIMM21 / Q9BVV7 / ENSMUSG00000024645 / ENSG00000075336 / ENSRNOG00000015142
TIMM29 / Q9BSF4 / ENSMUSG00000048429 / ENSG00000142444 / ENSRNOG00000009255
TIMM44 / O43615 / ENSMUSG00000002949 / ENSG00000104980 / ENSRNOG00000001058
TIMM50 / Q3ZCQ8 / ENSMUSG00000003438 / ENSG00000105197 / ENSRNOG00000037638
Homologs
Brain
Heart
Kidney
Liver
Lung
Testis
Organs
Bins
1
2
3
4
5
Species
Human
Mouse
Rat

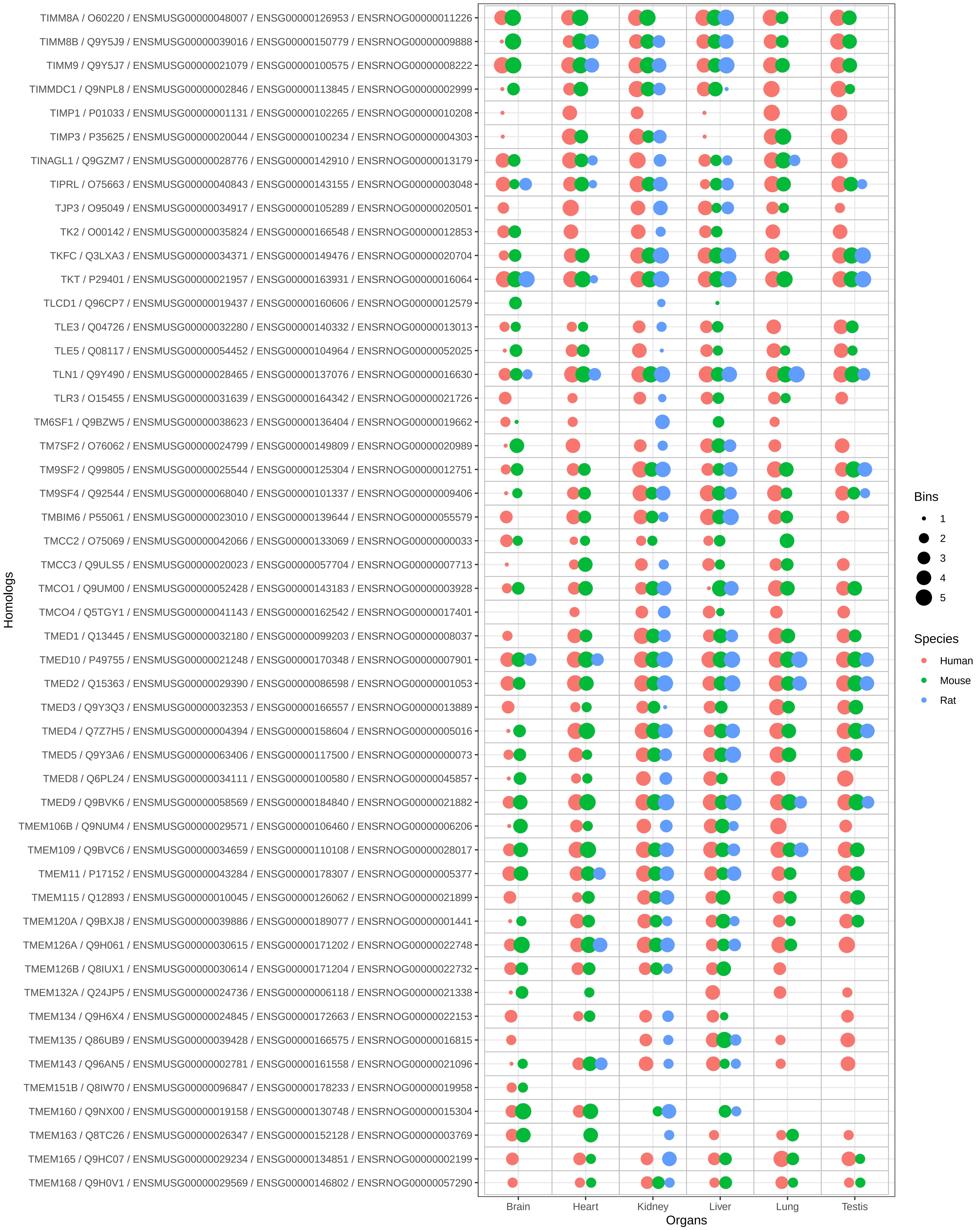
TIMM8A / O60220 / ENSMUSG00000048007 / ENSG00000126953 / ENSRNOG00000011226
TIMM8B / Q9Y5J9 / ENSMUSG00000039016 / ENSG00000150779 / ENSRNOG00000009888
TIMM9 / Q9Y5J7 / ENSMUSG00000021079 / ENSG00000100575 / ENSRNOG00000008222
TIMMDC1 / Q9NPL8 / ENSMUSG00000002846 / ENSG00000113845 / ENSRNOG00000002999
TIMP1 / P01033 / ENSMUSG00000001131 / ENSG00000102265 / ENSRNOG00000010208
TIMP3 / P35625 / ENSMUSG00000020044 / ENSG00000100234 / ENSRNOG00000004303
TINAGL1 / Q9GZM7 / ENSMUSG00000028776 / ENSG00000142910 / ENSRNOG00000013179
TIPRL / O75663 / ENSMUSG00000040843 / ENSG00000143155 / ENSRNOG00000003048
TJP3 / O95049 / ENSMUSG00000034917 / ENSG00000105289 / ENSRNOG00000020501
TK2 / O00142 / ENSMUSG00000035824 / ENSG00000166548 / ENSRNOG00000012853
TKFC / Q3LXA3 / ENSMUSG00000034371 / ENSG00000149476 / ENSRNOG00000020704
TKT / P29401 / ENSMUSG00000021957 / ENSG00000163931 / ENSRNOG00000016064
TLCD1 / Q96CP7 / ENSMUSG00000019437 / ENSG00000160606 / ENSRNOG00000012579
TLE3 / Q04726 / ENSMUSG00000032280 / ENSG00000140332 / ENSRNOG00000013013
TLE5 / Q08117 / ENSMUSG00000054452 / ENSG00000104964 / ENSRNOG00000052025
TLN1 / Q9Y490 / ENSMUSG00000028465 / ENSG00000137076 / ENSRNOG00000016630
TLR3 / O15455 / ENSMUSG00000031639 / ENSG00000164342 / ENSRNOG00000021726
TM6SF1 / Q9BZW5 / ENSMUSG00000038623 / ENSG00000136404 / ENSRNOG00000019662
TM7SF2 / O76062 / ENSMUSG00000024799 / ENSG00000149809 / ENSRNOG00000020989
TM9SF2 / Q99805 / ENSMUSG00000025544 / ENSG00000125304 / ENSRNOG00000012751
TM9SF4 / Q92544 / ENSMUSG00000068040 / ENSG00000101337 / ENSRNOG00000009406
TMBIM6 / P55061 / ENSMUSG00000023010 / ENSG00000139644 / ENSRNOG00000055579
TMCC2 / O75069 / ENSMUSG00000042066 / ENSG00000133069 / ENSRNOG00000000033
TMCC3 / Q9ULS5 / ENSMUSG00000020023 / ENSG00000057704 / ENSRNOG00000007713
TMCO1 / Q9UM00 / ENSMUSG00000052428 / ENSG00000143183 / ENSRNOG00000003928
TMCO4 / Q5TGY1 / ENSMUSG00000041143 / ENSG00000162542 / ENSRNOG00000017401
TMED1 / Q13445 / ENSMUSG00000032180 / ENSG00000099203 / ENSRNOG00000008037
TMED10 / P49755 / ENSMUSG00000021248 / ENSG00000170348 / ENSRNOG00000007901
TMED2 / Q15363 / ENSMUSG00000029390 / ENSG00000086598 / ENSRNOG00000001053
TMED3 / Q9Y3Q3 / ENSMUSG00000032353 / ENSG00000166557 / ENSRNOG00000013889
TMED4 / Q7Z7H5 / ENSMUSG00000004394 / ENSG00000158604 / ENSRNOG00000005016
TMED5 / Q9Y3A6 / ENSMUSG00000063406 / ENSG00000117500 / ENSRNOG00000000073
TMED8 / Q6PL24 / ENSMUSG00000034111 / ENSG00000100580 / ENSRNOG00000045857
TMED9 / Q9BVK6 / ENSMUSG00000058569 / ENSG00000184840 / ENSRNOG00000021882
TMEM106B / Q9NUM4 / ENSMUSG00000029571 / ENSG00000106460 / ENSRNOG00000006206
TMEM109 / Q9BVC6 / ENSMUSG00000034659 / ENSG00000110108 / ENSRNOG00000028017
TMEM11 / P17152 / ENSMUSG00000043284 / ENSG00000178307 / ENSRNOG00000005377
TMEM115 / Q12893 / ENSMUSG00000010045 / ENSG00000126062 / ENSRNOG00000021899
TMEM120A / Q9BXJ8 / ENSMUSG00000039886 / ENSG00000189077 / ENSRNOG00000001441
TMEM126A / Q9H061 / ENSMUSG00000030615 / ENSG00000171202 / ENSRNOG00000022748
TMEM126B / Q8IUX1 / ENSMUSG00000030614 / ENSG00000171204 / ENSRNOG00000022732
TMEM132A / Q24JP5 / ENSMUSG00000024736 / ENSG00000006118 / ENSRNOG00000021338
TMEM134 / Q9H6X4 / ENSMUSG00000024845 / ENSG00000172663 / ENSRNOG00000022153
TMEM135 / Q86UB9 / ENSMUSG00000039428 / ENSG00000166575 / ENSRNOG00000016815
TMEM143 / Q96AN5 / ENSMUSG00000002781 / ENSG00000161558 / ENSRNOG00000021096
TMEM151B / Q8IW70 / ENSMUSG00000096847 / ENSG00000178233 / ENSRNOG00000019958
TMEM160 / Q9NX00 / ENSMUSG00000019158 / ENSG00000130748 / ENSRNOG00000015304
TMEM163 / Q8TC26 / ENSMUSG00000026347 / ENSG00000152128 / ENSRNOG00000003769
TMEM165 / Q9HC07 / ENSMUSG00000029234 / ENSG00000134851 / ENSRNOG00000002199
TMEM168 / Q9H0V1 / ENSMUSG00000029569 / ENSG00000146802 / ENSRNOG00000057290
Homologs
Brain
Heart
Kidney
Liver
Lung
Testis
Organs
Bins
1
2
3
4
5
Species
Human
Mouse
Rat

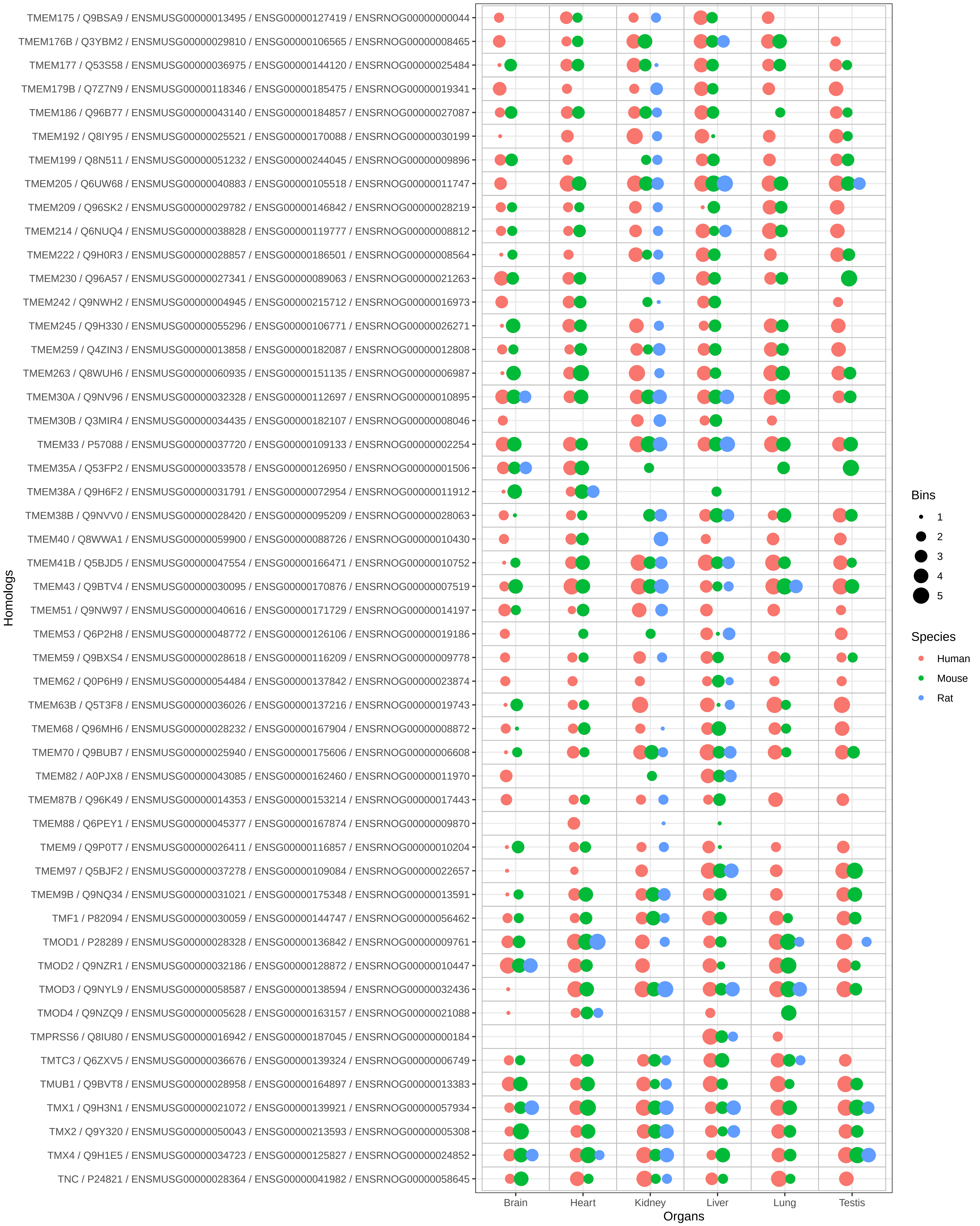
TMEM175 / Q9BSA9 / ENSMUSG00000013495 / ENSG00000127419 / ENSRNOG00000000044
TMEM176B / Q3YBM2 / ENSMUSG00000029810 / ENSG00000106565 / ENSRNOG00000008465
TMEM177 / Q53S58 / ENSMUSG00000036975 / ENSG00000144120 / ENSRNOG00000025484
TMEM179B / Q7Z7N9 / ENSMUSG00000118346 / ENSG00000185475 / ENSRNOG00000019341
TMEM186 / Q96B77 / ENSMUSG00000043140 / ENSG00000184857 / ENSRNOG00000027087
TMEM192 / Q8IY95 / ENSMUSG00000025521 / ENSG00000170088 / ENSRNOG00000030199
TMEM199 / Q8N511 / ENSMUSG00000051232 / ENSG00000244045 / ENSRNOG00000009896
TMEM205 / Q6UW68 / ENSMUSG00000040883 / ENSG00000105518 / ENSRNOG00000011747
TMEM209 / Q96SK2 / ENSMUSG00000029782 / ENSG00000146842 / ENSRNOG00000028219
TMEM214 / Q6NUQ4 / ENSMUSG00000038828 / ENSG00000119777 / ENSRNOG00000008812
TMEM222 / Q9H0R3 / ENSMUSG00000028857 / ENSG00000186501 / ENSRNOG00000008564
TMEM230 / Q96A57 / ENSMUSG00000027341 / ENSG00000089063 / ENSRNOG00000021263
TMEM242 / Q9NWH2 / ENSMUSG00000004945 / ENSG00000215712 / ENSRNOG00000016973
TMEM245 / Q9H330 / ENSMUSG00000055296 / ENSG00000106771 / ENSRNOG00000026271
TMEM259 / Q4ZIN3 / ENSMUSG00000013858 / ENSG00000182087 / ENSRNOG00000012808
TMEM263 / Q8WUH6 / ENSMUSG00000060935 / ENSG00000151135 / ENSRNOG00000006987
TMEM30A / Q9NV96 / ENSMUSG00000032328 / ENSG00000112697 / ENSRNOG00000010895
TMEM30B / Q3MIR4 / ENSMUSG00000034435 / ENSG00000182107 / ENSRNOG00000008046
TMEM33 / P57088 / ENSMUSG00000037720 / ENSG00000109133 / ENSRNOG00000002254
TMEM35A / Q53FP2 / ENSMUSG00000033578 / ENSG00000126950 / ENSRNOG00000001506
TMEM38A / Q9H6F2 / ENSMUSG00000031791 / ENSG00000072954 / ENSRNOG00000011912
TMEM38B / Q9NVV0 / ENSMUSG00000028420 / ENSG00000095209 / ENSRNOG00000028063
TMEM40 / Q8WWA1 / ENSMUSG00000059900 / ENSG00000088726 / ENSRNOG00000010430
TMEM41B / Q5BJD5 / ENSMUSG00000047554 / ENSG00000166471 / ENSRNOG00000010752
TMEM43 / Q9BTV4 / ENSMUSG00000030095 / ENSG00000170876 / ENSRNOG00000007519
TMEM51 / Q9NW97 / ENSMUSG00000040616 / ENSG00000171729 / ENSRNOG00000014197
TMEM53 / Q6P2H8 / ENSMUSG00000048772 / ENSG00000126106 / ENSRNOG00000019186
TMEM59 / Q9BXS4 / ENSMUSG00000028618 / ENSG00000116209 / ENSRNOG00000009778
TMEM62 / Q0P6H9 / ENSMUSG00000054484 / ENSG00000137842 / ENSRNOG00000023874
TMEM63B / Q5T3F8 / ENSMUSG00000036026 / ENSG00000137216 / ENSRNOG00000019743
TMEM68 / Q96MH6 / ENSMUSG00000028232 / ENSG00000167904 / ENSRNOG00000008872
TMEM70 / Q9BUB7 / ENSMUSG00000025940 / ENSG00000175606 / ENSRNOG00000006608
TMEM82 / A0PJX8 / ENSMUSG00000043085 / ENSG00000162460 / ENSRNOG00000011970
TMEM87B / Q96K49 / ENSMUSG00000014353 / ENSG00000153214 / ENSRNOG00000017443
TMEM88 / Q6PEY1 / ENSMUSG00000045377 / ENSG00000167874 / ENSRNOG00000009870
TMEM9 / Q9P0T7 / ENSMUSG00000026411 / ENSG00000116857 / ENSRNOG00000010204
TMEM97 / Q5BJF2 / ENSMUSG00000037278 / ENSG00000109084 / ENSRNOG00000022657
TMEM9B / Q9NQ34 / ENSMUSG00000031021 / ENSG00000175348 / ENSRNOG00000013591
TMF1 / P82094 / ENSMUSG00000030059 / ENSG00000144747 / ENSRNOG00000056462
TMOD1 / P28289 / ENSMUSG00000028328 / ENSG00000136842 / ENSRNOG00000009761
TMOD2 / Q9NZR1 / ENSMUSG00000032186 / ENSG00000128872 / ENSRNOG00000010447
TMOD3 / Q9NYL9 / ENSMUSG00000058587 / ENSG00000138594 / ENSRNOG00000032436
TMOD4 / Q9NZQ9 / ENSMUSG00000005628 / ENSG00000163157 / ENSRNOG00000021088
TMPRSS6 / Q8IU80 / ENSMUSG00000016942 / ENSG00000187045 / ENSRNOG00000000184
TMTC3 / Q6ZXV5 / ENSMUSG00000036676 / ENSG00000139324 / ENSRNOG00000006749
TMUB1 / Q9BVT8 / ENSMUSG00000028958 / ENSG00000164897 / ENSRNOG00000013383
TMX1 / Q9H3N1 / ENSMUSG00000021072 / ENSG00000139921 / ENSRNOG00000057934
TMX2 / Q9Y320 / ENSMUSG00000050043 / ENSG00000213593 / ENSRNOG00000005308
TMX4 / Q9H1E5 / ENSMUSG00000034723 / ENSG00000125827 / ENSRNOG00000024852
TNC / P24821 / ENSMUSG00000028364 / ENSG00000041982 / ENSRNOG00000058645
Homologs
Brain
Heart
Kidney
Liver
Lung
Testis
Organs
Bins
1
2
3
4
5
Species
Human
Mouse
Rat

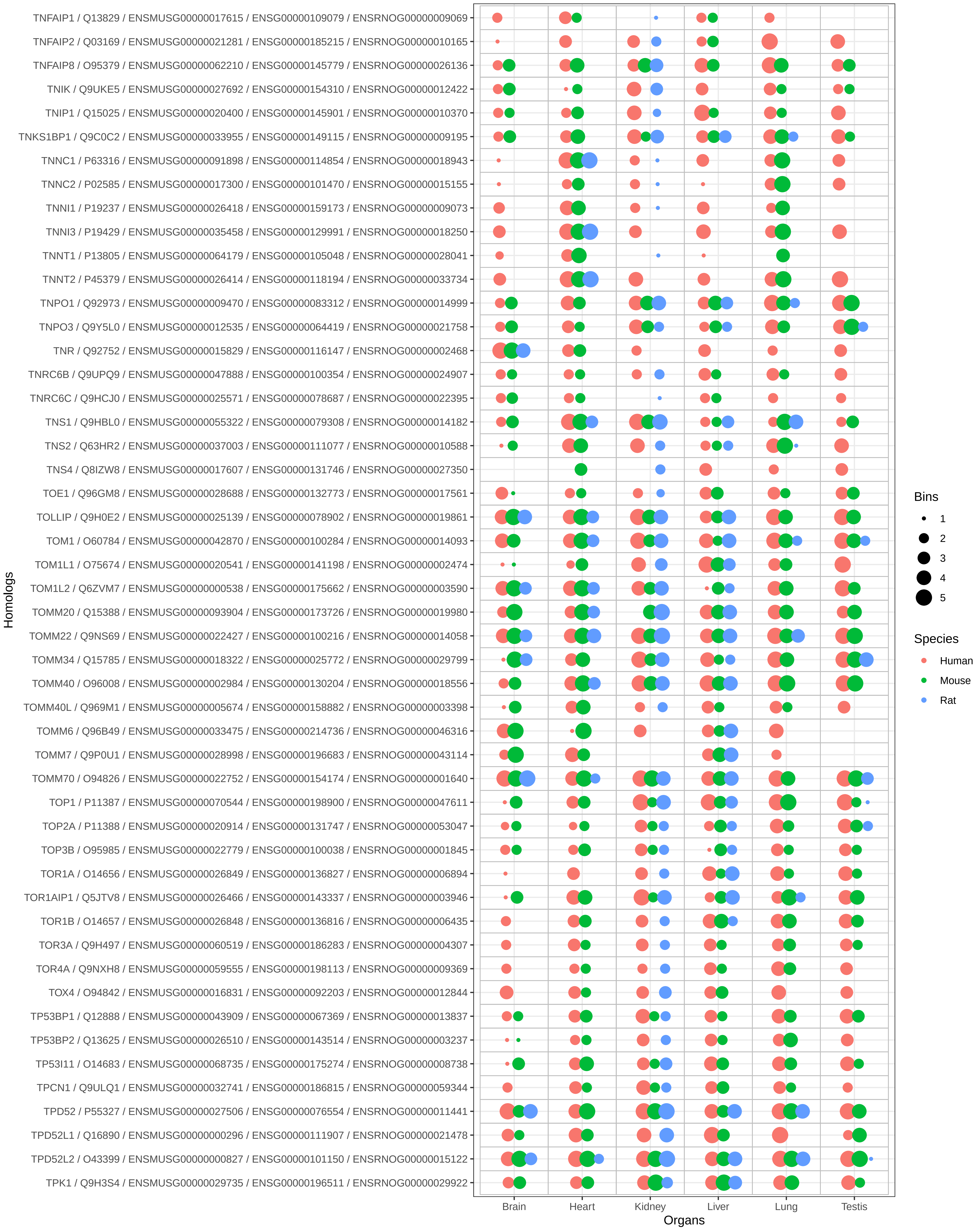
TNFAIP1 / Q13829 / ENSMUSG00000017615 / ENSG00000109079 / ENSRNOG00000009069
TNFAIP2 / Q03169 / ENSMUSG00000021281 / ENSG00000185215 / ENSRNOG00000010165
TNFAIP8 / O95379 / ENSMUSG00000062210 / ENSG00000145779 / ENSRNOG00000026136
TNIK / Q9UKE5 / ENSMUSG00000027692 / ENSG00000154310 / ENSRNOG00000012422
TNIP1 / Q15025 / ENSMUSG00000020400 / ENSG00000145901 / ENSRNOG00000010370
TNKS1BP1 / Q9C0C2 / ENSMUSG00000033955 / ENSG00000149115 / ENSRNOG00000009195
TNNC1 / P63316 / ENSMUSG00000091898 / ENSG00000114854 / ENSRNOG00000018943
TNNC2 / P02585 / ENSMUSG00000017300 / ENSG00000101470 / ENSRNOG00000015155
TNNI1 / P19237 / ENSMUSG00000026418 / ENSG00000159173 / ENSRNOG00000009073
TNNI3 / P19429 / ENSMUSG00000035458 / ENSG00000129991 / ENSRNOG00000018250
TNNT1 / P13805 / ENSMUSG00000064179 / ENSG00000105048 / ENSRNOG00000028041
TNNT2 / P45379 / ENSMUSG00000026414 / ENSG00000118194 / ENSRNOG00000033734
TNPO1 / Q92973 / ENSMUSG00000009470 / ENSG00000083312 / ENSRNOG00000014999
TNPO3 / Q9Y5L0 / ENSMUSG00000012535 / ENSG00000064419 / ENSRNOG00000021758
TNR / Q92752 / ENSMUSG00000015829 / ENSG00000116147 / ENSRNOG00000002468
TNRC6B / Q9UPQ9 / ENSMUSG00000047888 / ENSG00000100354 / ENSRNOG00000024907
TNRC6C / Q9HCJ0 / ENSMUSG00000025571 / ENSG00000078687 / ENSRNOG00000022395
TNS1 / Q9HBL0 / ENSMUSG00000055322 / ENSG00000079308 / ENSRNOG00000014182
TNS2 / Q63HR2 / ENSMUSG00000037003 / ENSG00000111077 / ENSRNOG00000010588
TNS4 / Q8IZW8 / ENSMUSG00000017607 / ENSG00000131746 / ENSRNOG00000027350
TOE1 / Q96GM8 / ENSMUSG00000028688 / ENSG00000132773 / ENSRNOG00000017561
TOLLIP / Q9H0E2 / ENSMUSG00000025139 / ENSG00000078902 / ENSRNOG00000019861
TOM1 / O60784 / ENSMUSG00000042870 / ENSG00000100284 / ENSRNOG00000014093
TOM1L1 / O75674 / ENSMUSG00000020541 / ENSG00000141198 / ENSRNOG00000002474
TOM1L2 / Q6ZVM7 / ENSMUSG00000000538 / ENSG00000175662 / ENSRNOG00000003590
TOMM20 / Q15388 / ENSMUSG00000093904 / ENSG00000173726 / ENSRNOG00000019980
TOMM22 / Q9NS69 / ENSMUSG00000022427 / ENSG00000100216 / ENSRNOG00000014058
TOMM34 / Q15785 / ENSMUSG00000018322 / ENSG00000025772 / ENSRNOG00000029799
TOMM40 / O96008 / ENSMUSG00000002984 / ENSG00000130204 / ENSRNOG00000018556
TOMM40L / Q969M1 / ENSMUSG00000005674 / ENSG00000158882 / ENSRNOG00000003398
TOMM6 / Q96B49 / ENSMUSG00000033475 / ENSG00000214736 / ENSRNOG00000046316
TOMM7 / Q9P0U1 / ENSMUSG00000028998 / ENSG00000196683 / ENSRNOG00000043114
TOMM70 / O94826 / ENSMUSG00000022752 / ENSG00000154174 / ENSRNOG00000001640
TOP1 / P11387 / ENSMUSG00000070544 / ENSG00000198900 / ENSRNOG00000047611
TOP2A / P11388 / ENSMUSG00000020914 / ENSG00000131747 / ENSRNOG00000053047
TOP3B / O95985 / ENSMUSG00000022779 / ENSG00000100038 / ENSRNOG00000001845
TOR1A / O14656 / ENSMUSG00000026849 / ENSG00000136827 / ENSRNOG00000006894
TOR1AIP1 / Q5JTV8 / ENSMUSG00000026466 / ENSG00000143337 / ENSRNOG00000003946
TOR1B / O14657 / ENSMUSG00000026848 / ENSG00000136816 / ENSRNOG00000006435
TOR3A / Q9H497 / ENSMUSG00000060519 / ENSG00000186283 / ENSRNOG00000004307
TOR4A / Q9NXH8 / ENSMUSG00000059555 / ENSG00000198113 / ENSRNOG00000009369
TOX4 / O94842 / ENSMUSG00000016831 / ENSG00000092203 / ENSRNOG00000012844
TP53BP1 / Q12888 / ENSMUSG00000043909 / ENSG00000067369 / ENSRNOG00000013837
TP53BP2 / Q13625 / ENSMUSG00000026510 / ENSG00000143514 / ENSRNOG00000003237
TP53I11 / O14683 / ENSMUSG00000068735 / ENSG00000175274 / ENSRNOG00000008738
TPCN1 / Q9ULQ1 / ENSMUSG00000032741 / ENSG00000186815 / ENSRNOG00000059344
TPD52 / P55327 / ENSMUSG00000027506 / ENSG00000076554 / ENSRNOG00000011441
TPD52L1 / Q16890 / ENSMUSG00000000296 / ENSG00000111907 / ENSRNOG00000021478
TPD52L2 / O43399 / ENSMUSG00000000827 / ENSG00000101150 / ENSRNOG00000015122
TPK1 / Q9H3S4 / ENSMUSG00000029735 / ENSG00000196511 / ENSRNOG00000029922
Homologs
Brain
Heart
Kidney
Liver
Lung
Testis
Organs
Bins
1
2
3
4
5
Species
Human
Mouse
Rat

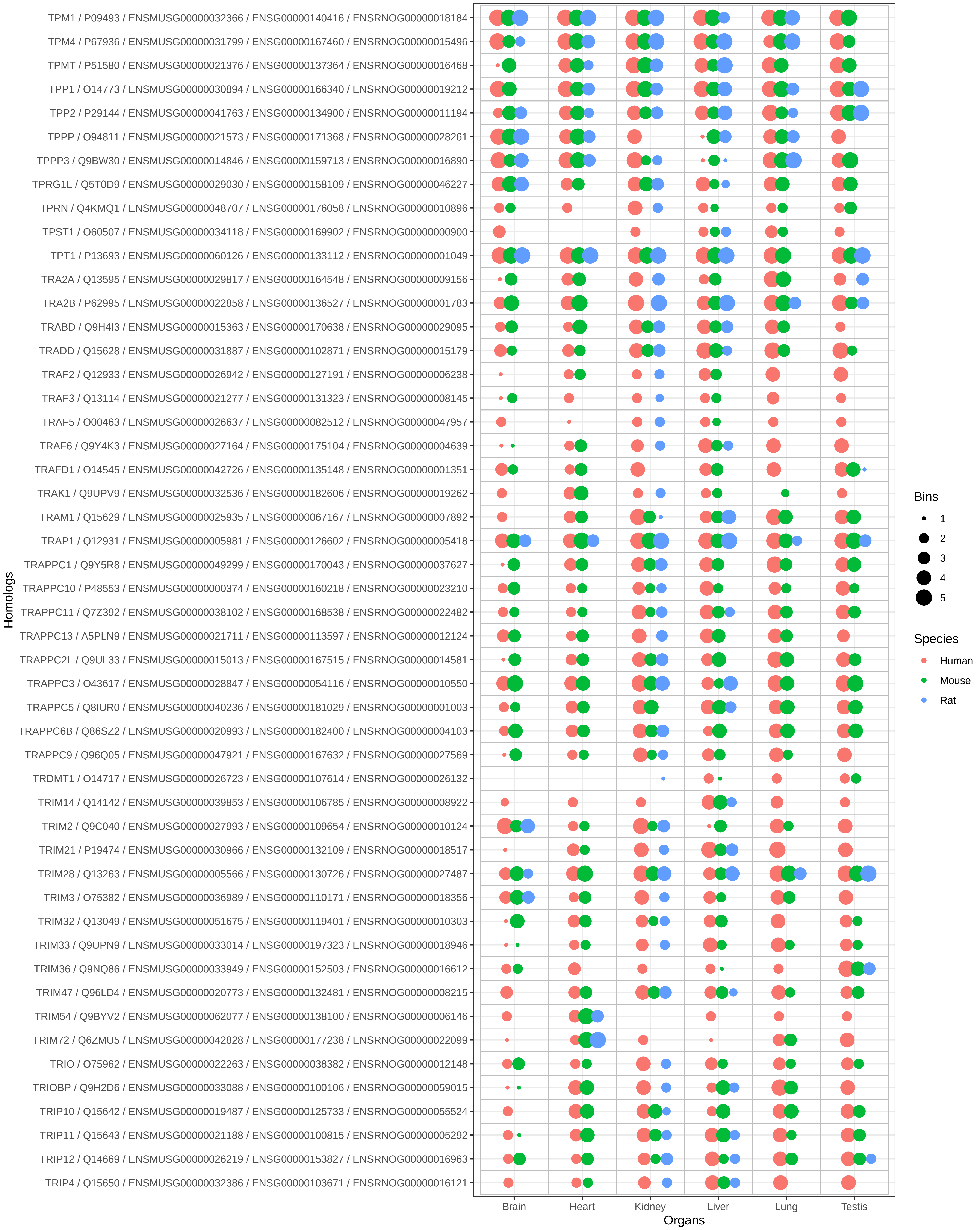
TPM1 / P09493 / ENSMUSG00000032366 / ENSG00000140416 / ENSRNOG00000018184
TPM4 / P67936 / ENSMUSG00000031799 / ENSG00000167460 / ENSRNOG00000015496
TPMT / P51580 / ENSMUSG00000021376 / ENSG00000137364 / ENSRNOG00000016468
TPP1 / O14773 / ENSMUSG00000030894 / ENSG00000166340 / ENSRNOG00000019212
TPP2 / P29144 / ENSMUSG00000041763 / ENSG00000134900 / ENSRNOG00000011194
TPPP / O94811 / ENSMUSG00000021573 / ENSG00000171368 / ENSRNOG00000028261
TPPP3 / Q9BW30 / ENSMUSG00000014846 / ENSG00000159713 / ENSRNOG00000016890
TPRG1L / Q5T0D9 / ENSMUSG00000029030 / ENSG00000158109 / ENSRNOG00000046227
TPRN / Q4KMQ1 / ENSMUSG00000048707 / ENSG00000176058 / ENSRNOG00000010896
TPST1 / O60507 / ENSMUSG00000034118 / ENSG00000169902 / ENSRNOG00000000900
TPT1 / P13693 / ENSMUSG00000060126 / ENSG00000133112 / ENSRNOG00000001049
TRA2A / Q13595 / ENSMUSG00000029817 / ENSG00000164548 / ENSRNOG00000009156
TRA2B / P62995 / ENSMUSG00000022858 / ENSG00000136527 / ENSRNOG00000001783
TRABD / Q9H4I3 / ENSMUSG00000015363 / ENSG00000170638 / ENSRNOG00000029095
TRADD / Q15628 / ENSMUSG00000031887 / ENSG00000102871 / ENSRNOG00000015179
TRAF2 / Q12933 / ENSMUSG00000026942 / ENSG00000127191 / ENSRNOG00000006238
TRAF3 / Q13114 / ENSMUSG00000021277 / ENSG00000131323 / ENSRNOG00000008145
TRAF5 / O00463 / ENSMUSG00000026637 / ENSG00000082512 / ENSRNOG00000047957
TRAF6 / Q9Y4K3 / ENSMUSG00000027164 / ENSG00000175104 / ENSRNOG00000004639
TRAFD1 / O14545 / ENSMUSG00000042726 / ENSG00000135148 / ENSRNOG00000001351
TRAK1 / Q9UPV9 / ENSMUSG00000032536 / ENSG00000182606 / ENSRNOG00000019262
TRAM1 / Q15629 / ENSMUSG00000025935 / ENSG00000067167 / ENSRNOG00000007892
TRAP1 / Q12931 / ENSMUSG00000005981 / ENSG00000126602 / ENSRNOG00000005418
TRAPPC1 / Q9Y5R8 / ENSMUSG00000049299 / ENSG00000170043 / ENSRNOG00000037627
TRAPPC10 / P48553 / ENSMUSG00000000374 / ENSG00000160218 / ENSRNOG00000023210
TRAPPC11 / Q7Z392 / ENSMUSG00000038102 / ENSG00000168538 / ENSRNOG00000022482
TRAPPC13 / A5PLN9 / ENSMUSG00000021711 / ENSG00000113597 / ENSRNOG00000012124
TRAPPC2L / Q9UL33 / ENSMUSG00000015013 / ENSG00000167515 / ENSRNOG00000014581
TRAPPC3 / O43617 / ENSMUSG00000028847 / ENSG00000054116 / ENSRNOG00000010550
TRAPPC5 / Q8IUR0 / ENSMUSG00000040236 / ENSG00000181029 / ENSRNOG00000001003
TRAPPC6B / Q86SZ2 / ENSMUSG00000020993 / ENSG00000182400 / ENSRNOG00000004103
TRAPPC9 / Q96Q05 / ENSMUSG00000047921 / ENSG00000167632 / ENSRNOG00000027569
TRDMT1 / O14717 / ENSMUSG00000026723 / ENSG00000107614 / ENSRNOG00000026132
TRIM14 / Q14142 / ENSMUSG00000039853 / ENSG00000106785 / ENSRNOG00000008922
TRIM2 / Q9C040 / ENSMUSG00000027993 / ENSG00000109654 / ENSRNOG00000010124
TRIM21 / P19474 / ENSMUSG00000030966 / ENSG00000132109 / ENSRNOG00000018517
TRIM28 / Q13263 / ENSMUSG00000005566 / ENSG00000130726 / ENSRNOG00000027487
TRIM3 / O75382 / ENSMUSG00000036989 / ENSG00000110171 / ENSRNOG00000018356
TRIM32 / Q13049 / ENSMUSG00000051675 / ENSG00000119401 / ENSRNOG00000010303
TRIM33 / Q9UPN9 / ENSMUSG00000033014 / ENSG00000197323 / ENSRNOG00000018946
TRIM36 / Q9NQ86 / ENSMUSG00000033949 / ENSG00000152503 / ENSRNOG00000016612
TRIM47 / Q96LD4 / ENSMUSG00000020773 / ENSG00000132481 / ENSRNOG00000008215
TRIM54 / Q9BYV2 / ENSMUSG00000062077 / ENSG00000138100 / ENSRNOG00000006146
TRIM72 / Q6ZMU5 / ENSMUSG00000042828 / ENSG00000177238 / ENSRNOG00000022099
TRIO / O75962 / ENSMUSG00000022263 / ENSG00000038382 / ENSRNOG00000012148
TRIOBP / Q9H2D6 / ENSMUSG00000033088 / ENSG00000100106 / ENSRNOG00000059015
TRIP10 / Q15642 / ENSMUSG00000019487 / ENSG00000125733 / ENSRNOG00000055524
TRIP11 / Q15643 / ENSMUSG00000021188 / ENSG00000100815 / ENSRNOG00000005292
TRIP12 / Q14669 / ENSMUSG00000026219 / ENSG00000153827 / ENSRNOG00000016963
TRIP4 / Q15650 / ENSMUSG00000032386 / ENSG00000103671 / ENSRNOG00000016121
Homologs
Brain
Heart
Kidney
Liver
Lung
Testis
Organs
Bins
1
2
3
4
5
Species
Human
Mouse
Rat

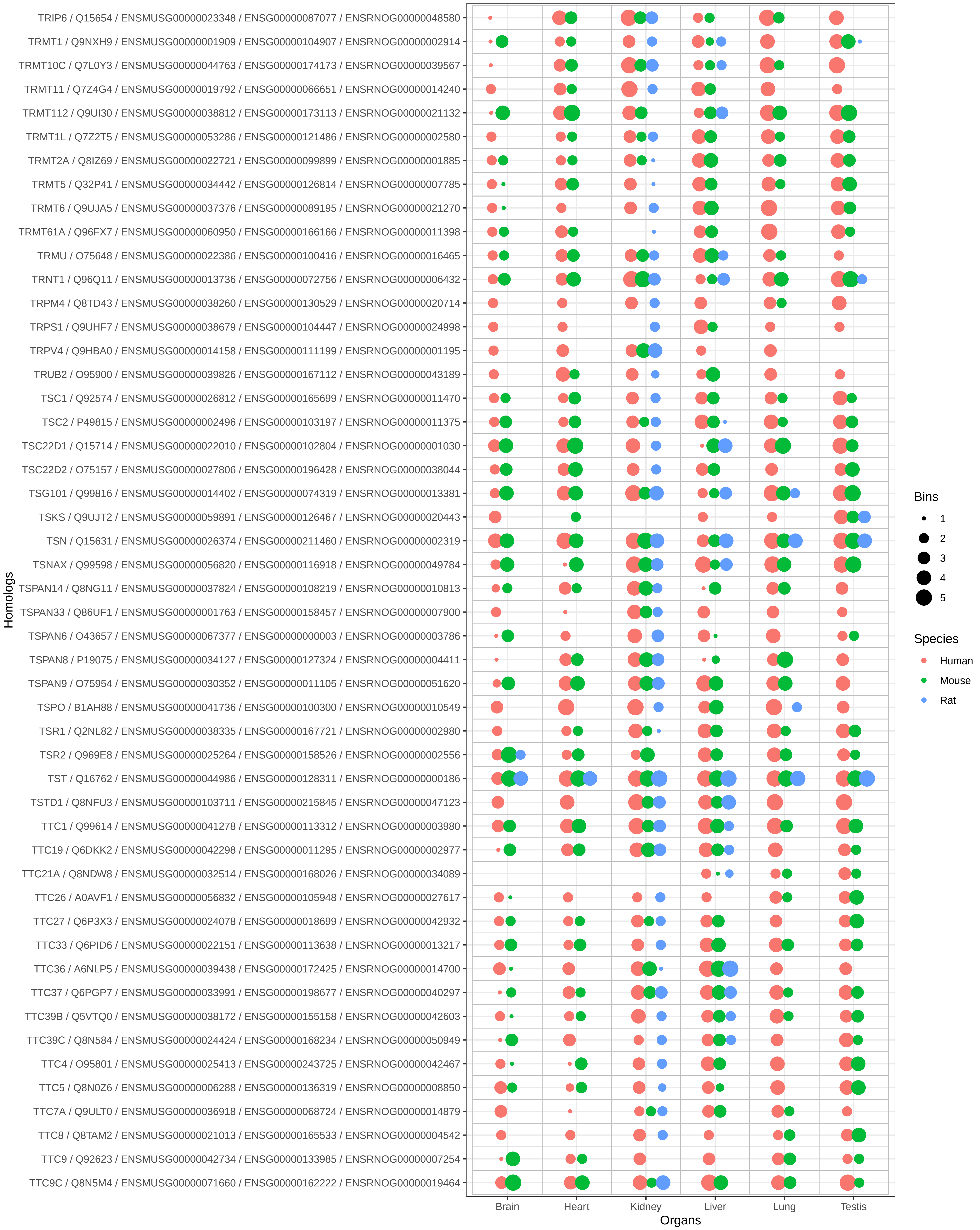
TRIP6 / Q15654 / ENSMUSG00000023348 / ENSG00000087077 / ENSRNOG00000048580
TRMT1 / Q9NXH9 / ENSMUSG00000001909 / ENSG00000104907 / ENSRNOG00000002914
TRMT10C / Q7L0Y3 / ENSMUSG00000044763 / ENSG00000174173 / ENSRNOG00000039567
TRMT11 / Q7Z4G4 / ENSMUSG00000019792 / ENSG00000066651 / ENSRNOG00000014240
TRMT112 / Q9UI30 / ENSMUSG00000038812 / ENSG00000173113 / ENSRNOG00000021132
TRMT1L / Q7Z2T5 / ENSMUSG00000053286 / ENSG00000121486 / ENSRNOG00000002580
TRMT2A / Q8IZ69 / ENSMUSG00000022721 / ENSG00000099899 / ENSRNOG00000001885
TRMT5 / Q32P41 / ENSMUSG00000034442 / ENSG00000126814 / ENSRNOG00000007785
TRMT6 / Q9UJA5 / ENSMUSG00000037376 / ENSG00000089195 / ENSRNOG00000021270
TRMT61A / Q96FX7 / ENSMUSG00000060950 / ENSG00000166166 / ENSRNOG00000011398
TRMU / O75648 / ENSMUSG00000022386 / ENSG00000100416 / ENSRNOG00000016465
TRNT1 / Q96Q11 / ENSMUSG00000013736 / ENSG00000072756 / ENSRNOG00000006432
TRPM4 / Q8TD43 / ENSMUSG00000038260 / ENSG00000130529 / ENSRNOG00000020714
TRPS1 / Q9UHF7 / ENSMUSG00000038679 / ENSG00000104447 / ENSRNOG00000024998
TRPV4 / Q9HBA0 / ENSMUSG00000014158 / ENSG00000111199 / ENSRNOG00000001195
TRUB2 / O95900 / ENSMUSG00000039826 / ENSG00000167112 / ENSRNOG00000043189
TSC1 / Q92574 / ENSMUSG00000026812 / ENSG00000165699 / ENSRNOG00000011470
TSC2 / P49815 / ENSMUSG00000002496 / ENSG00000103197 / ENSRNOG00000011375
TSC22D1 / Q15714 / ENSMUSG00000022010 / ENSG00000102804 / ENSRNOG00000001030
TSC22D2 / O75157 / ENSMUSG00000027806 / ENSG00000196428 / ENSRNOG00000038044
TSG101 / Q99816 / ENSMUSG00000014402 / ENSG00000074319 / ENSRNOG00000013381
TSKS / Q9UJT2 / ENSMUSG00000059891 / ENSG00000126467 / ENSRNOG00000020443
TSN / Q15631 / ENSMUSG00000026374 / ENSG00000211460 / ENSRNOG00000002319
TSNAX / Q99598 / ENSMUSG00000056820 / ENSG00000116918 / ENSRNOG00000049784
TSPAN14 / Q8NG11 / ENSMUSG00000037824 / ENSG00000108219 / ENSRNOG00000010813
TSPAN33 / Q86UF1 / ENSMUSG00000001763 / ENSG00000158457 / ENSRNOG00000007900
TSPAN6 / O43657 / ENSMUSG00000067377 / ENSG00000000003 / ENSRNOG00000003786
TSPAN8 / P19075 / ENSMUSG00000034127 / ENSG00000127324 / ENSRNOG00000004411
TSPAN9 / O75954 / ENSMUSG00000030352 / ENSG00000011105 / ENSRNOG00000051620
TSPO / B1AH88 / ENSMUSG00000041736 / ENSG00000100300 / ENSRNOG00000010549
TSR1 / Q2NL82 / ENSMUSG00000038335 / ENSG00000167721 / ENSRNOG00000002980
TSR2 / Q969E8 / ENSMUSG00000025264 / ENSG00000158526 / ENSRNOG00000002556
TST / Q16762 / ENSMUSG00000044986 / ENSG00000128311 / ENSRNOG00000000186
TSTD1 / Q8NFU3 / ENSMUSG00000103711 / ENSG00000215845 / ENSRNOG00000047123
TTC1 / Q99614 / ENSMUSG00000041278 / ENSG00000113312 / ENSRNOG00000003980
TTC19 / Q6DKK2 / ENSMUSG00000042298 / ENSG00000011295 / ENSRNOG00000002977
TTC21A / Q8NDW8 / ENSMUSG00000032514 / ENSG00000168026 / ENSRNOG00000034089
TTC26 / A0AVF1 / ENSMUSG00000056832 / ENSG00000105948 / ENSRNOG00000027617
TTC27 / Q6P3X3 / ENSMUSG00000024078 / ENSG00000018699 / ENSRNOG00000042932
TTC33 / Q6PID6 / ENSMUSG00000022151 / ENSG00000113638 / ENSRNOG00000013217
TTC36 / A6NLP5 / ENSMUSG00000039438 / ENSG00000172425 / ENSRNOG00000014700
TTC37 / Q6PGP7 / ENSMUSG00000033991 / ENSG00000198677 / ENSRNOG00000040297
TTC39B / Q5VTQ0 / ENSMUSG00000038172 / ENSG00000155158 / ENSRNOG00000042603
TTC39C / Q8N584 / ENSMUSG00000024424 / ENSG00000168234 / ENSRNOG00000050949
TTC4 / O95801 / ENSMUSG00000025413 / ENSG00000243725 / ENSRNOG00000042467
TTC5 / Q8N0Z6 / ENSMUSG00000006288 / ENSG00000136319 / ENSRNOG00000008850
TTC7A / Q9ULT0 / ENSMUSG00000036918 / ENSG00000068724 / ENSRNOG00000014879
TTC8 / Q8TAM2 / ENSMUSG00000021013 / ENSG00000165533 / ENSRNOG00000004542
TTC9 / Q92623 / ENSMUSG00000042734 / ENSG00000133985 / ENSRNOG00000007254
TTC9C / Q8N5M4 / ENSMUSG00000071660 / ENSG00000162222 / ENSRNOG00000019464
Homologs
Brain
Heart
Kidney
Liver
Lung
Testis
Organs
Bins
1
2
3
4
5
Species
Human
Mouse
Rat

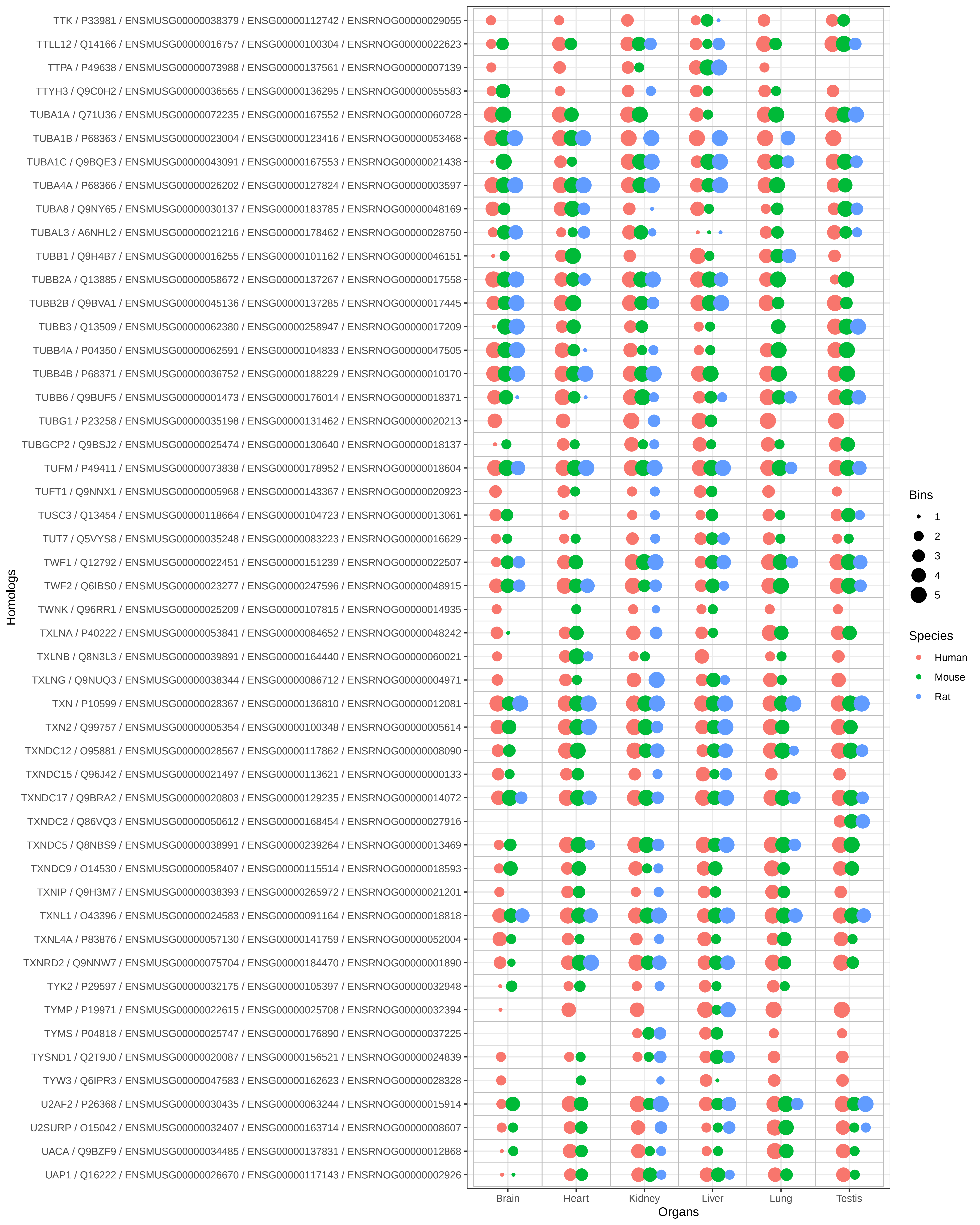
TTK / P33981 / ENSMUSG00000038379 / ENSG00000112742 / ENSRNOG00000029055
TTLL12 / Q14166 / ENSMUSG00000016757 / ENSG00000100304 / ENSRNOG00000022623
TTPA / P49638 / ENSMUSG00000073988 / ENSG00000137561 / ENSRNOG00000007139
TTYH3 / Q9C0H2 / ENSMUSG00000036565 / ENSG00000136295 / ENSRNOG00000055583
TUBA1A / Q71U36 / ENSMUSG00000072235 / ENSG00000167552 / ENSRNOG00000060728
TUBA1B / P68363 / ENSMUSG00000023004 / ENSG00000123416 / ENSRNOG00000053468
TUBA1C / Q9BQE3 / ENSMUSG00000043091 / ENSG00000167553 / ENSRNOG00000021438
TUBA4A / P68366 / ENSMUSG00000026202 / ENSG00000127824 / ENSRNOG00000003597
TUBA8 / Q9NY65 / ENSMUSG00000030137 / ENSG00000183785 / ENSRNOG00000048169
TUBAL3 / A6NHL2 / ENSMUSG00000021216 / ENSG00000178462 / ENSRNOG00000028750
TUBB1 / Q9H4B7 / ENSMUSG00000016255 / ENSG00000101162 / ENSRNOG00000046151
TUBB2A / Q13885 / ENSMUSG00000058672 / ENSG00000137267 / ENSRNOG00000017558
TUBB2B / Q9BVA1 / ENSMUSG00000045136 / ENSG00000137285 / ENSRNOG00000017445
TUBB3 / Q13509 / ENSMUSG00000062380 / ENSG00000258947 / ENSRNOG00000017209
TUBB4A / P04350 / ENSMUSG00000062591 / ENSG00000104833 / ENSRNOG00000047505
TUBB4B / P68371 / ENSMUSG00000036752 / ENSG00000188229 / ENSRNOG00000010170
TUBB6 / Q9BUF5 / ENSMUSG00000001473 / ENSG00000176014 / ENSRNOG00000018371
TUBG1 / P23258 / ENSMUSG00000035198 / ENSG00000131462 / ENSRNOG00000020213
TUBGCP2 / Q9BSJ2 / ENSMUSG00000025474 / ENSG00000130640 / ENSRNOG00000018137
TUFM / P49411 / ENSMUSG00000073838 / ENSG00000178952 / ENSRNOG00000018604
TUFT1 / Q9NNX1 / ENSMUSG00000005968 / ENSG00000143367 / ENSRNOG00000020923
TUSC3 / Q13454 / ENSMUSG00000118664 / ENSG00000104723 / ENSRNOG00000013061
TUT7 / Q5VYS8 / ENSMUSG00000035248 / ENSG00000083223 / ENSRNOG00000016629
TWF1 / Q12792 / ENSMUSG00000022451 / ENSG00000151239 / ENSRNOG00000022507
TWF2 / Q6IBS0 / ENSMUSG00000023277 / ENSG00000247596 / ENSRNOG00000048915
TWNK / Q96RR1 / ENSMUSG00000025209 / ENSG00000107815 / ENSRNOG00000014935
TXLNA / P40222 / ENSMUSG00000053841 / ENSG00000084652 / ENSRNOG00000048242
TXLNB / Q8N3L3 / ENSMUSG00000039891 / ENSG00000164440 / ENSRNOG00000060021
TXLNG / Q9NUQ3 / ENSMUSG00000038344 / ENSG00000086712 / ENSRNOG00000004971
TXN / P10599 / ENSMUSG00000028367 / ENSG00000136810 / ENSRNOG00000012081
TXN2 / Q99757 / ENSMUSG00000005354 / ENSG00000100348 / ENSRNOG00000005614
TXNDC12 / O95881 / ENSMUSG00000028567 / ENSG00000117862 / ENSRNOG00000008090
TXNDC15 / Q96J42 / ENSMUSG00000021497 / ENSG00000113621 / ENSRNOG00000000133
TXNDC17 / Q9BRA2 / ENSMUSG00000020803 / ENSG00000129235 / ENSRNOG00000014072
TXNDC2 / Q86VQ3 / ENSMUSG00000050612 / ENSG00000168454 / ENSRNOG00000027916
TXNDC5 / Q8NBS9 / ENSMUSG00000038991 / ENSG00000239264 / ENSRNOG00000013469
TXNDC9 / O14530 / ENSMUSG00000058407 / ENSG00000115514 / ENSRNOG00000018593
TXNIP / Q9H3M7 / ENSMUSG00000038393 / ENSG00000265972 / ENSRNOG00000021201
TXNL1 / O43396 / ENSMUSG00000024583 / ENSG00000091164 / ENSRNOG00000018818
TXNL4A / P83876 / ENSMUSG00000057130 / ENSG00000141759 / ENSRNOG00000052004
TXNRD2 / Q9NNW7 / ENSMUSG00000075704 / ENSG00000184470 / ENSRNOG00000001890
TYK2 / P29597 / ENSMUSG00000032175 / ENSG00000105397 / ENSRNOG00000032948
TYMP / P19971 / ENSMUSG00000022615 / ENSG00000025708 / ENSRNOG00000032394
TYMS / P04818 / ENSMUSG00000025747 / ENSG00000176890 / ENSRNOG00000037225
TYSND1 / Q2T9J0 / ENSMUSG00000020087 / ENSG00000156521 / ENSRNOG00000024839
TYW3 / Q6IPR3 / ENSMUSG00000047583 / ENSG00000162623 / ENSRNOG00000028328
U2AF2 / P26368 / ENSMUSG00000030435 / ENSG00000063244 / ENSRNOG00000015914
U2SURP / O15042 / ENSMUSG00000032407 / ENSG00000163714 / ENSRNOG00000008607
UACA / Q9BZF9 / ENSMUSG00000034485 / ENSG00000137831 / ENSRNOG00000012868
UAP1 / Q16222 / ENSMUSG00000026670 / ENSG00000117143 / ENSRNOG00000002926
Homologs
Brain
Heart
Kidney
Liver
Lung
Testis
Organs
Bins
1
2
3
4
5
Species
Human
Mouse
Rat

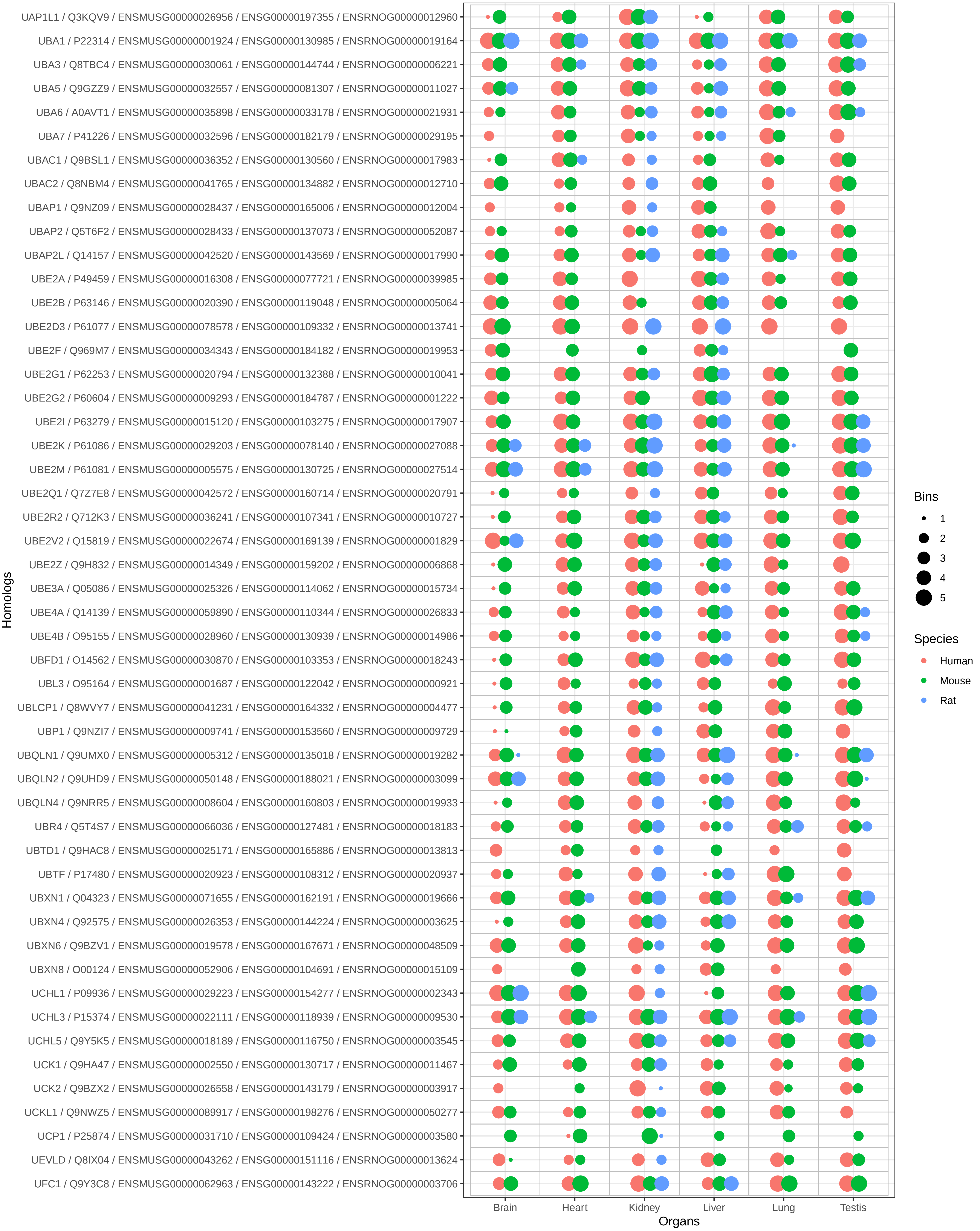

UAP1L1 / Q3KQV9 / ENSMUSG00000026956 / ENSG00000197355 / ENSRNOG00000012960
UBA1 / P22314 / ENSMUSG00000001924 / ENSG00000130985 / ENSRNOG00000019164
UBA3 / Q8TBC4 / ENSMUSG00000030061 / ENSG00000144744 / ENSRNOG00000006221
UBA5 / Q9GZZ9 / ENSMUSG00000032557 / ENSG00000081307 / ENSRNOG00000011027
UBA6 / A0AVT1 / ENSMUSG00000035898 / ENSG00000033178 / ENSRNOG00000021931
UBA7 / P41226 / ENSMUSG00000032596 / ENSG00000182179 / ENSRNOG00000029195
UBAC1 / Q9BSL1 / ENSMUSG00000036352 / ENSG00000130560 / ENSRNOG00000017983
UBAC2 / Q8NBM4 / ENSMUSG00000041765 / ENSG00000134882 / ENSRNOG00000012710
UBAP1 / Q9NZ09 / ENSMUSG00000028437 / ENSG00000165006 / ENSRNOG00000012004
UBAP2 / Q5T6F2 / ENSMUSG00000028433 / ENSG00000137073 / ENSRNOG00000052087
UBAP2L / Q14157 / ENSMUSG00000042520 / ENSG00000143569 / ENSRNOG00000017990
UBE2A / P49459 / ENSMUSG00000016308 / ENSG00000077721 / ENSRNOG00000039985
UBE2B / P63146 / ENSMUSG00000020390 / ENSG00000119048 / ENSRNOG00000005064
UBE2D3 / P61077 / ENSMUSG00000078578 / ENSG00000109332 / ENSRNOG00000013741
UBE2F / Q969M7 / ENSMUSG00000034343 / ENSG00000184182 / ENSRNOG00000019953
UBE2G1 / P62253 / ENSMUSG00000020794 / ENSG00000132388 / ENSRNOG00000010041
UBE2G2 / P60604 / ENSMUSG00000009293 / ENSG00000184787 / ENSRNOG00000001222
UBE2I / P63279 / ENSMUSG00000015120 / ENSG00000103275 / ENSRNOG00000017907
UBE2K / P61086 / ENSMUSG00000029203 / ENSG00000078140 / ENSRNOG00000027088
UBE2M / P61081 / ENSMUSG00000005575 / ENSG00000130725 / ENSRNOG00000027514
UBE2Q1 / Q7Z7E8 / ENSMUSG00000042572 / ENSG00000160714 / ENSRNOG00000020791
UBE2R2 / Q712K3 / ENSMUSG00000036241 / ENSG00000107341 / ENSRNOG00000010727
UBE2V2 / Q15819 / ENSMUSG00000022674 / ENSG00000169139 / ENSRNOG00000001829
UBE2Z / Q9H832 / ENSMUSG00000014349 / ENSG00000159202 / ENSRNOG00000006868
UBE3A / Q05086 / ENSMUSG00000025326 / ENSG00000114062 / ENSRNOG00000015734
UBE4A / Q14139 / ENSMUSG00000059890 / ENSG00000110344 / ENSRNOG00000026833
UBE4B / O95155 / ENSMUSG00000028960 / ENSG00000130939 / ENSRNOG00000014986
UBFD1 / O14562 / ENSMUSG00000030870 / ENSG00000103353 / ENSRNOG00000018243
UBL3 / O95164 / ENSMUSG00000001687 / ENSG00000122042 / ENSRNOG00000000921
UBLCP1 / Q8WVY7 / ENSMUSG00000041231 / ENSG00000164332 / ENSRNOG00000004477
UBP1 / Q9NZI7 / ENSMUSG00000009741 / ENSG00000153560 / ENSRNOG00000009729
UBQLN1 / Q9UMX0 / ENSMUSG00000005312 / ENSG00000135018 / ENSRNOG00000019282
UBQLN2 / Q9UHD9 / ENSMUSG00000050148 / ENSG00000188021 / ENSRNOG00000003099
UBQLN4 / Q9NRR5 / ENSMUSG00000008604 / ENSG00000160803 / ENSRNOG00000019933
UBR4 / Q5T4S7 / ENSMUSG00000066036 / ENSG00000127481 / ENSRNOG00000018183
UBTD1 / Q9HAC8 / ENSMUSG00000025171 / ENSG00000165886 / ENSRNOG00000013813
UBTF / P17480 / ENSMUSG00000020923 / ENSG00000108312 / ENSRNOG00000020937
UBXN1 / Q04323 / ENSMUSG00000071655 / ENSG00000162191 / ENSRNOG00000019666
UBXN4 / Q92575 / ENSMUSG00000026353 / ENSG00000144224 / ENSRNOG00000003625
UBXN6 / Q9BZV1 / ENSMUSG00000019578 / ENSG00000167671 / ENSRNOG00000048509
UBXN8 / O00124 / ENSMUSG00000052906 / ENSG00000104691 / ENSRNOG00000015109
UCHL1 / P09936 / ENSMUSG00000029223 / ENSG00000154277 / ENSRNOG00000002343
UCHL3 / P15374 / ENSMUSG00000022111 / ENSG00000118939 / ENSRNOG00000009530
UCHL5 / Q9Y5K5 / ENSMUSG00000018189 / ENSG00000116750 / ENSRNOG00000003545
UCK1 / Q9HA47 / ENSMUSG00000002550 / ENSG00000130717 / ENSRNOG00000011467
UCK2 / Q9BZX2 / ENSMUSG00000026558 / ENSG00000143179 / ENSRNOG00000003917
UCKL1 / Q9NWZ5 / ENSMUSG00000089917 / ENSG00000198276 / ENSRNOG00000050277
UCP1 / P25874 / ENSMUSG00000031710 / ENSG00000109424 / ENSRNOG00000003580
UEVLD / Q8IX04 / ENSMUSG00000043262 / ENSG00000151116 / ENSRNOG00000013624
UFC1 / Q9Y3C8 / ENSMUSG00000062963 / ENSG00000143222 / ENSRNOG00000003706
Homologs
Brain
Heart
Kidney
Liver
Lung
Testis
Organs
Bins
1
2
3
4
5
Species
Human
Mouse
Rat

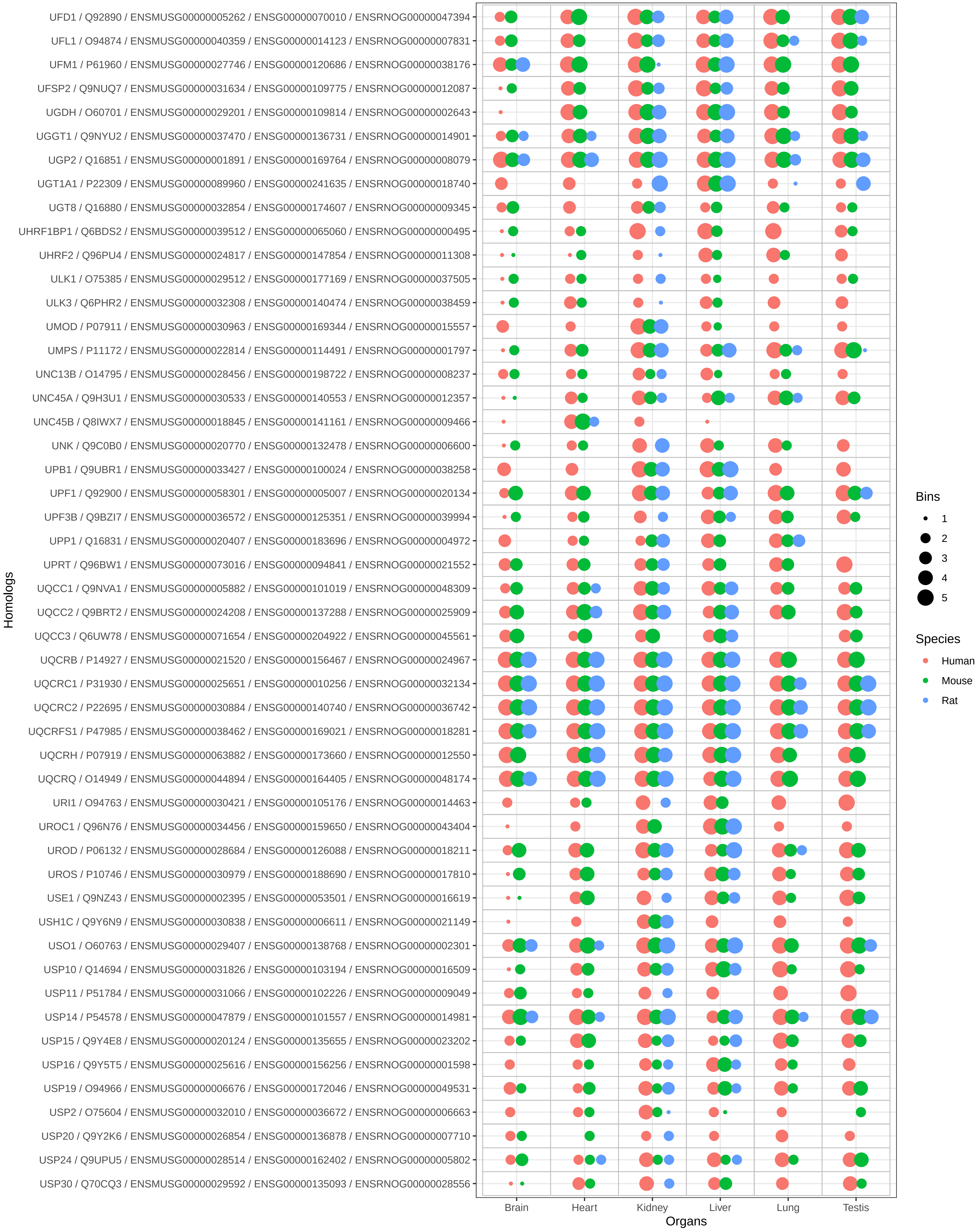
UFD1 / Q92890 / ENSMUSG00000005262 / ENSG00000070010 / ENSRNOG00000047394
UFL1 / O94874 / ENSMUSG00000040359 / ENSG00000014123 / ENSRNOG00000007831
UFM1 / P61960 / ENSMUSG00000027746 / ENSG00000120686 / ENSRNOG00000038176
UFSP2 / Q9NUQ7 / ENSMUSG00000031634 / ENSG00000109775 / ENSRNOG00000012087
UGDH / O60701 / ENSMUSG00000029201 / ENSG00000109814 / ENSRNOG00000002643
UGGT1 / Q9NYU2 / ENSMUSG00000037470 / ENSG00000136731 / ENSRNOG00000014901
UGP2 / Q16851 / ENSMUSG00000001891 / ENSG00000169764 / ENSRNOG00000008079
UGT1A1 / P22309 / ENSMUSG00000089960 / ENSG00000241635 / ENSRNOG00000018740
UGT8 / Q16880 / ENSMUSG00000032854 / ENSG00000174607 / ENSRNOG00000009345
UHRF1BP1 / Q6BDS2 / ENSMUSG00000039512 / ENSG00000065060 / ENSRNOG00000000495
UHRF2 / Q96PU4 / ENSMUSG00000024817 / ENSG00000147854 / ENSRNOG00000011308
ULK1 / O75385 / ENSMUSG00000029512 / ENSG00000177169 / ENSRNOG00000037505
ULK3 / Q6PHR2 / ENSMUSG00000032308 / ENSG00000140474 / ENSRNOG00000038459
UMOD / P07911 / ENSMUSG00000030963 / ENSG00000169344 / ENSRNOG00000015557
UMPS / P11172 / ENSMUSG00000022814 / ENSG00000114491 / ENSRNOG00000001797
UNC13B / O14795 / ENSMUSG00000028456 / ENSG00000198722 / ENSRNOG00000008237
UNC45A / Q9H3U1 / ENSMUSG00000030533 / ENSG00000140553 / ENSRNOG00000012357
UNC45B / Q8IWX7 / ENSMUSG00000018845 / ENSG00000141161 / ENSRNOG00000009466
UNK / Q9C0B0 / ENSMUSG00000020770 / ENSG00000132478 / ENSRNOG00000006600
UPB1 / Q9UBR1 / ENSMUSG00000033427 / ENSG00000100024 / ENSRNOG00000038258
UPF1 / Q92900 / ENSMUSG00000058301 / ENSG00000005007 / ENSRNOG00000020134
UPF3B / Q9BZI7 / ENSMUSG00000036572 / ENSG00000125351 / ENSRNOG00000039994
UPP1 / Q16831 / ENSMUSG00000020407 / ENSG00000183696 / ENSRNOG00000004972
UPRT / Q96BW1 / ENSMUSG00000073016 / ENSG00000094841 / ENSRNOG00000021552
UQCC1 / Q9NVA1 / ENSMUSG00000005882 / ENSG00000101019 / ENSRNOG00000048309
UQCC2 / Q9BRT2 / ENSMUSG00000024208 / ENSG00000137288 / ENSRNOG00000025909
UQCC3 / Q6UW78 / ENSMUSG00000071654 / ENSG00000204922 / ENSRNOG00000045561
UQCRB / P14927 / ENSMUSG00000021520 / ENSG00000156467 / ENSRNOG00000024967
UQCRC1 / P31930 / ENSMUSG00000025651 / ENSG00000010256 / ENSRNOG00000032134
UQCRC2 / P22695 / ENSMUSG00000030884 / ENSG00000140740 / ENSRNOG00000036742
UQCRFS1 / P47985 / ENSMUSG00000038462 / ENSG00000169021 / ENSRNOG00000018281
UQCRH / P07919 / ENSMUSG00000063882 / ENSG00000173660 / ENSRNOG00000012550
UQCRQ / O14949 / ENSMUSG00000044894 / ENSG00000164405 / ENSRNOG00000048174
URI1 / O94763 / ENSMUSG00000030421 / ENSG00000105176 / ENSRNOG00000014463
UROC1 / Q96N76 / ENSMUSG00000034456 / ENSG00000159650 / ENSRNOG00000043404
UROD / P06132 / ENSMUSG00000028684 / ENSG00000126088 / ENSRNOG00000018211
UROS / P10746 / ENSMUSG00000030979 / ENSG00000188690 / ENSRNOG00000017810
USE1 / Q9NZ43 / ENSMUSG00000002395 / ENSG00000053501 / ENSRNOG00000016619
USH1C / Q9Y6N9 / ENSMUSG00000030838 / ENSG00000006611 / ENSRNOG00000021149
USO1 / O60763 / ENSMUSG00000029407 / ENSG00000138768 / ENSRNOG00000002301
USP10 / Q14694 / ENSMUSG00000031826 / ENSG00000103194 / ENSRNOG00000016509
USP11 / P51784 / ENSMUSG00000031066 / ENSG00000102226 / ENSRNOG00000009049
USP14 / P54578 / ENSMUSG00000047879 / ENSG00000101557 / ENSRNOG00000014981
USP15 / Q9Y4E8 / ENSMUSG00000020124 / ENSG00000135655 / ENSRNOG00000023202
USP16 / Q9Y5T5 / ENSMUSG00000025616 / ENSG00000156256 / ENSRNOG00000001598
USP19 / O94966 / ENSMUSG00000006676 / ENSG00000172046 / ENSRNOG00000049531
USP2 / O75604 / ENSMUSG00000032010 / ENSG00000036672 / ENSRNOG00000006663
USP20 / Q9Y2K6 / ENSMUSG00000026854 / ENSG00000136878 / ENSRNOG00000007710
USP24 / Q9UPU5 / ENSMUSG00000028514 / ENSG00000162402 / ENSRNOG00000005802
USP30 / Q70CQ3 / ENSMUSG00000029592 / ENSG00000135093 / ENSRNOG00000028556
Homologs
Brain
Heart
Kidney
Liver
Lung
Testis
Organs
Bins
1
2
3
4
5
Species
Human
Mouse
Rat

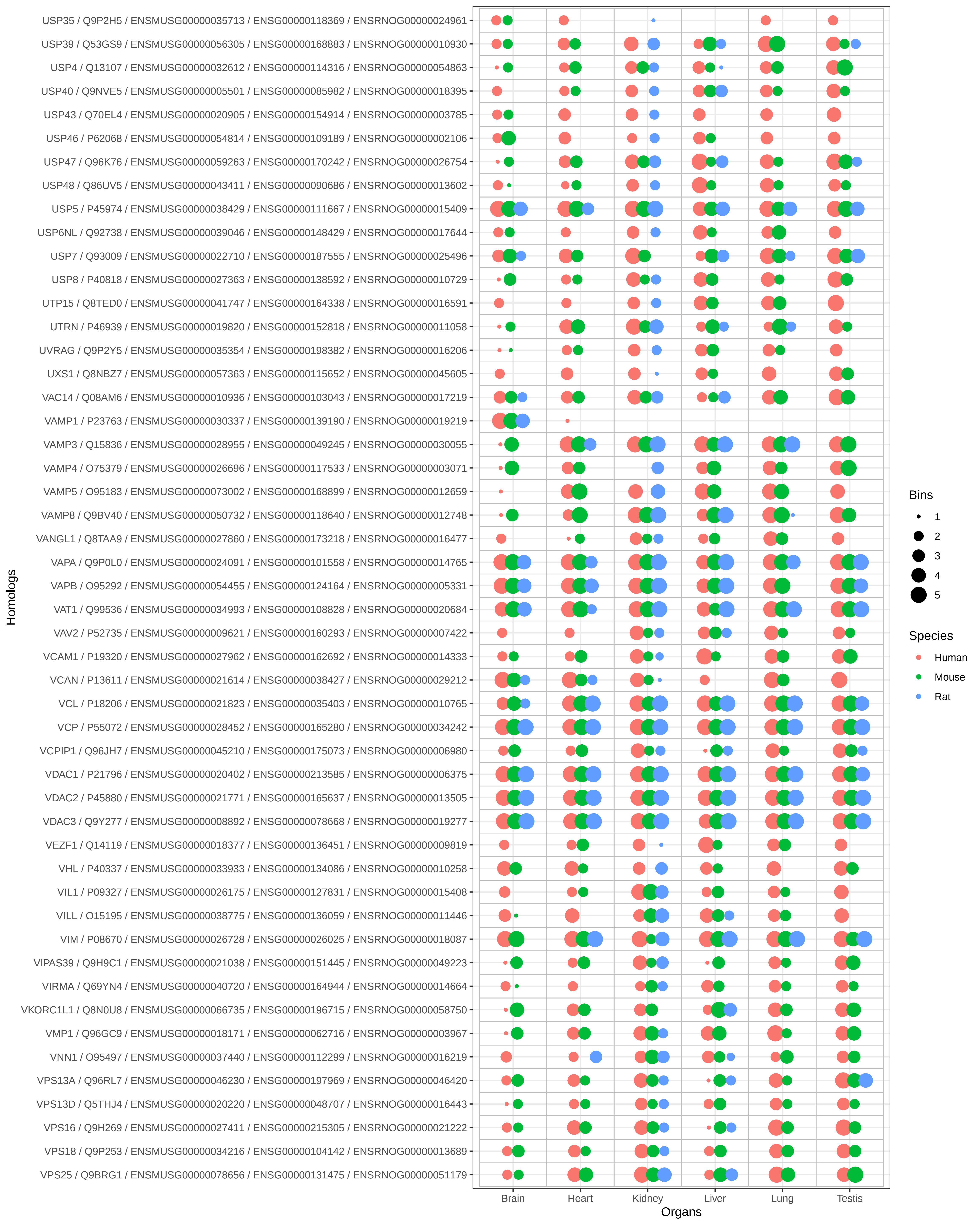
USP35 / Q9P2H5 / ENSMUSG00000035713 / ENSG00000118369 / ENSRNOG00000024961
USP39 / Q53GS9 / ENSMUSG00000056305 / ENSG00000168883 / ENSRNOG00000010930
USP4 / Q13107 / ENSMUSG00000032612 / ENSG00000114316 / ENSRNOG00000054863
USP40 / Q9NVE5 / ENSMUSG00000005501 / ENSG00000085982 / ENSRNOG00000018395
USP43 / Q70EL4 / ENSMUSG00000020905 / ENSG00000154914 / ENSRNOG00000003785
USP46 / P62068 / ENSMUSG00000054814 / ENSG00000109189 / ENSRNOG00000002106
USP47 / Q96K76 / ENSMUSG00000059263 / ENSG00000170242 / ENSRNOG00000026754
USP48 / Q86UV5 / ENSMUSG00000043411 / ENSG00000090686 / ENSRNOG00000013602
USP5 / P45974 / ENSMUSG00000038429 / ENSG00000111667 / ENSRNOG00000015409
USP6NL / Q92738 / ENSMUSG00000039046 / ENSG00000148429 / ENSRNOG00000017644
USP7 / Q93009 / ENSMUSG00000022710 / ENSG00000187555 / ENSRNOG00000025496
USP8 / P40818 / ENSMUSG00000027363 / ENSG00000138592 / ENSRNOG00000010729
UTP15 / Q8TED0 / ENSMUSG00000041747 / ENSG00000164338 / ENSRNOG00000016591
UTRN / P46939 / ENSMUSG00000019820 / ENSG00000152818 / ENSRNOG00000011058
UVRAG / Q9P2Y5 / ENSMUSG00000035354 / ENSG00000198382 / ENSRNOG00000016206
UXS1 / Q8NBZ7 / ENSMUSG00000057363 / ENSG00000115652 / ENSRNOG00000045605
VAC14 / Q08AM6 / ENSMUSG00000010936 / ENSG00000103043 / ENSRNOG00000017219
VAMP1 / P23763 / ENSMUSG00000030337 / ENSG00000139190 / ENSRNOG00000019219
VAMP3 / Q15836 / ENSMUSG00000028955 / ENSG00000049245 / ENSRNOG00000030055
VAMP4 / O75379 / ENSMUSG00000026696 / ENSG00000117533 / ENSRNOG00000003071
VAMP5 / O95183 / ENSMUSG00000073002 / ENSG00000168899 / ENSRNOG00000012659
VAMP8 / Q9BV40 / ENSMUSG00000050732 / ENSG00000118640 / ENSRNOG00000012748
VANGL1 / Q8TAA9 / ENSMUSG00000027860 / ENSG00000173218 / ENSRNOG00000016477
VAPA / Q9P0L0 / ENSMUSG00000024091 / ENSG00000101558 / ENSRNOG00000014765
VAPB / O95292 / ENSMUSG00000054455 / ENSG00000124164 / ENSRNOG00000005331
VAT1 / Q99536 / ENSMUSG00000034993 / ENSG00000108828 / ENSRNOG00000020684
VAV2 / P52735 / ENSMUSG00000009621 / ENSG00000160293 / ENSRNOG00000007422
VCAM1 / P19320 / ENSMUSG00000027962 / ENSG00000162692 / ENSRNOG00000014333
VCAN / P13611 / ENSMUSG00000021614 / ENSG00000038427 / ENSRNOG00000029212
VCL / P18206 / ENSMUSG00000021823 / ENSG00000035403 / ENSRNOG00000010765
VCP / P55072 / ENSMUSG00000028452 / ENSG00000165280 / ENSRNOG00000034242
VCPIP1 / Q96JH7 / ENSMUSG00000045210 / ENSG00000175073 / ENSRNOG00000006980
VDAC1 / P21796 / ENSMUSG00000020402 / ENSG00000213585 / ENSRNOG00000006375
VDAC2 / P45880 / ENSMUSG00000021771 / ENSG00000165637 / ENSRNOG00000013505
VDAC3 / Q9Y277 / ENSMUSG00000008892 / ENSG00000078668 / ENSRNOG00000019277
VEZF1 / Q14119 / ENSMUSG00000018377 / ENSG00000136451 / ENSRNOG00000009819
VHL / P40337 / ENSMUSG00000033933 / ENSG00000134086 / ENSRNOG00000010258
VIL1 / P09327 / ENSMUSG00000026175 / ENSG00000127831 / ENSRNOG00000015408
VILL / O15195 / ENSMUSG00000038775 / ENSG00000136059 / ENSRNOG00000011446
VIM / P08670 / ENSMUSG00000026728 / ENSG00000026025 / ENSRNOG00000018087
VIPAS39 / Q9H9C1 / ENSMUSG00000021038 / ENSG00000151445 / ENSRNOG00000049223
VIRMA / Q69YN4 / ENSMUSG00000040720 / ENSG00000164944 / ENSRNOG00000014664
VKORC1L1 / Q8N0U8 / ENSMUSG00000066735 / ENSG00000196715 / ENSRNOG00000058750
VMP1 / Q96GC9 / ENSMUSG00000018171 / ENSG00000062716 / ENSRNOG00000003967
VNN1 / O95497 / ENSMUSG00000037440 / ENSG00000112299 / ENSRNOG00000016219
VPS13A / Q96RL7 / ENSMUSG00000046230 / ENSG00000197969 / ENSRNOG00000046420
VPS13D / Q5THJ4 / ENSMUSG00000020220 / ENSG00000048707 / ENSRNOG00000016443
VPS16 / Q9H269 / ENSMUSG00000027411 / ENSG00000215305 / ENSRNOG00000021222
VPS18 / Q9P253 / ENSMUSG00000034216 / ENSG00000104142 / ENSRNOG00000013689
VPS25 / Q9BRG1 / ENSMUSG00000078656 / ENSG00000131475 / ENSRNOG00000051179
Homologs
Brain
Heart
Kidney
Liver
Lung
Testis
Organs
Bins
1
2
3
4
5
Species
Human
Mouse
Rat

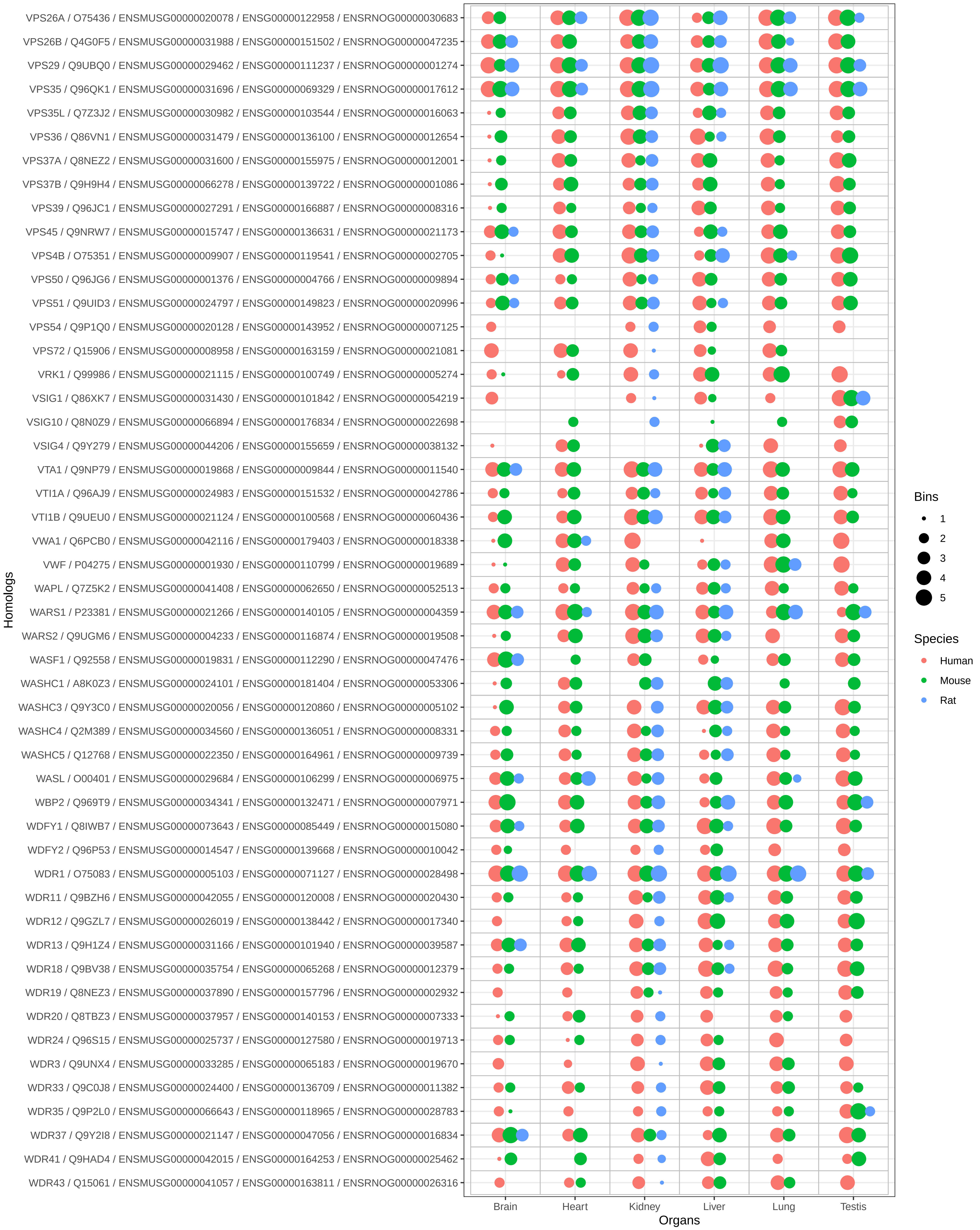
VPS26A / O75436 / ENSMUSG00000020078 / ENSG00000122958 / ENSRNOG00000030683
VPS26B / Q4G0F5 / ENSMUSG00000031988 / ENSG00000151502 / ENSRNOG00000047235
VPS29 / Q9UBQ0 / ENSMUSG00000029462 / ENSG00000111237 / ENSRNOG00000001274
VPS35 / Q96QK1 / ENSMUSG00000031696 / ENSG00000069329 / ENSRNOG00000017612
VPS35L / Q7Z3J2 / ENSMUSG00000030982 / ENSG00000103544 / ENSRNOG00000016063
VPS36 / Q86VN1 / ENSMUSG00000031479 / ENSG00000136100 / ENSRNOG00000012654
VPS37A / Q8NEZ2 / ENSMUSG00000031600 / ENSG00000155975 / ENSRNOG00000012001
VPS37B / Q9H9H4 / ENSMUSG00000066278 / ENSG00000139722 / ENSRNOG00000001086
VPS39 / Q96JC1 / ENSMUSG00000027291 / ENSG00000166887 / ENSRNOG00000008316
VPS45 / Q9NRW7 / ENSMUSG00000015747 / ENSG00000136631 / ENSRNOG00000021173
VPS4B / O75351 / ENSMUSG00000009907 / ENSG00000119541 / ENSRNOG00000002705
VPS50 / Q96JG6 / ENSMUSG00000001376 / ENSG00000004766 / ENSRNOG00000009894
VPS51 / Q9UID3 / ENSMUSG00000024797 / ENSG00000149823 / ENSRNOG00000020996
VPS54 / Q9P1Q0 / ENSMUSG00000020128 / ENSG00000143952 / ENSRNOG00000007125
VPS72 / Q15906 / ENSMUSG00000008958 / ENSG00000163159 / ENSRNOG00000021081
VRK1 / Q99986 / ENSMUSG00000021115 / ENSG00000100749 / ENSRNOG00000005274
VSIG1 / Q86XK7 / ENSMUSG00000031430 / ENSG00000101842 / ENSRNOG00000054219
VSIG10 / Q8N0Z9 / ENSMUSG00000066894 / ENSG00000176834 / ENSRNOG00000022698
VSIG4 / Q9Y279 / ENSMUSG00000044206 / ENSG00000155659 / ENSRNOG00000038132
VTA1 / Q9NP79 / ENSMUSG00000019868 / ENSG00000009844 / ENSRNOG00000011540
VTI1A / Q96AJ9 / ENSMUSG00000024983 / ENSG00000151532 / ENSRNOG00000042786
VTI1B / Q9UEU0 / ENSMUSG00000021124 / ENSG00000100568 / ENSRNOG00000060436
VWA1 / Q6PCB0 / ENSMUSG00000042116 / ENSG00000179403 / ENSRNOG00000018338
VWF / P04275 / ENSMUSG00000001930 / ENSG00000110799 / ENSRNOG00000019689
WAPL / Q7Z5K2 / ENSMUSG00000041408 / ENSG00000062650 / ENSRNOG00000052513
WARS1 / P23381 / ENSMUSG00000021266 / ENSG00000140105 / ENSRNOG00000004359
WARS2 / Q9UGM6 / ENSMUSG00000004233 / ENSG00000116874 / ENSRNOG00000019508
WASF1 / Q92558 / ENSMUSG00000019831 / ENSG00000112290 / ENSRNOG00000047476
WASHC1 / A8K0Z3 / ENSMUSG00000024101 / ENSG00000181404 / ENSRNOG00000053306
WASHC3 / Q9Y3C0 / ENSMUSG00000020056 / ENSG00000120860 / ENSRNOG00000005102
WASHC4 / Q2M389 / ENSMUSG00000034560 / ENSG00000136051 / ENSRNOG00000008331
WASHC5 / Q12768 / ENSMUSG00000022350 / ENSG00000164961 / ENSRNOG00000009739
WASL / O00401 / ENSMUSG00000029684 / ENSG00000106299 / ENSRNOG00000006975
WBP2 / Q969T9 / ENSMUSG00000034341 / ENSG00000132471 / ENSRNOG00000007971
WDFY1 / Q8IWB7 / ENSMUSG00000073643 / ENSG00000085449 / ENSRNOG00000015080
WDFY2 / Q96P53 / ENSMUSG00000014547 / ENSG00000139668 / ENSRNOG00000010042
WDR1 / O75083 / ENSMUSG00000005103 / ENSG00000071127 / ENSRNOG00000028498
WDR11 / Q9BZH6 / ENSMUSG00000042055 / ENSG00000120008 / ENSRNOG00000020430
WDR12 / Q9GZL7 / ENSMUSG00000026019 / ENSG00000138442 / ENSRNOG00000017340
WDR13 / Q9H1Z4 / ENSMUSG00000031166 / ENSG00000101940 / ENSRNOG00000039587
WDR18 / Q9BV38 / ENSMUSG00000035754 / ENSG00000065268 / ENSRNOG00000012379
WDR19 / Q8NEZ3 / ENSMUSG00000037890 / ENSG00000157796 / ENSRNOG00000002932
WDR20 / Q8TBZ3 / ENSMUSG00000037957 / ENSG00000140153 / ENSRNOG00000007333
WDR24 / Q96S15 / ENSMUSG00000025737 / ENSG00000127580 / ENSRNOG00000019713
WDR3 / Q9UNX4 / ENSMUSG00000033285 / ENSG00000065183 / ENSRNOG00000019670
WDR33 / Q9C0J8 / ENSMUSG00000024400 / ENSG00000136709 / ENSRNOG00000011382
WDR35 / Q9P2L0 / ENSMUSG00000066643 / ENSG00000118965 / ENSRNOG00000028783
WDR37 / Q9Y2I8 / ENSMUSG00000021147 / ENSG00000047056 / ENSRNOG00000016834
WDR41 / Q9HAD4 / ENSMUSG00000042015 / ENSG00000164253 / ENSRNOG00000025462
WDR43 / Q15061 / ENSMUSG00000041057 / ENSG00000163811 / ENSRNOG00000026316
Homologs
Brain
Heart
Kidney
Liver
Lung
Testis
Organs
Bins
1
2
3
4
5
Species
Human
Mouse
Rat

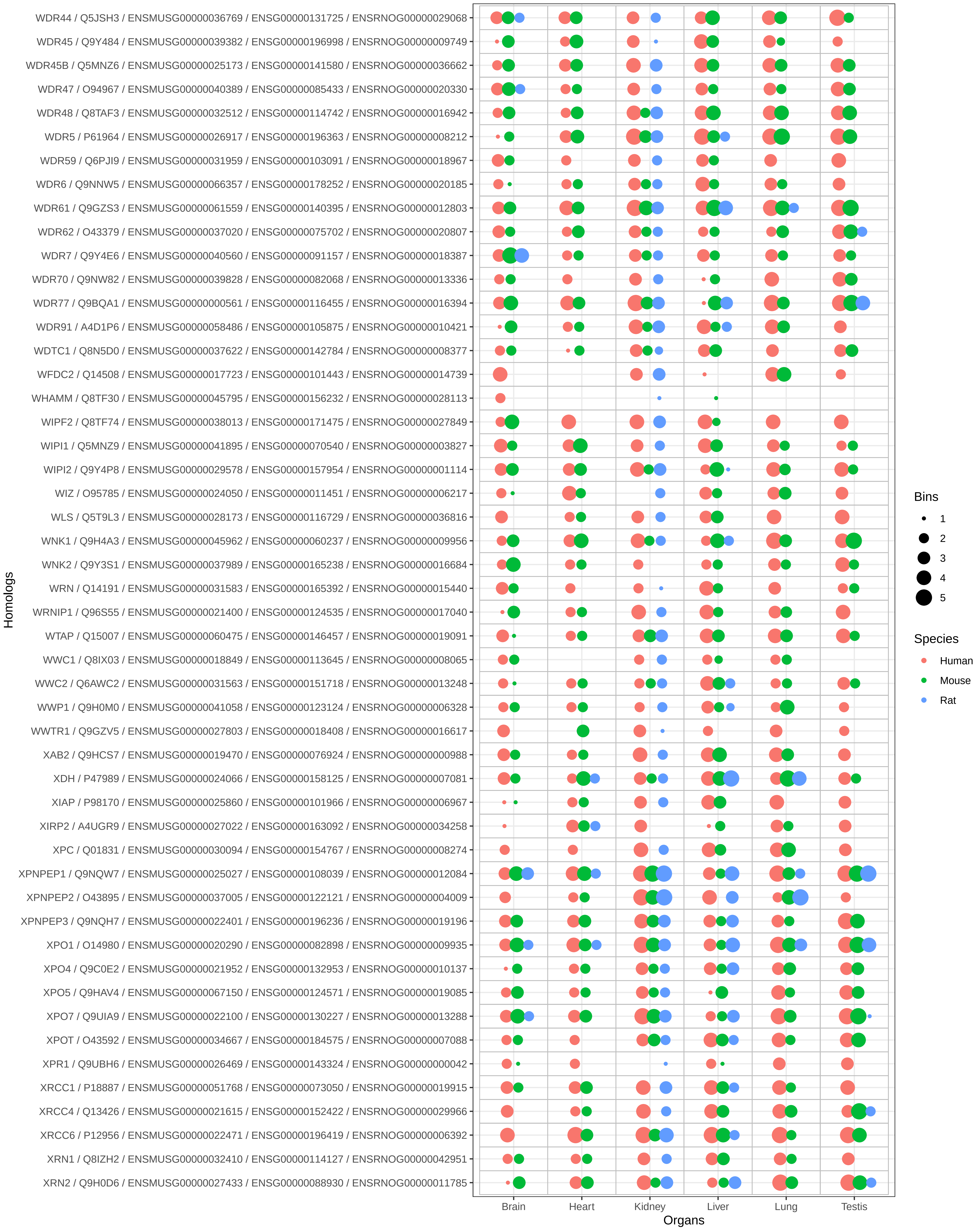
WDR44 / Q5JSH3 / ENSMUSG00000036769 / ENSG00000131725 / ENSRNOG00000029068
WDR45 / Q9Y484 / ENSMUSG00000039382 / ENSG00000196998 / ENSRNOG00000009749
WDR45B / Q5MNZ6 / ENSMUSG00000025173 / ENSG00000141580 / ENSRNOG00000036662
WDR47 / O94967 / ENSMUSG00000040389 / ENSG00000085433 / ENSRNOG00000020330
WDR48 / Q8TAF3 / ENSMUSG00000032512 / ENSG00000114742 / ENSRNOG00000016942
WDR5 / P61964 / ENSMUSG00000026917 / ENSG00000196363 / ENSRNOG00000008212
WDR59 / Q6PJI9 / ENSMUSG00000031959 / ENSG00000103091 / ENSRNOG00000018967
WDR6 / Q9NNW5 / ENSMUSG00000066357 / ENSG00000178252 / ENSRNOG00000020185
WDR61 / Q9GZS3 / ENSMUSG00000061559 / ENSG00000140395 / ENSRNOG00000012803
WDR62 / O43379 / ENSMUSG00000037020 / ENSG00000075702 / ENSRNOG00000020807
WDR7 / Q9Y4E6 / ENSMUSG00000040560 / ENSG00000091157 / ENSRNOG00000018387
WDR70 / Q9NW82 / ENSMUSG00000039828 / ENSG00000082068 / ENSRNOG00000013336
WDR77 / Q9BQA1 / ENSMUSG00000000561 / ENSG00000116455 / ENSRNOG00000016394
WDR91 / A4D1P6 / ENSMUSG00000058486 / ENSG00000105875 / ENSRNOG00000010421
WDTC1 / Q8N5D0 / ENSMUSG00000037622 / ENSG00000142784 / ENSRNOG00000008377
WFDC2 / Q14508 / ENSMUSG00000017723 / ENSG00000101443 / ENSRNOG00000014739
WHAMM / Q8TF30 / ENSMUSG00000045795 / ENSG00000156232 / ENSRNOG00000028113
WIPF2 / Q8TF74 / ENSMUSG00000038013 / ENSG00000171475 / ENSRNOG00000027849
WIPI1 / Q5MNZ9 / ENSMUSG00000041895 / ENSG00000070540 / ENSRNOG00000003827
WIPI2 / Q9Y4P8 / ENSMUSG00000029578 / ENSG00000157954 / ENSRNOG00000001114
WIZ / O95785 / ENSMUSG00000024050 / ENSG00000011451 / ENSRNOG00000006217
WLS / Q5T9L3 / ENSMUSG00000028173 / ENSG00000116729 / ENSRNOG00000036816
WNK1 / Q9H4A3 / ENSMUSG00000045962 / ENSG00000060237 / ENSRNOG00000009956
WNK2 / Q9Y3S1 / ENSMUSG00000037989 / ENSG00000165238 / ENSRNOG00000016684
WRN / Q14191 / ENSMUSG00000031583 / ENSG00000165392 / ENSRNOG00000015440
WRNIP1 / Q96S55 / ENSMUSG00000021400 / ENSG00000124535 / ENSRNOG00000017040
WTAP / Q15007 / ENSMUSG00000060475 / ENSG00000146457 / ENSRNOG00000019091
WWC1 / Q8IX03 / ENSMUSG00000018849 / ENSG00000113645 / ENSRNOG00000008065
WWC2 / Q6AWC2 / ENSMUSG00000031563 / ENSG00000151718 / ENSRNOG00000013248
WWP1 / Q9H0M0 / ENSMUSG00000041058 / ENSG00000123124 / ENSRNOG00000006328
WWTR1 / Q9GZV5 / ENSMUSG00000027803 / ENSG00000018408 / ENSRNOG00000016617
XAB2 / Q9HCS7 / ENSMUSG00000019470 / ENSG00000076924 / ENSRNOG00000000988
XDH / P47989 / ENSMUSG00000024066 / ENSG00000158125 / ENSRNOG00000007081
XIAP / P98170 / ENSMUSG00000025860 / ENSG00000101966 / ENSRNOG00000006967
XIRP2 / A4UGR9 / ENSMUSG00000027022 / ENSG00000163092 / ENSRNOG00000034258
XPC / Q01831 / ENSMUSG00000030094 / ENSG00000154767 / ENSRNOG00000008274
XPNPEP1 / Q9NQW7 / ENSMUSG00000025027 / ENSG00000108039 / ENSRNOG00000012084
XPNPEP2 / O43895 / ENSMUSG00000037005 / ENSG00000122121 / ENSRNOG00000004009
XPNPEP3 / Q9NQH7 / ENSMUSG00000022401 / ENSG00000196236 / ENSRNOG00000019196
XPO1 / O14980 / ENSMUSG00000020290 / ENSG00000082898 / ENSRNOG00000009935
XPO4 / Q9C0E2 / ENSMUSG00000021952 / ENSG00000132953 / ENSRNOG00000010137
XPO5 / Q9HAV4 / ENSMUSG00000067150 / ENSG00000124571 / ENSRNOG00000019085
XPO7 / Q9UIA9 / ENSMUSG00000022100 / ENSG00000130227 / ENSRNOG00000013288
XPOT / O43592 / ENSMUSG00000034667 / ENSG00000184575 / ENSRNOG00000007088
XPR1 / Q9UBH6 / ENSMUSG00000026469 / ENSG00000143324 / ENSRNOG00000000042
XRCC1 / P18887 / ENSMUSG00000051768 / ENSG00000073050 / ENSRNOG00000019915
XRCC4 / Q13426 / ENSMUSG00000021615 / ENSG00000152422 / ENSRNOG00000029966
XRCC6 / P12956 / ENSMUSG00000022471 / ENSG00000196419 / ENSRNOG00000006392
XRN1 / Q8IZH2 / ENSMUSG00000032410 / ENSG00000114127 / ENSRNOG00000042951
XRN2 / Q9H0D6 / ENSMUSG00000027433 / ENSG00000088930 / ENSRNOG00000011785
Homologs
Brain
Heart
Kidney
Liver
Lung
Testis
Organs
Bins
1
2
3
4
5
Species
Human
Mouse
Rat

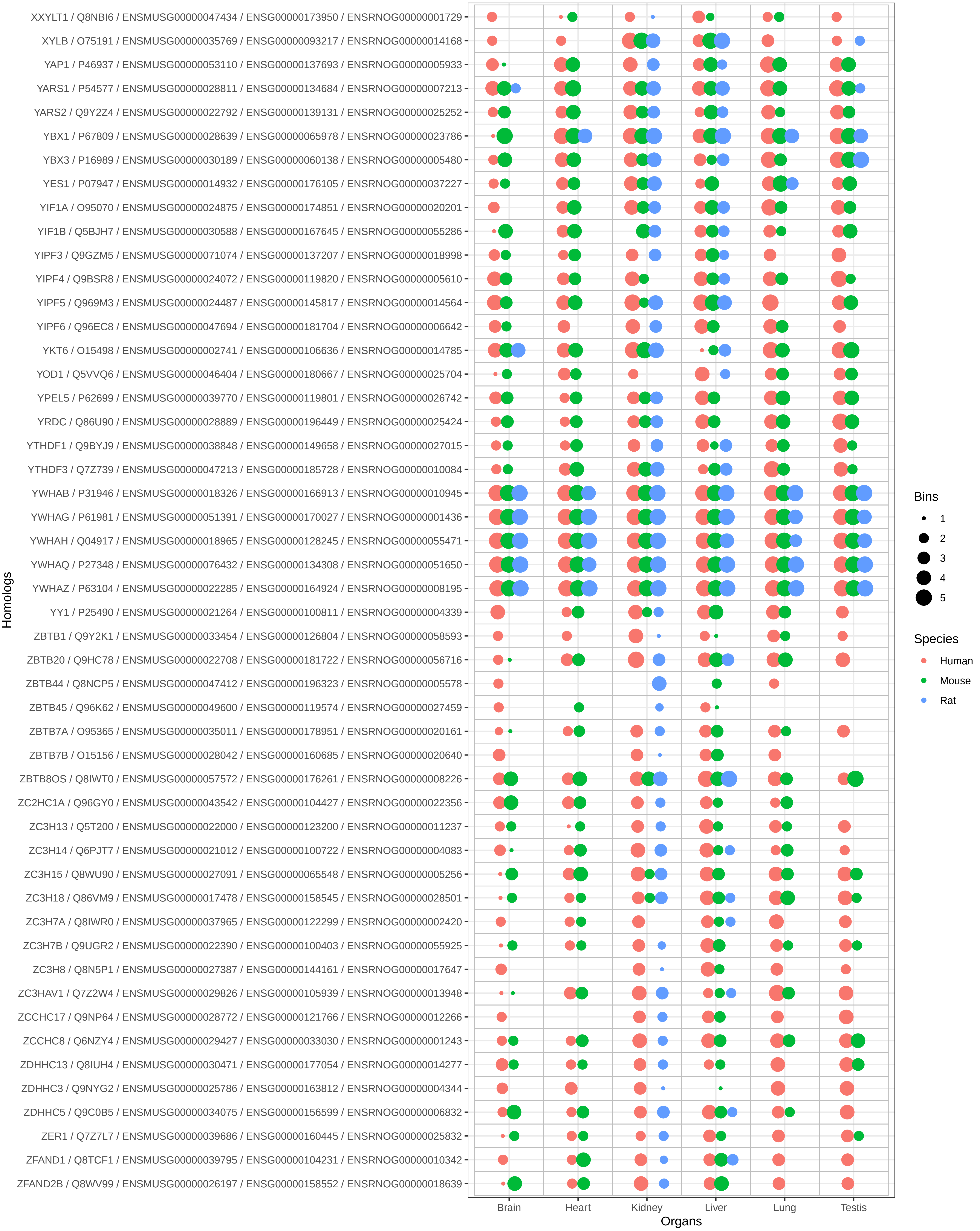
XXYLT1 / Q8NBI6 / ENSMUSG00000047434 / ENSG00000173950 / ENSRNOG00000001729
XYLB / O75191 / ENSMUSG00000035769 / ENSG00000093217 / ENSRNOG00000014168
YAP1 / P46937 / ENSMUSG00000053110 / ENSG00000137693 / ENSRNOG00000005933
YARS1 / P54577 / ENSMUSG00000028811 / ENSG00000134684 / ENSRNOG00000007213
YARS2 / Q9Y2Z4 / ENSMUSG00000022792 / ENSG00000139131 / ENSRNOG00000025252
YBX1 / P67809 / ENSMUSG00000028639 / ENSG00000065978 / ENSRNOG00000023786
YBX3 / P16989 / ENSMUSG00000030189 / ENSG00000060138 / ENSRNOG00000005480
YES1 / P07947 / ENSMUSG00000014932 / ENSG00000176105 / ENSRNOG00000037227
YIF1A / O95070 / ENSMUSG00000024875 / ENSG00000174851 / ENSRNOG00000020201
YIF1B / Q5BJH7 / ENSMUSG00000030588 / ENSG00000167645 / ENSRNOG00000055286
YIPF3 / Q9GZM5 / ENSMUSG00000071074 / ENSG00000137207 / ENSRNOG00000018998
YIPF4 / Q9BSR8 / ENSMUSG00000024072 / ENSG00000119820 / ENSRNOG00000005610
YIPF5 / Q969M3 / ENSMUSG00000024487 / ENSG00000145817 / ENSRNOG00000014564
YIPF6 / Q96EC8 / ENSMUSG00000047694 / ENSG00000181704 / ENSRNOG00000006642
YKT6 / O15498 / ENSMUSG00000002741 / ENSG00000106636 / ENSRNOG00000014785
YOD1 / Q5VVQ6 / ENSMUSG00000046404 / ENSG00000180667 / ENSRNOG00000025704
YPEL5 / P62699 / ENSMUSG00000039770 / ENSG00000119801 / ENSRNOG00000026742
YRDC / Q86U90 / ENSMUSG00000028889 / ENSG00000196449 / ENSRNOG00000025424
YTHDF1 / Q9BYJ9 / ENSMUSG00000038848 / ENSG00000149658 / ENSRNOG00000027015
YTHDF3 / Q7Z739 / ENSMUSG00000047213 / ENSG00000185728 / ENSRNOG00000010084
YWHAB / P31946 / ENSMUSG00000018326 / ENSG00000166913 / ENSRNOG00000010945
YWHAG / P61981 / ENSMUSG00000051391 / ENSG00000170027 / ENSRNOG00000001436
YWHAH / Q04917 / ENSMUSG00000018965 / ENSG00000128245 / ENSRNOG00000055471
YWHAQ / P27348 / ENSMUSG00000076432 / ENSG00000134308 / ENSRNOG00000051650
YWHAZ / P63104 / ENSMUSG00000022285 / ENSG00000164924 / ENSRNOG00000008195
YY1 / P25490 / ENSMUSG00000021264 / ENSG00000100811 / ENSRNOG00000004339
ZBTB1 / Q9Y2K1 / ENSMUSG00000033454 / ENSG00000126804 / ENSRNOG00000058593
ZBTB20 / Q9HC78 / ENSMUSG00000022708 / ENSG00000181722 / ENSRNOG00000056716
ZBTB44 / Q8NCP5 / ENSMUSG00000047412 / ENSG00000196323 / ENSRNOG00000005578
ZBTB45 / Q96K62 / ENSMUSG00000049600 / ENSG00000119574 / ENSRNOG00000027459
ZBTB7A / O95365 / ENSMUSG00000035011 / ENSG00000178951 / ENSRNOG00000020161
ZBTB7B / O15156 / ENSMUSG00000028042 / ENSG00000160685 / ENSRNOG00000020640
ZBTB8OS / Q8IWT0 / ENSMUSG00000057572 / ENSG00000176261 / ENSRNOG00000008226
ZC2HC1A / Q96GY0 / ENSMUSG00000043542 / ENSG00000104427 / ENSRNOG00000022356
ZC3H13 / Q5T200 / ENSMUSG00000022000 / ENSG00000123200 / ENSRNOG00000011237
ZC3H14 / Q6PJT7 / ENSMUSG00000021012 / ENSG00000100722 / ENSRNOG00000004083
ZC3H15 / Q8WU90 / ENSMUSG00000027091 / ENSG00000065548 / ENSRNOG00000005256
ZC3H18 / Q86VM9 / ENSMUSG00000017478 / ENSG00000158545 / ENSRNOG00000028501
ZC3H7A / Q8IWR0 / ENSMUSG00000037965 / ENSG00000122299 / ENSRNOG00000002420
ZC3H7B / Q9UGR2 / ENSMUSG00000022390 / ENSG00000100403 / ENSRNOG00000055925
ZC3H8 / Q8N5P1 / ENSMUSG00000027387 / ENSG00000144161 / ENSRNOG00000017647
ZC3HAV1 / Q7Z2W4 / ENSMUSG00000029826 / ENSG00000105939 / ENSRNOG00000013948
ZCCHC17 / Q9NP64 / ENSMUSG00000028772 / ENSG00000121766 / ENSRNOG00000012266
ZCCHC8 / Q6NZY4 / ENSMUSG00000029427 / ENSG00000033030 / ENSRNOG00000001243
ZDHHC13 / Q8IUH4 / ENSMUSG00000030471 / ENSG00000177054 / ENSRNOG00000014277
ZDHHC3 / Q9NYG2 / ENSMUSG00000025786 / ENSG00000163812 / ENSRNOG00000004344
ZDHHC5 / Q9C0B5 / ENSMUSG00000034075 / ENSG00000156599 / ENSRNOG00000006832
ZER1 / Q7Z7L7 / ENSMUSG00000039686 / ENSG00000160445 / ENSRNOG00000025832
ZFAND1 / Q8TCF1 / ENSMUSG00000039795 / ENSG00000104231 / ENSRNOG00000010342
ZFAND2B / Q8WV99 / ENSMUSG00000026197 / ENSG00000158552 / ENSRNOG00000018639
Homologs
Brain
Heart
Kidney
Liver
Lung
Testis
Organs
Bins
1
2
3
4
5
Species
Human
Mouse
Rat

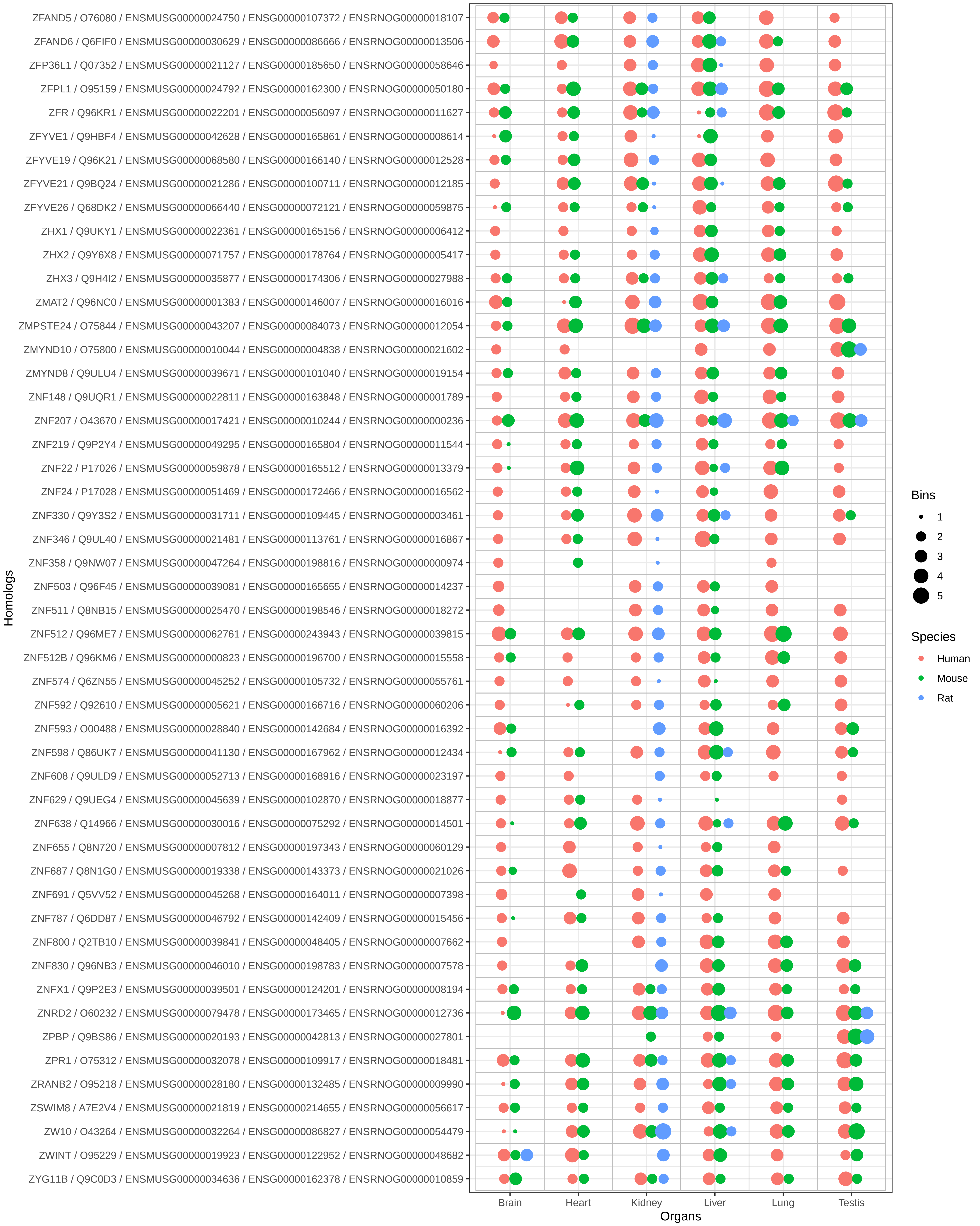
ZFAND5 / O76080 / ENSMUSG00000024750 / ENSG00000107372 / ENSRNOG00000018107
ZFAND6 / Q6FIF0 / ENSMUSG00000030629 / ENSG00000086666 / ENSRNOG00000013506
ZFP36L1 / Q07352 / ENSMUSG00000021127 / ENSG00000185650 / ENSRNOG00000058646
ZFPL1 / O95159 / ENSMUSG00000024792 / ENSG00000162300 / ENSRNOG00000050180
ZFR / Q96KR1 / ENSMUSG00000022201 / ENSG00000056097 / ENSRNOG00000011627
ZFYVE1 / Q9HBF4 / ENSMUSG00000042628 / ENSG00000165861 / ENSRNOG00000008614
ZFYVE19 / Q96K21 / ENSMUSG00000068580 / ENSG00000166140 / ENSRNOG00000012528
ZFYVE21 / Q9BQ24 / ENSMUSG00000021286 / ENSG00000100711 / ENSRNOG00000012185
ZFYVE26 / Q68DK2 / ENSMUSG00000066440 / ENSG00000072121 / ENSRNOG00000059875
ZHX1 / Q9UKY1 / ENSMUSG00000022361 / ENSG00000165156 / ENSRNOG00000006412
ZHX2 / Q9Y6X8 / ENSMUSG00000071757 / ENSG00000178764 / ENSRNOG00000005417
ZHX3 / Q9H4I2 / ENSMUSG00000035877 / ENSG00000174306 / ENSRNOG00000027988
ZMAT2 / Q96NC0 / ENSMUSG00000001383 / ENSG00000146007 / ENSRNOG00000016016
ZMPSTE24 / O75844 / ENSMUSG00000043207 / ENSG00000084073 / ENSRNOG00000012054
ZMYND10 / O75800 / ENSMUSG00000010044 / ENSG00000004838 / ENSRNOG00000021602
ZMYND8 / Q9ULU4 / ENSMUSG00000039671 / ENSG00000101040 / ENSRNOG00000019154
ZNF148 / Q9UQR1 / ENSMUSG00000022811 / ENSG00000163848 / ENSRNOG00000001789
ZNF207 / O43670 / ENSMUSG00000017421 / ENSG00000010244 / ENSRNOG00000000236
ZNF219 / Q9P2Y4 / ENSMUSG00000049295 / ENSG00000165804 / ENSRNOG00000011544
ZNF22 / P17026 / ENSMUSG00000059878 / ENSG00000165512 / ENSRNOG00000013379
ZNF24 / P17028 / ENSMUSG00000051469 / ENSG00000172466 / ENSRNOG00000016562
ZNF330 / Q9Y3S2 / ENSMUSG00000031711 / ENSG00000109445 / ENSRNOG00000003461
ZNF346 / Q9UL40 / ENSMUSG00000021481 / ENSG00000113761 / ENSRNOG00000016867
ZNF358 / Q9NW07 / ENSMUSG00000047264 / ENSG00000198816 / ENSRNOG00000000974
ZNF503 / Q96F45 / ENSMUSG00000039081 / ENSG00000165655 / ENSRNOG00000014237
ZNF511 / Q8NB15 / ENSMUSG00000025470 / ENSG00000198546 / ENSRNOG00000018272
ZNF512 / Q96ME7 / ENSMUSG00000062761 / ENSG00000243943 / ENSRNOG00000039815
ZNF512B / Q96KM6 / ENSMUSG00000000823 / ENSG00000196700 / ENSRNOG00000015558
ZNF574 / Q6ZN55 / ENSMUSG00000045252 / ENSG00000105732 / ENSRNOG00000055761
ZNF592 / Q92610 / ENSMUSG00000005621 / ENSG00000166716 / ENSRNOG00000060206
ZNF593 / O00488 / ENSMUSG00000028840 / ENSG00000142684 / ENSRNOG00000016392
ZNF598 / Q86UK7 / ENSMUSG00000041130 / ENSG00000167962 / ENSRNOG00000012434
ZNF608 / Q9ULD9 / ENSMUSG00000052713 / ENSG00000168916 / ENSRNOG00000023197
ZNF629 / Q9UEG4 / ENSMUSG00000045639 / ENSG00000102870 / ENSRNOG00000018877
ZNF638 / Q14966 / ENSMUSG00000030016 / ENSG00000075292 / ENSRNOG00000014501
ZNF655 / Q8N720 / ENSMUSG00000007812 / ENSG00000197343 / ENSRNOG00000060129
ZNF687 / Q8N1G0 / ENSMUSG00000019338 / ENSG00000143373 / ENSRNOG00000021026
ZNF691 / Q5VV52 / ENSMUSG00000045268 / ENSG00000164011 / ENSRNOG00000007398
ZNF787 / Q6DD87 / ENSMUSG00000046792 / ENSG00000142409 / ENSRNOG00000015456
ZNF800 / Q2TB10 / ENSMUSG00000039841 / ENSG00000048405 / ENSRNOG00000007662
ZNF830 / Q96NB3 / ENSMUSG00000046010 / ENSG00000198783 / ENSRNOG00000007578
ZNFX1 / Q9P2E3 / ENSMUSG00000039501 / ENSG00000124201 / ENSRNOG00000008194
ZNRD2 / O60232 / ENSMUSG00000079478 / ENSG00000173465 / ENSRNOG00000012736
ZPBP / Q9BS86 / ENSMUSG00000020193 / ENSG00000042813 / ENSRNOG00000027801
ZPR1 / O75312 / ENSMUSG00000032078 / ENSG00000109917 / ENSRNOG00000018481
ZRANB2 / O95218 / ENSMUSG00000028180 / ENSG00000132485 / ENSRNOG00000009990
ZSWIM8 / A7E2V4 / ENSMUSG00000021819 / ENSG00000214655 / ENSRNOG00000056617
ZW10 / O43264 / ENSMUSG00000032264 / ENSG00000086827 / ENSRNOG00000054479
ZWINT / O95229 / ENSMUSG00000019923 / ENSG00000122952 / ENSRNOG00000048682
ZYG11B / Q9C0D3 / ENSMUSG00000034636 / ENSG00000162378 / ENSRNOG00000010859
Homologs
Brain
Heart
Kidney
Liver
Lung
Testis
Organs
Bins
1
2
3
4
5
Species
Human
Mouse
Rat

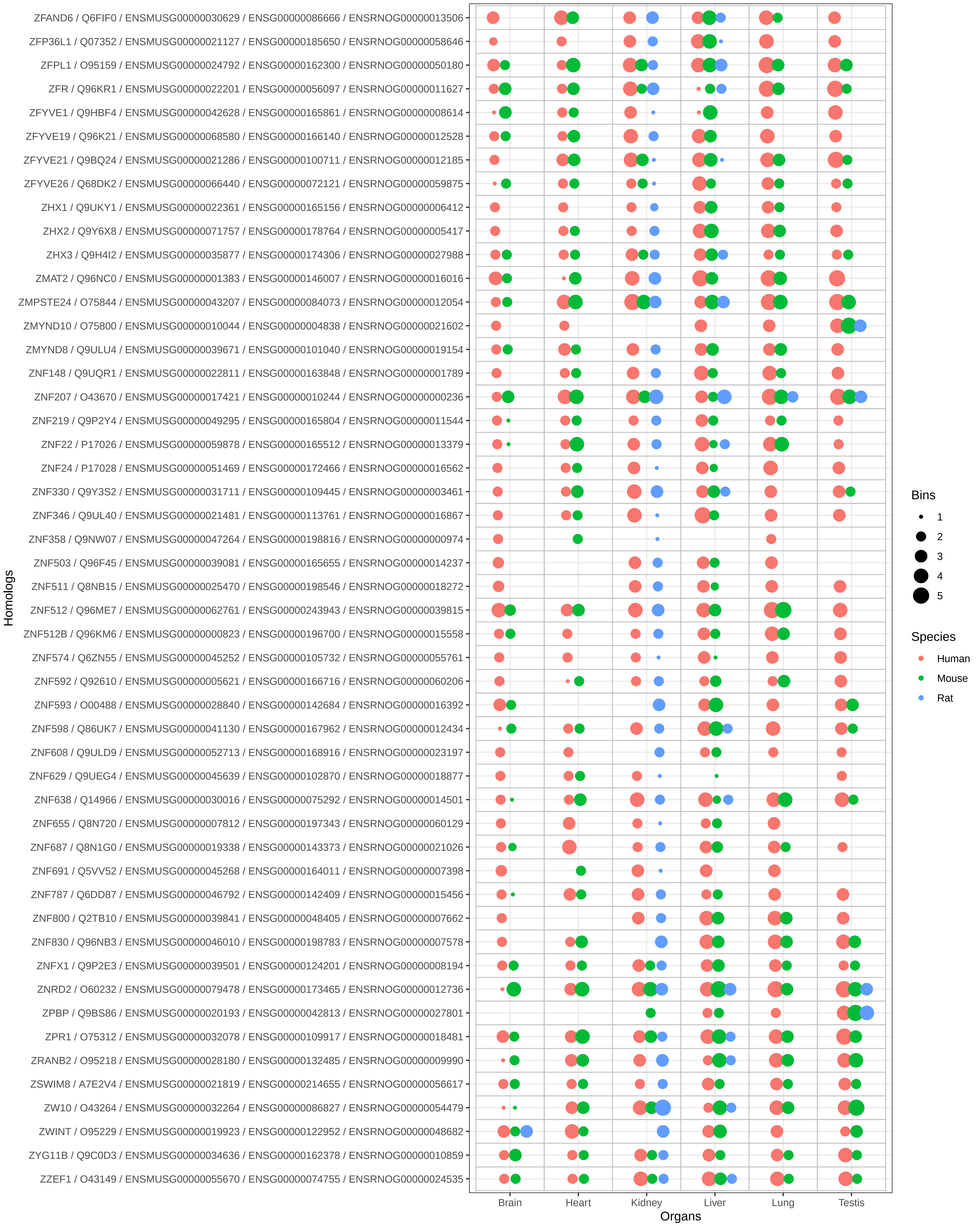
ZFAND6 / Q6FIF0 / ENSMUSG00000030629 / ENSG00000086666 / ENSRNOG00000013506
ZFP36L1 / Q07352 / ENSMUSG00000021127 / ENSG00000185650 / ENSRNOG00000058646
ZFPL1 / O95159 / ENSMUSG00000024792 / ENSG00000162300 / ENSRNOG00000050180
ZFR / Q96KR1 / ENSMUSG00000022201 / ENSG00000056097 / ENSRNOG00000011627
ZFYVE1 / Q9HBF4 / ENSMUSG00000042628 / ENSG00000165861 / ENSRNOG00000008614
ZFYVE19 / Q96K21 / ENSMUSG00000068580 / ENSG00000166140 / ENSRNOG00000012528
ZFYVE21 / Q9BQ24 / ENSMUSG00000021286 / ENSG00000100711 / ENSRNOG00000012185
ZFYVE26 / Q68DK2 / ENSMUSG00000066440 / ENSG00000072121 / ENSRNOG00000059875
ZHX1 / Q9UKY1 / ENSMUSG00000022361 / ENSG00000165156 / ENSRNOG00000006412
ZHX2 / Q9Y6X8 / ENSMUSG00000071757 / ENSG00000178764 / ENSRNOG00000005417
ZHX3 / Q9H4I2 / ENSMUSG00000035877 / ENSG00000174306 / ENSRNOG00000027988
ZMAT2 / Q96NC0 / ENSMUSG00000001383 / ENSG00000146007 / ENSRNOG00000016016
ZMPSTE24 / O75844 / ENSMUSG00000043207 / ENSG00000084073 / ENSRNOG00000012054
ZMYND10 / O75800 / ENSMUSG00000010044 / ENSG00000004838 / ENSRNOG00000021602
ZMYND8 / Q9ULU4 / ENSMUSG00000039671 / ENSG00000101040 / ENSRNOG00000019154
ZNF148 / Q9UQR1 / ENSMUSG00000022811 / ENSG00000163848 / ENSRNOG00000001789
ZNF207 / O43670 / ENSMUSG00000017421 / ENSG00000010244 / ENSRNOG00000000236
ZNF219 / Q9P2Y4 / ENSMUSG00000049295 / ENSG00000165804 / ENSRNOG00000011544
ZNF22 / P17026 / ENSMUSG00000059878 / ENSG00000165512 / ENSRNOG00000013379
ZNF24 / P17028 / ENSMUSG00000051469 / ENSG00000172466 / ENSRNOG00000016562
ZNF330 / Q9Y3S2 / ENSMUSG00000031711 / ENSG00000109445 / ENSRNOG00000003461
ZNF346 / Q9UL40 / ENSMUSG00000021481 / ENSG00000113761 / ENSRNOG00000016867
ZNF358 / Q9NW07 / ENSMUSG00000047264 / ENSG00000198816 / ENSRNOG00000000974
ZNF503 / Q96F45 / ENSMUSG00000039081 / ENSG00000165655 / ENSRNOG00000014237
ZNF511 / Q8NB15 / ENSMUSG00000025470 / ENSG00000198546 / ENSRNOG00000018272
ZNF512 / Q96ME7 / ENSMUSG00000062761 / ENSG00000243943 / ENSRNOG00000039815
ZNF512B / Q96KM6 / ENSMUSG00000000823 / ENSG00000196700 / ENSRNOG00000015558
ZNF574 / Q6ZN55 / ENSMUSG00000045252 / ENSG00000105732 / ENSRNOG00000055761
ZNF592 / Q92610 / ENSMUSG00000005621 / ENSG00000166716 / ENSRNOG00000060206
ZNF593 / O00488 / ENSMUSG00000028840 / ENSG00000142684 / ENSRNOG00000016392
ZNF598 / Q86UK7 / ENSMUSG00000041130 / ENSG00000167962 / ENSRNOG00000012434
ZNF608 / Q9ULD9 / ENSMUSG00000052713 / ENSG00000168916 / ENSRNOG00000023197
ZNF629 / Q9UEG4 / ENSMUSG00000045639 / ENSG00000102870 / ENSRNOG00000018877
ZNF638 / Q14966 / ENSMUSG00000030016 / ENSG00000075292 / ENSRNOG00000014501
ZNF655 / Q8N720 / ENSMUSG00000007812 / ENSG00000197343 / ENSRNOG00000060129
ZNF687 / Q8N1G0 / ENSMUSG00000019338 / ENSG00000143373 / ENSRNOG00000021026
ZNF691 / Q5VV52 / ENSMUSG00000045268 / ENSG00000164011 / ENSRNOG00000007398
ZNF787 / Q6DD87 / ENSMUSG00000046792 / ENSG00000142409 / ENSRNOG00000015456
ZNF800 / Q2TB10 / ENSMUSG00000039841 / ENSG00000048405 / ENSRNOG00000007662
ZNF830 / Q96NB3 / ENSMUSG00000046010 / ENSG00000198783 / ENSRNOG00000007578
ZNFX1 / Q9P2E3 / ENSMUSG00000039501 / ENSG00000124201 / ENSRNOG00000008194
ZNRD2 / O60232 / ENSMUSG00000079478 / ENSG00000173465 / ENSRNOG00000012736
ZPBP / Q9BS86 / ENSMUSG00000020193 / ENSG00000042813 / ENSRNOG00000027801
ZPR1 / O75312 / ENSMUSG00000032078 / ENSG00000109917 / ENSRNOG00000018481
ZRANB2 / O95218 / ENSMUSG00000028180 / ENSG00000132485 / ENSRNOG00000009990
ZSWIM8 / A7E2V4 / ENSMUSG00000021819 / ENSG00000214655 / ENSRNOG00000056617
ZW10 / O43264 / ENSMUSG00000032264 / ENSG00000086827 / ENSRNOG00000054479
ZWINT / O95229 / ENSMUSG00000019923 / ENSG00000122952 / ENSRNOG00000048682
ZYG11B / Q9C0D3 / ENSMUSG00000034636 / ENSG00000162378 / ENSRNOG00000010859
ZZEF1 / O43149 / ENSMUSG00000055670 / ENSG00000074755 / ENSRNOG00000024535
Homologs
Brain
Heart
Kidney
Liver
Lung
Testis
Organs
Bins
1
2
3
4
5
Species
Human
Mouse
Rat
